# In-depth computational analysis of natural and artificial carbon fixation pathways

**DOI:** 10.1101/2021.01.05.425423

**Authors:** Hannes Löwe, Andreas Kremling

**Affiliations:** Systems Biotechnology, Technical University of Munich

## Abstract

In the recent years, engineering new-to-nature CO2 and C1 fixing metabolic pathways made a leap forward. These new, artificial pathways promise higher yields and activity than natural ones like the Calvin-Benson-Bassham cycle. The question remains how to best predict their *in vivo* performance and what actually makes one pathway “better” than another.

In this context, we explore aerobic carbon fixation pathways by a computational approach and compare them based on their ATP-efficiency and specific activity considering the kinetics and thermodynamics of the reactions. Beside natural pathways, this included the artificial Reductive Glycine Pathway, the CETCH cycle and two completely new cycles with superior stoichiometry: The Reductive Citramalyl-CoA cycle and the 2-Hydroxyglutarate-Reverse Tricarboxylic Acid cycle. A comprehensive kinetic data set was collected for all enzymes of all pathways and missing kinetic data was sampled with the Parameter Balancing algorithm. Kinetic and thermodynamic data were fed to the Enzyme Cost Minimization algorithm to check for respective inconsistencies and calculate pathway specific activities.

We found that the Reductive Glycine Pathway, the CETCH cycle and the new Reductive Citramalyl-CoA cycle were predicted to have higher ATP-efficiencies and specific activities than the natural cycles. The Calvin Cycle performed better than previously thought, however. It can be concluded that the weaker overall characteristics in the design of the Calvin Cycle might be compensated by other benefits like robustness, low nutrient demand and a good compatibility with the host’s physiological requirements. Nevertheless, the artificial carbon fixation cycles hold great potential for future applications in Industrial Biotechnology and Synthetic Biology.

## Introduction

As of 2020, most of the earth’s surface is cultivated by mankind: Around 50% of the habitable land is actually agricultural land with 37% of the remaining land being forests (with a decreasing trend) (World bank & Food and Agriculture Organization of the UN, 2016). In order to produce more crop to feed a growing population or to fuel the bioeconomy, it is clear that either natural ecosystems have to be converted to agricultural land or agriculture has to be intensified, both with potentially tremendous downsides for nature. Crop plant areal productivity is finally limited by photosynthetic efficiency that is rather low: only around 2.7-3.7% of the incoming sunlight is converted to a usable product (Zhu et al., 2008) and this only during the growth season. Among other reasons, the low efficiency has been attributed to the low turnover of the key enzyme, Ribulose-1,5-Bisphosphate Carboxylase/Oxygenase (RuBisCO), and its tendency to use oxygen in a wasteful side reaction. The product of this reaction, phosphoglycolate, has to be recovered in an energetically wasteful process called photorespiration. Another reason for low productivity was found in the suboptimal efficiency of the Reductive Pentose Phosphate cycle, better known as the Calvin-Benson-Bassham cycle (CBB cycle) which is the central carbon fixation pathway in plants, green algae and cyanobacteria. In nature, some carboxylating enzymes can be found that have supposedly better characteristics than RuBisCO (Bar-Even et al., 2010) which led to the question why evolution developed photosynthesis the way it is. This has inspired researchers to explore whether the dark reaction of photosynthesis can be augmented to support higher productivity by mainly four approaches: i) Enzyme engineering of RuBisCO to increase its activity (Wilson & Whitney, 2017), ii) Implementation of CO2 concentration mechanisms to reduce photorespiration (A. I. Flamholz et al., 2020; Long et al., 2018), iii) Replacing C3 with C4 photosynthesis or with completely new carbon fixation pathways, like the CETCH cycle (Schwander et al., 2016), iv) Installing more efficient pathways for the recycling of phosphoglycolate during photorespiration (South et al., 2019; Trudeau et al., 2018). A more detailed overview of the efforts to improve photosynthesis can be found in (Bar-Even, 2018).

A further option to circumvent the limitations of the areal productivity of photosynthesis is to use alternative sources of energy and carbon, like the C1 compounds formic acid, methanol or a mixture of H2 and CO2 gas. These electron and carbon sources can be derived from waste/biomass gasification or electrolysis which increases the areal productivity compared to crop-based primary production by an order of magnitude (Nangle et al., 2020). For the assimilation of these compounds, nature also has evolved specialized microorganisms with dedicated pathways like the Serine cycle or the Ribulose Monophosphate (RuMP) cycle that are thought to be more efficient than the traditional CBB cycle (Claassens et al., 2019). For biotechnological purposes, it was tried to implant these pathways into *Escherichia coli* with mixed success (He et al., 2018; Keller et al., 2020; Yu & Liao, 2018). Additionally, multiple optimized artificial pathways for C1-assimilation have been proposed and their basic working principle could be shown (Chou et al., 2019; He et al., 2020; Siegel et al., 2015). Among them, the Reductive Glycine Pathway (rGlyP) that was developed by the group of Arren Bar-Even and others, shines as it is the most efficient aerobic formate-dependent pathway and the only pathway to be integrated *in vivo* to support growth with C1-compounds as the sole carbon and energy source (Claassens et al., 2020; Kim et al., 2020). This work is dedicated to Arren Bar-Even, a visionary in pathway design and metabolic engineering, who passed away far too early in the zenith of his scientific achievements.

Throughout this article, we will call biochemical pathways simply “pathways”, which may be linear biochemical pathways or cycles, for simplification. When comparing different pathways, the new artificial designs presented in former studies were usually thought to be better because of the following reasons: i) a lower demand of ATP to form a product (Bar-Even et al., 2010; He et al., 2020; Schwander et al., 2016), ii) key enzymes with a higher activity (Bar-Even et al., 2010; Cotton et al., 2018), iii) a higher mean thermodynamic driving force (i.e. free energy *ΔG*) of the reactions (Hädicke et al., 2018; Satanowski et al., 2020; Siegel et al., 2015), or iv) a higher affinity for CO2 (Bar-Even et al., 2010; Cotton et al., 2018). All these criteria have limitations, however, and there hasn’t been a conclusive comparison to our knowledge that systematically took into account the enzyme kinetics apart from the forward rate constant. As more efficient pathways are supposed to have lower energy losses and thus usually operate closer to thermodynamic equilibrium, reversibility of reactions accounts for a higher demand for the respective enzymes. Additionally, kinetic undersaturation will limit a pathway’s activity with RuBisCO’s affinity for CO2 being a prominent example. (Noor et al., 2016) developed a method to integrate these additional thermodynamic and kinetic constraints by means of the Enzyme Cost Minimization (ECM) algorithm. This algorithm predicts pathway activities if thermodynamic and kinetic data are available. Missing kinetic parameters can be estimated using the Parameter Balancing algorithm (Lubitz et al., 2010) to yield a complete and consistent set of parameters for each reaction.

In this work, we made an attempt to objectively compare natural occurring and artificial pathways considering their product specific energy costs projected to ATP demand and the pathway specific activity. Only oxygen-tolerant pathways were considered since data for most of the anaerobic pathway is scarce and they will not be able to produce ATP-intensive metabolites without by-product formation (Bertsch & Müller, 2015; Takors et al., 2018). By recombining reactions of existing pathways and by adding reactions that were introduced in recent publications, we present 2 new artificial pathways: the reductive Citramalyl-CoA Cycle (rCCC) and the 2-Hydroxyglutarate-Reductive Tricarboxylic Acid Cycle (2-HG-rTCA). Both were designed to perform better than all known pathways in terms of ATP-efficiency. Finally, we compare natural and artificial pathways using the ECM algorithm and draw conclusions about evolution, design principles and future perspectives of these pathways.

## Material & Methods

### Data collection

In order to quantify the pathway specific activities of natural and artificial CO2- and C1-fixing pathways, a comprehensive data set was collected from different sources which was then used for further calculations. The central sources of enzymatic data and kinetic data in particular are listed in Table 1 specifying which kinetic data was extracted from each source. For most enzymes, kinetic data exists for enzymes from various organisms under varying conditions. We chose parameters of optimal enzymes (i.e. those that allow for the highest activity) to explore the potential of each pathway. These enzymes could be derived from any organisms in all domains of life. All primary sources were manually curated from original publications as parameters in BRENDA are often stored with wrong units. The complete workflow is illustrated in Figure 1.

**Table 1:** Data sources for kinetic data used in this work

| Database/source | Extracted data |
| --- | --- |
| KEGG (Kanehisa & Goto, 2000) | Reaction stoichiometries<br>Systematic names of enzymes |
| eQuilibrator (A. Flamholz et al., 2012) | Equilibrium constants and Gibbs free energies under standard conditions $\Delta_r G^\circ$ and respective uncertainties (pH 7.5, Ionic strength of 0.25 M)<br>Templates for SBTAB-files using the ECM pathway analysis platform |
| BRENDA (Jeske et al., 2019) | Starting point for kinetic data search, using the primary sources linked with kinetic data<br>Direct source of kinetic data when no data was available for a certain enzyme |
| UniProt (Bateman, 2019) | Protein molecular weights if not specified in primary source |
| Primary literature focused on enzyme characterization* | • Main source for enzymatic parameters $K_m$, $k_{cat}^+$ and $k_{cat}^-$<br>• Molecular weight of enzymes if available |

\* details are listed in supplementary Table S1

**Figure 1:**
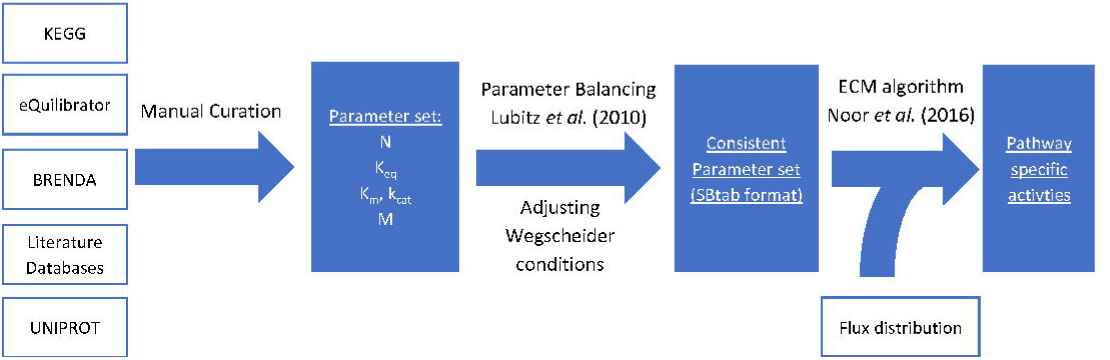
Schematic workflow that was applied to calculate the pathway specific activities from the kinetic parameter set and the flux distributions of each pathway. Box with white-colored filling indicate input data(-sources), blue boxes indicate processed data. N: stoichiometric matrix, Keq: equilibrium constants, Km: Michaelis constants, kcat: forward rate constants, M: molecular weight.

During data collection, kinetic parameter sets with more data for Michaelis constants and those with forward and backward rate constants were preferred over sets with less measured data. The most important parameters which are decisive for a good estimate are the rate constant of the direction the reaction is supposed to operate and the *K*_*m*_ of the primary substrate, i.e. usually the carbon intermediate in the respective pathway.

All kinetic data were brought together in an SBtab model file and SBtab data file as specified in (Lubitz et al., 2016; Noor et al., 2016). The templates of the respective SBtab files were provided by the ECM pathway analysis platform on the eQuilibrator website. (http://equilibrator.weizmann.ac.il/pathway/) An additional column for parameter uncertainties was added and table headings were slightly modified to match the current implementation of the Metabolic Network Toolbox (https://github.com/liebermeister/metabolic-network-toolbox) in MATLAB that was used to process the data. The final SBtab files are included as part of the MATLAB files in supplementary information.

### Implementation of quinone dependent reactions

During data collection, it had to be decided which enzymes are used for each reaction as there are cases in which a reaction with identical main substrate and product is catalyzed by different enzymes. The only difference in these cases might be the use of different cofactors or whether the reaction is coupled to ion translocation through the membrane. This was especially important for the electron transport chain. It is known that depending on the organism and the conditions, different variants of the electron transport chain take place (Borisov et al., 2011). Additionally, different quinones with varying redox potential might be used. In the following section, we present how the quinone-dependent reactions were integrated into the data structure.

For the analysis of pathways, quinones were treated as cofactors like other electron carriers, e.g. NADH and were not regenerated in the reaction stoichiometries. The only quinone dependent enzymes that were integrated in the data structure are oxidoreductases that turn double bonds to single bonds and *vice versa* like the succinate dehydrogenase complex. In the Citric Acid Cycle, the direction of this enzyme is towards the oxidation of succinate which is usually accompanied by ubiquinone reduction due to the high redox potential of the ubiquinone(UQ)/ubiquinol(UQH2) redox couple (Cecchini et al., 2002). To poise the reaction in the reverse, reductive direction, bacteria, e.g. *E. coli* (Cecchini et al., 2002), usually use quinones with a lower redox potential like the menaquinone(MQ)/menaquinol(MQH2) couple. In *Bacillus subtilis*, this menaquinone is used for the oxidation of succinate accompanied by the translocation of 2 protons from the periplasm to the cytosol which has been shown to be reversible (Schnorpfeil et al., 2001). Therefore, different pathways that use the enzyme in either direction would be able to operate with this enzyme which is why menaquinone and kinetic data of the *B. subtilis* enzyme was chosen for the reaction in the data structure. To account for the 2 protons that are pumped by the enzyme through the membrane, the equilibrium constant of this reaction was adjusted since the proton gradient was not included in the equilibrium constant taken from eQuilibrator. The change in the equilibrium constant was calculated according to the following formula, assuming a membrane potential Δ*ψ* of −150 mV:

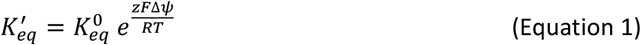

With 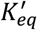: The adjusted equilibrium constant, 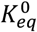: Original equilibrium constant provided by eQuilibrator, *F*: Faraday Constant, *z*: Number of protons transferred from cytosol to periplasm, Δ*ψ*: Transmembrane potential, R: General Gas Constant, *T*: Temperature

Depending on the direction of proton translocation, the exponent of the exponential function in Equation 1 will have a positive or negative exponent (per definition: z is positive for transport from cytosol to periplasm). When the reaction is using succinate as substrate and fumarate as product, the equilibrium constant has to be higher than without considering the proton gradient (2 protons from the periplasm are transferred to the cytoplasm) and hence the exponent is positive. It has to be noted that ubiquinone dependent succinate dehydrogenase also shows reversibility (to a smaller extent however), so using menaquinone in the stoichiometry is just one option.

Apart from the succinate dehydrogenase, only one other type reaction with quinones was implemented: Acyl-CoA dehydrogenases that desaturate carbon-carbon single bonds, as found e.g. in β-oxidation. These enzymes were assumed to channel the electrons to the electron transport chain at the level of ubiquinone which is an exergonic reaction under physiological conditions.

### Kinetic parameter estimation by Parameter Balancing

The SBtab files with the collected data were used as the input for the Parameter Balancing algorithm to estimate the missing kinetic parameters. To this end, a kinetic model for the reaction has to be defined. Since the exact reaction mechanisms are not known for every enzyme, a modular rate law for the flux v was used that is flexible enough to give good estimations (Liebermeister et al., 2010):

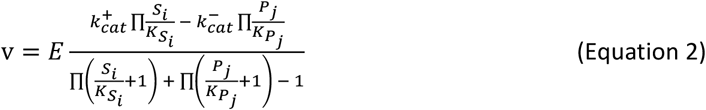

with *E*: enzyme concentration, *S*_*i*_: concentration of substrate *i, P*_*j*_: concentration of product *j*, 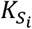: half-saturation constant of substrate *i*, 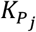: half-saturation constant of product *j*,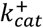: forward rate constant, 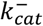: backward rate constant

Following the derivation by (Noor et al., 2016), this can be refactored to:

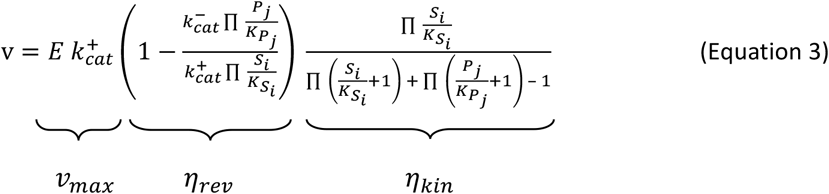

*v*_*max*_ is the maximal velocity of the reaction or the “capacity” of the enzyme that is decreased by the reversibility factor *η*_*rev*_ and the kinetic factor *η*_*kin*_. These factors are numbers between 0 and 1. This holds for positive fluxes – for negative fluxes, the equation is refactored to:

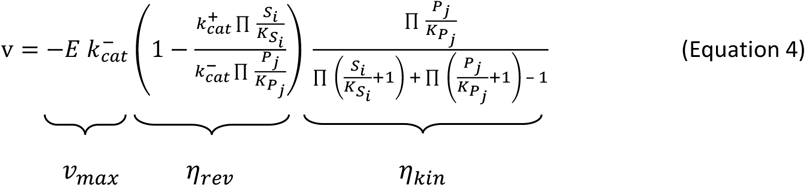

The conditional factors give information on why reactions are not running at their maximum capacity: A low reversibility factor *η*_*rev*_ indicates that the thermodynamics are limiting the reaction velocity, e.g. because of a low equilibrium constant. A low kinetic factor *η*_*kin*_ indicates a kinetic limitation because of substrate or product saturation (Noor et al., 2016). This kinetic law was used as a template to estimate the remaining parameters with the Parameter Balancing algorithm.

Therefore, the data from the SBtab data file was loaded into MATLAB and each reaction was balanced alone without considering the other reactions in the network using the MATLAB implementation of Parameter Balancing. The equilibrium constants were not adjusted, but kept fixed during balancing to ensure that Wegscheider relationships are still fulfilled after balancing. This was done by setting the option “fix_Keq_in_sampling” to 1. After parameter balancing, the resulting parameters should fulfil the Haldane relationship which can be formulated as follows (Noor et al., 2016):

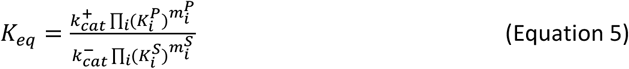

with *K*_*eq*_: equilibrium constant, *k*_*cat*_: forward and backward rate constants, 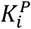: Michaelis constant of product i, 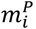: stoichiometric factor of product i, 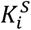: Michaelis constant of substrate i,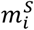: stoichiometric factor of substrate i.

For some reactions the algorithm did not produce consistent sets of parameters because the data that were used as a basis, were contradictory with respect to the equilibrium constants of the reactions. This would typically happen if forward and backward kinetic constants were both available. In these cases, the uncertainties of the measured constants as well as the parameter prior uncertainties of kinetic rate constants and Michaelis constants were iteratively increased and balanced until the Haldane relationships were fulfilled. The Parameter Balancing algorithm did not only give estimates for parameter values, but also for their uncertainties. Both were stored in data structures for further calculations. After this step, the data structure with the network kinetics contained a complete set of parameters for all reactions and all reactants. The corresponding MATLAB code is included as part of the MATLAB files in supplementary information.

### Correction of the kinetics of the C5-carboxylic acid subnetwork

For some reactions in the data structure, estimates of the equilibrium constants that were derived from eQuilibrator had a very high uncertainty. This was especially pronounced for the reactions that formed or cleaved C5-dicarboxylic acids, for instance the reaction of β-methylmalyl-CoA lyase:

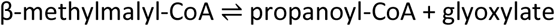

eQuilibrator predicts a free energy of *Δ*_*r*_*G*^*0*^ = 15.9 ± 16.4 kJ/mol for this reaction translating to a standard deviation of a factor of around 750 in the equilibrium constant, which has a considerable impact on the directionality of the reactions. For these reactions, the equilibrium constants were adjusted by using the kinetic information that was available for these reaction: As the equilibrium constant is mathematically related to the other parameters (compare Equation 5), this could be done by estimating a new equilibrium constant for β-methylmalyl-CoA lyase by Parameter Balancing (a complete set of parameters was determined for this reaction (Cecchini et al., 2002)). Next, the other reactions of C5-carboxylic acids with high uncertainties were corrected by the relative change of the new estimate for the equilibrium constant compared to the original constant of β-methylmalyl-CoA lyase. This was necessary to fulfil Wegscheider conditions that were otherwise violated by the change of the equilibrium constant of β-methylmalyl-CoA lyase. By just changing the equilibrium constants of the reactions with high uncertainties, the Haldane relationship in Equation 5 did not hold anymore, however, since only the left side of the equation was changed. A conservative way to adjust the right side of Equation 5 is to modify those parameters that are estimated instead of the values that were actually experimentally determined. As an example: If the forward reaction rate 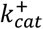of a certain reaction is known from experimental data and the equilibrium constant is corrected to be twice as high as estimated by eQuilibrator, the Haldane relationship can be fulfilled by dividing the backward reaction rate 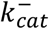by 2.

All reactions that were affected by the change of the equilibrium constant of β-methylmalyl-CoA lyase, were adjusted by the before mentioned approach. After the corrections, the reactions again fulfilled Wegscheider conditions and Haldane relationships.

### Determination of pathway specific activities

To calculate the pathway specific activity of a selected pathway, the MATLAB implementation of the ECM algorithm was used (Noor et al., 2016). The MATLAB function for ECM needs four arguments: i) a biochemical network containing the reaction stoichiometries, ii) the kinetic parameters of the reactions, iii) a flux distribution representing the selected pathway and iv) an options data structure which can be used to specify upper and lower bounds for metabolites, cost weights for enzyme concentrations as well as further options.

These four arguments were provided as follows: The reaction stoichiometries (i) were specified in the SBtab model file that was assembled during data collection. The stoichiometries were loaded into a data structure with the Metabolic Network Toolbox. The data structure with the kinetic parameters was obtained by Parameter Balancing (for details, compare the respective section in Material&Methods). For each pathway, a flux distribution (iii) was set up as a vector that reflects fluxes in the selected pathway. For the upper and lower bounds of metabolite concentrations (iv), the standard bounds provided by the ECM pathway analysis platform of eQuilibrator were taken and slightly modified with values from the literature to represent a realistic profile of metabolite concentrations (detailed values for metabolite bounds can be found in the supplementary information in Table S2). As enzyme cost weights (iv), the molecular weights of the enzymes were taken as it was assumed that bigger enzymes will be costlier to synthetize for the cells (Kaleta et al., 2013; Noor et al., 2016). The molecular weights of enzymes are stored in a table in the SBtab model file and were loaded into a data structure.

The ECM algorithm will return the minimal enzyme concentrations to support the given flux distribution and thereby offer a measure of pathway activity per milligram of total enzyme. In addition, the enzyme capacities, reversibility and kinetic factors (compare Equation 3) are returned by the algorithm. With these factors, it can be evaluated if a certain reaction is thermodynamically or kinetically limited as described in the section on Parameter Balancing.

Uncertainties were estimated by a Monte Carlo method, where in every iteration the input parameters (i.e. the kinetic parameters) were randomly varied according to their geometric standard deviations assuming a log-normal distribution. The uncertainties of kinetic parameters were derived during Parameter Balancing as described in the respective section. The final values that are presented in the results section are the mean values and standard deviations of 100 iterations.

### Calculation of projected ATP costs of pathways

To compare the pathways in terms of efficiency, a common criterion for all pathways had to be found as they use different cofactors. This was done by converting all costs of a pathway into projected ATP demand. In reactions that produce AMP, it was assumed that two equivalents of ATP are required for its regeneration to ATP. NADH can be oxidized by the respiratory chain to yield a total of 2.5 ATP during optimal aerobic respiration (Berg et al., 2010) which represents an optimistic estimate. FADH2 which is a cofactor of the succinate dehydrogenase complex is supposed to yield 1.5 ATP (Berg et al., 2010), the same should be true for electrons derived from the oxidation of FADH2-dependent acyl-CoA dehydrogenases that are found in β-oxidation. This results in a yield of 1.5 ATP per ubiquinol as it is the direct acceptor of electrons from FADH2.

The NADH:ubiquinone oxidoreductase complex is supposed to transport 4 protons from the cytosol to the periplasm, equaling 1 ATP (Friedrich, 1998). During fumarate respiration in *E. coli*, the proton translocating NADH dehydrogenase (type I) is used (Tran et al., 1997) which leads to the assumption that the maximal achievable stoichiometry would be:

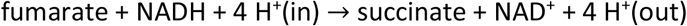

Assuming a H^+^/ATP ratio of the ATP synthase of 4, this results in the generation of 1 ATP. In other words: the cost for one menaquinol or ubiquinol on the substrate side is −1 ATP with the implicit condition that the electrons carried by the quinol are derived from NADH. The difference in the costs of NADPH and NADH depends on the ratio of their reduced and oxidized form, so for reasons of simplification, it was assumed that both have the same cost. One could argue though that NADPH is worth more ATP as it is usually more reduced than NADH (i.e. the NADPH/NADP^+^ ratio is thought to be a lot higher than the NADH/NAD^+^ ratio in bacteria (Sauer et al., 2004)). Therefore, the free energy of the oxidation of NADPH will be higher in many cases.

It also has to be stated that different organisms use different respiratory chains (Borisov et al., 2011; Shepherd & Poole, 2013) and the stoichiometry of ATP synthase may also vary (e.g. in chloroplasts the H^+^/ATP ratio is rather 4.67 than 4 (Davis & Kramer, 2020)). Therefore, the projected ATP cost represent rather optimistic estimates, i.e. they might be underestimated. This does not influence the overall comparison between pathways, however.

## Results

### Overview of the natural and artificial pathways used in this study

Seven pathways were chosen from the literature and implemented by adding their stoichiometries and relevant kinetic and thermodynamic parameters to the SBtab model file and SBtab data file as described in Material and Methods. An overview of these pathways is given in Figure 2.

**Figure 2:**
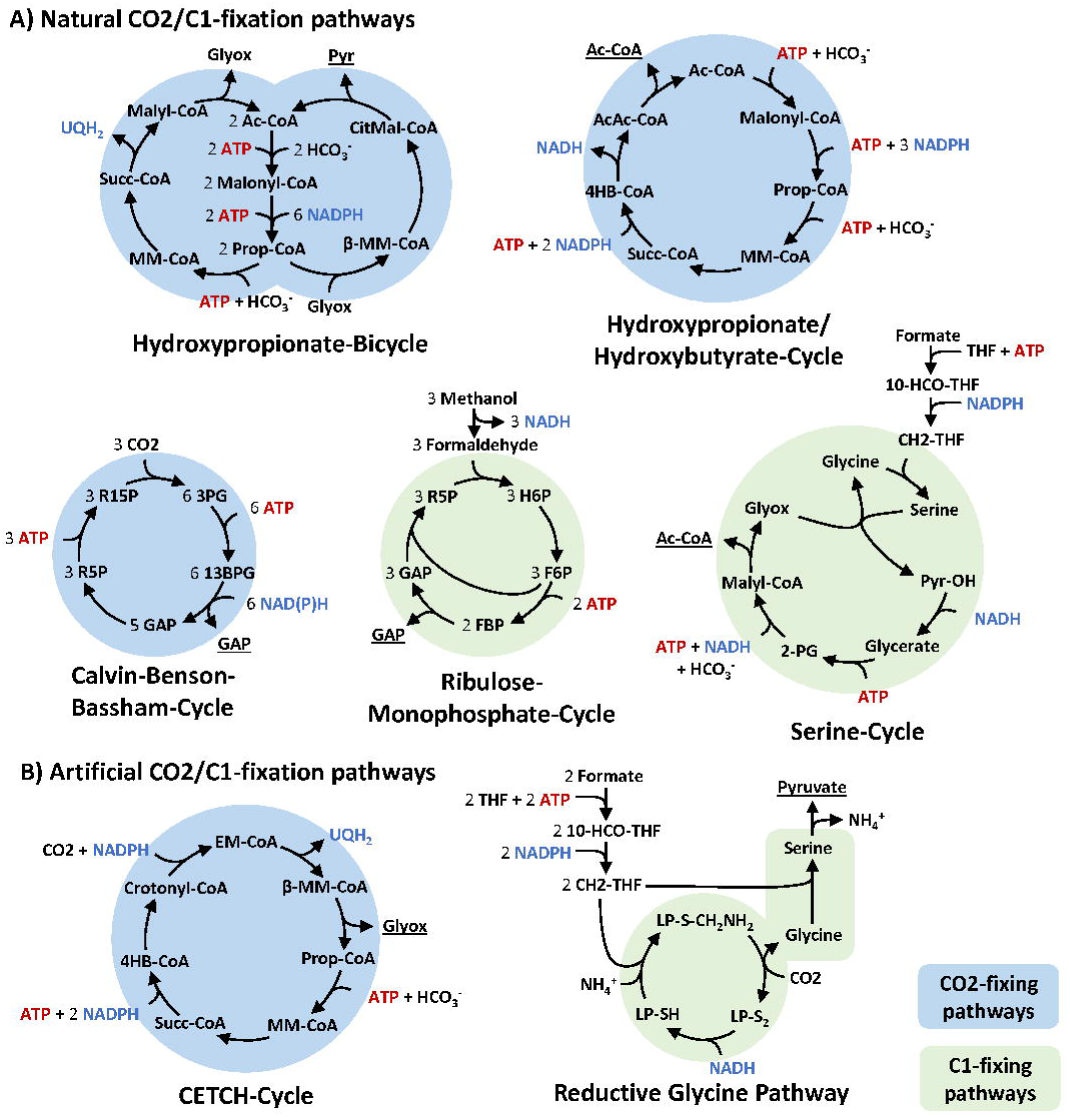
Overview of natural occurring and artificial CO2 and C1-fixing pathways that were integrated in the model. For simplification, reaction arrows can include multiple reactions and skip some metabolites. ADP, AMP, phosphate, water, and oxidized forms of electron carriers were also left out to improve clarity. Primary products of the pathways are underlined. **A)** Natural Pathways. **B)** Artificial Pathways. Abbreviations: Ac-CoA, acetyl-CoA; Prop-CoA, propanoyl-CoA; β-MM-CoA, β-methylmalyl-CoA; CitMal-CoA, citramalyl-CoA; Pyr, pyruvate; MM-CoA, methylmalonyl-CoA; Succ-CoA, succinyl-CoA; Glyox, glyoxylate; 4HB-CoA, 4-hydroxybutyrate; AcAc-CoA, acetoacetyl-CoA; R15P, ribulose-1,5-bisphosphate; 3PG, 3-phosphoglycerate; 13BPG, 1,3-bisphophoglycerate; GAP, glyceraldehyde-3-phosphate; R5P, ribulose-5-phosphate; H6P, hexulose-6-phosphate; OH-Pyr, hydroxypyruvate; 2-PG, 2-phosphoglycerate; 10-CHO-THF, 10-formyltetrahydrofolate; CH2-THF, 5,10-methylenetetrahydrofolate; LP-S_2_, [glycine-cleavage complex H protein]-N6-lipoyl-L-lysine; LP-S-CH_2_NH_2_, [glycine-cleavage complex H protein]-S-aminomethyl-N6-dihydrolipoyl-L-lysine; LP-SH, [glycine-cleavage complex H protein]-dihydrolipoyl-L-lysine.

We included three natural occurring, oxygen insensitive CO2-fixing pathways: The Calvin-Benson-Bassham cycle (CBB cycle), the Hydroxypropionate Bicycle (HP Bicycle) and the 3-Hydroxypropionate/4-Hydroxybutyrate Cycle (3-HP/4-HB cycle). For the 3-HP/4-HB Cycle, two different variants have been described (Könneke et al., 2014), both of which were integrated. They were designated as Crenarchaeal or Thaumarchaeal, respectively. Additionally, two C1-fixing pathways were added to the SBtab model file: The Serine Cycle that is known for its efficient assimilation of formate and the Ribulose-Monophosphate Cycle (RuMP cycle) that uses methanol as the main substrate. The RuMP cycle exists in various variants. We integrated the most efficient one that is using a NAD-dependent methanol dehydrogenase as found in the thermophilic *Bacillus methanolicus*.

Apart from the natural pathways, two artificial pathways were also included in the SBtab files: The CETCH Cycle and the linear Reductive Glycine Pathway (rGlyP). Although there is a plethora of artificial pathways that have been proposed in the past, these two pathways have very favorable characteristics: The CETCH cycle is one of the few artificial CO2-fixing pathway that have been implemented in vitro or in vivo (Schwander et al., 2016), the other being the recently presented Gnd– Entner–Doudoroff cycle (GED cycle) (Satanowski et al., 2020) which can be seen as a variant of the CBB cycle. The GED pathway relies on the phosphogluconate dehydrogenase that will only operate with high CO2 concentrations, is not substantially faster than RuBisCO (especially RuBisCO variants of C4 plants or cyanobacteria with higher *K*_*m*_ for CO2) and the overall pathway stoichiometry is identical to the CBB cycle. Still the GED cycle might bear great potential in the future if more efficient variants of Phosphogluconate dehydrogenase can be found or engineered as it avoids photorespiration. In this study, however, it was not considered. The CETCH cycle on the other hand can operate at lower CO2 concentrations and might theoretically be faster and more efficient than the CBB cycle (Schwander et al., 2016).

The other artificial pathway from the literature that was included in this study, the Reductive Glycine Pathway is the only artificial C1-fixing pathway that has been implemented in vivo to support growth on formate as the sole carbon and energy source and is also the potentially most efficient pathway to support formatotrophic growth. Exotic pathways that rely on high concentrations of formaldehyde (Chou et al., 2019; He et al., 2020; Siegel et al., 2015) and pathways that are variations of existing pathways (Yang et al., 2019; Yu & Liao, 2018) were also not considered.

The detailed pathway maps that were integrated here, as well as the corresponding metabolites and enzymes, the respective kinetic data and their literature sources can be found in the supplementary information in supplementary Figures S1-8 and Table S1.

### Design of new pathways related to the reverse TCA and integration into the SBtab model file

In addition to pathways that were described by other authors, we also designed two completely new pathways that are insensitive to oxygen and show superior efficiency: The reductive Citramalyl-CoA Cycle (rCCC) and the 2-Hydroxyglutarate-reverse TCA (2-HG-rTCA) that are depicted in detail in Figure 3. Both pathways share a common structure with the reverse TCA cycle that is regarded as one of the most efficient pathways for CO2 fixation (Bar-Even et al., 2010; Siegel et al., 2015), but is limited to a very specialized group of anaerobic or microaerophilic organisms. The rCCC is a composite of existing pathways: it combines one part of the reverse TCA (from oxaloacetate to succinyl-CoA), one part of the CETCH Cycle (from succinyl-CoA to mesaconyl-C1-CoA) and one part of the HP Bicycle (from mesaconyl-C1-CoA to pyruvate and acetyl-CoA). The cycle is closed by pyruvate carboxylase, a carboxylating enzyme with high activity that is supposed to be superior to RuBisCO (Bar-Even et al., 2010; Cotton et al., 2018). The cycle only needs 3 ATP to produce 1 acetyl-CoA and is thus even more efficient than the Thaumarchaeal 3-HP/4-HB cycle that was described as the most ATP-efficient aerobic CO2-fixing pathway (Könneke et al., 2014) and needs 4 ATP to produce 1 acetyl-CoA. The 2-HG-rTCA, the second pathway that was designed, has an even higher overlap with the reverse TCA (from ketoglutarate to succinyl-CoA in the reductive direction). Hence, the only reaction that is missing from the rTCA is the ferredoxin dependent 2-oxoglutarate synthase which is also the only strictly oxygen-sensitive enzyme of the cycle. This carboxylating enzyme was replaced in the 2-HG-rTCA by an enzyme that extends the C4-carboxylic acid not by CO2, but with formyl-CoA in the following fashion:

**Figure 3:**
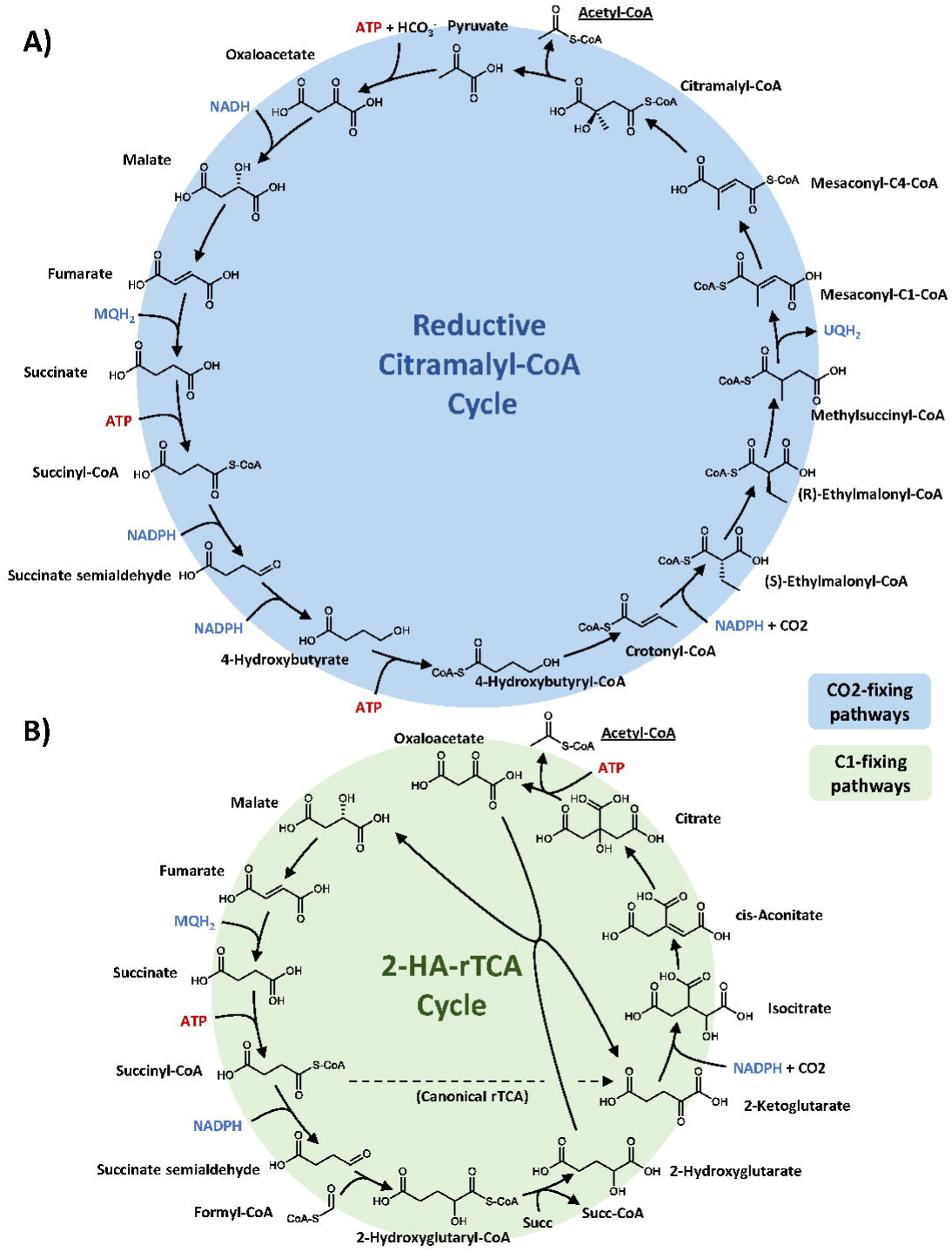
Overview of the reductive Citramalyl-CoA Cycle (rCCC) and the 2-Hydroxyglutarate-reverse TCA cycle (2-HG-rTCA Cycle) that were designed in this work. For simplification, reaction arrows can include multiple reactions and skip some metabolites. ADP, AMP, phosphate, water, and oxidized forms of electron carriers were also left out to improve clarity. Abbreviations: Succ-CoA, succinyl-CoA; succ, succinate.

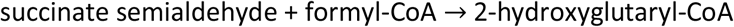

This reaction is catalyzed by a 2-hydroxyacyl-CoA lyase, an enzyme that has recently gained attention as an enzyme to assimilate C1-bodies (Chou et al., 2019). In their original work, Chou *et al*. use this enzyme to extend formaldehyde with formyl-CoA. Beside the high toxicity of formaldehyde, the enzyme also has a very high *K*_*m*_ for this substrate, making it less than ideal. Longer chain length aldehydes show lower *K*_*m*_ values which help to limit toxic aldehyde concentrations. We assumed the *K*_*m*_ value and turnover number for succinate semialdehyde to be the geometric mean of propanal and pentanal since succinate semialdehyde has 4 carbon atoms. For future applications, this is probably a conservative estimate since the enzyme was not yet engineered for the shorter chained substrates and is actually supposed to catalyze the reverse reaction. So higher affinities and turnover numbers for succinic semialdehyde should be possible. In the 2-HG-rTCA cycle, 2-hydroxyglutaryl-CoA is then supposed to transfer its CoA group to an acceptor carboxylic acid. Here, it was assumed to be transferred to succinate. For the kinetics, it was estimated that similar kinetic parameters as determined for succinyl-CoA:malate CoA-transferase can be achieved. In a recent study, it could be demonstrated that highly active 2-hydroxyacyl-CoA transferases can be found in nature (Zhang et al., 2019), so from a design perspective, this should not be a problematic reaction step. Next, 2-hydroxyglutarate is oxidized to 2-ketoglutarate. This thermodynamically difficult step (at least with NAD^+^ as the oxidizing agent) is catalyzed by the lactate-malate transhydrogenase that also accepts 2-hydroxyglutarate as a substrate, with a lower specificity and rate, however. Again, this is a conservative estimation as the enzyme has not yet been optimized for the substrates. From 2-ketoglutarate on, the pathway follows the traditional reverse TCA.

The detailed pathway maps, as well as the corresponding enzymes, the respective kinetic data and their sources can be found in the supplementary information in Figure S4 and Table S1.

### Design and integration of a new phosphatase-less Calvin-Benson-Bassham cycle variant

In order to analyze and to understand the natural design of the CBB cycle, a hypothetically augmented version of the pathway was designed that uses 2 mol ATP less for the production of one mol C3 sugar. This is achieved by replacing the irreversible phosphatase reactions with phosphotransferase reactions in the following fashion:

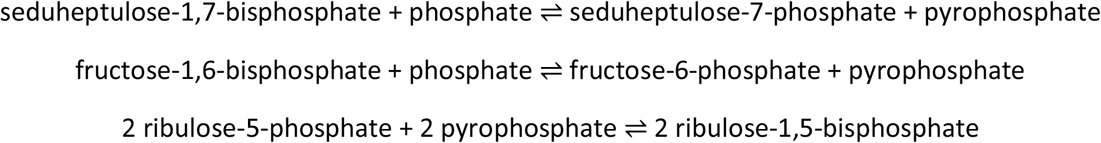

This net phosphate transfers are exergonic under standard conditions with Δ_r_G’° = −11±4 kJ/mol for the transfer of phosphate from seduheptulose-bisphosphate to ribulose-5-phosphate, and with Δ_r_G’° = − 4±5 kJ/mol for transfer of phosphate from fructose-bisphosphate to ribulose-5-phosphate. A simplified pathway map of this imaginary cycle is illustrated in Figure 4 and a more detailed map specifying the enzymes for each reaction can be found in the supplementary information in Figure S5.

**Figure 4:**
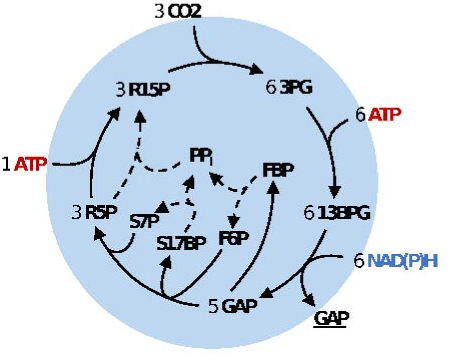
Overview of the phosphatase-less Calvin-Benson-Bassham Cycle variant that was designed in this work. Dashed lines represent the phosphotransferase reactions that were added which replace the phosphatase reaction and transfer 2 phosphate groups to ribulose-5-phosphate (R5P). For simplification, reaction arrows can include multiple reactions and skip some metabolites. ADP, AMP, phosphate, water, and oxidized forms of electron carriers were also left out to improve clarity. Abbreviations: R15P, ribulose-1,5-bisphosphate; 3PG, 3-phosphoglycerate; 13BPG, 1,3-bisphosphoglycerate; GAP, glyceraldehyde-3-phosphate; FBP, fructose-1,6-bisphosphate; F6P, fructose-6-phosphate; S17BP, seduheptulose-1,7-bisphosphate; PP_i_, pyrophyosphate; S7P, seduheptulose-7-phosphate; R5P, ribulose-5-phosphate.

Enzymes were found that are able to catalyze the described phosphate transfer reactions via pyrophosphate (Reshetnikov et al., 2008) in a promiscuous fashion. Enzymes that can directly transfer the phosphate group from one sugar to the next might also appear in nature and thus might avoid regulatory problems associated with pyrophosphate.

### Connecting products of pathways with each other in a modular fashion

The primary products of the pathways presented in the last sections are different for each pathway:

- The 3-HP/4-HB Cycle, the rCCC, the 2-HG-rTCA and the Serine Cycle produce acetyl-CoA
- The CBB cycle and RuMP Cycle produce glyceraldehyde-3-phosphate
- The HP Bicycle and reductive Glycine Pathway produce pyruvate
- The CETCH Cycle produces glyoxylate as the primary product

This is a challenge for the comparison of the pathways as it is unclear how the “value” of a produced metabolite relates to another. To solve this, pathways were added to the SBtab model file that connect the primary products with each other. These additional connecting pathways will be called “connecting modules” as the can be modularly combined with the original primary pathways. The connecting modules that were implemented are described in the following sections and summed up with their net stoichiometries in Table 2. Again, the respective reactions, their stoichiometries and kinetic parameters were added to the SBtab model and data files. Some of these connections are obvious at first: Pyruvate can be turned into glyceraldehyde-3-phosphate via gluconeogenesis, for instance. Glyceraldehyde-3-phosphate can inversely be converted back to pyruvate or oxaloacetate via the last steps of glycolysis.

**Table 2:** Modules connecting pathways that were implemented in the SBtab model file

| Originating Pathway<sup>e</sup> | Stoichiometry (main substrates underlined, main products bold) | $\Delta_r G^0$, kJ/mol |
| --- | --- | --- |
| **Glyoxylate converting modules** |  |  |
| β-Hydroxyaspartate cycle | $2 \text{ Glyoxylate} + \text{NADH} \rightarrow \text{Oxaloacetate} + \text{NAD}^+$ | $-71 \pm 6$ |
| HP-Bicycle (modified) | $\text{Glyoxylate} + 3 \text{ NADPH} + 2 \text{ HCO}_3^- + \text{ATP} \rightarrow \text{Oxaloacetate} + 3 \text{ NADP}^+ + \text{ADP} + \text{P}_i$ | $-28 \pm 13$ |
| Reverse Glyoxylate Shunt | $\text{Glyoxylate} + \text{ATP} + \text{MQH}_2 + \text{NADH} + \text{CoA} + 2 \text{ H}^+_{\text{in}} \rightarrow \text{Acetyl-CoA} + \text{ADP} + \text{P}_i + \text{MQ} + \text{NAD}^+ + 2 \text{ H}^+_{\text{out}}$ | $-15 \pm 7^a$ |
| Serine Cycle | $\text{Glyoxylate} + \text{Formate} + \text{NADH} + \text{NADPH} + \text{HCO}_3^- + 2 \text{ ATP} \rightarrow \text{Oxaloacetate} + \text{NAD}^+ + \text{NADP}^+ + 2 \text{ ADP} + 2 \text{ P}_i$ | $-76 \pm 8$ |
| **Acetyl-CoA converting modules** |  |  |
| Ethylmalonyl-CoA pathway (modified) | $2 \text{ Acetyl-CoA} + 2 \text{ NADPH} + 0.5 \text{ NAD}^+ + \text{UQ} + \text{MQ} + \text{HCO}_3^- + \text{CO}_2 + 2 \text{ H}^+_{\text{out}} \rightarrow 1.5 \text{ Oxaloacetate} + 2 \text{ NADP}^+ + 0.5 \text{ NADH} + \text{UQH}_2 + \text{MQH}_2 + 2 \text{ CoA} + 2 \text{ H}^+_{\text{in}}$ | $-65 \pm 13^a$ |
| HP/HB-cycle (modified)<sup>b</sup> | $\text{Acetyl-CoA} + 3 \text{ ATP} + 3 \text{ NADPH} + \text{NAD}^+ + \text{MQ} + 2 \text{ HCO}_3^- + 2 \text{ H}^+_{\text{out}} \rightarrow \text{Oxaloacetate} + 3 \text{ ADP} + 3 \text{ P}_i + 3 \text{ NADP}^+ + \text{NADH} + \text{MQH}_2 + \text{CoA} + 2 \text{ H}^+_{\text{in}}$ | $-101 \pm 14^a$ |
| Red. Citramalyl/Ethyl-malonyl-CoA pathway | $\text{Acetyl-CoA} + \text{HCO}_3^- + 2 \text{ NADPH} + \text{UQ} \rightarrow \text{Pyruvate} + 2 \text{ NADP}^+ + \text{UQH}_2 + \text{CoA}$ | $-39 \pm 9$ |
| **Pyruvate converting modules<sup>c</sup>** |  |  |
| 4-HB pathway<sup>b</sup> (+Pyruvate carboxylase) | $\text{Pyruvate} + 2 \text{ CoA} + 3 \text{ ATP} + \text{HCO}_3^- + \text{MQH}_2 + 2 \text{ NADPH} + 2 \text{ H}^+_{\text{in}} \rightarrow 2 \text{ Acetyl-CoA} + 3 \text{ ADP} + 3 \text{ P}_i + \text{MQ} + 2 \text{ NADP}^+ + 2 \text{ H}^+_{\text{out}}$ | $-42 \pm 10$ |
| MCG-like cycle<sup>d</sup> (+Pyruvate carboxylase) | $\text{Pyruvate} + 3 \text{ ATP} + \text{HCO}_3^- + 2 \text{ CoA} + 3 \text{ NADH} \rightarrow 2 \text{ Acetyl-CoA} + 3 \text{ ADP} + 3 \text{ P}_i + 3 \text{ NAD}^+$ | $-115 \pm 8$ |
| **General modules** |  |  |
| Glycolysis (starting from GAP) | $\text{Glyceraldehyde-3-phosphate} + \text{ADP} + \text{NAD(P)}^+ + \text{HCO}_3^- \rightarrow \text{Oxaloacetate} + \text{NAD(P)H} + \text{ATP}$ | $-49 \pm 6$ |
| Gluconeogenesis (starting from oxaloacetate) | $\text{Oxaloacetate} + 2 \text{ ATP} + \text{NADH} \rightarrow \text{Glyceraldehyde-3-phosphate} + 2 \text{ ADP} + \text{P}_i + \text{NAD}^+ + \text{CO}_2$ | $28 \pm 6$ |

<sup>a</sup> the free energy was calculated assuming a transmembrane potential of -150 mV
<sup>b</sup> this module exists in two variants (like the pathway it is derived from), depending on whether a AMP or ADP producing 3-Hydroxypropyl-CoA/4-Hydroxybutyryl-CoA synthase is used or not. In the here presented form it corresponds the Crenarchaeal form. Both variants were considered, however, for the following analyses
<sup>c</sup> in addition to the modules listed here, single reactions such as the pyruvate dehydrogenase complex or the pyruvate carboxylase were also used to connect metabolites
<sup>d</sup> this cycle has the same working principle as the malyl-CoA-glycerate cycle (MCG cycle) presented in (Yu et al., 2018). Instead of turning 2 glyoxylate into glycerate which are then converted to oxaloacetate, this cycle directly uses the β-Hydroxyaspartate cycle to turn 2 glyoxylate into oxaloacetate. A detailed pathway map can be found in supplementary Figure S8B.
<sup>o</sup> original pathway in this context relates to the pathway in which the respective enzymes are usually found. Many of the connecting modules are subnetworks of the full, original pathways.

For other primary pathway products, the situation is more difficult, however: Assimilation of acetyl-CoA to C4 dicarboxylic acids is done by many organisms with the Glyoxylate Shunt that follows the stoichiometry:

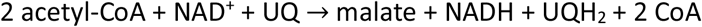

where UQ is ubiquinone and UQ_2_ is ubiquinol.

This pathway is oxidative (i.e. electrons are freed) which is contrary to the reductive nature of carbon fixation. Besides, for gluconeogenesis two equivalents of CO2 are produced per molecule of formed C6 sugar using the Glyoxylate Shunt. This is why some bacteria use different pathways for the assimilation of acetyl-CoA that are reductive and even fix CO2, like the Ethylmalonyl-CoA pathway. Bacteria with the 3-HP/4-HB pathway can turn acetyl-CoA to succinyl-CoA with the 3-HP route that is an intrinsic part of their cycle. In case of the rCCC, a part of the pathway can be used for the conversion of acetyl-CoA to pyruvate if 3 reactions from the Ethylmalonyl-CoA pathway are added as showcased in Figure 5.

**Figure 5:**
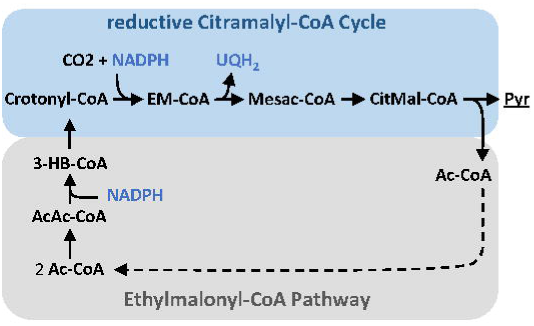
Design of a pathway to convert acetyl-CoA (Ac-CoA) to pyruvate (Pyr) using parts of the reductive Citramalyl-CoA Cycle (blue) and the Ethylmalonyl-CoA Pathway (grey). The dashed arrow denotes the possible recycling of acetyl-CoA for the first reaction of the Ethylmalonyl-CoA pathway. Abbreviations: EM-CoA, ethylmalonyl-CoA; Mesac-CoA, mesaconyl-CoA; CitMal-CoA, citramalyl-CoA; Ac-CoA, acetyl-CoA; Pyr, pyruvate; AcAc-CoA, acetoacetyl-CoA; 3-HB-CoA, 3-hydroxybutyryl-CoA.

This pathway module does not need ATP. Following the calculation of projected ATP costs presented in Material and Methods, the net transfer of 2 electrons from NADPH to ubiquinone comes at a projected cost of 1 ATP, however. This is on par with the most efficient modules introduced by Bar-Even *et al*. (2010) in their seminal work almost a decade ago as part of the MOG pathways. In contrast to their approach that uses an oxygen-sensitive aminomutase or acrylyl-CoA hydratase, the design presented in Figure 5 is assembled purely from oxygen-insensitive reactions that appear in actual carbon fixation pathways in nature.

For the conversion of glyoxylate to other metabolites, microorganisms employ different strategies: glyoxylate can be turned to glycine and then sequestered by the glycine cleavage system to finally yield glycerate (Douce et al., 2001), alternatively 2 mol of glyoxylate can be converted to 1 mol of glycerate and CO2 with tartronate-semialdehyde as an intermediate (Bar-Even et al., 2010; Grostern et al., 2012). Recently, it was found that glyoxylate can also be sequestered to oxaloacetate by the β-Hydroxyaspartate Cycle (Schada von Borzyskowski et al., 2019). Both glycerate and oxaloacetate can easily be converted to phosphoenolpyruvate at the cost of one ATP, but glycerate needs one ATP to react to oxaloacetate (via 2-phosphoglycerate). Therefore, only the β-Hydroxyaspartate cycle was implemented in the SBtab model file. In addition to natural pathways, the reverse Gloxylate Shunt introduced by (Mainguet et al., 2013) was also implemented as a potentially very efficient pathway that turns glyoxylate to acetyl-CoA.

Another interesting case is the conversion of pyruvate or C4 dicarboxylic acids to acetyl-CoA. Plants use the pyruvate dehydrogenase complex, an oxidative decarboxylation reaction, which is in contrast to the reductive carbon fixing CBB cycle. To capitalize on this potential, it was proposed to rather split a C4 body into two C2 bodies, e.g. in the synthetic malyl-CoA-glycerate carbon fixation pathway (MCG pathway) (Yu et al., 2018). In this pathway, malate is split into acetyl-CoA and glyoxylate. Glyoxylate is then fixed to glycerate as described earlier. Finally, malate can be regenerated from glycerate. We implemented a variant of this pathway in the SBtab model file that uses the β-Hydroxyaspartate Cycle for glyoxylate assimilation instead which results in a cycle that needs 1 mol of ATP less than the original pathway per 1 mol of acetyl-CoA. A detailed map of this MCG-like pathway can be found in the supplementary information in Figure S8B. Nature also has invented a way to convert C4 dicarboxylic acids to 2 acetyl-CoA as part of the 3-HP/4-HB cycle (compare Figure 2), starting from succinyl-CoA and ending in two molecules of acetyl-CoA. Both pathway modules were added to the SBtab model file. Detailed pathway maps of the connecting modules with the respective enzymes can be found in Supplementary Figures S6-S9.

### Adjustment of the equilibrium constants of the C5 dicarboxylic acid platform

After collecting the kinetic data for all reactions described in the previous sections, we next handled the problem of high uncertainties of the kinetic parameters of some critical reactions in some of the pathways. In specific, eQuilibrator had difficulties to predict the free formation energies of certain C5 dicarboxylic acids that are crucial for some of the presented pathways. This resulted in poor prediction for the crotonyl-CoA reductase/carboxylase (Δ_r_G’° = −43±17 kJ/mol), the β-methylmalyl-CoA lyase (Δ_r_G’° = +16±16 kJ/mol) and citramalyl-CoA lyase (Δ_r_G’° = +11±15 kJ/mol) as shown in Figure 6. These reactions correspond to the in- and outlet of the C5 dicarboxylic acid platform which are thus coupled by Wegscheider conditions. In the case of β-methylmalyl-CoA lyase, a complete set of kinetic parameters was measured by Erb *et al*. (2010). The parameters could be used to estimate a new value for the equilibrium constant using the Haldane relationship (compare Equation 5) by Parameter Balancing (see Material and Methods for details). An equilibrium constant of *K*_*eq*_ = 11.8±1.5 (geometric standard deviation) was calculated by the algorithm, corresponding to a standard free reaction energy of Δ_r_G’° = +6.1±1.0 kJ/mol. Based on the measured kinetic data, this reaction is thus considerably more exergonic than predicted by eQuilibrator which is in favor of the thermodynamics for the Ethylmalonyl-CoA pathway and the CETCH Cycle that use this reaction for β-methylmalyl-CoA cleavage. If only the equilibrium constant of the β-methylmalyl-CoA lyase is changed, the network will not satisfy the Wegscheider conditions anymore. Fortunately, the C5 dicarboxylic acid platform is enclosed by a small set of reactions, so it sufficed to adjust the free energies and equilibrium constants of crotonyl-CoA reductase/carboxylase (new Δ_r_G’° = −33 kJ/mol) and citramalyl-CoA lyase (new Δ_r_G’° = +1 kJ/mol) as described in Material and Methods. The original uncertainties in these reactions were actually higher than the change applied to them which justifies the correction. For the crotonyl-CoA reductase/carboxylase, the change makes hardly any difference since the reaction stays highly exergonic. The adjustment for citramalyl-CoA lyase change the thermodynamics in favor of the Reductive Citramalyl-CoA Cycle.

**Figure 6:**
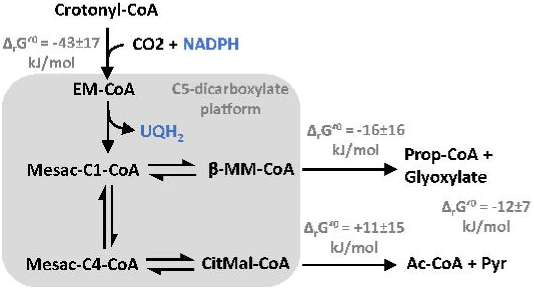
Schematic overview of the C5 dicarboxylic acid platform. The free reaction energies as predicted by eQuilibrator (pH 7.5, I = 0.25 M) are indicated for the influx and efflux reactions of the platform. Note the high uncertainties of the free energies. Additionally, the free energy of the hypothetical conversion of propionyl-CoA and glyoxylate to pyruvate and acetyl-CoA is indicated (dashed reaction arrow). Abbreviations: EM-CoA, ethylmalonyl-CoA; Mesac-C1-CoA, mesaconyl-C1-CoA; β-MM-CoA, β-methylmalyl-CoA; Prop-CoA, propanoyl-CoA; Mesac-C4-CoA, mesaconyl-C4-CoA; CitMal-CoA, citramalyl-CoA; Ac-CoA, acetyl-CoA; Pyr, pyruvate.

Figure 6 also illustrates the thermodynamics of the conversion of propionyl-CoA and glyoxylate to pyruvate and acetyl-CoA in the Hydroxypropionate Bicycle which is exergonic overall. It is also clear that when comparing the two exits of the C5 dicarboxylic acid platform, the route towards acetyl-CoA and pyruvate is more exergonic in total which gives the rCCC a slight edge over the CETCH cycle on a thermodynamic level.

### Comparison of pathway specific activities and efficiencies of CO2 and C1 fixing pathways

Many pathway designs presented in this work are supposed to be superior to previous pathways, but how can we actually predict whether or not these pathways are feasible or even better apart from their stoichiometry? To rule out that pathways are thermodynamically and kinetically not sound, the ECM algorithm was used which is able to factor in the thermodynamics and kinetics of the reactions based on all parameters that were collected. After assembly of all pathways and connecting modules, the SBtab model file comprised 96 reactions with 94 metabolites and 526 enzymatic parameters (before Parameter Balancing). The data file was then processed by Parameter Balancing to yield a complete and consistent parameter set for all reactions. Next, we evaluated the pathway specific activities of all pathways with ECM towards the production of a certain metabolite.

First, all pathways were compared for the production of glyceraldehyde-3-phosphate as shown in Figure 7. The “(w/o SHMT)” label means that the serine hydroxymethyltransferase was not included when showing the pathway specific activity. As already noted by others (Šmejkalová et al., 2010), the kinetics of this enzyme seem to be estimated inaccurately by the usual assays which is why the enzyme accounted for a major fraction of enzyme demand in the respective pathways. The pathway specific activity of the CBB cycle variants was in the order of 0.25 µmol min^-1^ mg^-1^ which is in line with the estimation by (Bar-Even et al., 2010). All pathways were predicted to have pathway activities in the same order of magnitude except the RuMP cycle which is especially low, and the rGlyP pathway (w/o SHMT) which is exceptionally high. The main problem of the RuMP cycle is the methanol dehydrogenase reaction which is thermodynamically and kinetically very unfavorable (compare also Figure 8). Also, the forward rate constant at a temperature of 37°C was taken which is a lot lower than the temperature of its native host (Brautaset et al., 2007). Apart from the unfavorable kinetics of the methanol dehydrogenase, the RuMP cycle was only thermodynamically feasible if the upper and lower bounds for NAD/NADH and NADP/NADPH were extended further by a factor of 10. It remains enigmatic how microorganisms like *Bacillus methanolicus* run this cycle, but the high temperature requirement might play a role here.

**Figure 7:**
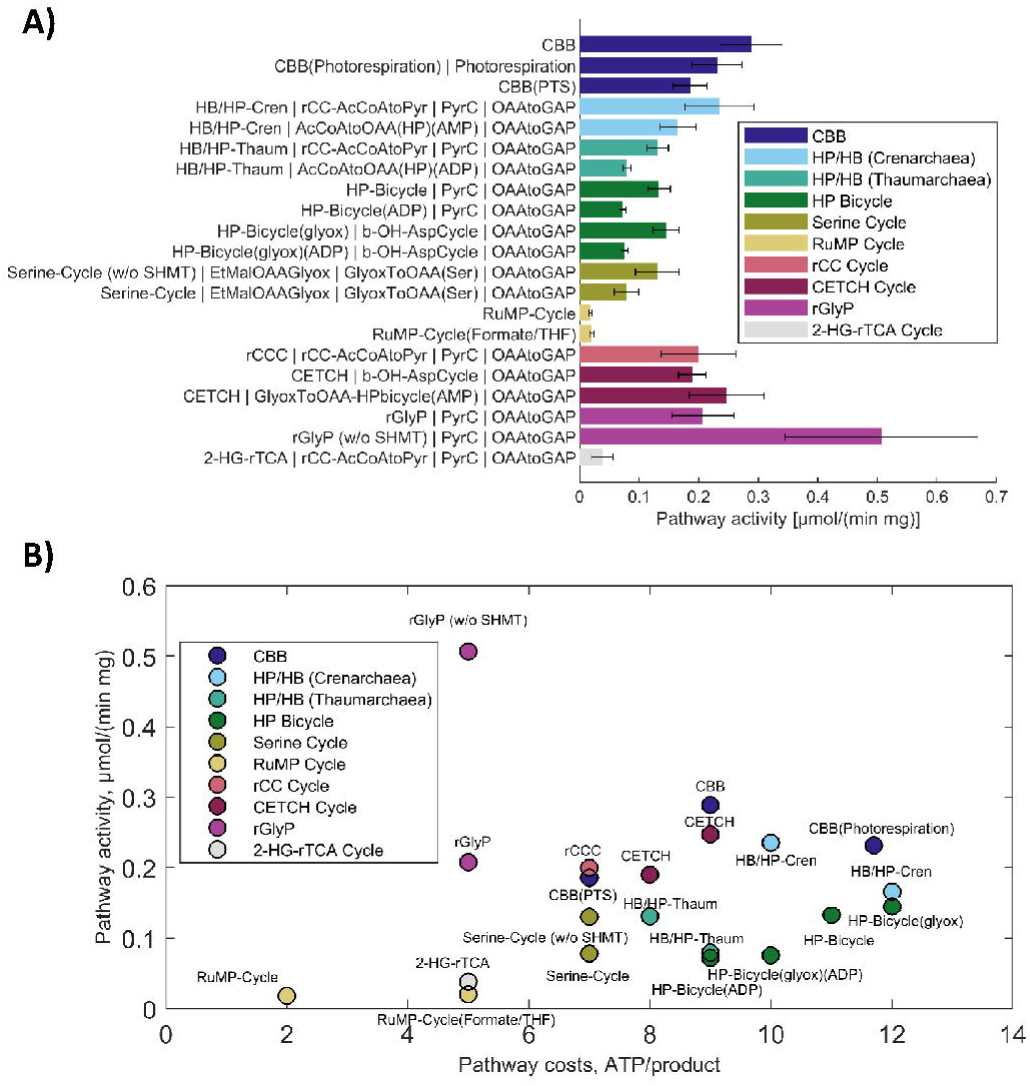
Comparison of natural and artificial carbon fixation pathways with the Enzyme Cost Minimization algorithm. The final product of choice was glyceraldehyde-3-phosphate in this case. (w/o SHMT) indicates that the cost of the serine hydroxymethyltransferase are not included in the diagram. The concentration of CO2 was assumed to not exceed 10 µM, corresponding approximately to air-saturation. **A)** Pathway specific activities with standard deviations and the full name of the main pathways and the connecting modules to transform their primary product to glyceraldehyde-3-phosphate. **B)** Pathway specific activities compared to the projected ATP costs of the pathway to produce glyceraldehyde-3-phosphate. The data labels only specify the main pathway because of limited space.

**Figure 8:**
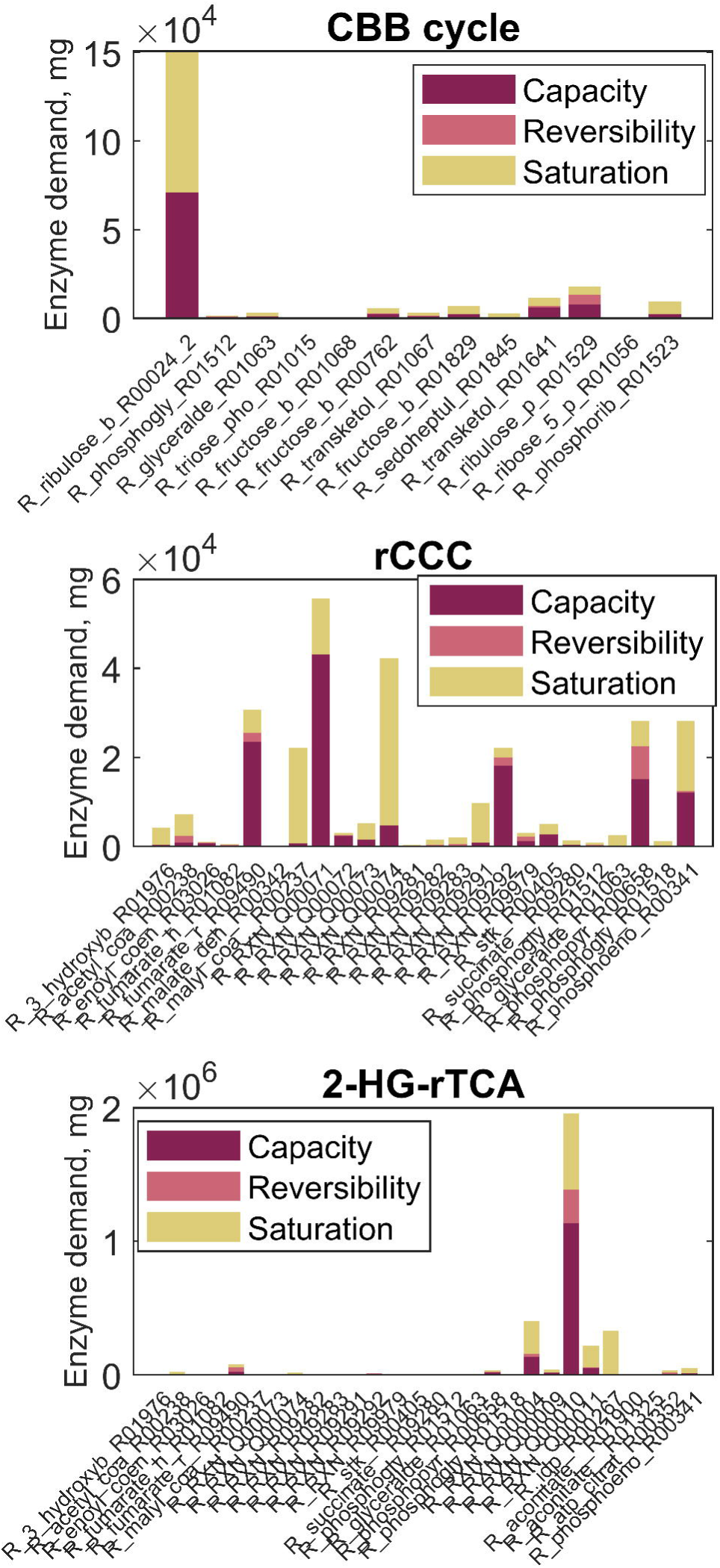
Enzyme demands to sustain a total pathway activity of 1 mmol product per second. Contribution of the capacity, the reversibility and the saturation with substrates or products of each reaction to the demand of enzymes for the CBB cycle, the rCCC and the 2-HG-rTCA. The figure follows the wording of (Noor et al., 2016). “Capacity”: demand of enzyme caused by a limitation by the catalytic rate constant. “Reversibility”: extra amount of enzyme needed because of a backward flux. “Saturation”: additional enzyme necessary because of undersaturation with a substrate or oversaturation with a product. The values present the optimized state as predicted by the ECM algorithm assuming a CO2 concentration of 10 µM. Glyceraldehyde-3-phosphate was chosen as a product for the cycles. Reaction names correspond to their identifiers in the SBtab model file (Supplementary file Reactions_Composite17_model.tsv)

The rGlyP (w/o SHMT) on the other hand showed a pathway specific activity around twice as high as the next best pathway, albeit with high uncertainty. This can be attributed to the small number of reactions steps and the favorable thermodynamics and kinetics of the rGlyP although it has to be pointed out that the kinetic parameters from the literature were mainly based on activity measurements with unnatural substrates. Kinetic data of the assembled glycine cleavage system, especially in the carboxylating direction is sparse. Additionally, eQuilibrator had trouble to estimate the free energy of the carboxylation reaction that is catalyzed by the P-protein (Δ_r_G’° = −17.3±15.9 kJ/mol):

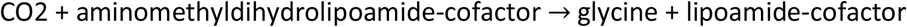

As this is one of the key reactions and we also did not find a value for the forward kinetic constant, this standard deviation has a crucial impact on the total estimation of pathway specific activity. *In vivo* the rGlyP will only work with high CO2 concentration (Claassens et al., 2020; Kim et al., 2020) which also does not match with the results of the analysis here. All in all, the estimation for the rGlyP are therefore not completely trustworthy which stresses how urgently kinetic data for this system is needed to make accurate predictions.

The 2-HG-rTCA cycle in conjunction with the acetyl-CoA conversion module from the rCCC has a very low requirement of 3 ATP per product equal to the rGlyP. Although it has a lower pathway activity, it is still promising, considering that conservative assumptions were made about the kinetic parameters of the enzymes. Comparing the CO2-fixing pathways, the rCCC, the CETCH cycle and the Crenarchaeal 3-HB/4-HB cycle showed pathway specific activities equal to the CBB cycle with 10% of photorespiration, but with a lower ATP demand per product.

Looking at the single contributions of the reactions to the enzyme demand, the Enzyme Cost Minimization algorithm makes consistent prediction as can be seen in Figure 8. For the CBB cycle, RuBisCO (“R_ribulose_b_R00024_2”) is the enzyme with the highest demand in the cycle where half of the demand is due to the relatively low turnover number (represented by “Capacity”) and half is caused by undersaturation with CO2 (represented by “Saturation”). In comparison to the CBB, the new rCCC and the 2-HG-rTCA cycle (Figure 8) do not have a single limiting enyzme, but the enzyme demand is more evenly distributed among the enzymes. For the rCCC the enzyme with the highest demand is the 4-hydroxybutyrate---CoA ligase (ADP-forming) with the number Q00071. This enzyme is supposed to be slower and thermodynamically more challenging than the AMP-forming variant (Könneke et al., 2014) and is thus likely to be a bottleneck for the pathway. Considering the 2-HG-rTCA cycle, the enzyme with the highest demand is the succinyl-CoA:2-hydroxyglutarate CoA-transferase (identifier Q00010).

As noted before, this enzyme activity is just a side-reaction of the succinyl-CoA:(S)-malate CoA-transferase and thus with a more specific enzyme, the pathway activity will be higher. In the 2-HG-rTCA cycle, the isocitrate dehydrogenase (carboxylating) is severly limited by CO2 (K_m_ of 1.3 mM) and is therefore the second costliest enzyme in the cycle which could be solved in a bioreactor fed with a higher CO2 concentration. Taken together, the estimates for the new pathways show a lot of potential for the new artificial pathways. The contributions of each reaction on the other pathways can be found in the supplementary information in Figures S10-S16.

Glyceraldehyde-3-phosphate is the main product of the CBB cycle which is why this pathway was likely favored when choosing it for comparison. Therefore, the same analysis was repeated using acetyl-CoA as the product of choice which is shown in Figure 9. As expected, the acetyl-CoA producing pathways performed better in this case compared to the glyceraldehyde-3-phosphate producing ones (mainly the CBB cycle since the RuMP cycle again showed low activity).

**Figure 9:**
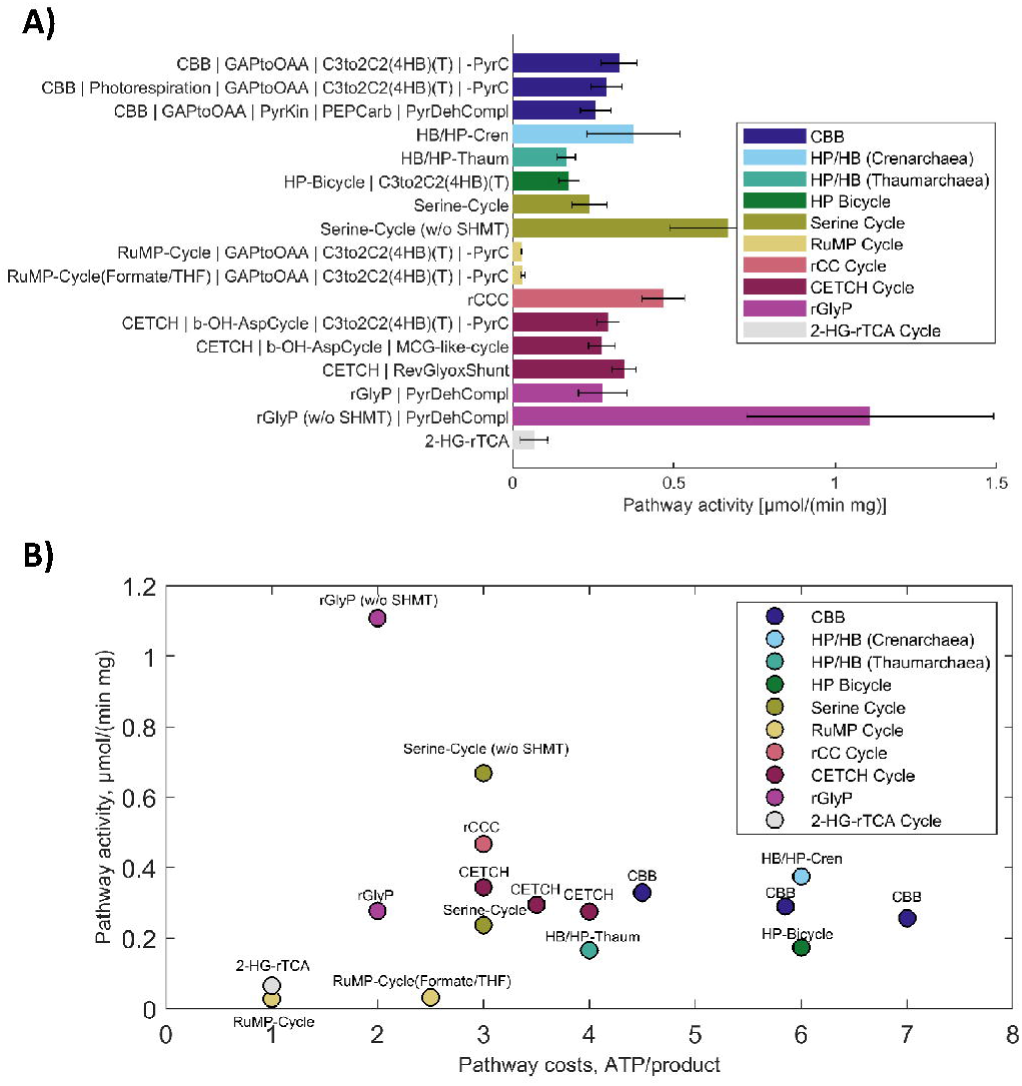
Comparison of natural and artificial carbon fixation pathways with the Enzyme Cost Minimization algorithm. The final product of choice was acetyl-CoA in this case. (w/o SHMT) indicates that the cost of the serine hydroxymethyltransferase are not included in the diagram. The concentration of CO2 was assumed to not exceed 10 µM, corresponding approximately to air-saturation. **A)** Pathway specific activities with standard deviations and the full name of the main pathways and the connecting modules to transform their primary product to acetyl-CoA. **B)** Pathway specific activities compared to the projected ATP costs of the pathway to produce acetyl-CoA.

The general trend of pathway specific activities is not influenced by the choice of product: Again, the rCCC and CETCH cycle outperformed the CBB cycle in terms of efficiency and activity. The rGlyP again showed the highest and the RuMP Cycle the lowest activity.

Since the CBB cycle is mainly limited by RuBisCO, lowering the CO2 concentration has a dramatic impact on its pathway specific activity which is less of a concern in alternative CO2-fixing pathways (compare Supplementary Figures S10-S16). Because of diffusional limitations, the steady state CO2 concentrations in the cell will be lower than under air saturation. Consequently, alternative pathways are especially better under low CO2 concentrations and the comparisons here are slightly in favor of the CBB cycle when working with the CO2 concentration of air.

## Discussion

### Artificial pathways are predicted to outperform natural ones

In the simulations described in the last section, it could be clearly seen that the artificial carbon fixing pathways have both higher activity and are more energy efficient: The rCCC and CETCH Cycle outperform the CBB cycle and the rGlyP is superior to the Serine Cycle. Again, the results of the rGlyP have to be interpreted with caution as critical kinetic parameters were not available for this pathway. Therefore, these artificial pathways have high potential for the implementation in living organisms to create chassis organisms for the fixation of CO2 and C1-compounds in Industrial Biotechnology. The CO2 fixing pathways might also be implemented in crop plants and thus boost their productivity which can be regarded as the Holy Grail of plant biotechnology. To achieve this, the plant and chloroplast genome would have to be radically changed on a metabolic and regulatory level which as of today, seems out of range of the methods of Synthetic Biology. First applications will consequently rather be found be in microorganisms that can be manipulated easier and cultivated under controlled conditions.

### The performance of the Calvin-Benson-Bassham cycle has been underestimated by most studies

Apart from the Reductive Glycine Pathway which was difficult to predict because of a lack of kinetic data, most pathways actually have a pathway specific activity in the same order of magnitude (compare Figure 7). This was primarily seen when glyceraldehyde-3-phosphate was chosen as the product and is in contrast to previous reports that concentrated on the weaknesses of the CBB cycle (Bar-Even et al., 2010; Claassens, 2017; Erb & Zarzycki, 2018; Stoffel et al., 2019). In fact, glyceraldehyde-3-phosphate as a product is a realistic case for comparison since plants rely on sucrose as a transport sugar. As sucrose is chemically relatively inert, it can be used in high concentrations in the conducting vascular cells of plants in which sucrose accumulates up to a concentration of one molar (Geiger, 2011). This “high voltage” power transmission of the plant can hardly be realized with fats or acids. Therefore, the advantages that other carbon fixation pathways show for the production for non-sugar products might not be relevant for plants. In addition, the CBB cycle generates sugars that can be directly used as precursor for different biomolecules, like ribose that is part DNA and RNA. It also has to be noted that only C3 photosynthesis was included in the analysis which is strongly limited by low CO2 concentration and low RuBisCO activity (Figure 8). Using CO2 concentration mechanisms and RuBisCO variants of C4 plants with a higher *K*_*m*_ and *k*_*cat*_, the pathway specific activity would be significantly higher and thus probably surpassing even the best artificial pathways. Taken together, the CBB cycle is not as bad as previously thought, according to our calculations, and there is a good reason that most of the carbon fixation on this planet is facilitated by this pathway. After all, RuBisCO might not be as “bad” as previously thought as others have already noted (Bathellier et al., 2018).

### Plant-like photosynthesis is not designed to be as ATP-efficient as possible

Although the CBB cycle has a higher demand of ATP for one mol of a specific product than other pathways, this difference is actually not very high: the free burning energy of glyceraldehyde-3-phosphate is around −1500 kJ/mol, for instance, while the energy of ATP hydrolysis in around −40 kJ/mol. So the energy that is saved by more efficient pathways might not be as important as other factors which will be discussed in the next section.

In fact, there are indications that ATP-efficiency is not a decisive factor for the design of carbon fixation in photosynthetic organisms. The phosphatase-less CBB cycle that we introduced in this work is a good example for this. With its modified reactions, the CBB cycle would save 2 ATP per formed C3 sugar. In addition, the pathway also didn’t lose much of its activity although enzymes with promiscuous activity were taken that were not specialized to run this cycle. Why has nature not made this “easy” step in evolution to optimize the CBB cycle? While it might well be possible that there are actually organisms that use this modified cycle which just haven’t been found yet, there is no evidence so far that this variant is used in photosynthetic organisms. Not having the irreversible phosphatase reactions might disrupt the regulation and/or robustness of the cycle which might be more important.

A second example that ATP-efficiency is not the main driver in the evolution of photosynthesis can be found in C4-plant metabolism. C4 plants use a CO2 concentration mechanism that gives these plants a selective advantage in certain climates (Atkinson et al., 2016). CO2 concentration in the mesophyll cells can be achieved by the actions of PEP synthase, PEP carboxylase and malate dehydrogenase (malic enzyme variant of C4 photosynthesis):

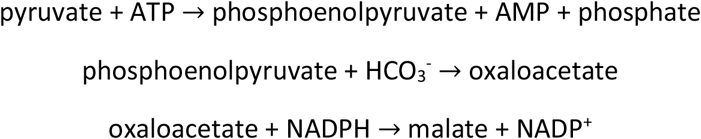

This set of reactions is quite exothermic in total with Δ_r_G’° = −47±7 kJ/mol. Malate is then shuttled to the bundle sheath cells where CO2 is released by malic enzyme and refixed by the CBB cycle. Although a total minimum of 15 ATP is necessary for the formation of one C3 sugar, C4 plants easily grow faster than C3 plants in most situations. Additionally, thinking of optimal design, the first steps in CO2 concentration mechanism could also be catalyzed by pyruvate carboxylase:

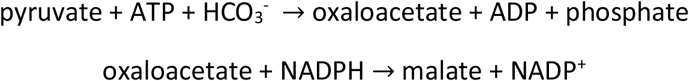

These reactions are practically irreversible (Δ_r_G’° = −35±7 kJ/mol) as well with the difference that one ATP is saved and no pyrophosphate is produced. In total, using pyruvate carboxylase instead of the natural occurring enzymes would hence save 3 ATP per C3 sugar in C4 plants. Again, this variant has not been found in nature although all enzymes exist in plants.

Therefore, it seems that plants prefer to have a slight edge in driving force or activity instead of being more efficient ATP-wise.

### Unconsidered factors in pathway comparison

Just judging the pathways by their projected ATP costs and pathways activities is like judging food just by its nutritional value and shelf price. While those are important factors they are not necessarily decisive. One unconsidered factor in this context are the vitamins: All pathways need certain cofactors to function as listed in Table 3. Interestingly, cobalamin (vitamin B12) is necessary for all CO2-fixing pathways except for the CBB cycle. Being the most structurally complex of all vitamins, B12 biosynthesis is a complex pathway that does not occur in plants and animals. In addition, it also depends on cobalt. The costs associated to its synthesis were not taken into account in this work.

**Table 3:**
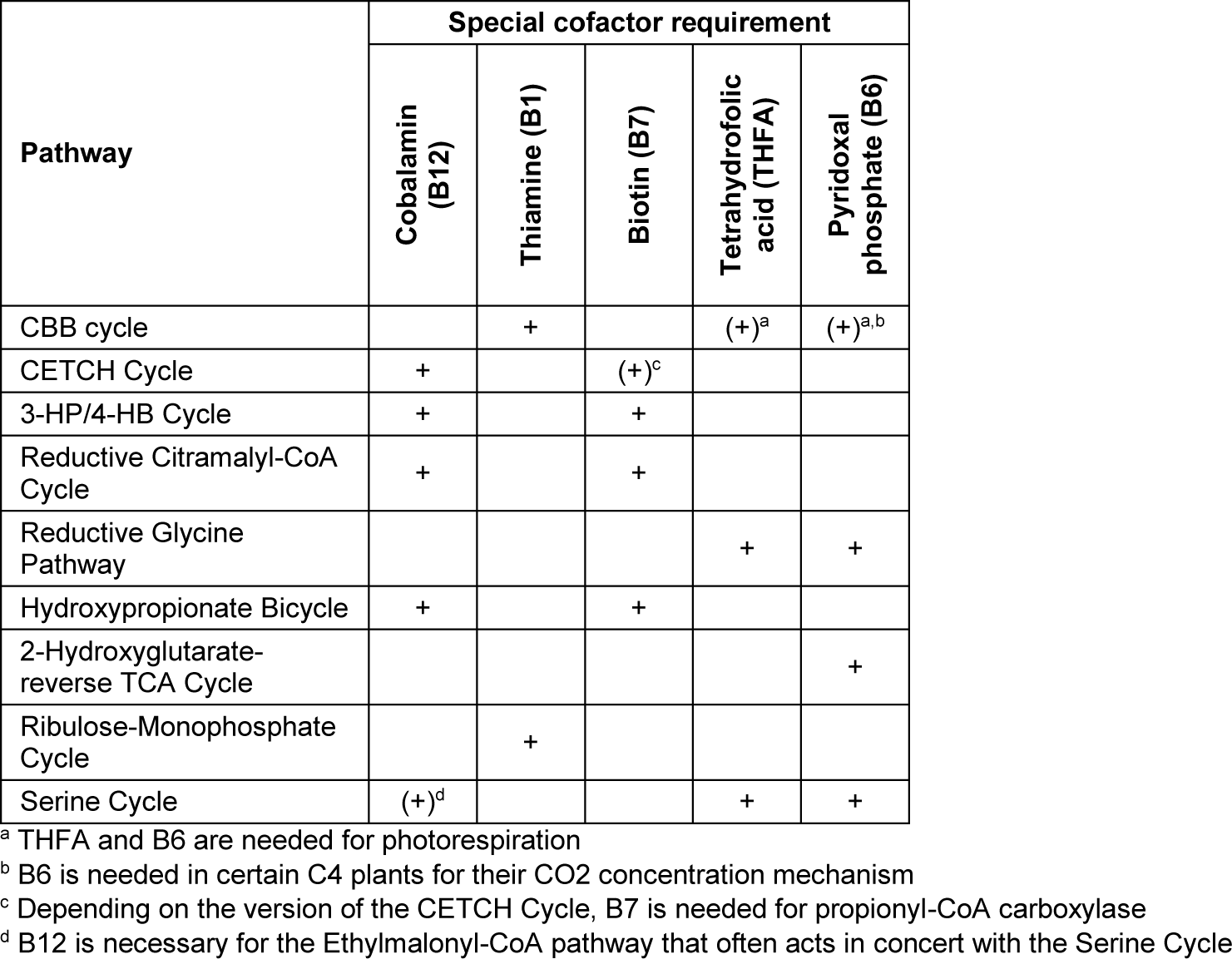
Special cofactor requirement of the different pathways

The other vitamins are structurally more compact, less demanding in their biosynthesis, independent of metal ions and are needed for many other vital reactions in the metabolism as well. In the future, this requirement could be addressed by adding B12 as a metabolite and its biosynthesis to the SBtab model file if adequate data is available. This will help to evaluate whether or not B12 requirement has a critical impact.

Other unconsidered factors in the analysis presented in this work are the controllability and robustness of the pathways. Previous works have shown that these factors are an essential part of the design principles in nature (Bromig et al., 2020; Lee et al., 2014; Whitacre, 2012). Allosteric control of enzymes was also not included in the reaction kinetics. More dedicated models and algorithms that combine pathway performance and criteria for the robustness could be established in the future. This way, good trade-offs between the two criteria could be predicted in artificial pathways, or be explained in natural occurring pathways. New efforts in the design of artificial allosteric control of enzymes (Chen et al., 2015; Cross et al., 2013) could even make possible the prediction and design of a regulatory layer on top of the reactivity layer in the network to build pathway designs that are optimal for both criteria.

So far, metabolite toxicity has also not been addressed in this work. Toxic metabolites in the pathways include for example aldehydes like glyoxylate, succinate semialdehyde, malonate semialdehyde, but also propionyl-CoA. To include this in the future, tighter constraints on these toxic compounds may be implemented, possibly based on concentrations that are actually measured within cells with a respective metabolism.

### Predictive power of the results

While the predictions made in this work match with observations of natural behavior, there were clear limitations in the purely computational approach. This mainly stems from uncertainty in the original data that was used to implement the kinetics of the reactions. eQuilibrator gave only rough estimates for some important reactions in the pathways. Moreover, for some sensitive parameters, there were only single studies available in which the enzymes were characterized. The ECM algorithm can predict, however, which reactions will be likely limiting the pathways and thereby gives good suggestions for enzymes that need to be better characterized. This stresses how important enzymology is for the understanding and the design of biochemical pathways as has also been noted by Tobias Erb recently (Erb, 2019). In fact, the predictions made in this work are only possible because of the thorough characterization of all the enzymes by others. In the era of big data, it might be even beneficial to provide the raw data of enzyme kinetics studies instead of only catalytic constants and Michaelis constants in the future. The raw data could be processed by dedicated algorithms which would improve the quality of their predictions.

Regarding the equilibrium constants, it would be very advantageous for further calculations to experimentally determine the constants of uncertain reactions in an environment that resembles physiological conditions. This data is especially needed for the glycine cleavage system and the C5 carboxylic acid platform as indicated in the results section of this work. The Parameter Balancing algorithm should be adopted to integrate omics data to make use of *in vivo* data which might be very different from *in vitro* enzyme kinetics as has been seen for the Ethylmalonyl-CoA pathway in this study. In this sense, modelling approaches that are able to quantify uncertainties and identify missing data are ideal. This way, models and simulations can be iteratively improved to get closer and closer to the biological reality and a deeper understanding of metabolism and design principles of natural pathways.

## Conclusion

In this work, artificially designed carbon fixing pathways have been subjected to a trial of fire with a model based approach by testing their thermodynamic and kinetic consistency when compared to natural pathways. Indeed, the artificial pathways performed better in terms of pathway specific activities and ATP-efficiency, underpinning their future potential for applications in biotechnology. The CBB cycle compared better with the artificial pathways than previously thought, however. We also designed two new pathways, the Reductive Citramalyl-CoA Cycle and the 2-Hydroxyglutarate-Reverse TCA Cycle, that show optimal characteristics and are good candidates to be implemented *in vivo* in the future. The improved yields and activities these pathways promise will enable the creation of highly efficient platform microorganisms for a future bio-based economy.

### CRediT authorship contribution statement

H. Löwe: Conceptualization, Methodology, Validation, Visualization, Investigation, Writing − original draft, Writing - review & editing. A. Kremling: Funding acquisition, Project administration, Supervision, Writing - review & editing.

## Supporting information

SBtab model file

SBtab data file

MATLAB files

## Acknowledgements

We like to thank Wolfram Liebermeister for the fruitful discussion and the support with setting up the MATLAB implementations of the toolboxes that were used in this study. We also thank Franziska Kratzl for her suggestions to the manuscript.

