## Supplementary Information for "In-depth computational analysis of natural and artificial carbon fixation pathways"

#### 1. Detailed pathway maps of all pathways with respective enzymes

##### 1.1. Artificial and natural CO2/C1 fixing pathways

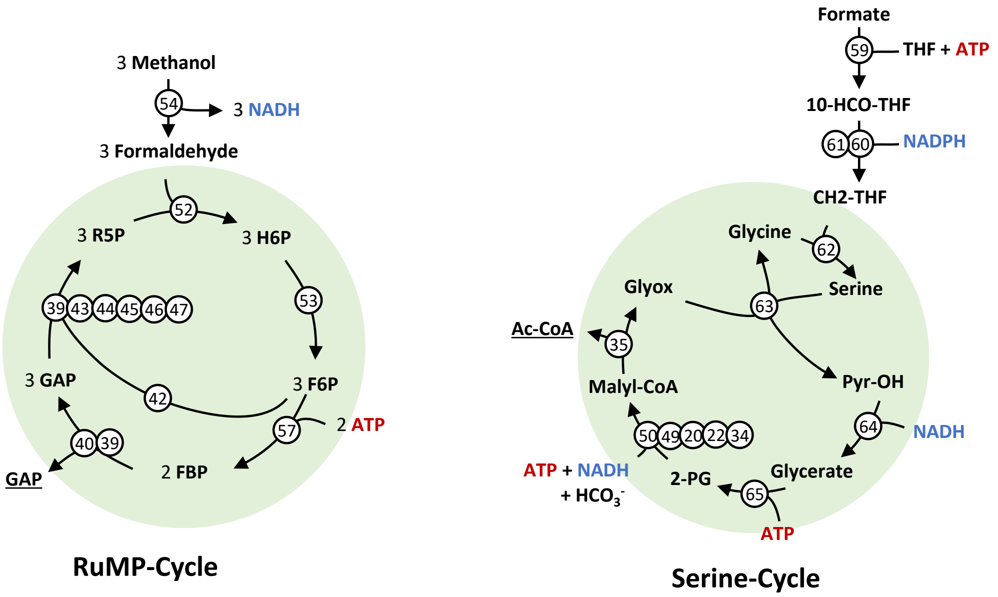


Figure S1: Flux maps of natural C1-fixing pathways, in this case methanol (RuMP-Cycle) and formate (Serine-Cycle). For simplification, reaction arrows can include multiple reactions and skip some metabolites. ADP, AMP, phosphate, water, and oxidized forms of electron carriers were also left out to improve clarity. Reactions are numbered and each number specifies the respective enzyme according to Table S1. Abbreviations: Ac-CoA, acetyl-CoA; Glyox, glyoxylate; GAP, glyceraldehyde-3-phosphate; R5P, ribulose-5-phosphate; H6P, hexulose-6-phosphate; OH-Pyr, hydroxypyruvate; 2-PG, 2-phosphoglycerate; 10-CHO-THF, 10-formyltetrahydrofolate; CH2-THF, 5,10-methylenetetrahydrofolate; LP-$S_2$, [glycine-cleavage complex H protein]-N6-lipoyl-L-lysine; LP-S-$CH_2NH_2$, [glycine-cleavage complex H protein]-S-aminomethyl-N6-dihydrolipoyl-L-lysine; LP-SH, [glycine-cleavage complex H protein]- dihydrolipoyl-L-lysine.

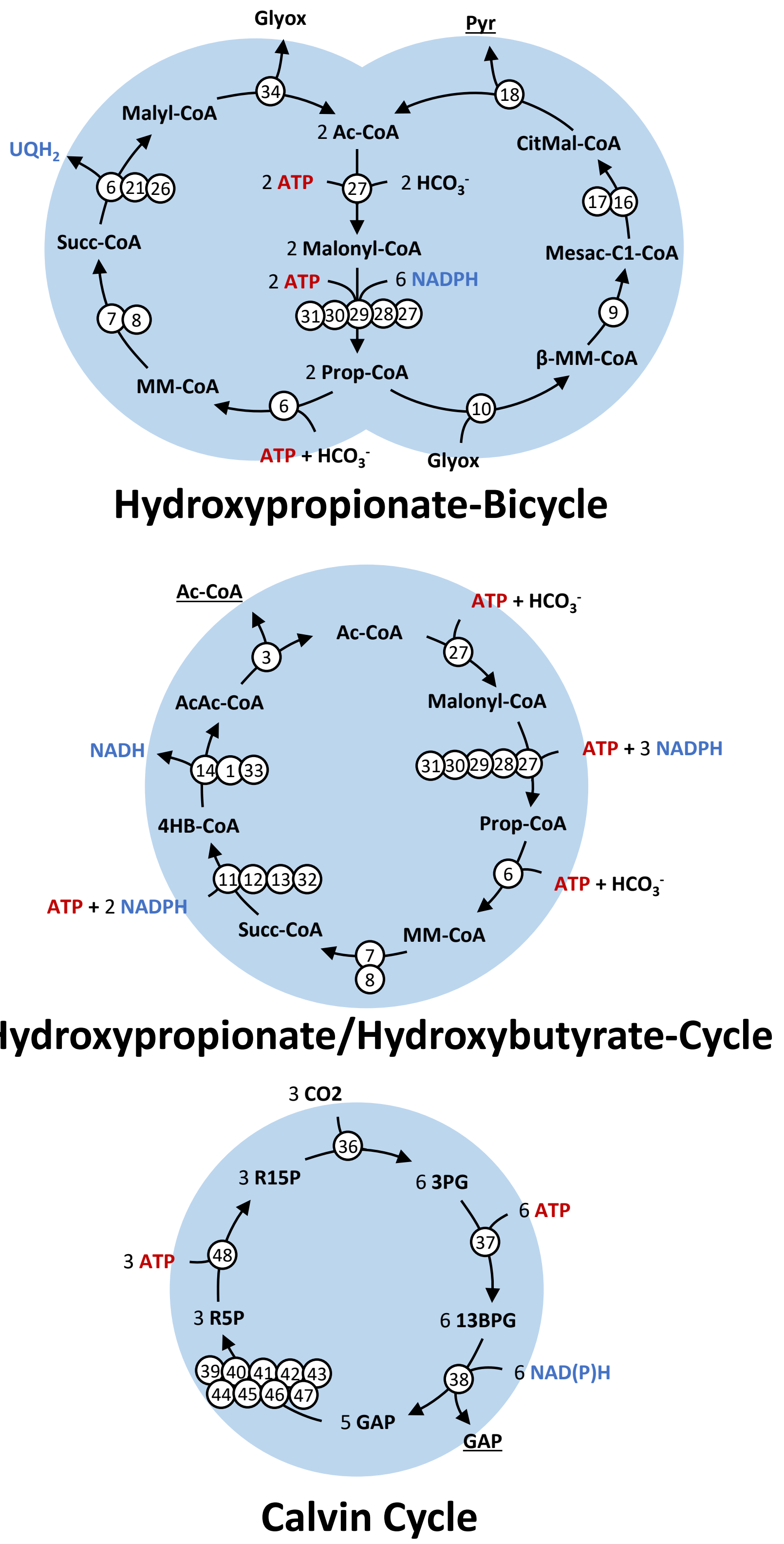


Figure S2: Flux maps of natural CO2-fixing pathways. For simplification, reaction arrows can include multiple reactions and skip some metabolites. ADP, AMP, phosphate, water, and oxidized forms of electron carriers were also left out to improve clarity. Reactions are numbered and each number specifies the respective enzyme according to Table S1. Abbreviations: Ac-CoA, acetyl-CoA; Prop-CoA, propanoyl-CoA; β-MM-CoA, β-methylmalyl-CoA; CitMal-CoA, citramalyl-CoA; Pyr, pyruvate; MM-CoA, methylmalonyl-CoA; Succ-CoA, succinyl-CoA; Glyox, glyoxylate; 4HB-CoA, 4-hydroxybutyrate; AcAc-CoA, acetoacetyl-CoA; R15P, ribulose-1,5-bisphosphate; 3PG, 3-phosphoglycerate; 13BPG, 1,3-bisphophoglycerate; GAP, glyceraldehyde-3-phosphate; R5P, ribulose-5-phosphate.

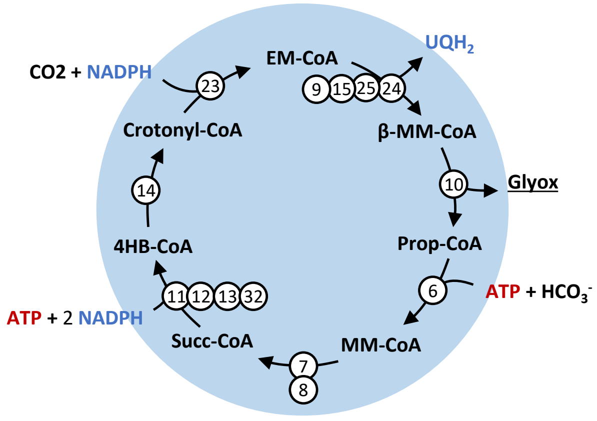


**CETCH-Cycle**

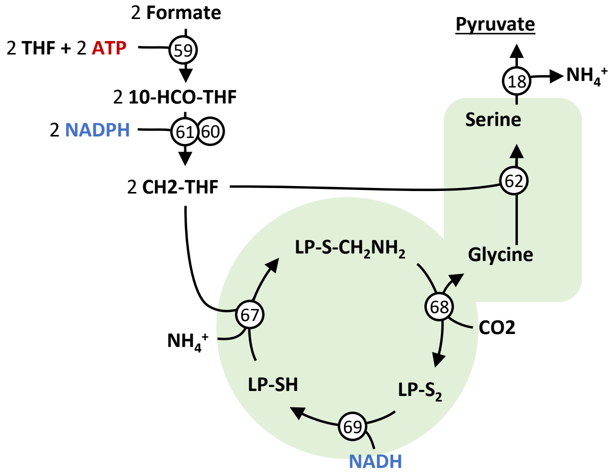


**Reductive Glycine Pathway**

Figure S3: Flux maps of artificial CO2- or C1-fixing pathways established in earlier works. For simplification, reaction arrows can include multiple reactions and skip some metabolites. ADP, AMP, phosphate, water, and oxidized forms of electron carriers were also left out to improve clarity. Reactions are numbered and each number specifies the respective enzyme according to Table S1. Abbreviations: Ac-CoA, acetyl-CoA; Prop-CoA, propanoyl-CoA; β-MM-CoA, β-methylmalyl-CoA; Pyr, pyruvate; MM-CoA, methylmalonyl-CoA; Succ-CoA, succinyl-CoA; Glyox, glyoxylate; 4HB-CoA, 4-hydroxybutyrate; OH-Pyr, hydroxypyruvate; 10-CHO-THF, 10-formyltetrahydrofolate; CH2-THF, 5,10-methylenetetrahydrofolate; LP-$S_2$, [glycine-cleavage complex H protein]-N6-lipoyl-L-lysine; LP-S-$CH_2NH_2$, [glycine-cleavage complex H protein]-S-aminomethyl-N6-dihydrolipoyl-L-lysine; LP-SH, [glycine-cleavage complex H protein]- dihydrolipoyl-L-lysine.

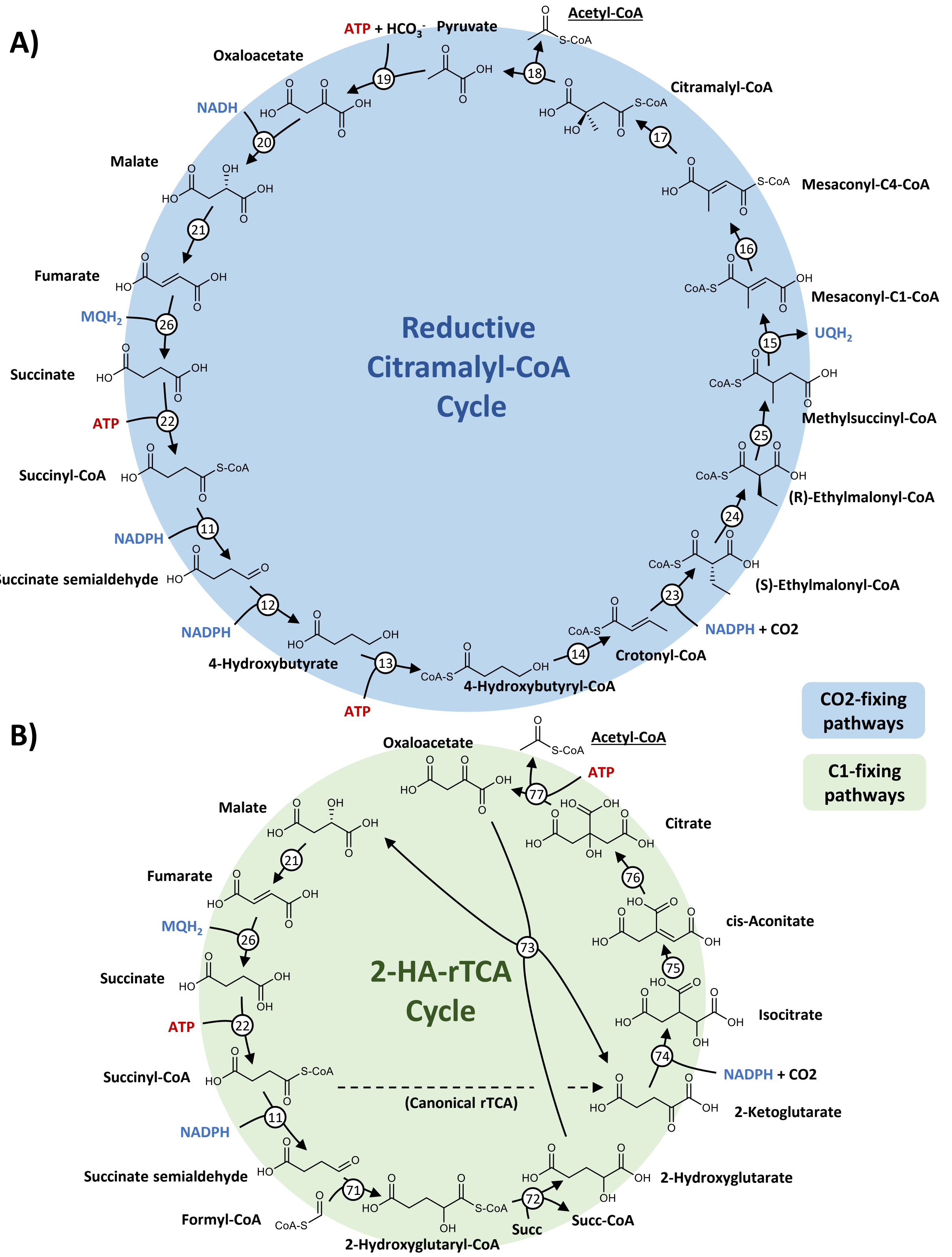


Figure S4: Flux maps of the rCCC and 2-HA-rTCA Cycle. ADP, AMP, phosphate, water, and oxidized forms of electron carriers were also left out to improve clarity. Reactions are numbered and each number specifies the respective enzyme according to Table S1. Abbreviations: Succ, succinate; Succ-CoA, succinyl-CoA.

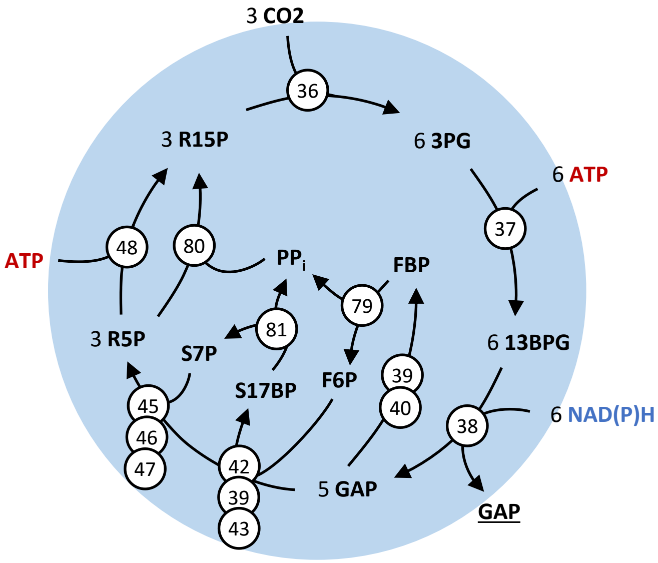


**Phosphatase-less Calvin Cycle**

Figure S5: Flux maps of the hypothetical phosphatase-less Calvin-Benson-Bassham Cycle. For simplification, reaction arrows can include multiple reactions and skip some metabolites. ADP, AMP, phosphate, water, and oxidized forms of electron carriers were also left out to improve clarity. Reactions are numbered and each number specifies the respective enzyme according to Table S1. Abbreviations: R15P, ribulose-1,5-bisphosphate; 3PG, 3-phosphoglycerate; 13BPG, 1,3-bisphosphoglycerate; GAP, glyceraldehyde-3-phosphate; FBP, fructose-1,6-bisphosphate; F6P, fructose-6-phosphate; S17BP, seduheptulose-1,7-bisphosphate; PPi, pyrophyosphate; S7P, seduheptulose-7-phosphate; R5P, ribulose-5-phosphate.

#### 1.2. Pathway modules connecting central metabolites

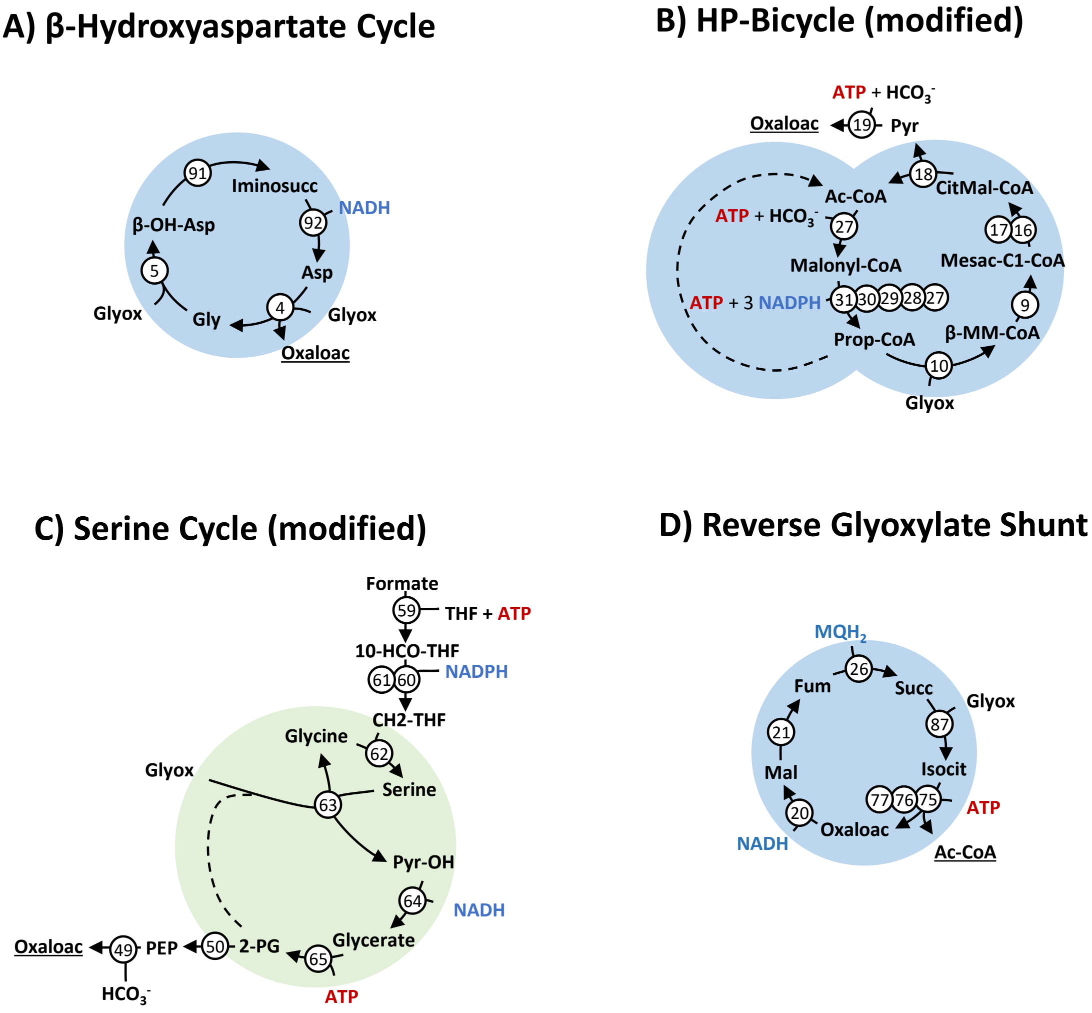


Figure S6: Flux maps of pathway subnetworks that are modules for conversion of **glyoxylate to acetyl-CoA or oxaloacetate**. For simplification, reaction arrows can include multiple reactions and skip some metabolites. ADP, AMP, phosphate, water, and oxidized forms of electron carriers were also left out to improve clarity. Reactions are numbered and each number specifies the respective enzyme according to Table S1. Abbreviations: Iminosucc, iminosuccinate; β-OH-Asp, β-hydroxyaspartate; Asp, aspartate; Glyox, glyoxylate; Oxaloac, oxaloacetate; Gly, glycine; Pyr, pyruvate; CitMal-CoA, citramalyl-CoA; Ac-CoA, acetyl-CoA; Prop-CoA, propanoyl-CoA; β-MM-CoA, β-methylmalyl-CoA; Mesac-C1-CoA, mesaconyl-C1-CoA; Fum, fumarate; Succ, succinate; Isocit, isocitrate; Mal, malate; PEP, phosphoenolpyruvate; OH-Pyr, hydroxypyruvate; 2-PG, 2-phosphoglycerate; 10-CHO-THF, 10-formyltetrahydrofolate; CH2-THF, 5,10-methylenetetrahydrofolate; LP-$S_2$, [glycine-cleavage complex H protein]-N6-lipoyl-L-lysine; LP-S-$CH_2NH_2$, [glycine-cleavage complex H protein]-S-aminomethyl-N6-dihydrolipoyl-L-lysine; LP-SH, [glycine-cleavage complex H protein]- dihydrolipoyl-L-lysine.

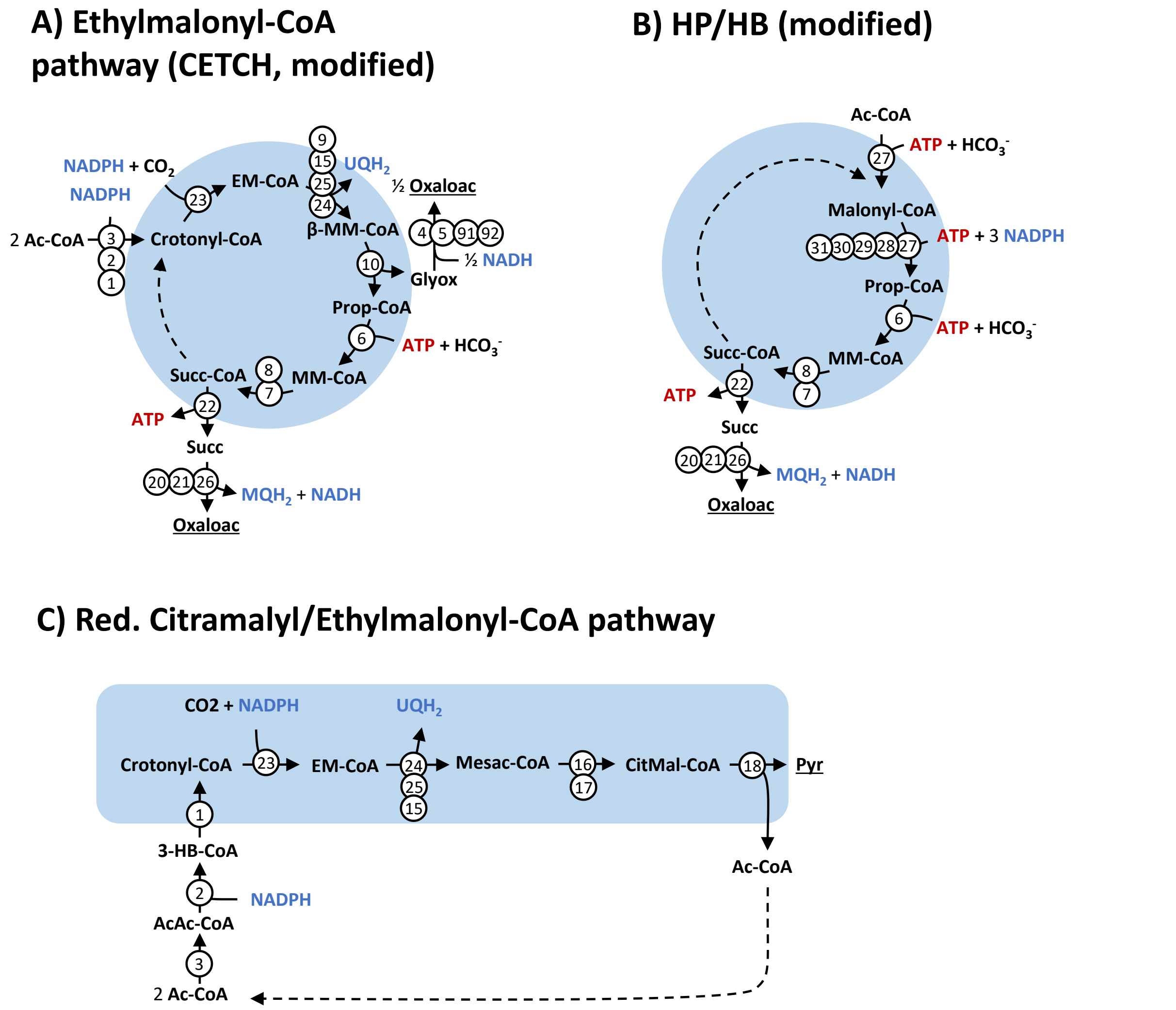


Figure S7: Flux maps of pathway subnetworks that are modules for conversion of **acetyl-CoA to pyruvate or oxaloacetate**. For simplification, reaction arrows can include multiple reactions and skip some metabolites. ADP, AMP, phosphate, water, and oxidized forms of electron carriers were also left out to improve clarity. Reactions are numbered and each number specifies the respective enzyme according to Table S1. Abbreviations: Glyox, glyoxylate; Oxaloac, oxaloacetate; Pyr, pyruvate; EM-CoA, ethylmalonyl-CoA; CitMal-CoA, citramalyl-CoA; Ac-CoA, acetyl-CoA; Prop-CoA, propanoyl-CoA; β-MM-CoA, β-methylmalyl-CoA; MM-CoA, methylmalonyl-CoA; Succ-CoA, succinyl-CoA; Mesac-CoA, mesaconyl-CoA; Succ, succinate; AcAc-CoA, acetoacetyl-CoA; 3-HB-CoA, 3-hydroxybutanoyl-CoA; 10-CHO-THF, 10-formyltetrahydrofolate; CH2-THF, 5,10-methylenetetrahydrofolate; LP-$S_2$, [glycine-cleavage complex H protein]-N6-lipoyl-L-lysine; LP-S-$CH_2NH_2$, [glycine-cleavage complex H protein]-S-aminomethyl-N6-dihydrolipoyl-L-lysine; LP-SH, [glycine-cleavage complex H protein]-dihydrolipoyl-L-lysine.

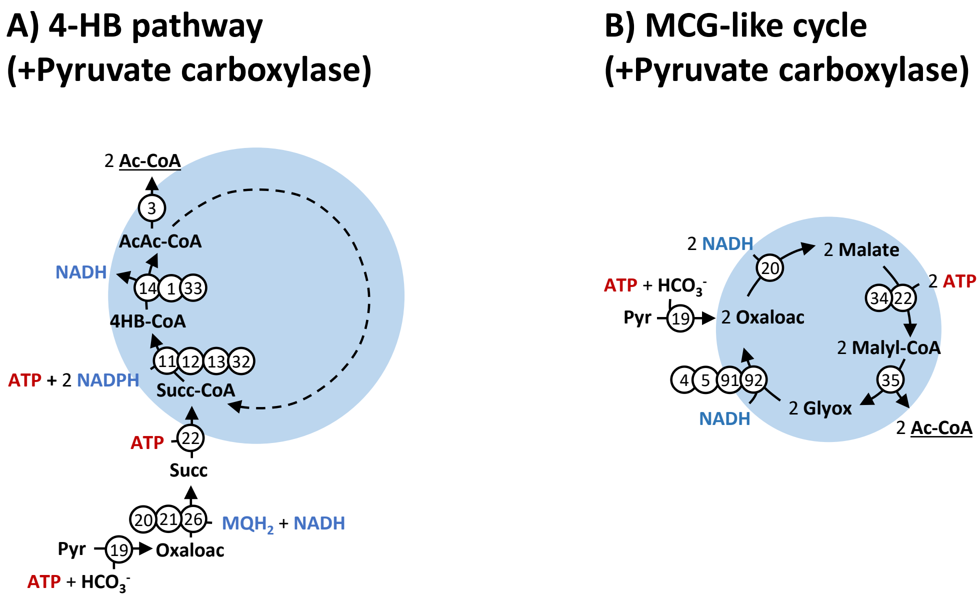


Figure S8: Flux maps of pathway subnetworks that are modules for conversion of **one mol pyruvate or oxaloacetate to 2 mol acetyl-CoA**. For simplification, reaction arrows can include multiple reactions and skip some metabolites. ADP, AMP, phosphate, water, and oxidized forms of electron carriers were also left out to improve clarity. Reactions are numbered and each number specifies the respective enzyme according to Table S1. Abbreviations: Glyox, glyoxylate; Oxaloac, oxaloacetate; Pyr, pyruvate; Ac-CoA, acetyl-CoA; Prop-CoA, propanoyl-CoA; Succ-CoA, succinyl-CoA; Malate, malate; Mesac-CoA, mesaconyl-CoA; Succ, succinate; AcAc-CoA, acetoacetyl-CoA; 4HB-CoA, 4-hydroxybutanoyl-CoA.

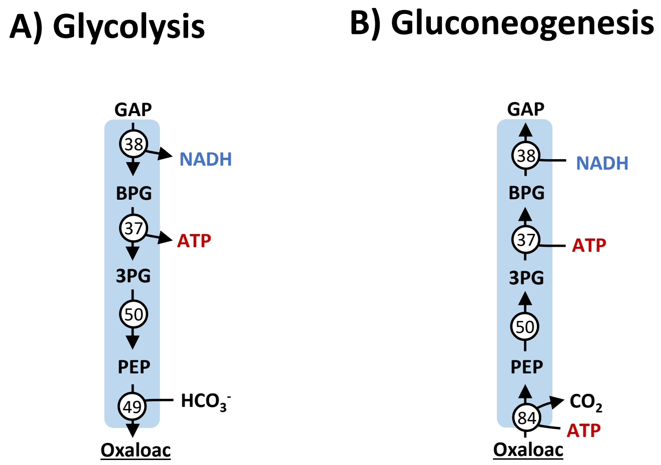


Figure S9: Flux maps of pathway subnetworks that are modules for conversion **of glyceraldehyde-3-phosphate to oxaloacetate (part of the glycolysis) and vice versa (gluconeogenesis)**. For simplification, reaction arrows can include multiple reactions and skip some metabolites. ADP, AMP, phosphate, water, and oxidized forms of electron carriers were also left out to improve clarity. Reactions are numbered and each number specifies the respective enzyme according to Table S1. Abbreviations: GAP, glyceraldehyde-3-phosphate; BPG, 1,3-bisphosphoglycerate; 3PG, 3-phosphoglycerate; PEP, phosphoenolpyruvate; Oxaloac, oxaloacetate.

### 2. Data and parameters used in this study with references

#### 2.1. Kinetic parameters of enzymes

Table S1: Kinetic parameters used in this study with their uncertainty and sources

| Enzyme name/ Parameter | Value | Uncer-tainty | Unit | Comment | Source | # |
|---|---|---|---|---|---|---|
| **(3S)-3-hydroxyacyl-CoA hydro-lyase** | | | | | | **1** |
| Stoichiometry: (S)-3-hydroxybutanoyl-CoA(aq) ⇌ crotonoyl-CoA(aq) + H2O(l) | | | | | | |
| $K_{eq}$ | 0.2155 | | | | eQuilibrator | |
| M | 15355 | | g mol$^{-1}$ | | UniProt | |
| Backward reaction: | | | | | | |
| $K_m$ (crotonyl-CoA) | 9.26 | | µM | | (Moskowitz & Merrick, 1969) | |
| $K_{cat}^-$ | 49.14 | | s$^{-1}$ | | (Moskowitz & Merrick, 1969) | |
| **(S)-3-hydroxyacyl-CoA:NADP+ oxidoreductase** | | | | | | **2** |
| Stoichiometry: (S)-3-hydroxybutanoyl-CoA(aq) + NADP+(aq) ⇌ acetoacetyl-CoA(aq) + NADPH(aq) | | | | | | |
| $K_{eq}$ | $2.814 \cdot 10^{-3}$ | | | | eQuilibrator | |
| M | 141000 | | g mol$^{-1}$ | | (Belova et al., 1997) | |
| Backward reaction: | | | | | | |
| $K_m$ (acetoacetyl-CoA) | 11.6 | | µM | | (Belova et al., 1997) | |
| $K_m$ (NADPH) | 41 | | µM | | (Belova et al., 1997) | |
| $K_{cat}^-$ | 432.4 | | s$^{-1}$ | | (Belova et al., 1997) | |
| **acetyl-CoA:acetyl-CoA C-acetyltransferase** | | | | | | **3** |
| Stoichiometry: 2 acetyl-CoA(aq) ⇌ CoA(aq) + acetoacetyl-CoA(aq) | | | | | | |
| $K_{eq}$ | $2.77 \cdot 10^{-5}$ | | | | eQuilibrator | |
| M | 86497 | | g mol$^{-1}$ | | UniProt | |
| Forward reaction: | | | | | | |
| $K_m$ (acetyl-CoA) | 600 | | µM | | (Hedl et al., 2002) | |
| $K_{cat}^+$ | 122.5 | | s$^{-1}$ | | (Hedl et al., 2002) | |
| Backward reaction: | | | | | | |
| $K_m$ (acetoacetyl-CoA) | 88 | | µM | | (Hedl et al., 2002) | |
| $K_m$ (CoA) | 10 | | µM | | (Hedl et al., 2002) | |
| $K_{cat}^-$ | 1802 | | s$^{-1}$ | | (Hedl et al., 2002) | |
| **L-aspartate:glyoxylate aminotransferase** | | | | | | **4** |
| Stoichiometry: aspartate(aq) + glyoxylate(aq) ⇌ oxaloacetate(aq) + glycine(aq) | | | | | | |
| $K_{eq}$ | 9.546 | | | | eQuilibrator | |
| M | 42507 | | g mol$^{-1}$ | | (Schada von Borzyskowski et al., 2019) | |
| Forward reaction: | | | | | | |
| $K_m$ (aspartate) | 2500 | 100 | µM | | (Schada von Borzyskowski et al., 2019) | |
| $K_m$ (glyoxylate) | 430 | 20 | µM | | (Schada von Borzyskowski et al., 2019) | |
| $K_{cat}^+$ | 57 | 1 | s$^{-1}$ | | (Schada von Borzyskowski et al., 2019) | |
| Backward reaction: | | | | | | |
| $K_m$ (oxaloacetate) | 2900 | 270 | µM | | (Schada von Borzyskowski et al., 2019) | |
| $K_m$ (glycine) | 9500 | 400 | µM | | (Schada von Borzyskowski et al., 2019) | |
| $K_{cat}^-$ | 0.76 | 0.02 | s$^{-1}$ | | (Schada von Borzyskowski et al., 2019) | |
| **(2R,3S)-beta-Hydroxyasparate glyoxylate-lyase (glycine-forming)** | | | | | | **5** |
| Stoichiometry: 3-hydroxyaspartate(aq) ⇌ glycine(aq) + glyoxylate(aq) | | | | | | |
| $K_{eq}$ | 0.536 | | | | eQuilibrator | |
| M | 41786 | | g mol$^{-1}$ | | (Schada von Borzyskowski et al., 2019) | |
| Forward reaction: | | | | | | |
| $K_m$ (3-hydroxyaspartate) | 280 | 30 | µM | | (Schada von Borzyskowski et al., 2019) | |
| $K_{cat}^+$ | 33 | 1 | s$^{-1}$ | | (Schada von Borzyskowski et al., 2019) | |
| Backward reaction: | | | | | | |
| $K_m$ (glyoxylate) | 230 | 30 | µM | | (Schada von Borzyskowski et al., 2019) | |
| $K_m$ (glycine) | 4310 | 340 | µM | | (Schada von Borzyskowski et al., 2019) | |
| $K_{cat}^-$ | 89 | 4 | s$^{-1}$ | | (Schada von Borzyskowski et al., 2019) | |
| **propanoyl-CoA:carbon-dioxide ligase (ADP-forming)** | | | | | | **6** |
| Stoichiometry: ATP(aq) + propanoyl-CoA(aq) + HCO3-(aq) ⇌ ADP(aq) + phosphate(aq) + (S)-methylmalonyl-CoA(aq) | | | | | | |
| $K_{eq}$ | 16.20 | | | | eQuilibrator | |
| M | 510000 | | g mol$^{-1}$ | | (Kimura et al., 1998) | |
| Forward reaction: | | | | | | |
| $K_m$ (ATP) | 36 | | µM | | (Kimura et al., 1998) | |
| $K_m$ (propanoyl-CoA) | 32 | | µM | | (Kimura et al., 1998) | |
| $K_m$ ($HCO_3^-$) | 1130 | | µM | | (Kimura et al., 1998) | |
| $K_{cat}^+$ | 120.1 | | s$^{-1}$ | | (Kimura et al., 1998) | |

Table S1 (continued)

| Enzyme name/ Parameter | Value | Uncer-tainty | Unit | Comment | Source | # |
|---|---|---|---|---|---|---|
| **methylmalonyl-CoA epimerase** | | | | | | **7** |
| Stoichiometry: (R)-methylmalonyl-CoA(aq) ⇌ (S)-methylmalonyl-CoA(aq) | | | | | | |
| $K_{eq}$ | 1 | | | | eQuilibrator | |
| M | 16081 | | g mol$^{-1}$ | | (Dayem et al., 2002) | |
| Forward reaction: | | | | | | |
| $K_m$ ((R)-methylmalonyl-CoA) | 38 | | µM | | (Dayem et al., 2002) | |
| $K_{cat}^{+}$ | 150 | | s$^{-1}$ | | (Dayem et al., 2002) | |
| **(R)-methylmalonyl-CoA CoA-carbonylmutase** | | | | | | **8** |
| Stoichiometry: (R)-methylmalonyl-CoA(aq) ⇌ succinyl-CoA(aq) | | | | | | |
| $K_{eq}$ | 22.23 | | | | eQuilibrator | |
| M | 141774 | | g mol$^{-1}$ | | UniProt | |
| Forward reaction: | | | | | | |
| $K_m$ ((R)-methylmalonyl-CoA) | 86 | 13 | µM | | (Padovani & Banerjee, 2006) | |
| $K_{cat}^{+}$ | 255 | 17 | s$^{-1}$ | | (Padovani & Banerjee, 2006) | |
| **(2R,3S)-2-methylmalyl-CoA hydro-lyase (2-methylfumaryl-CoA-forming)** | | | | | | **9** |
| Stoichiometry: L-erythro-3-methylmalyl-CoA(aq) ⇌ mesaconyl-C1-CoA(aq) + H2O(l) | | | | | | |
| $K_{eq}$ | 3.668 | | | | eQuilibrator | |
| M | 43000 | | g mol$^{-1}$ | | (Borjian et al., 2017) | |
| Forward reaction: | | | | | | |
| $K_m$ (L-erythro-3-methylmalyl-CoA) | 350 | 50 | µM | | (Borjian et al., 2017) | |
| $K_{cat}^{+}$ | 191 | 9 | s$^{-1}$ | | (Borjian et al., 2017) | |
| Backward reaction: | | | | | | |
| $K_m$ (mesaconyl-C1-CoA) | 1000 | 10 | µM | | (Borjian et al., 2017) | |
| $K_{cat}^{-}$ | 14.8 | 0.5 | s$^{-1}$ | | (Borjian et al., 2017) | |
| **L-erythro-3-methylmalyl-CoA glyoxylate-lyase (propanoyl-CoA-forming)** | | | | | | **10** |
| Stoichiometry: L-erythro-3-methylmalyl-CoA(aq) ⇌ propanoyl-CoA(aq) + glyoxylate(aq) | | | | | | |
| $K_{eq}$ | $1.622 \cdot 10^{-3}$ | | | | eQuilibrator | |
| M | 36800 | | g mol$^{-1}$ | | (Erb et al., 2010) | |
| Forward reaction: | | | | | | |
| $K_m$ (L-erythro-3-methylmalyl-CoA) | 10 | | µM | | (Erb et al., 2010) | |
| $K_{cat}^{+}$ | 2.76 | | s$^{-1}$ | | (Erb et al., 2010) | |
| Backward reaction: | | | | | | |
| $K_m$ (propanoyl-CoA) | 0.2 | | µM | | (Erb et al., 2010) | |
| $K_m$ (glyoxylate) | 4.1 | | µM | | (Erb et al., 2010) | |
| $K_{cat}^{-}$ | 12.27 | | s$^{-1}$ | | (Erb et al., 2010) | |
| **succinate semialdehyde:NADP+ oxidoreductase (CoA-acylating)** | | | | | | **11** |
| Stoichiometry: succinate semialdehyde(aq) + CoA(aq) + NADP+(aq) ⇌ succinyl-CoA(aq) + NADPH(aq) | | | | | | |
| $K_{eq}$ | 215.0 | | | | eQuilibrator | |
| M | | | g mol$^{-1}$ | | (Yoshida et al., 2016) | |
| Backward reaction: | | | | | | |
| $K_m$ (succinyl-CoA) | 550 | 130 | µM | | (Yoshida et al., 2016) | |
| $K_m$ (NADPH) | 1420 | 190 | µM | | (Yoshida et al., 2016) | |
| $K_{cat}^{-}$ | 109 | 12 | s$^{-1}$ | | (Yoshida et al., 2016) | |
| **4-hydroxybutanoate:NADP+ oxidoreductase** | | | | | | **12** |
| Stoichiometry: 4-hydroxybutanoic acid(aq) + NADP+(aq) ⇌ succinate semialdehyde(aq) + NADPH(aq) | | | | | | |
| $K_{eq}$ | 3.726 | | | | eQuilibrator | |
| M | 134000 | | g mol$^{-1}$ | | (Meyer et al., 2015) | |
| Backward reaction: | | | | | | |
| $K_m$ (succinate semialdehyde) | 5100 | 600 | µM | | (Meyer et al., 2015) | |
| $K_m$ (NADPH) | 1420 | 190 | µM | | (Meyer et al., 2015) | |
| $K_{cat}^{-}$ | 666 | 89 | s$^{-1}$ | | (Meyer et al., 2015) | |
| **4-hydroxybutyrate---CoA ligase (ADP-forming)** | | | | | | **13** |
| Stoichiometry: ATP(aq) + 4-hydroxybutanoate(aq) + CoA(aq) ⇌ ADP(aq) + Phosphate(aq) + 4-hydroxybutanoyl-CoA(aq) | | | | | | |
| $K_{eq}$ | 971.9 | | | | eQuilibrator | |
| M | 75600 | | g mol$^{-1}$ | | (Könneke et al., 2014) | |
| Forward reaction: | | | | | | |
| $K_m$ (4-hydroxybutyrate) | 370 | 60 | µM | | (Könneke et al., 2014) | |
| $K_m$ (ATP) | 220 | 70 | µM | | (Könneke et al., 2014) | |
| $K_m$ (CoA) | 160 | 50 | µM | | (Könneke et al., 2014) | |
| $K_{cat}^{+}$ | 1.76 | 0.13 | s$^{-1}$ | | (Könneke et al., 2014) | |

Table S1 (continued)

| Enzyme name/ Parameter | Value | Uncer-tainty | Unit | Comment | Source | # |
|---|---|---|---|---|---|---|
| **4-hydroxybutanoyl-CoA hydro-lyase** | | | | | | **14** |
| Stoichiometry: 4-hydroxybutanoyl-CoA(aq) ⇌ crotonyl-CoA(aq) + H2O(l) | | | | | | |
| $K_{eq}$ | 22.72 | | | | eQuilibrator | |
| M | 56765 | | g $mol^{-1}$ | | Könneke et al., 2014), UniProt | |
| Forward reaction: | | | | | | |
| $K_m$ (4-hydroxybutyryl-CoA) | 60 | 20 | µM | | (Könneke et al., 2014) | |
| $K_{cat}^{+}$ | 23.7 | 1.9 | $s^{-1}$ | | (Könneke et al., 2014) | |
| **(2S)-methylsuccinyl-CoA:electron-transfer flavoprotein oxidoreductase** | | | | | | **15** |
| Stoichiometry: methylsuccinyl-CoA(aq) + ubiquinone(aq) ⇌ mesaconyl-C1-CoA(aq) + ubinquinol(aq) | | | | | | |
| $K_{eq}$ | 28000 | | | | eQuilibrator | |
| M | 60000 | | g $mol^{-1}$ | | (Schwander et al., 2018) | |
| Forward reaction: | | | | | | |
| $K_m$ (methylsuccinyl-CoA) | 80 | 5 | µM | Electron acceptor unknown | (Schwander et al., 2018) | |
| $K_{cat}^{+}$ | 82.3 | 1.4 | $s^{-1}$ | | (Schwander et al., 2018) | |
| **2-methylfumaryl-CoA 1,4-CoA-mutase** | | | | | | **16** |
| Stoichiometry: mesaconyl-C1-CoA(aq) ⇌ mesaconyl-C4-CoA(aq) | | | | | | |
| $K_{eq}$ | 1 | | | | eQuilibrator | |
| M | 47000 | | g $mol^{-1}$ | | (Zarzycki et al., 2009) | |
| Forward reaction: | | | | | | |
| $K_m$ (mesaconyl-C1-CoA) | 240 | | µM | T = 55°C | (Zarzycki et al., 2009) | |
| $K_{cat}^{+}$ | 407.3 | | $s^{-1}$ | T = 55°C | (Zarzycki et al., 2009) | |
| **(S)-citramalyl-CoA hydro-lyase (3-methylfumaryl-CoA-forming)** | | | | | | **17** |
| Stoichiometry: (3S)-citramalyl-CoA(aq) ⇌ mesaconyl-C4-CoA(aq) + H2O(l) | | | | | | |
| $K_{eq}$ | 0.2782 | | | | eQuilibrator | |
| M | 33000 | | g $mol^{-1}$ | | (Zarzycki et al., 2009) | |
| Backward reaction: | | | | | | |
| $K_m$ (mesaconyl-C4-CoA) | 75 | | µM | T = 55°C | (Zarzycki et al., 2009) | |
| $K_{cat}^{-}$ | 522.5 | | $s^{-1}$ | T = 55°C | (Zarzycki et al., 2009) | |
| **(3S)-citramalyl-CoA pyruvate-lyase (acetyl-CoA-forming)** | | | | | | **18** |
| Stoichiometry: (3S)-citramalyl-CoA(aq) ⇌ acetyl-CoA(aq) + pyruvate(aq) | | | | | | |
| $K_{eq}$ | $1.293 \cdot 10^{-2}$ | | | | eQuilibrator | |
| M | 29022 | | g $mol^{-1}$ | | (Sasikaran et al., 2014) | |
| Forward reaction: | | | | | | |
| $K_m$ (citramalyl-CoA) | 30 | 9 | µM | | (Sasikaran et al., 2014) | |
| $K_{cat}^{+}$ | 82 | 10 | $s^{-1}$ | | (Sasikaran et al., 2014) | |
| **pyruvate:carbon-dioxide ligase (ADP-forming)** | | | | | | **19** |
| Stoichiometry: ATP(aq) + pyruvate(aq) + HCO3-(aq) ⇌ ADP(aq) + phosphate(aq) + oxaloacetate(aq) | | | | | | |
| $K_{eq}$ | 24.68 | | | | eQuilibrator | |
| M | 130000 | | g $mol^{-1}$ | | (Gurr & Jones, 1977) | |
| Forward reaction: | | | | | | |
| $K_m$ (pyruvate) | 1860 | | µM | | (Gurr & Jones, 1977) | |
| $K_m$ (ATP) | 80 | | µM | | (Gurr & Jones, 1977) | |
| $K_m$ ($HCO_3^-$) | 330 | | µM | | (Gurr & Jones, 1977) | |
| $K_{cat}^{+}$ | 58.5 | | $s^{-1}$ | | (Gurr & Jones, 1977) | |
| **(S)-malate:NAD+ oxidoreductase** | | | | | | **20** |
| Stoichiometry: (S)-malate(aq) + NAD+(aq) ⇌ oxaloacetate(aq) + NADH(aq) | | | | | | |
| $K_{eq}$ | $2.11 \cdot 10^{-5}$ | | | | eQuilibrator | |
| M | 40900 | | g $mol^{-1}$ | | (Muslin et al., 1995) | |
| Forward reaction: | | | | | | |
| $K_m$ (malate) | 2600 | 200 | µM | | (Muslin et al., 1995) | |
| $K_m$ ($NAD^+$) | 260 | 30 | µM | | (Muslin et al., 1995) | |
| $K_{cat}^{+}$ | 21 | | $s^{-1}$ | | (Muslin et al., 1995) | |
| Backward reaction: | | | | | | |
| $K_m$ (oxaloacetate) | 49 | 3 | µM | | (Muslin et al., 1995) | |
| $K_m$ (NADH) | 61 | 2 | µM | | (Muslin et al., 1995) | |
| $K_{cat}^{-}$ | 900 | | $s^{-1}$ | | (Muslin et al., 1995) | |

Table S1 (continued)

| Enzyme name/ Parameter | Value | Uncer-tainty | Unit | Comment | Source | # |
|---|---|---|---|---|---|---|
| **(S)-malate hydro-lyase (fumarate-forming)** | | | | | | **21** |
| Stoichiometry: (S)-malate(aq) ⇌ fumarate(aq) + H2O(l) | | | | | | |
| $K_{eq}$ | 0.2492 | | | | eQuilibrator | |
| M | 60000 | | g mol$^{-1}$ | | (Ueda et al., 1991) | |
| Forward reaction: | | | | | | |
| $K_m$ (malate) | 700 | 100 | µM | | (Ueda et al., 1991) | |
| $K_{cat}^+$ | 720 | 40 | s$^{-1}$ | | (Ueda et al., 1991) | |
| Backward reaction: | | | | | | |
| $K_m$ (fumarate) | 460 | 80 | µM | | (Ueda et al., 1991) | |
| $K_{cat}^-$ | 1900 | 100 | s$^{-1}$ | | (Ueda et al., 1991) | |
| **succinate:CoA ligase (ADP-forming)** | | | | | | **22** |
| Stoichiometry: ATP(aq) + succinate(aq) + CoA(aq) ⇌ ADP(aq) + phosphate(aq) + succinyl-CoA(aq) | | | | | | |
| $K_{eq}$ | 1.745 | | | | eQuilibrator | |
| M | 71170 | | g mol$^{-1}$ | | (Nolte et al., 2014) | |
| Forward reaction: | | | | | | |
| $K_m$ (succinate) | 141 | 3 | µM | | (Nolte et al., 2014) | |
| $K_m$ (ATP) | 55 | 2 | µM | | (Nolte et al., 2014) | |
| $K_m$ (CoA) | 58 | 5 | µM | | (Nolte et al., 2014) | |
| $K_{cat}^+$ | 26.7 | 1.1 | s$^{-1}$ | | (Nolte et al., 2014) | |
| **(2S)-ethylmalonyl-CoA:NADP+ oxidoreductase (decarboxylating)** | | | | | | **23** |
| Stoichiometry: (2S)-ethylmalonyl-CoA(aq) + NADP+(aq) ⇌ crotonyl-CoA(aq) + CO2(aq) + NADPH(aq) | | | | | | |
| $K_{eq}$ | $2.5 \cdot 10^{-8}$ | | | | eQuilibrator | |
| M | 48476 | | g mol$^{-1}$ | | UniProt | |
| Backward reaction: | | | | | | |
| $K_m$ (crotonyl-CoA) | 21 | 2 | µM | | (Stoffel et al., 2019) | |
| $K_m$ (NADPH) | 37 | 4 | µM | | (Stoffel et al., 2019) | |
| $K_m$ (CO2) | 90 | 1 | µM | | (Stoffel et al., 2019) | |
| $K_{cat}^-$ | 103 | 3 | s$^{-1}$ | | (Stoffel et al., 2019) | |
| **(2S)-ethylmalonyl-CoA epimerase** | | | | | | **24** |
| Stoichiometry: (2S)-ethylmalonyl-CoA(aq) ⇌ (2R)-ethylmalonyl-CoA(aq) | | | | | | |
| $K_{eq}$ | 1 | | | | eQuilibrator | |
| M | 14260 | | g mol$^{-1}$ | | UniProt | |
| Forward reaction: | | | | | | |
| $K_m$ (S-ethylmalonyl-CoA) | 40 | | µM | | (Erb et al., 2008) | |
| $K_{cat}^+$ | 26.14 | | s$^{-1}$ | | (Erb et al., 2008) | |
| **(2R)-ethylmalonyl-CoA CoA-carbonylmutase** | | | | | | **25** |
| Stoichiometry: (2R)-Ethylmalonyl-CoA(aq) ⇌ Methylsuccinyl-CoA(aq) | | | | | | |
| $K_{eq}$ | 1 | | | | eQuilibrator | |
| M | 74000 | | g mol$^{-1}$ | | (Erb et al., 2008) | |
| Forward reaction: | | | | | | |
| $K_m$ (R-ethylmalonyl-CoA) | 60 | | µM | | (Erb et al., 2008) | |
| $K_{cat}^+$ | 8.63 | | s$^{-1}$ | | (Erb et al., 2008) | |
| **succinate:menaquinone oxidoreductase** | | | | | | **26** |
| Stoichiometry: succinate(aq) + menaquinone(aq) + 2 H+(out) ⇌ fumarate(aq) + menaquinol(aq) + 2 H+(in) | | | | | | |
| $K_{eq}$ | 286.8 | | | Assuming a membrane potential of 0.15 V & translocation of 2 $H^+$ | eQuilibrator + calculation | |
| M | 116499 | | g mol$^{-1}$ | | UniProt(*B. subtilis* protein) | |
| Forward reaction: | | | | | | |
| $K_m$ (succinate) | 900 | | µM | *B. subtilis* | (Hederstedt & Heden, 1989) | |
| $K_m$ (menaquinone) | 1.5 | | µM | *E. coli* (no data *for B. subtilis*) | (Maklashina & Cecchini, 1999) | |
| $K_{cat}^+$ | 94 | | s$^{-1}$ | *B. subtilis* | (Hederstedt & Heden, 1989) | |
| Backward reaction: | | | | | | |
| $K_m$ (fumarate) | 20 | 2 | µM | *E. coli* (no data *for B. subtilis*) | (Maklashina & Cecchini, 1999) | |
| $K_m$ (menaquinol) | 5.4 | | µM | *E. coli* (no data *for B. subtilis*) | (Maklashina & Cecchini, 1999) | |
| $K_{cat}^-$ | 23.5 | | s$^{-1}$ | *B. subtilis*, extrapolated from cell extract activity and protein mass fraction | (Schnorpfeil et al., 2001) + calculations | |

Table S1 (continued)

| **Enzyme name/ Parameter** | **Value** | **Uncer-tainty** | **Unit** | **Comment** | **Source** | **#** |
|---|---|---|---|---|---|---|
| **acetyl-CoA:hydrogencarbonate ligase (ADP-forming)** | | | | | | **27** |
| Stoichiometry: acetyl-CoA(aq) + $HCO3^-$(aq) + ATP(aq) <=> malonyl-CoA(aq) + ADP(aq) + phosphate(aq) | | | | | | |
| $K_{eq}$ | 33.05 | | | | eQuilibrator | |
| M | 2270000 | | g $mol^{-1}$ | | (Price et al., 2003) | |
| Forward reaction: | | | | | | |
| $K_m$ (acetyl-CoA) | 120 | 10 | µM | | (Price et al., 2003) | |
| $K_m$ (ATP) | 80 | 10 | µM | | (Price et al., 2003) | |
| $K_m$ ($HCO_3^-$) | 1009 | 70 | µM | | (Price et al., 2003) | |
| $K_{cat}^+$ | 8210 | | $s^{-1}$ | | (Price et al., 2003) | |
| **malonyl-CoA:NADP+ oxidoreductase (3-Hydroxypropanoate-forming)** | | | | | | **28** |
| Stoichiometry: 3-hydroxypropanoate(aq) + CoA(aq) + 2 NADP+(aq) ⇌ malonyl-CoA(aq) + 2 NADPH(aq) | | | | | | |
| $K_{eq}$ | 0.02967 | | | | eQuilibrator | |
| M | 300000 | | g $mol^{-1}$ | | (Hügler et al., 2002) | |
| Backward reaction: | | | | | | |
| $K_m$ (malonyl-CoA) | 30 | | µM | | (Hügler et al., 2002) | |
| $K_m$ (NADPH) | 25 | | µM | | (Hügler et al., 2002) | |
| $K_{cat}^-$ | 50 | | $s^{-1}$ | | (Hügler et al., 2002) | |
| **hydroxypropanoate:CoA ligase (AMP-forming)/acrylyl-CoA:NADP+ oxidoreductase (propanoyl-CoA-forming)** | | | | | | **29** |
| Stoichiometry: 3-hydroxypropanoate(aq) + ATP(aq) + CoA(aq) + NADPH(aq) ⇌ propanoyl-CoA(aq) + AMP(aq) + $PP_i$(aq) + NADP+(aq) | | | | | | |
| $K_{eq}$ | $2.04 \cdot 10^{10}$ | | | | eQuilibrator | |
| M | 200000 | | g $mol^{-1}$ | | (Bernhardsgrütter et al., 2018) | |
| Forward reaction: | | | | | | |
| $K_m$ (3-hydroxypropionate) | 200 | 20 | µM | | (Bernhardsgrütter et al., 2018) | |
| $K_m$ (ATP) | 340 | 140 | µM | | (Bernhardsgrütter et al., 2018) | |
| $K_m$ (CoA) | 220 | 50 | µM | | (Bernhardsgrütter et al., 2018) | |
| $K_m$ (NADPH) | 20 | 3 | µM | | (Bernhardsgrütter et al., 2018) | |
| $K_{cat}^+$ | 5.1 | 0.2 | $s^{-1}$ | | (Bernhardsgrütter et al., 2018) | |
| **hydroxypropanoate:CoA ligase (ADP-forming)** | | | | | | **30** |
| Stoichiometry: 3-hydroxypropanoate(aq) + ATP(aq) + CoA(aq) ⇌ 3-hydroxypropanoyl-CoA(aq) + ADP(aq) + phosphate(aq) + NADP(aq) | | | | | | |
| $K_{eq}$ | 0.01104 | | | | eQuilibrator | |
| M | 76000 | | g $mol^{-1}$ | | (Könneke et al., 2014) | |
| Forward reaction: | | | | | | |
| $K_m$ (3-hydroxypropionate) | 1200 | 200 | µM | | (Könneke et al., 2014) | |
| $K_m$ (ATP) | 600 | 100 | µM | | (Könneke et al., 2014) | |
| $K_m$ (CoA) | 160 | 90 | µM | | (Könneke et al., 2014) | |
| $K_{cat}^+$ | 0.747 | | $s^{-1}$ | | (Könneke et al., 2014) | |
| Backward reaction: | | | | | | |
| $K_m$ (phosphate) | 5000 | 5000 | µM | A parameter prior of 5 mM for phosphate was used | This study | |
| **propanoyl-CoA:NADP+ oxidoreductase/ hydroxypropanoyl-CoA dehydratase** | | | | | | **31** |
| Stoichiometry: propanoyl-CoA(aq) + NADP+(aq) + H2O(l) ⇌ 3-hydroxypropanoyl-CoA(aq) + NADPH(aq) | | | | | | |
| $K_{eq}$ | $6.87 \cdot 10^{-11}$ | | | | eQuilibrator | |
| M | 133000 | | g $mol^{-1}$ | | (Bernhardsgrütter et al., 2018), only 2/3 of the enzyme are needed | |
| Backward reaction: | | | | | | |
| $K_m$ (3-hydroxypropanoyl-CoA) | 200 | 20 | µM | It was assumed that the Km would be in the order of 3-hydroxypropanoate of the original enzyme | (Bernhardsgrütter et al., 2018) | |
| $K_m$ (NADPH) | 20 | 3 | µM | | (Bernhardsgrütter et al., 2018) | |
| $K_{cat}^-$ | 11.7 | 1 | $s^{-1}$ | | (Bernhardsgrütter et al., 2018) | |
| **4-hydroxybutyrate---CoA ligase (AMP-forming)** | | | | | | **32** |
| Stoichiometry: ATP(aq) + 4-hydroxybutanoate(aq) + CoA(aq) ⇌ AMP(aq) + diphosphate(aq) + 4-hydroxybutanoyl-CoA(aq) | | | | | | |
| $K_{eq}$ | $1.237 \cdot 10^5$ | | | | eQuilibrator | |
| M | 64500 | | g $mol^{-1}$ | | (Hawkins et al., 2014) | |
| Forward reaction: | | | | | | |
| $K_m$ (4-hydroxybutanoate) | 2000 | 400 | µM | T = 70°C | (Hawkins et al., 2014) | |
| $K_{cat}^+$ | 1.82 | 0.12 | $s^{-1}$ | T = 70°C | (Hawkins et al., 2014) | |

Table S1 (continued)

| Enzyme name/ Parameter | Value | Uncer-tainty | Unit | Comment | Source | # |
|---|---|---|---|---|---|---|
| **(S)-3-hydroxyacyl-CoA:NAD+ oxidoreductase** | | | | | | **33** |
| Stoichiometry: (S)-3-hydroxybutanoyl-CoA(aq) + NAD+(aq) ⇌ acetoacetyl-CoA(aq) + NADH(aq) | | | | | | |
| $K_{eq}$ | $3.043 \cdot 10^{-3}$ | | | | eQuilibrator | |
| M | 79594 | | g $mol^{-1}$ | | UniProt | |
| Forward reaction: | | | | | | |
| $K_m$ ($NAD^+$) | 830 | | µM | | (Shimakata et al., 1979) | |
| Backward reaction: | | | | | | |
| $K_m$ (acetoacetyl-CoA) | 66 | | µM | | (Binstock & Schulz, 1981) | |
| $K_m$ (NADH) | 33 | | µM | | (Shimakata et al., 1979) | |
| $K_{cat}^-$ | 84.9 | | $s^{-1}$ | | (Binstock & Schulz, 1981) | |
| **succinyl-CoA:(S)-malate CoA-transferase** | | | | | | **34** |
| Stoichiometry: succinyl-CoA(aq) + (S)-malate(aq) ⇌ succinate(aq) + (r,s)-malyl-coa(aq) | | | | | | |
| $K_{eq}$ | 6.877 | | | | eQuilibrator | |
| M | 90800 | | g $mol^{-1}$ | | (Friedmann et al., 2006) | |
| Forward reaction: | | | | | | |
| $K_m$ (succinyl-CoA) | 500 | 100 | µM | T = 55°C | (Friedmann et al., 2006) | |
| $K_m$ ((S)-malate) | 1300 | 300 | µM | T = 55°C | (Friedmann et al., 2006) | |
| $K_{cat}^+$ | 11.35 | | $s^{-1}$ | T = 55°C | (Friedmann et al., 2006) | |
| **L-malyl-CoA glyoxylate-lyase (acetyl-CoA-forming)** | | | | | | **35** |
| Stoichiometry: L-malyl-CoA(aq) ⇌ acetyl-CoA(aq) + glyoxylate(aq) | | | | | | |
| $K_{eq}$ | $2.801 \cdot 10^{-3}$ | | | | eQuilibrator | |
| M | 204000 | | g $mol^{-1}$ | | (Meister et al., 2005) | |
| Forward reaction: | | | | | | |
| $K_m$ ((r,s)-malyl-coa(aq)) | 15 | 3 | µM | | (Meister et al., 2005) | |
| $K_{cat}^+$ | 19.38 | 0.68 | $s^{-1}$ | | (Meister et al., 2005) | |
| Backward reaction: | | | | | | |
| $K_m$ (acetyl-CoA) | 140 | 10 | µM | | (Meister et al., 2005) | |
| $K_{cat}^-$ | 126 | 3 | $s^{-1}$ | | (Meister et al., 2005) | |
| **3-phospho-D-glycerate carboxy-lyase (dimerizing; D-ribulose-1,5-bisphosphate-forming)** | | | | | | **36** |
| Stoichiometry: 2 3-phospho-D-glycerate(aq) ⇌ D-ribulose-1,5-bisphosphate(aq) + CO2(aq) + H2O(l) | | | | | | |
| $K_{eq}$ | $8.14 \cdot 10^{-7}$ | | | | eQuilibrator | |
| M | 72268 | | g $mol^{-1}$ | | UniProt | |
| Backward reaction: | | | | | | |
| $K_m$ (ribulose-1,5-bisphosphate) | 30 | 4 | µM | | (Carmo-Silva et al., 2010) | |
| $K_m$ (CO2) | 10.9 | 0.9 | µM | | (Nakano et al., 2010) | |
| $K_{cat}^-$ | 3.06 | 0.08 | $s^{-1}$ | | (Carmo-Silva et al., 2010) | |
| **ATP:3-phospho-D-glycerate 1-phosphotransferase** | | | | | | **37** |
| Stoichiometry: ATP(aq) + 3-phospho-D-glycerate(aq) ⇌ ADP(aq) + 3-phospho-D-glyceroyl phosphate(aq) | | | | | | |
| $K_{eq}$ | $5.257 \cdot 10^{-4}$ | | | | eQuilibrator | |
| M | 41784 | | g $mol^{-1}$ | | UniProt | |
| Forward reaction: | | | | | | |
| $K_m$ (3-phosphoglycerate) | 180 | 20 | µM | | (Tsukamoto et al., 2013) | |
| $K_m$ (ATP) | 210 | 12 | µM | | (Tsukamoto et al., 2013) | |
| $K_{cat}^+$ | 403 | 19 | $s^{-1}$ | | (Tsukamoto et al., 2013) | |
| **D-glyceraldehyde-3-phosphate:NADP+ oxidoreductase (phosphorylating)** | | | | | | **38** |
| Stoichiometry: D-glyceraldehyde 3-phosphate(aq) + phosphate(aq) + NADP+(aq) ⇌ 3-phospho-D-glyceroyl phosphate(aq) + NADPH(aq) | | | | | | |
| $K_{eq}$ | 0.4176 | | | | eQuilibrator | |
| M | 150000 | | g $mol^{-1}$ | | (Erales et al., 2008) | |
| Forward reaction: | | | | | | |
| $K_m$ (phosphate) | 5000 | 5000 | µM | | This work | |
| Backward reaction: | | | | | | |
| $K_m$ (1,3-bisphosphoglycerate) | 210 | 12 | µM | | (Erales et al., 2008) | |
| $K_m$ (NADPH) | 140 | 6 | µM | | (Erales et al., 2008) | |
| $K_{cat}^-$ | 815 | 18 | $s^{-1}$ | | (Erales et al., 2008) | |
| **D-glyceraldehyde-3-phosphate aldose-ketose-isomerase** | | | | | | **39** |
| Stoichiometry: D-glyceraldehyde 3-phosphate(aq) ⇌ glycerone phosphate(aq) | | | | | | |
| $K_{eq}$ | 9.354 | | | | eQuilibrator | |
| M | 28213 | | g $mol^{-1}$ | | (Mathur et al., 2006) | |
| Forward reaction: | | | | | | |
| $K_m$ (glyceraldehyde-3-phosphate) | 84 | | µM | | (Mathur et al., 2006) | |
| $K_{cat}^+$ | 2680 | | $s^{-1}$ | | (Mathur et al., 2006) | |

Table S1 (continued)

| Enzyme name/ Parameter | Value | Uncer-tainty | Unit | Comment | Source | # |
|---|---|---|---|---|---|---|
| **D-fructose-1,6-bisphosphate D-glyceraldehyde-3-phosphate-lyase (glycerone-phosphate-forming)** | | | | | | **40** |
| Stoichiometry: D-fructose-1,6-bisphosphate(aq) ⇌ glycerone phosphate(aq) + D-glyceraldehyde 3-phosphate(aq) | | | | | | |
| $K_{eq}$ | $1.591 \cdot 10^{-4}$ | | | | eQuilibrator | |
| M | 66000 | | g $mol^{-1}$ | | (Nakahara et al., 2003) | |
| Forward reaction: | | | | | | |
| $K_m$ (fructose-1,6-bisphosphate) | 7 | | µM | | (Nakahara et al., 2003) | |
| $K_{cat}^{+}$ | 15.8 | | $s^{-1}$ | | (Nakahara et al., 2003) | |
| **D-fructose-1,6-bisphosphate 1-phosphohydrolase** | | | | | | **41** |
| Stoichiometry: D-fructose-1,6-bisphosphate(aq) + H2O(l) ⇌ D-fructose-6-phosphate(aq) + phosphate(aq) | | | | | | |
| $K_{eq}$ | 41.47 | | | | eQuilibrator | |
| M | 36520 | | g $mol^{-1}$ | | (Kelley-Loughnane et al., 2002) | |
| Forward reaction: | | | | | | |
| $K_m$ (fructose-1,6-bisphosphate) | 16 | 2 | µM | | (Kelley-Loughnane et al., 2002) | |
| $K_{cat}^{+}$ | 14.6 | 0.8 | $s^{-1}$ | | (Kelley-Loughnane et al., 2002) | |
| **D-fructose 6-phosphate:D-glyceraldehyde-3-phosphate glycolaldehyde transferase** | | | | | | **42** |
| Stoichiometry: D-fructose-6-phosphate(aq) + D-glyceraldehyde 3-phosphate(aq) ⇌ D-erythrose 4-phosphate(aq) + D-xylulose-5-phosphate(aq) | | | | | | |
| $K_{eq}$ | $1.673 \cdot 10^{-2}$ | | | | eQuilibrator | |
| M | 74143 | | g $mol^{-1}$ | | (Sprenger et al., 1995) | |
| Forward reaction: | | | | | | |
| $K_m$ (fructose-6-phosphate) | 1100 | | µM | | (Sprenger et al., 1995) | |
| $K_m$ (glyceraldehyde-3-phosphate) | 2300 | | µM | | (Sprenger et al., 1995) | |
| $K_{cat}^{+}$ | 62.28 | | $s^{-1}$ | | (Sprenger et al., 1995) | |
| Backward reaction: | | | | | | |
| $K_m$ (erythrose-4-phosphate) | 90 | | µM | | (Sprenger et al., 1995) | |
| $K_m$ (xylulose-5-phosphate) | 160 | | µM | | (Sprenger et al., 1995) | |
| $K_{cat}^{-}$ | 135.9 | | $s^{-1}$ | | (Sprenger et al., 1995) | |
| **sedoheptulose 1,7-bisphosphate D-glyceraldehyde-3-phosphate-lyase** | | | | | | **43** |
| Stoichiometry: sedoheptulose-1,7-bisphosphate(aq) ⇌ glycerone phosphate(aq) + D-erythrose 4-phosphate(aq) | | | | | | |
| $K_{eq}$ | $3.371 \cdot 10^{-3}$ | | | | eQuilibrator | |
| M | 33000 | | g $mol^{-1}$ | | (Nakahara et al., 2003) | |
| Forward reaction: | | | | | | |
| $K_m$ (sedoheptulose-1,7-bisphosphate) | 47 | | µM | | (Nakahara et al., 2003) | |
| $K_{cat}^{+}$ | 1.746 | | $s^{-1}$ | | (Nakahara et al., 2003) | |
| **sedoheptulose-1,7-bisphosphate 1-phosphohydrolase** | | | | | | **44** |
| Stoichiometry: sedoheptulose-1,7-bisphosphate(aq) + H2O(l) ⇌ sedoheptulose-7-phosphate(aq) + phosphate(aq) | | | | | | |
| $K_{eq}$ | 669.4 | | | | eQuilibrator | |
| M | 42081 | | g $mol^{-1}$ | | UniProt | |
| Forward reaction: | | | | | | |
| $K_m$ (sedoheptulose-1,7-bisphosphate) | 50 | | µM | | (Cadet & Meunier, 1988) | |
| $K_{cat}^{+}$ | 81 | | $s^{-1}$ | | (Cadet & Meunier, 1988) | |
| **sedoheptulose-7-phosphate:D-glyceraldehyde-3-phosphate glycolaldehydetransferase** | | | | | | **45** |
| Stoichiometry: sedoheptulose-7-phosphate(aq) + D-glyceraldehyde 3-phosphate(aq) ⇌ D-ribose-5-phosphate(aq) + D-xylulose-5-phosphate(aq) | | | | | | |
| $K_{eq}$ | 0.2104 | | | | eQuilibrator | |
| M | 74143 | | g $mol^{-1}$ | | (Sprenger et al., 1995) | |
| Forward reaction: | | | | | | |
| $K_m$ (sedoheptulose-7-phosphate) | 4000 | | µM | | (Sprenger et al., 1995) | |
| $K_m$ (glyceraldehyde 3-phosphate) | 2100 | | µM | | (Sprenger et al., 1995) | |
| $K_{cat}^{+}$ | 7.414 | | $s^{-1}$ | | (Sprenger et al., 1995) | |
| Backward reaction: | | | | | | |
| $K_m$ (ribose-5-phosphate) | 1500 | | µM | | (Sprenger et al., 1995) | |
| $K_m$ (xylulose-5-phosphate) | 160 | | µM | | (Sprenger et al., 1995) | |
| **D-ribulose-5-phosphate 3-epimerase** | | | | | | **46** |
| Stoichiometry: D-ribulose-5-phosphate(aq) ⇌ D-xylulose-5-phosphate(aq) | | | | | | |
| $K_{eq}$ | 3.897 | | | | eQuilibrator | |
| M | 23334 | | g $mol^{-1}$ | | UniProt | |
| Forward reaction: | | | | | | |
| $K_m$ (ribulose-5-phosphate) | 56.0 | 4.4 | µM | | (Le et al., 2017) | |
| $K_{cat}^{+}$ | 42.8 | 1.3 | $s^{-1}$ | | (Le et al., 2017) | |

Table S1 (continued)

| Enzyme name/ Parameter | Value | Uncer-tainty | Unit | Comment | Source | # |
|---|---|---|---|---|---|---|
| **D-ribose-5-phosphate aldose-ketose-isomerase** | | | | | | **47** |
| Stoichiometry: D-ribose-5-phosphate(aq) ⇌ D-ribulose-5-phosphate(aq) | | | | | | |
| $K_{eq}$ | 0.4421 | | | | eQuilibrator | |
| M | 45720 | | g $mol^{-1}$ | | (Zhang et al., 2003), UniProt | |
| Forward reaction: | | | | | | |
| $K_m$ (ribose-5-phosphate) | 3100 | 200 | µM | | (Zhang et al., 2003) | |
| $K_{cat}^{+}$ | 2100 | 300 | $s^{-1}$ | | (Zhang et al., 2003) | |
| **ATP:D-ribulose-5-phosphate 1-phosphotransferase** | | | | | | **48** |
| Stoichiometry: ATP + D-ribulose 5-phosphate = ADP + D-ribulose 1,5-bisphosphate | | | | | | |
| $K_{eq}$ | 9583 | | | | eQuilibrator | |
| M | 178000 | | g $mol^{-1}$ | | (Wadano et al., 1998) | |
| Forward reaction: | | | | | | |
| $K_m$ (ATP) | 90 | | µM | | (Wadano et al., 1998) | |
| $K_m$ (ribulose 5-phosphate) | 270 | | µM | | (Wadano et al., 1998) | |
| $K_{cat}^{+}$ | 215.4 | | $s^{-1}$ | | (Wadano et al., 1998) | |
| **phosphate:oxaloacetate carboxy-lyase (adding phosphate, phosphoenolpyruvate-forming)** | | | | | | **49** |
| Stoichiometry: phosphate(aq) + oxaloacetate(aq) ⇌ phosphoenolpyruvate(aq) + $HCO_3^-$(aq) | | | | | | |
| $K_{eq}$ | $1.863 \cdot 10^{-6}$ | | | | eQuilibrator | |
| M | 114000 | | g $mol^{-1}$ | | (Chang et al., 2014) | |
| Backward reaction: | | | | | | |
| $K_m$ (phosphoenolpyruvate) | 790 | 90 | µM | | (Chang et al., 2014) | |
| $K_m$ ($HCO_3^-$) | 169 | | µM | | (Chang et al., 2014) | |
| $K_{cat}^{-}$ | 42 | 3 | $s^{-1}$ | | (Chang et al., 2014) | |
| **2-phospho-D-glycerate hydro-lyase (phosphoenolpyruvate-forming)** | | | | | | **50** |
| Stoichiometry: 2-phospho-D-glycerate(aq) ⇌ phosphoenolpyruvate(aq) + H2O(l) | | | | | | |
| $K_{eq}$ | 5.191 | | | | eQuilibrator | |
| M | 46000 | | g $mol^{-1}$ | | (Zadvornyy et al., 2015) | |
| Forward reaction: | | | | | | |
| $K_m$ (phospho-D-glycerate) | 160 | 10 | µM | | (Zadvornyy et al., 2015) | |
| $K_{cat}^{+}$ | 95 | 4 | $s^{-1}$ | | (Zadvornyy et al., 2015) | |
| **D-phosphoglycerate 2,3-phosphomutase (2,3-diphosphoglycerate-independent)** | | | | | | **51** |
| Stoichiometry: 2-phospho-D-glycerate(aq) ⇌ 3-phospho-D-glycerate(aq) | | | | | | |
| $K_{eq}$ | 5.300 | | | | eQuilibrator | |
| M | 64000 | | g $mol^{-1}$ | | (GRAÑA et al., 1989) | |
| Forward reaction: | | | | | | |
| $K_m$ (2-phospho-D-glycerate) | 369 | 20 | µM | | (GRAÑA et al., 1989) | |
| Backward reaction: | | | | | | |
| $K_m$ (3-phospho-D-glycerate) | 324 | 20 | µM | | (GRAÑA et al., 1989) | |
| $K_{cat}^{-}$ | 216 | | $s^{-1}$ | | (GRAÑA et al., 1989) | |
| **D-arabino-hex-3-ulose-6-phosphate formaldehyde-lyase (D-ribulose-5-phosphate-forming)** | | | | | | **52** |
| Stoichiometry: D-arabino-hex-3-ulose-6-phosphate(aq) ⇌ D-ribulose-5-phosphate(aq) + formaldehyde(aq) | | | | | | |
| $K_{eq}$ | $2.312 \cdot 10^{-5}$ | | | | eQuilibrator | |
| M | 27000 | | g $mol^{-1}$ | | (Arfman et al., 1990) | |
| Backward reaction: | | | | | | |
| $K_m$ (ribulose-5-phosphate) | 450 | | µM | | (Arfman et al., 1990) | |
| $K_m$ (formaldehyde) | 147 | | µM | | (Arfman et al., 1990) | |
| $K_{cat}^{-}$ | 216 | | $s^{-1}$ | | (Arfman et al., 1990) | |
| **D-arabino-hex-3-ulose-6-phosphate isomerase** | | | | | | **53** |
| Stoichiometry: D-arabino-hex-3-ulose-6-phosphate(aq) ⇌ D-fructose-6-phosphate(aq) | | | | | | |
| $K_{eq}$ | 305.9 | | | | eQuilibrator | |
| M | 67000 | | g $mol^{-1}$ | | (Ferenci et al., 1974) | |
| Forward reaction: | | | | | | |
| $K_m$ (D-arabino-hex-3-ulose-6-phosphate) | 100 | | µM | | (Ferenci et al., 1974) | |
| $K_{cat}^{+}$ | 1742 | | $s^{-1}$ | | (Ferenci et al., 1974) | |
| Backward reaction: | | | | | | |
| $K_m$ (fructose-6-phosphate) | 1100 | | µM | | (Ferenci et al., 1974) | |
| **methanol:NAD+ oxidoreductase** | | | | | | **54** |
| Stoichiometry: methanol(aq) + NAD+(aq) ⇌ formaldehyde(aq) + NADH(aq) | | | | | | |
| $K_{eq}$ | $4.368 \cdot 10^{-6}$ | | | | eQuilibrator | |
| M | 40700 | | g $mol^{-1}$ | | (Wu et al., 2016) | |
| Forward reaction: | | | | | | |
| $K_m$ (methanol) | 21600 | 1500 | µM | | (Wu et al., 2016) | |
| $K_{cat}^{+}$ | 0.06 | 0.003 | $s^{-1}$ | Activity scaled down to appropriate pH | (Wu et al., 2016) | |

Table S1 (continued)

| Enzyme name/ Parameter | Value | Uncer-tainty | Unit | Comment | Source | # |
|---|---|---|---|---|---|---|
| **formaldehyde:NADP+ oxidoreductase (CoA-acetylating)** | | | | | | **55** |
| Stoichiometry: formaldehyde(aq) + CoA(aq) + NADP+(aq) ⇌ formyl-CoA(aq) + NADPH(aq) | | | | | | |
| $K_{eq}$ | $5.737 \cdot 10^{5}$ | | | | eQuilibrator | |
| M | 52000 | | g mol$^{-1}$ | | (Chou et al., 2019) | |
| Forward reaction: | | | | | | |
| $K_m$ (formaldehyde) | 10400 | 2200 | µM | | (Chou et al., 2019) | |
| $K_{cat}^{+}$ | 0.99 | 0.07 | s$^{-1}$ | | (Chou et al., 2019) | |
| **formyl-CoA:oxalate CoA-transferase** | | | | | | **56** |
| Stoichiometry: formyl-CoA(aq) + succinate(aq) ⇌ formate(aq) + succinyl-CoA(aq) | | | | | | |
| $K_{eq}$ | 0.01888 | | | | eQuilibrator | |
| M | 44000 | | g mol$^{-1}$ | | (Baetz & Allison, 1990) | |
| Forward reaction: | | | | | | |
| $K_m$ (formyl-CoA) | 3000 | 500 | µM | | (Baetz & Allison, 1990) | |
| $K_m$ (succinate) | 2300 | 600 | µM | | (Baetz & Allison, 1990) | |
| $K_{cat}^{+}$ | 14.08 | | s$^{-1}$ | | (Baetz & Allison, 1990) | |
| **ATP:D-fructose-6-phosphate 1-phosphotransferase** | | | | | | **57** |
| Stoichiometry: ATP(aq) + D-fructose-6-phosphate(aq) ⇌ ADP(aq) + D-fructose-1,6-bisphosphate(aq) | | | | | | |
| $K_{eq}$ | 1997 | | | | eQuilibrator | |
| M | 37000 | | g mol$^{-1}$ | | (Kotlarz & Buc, 1982) | |
| Forward reaction: | | | | | | |
| $K_m$ (ATP) | 50 | | µM | | (Kotlarz & Buc, 1982) | |
| $K_m$ (fructose-6-phosphate) | 11 | | µM | | (Kotlarz & Buc, 1982) | |
| $K_{cat}^{+}$ | 126.4 | | s$^{-1}$ | | (Kotlarz & Buc, 1982) | |
| **L-serine formaldehyde-lyase (glycine-forming)** | | | | | | **58** |
| Stoichiometry: L-serine(aq) ⇌ glycine(aq) + formaldehyde(aq) | | | | | | |
| $K_{eq}$ | $4.855 \cdot 10^{-4}$ | | | | eQuilibrator | |
| M | 48000 | | g mol$^{-1}$ | | (MIYAZAKI et al., 1987) | |
| Forward reaction: | | | | | | |
| $K_m$ (serine) | 150 | | µM | | (MIYAZAKI et al., 1987) | |
| $K_{cat}^{+}$ | 10.8 | | s$^{-1}$ | | (MIYAZAKI et al., 1987) | |
| Backward reaction: | | | | | | |
| $K_m$ (glycine) | 46 | | µM | | (MIYAZAKI et al., 1987) | |
| $K_m$ (formaldehyde) | 4800 | | µM | | (MIYAZAKI et al., 1987) | |
| $K_{cat}^{-}$ | 264 | | s$^{-1}$ | | (MIYAZAKI et al., 1987) | |
| **Formate:tetrahydrofolate ligase (ADP-forming)** | | | | | | **59** |
| Stoichiometry: tetrahydrofolate(aq) + formate(aq) + ATP(aq) ⇌ ADP(aq) + orthophosphate(aq) + 10-formyltetrahydrofolate(aq) | | | | | | |
| $K_{eq}$ | 3.696 | | | | eQuilibrator | |
| M | 240000 | | g mol$^{-1}$ | | (Marx et al., 2003) | |
| Forward reaction: | | | | | | |
| $K_m$ (tetrahydrofolate) | 800 | | µM | | (Marx et al., 2003) | |
| $K_m$ (formate) | 22000 | | µM | | (Marx et al., 2003) | |
| $K_m$ (ATP) | 21 | | µM | | (Marx et al., 2003) | |
| $K_{cat}^{+}$ | 468 | | s$^{-1}$ | | (Marx et al., 2003) | |
| **5,10-Methenyltetrahydrofolate 5-hydrolase (decyclizing)** | | | | | | **60** |
| Stoichiometry: 5,10-methenyltetrahydrofolate(aq) + H2O(l) ⇌ 10-formyltetrahydrofolate(aq) | | | | | | |
| $K_{eq}$ | 10.93 | | | | eQuilibrator | |
| M | 22000 | | g mol$^{-1}$ | | (Pomper et al., 1999) | |
| Forward reaction: | | | | | | |
| $K_m$ (5,10-methenyltetra-hydrofolate) | 35 | 4 | µM | | (Pomper et al., 1999) | |
| $K_{cat}^{+}$ | 231 | | s$^{-1}$ | | (Pomper et al., 1999) | |
| **5,10-methylenetetrahydrofolate:NADP+ oxidoreductase** | | | | | | **61** |
| Stoichiometry: 5,10-methylenetetrahydrofolate(aq) + NADP+(aq) ⇌ 5,10-methenyltetrahydrofolate(aq) + NADPH(aq) | | | | | | |
| $K_{eq}$ | 0.07062 | | | | eQuilibrator | |
| M | 60000 | | g mol$^{-1}$ | | (Dev & Harvey, 1978) | |
| Forward reaction: | | | | | | |
| $K_m$ (NADP) | 53 | 5 | µM | | (Dev & Harvey, 1978) | |
| $K_m$ (5,10-methylenetetra-hydrofolate) | 35 | 4 | µM | | (Dev & Harvey, 1978) | |
| $K_{cat}^{+}$ | 97.6 | 6 | s$^{-1}$ | | (Dev & Harvey, 1978) | |

Table S1 (continued)

| Enzyme name/ Parameter | Value | Uncer-tainty | Unit | Comment | Source | # |
|---|---|---|---|---|---|---|
| **5,10-methylenetetrahydrofolate:glycine hydroxymethyltransferase** | | | | | | **62** |
| Stoichiometry: 5,10-methylenetetrahydrofolate(aq) + glycine(aq) + H2O(l) ⇌ tetrahydrofolate(aq) + L-serine(aq) | | | | | | |
| $K_{eq}$ | 0.06931 | | | | eQuilibrator | |
| M | 94000 | | g $mol^{-1}$ | | (Schirch et al., 1985) | |
| Forward reaction: | | | | | | |
| $K_m$ (5,10-methylenetetra-hydrofolate) | 33 | | µM | | (Schirch et al., 1985) | |
| $K_m$ (glycine) | 850 | | µM | | (Schirch et al., 1985) | |
| Backward reaction: | | | | | | |
| $K_m$ (tetrahydrofolate) | 80 | | µM | | (Schirch et al., 1985) | |
| $K_m$ (serine) | 800 | | µM | | (Schirch et al., 1985) | |
| $K_{cat}^-$ | 21.3 | | $s^{-1}$ | | (Schirch et al., 1985) | |
| **L-serine:glyoxylate aminotransferase** | | | | | | **63** |
| Stoichiometry: L-serine(aq) + glyoxylate(aq) ⇌ 3-hydroxypyruvate(aq) + glycine(aq) | | | | | | |
| $K_{eq}$ | 5.502 | | | | eQuilibrator | |
| M | 150000 | | g $mol^{-1}$ | | (IZUMI et al., 1990) | |
| Forward reaction: | | | | | | |
| $K_m$ (serine) | 280 | 20 | µM | | (Karsten et al., 2001) | |
| $K_m$ (glyoxylate) | 230 | | µM | | (IZUMI et al., 1990) | |
| $K_{cat}^+$ | 173.75 | | $s^{-1}$ | | (IZUMI et al., 1990) | |
| **D-glycerate:NAD+ oxidoreductase** | | | | | | **64** |
| Stoichiometry: D-glycerate + NAD+ = hydroxypyruvate + NADH + H+ | | | | | | |
| $K_{eq}$ | $1.132 \cdot 10^{-5}$ | | | | eQuilibrator | |
| M | 71000 | | g $mol^{-1}$ | | (Chistoserdova & Lidstrom, 1991) | |
| Forward reaction: | | | | | | |
| $K_m$ (glycerate) | 2600 | | µM | | (Chistoserdova & Lidstrom, 1991) | |
| $K_{cat}^+$ | 11.98 | | $s^{-1}$ | | (Chistoserdova & Lidstrom, 1991) | |
| Backward reaction: | | | | | | |
| $K_m$ (hydroxypyruvate) | 40 | | µM | | (Chistoserdova & Lidstrom, 1991) | |
| $K_m$ (NADH) | 100 | | µM | | (Chistoserdova & Lidstrom, 1991) | |
| $K_{cat}^-$ | 798.8 | | $s^{-1}$ | | (Chistoserdova & Lidstrom, 1991) | |
| **ATP:D-glycerate 2-phosphotransferase** | | | | | | **65** |
| Stoichiometry: ATP(aq) + D-glycerate(aq) ⇌ ADP(aq) + 2-phospho-D-glycerate(aq) | | | | | | |
| $K_{eq}$ | 49.65 | | | | eQuilibrator | |
| M | 39000 | | g $mol^{-1}$ | | UniProt | |
| Forward reaction: | | | | | | |
| $K_m$ (ATP) | 121 | 16 | µM | | (Bartsch et al., 2008) | |
| $K_m$ (glycerate) | 86 | 1 | µM | | (Bartsch et al., 2008) | |
| $K_{cat}^+$ | 302 | 7 | $s^{-1}$ | | (Bartsch et al., 2008) | |
| **(S)-methylmalonyl-CoA:pyruvate carboxytransferase** | | | | | | **66** |
| Stoichiometry: malonyl-CoA(aq) + pyruvate(aq) ⇌ acetyl-CoA(aq) + oxaloacetate(aq) | | | | | | |
| $K_{eq}$ | 0.7468 | | | | eQuilibrator | |
| M | 670000 | | g $mol^{-1}$ | | (Wood et al., 1969) | |
| Forward reaction: | | | | | | |
| $K_m$ (malonyl-CoA) | 350 | | µM | | (Wood et al., 1969) | |
| $K_m$ (pyruvate) | 760 | | µM | | (Wood et al., 1969) | |
| Backward reaction: | | | | | | |
| $K_m$ (acetyl-CoA) | 560 | | µM | | (Wood et al., 1969) | |
| $K_m$ (oxaloacetate) | 570 | | µM | | (Wood et al., 1969) | |
| $K_{cat}^-$ | 233 | | $s^{-1}$ | | (Wood et al., 1969) | |
| **[protein]-S8-aminomethyldihydrolipoyllysine:tetrahydrofolate aminomethyltransferase (ammonia-forming) (T-protein)** | | | | | | **67** |
| Stoichiometry: tetrahydrofolate(aq) + S(8)-aminomethyldihydrolipoamide(aq) ⇌ $NH_3$(aq) + 5,10-methylenetetrahydrofolate(aq) + dihydrolipoamide(aq) | | | | | | |
| $K_{eq}$ | 0.3198 | | | | eQuilibrator | |
| M | 40147 | | g $mol^{-1}$ | | UniProt | |
| Backward reaction: | | | | | | |
| $K_m$ ($NH_3$) | 9.2 | | µM | | (Okamura-Ikeda et al., 2003) | |
| $K_m$ (5,10-methylenetetra-hydrofolate) | 88.1 | | µM | | (Okamura-Ikeda et al., 2003) | |
| $K_m$ (dihydrolipoamide) | 0.69 | | µM | | (Okamura-Ikeda et al., 2003) | |
| $K_{cat}^-$ | 19.4 | | $s^{-1}$ | | (Okamura-Ikeda et al., 2003) | |

Table S1 (continued)

| Enzyme name/ Parameter | Value | Uncer-tainty | Unit | Comment | Source | # |
|---|---|---|---|---|---|---|
| **glycine:H-protein-lipoyllysine oxidoreductase (decarboxylating, acceptor-amino-methylating) (P-protein)** | | | | | | **68** |
| Stoichiometry: glycine(aq) + lipoamide(aq) ⇌ CO2(aq) + S(8)-aminomethyldihydrolipoamide(aq) | | | | | | |
| $K_{eq}$ | $9.202 \cdot 10^{-4}$ | | | | eQuilibrator | |
| M | 100000 | | g $mol^{-1}$ | | (Fujiwara & Motokawa, 1983) | |
| Forward reaction: | | | | | | |
| $K_m$ (glycine) | 5800 | | µM | | (Fujiwara & Motokawa, 1983) | |
| $K_m$ (lipoamide) | 3.4 | | µM | | (Fujiwara & Motokawa, 1983) | |
| $K_{cat}^{+}$ | 6.8 | | $s^{-1}$ | | (Fujiwara & Motokawa, 1983) | |
| Backward reaction: | | | | | | |
| $K_m$ (CO2) | 5400 | | µM | | (Fujiwara & Motokawa, 1983) | |
| $K_m$ (S(8)-aminomethyl-dihydrolipoamide) | 1.6 | | µM | | (Fujiwara & Motokawa, 1983) | |
| **protein-N6-(dihydrolipoyl)lysine:NAD+ oxidoreductase (L-protein)** | | | | | | **69** |
| Stoichiometry: lipoamide(aq) + NADH(aq) ⇌ dihydrolipoamide(aq) + NAD+(aq) | | | | | | |
| $K_{eq}$ | 1.907 | | | | eQuilibrator | |
| M | 126000 | | g $mol^{-1}$ | | (Neuburger et al., 2000) | |
| Backward reaction: | | | | | | |
| $K_m$ (dihydrolipoamide) | 27 | 3 | µM | | (Neuburger et al., 2000) | |
| $K_{cat}^{-}$ | 345 | 10 | $s^{-1}$ | | (Neuburger et al., 2000) | |
| **malyl-CoA lyase (formyl-CoA forming)** | | | | | | **70** |
| Stoichiometry: malonate semialdehyde(aq) + formyl-CoA(aq) ⇌ malyl-CoA(aq) | | | | | | |
| $K_{eq}$ | 110.8 | | | | eQuilibrator | |
| M | 59116 | | g $mol^{-1}$ | | (Chou et al., 2019) | |
| Forward reaction: | | | | | | |
| $K_m$ (malonate semialdehyde) | 16000 | 4000 | µM | It was assumed that malonate semi-aldehyde behaves similar to propanal | (Chou et al., 2019) | |
| $K_m$ (formyl-CoA) | 200 | 50 | µM | | (Chou et al., 2019) | |
| $K_{cat}^{+}$ | 4.7 | 0.4 | $s^{-1}$ | | (Chou et al., 2019) | |
| **2-hydroxyglutaryl-CoA lyase (formyl-CoA forming)** | | | | | | **71** |
| Stoichiometry: succinic semialdehyde(aq) + formyl-CoA(aq) ⇌ 2-hydroxyglutaryl-CoA(aq) | | | | | | |
| $K_{eq}$ | 105.1 | | | | eQuilibrator | |
| M | 59116 | | g $mol^{-1}$ | | (Chou et al., 2019) | |
| Forward reaction: | | | | | | |
| $K_m$ (succinic semialdehyde) | 2500 | 600 | µM | It was assumed that succinic semialde-hyde behaves like a mixture of propanal and pentanal | (Chou et al., 2019) | |
| $K_m$ (formyl-CoA) | 200 | 50 | µM | | (Chou et al., 2019) | |
| $K_{cat}^{+}$ | 3.1 | 0.4 | $s^{-1}$ | | (Chou et al., 2019) | |
| **succinyl-CoA:2-hydroxyglutarate CoA-transferase** | | | | | | **72** |
| Stoichiometry: 2-hydroxyglutaryl-CoA(aq) + succinate(aq) ⇌ 2-hydroxyglutarate(aq) + succinyl-CoA(aq) | | | | | | |
| $K_{eq}$ | 0.2827 | | | | eQuilibrator | |
| M | 90800 | | g $mol^{-1}$ | | (Friedmann et al., 2006) | |
| Backward reaction: | | | | | | |
| $K_m$ (2-hydroxyglutarate) | 1300 | 300 | µM | It was assumed that 2-hydroxyglutarate behaves similar to 2-hydroxysuccinate (malate). Note that the CoA will be transferred to the other carboxy group | (Friedmann et al., 2006) | |
| $K_m$ (succinyl-CoA) | 500 | 100 | µM | | (Friedmann et al., 2006) | |
| $K_{cat}^{-}$ | 11.35 | | $s^{-1}$ | | (Friedmann et al., 2006) | |

Table S1 (continued)

| Enzyme name/ Parameter | Value | Uncer-tainty | Unit | Comment | Source | # |
|---|---|---|---|---|---|---|
| **2-hydroxyglutarate oxaloacetate transhydrogenase** | | | | | | **73** |
| Stoichiometry: oxaloacetate(aq) + 2-hydroxyglutarate(aq) ⇌ malate(aq) + alpha-ketoglutarate(aq) | | | | | | |
| $K_{eq}$ | 22.58 | | | | eQuilibrator | |
| M | 99000 | | g $mol^{-1}$ | | (Allen, 1966) | |
| Forward reaction: | | | | | | |
| $K_m$ (oxaloacetate) | 50 | | µM | | (Allen, 1966) | |
| $K_m$ (2-hydroxyglutarate) | 1400 | | µM | | (Allen, 1966) | |
| $K_{cat}^+$ | 1.98 | | $s^{-1}$ | | (Allen, 1966) | |
| Backward reaction: | | | | | | |
| $K_m$ (malate) | 1400 | | µM | | (Allen, 1966) | |
| $K_m$ (alpha-ketoglutarate) | 50 | | µM | | (Allen, 1966) | |
| **isocitrate:NADP+ oxidoreductase (decarboxylating)** | | | | | | **74** |
| Stoichiometry: isocitrate(aq) + NADP+(aq) ⇌ 2-oxoglutarate(aq) + CO2(aq) + NADPH(aq) | | | | | | |
| $K_{eq}$ | 0.08955 | | | | eQuilibrator | |
| M | 80000 | | g $mol^{-1}$ | | (Kanao et al., 2002) | |
| Forward reaction: | | | | | | |
| $K_m$ (isocitrate) | 45 | 13 | µM | | (Kanao et al., 2002) | |
| $K_m$ (NADP) | 27 | 10 | µM | | (Kanao et al., 2002) | |
| $K_{cat}^+$ | 200 | 8 | $s^{-1}$ | | (Kanao et al., 2002) | |
| Backward reaction: | | | | | | |
| $K_m$ (2-oxoglutarate) | 1100 | 500 | µM | | (Kanao et al., 2002) | |
| $K_m$ (CO2) | 1300 | 300 | µM | | (Kanao et al., 2002) | |
| $K_{cat}^-$ | 51 | 12 | $s^{-1}$ | | (Kanao et al., 2002) | |
| **isocitrate hydro-lyase (cis-aconitate-forming)** | | | | | | **75** |
| Stoichiometry: isocitrate(aq) ⇌ cis-aconitate(aq) + H2O(l) | | | | | | |
| $K_{eq}$ | 0.7572 | | | | eQuilibrator | |
| M | 102000 | | g $mol^{-1}$ | | (Baumgart & Bott, 2011) | |
| Forward reaction: | | | | | | |
| $K_m$ (isocitrate) | 552 | 302 | µM | | (Baumgart & Bott, 2011) | |
| $K_{cat}^+$ | 28.05 | | $s^{-1}$ | | (Baumgart & Bott, 2011) | |
| Backward reaction: | | | | | | |
| $K_m$ (cis-aconitate) | 18.5 | 3.4 | µM | | (Baumgart & Bott, 2011) | |
| $K_{cat}^-$ | 69.02 | | $s^{-1}$ | | (Baumgart & Bott, 2011) | |
| **citrate hydro-lyase (cis-aconitate-forming)** | | | | | | **76** |
| Stoichiometry: citrate(aq) ⇌ cis-aconitate(aq) + H2O(l) | | | | | | |
| $K_{eq}$ | 0.03482 | | | | eQuilibrator | |
| M | 102000 | | g $mol^{-1}$ | | (Baumgart & Bott, 2011) | |
| Forward reaction: | | | | | | |
| $K_m$ (citrate) | 480 | 20 | µM | | (Baumgart & Bott, 2011) | |
| $K_{cat}^+$ | 19.04 | | $s^{-1}$ | | (Baumgart & Bott, 2011) | |
| Backward reaction: | | | | | | |
| $K_m$ (cis-aconitate) | 18.5 | 3.4 | µM | | (Baumgart & Bott, 2011) | |
| $K_{cat}^-$ | 69.02 | | $s^{-1}$ | | (Baumgart & Bott, 2011) | |
| **acetyl-CoA:oxaloacetate C-acetyltransferase [(pro-S)-carboxymethyl-forming, ADP-phosphorylating]** | | | | | | **77** |
| Stoichiometry: ATP(aq) + citrate(aq) + CoA(aq) ⇌ ADP(aq) + orthophosphate(aq) + acetyl-CoA(aq) + oxaloacetate(aq) | | | | | | |
| $K_{eq}$ | 0.01334 | | | | eQuilibrator | |
| M | 623000 | | g $mol^{-1}$ | | (Houston & Nimmo, 1984) | |
| Forward reaction: | | | | | | |
| $K_m$ (ATP) | 222 | 28 | µM | | (Ranganathan et al., 1980) | |
| $K_m$ (citrate) | 40 | 10 | µM | | (Ranganathan et al., 1980) | |
| $K_m$ (CoA) | 2.6 | | µM | | (Ranganathan et al., 1980) | |
| $K_{cat}^+$ | 141.2 | | $s^{-1}$ | | (Houston & Nimmo, 1984) | |
| **L-serine ammonia-lyase (pyruvate-forming)** | | | | | | **18** |
| Stoichiometry: L-serine(aq) ⇌ pyruvate(aq) + NH3(aq) | | | | | | |
| $K_{eq}$ | $1.213 \cdot 10^5$ | | | | eQuilibrator | |
| M | 48236 | | g $mol^{-1}$ | | (Gannon et al., 1977) | |
| Forward reaction: | | | | | | |
| $K_m$ (serine) | 7000 | | µM | | (Gannon et al., 1977) | |
| $K_{cat}^+$ | 530.6 | | $s^{-1}$ | | (Gannon et al., 1977) | |

Table S1 (continued)

| Enzyme name/ Parameter | Value | Uncer-tainty | Unit | Comment | Source | # |
|---|---|---|---|---|---|---|
| **malonate semialdehyde:NADP+ oxidoreductase (malonate semialdehyde-forming)** | | | | | | **78** |
| Stoichiometry: malonyl-CoA(aq) + NADPH(aq) ⇌ malonate semialdehyde(aq) + NADP(aq) + CoA(aq) | | | | | | |
| $K_{eq}$ | $9.017 \cdot 10^{-3}$ | | | | eQuilibrator | |
| M | 300000 | | g $mol^{-1}$ | | (Hügler et al., 2002) | |
| Forward reaction: | | | | | | |
| $K_m$ (malonyl-CoA) | 30 | | µM | | (Hügler et al., 2002) | |
| $K_m$ (NADPH) | 25 | | µM | | (Hügler et al., 2002) | |
| $K_{cat}^+$ | 50 | | $s^{-1}$ | | (Hügler et al., 2002) | |
| **diphosphate---fructose-6-phosphate 1-phosphotransferase** | | | | | | **79** |
| Stoichiometry: fructose-1,6-bisphosphate(aq) + phosphate(aq) ⇌ fructose-6-phosphate(aq) + pyrophosphate(aq) | | | | | | |
| $K_{eq}$ | 0.2221 | | | | eQuilibrator | |
| M | 90000 | | g $mol^{-1}$ | | (Reshetnikov et al., 2008) | |
| Forward reaction: | | | | | | |
| $K_m$ (fructose-1,6-bisphosphate) | 328 | | µM | | (Reshetnikov et al., 2008) | |
| $K_m$ (phosphate) | 8690 | | µM | | (Reshetnikov et al., 2008) | |
| $K_{cat}^+$ | 13.5 | | $s^{-1}$ | | (Reshetnikov et al., 2008) | |
| Backward reaction: | | | | | | |
| $K_m$ (fructose-6-phosphate) | 2270 | | µM | | (Reshetnikov et al., 2008) | |
| $K_m$ (pyrophosphate) | 27 | | µM | | (Reshetnikov et al., 2008) | |
| $K_{cat}^-$ | 11.4 | | $s^{-1}$ | | (Reshetnikov et al., 2008) | |
| **diphosphate---ribulose-5-phosphate 1-phosphotransferase** | | | | | | **80** |
| Stoichiometry: ribulose-1,5-bisphosphate(aq) + phosphate(aq) ⇌ ribulose-5-phosphate(aq) + pyrophosphate(aq) | | | | | | |
| $K_{eq}$ | 0.02098 | | | | eQuilibrator | |
| M | 90000 | | g $mol^{-1}$ | | (Reshetnikov et al., 2008) | |
| Forward reaction: | | | | | | |
| $K_m$ (phosphate) | 8690 | | µM | | (Reshetnikov et al., 2008) | |
| $K_{cat}^+$ | 6.3 | | $s^{-1}$ | | (Reshetnikov et al., 2008) | |
| Backward reaction: | | | | | | |
| $K_m$ (pyrophosphate) | 27 | | µM | | (Reshetnikov et al., 2008) | |
| $K_{cat}^-$ | 4.5 | | $s^{-1}$ | | (Reshetnikov et al., 2008) | |
| **diphosphate---Sedoheptulose-7-phosphate 1-phosphotransferase** | | | | | | **81** |
| Stoichiometry: sedoheptulose-bisphosphate(aq) + phosphate(aq) ⇌ sedoheptulose-7-phosphate(aq) + pyrophosphate(aq) | | | | | | |
| $K_{eq}$ | 1.625 | | | | eQuilibrator | |
| M | 90000 | | g $mol^{-1}$ | | (Reshetnikov et al., 2008) | |
| Forward reaction: | | | | | | |
| $K_m$ (phosphate) | 8690 | | µM | | (Reshetnikov et al., 2008) | |
| $K_{cat}^+$ | 33.75 | | $s^{-1}$ | | (Reshetnikov et al., 2008) | |
| Backward reaction: | | | | | | |
| $K_m$ (sedoheptulose-7-phosphate) | 30 | | µM | | (Reshetnikov et al., 2008) | |
| $K_m$ (pyrophosphate) | 27 | | µM | | (Reshetnikov et al., 2008) | |
| $K_{cat}^-$ | 46.5 | | $s^{-1}$ | | (Reshetnikov et al., 2008) | |
| **ATP:pyruvate, water phosphotransferase (PEP synthase)** | | | | | | **82** |
| Stoichiometry: ATP(aq) + pyruvate(aq) + H2O(l) ⇌ AMP(aq) + phosphoenolpyruvate(aq) + phosphate(aq) | | | | | | |
| $K_{eq}$ | 2.410 | | | | eQuilibrator | |
| M | 77000 | | g $mol^{-1}$ | | (NARINDRASORASAK & BRIDGER, 1977) | |
| Forward reaction: | | | | | | |
| $K_m$ (pyruvate) | 83 | | µM | | (Berman & Cohn, 1970) | |
| $K_m$ (ATP) | 24 | | µM | | (Berman & Cohn, 1970) | |
| $K_{cat}^+$ | 10.27 | | $s^{-1}$ | | (NARINDRASORASAK & BRIDGER, 1977) | |
| **malate dehydrogenase (oxaloacetate-decarboxylating) (NADP+)** | | | | | | **83** |
| Stoichiometry: malate(aq) + NADP(aq) ⇌ pyruvate(aq) + CO2(aq) + NADPH(aq) | | | | | | |
| $K_{eq}$ | 0.004876 | | | | eQuilibrator | |
| M | 330000 | | g $mol^{-1}$ | | (Ziegler, 1974) | |
| Forward reaction: | | | | | | |
| $K_m$ (malate) | 700 | | µM | | (Häusler et al., 1987) | |
| $K_m$ (NADP) | 15 | | µM | | (Ziegler, 1974) | |
| $K_{cat}^+$ | 346.5 | | $s^{-1}$ | | (Häusler et al., 1987) | |
| Backward reaction: | | | | | | |
| $K_m$ (pyruvate) | 3000 | | µM | | (Ziegler, 1974) | |
| $K_m$ (CO2) | 1200 | | µM | | (Häusler et al., 1987) | |
| $K_m$ (NADPH) | 45 | | µM | | (Ziegler, 1974) | |

Table S1 (continued)

| Enzyme name/ Parameter | Value | Uncer-tainty | Unit | Comment | Source | # |
|---|---|---|---|---|---|---|
| **ATP:oxaloacetate carboxy-lyase (transphosphorylating; phosphoenolpyruvate-forming)** | | | | | | **84** |
| Stoichiometry: oxaloacetate(aq) + ATP(aq) ⇌ phosphoenolpyruvate(aq) + ADP(aq) + $CO_2$(aq) | | | | | | |
| $K_{eq}$ | 0.01152 | | | | eQuilibrator | |
| M | 70682 | | g $mol^{-1}$ | | UniProt | |
| Forward reaction: | | | | | | |
| $K_m$ (oxaloacetate) | 156 | 2 | µM | | (Chen et al., 2002) | |
| $K_m$ (ATP) | 25.7 | 3.5 | µM | | (Chen et al., 2002) | |
| $K_{cat}^{+}$ | 47 | 6 | $s^{-1}$ | | (Chen et al., 2002) | |
| Backward reaction: | | | | | | |
| $K_m$ (phosphoenolpyruvate) | 2300 | 250 | µM | | (Chen et al., 2002) | |
| $K_m$ (ADP) | 36000 | 3300 | µM | | (Chen et al., 2002) | |
| $K_m$ ($CO_2$) | 55 | 18 | µM | | (Chen et al., 2002) | |
| $K_{cat}^{-}$ | 22 | 2 | $s^{-1}$ | | (Chen et al., 2002) | |
| **Pyruvate:NAD+-oxidoreductase (decarboxylating, acceptor-acetylating) (pyruvate dehydrogenase complex)** | | | | | | **85** |
| Stoichiometry: pyruvate(aq) + CoA(aq) + NAD+(aq) ⇌ acetyl-CoA(aq) + $CO_2$(aq) + NADH(aq) | | | | | | |
| $K_{eq}$ | $1.732 \cdot 10^{6}$ | | | | eQuilibrator | |
| M | 200000 | | g $mol^{-1}$ | | (SAUMWEBER et al., 1981) | |
| Forward reaction: | | | | | | |
| $K_m$ (pyruvate) | 73 | | µM | | (SAUMWEBER et al., 1981) | |
| $K_m$ (NAD+) | 70 | | µM | | (Wang et al., 2010)Wa | |
| $K_{cat}^{+}$ | 5970 | | $s^{-1}$ | | (SAUMWEBER et al., 1981) | |
| **ATP:pyruvate 2-O-phosphotransferase** | | | | | | **86** |
| Stoichiometry: phosphoenolpyruvate(aq) + ADP(aq) ⇌ pyruvate(aq) + ATP(aq) | | | | | | |
| $K_{eq}$ | $2.174 \cdot 10^{3}$ | | | | eQuilibrator | |
| M | 185000 | | g $mol^{-1}$ | | (Abbe & Yamada, 1982) | |
| Forward reaction: | | | | | | |
| $K_m$ (phosphoenolpyruvate) | 220 | | µM | | (Abbe & Yamada, 1982) | |
| $K_m$ (ADP) | 390 | | µM | | (Abbe & Yamada, 1982) | |
| $K_{cat}^{+}$ | 363.8 | | $s^{-1}$ | | (Abbe & Yamada, 1982) | |
| **isocitrate glyoxylate-lyase (succinate-forming)** | | | | | | **87** |
| Stoichiometry: succinate(aq) + glyoxylate(aq) ⇌ isocitrate(aq) | | | | | | |
| $K_{eq}$ | 43.42 | | | | eQuilibrator | |
| M | 177000 | | g $mol^{-1}$ | | (MacKintosh & Nimmo, 1988) | |
| Forward reaction: | | | | | | |
| $K_m$ (succinate) | 590 | | µM | | (MacKintosh & Nimmo, 1988) | |
| $K_m$ (glyoxylate) | 130 | | µM | | (MacKintosh & Nimmo, 1988) | |
| Backward reaction: | | | | | | |
| $K_m$ (isocitrate) | 63 | 4 | µM | | (MacKintosh & Nimmo, 1988) | |
| $K_{cat}^{-}$ | 110.6 | | $s^{-1}$ | | (MacKintosh & Nimmo, 1988) | |
| **(S)-2-hydroxy carboxylate:oxygen 2-oxidoreductase** | | | | | | **88** |
| Stoichiometry: glycolate(aq) + $O_2$(aq) ⇌ glyoxylate(aq) + $H_2O_2$(aq) | | | | | | |
| $K_{eq}$ | $3.737 \cdot 10^{15}$ | | | | eQuilibrator | |
| M | 40924 | | g $mol^{-1}$ | | (Pennati & Gadda, 2009) | |
| Forward reaction: | | | | | | |
| $K_m$ (glycolate) | 200 | 10 | µM | | (Pennati & Gadda, 2009) | |
| $K_m$ ($O_2$) | 440 | 20 | µM | | (Pennati & Gadda, 2009) | |
| $K_{cat}^{+}$ | 20 | 0.4 | $s^{-1}$ | | (Pennati & Gadda, 2009) | |
| **3-phospho-D-glycerate carboxy-lyase (dimerizing; D-ribulose-1,5-bisphosphate-forming)** | | | | | | **89** |
| Stoichiometry: ribulose-1,5-bisphosphate(aq) + $O_2$(aq) ⇌ 3-phosphoglycerate(aq) + 2-phosphoglycolate(aq) | | | | | | |
| $K_{eq}$ | $2.389 \cdot 10^{92}$ | | | | eQuilibrator | |
| M | 72268 | | g $mol^{-1}$ | | (Carmo-Silva et al., 2010) | |
| Forward reaction: | | | | | | |
| $K_m$ (ribulose-1,5-bisphosphate) | 100 | | µM | | (Carmo-Silva et al., 2010) | |
| $K_m$ ($O_2$) | 341 | 33 | µM | | (Carmo-Silva et al., 2010) | |
| $K_{cat}^{+}$ | 0.95 | 0.04 | $s^{-1}$ | | (Carmo-Silva et al., 2010) | |
| **2-phosphoglycolate phosphohydrolase** | | | | | | **90** |
| Stoichiometry: 2-phosphoglycolate(aq) + $H_2O$(l) ⇌ glycolate(aq) + phosphate(aq) | | | | | | |
| $K_{eq}$ | $1.030 \cdot 10^{5}$ | | | | eQuilibrator | |
| M | 33012 | | g $mol^{-1}$ | | UniProt | |
| Forward reaction: | | | | | | |
| $K_m$ (2-phosphoglycolate) | 26 | | µM | | (Husic & Tolbert, 1984) | |
| $K_{cat}^{+}$ | 41.65 | | $s^{-1}$ | | (Husic & Tolbert, 1984) | |

Table S1 (continued)

| **Enzyme name/ Parameter** | **Value** | **Uncer-tainty** | **Unit** | **Comment** | **Source** | **#** |
|---|---|---|---|---|---|---|
| **(2R,3S)-beta-Hydroxyasparate hydro-lyase (Iminosuccinate forming)** | | | | | | **91** |
| Stoichiometry: 3-hydroxyaspartate(aq) ⇌ iminosuccinate(aq) + H2O(l) | | | | | | |
| $K_{eq}$ | 3.343 | | | | eQuilibrator | |
| M | 34015 | | g $mol^{-1}$ | | (Schada von Borzyskowski et al., 2019), UniProt | |
| Forward reaction: | | | | | | |
| $K_m$ (3-hydroxyaspartate) | 200 | 20 | µM | | (Schada von Borzyskowski et al., 2019) | |
| $K_{cat}^{+}$ | 35 | 1 | $s^{-1}$ | | (Schada von Borzyskowski et al., 2019) | |
| **aspartate:NAD+ oxidoreductase** | | | | | | **92** |
| Stoichiometry: iminosuccinate(aq) + NADH(aq) ⇌ aspartate(aq) + NAD+(aq) | | | | | | |
| $K_{eq}$ | $4.385 \cdot 10^{10}$ | | | | eQuilibrator | |
| M | 33655 | | g $mol^{-1}$ | | (Schada von Borzyskowski et al., 2019), UniProt | |
| Forward reaction: | | | | | | |
| $K_m$ (iminosuccinate) | 90 | 10 | µM | | (Schada von Borzyskowski et al., 2019) | |
| $K_m$ (NADH) | 20 | 3 | µM | | (Schada von Borzyskowski et al., 2019) | |
| $K_{cat}^{+}$ | 201 | 10 | $s^{-1}$ | | (Schada von Borzyskowski et al., 2019) | |

#### 2.2. Concentration bounds for metabolites

Table S2: Concentration ranges used for the Enzyme Cost Minimization algorithm

| Compound name | Compound ID | KEGG entry | Lower bound, [M] | Upper bound, [M] | Comment |
|---|---|---|---|---|---|
| H2O | C_h2o | C00001 | 1 | 1 | |
| ATP | C_atp | C00002 | 0.005 | 0.005 | ATP/ADP ratio of 100 |
| NAD | C_nad | C00003 | 0.001 | 0.001 | |
| NADH | C_nadh | C00004 | 0.00001 | 0.00001 | |
| NADPH | C_nadph | C00005 | 0.001 | 0.001 | |
| NADP | C_nadp | C00006 | 1.00E-05 | 1.00E-05 | |
| ADP | C_adp | C00008 | 0.00005 | 0.00005 | ATP/ADP ratio of 100 |
| phosphate | C_orthophosphate | C00009 | 0.01 | 0.01 | |
| CoA | C_coa | C00010 | 0.001 | 0.001 | |
| CO2 | C_co2 | C00011 | 1.00E-05 | 1.00E-05 | Order of magnitude of air saturated water (~350-400 ppm CO2) |
| pyrophosphate | C_diphosphate | C00013 | 0.001 | 0.001 | |
| NH3 | C_nh3 | C00014 | 0.0001 | 0.01 | |
| AMP | C_amp | C00020 | 0.0005 | 0.0005 | |
| pyruvate | C_pyruvate | C00022 | 1.00E-06 | 0.01 | |
| acetyl-CoA | C_acetyl_coa | C00024 | 1.00E-06 | 0.01 | |
| 2-oxoglutarate | C_2_oxoglutarate | C00026 | 1.00E-06 | 0.01 | |
| oxaloacetate | C_oxaloacetate | C00036 | 1.00E-06 | 0.01 | |
| glycine | C_glycine | C00037 | 1.00E-06 | 0.01 | |
| succinate | C_succinate | C00042 | 1.00E-06 | 0.01 | |
| glyoxylate | C_glyoxylate | C00048 | 1.00E-06 | 0.01 | |
| aspartate | C_l_aspartate | C00049 | 1.00E-06 | 0.01 | |
| formate | C_formate | C00058 | 1.00E-06 | 0.01 | |
| serine | C_l_serine | C00065 | 1.00E-06 | 0.01 | |
| formaldehyde | C_formaldehyde | C00067 | 1.00E-09 | 0.01 | |
| phosphoenolpyruvate | C_phosphoenolpyruvate | C00074 | 1.00E-06 | 0.01 | |
| malonyl-CoA | C_malonyl_coa | C00083 | 1.00E-06 | 0.01 | |
| fructose-6-phosphate | C_d_fructose_6_phosphoric_acid | C00085 | 1.00E-06 | 0.01 | |
| succinyl-CoA | C_succinyl_coa | C00091 | 1.00E-06 | 0.01 | |
| propanoyl-CoA | C_propanoyl_coa | C00100 | 1.00E-06 | 0.01 | |
| tetrahydrofolate | C_tetrahydrofolate | C00101 | 1.00E-06 | 0.01 | |
| glycerone phosphate | C_glycerone_phosphate | C00111 | 1.00E-06 | 0.01 | |
| ribose-5-phosphate | C_d_ribose_5_phosphate | C00117 | 1.00E-06 | 0.01 | |
| glyceraldehyde-3-phosphate | C_d_glyceraldehyde_3_phosphate | C00118 | 1.00E-06 | 0.01 | |
| fumarate | C_fumarate | C00122 | 1.00E-06 | 0.01 | |
| methanol | C_methanol | C00132 | 1.00E-06 | 0.1 | Methanol can be used at high concentrations in a bioreactor |
| 5,10-methylenetetrahydrofolate | C_510_methylenetetrahydrofolate | C00143 | 1.00E-06 | 0.01 | |
| malate | C_s_malate | C00149 | 1.00E-06 | 0.01 | |
| citrate | C_citrate | C00158 | 1.00E-06 | 0.01 | |
| hydroxypyruvate | C_hydroxypyruvate | C00168 | 1.00E-06 | 0.01 | |

Table S2 (continued)

| Compound name | Compound ID | KEGG entry | Lower bound, [M] | Upper bound, [M] | Comment |
|---|---|---|---|---|---|
| 3-phosphoglycerate | C_3_phospho_d_glycerate | C00197 | 1.00E-06 | 0.01 | |
| ribulose-5-phosphate | C_d_ribulose_5_phosphate | C00199 | 1.00E-06 | 0.01 | |
| malonate semialdehyde | C_3_oxopropanoate | C00222 | 1.00E-06 | 0.01 | |
| xylulose-5-phosphate | C_d_xylulose_5_phosphate | C00231 | 1.00E-06 | 0.01 | |
| succinate semialdehyde | C_succinate_semialdehyde | C00232 | 1.00E-06 | 0.01 | |
| 10-formyltetrahydrofolate | C_10_formyltetrahydrofolate | C00234 | 1.00E-06 | 0.01 | |
| 1,3-bisphosphoglycerate | C_3_phospho_d_glyceroyl_phosphate | C00236 | 1.00E-06 | 0.01 | |
| lipoamide | C_lipoamide | C00248 | 1.00E-08 | 0.0005 | Bounds for lipoamide are lower as it is supposed to be attached to the H-protein |
| glycerate | C_d_glycerate | C00258 | 1.00E-06 | 0.01 | |
| erythrose-4-phosphate | C_d_erythrose_4_phosphate | C00279 | 1.00E-06 | 0.01 | |
| isocitrate | C_isocitrate | C00311 | 1.00E-06 | 0.01 | |
| acetoacetyl-CoA | C_acetoacetyl_coa | C00332 | 1.00E-06 | 0.01 | |
| fructose-1,6-bisphosphate | C_d_fructose_16_bisphosphate | C00354 | 1.00E-06 | 0.01 | |
| ubiquinol | C_ubiquinol | C00390 | 1.00E-07 | 1.00E-04 | Quinones are located in the inner membrane and thus the concentration is assumed to be lower |
| ubiquinone | C_ubiquinone | C00399 | 1.00E-07 | 1.00E-04 | |
| cis-aconitate | C_cis_aconitate | C00417 | 1.00E-06 | 0.01 | |
| 5,10-methenyltetrahydrofolate | C_510_methenyltetrahydrofolate | C00445 | 1.00E-06 | 0.01 | |
| seduheptulose-1,7-bisphosphate | C_d_altro_heptulose_17_biphosphate | C00447 | 1.00E-06 | 0.01 | |
| benzoquinone | C_p_benzoquinone | C00472 | 1.00E-06 | 0.01 | |
| hydroquinone | C_hydroquinone | C00530 | 1.00E-06 | 0.01 | |
| dihydrolipoamide | C_dihydrolipoamide | C00579 | 1.00E-08 | 0.0005 | Bounds for lipoamide are lower as it is supposed to be attached to the H-protein |
| 2-phosphoglycerate | C_2_phospho_d_glycerate | C00631 | 1.00E-06 | 0.01 | |
| S-methylmalonyl-CoA | C_s_methylmalonyl_coa | C00683 | 1.00E-06 | 0.01 | |
| formyl-CoA | C_formyl_coa | C00798 | 1.00E-06 | 0.01 | |
| menaquinone | C_menaquinone | C00828 | 1.00E-07 | 1.00E-04 | Quinones are located in the inner membrane and thus the concentration is assumed to be lower |
| crotonyl-CoA | C_crotonoyl_coa | C00877 | 1.00E-06 | 0.01 | |
| acrylyl-CoA | C_propenoyl_coa | C00894 | 1.00E-06 | 0.01 | |
| 4-hydroxybutanoate | C_4_hydroxybutanoic_acid | C00989 | 1.00E-06 | 0.01 | |
| citramalyl-CoA | C_citramalyl_coa | C01011 | 1.00E-06 | 0.01 | |
| 3-hydroxypropanoate | C_3_hydroxypropanoate | C01013 | 1.00E-06 | 0.01 | |
| 3S-hydroxybutanoyl-CoA | C_s_3_hydroxybutanoyl_coa | C01144 | 1.00E-06 | 0.01 | |
| ribulose-1,5-bisphosphate | C_d_ribulose_15_biphosphate | C01182 | 1.00E-06 | 0.01 | |
| R-Methylmalonyl-CoA | C_r_methylmalonyl_coa | C01213 | 1.00E-06 | 0.01 | |
| $HCO_3^-$ | C_hco3_ | C01353 | 1.00E-06 | 0.00026 | Order of magnitude of air saturated water (~350-400 ppm CO2) |
| 2-hydroxyglutarate | C_2_hydroxyglutarate | C02630 | 1.00E-06 | 0.01 | |

Table S2 (continued)

| **Compound name** | **Compound ID** | **KEGG entry** | **Lower bound, [M]** | **Upper bound, [M]** | **Comment** |
|---|---|---|---|---|---|
| 2-hydroxyglutaryl-CoA | C_2_hydroxyglutaryl_coa | C03058 | 1.00E-06 | 0.01 | |
| β-hydroxyasparatate | C_erythro_3_hydroxy_ls_aspartate | C03961 | 1.00E-06 | 0.01 | |
| malyl-CoA | C_3s_3_carboxy_3_hydroxypropanoyl_coa | C04348 | 1.00E-06 | 0.01 | |
| seduheptulose-7-phosphate | C_d_sedoheptulose_7_phosphate | C05382 | 1.00E-06 | 0.01 | |
| 3-hydroxypropanoyl-CoA | C_3_hydroxypropionyl_coa | C05668 | 1.00E-06 | 0.01 | |
| menaquinol | C_menaquinol | C05819 | 1.00E-07 | 1.00E-04 | Quinones are located in the inner membrane and thus the concentration is assumed to be lower |
| iminoaspartate | C_iminoaspartate | C05840 | 1.00E-06 | 0.01 | |
| hexulose-6-phosphate | C_d_arabino_6_phospho_hex_3_ulose | C06019 | 1.00E-06 | 0.01 | |
| β-methylmalyl-CoA | C_l_erythro_3_methylmalyl_coa | C06027 | 1.00E-06 | 0.01 | |
| mesaconyl-C1-CoA | C_mesaconyl_coa | C06028 | 1.00E-06 | 0.01 | |
| 4-hydroxybutanoyl-CoA | C_4_hydroxybutyryl_coa | C11062 | 1.00E-06 | 0.01 | |
| S-ethylmalonyl-CoA | C_s_ethylmalonyl_coa | C18026 | 1.00E-06 | 0.01 | |
| mesaconyl-C4-CoA | C_mesaconyl_c4_coa | C18323 | 1.00E-06 | 0.01 | |
| methylsuccinyl-CoA | C_methylsuccinyl_coa | C18324 | 1.00E-06 | 0.01 | |
| R-ethylmalonyl-CoA | C_r_ethylmalonyl_coa | C20238 | 1.00E-06 | 0.01 | |
| S(8)-aminomethyldihydrolipoamide | C_s8_aminomethyldihydrolipoamide | C90640 | 1.00E-08 | 0.0005 | Bounds for lipoamide are lower as it is supposed to be attached to the H-protein |
| O2 | C_oxygen | C00007 | 0.000273 | 0.000273 | |
| H2O2 | C_hydrogen_peroxide | C00027 | 1.00E-06 | 0.01 | |
| glycolate | C_glycolate | C00160 | 1.00E-06 | 0.01 | |
| 2-phosphoglycolate | C_2_phosphoglycolate | C00988 | 1.00E-06 | 0.01 | |

#### 3. Detailed pathway analysis with single enzyme contributions

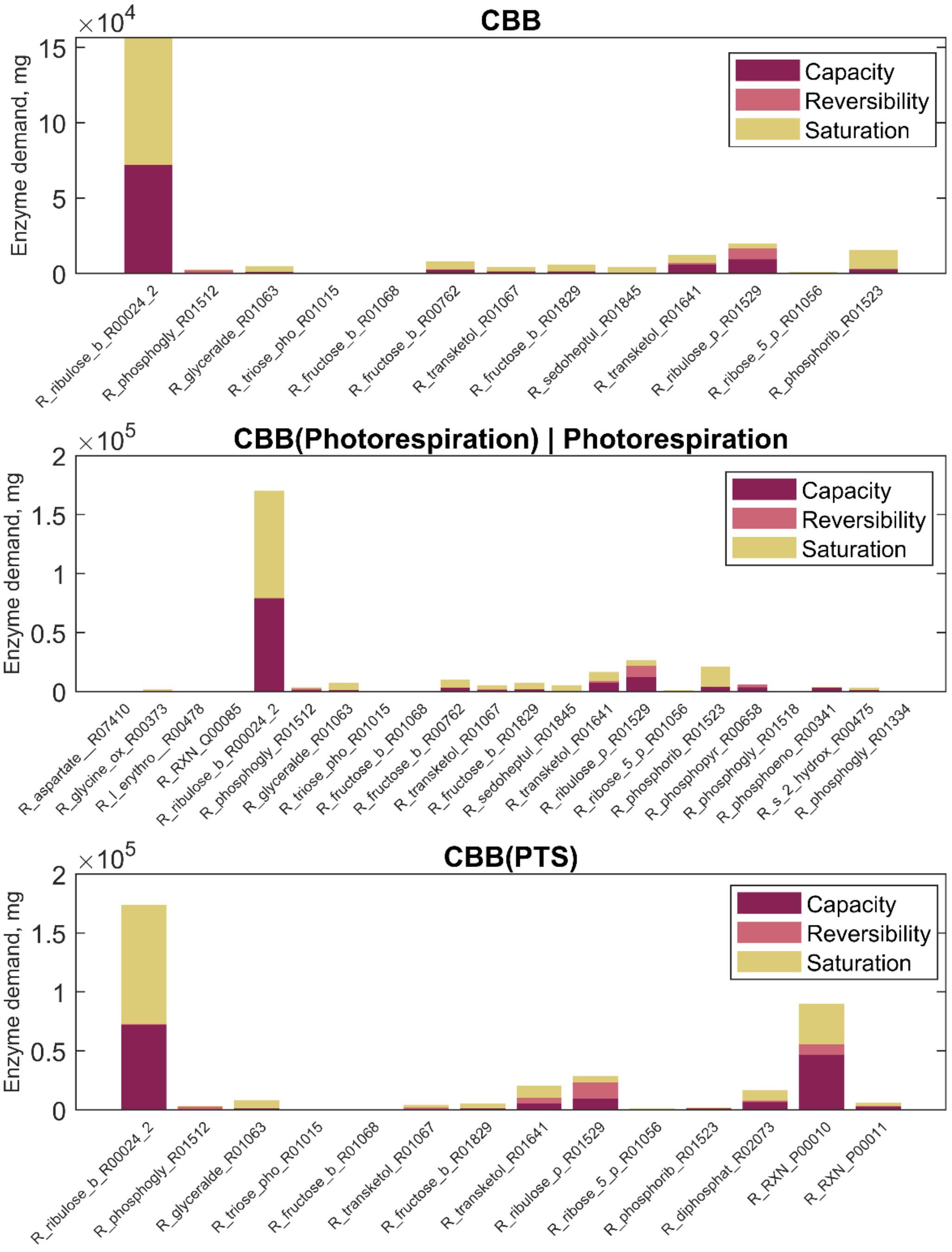


Figure S10: Enzyme demands to sustain a total pathway activity of 1 mmol product per second. Contribution of the capacity, the reversibility and the saturation with substrates or products of each reaction to the demand of enzymes. The figure follows the wording of (Noor et al., 2016). “Capacity”: demand of enzyme caused by a limitation by the catalytic rate constant. “Reversibility”: extra amount of enzyme needed because of a backward flux. “Saturation”: additional enzyme necessary because of undersaturation with a substrate or oversaturation with a product. The values present the optimized state as predicted by the ECM algorithm assuming a $CO_2$ concentration of 10 µM. Glyceraldehyde-3-phosphate was chosen as a product for the cycles in this case. Reaction names correspond to their identifiers in the SBtab model file (Supplementary file Reactions_Composite17_model.tsv)

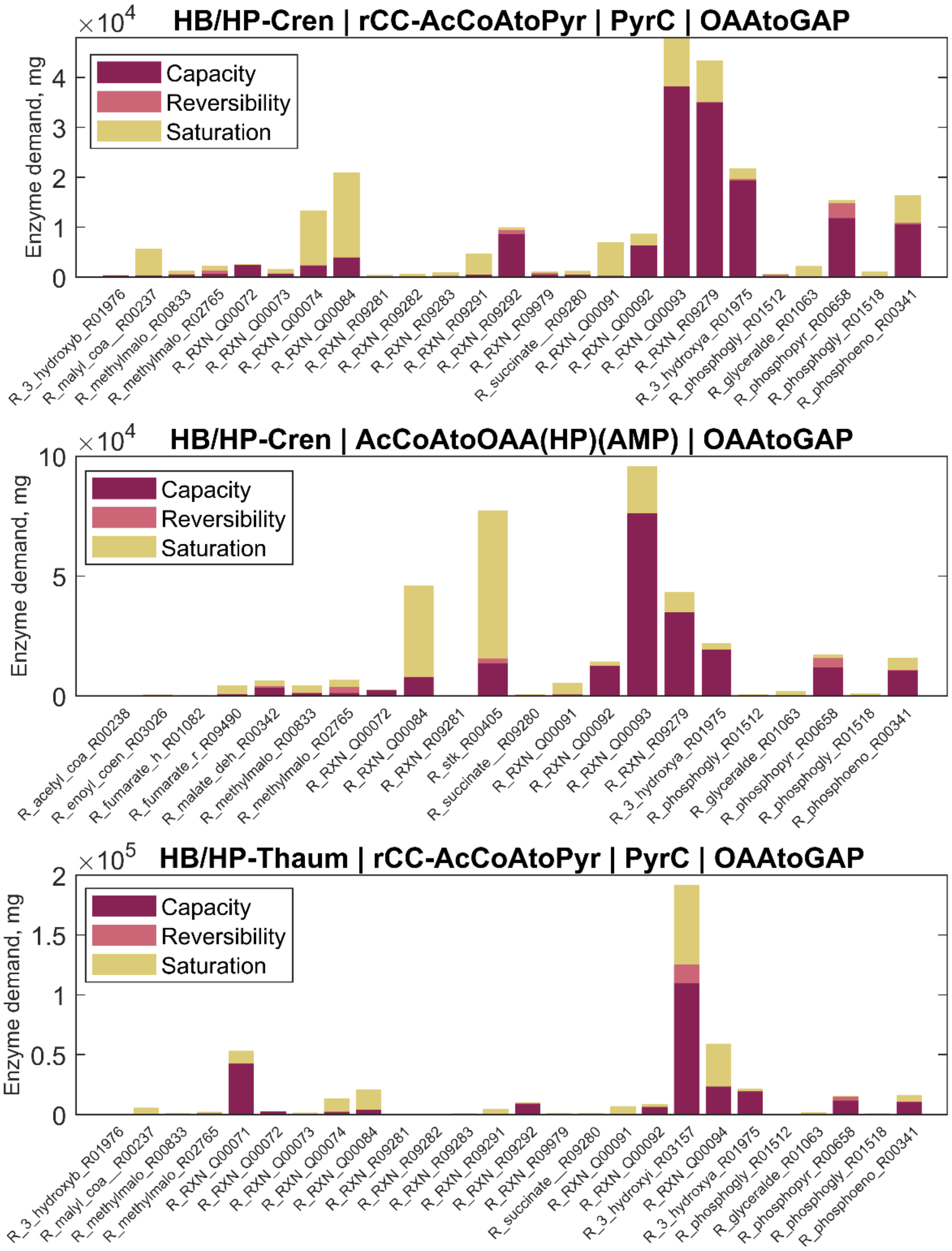


Figure S11: Enzyme demands to sustain a total pathway activity of 1 mmol product per second. Contribution of the capacity, the reversibility and the saturation with substrates or products of each reaction to the demand of enzymes. The figure follows the wording of (Noor et al., 2016). "Capacity": demand of enzyme caused by a limitation by the catalytic rate constant. "Reversibility": extra amount of enzyme needed because of a backward flux. "Saturation": additional enzyme necessary because of undersaturation with a substrate or oversaturation with a product. The values present the optimized state as predicted by the ECM algorithm assuming a $CO_2$ concentration of 10 µM. Glyceraldehyde-3-phosphate was chosen as a product for the cycles in this case. Reaction names correspond to their identifiers in the SBtab model file (Supplementary file Reactions_Composite17_model.tsv)

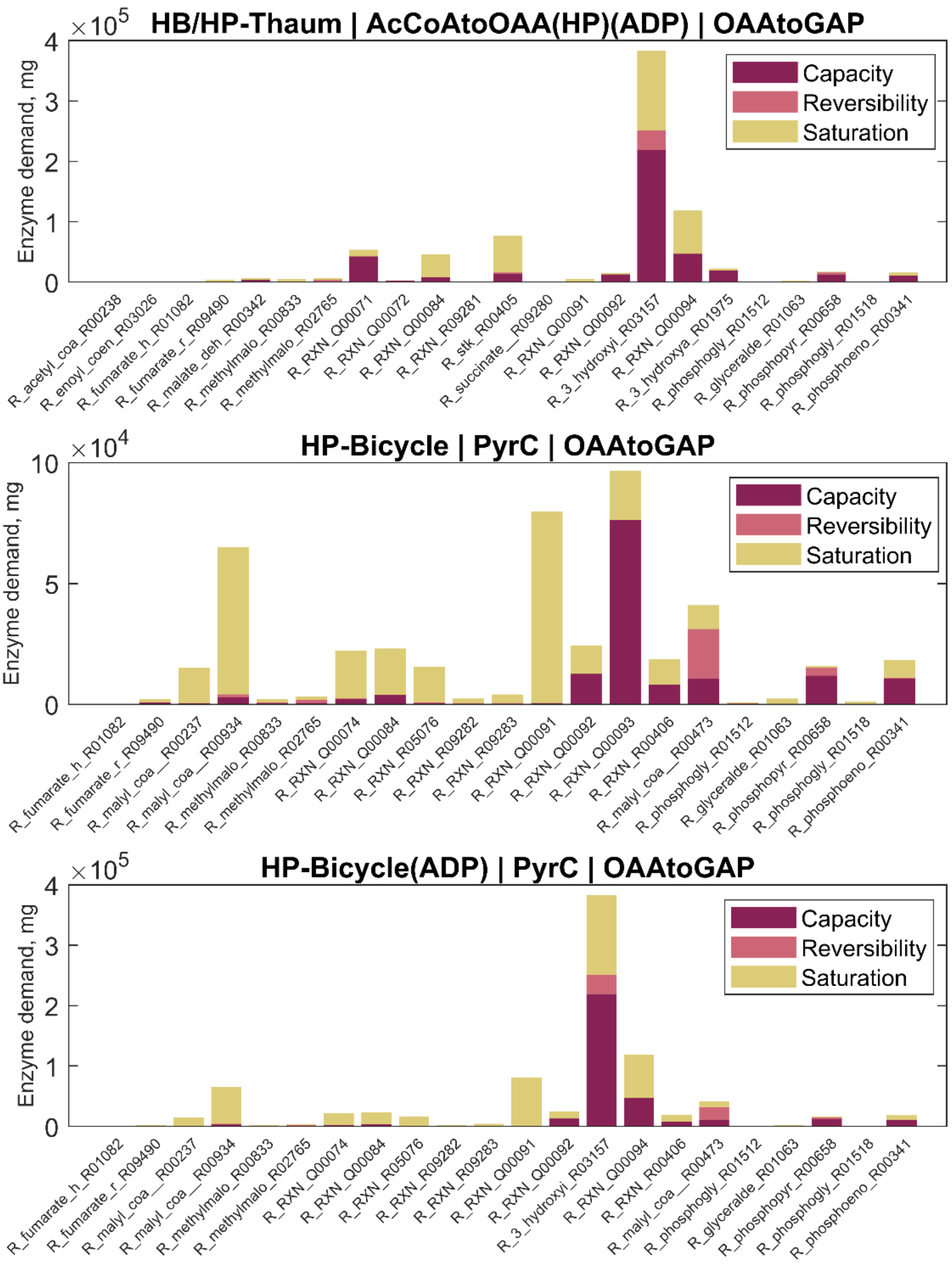


Figure S12: Enzyme demands to sustain a total pathway activity of 1 mmol product per second. Contribution of the capacity, the reversibility and the saturation with substrates or products of each reaction to the demand of enzymes. The figure follows the wording of (Noor et al., 2016). “Capacity”: demand of enzyme caused by a limitation by the catalytic rate constant. “Reversibility”: extra amount of enzyme needed because of a backward flux. “Saturation”: additional enzyme necessary because of undersaturation with a substrate or oversaturation with a product. The values present the optimized state as predicted by the ECM algorithm assuming a CO2 concentration of 10 µM. Glyceraldehyde-3-phosphate was chosen as a product for the cycles in this case. Reaction names correspond to their identifiers in the SBtab model file (Supplementary file Reactions_Composite17_model.tsv)

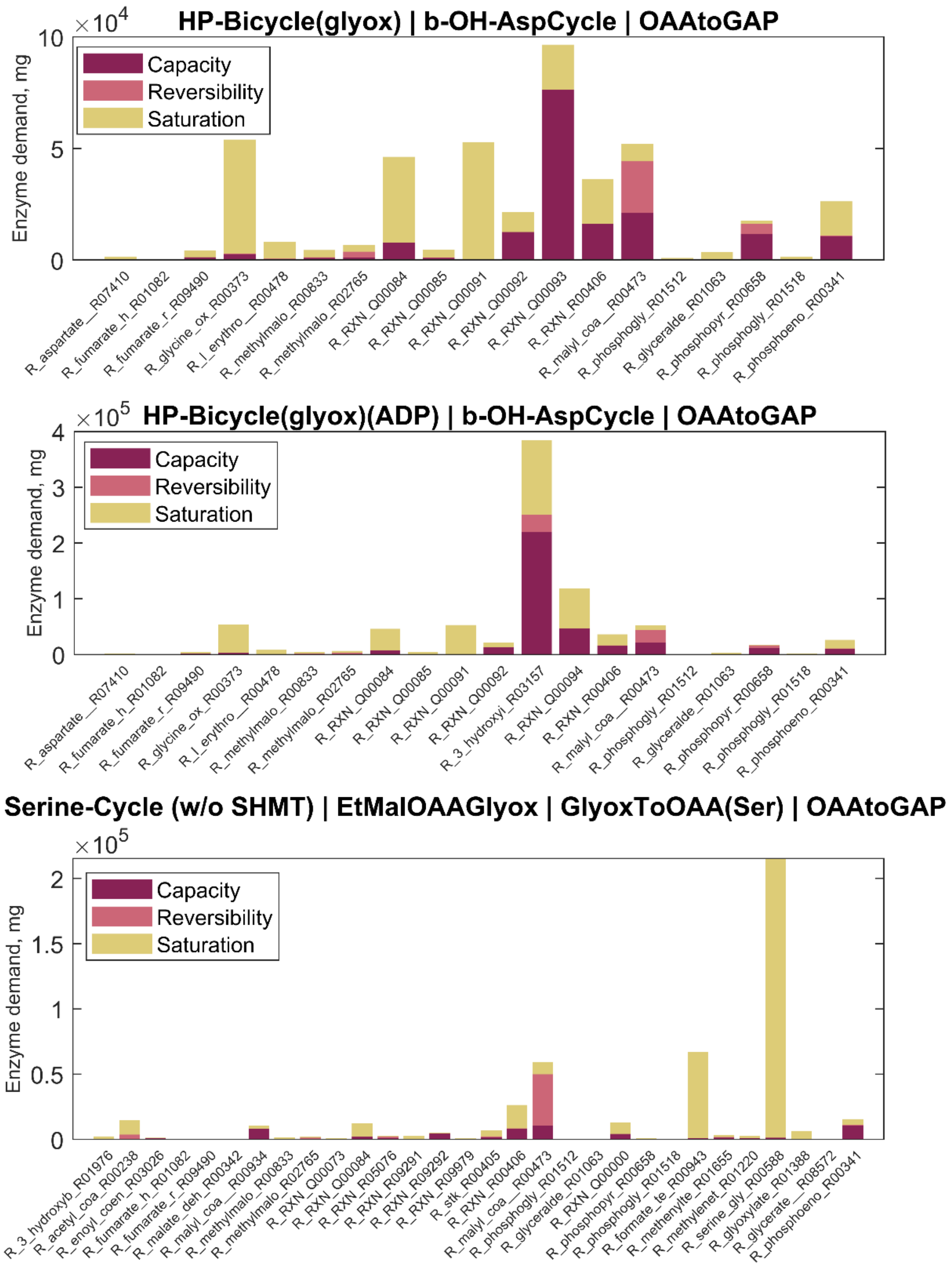


Figure S13: Enzyme demands to sustain a total pathway activity of 1 mmol product per second. Contribution of the capacity, the reversibility and the saturation with substrates or products of each reaction to the demand of enzymes. The figure follows the wording of (Noor et al., 2016). “Capacity”: demand of enzyme caused by a limitation by the catalytic rate constant. “Reversibility”: extra amount of enzyme needed because of a backward flux. “Saturation”: additional enzyme necessary because of undersaturation with a substrate or oversaturation with a product. The values present the optimized state as predicted by the ECM algorithm assuming a $CO_2$ concentration of 10 µM. Glyceraldehyde-3-phosphate was chosen as a product for the cycles in this case. Reaction names correspond to their identifiers in the SBtab model file (Supplementary file Reactions_Composite17_model.tsv)

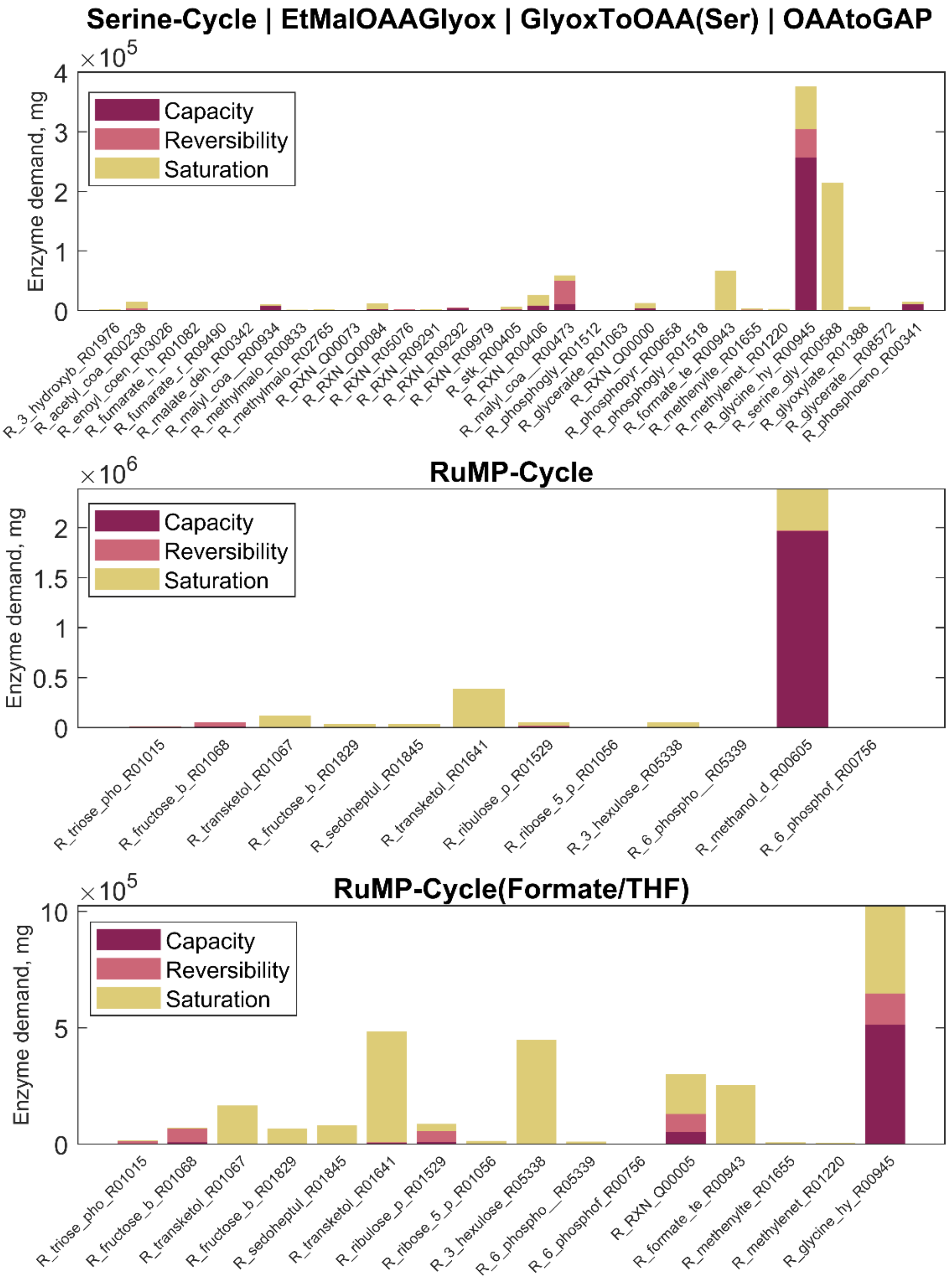


Figure S14: Enzyme demands to sustain a total pathway activity of 1 mmol product per second. Contribution of the capacity, the reversibility and the saturation with substrates or products of each reaction to the demand of enzymes. The figure follows the wording of (Noor et al., 2016). “Capacity”: demand of enzyme caused by a limitation by the catalytic rate constant. “Reversibility”: extra amount of enzyme needed because of a backward flux. “Saturation”: additional enzyme necessary because of undersaturation with a substrate or oversaturation with a product. The values present the optimized state as predicted by the ECM algorithm assuming a $CO_2$ concentration of 10 µM. Glyceraldehyde-3-phosphate was chosen as a product for the cycles in this case. Reaction names correspond to their identifiers in the SBtab model file (Supplementary file Reactions_Composite17_model.tsv)

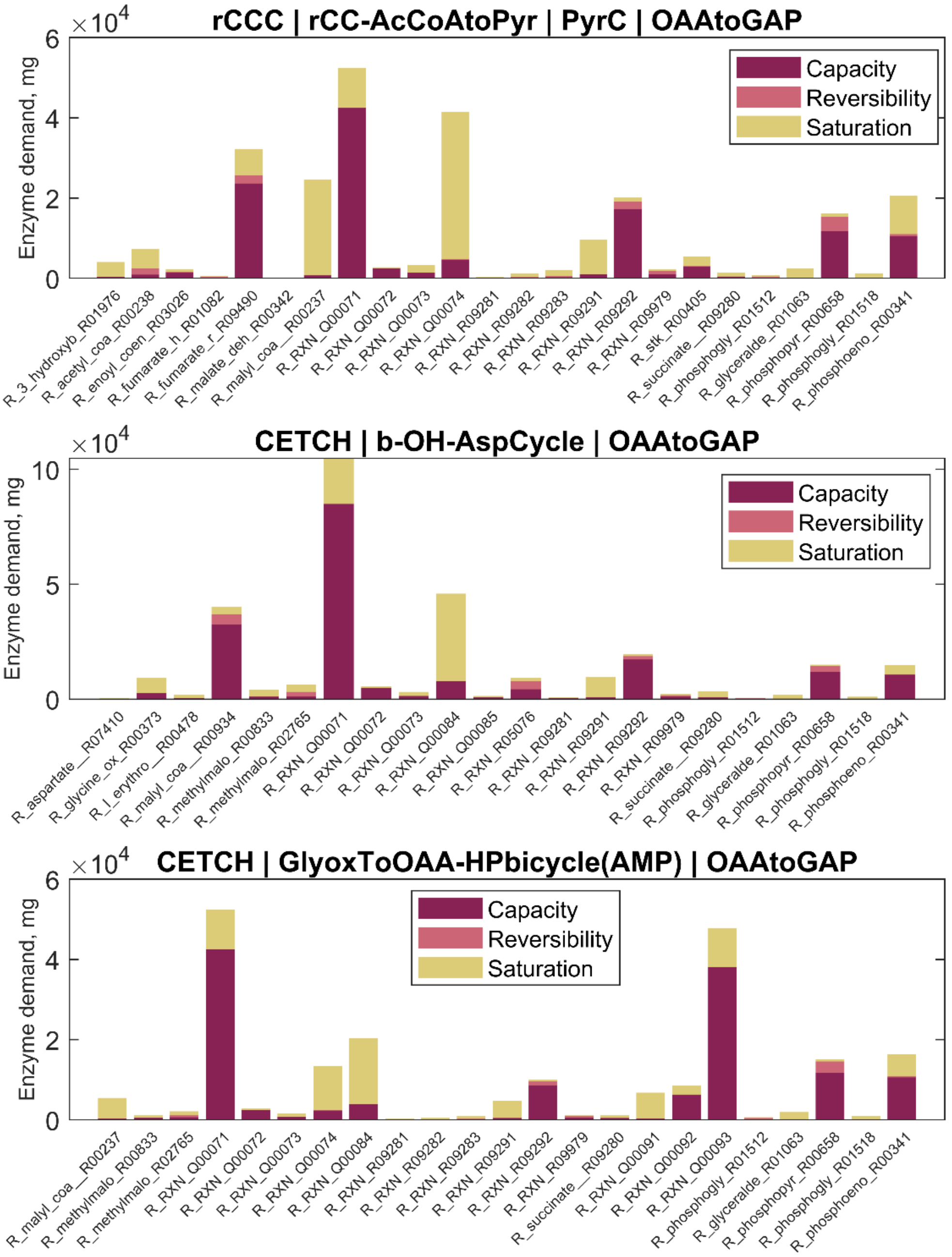


Figure S15: Enzyme demands to sustain a total pathway activity of 1 mmol product per second. Contribution of the capacity, the reversibility and the saturation with substrates or products of each reaction to the demand of enzymes. The figure follows the wording of (Noor et al., 2016). "Capacity": demand of enzyme caused by a limitation by the catalytic rate constant. "Reversibility": extra amount of enzyme needed because of a backward flux. "Saturation": additional enzyme necessary because of undersaturation with a substrate or oversaturation with a product. The values present the optimized state as predicted by the ECM algorithm assuming a CO2 concentration of 10 µM. Glyceraldehyde-3-phosphate was chosen as a product for the cycles in this case. Reaction names correspond to their identifiers in the SBtab model file (Supplementary file Reactions_Composite17_model.tsv)

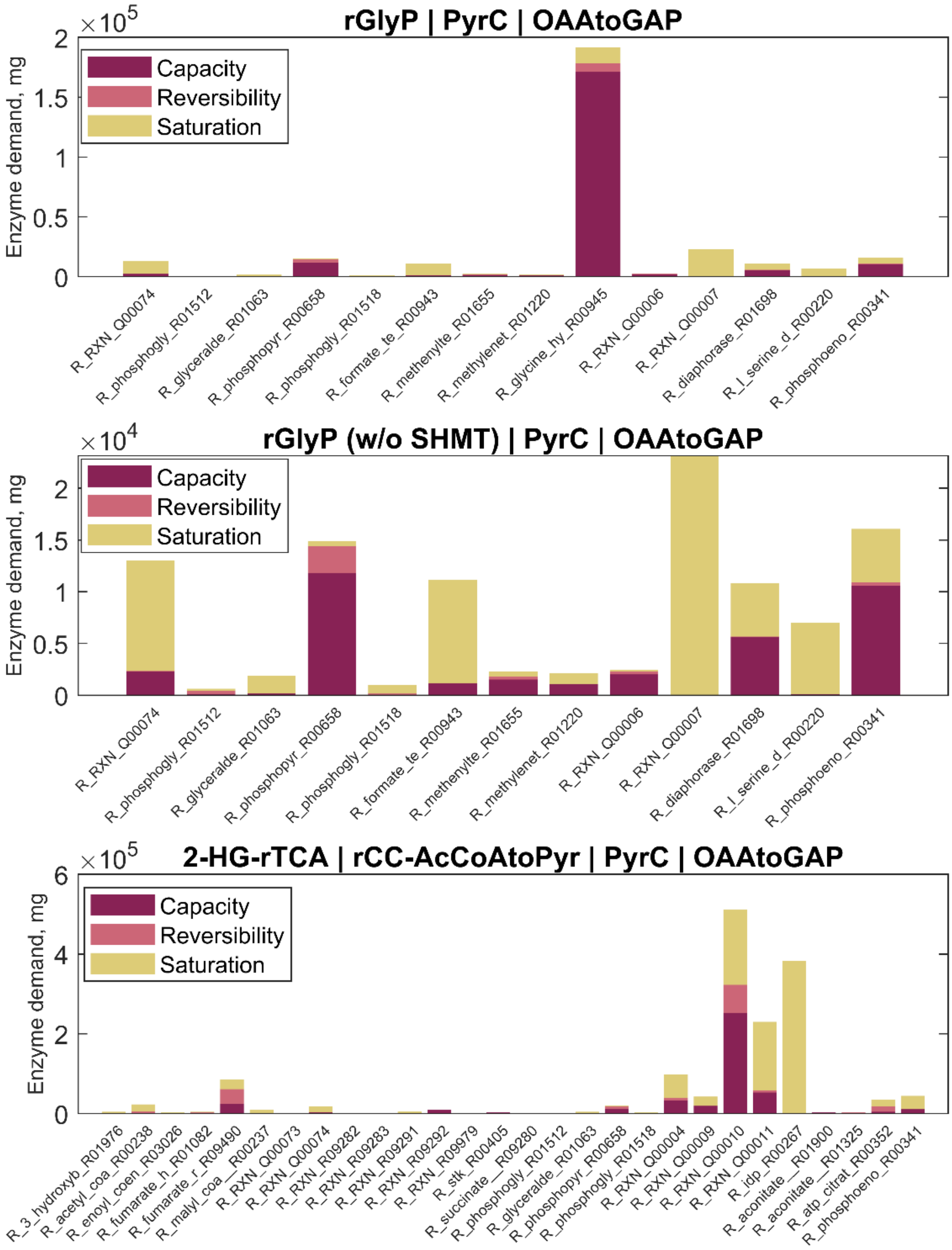


Figure S16: Enzyme demands to sustain a total pathway activity of 1 mmol product per second. Contribution of the capacity, the reversibility and the saturation with substrates or products of each reaction to the demand of enzymes. The figure follows the wording of (Noor et al., 2016). “Capacity”: demand of enzyme caused by a limitation by the catalytic rate constant. “Reversibility”: extra amount of enzyme needed because of a backward flux. “Saturation”: additional enzyme necessary because of undersaturation with a substrate or oversaturation with a product. The values present the optimized state as predicted by the ECM algorithm assuming a $CO_2$ concentration of 10 µM. Glyceraldehyde-3-phosphate was chosen as a product for the cycles in this case. Reaction names correspond to their identifiers in the SBtab model file (Supplementary file Reactions_Composite17_model.tsv)

**References (Supplement only)**


Abbe, K., & Yamada, T. (1982). Purification and properties of pyruvate kinase from Streptococcus mutans. *Journal of Bacteriology*, *149*(1), 299–305. https://doi.org/10.1128/jb.149.1.299-305.1982

Allen, S. H. . (1966). The isolation and characterization of malate-lactate transhydrogenase from *Micrococcus lactilyticus*. *The Journal of Biological Chemistry*, *241(22)*(Nov 25), 5266–5275.

Arfman, N., Bystrykh, L., Govorukhina, N. I., & Dijkhuizen, L. (1990). 3-Hexulose-6-phosphate synthase from thermotolerant methylotroph Bacillus C1. *Methods in Enzymology*, *188*(C), 391–397. https://doi.org/10.1016/0076-6879(90)88062-F

Baetz, A. L., & Allison, M. J. (1990). Purification and characterization of formyl-coenzyme A transferase from Oxalobacter formigenes. *Journal of Bacteriology*, *172*(7), 3537–3540. https://doi.org/10.1128/jb.172.7.3537-3540.1990

Bartsch, O., Hagemann, M., & Bauwe, H. (2008). Only plant-type (GLYK) glycerate kinases produce <scp>d</scp> -glycerate 3-phosphate. *FEBS Letters*, *582*(20), 3025–3028. https://doi.org/10.1016/j.febslet.2008.07.038

Baumgart, M., & Bott, M. (2011). Biochemical characterisation of aconitase from Corynebacterium glutamicum. *Journal of Biotechnology*, *154*(2–3), 163–170. https://doi.org/10.1016/j.jbiotec.2010.07.002

Belova, L. L., Sokolov, A. P., Sidorov, I. A., & Trotsenko, Y. A. (1997). Purification and characterization of NADPH-dependent acetoacetyl-CoA reductase from Methylobacterium extorquens. *FEMS Microbiology Letters*, *156*(2), 275–279. https://doi.org/10.1016/S0378-1097(97)00441-2

Berman, K. M., & Cohn, M. (1970). Phosphoenolpyruvate synthetase of *Escherichia coli*. Purification, some properties, and the role of divalent metal ions. *Journal of Biological Chemistry*, *245(20)*(Oct 25), 5309–5318.

Bernhardsgrütter, I., Vögeli, B., Wagner, T., Peter, D. M., Cortina, N. S., Kahnt, J., Bange, G., Engilberge, S., Girard, E., Riobé, F., Maury, O., Shima, S., Zarzycki, J., & Erb, T. J. (2018). The multicatalytic compartment of propionyl-CoA synthase sequesters a toxic metabolite. *Nature Chemical Biology*, *14*(12), 1127–1132. https://doi.org/10.1038/s41589-018-0153-x

Binstock, J. F., & Schulz, H. (1981). Fatty Acid Oxidation Complex from Escherichia coli. *Methods in Enzymology*, *71*(C), 403–411. https://doi.org/10.1016/0076-6879(81)71051-6

Borjian, F., Johnsen, U., Schönheit, P., & Berg, I. A. (2017). Succinyl-CoA: Mesaconate CoA-transferase and mesaconyl-CoA hydratase, enzymes of the methylaspartate cycle in Haloarcula hispanica. *Frontiers in Microbiology*, *8*(SEP). https://doi.org/10.3389/fmicb.2017.01683

Cadet, F., & Meunier, J. C. (1988). pH and kinetic studies of chloroplast sedoheptulose-1,7-bisphosphatase from spinach (Spinacia oleracea). *The Biochemical Journal*, *253*(1), 249–254. https://doi.org/10.1042/bj2530249

Carmo-Silva, A. E., Keys, A. J., Andralojc, P. J., Powers, S. J., Arrabaça, M. C., & Parry, M. A. J. (2010). Rubisco activities, properties, and regulation in three different C4 grasses under drought. *Journal of Experimental Botany*, *61*(9), 2355–2366. https://doi.org/10.1093/jxb/erq071

Chang, K. S., Jeon, H., Seo, S., Lee, Y., & Jin, E. S. (2014). Improvement of the phosphoenolpyruvate carboxylase activity of Phaeodactylum tricornutum PEPCase 1 through protein engineering. *Enzyme and Microbial Technology*, *60*, 64–71. https://doi.org/10.1016/j.enzmictec.2014.04.007

Chen, Z. H., Walker, R. P., Acheson, R. M., & Leegood, R. C. (2002). Phosphoenolpyruvate carboxykinase assayed at physiological concentrations of metal ions has a high affinity for CO2. *Plant Physiology*, *128*(1), 160–164. https://doi.org/10.1104/pp.010431

Chistoserdova, L. V., & Lidstrom, M. E. (1991). Purification and characterization of hydroxypyruvate reductase from the facultative methylotroph Methylobacterium extorquens AM1. *Journal of Bacteriology*, *173*(22), 7228–7232. https://doi.org/10.1128/jb.173.22.7228-7232.1991

Chou, A., Clomburg, J. M., Qian, S., & Gonzalez, R. (2019). 2-Hydroxyacyl-CoA lyase catalyzes acyloin condensation for one-carbon bioconversion. *Nature Chemical Biology*, *15*(9), 900–906. https://doi.org/10.1038/s41589-019-0328-0

Dayem, L. C., Carney, J. R., Santi, D. V., Pfeifer, B. A., Khosla, C., & Kealey, J. T. (2002). Metabolic engineering of a methylmalonyl-CoA mutase-epimerase pathway for complex polyketide biosynthesis in Escherichia coli. *Biochemistry*, *41*(16), 5193–5201. https://doi.org/10.1021/bi015593k

Dev, I. K., & Harvey, R. J. (1978). A complex of N5,N10-methylentetrahydrofolate dehydrogenase and N5,N10-methenyltetrahydrofolate cyclohydrolase in Escherichia coli. Purification subunit structure, and allosteric inhibition by N10-formyltetrahydrofolate. *Journal of Biological Chemistry*, *253*(12), 4245–4253. https://doi.org/10.1016/S0021-9258(17)34711-7

Erales, J., Gontero, B., & Maberly, S. C. (2008). SPECIFICITY AND FUNCTION OF GLYCERALDEHYDE-3-PHOSPHATE DEHYDROGENASE IN A FRESHWATER DIATOM, *ASTERIONELLA FORMOSA* (BACILLARIOPHYCEAE) [1]. *Journal of Phycology*, *44*(6), 1455–1464. https://doi.org/10.1111/j.1529-8817.2008.00600.x

Erb, T. J., Frerichs-Revermann, L., Fuchs, G., & Alber, B. E. (2010). The apparent malate synthase activity of Rhodobacter sphaeroides is due to two paralogous enzymes, (3S)-malyl-coenzyme A (CoA)/β-methylmalyl-CoA lyase and (3S)-malyl-CoA thioesterase. *Journal of Bacteriology*, *192*(5), 1249–1258. https://doi.org/10.1128/JB.01267-09

Erb, T. J., Rétey, J., Fuchs, G., & Alber, B. E. (2008). Ethylmalonyl-CoA mutase from Rhodobacter sphaeroides defines a new subclade of coenzyme B12-dependent Acyl-CoA mutases. *Journal of Biological Chemistry*, *283*(47), 32283–32293. https://doi.org/10.1074/jbc.M805527200

Ferenci, T., Strom, T., & Quayle, J. R. (1974). Purification and properties of 3 hexulose phosphate synthase and phospho 3 hexuloisomerase from Methylococcus capsulatus. *Biochemical Journal*, *144*(3), 477–486. https://doi.org/10.1042/bj1440477

Friedmann, S., Steindorf, A., Alber, B. E., & Fuchs, G. (2006). Properties of succinyl-coenzyme A:L-malate coenzyme A transferase and its role in the autotrophic 3-hydroxypropionate cycle of Chloroflexus aurantiacus. *Journal of Bacteriology*, *188*(7), 2646–2655. https://doi.org/10.1128/JB.188.7.2646-2655.2006

Fujiwara, K., & Motokawa, Y. (1983). Mechanism of the glycine cleavage reaction. Steady state kinetic studies of the P-protein-catalyzed reaction. *The Journal of Biological Chemistry*, *258(13)*(July 10), 8156–8162.

Gannon, F., Bridgeland, E. S., & Jones, K. M. (1977). L Serine dehydratase from Arthrobacter globiformis. *Biochemical Journal*, *161*(2), 345–355. https://doi.org/10.1042/bj1610345

GRAÑA, X., UREÑA, J., LUDEVID, D., CARRERAS, J., & CLIMENT, F. (1989). Purification, characterization and immunological properties of 2,3-bisphosphoglycerate-independent phosphoglycerate mutase from maize (Zea mays) seeds. *European Journal of Biochemistry*, *186*(1–2), 149–153. https://doi.org/10.1111/j.1432-1033.1989.tb15189.x

Gurr, J. A., & Jones, K. M. (1977). Purification and characterization of pyruvate carboxylase from Arthrobacter globiformis. *Archives of Biochemistry and Biophysics*, *179*(2), 444–455.

https://doi.org/10.1016/0003-9861(77)90132-1

Häusler, R. E., Holtum, J. A. M., & Latzko, E. (1987). CO2 is the inorganic carbon substrate of NADP malic enzymes from *Zea mays* and from wheat germ. *European Journal of Biochemistry*, *163*(3), 619–626. https://doi.org/10.1111/j.1432-1033.1987.tb10911.x

Hawkins, A. B., Adams, M. W. W., & Kelly, R. M. (2014). Conversion of 4-hydroxybutyrate to acetyl coenzyme A and its anapleurosis in the Metallosphaera sedula 3-hydroxypropionate/4-hydroxybutyrate carbon fixation pathway. *Applied and Environmental Microbiology*, *80*(8), 2536–2545. https://doi.org/10.1128/AEM.04146-13

Hederstedt, L., & Heden, L. O. (1989). New properties of Bacillus subtilis succinate dehydrogenase altered at the active site. The apparent active site thiol of succinate oxidoreductases is dispensable for succinate oxidation. *Biochemical Journal*, *260*(2), 491–497. https://doi.org/10.1042/bj2600491

Hedl, M., Sutherlin, A., Imogen Wilding, E., Mazzulla, M., McDevitt, D., Lane, P., Burgner, J. W., Lehnbeuter, K. R., Stauffacher, C. V., Gwynn, M. N., & Rodwell, V. W. (2002). Enterococcus faecalis acetoacetyl-coenzyme A thiolase/3-hydroxy-3-methylglutaryl-coenzyme A reductase, a dual-function protein of isopentenyl diphosphate biosynthesis. *Journal of Bacteriology*, *184*(8), 2116–2122. https://doi.org/10.1128/JB.184.8.2116-2122.2002

Houston, B., & Nimmo, H. G. (1984). Purification and some kinetic properties of rat liver ATP citrate lyase. *Biochemical Journal*, *224*(2), 437–443. https://doi.org/10.1042/bj2240437

Hügler, M., Menendez, C., Schägger, H., & Fuchs, G. (2002). Malonyl-coenzyme a reductase from Chloroflexus aurantiacus, a key enzyme of the 3-hydroxypropionate cycle for autotrophic CO2 fixation. *Journal of Bacteriology*, *184*(9), 2404–2410. https://doi.org/10.1128/JB.184.9.2404-2410.2002

Husic, H. D., & Tolbert, N. E. (1984). Anion and divalent cation activation of phosphoglycolate phosphatase from leaves. *Archives of Biochemistry and Biophysics*, *229*(1), 64–72. https://doi.org/10.1016/0003-9861(84)90130-9

IZUMI, Y., YOSHIDA, T., & YAMADA, H. (1990). Purification and characterization of serine-glyoxylate aminotransferase from a serine-producing methylotroph, Hyphomicrobium methylovorum GM2. *European Journal of Biochemistry*, *190*(2), 285–290. https://doi.org/10.1111/j.1432-1033.1990.tb15574.x

Kanao, T., Kawamura, M., Fukui, T., Atomi, H., & Imanaka, T. (2002). Characterization of isocitrate dehydrogenase from the green sulfur bacterium *Chlorobium limicola*. *European Journal of Biochemistry*, *269*(7), 1926–1931. https://doi.org/10.1046/j.1432-1033.2002.02849.x

Karsten, W. E., Cook, P. F., Ohshiro, T., & Izumi, Y. (2001). Initial velocity, spectral, and pH studies of the serine-glyoxylate aminotransferase from Hyphomicrobiuim methylovorum. *Archives of Biochemistry and Biophysics*, *388*(2), 267–275. https://doi.org/10.1006/abbi.2001.2294

Kelley-Loughnane, N., Biolsi, S. A., Gibson, K. M., Lu, G., Hehir, M. J., Phelan, P., & Kantrowitz, E. R. (2002). Purification, kinetic studies, and homology model of Escherichia coli fructose-1,6-bisphosphatase. *Biochimica et Biophysica Acta - Protein Structure and Molecular Enzymology*, *1594*(1), 6–16. https://doi.org/10.1016/S0167-4838(01)00261-8

Kimura, Y., Kojyo, T., Kimura, I., & Sato, M. (1998). Propionyl-coA carboxylase of Myxococcus xanthus: Catalytic properties and function in developing cells. *Archives of Microbiology*, *170*(3), 179–184. https://doi.org/10.1007/s002030050631

Könneke, M., Schubert, D. M., Brown, P. C., Hügler, M., Standfest, S., Schwander, T., Schada Von Borzyskowski, L., Erb, T. J., Stahl, D. A., & Berg, I. A. (2014). Ammonia-oxidizing archaea use the most energy-efficient aerobic pathway for CO2 fixation. *Proceedings of the National Academy of Sciences of the United States of America*, *111*(22), 8239–8244. https://doi.org/10.1073/pnas.1402028111

Kotlarz, D., & Buc, H. (1982). Phosphofructokinases from Escherichia coli. *Methods in Enzymology*, *90*(C), 60–70. https://doi.org/10.1016/S0076-6879(82)90107-0

Le, S. B., Heggeset, T. M. B., Haugen, T., Nærdal, I., & Brautaset, T. (2017). 6-Phosphofructokinase and ribulose-5-phosphate 3-epimerase in methylotrophic Bacillus methanolicus ribulose monophosphate cycle. *Applied Microbiology and Biotechnology*, *101*(10), 4185–4200. https://doi.org/10.1007/s00253-017-8173-0

MacKintosh, C., & Nimmo, H. G. (1988). Purification and regulatory properties of isocitrate lyase from Escherichia coli ML308. *The Biochemical Journal*, *250*(1), 25–31. https://doi.org/10.1042/bj2500025

Maklashina, E., & Cecchini, G. (1999). Comparison of catalytic activity and inhibitors of quinone reactions of succinate dehydrogenase (succinate-ubiquinone oxidoreductase) and fumarate reductase (menaquinol-fumarate oxidoreductase) from Escherichia coli. *Archives of Biochemistry and Biophysics*, *369*(2), 223–232. https://doi.org/10.1006/abbi.1999.1359

Marx, C. J., Laukel, M., Vorholt, J. A., & Lidstrom, M. E. (2003). Purification of the Formate-Tetrahydrofolate Ligase from Methylobacterium extorquens AM1 and Demonstration of Its Requirement for Methylotrophic Growth. *Journal of Bacteriology*, *185*(24), 7169–7175. https://doi.org/10.1128/JB.185.24.7169-7175.2003

Mathur, D., Malik, G., & Garg, L. C. (2006). Biochemical and functional characterization of triosephosphate isomerase from Mycobacterium tuberculosis H37Rv. *FEMS Microbiology Letters*, *263*(2), 229–235. https://doi.org/10.1111/j.1574-6968.2006.00420.x

Meister, M., Saum, S., Alber, B. E., & Fuchs, G. (2005). L-malyl-coenzyme A/β-methylmalyl-coenzyme A lyase is involved in acetate assimilation of the isocitrate lyase-negative bacterium Rhodobacter capsulatus. *Journal of Bacteriology*, *187*(4), 1415–1425. https://doi.org/10.1128/JB.187.4.1415-1425.2005

Meyer, M., Schweiger, P., & Deppenmeier, U. (2015). Succinic semialdehyde reductase Gox1801 from Gluconobacter oxydans in comparison to other succinic semialdehyde-reducing enzymes. *Applied Microbiology and Biotechnology*, *99*(9), 3929–3939. https://doi.org/10.1007/s00253-014-6191-8

MIYAZAKI, S. S., TOKI, S., IZUMI, Y., & YAMADA, H. (1987). Purification and characterization of a serine hydroxymethyltransferase from an obligate methylotroph, Hyphomicrobium methylovorum GM2. *European Journal of Biochemistry*, *162*(3), 533–540. https://doi.org/10.1111/j.1432-1033.1987.tb10672.x

Moskowitz, G. J., & Merrick, J. M. (1969). Metabolism of Poly-β-hydroxybutyrate. II. Enzymatic Synthesis of D-(-)-β-Hydroxybutyryl Coenzyme A by an Enoyl Hydrase from Rhodospirillum rubrum. *Biochemistry*, *8*(7), 2748–2755. https://doi.org/10.1021/bi00835a009

Muslin, E. H., Li, D., Stevens, F. J., Donnelly, M., Schiffer, M., & Anderson, L. E. (1995). Engineering a domain-locking disulfide into a bacterial malate dehydrogenase produces a redox-sensitive enzyme. *Biophysical Journal*, *68*(6), 2218–2223. https://doi.org/10.1016/S0006-3495(95)80430-3

Nakahara, K., Yamamoto, H., Miyake, C., & Yokota, A. (2003). Purification and characterization of class-I and class-II fructose-1,6-bisphosphate aldolases from the cyanobacterium Synechocystis sp. PCC6803. *Plant and Cell Physiology*, *44*(3), 326–333. https://doi.org/10.1093/pcp/pcg044

Nakano, T., Ashida, H., Mizohata, E., Matsumura, H., & Yokota, A. (2010). An evolutionally conserved Lys122 is essential for function in Rhodospirillum rubrum bona fide RuBisCO and Bacillus subtilis RuBisCO-like protein. *Biochemical and Biophysical Research Communications*, *392*(2), 212–216. https://doi.org/10.1016/j.bbrc.2010.01.017

NARINDRASORASAK, S., & BRIDGER, W. A. (1977). Phosphoenolpyruvate Synthetase of *Escherichia coli*: molecular weight, subunit composition, and identification of phosphohistidine in phosphoenzyme intermediate. *Journal of Biological Chemistry*, *252(10)*(May 25), 3121–3127.

Neuburger, M., Polidori, A. M., Piètre, E., Faure, M., Jourdain, A., Bourguignon, J., Pucci, B., & Douce, R. (2000). Interaction between the lipoamide-containing H-protein and the lipoamide dehydrogenase (L-protein) of the glycine decarboxylase multienzyme system. *European Journal of Biochemistry*, *267*(10), 2882–2889. https://doi.org/10.1046/j.1432-1327.2000.01301.x

Nolte, J. C., Schürmann, M., Schepers, C. L., Vogel, E., Wübbeler, J. H., & Steinbüchel, A. (2014). Novel characteristics of succinate coenzyme a (succinate-coa) ligases: Conversion of malate to malyl-coa and coa-thioester formation of succinate analogues in vitro. *Applied and Environmental Microbiology*, *80*(1), 166–176. https://doi.org/10.1128/AEM.03075-13

Okamura-Ikeda, K., Kameoka, N., Fujiwara, K., & Motokawa, Y. (2003). Probing the H-protein-induced conformational change and the function of the N-terminal region of Escherichia coli T-protein of the glycine cleavage system by limited proteolysis. *Journal of Biological Chemistry*, *278*(12), 10067–10072. https://doi.org/10.1074/jbc.M210853200

Padovani, D., & Banerjee, R. (2006). Assembly and protection of the radical enzyme, methylmalonyl-CoA mutase, by its chaperone. *Biochemistry*, *45*(30), 9300–9306. https://doi.org/10.1021/bi0604532

Pennati, A., & Gadda, G. (2009). Involvement of ionizable groups in catalysis of human liver glycolate oxidase. *Journal of Biological Chemistry*, *284*(45), 31214–31222. https://doi.org/10.1074/jbc.M109.040063

Pomper, B. K., Vorholt, J. A., Chistoserdova, L., Lidstrom, M. E., & Thauer, R. K. (1999). A methenyl tetrahydromethanopterin cyclohydrolase and a methenyl tetrahydrofolate cyclohydrolase in Methylobacterium extorquens AM1. *European Journal of Biochemistry*, *261*(2), 475–480. https://doi.org/10.1046/j.1432-1327.1999.00291.x

Price, L. J., Herbert, D., Moss, S. R., Cole, D. J., & Harwood, J. L. (2003). Graminicide insensitivity correlates with herbicide-binding co-operativity on acetyl-CoA carboxylase isoforms. *Biochemical Journal*, *375*(2), 415–423. https://doi.org/10.1042/BJ20030665

Ranganathan, N. S., Srere, P. A., & Linn, T. C. (1980). Comparison of phospho- and dephospho-ATP citrate lyase. *Archives of Biochemistry and Biophysics*, *204*(1), 52–58. https://doi.org/10.1016/0003-9861(80)90006-5

Reshetnikov, A. S., Rozova, O. N., Khmelenina, V. N., Mustakhimov, I. I., Beschastny, A. P., Murrell, J. C., & Trotsenko, Y. A. (2008). Characterization of the pyrophosphate-dependent 6-phosphofructokinase from Methylococcus capsulatus Bath. *FEMS Microbiology Letters*, *288*(2), 202–210. https://doi.org/10.1111/j.1574-6968.2008.01366.x

Sasikaran, J., Ziemski, M., Zadora, P. K., Fleig, A., & Berg, I. A. (2014). Bacterial itaconate degradation promotes pathogenicity. *Nature Chemical Biology*, *10*(5), 371–377. https://doi.org/10.1038/nchembio.1482

SAUMWEBER, H., BINDER, R., & BISSWANGER, H. (1981). Pyruvate Dehydrogenase Component of the Pyruvate Dehydrogenase Complex from Escherichia coli K12. Purification and Characterization. *European Journal of Biochemistry*, *114*(2), 407–411. https://doi.org/10.1111/j.1432-

1033.1981.tb05161.x

Schada von Borzyskowski, L., Severi, F., Krüger, K., Hermann, L., Gilardet, A., Sippel, F., Pommerenke, B., Claus, P., Cortina, N. S., Glatter, T., Zauner, S., Zarzycki, J., Fuchs, B. M., Bremer, E., Maier, U. G., Amann, R. I., & Erb, T. J. (2019). Marine Proteobacteria metabolize glycolate via the β-hydroxyaspartate cycle. *Nature*, *575*(7783), 500–504. https://doi.org/10.1038/s41586-019-1748-4

Schirch, V., Hopkins, S., Villar, E., & Angelaccio, S. (1985). Serine hydroxymethyltransferase from Escherichia coli: Purification and properties. *Journal of Bacteriology*, *163*(1), 1–7. https://doi.org/10.1128/jb.163.1.1-7.1985

Schnorpfeil, M., Janausch, I. G., Biel, S., Kröger, A., & Unden, G. (2001). Generation of a proton potential by succinate dehydrogenase of *Bacillus subtilis* functioning as a fumarate reductase. *European Journal of Biochemistry*, *268*(10), 3069–3074. https://doi.org/10.1046/j.1432-1327.2001.02202.x

Schwander, T., McLean, R., Zarzycki, J., & Erb, T. J. (2018). Structural basis for substrate specificity of methylsuccinyl-CoA dehydrogenase, an unusual member of the acyl-CoA dehydrogenase family. *Journal of Biological Chemistry*, *293*(5), 1702–1712. https://doi.org/10.1074/jbc.RA117.000764

Shimakata, T., Fujita, Y., & Kusaka, T. (1979). Purification and characterization of 3-hydroxyacyl-coa dehydrogenase of Mycobacterium smegmatis. *Journal of Biochemistry*, *86*(5), 1191–1198. https://doi.org/10.1093/oxfordjournals.jbchem.a132634

Sprenger, G. A., Schorken, U., Sprenger, G., & Sahm, H. (1995). Transketolase a of Escherichia coli K12. Purification and Properties of the Enzyme from Recombinant Strains. *European Journal of Biochemistry*, *230*(2), 525–532. https://doi.org/10.1111/j.1432-1033.1995.0525h.x

Stoffel, G. M. M., Saez, D. A., DeMirci, H., Vögeli, B., Rao, Y., Zarzycki, J., Yoshikuni, Y., Wakatsuki, S., Vöhringer-Martinez, E., & Erb, T. J. (2019). Four amino acids define the CO2 binding pocket of enoyl-CoA carboxylases/reductases. *Proceedings of the National Academy of Sciences of the United States of America*, *116*(28), 13964–13969. https://doi.org/10.1073/pnas.1901471116

Tsukamoto, Y., Fukushima, Y., Hara, S., & Hisabori, T. (2013). Redox control of the activity of phosphoglycerate kinase in synechocystis sp. PCC6803. *Plant and Cell Physiology*, *54*(4), 484–491. https://doi.org/10.1093/pcp/pct002

Ueda, Y., Yumoto, N., Tokushige, M., Fukui, K., & Ohya-nishiguchi, H. (1991). Purification and characterization of two types of fumarase from Escherichia coli. *Journal of Biochemistry*, *109*(5), 728–733. https://doi.org/10.1093/oxfordjournals.jbchem.a123448

Wadano, A., Nishikawa, K., Hirahashi, T., Satoh, R., & Iwaki, T. (1998). Reaction mechanism of phosphoribulokinase from a cyanobacterium, Synechococcus PCC7942. *Photosynthesis Research*, *56*(1), 27–33. https://doi.org/10.1023/A:1005979801741

Wang, Q., Ou, M. S., Kim, Y., Ingram, L. O., & Shanmugam, K. T. (2010). Metabolic flux control at the pyruvate node in an anaerobic escherichia coli strain with an active pyruvate dehydrogenase. *Applied and Environmental Microbiology*, *76*(7), 2107–2114. https://doi.org/10.1128/AEM.02545-09

Wood, H. G., Jacobson, B., Gerwin, B. I., & Northrop, D. B. (1969). [36] Oxaloacetate transcarboxylase from Propionibacterium. *Methods in Enzymology*, *13*(C), 215–230. https://doi.org/10.1016/0076-6879(69)13041-4

Wu, T. Y., Chen, C. T., Liu, J. T. J., Bogorad, I. W., Damoiseaux, R., & Liao, J. C. (2016). Characterization and evolution of an activator-independent methanol dehydrogenase from Cupriavidus necator N-1. *Applied Microbiology and Biotechnology*, *100*(11), 4969–4983.

https://doi.org/10.1007/s00253-016-7320-3

Yoshida, Y., Sato, M., Kezuka, Y., Hasegawa, Y., Nagano, K., Takebe, J., & Yoshimura, F. (2016). Acyl-CoA reductase PGN-0723 utilizes succinyl-CoA to generate succinate semialdehyde in a butyrate-producing pathway of Porphyromonas gingivalis. *Archives of Biochemistry and Biophysics*, *596*, 138–148. https://doi.org/10.1016/j.abb.2016.03.014

Zadvornyy, O. A., Boyd, E. S., Posewitz, M. C., Zorin, N. A., & Peters, J. W. (2015). Biochemical and structural characterization of enolase from Chloroflexus aurantiacus: Evidence for a thermophilic origin. *Frontiers in Bioengineering and Biotechnology*, *3*(JUN), 74. https://doi.org/10.3389/fbioe.2015.00074

Zarzycki, J., Brecht, V., Müller, M., & Fuchs, G. (2009). Identifying the missing steps of the autotrophic 3-hydroxypropionate CO2 fixation cycle in Chloroflexus aurantiacus. *Proceedings of the National Academy of Sciences of the United States of America*, *106*(50), 21317–21322. https://doi.org/10.1073/pnas.0908356106

Zhang, R. G., Andersson, C. E., Savchenko, A., Skarina, T., Evdokimova, E., Beasley, S., Arrowsmith, C. H., Edwards, A. M., Joachimiak, A., & Mowbray, S. L. (2003). Structure of Escherichia coli ribose-5-phosphate isomerase: A ubiquitous enzyme of the pentose phosphate pathway and the Calvin cycle. *Structure*, *11*(1), 31–42. https://doi.org/10.1016/S0969-2126(02)00933-4

Ziegler, I. (1974). Malate dehydrogenase in zea may: Properties and inhibition by sulfite. *BBA - Enzymology*, *364*(1), 28–37. https://doi.org/10.1016/0005-2744(74)90129-6
