## Supplementary material for "In-depth computational analysis of natural and artificial carbon fixation pathways": MATLAB files: AdjustKineticDataKeq.m

```
function kinetic_data_out = AdjustKineticDataKeq(kinetic_data_in, Keq, Keq0, kcat_adj_mode)
    for i=1:length(Keq0)
        if(abs(log(Keq(i))-log(Keq0(i)))> 0.1)
            % conservative case: only lower reaction constants:
            pf = Keq0(i)/Keq(i);
            if(kcat_adj_mode == 1 || isnan(kinetic_data_in.Kcatf.mean(i)) || isnan(kinetic_data_in.Kcatr.mean(i))) % if one reaction rate is given, change the rates one-sided like in the conservative case
                if(Keq(i) > Keq(i))
                    if(~isnan(kinetik_data_in.Kcatf.mean(i)))
                        kinetic_data_in.Kcatf.mean(i) = kinetic_data_in.Kcatf.mean(i)*pf;
                        kinetic_data_in.Kcatf.std(i) = kinetic_data_in.Kcatf.std(i)*pf;
                    else
                        kinetic_data_in.Kcatr.mean(i) = kinetic_data_in.Kcatr.mean(i)/pf;
                        kinetic_data_in.Kcatr.std(i) = kinetic_data_in.Kcatr.std(i)/pf;
                    end
                else
                    if(~isnan(kinetic_data_in.Kcatr.mean(i)))
                        kinetic_data_in.Kcatr.mean(i) = kinetic_data_in.Kcatr.mean(i)/pf;
                        kinetic_data_in.Kcatr.std(i) = kinetic_data_in.Kcatr.std(i)/pf;
                    else
                        kinetic_data_in.Kcatf.mean(i) = kinetic_data_in.Kcatf.mean(i)*pf;
                        kinetic_data_in.Kcatf.std(i) = kinetic_data_in.Kcatf.std(i)*pf;
                    end
                end
            else 
                pf = sqrt(pf);
                kinetic_data_in.Kcatf.mean(i) = kinetic_data_in.Kcatf.mean(i)*pf;
                kinetic_data_in.Kcatf.std(i) = kinetic_data_in.Kcatf.std(i)*pf;
                kinetic_data_in.Kcatr.mean(i) = kinetic_data_in.Kcatr.mean(i)/pf;
                kinetic_data_in.Kcatf.std(i) = kinetic_data_in.Kcatf.std(i)/pf;
            end
        end
        kinetic_data_in.Kcatf.mean_ln(i) = lognormal_normal2log(kinetic_data_in.Kcatf.mean(i),kinetic_data_in.Kcatf.std(i));
        kinetic_data_in.Kcatf.median(i) = exp(kinetic_data_in.Kcatf.mean_ln(i));
        kinetic_data_in.Kcatr.mean_ln(i) = lognormal_normal2log(kinetic_data_in.Kcatr.mean(i),kinetic_data_in.Kcatr.std(i));
        kinetic_data_in.Kcatr.median(i) = exp(kinetic_data_in.Kcatr.mean_ln(i));
    end
    kinetic_data_out = kinetic_data_in;
end
```
