## Supplementary material for "In-depth computational analysis of natural and artificial carbon fixation pathways": MATLAB files: CalcProjATPcosts.m

```
function pathway_cost = CalcProjATPcosts(reaction_names, fluxes, network)
    vv = zeros(length(network.reaction_names),1);

    for i = 1:length(reaction_names)
        index = find(strcmp(network.reaction_names, reaction_names{i}));
        vv(index(1)) = fluxes(i);
    end
    flux = network.N*vv;
    o_str = TotalStoichiometryAsString(network, reaction_names, fluxes);
    fprintf('Total stoichiometry: %s\n', o_str);

    % Calculate ATP costs of pathways:
    i_AMP = find(strcmp(network.metabolites,'C_amp'));
    i_ADP = find(strcmp(network.metabolites,'C_adp'));
    i_Mena = find(strcmp(network.metabolites,'C_menaquinol'));
    i_Ubi = find(strcmp(network.metabolites,'C_ubiquinol'));
    pathway_cost = flux(i_AMP)*2 + flux(i_ADP) + flux(i_Mena) + flux(i_Ubi);
end
```
