## Supplementary material for "In-depth computational analysis of natural and artificial carbon fixation pathways": MATLAB files: calculate_rev_sat_HL.m

```
function [u_cap,eta_sat,eta_rev] = calculate_rev_sat_HL(kinetic_law, N, W, ind_ext, u, c, KA, KI, KM, KV, Keq, h, Mplus, Mminus, Wplus, Wminus, nm, nr)

% [v, v_plus, v_minus] = modular_velocities(kinetic_law,N,W,ind_ext,u,c,KA,KI,KM,KV,Keq,h,Mplus, Mminus, Wplus, Wminus, nm, nr)

c(find(c<10^-14)) = 10^-14;

%% Global variables to speed up this function; see ecm_one_run.m and convex_parameter_estimation.m
global global_structure_matrices 

if exist('global_structure_matrices','var'),
  global Mplus Mminus Wplus Wminus nm nr ind_M ind_Wp ind_Wm
end

if isempty(Mplus),
  [Mplus, Mminus, Wplus, Wminus, nm, nr, N_int,ind_M,ind_Wp,ind_Wm] = make_structure_matrices(N,W,ind_ext,h);
end

[log_alpha_A, log_beta_A] = k_to_log_alpha(KA,c);
[log_alpha_I            ] = k_to_log_alpha(KI,c);
[Kplus,Kminus]            = ms_compute_Kcat(N,KM,KV,Keq);

log_c_by_k        = repmat(log(c)',nr,1);
log_c_by_k(ind_M) = log_c_by_k(ind_M) - log(KM(ind_M)+10^-15);

theta_plus  = exp( sum( Mplus  .* log_c_by_k, 2) );
theta_minus = exp( sum( Mminus .* log_c_by_k, 2) );

switch kinetic_law,  
  case {'cs','ms'},
    psi_plus    = exp( sum( Mplus  .* log(1+exp(log_c_by_k)) , 2) );
    psi_minus   = exp( sum( Mminus .* log(1+exp(log_c_by_k)) , 2) );    
end

% complete allosteric regulation
regulation_term = exp( sum( Wplus .* log_beta_A + Wminus .* log_alpha_I, 2) );

switch kinetic_law,  
  case 'cs', D = psi_plus + psi_minus - 1;
  case 'ms', D = psi_plus .* psi_minus;
  case 'ds', D = theta_plus + theta_minus + 1;
  case 'rp', D = 1;
  case 'fd', D = sqrt(theta_plus .* theta_minus);
end

v_plus  = u .* regulation_term .* Kplus  .* theta_plus  ./ D;
v_minus = u .* regulation_term .* Kminus .* theta_minus ./ D;

v_plus  = real(full(v_plus));
v_minus = real(full(v_minus));
v       = v_plus - v_minus;

eta_rev = zeros(length(v),1);
eta_sat = zeros(length(v),1);
u_cap = zeros(length(v),1);

eta_rev(v>0) = 1 - (Kminus(v>0) .* theta_minus(v>0)./Kplus(v>0)./theta_plus(v>0));
eta_sat(v>0) = theta_plus(v>0)./D(v>0);
u_cap(v>0) = v(v>0)./Kplus(v>0);

eta_rev(v<0) = 1 - (Kplus(v<0) .* theta_plus(v<0)./theta_minus(v<0))./Kminus(v<0);
eta_sat(v<0) = theta_minus(v<0)./D(v<0);
u_cap(v<0) = -v(v<0)./Kminus(v<0);
```
