## Supplementary material for "In-depth computational analysis of natural and artificial carbon fixation pathways": MATLAB files: CapRevSat_Plot.m

```
function CapRevSat_Plot(str_title,cycle_ids,Cap_Cell,Cap_Rev_Cell,Cap_Rev_Sat_Cell,SubNetReactNames_Cell,iCycles,xx,yy,alt_titles)

bf_colormap =   [51 34 136; % Calvin cycle
                136 204 238; % HP/HB cycle (Cren)
                68 170 153; % HP/HB cycle (Thaum)
                17 119 51; % HP bicycle
                153 153 51; % Serine cycle
                221 204 119; % RuMP cycle
                204 102 119; % rCC cycle
                136 34 85; % CETCH cycle
                170 68 153; % rGC pathw.
                221 221 221]/255; % 2-HA/rTCA
            
figure
if(~isempty(str_title))
    sgtitle(str_title);
end

cycle_names = cell(numel(cycle_ids),1);

for i_ECM = 1:numel(iCycles)
    cycle = cycle_ids{iCycles(i_ECM)};
    %cycle_names{i_ECM} = ['{\fontfamily{cmss}\selectfont \textbf{' cycle{1} '}'];
    cycle_names{i_ECM} = cycle{1};
    for i = 2:(numel(cycle)/2)
        cycle_names{i_ECM} = [cycle_names{i_ECM} ' | ' cycle{i*2-1}];
    end
    %cycle_names{i_ECM} = [cycle_names{i_ECM} '}'];
    M = [Cap_Cell{iCycles(i_ECM)}, Cap_Rev_Cell{iCycles(i_ECM)} - Cap_Cell{iCycles(i_ECM)}, Cap_Rev_Sat_Cell{iCycles(i_ECM)} - Cap_Rev_Cell{iCycles(i_ECM)}];
    subplot(yy, xx, i_ECM);
    h = bar(1:length(Cap_Cell{iCycles(i_ECM)}),M, 'stacked','EdgeColor','none','BarWidth',0.8);
    my_colormap = bf_colormap([8 7 6],1:3);%[1-((1-bf_colormap(5,1:3))); 1-((1-bf_colormap(5,1:3))*0.8); 1-((1-bf_colormap(5,1:3))*0.6)];%[0.35 0.35 0.9; 0.8 0.2 0.7; 1 0.3 0.2];
    for it = 1:length(h)
      h(it).FaceColor = 'Flat';
      h(it).CData = my_colormap(it,:);
    end
    ylabel('Enzyme demand, mg','FontSize',10);
    set(gca,'Xtick',1:length(Cap_Cell{iCycles(i_ECM)}),'xticklabel',SubNetReactNames_Cell{iCycles(i_ECM)},'TickLabelInterpreter','none');
    xtickangle(45);
    legend('Capacity','Reversibility','Saturation','Location','SouthWest');
    if(isempty(alt_titles))
        title(cycle_names{i_ECM});
    else
        title(alt_titles{i_ECM});
    end
    ax = gca;
    ax.XAxis.TickLength = [0 0];
    ax.XAxis.FontSize = 7;
end            
            

end
```
