## Supplementary material for "In-depth computational analysis of natural and artificial carbon fixation pathways": MATLAB files: CarbonFixationPBECMCompareFinal.m

```
%% CarbonFixationPBECMCompareFinal
% this script is split in two parts: the first part loads the input data
% and applies parameter balancing. In the second part, the ECM algorithm is
% used to make predictions about pathway specific activities

% define SBtab model and data file locations and names:
modelfile = 'reactions_Composite17_model.tsv';%modelfile = 'reactions_Composite_model.tsv';%'CETCH_SuCrEtCi_Comp3_model.tsv';
datafile = 'reactions_Composite17_data.tsv';%datafile = 'reactions_Composite_data.tsv';%'CETCH_SuCrEtCi_Comp3_data.tsv';

% ------------------------ Parameter Balancing begin ---------------------
% ------------------------------------------------------------------------
% set options for Parameter Balancing
pb_options = parameter_balancing_options;

pb_options.enforce_flux_directions    = 0;
pb_options.adjust_to_fluxes           = 0;
pb_options.preferred_data_element_ids = 'sbml';
% set this option to use the same settings as n the python version:
pb_options.use_python_version_defaults = 1;
pb_options.fix_Keq_in_sampling = 1; % note that reactions where substrate and product catalytic constant are known will still produce different Keq values.

parameter_prior = parameter_balancing_prior([],pb_options.parameter_prior_file,1); 
parameter_prior = pb_parameter_prior_adjust(parameter_prior, pb_options);
% Adjust parameter prior to support higher and lower Keq values:
ind_Keq = find(strcmp(parameter_prior.Symbol,'Keq'));
parameter_prior.UpperBound(ind_Keq) = {num2str(10^100)};
parameter_prior.LowerBound(ind_Keq) = {num2str(10^-12)};

% Load original kinetic data:
my_sbtab_model      = sbtab_document_load_from_one(modelfile);
options = struct;
warnings = '';
network  = sbtab_to_network(my_sbtab_model,options);
[model_quantities, basic_quantities, data_quantities, pseudo_quantities] = parameter_balancing_quantities(parameter_prior, network, pb_options);
kinetic_data = kinetic_data_load(data_quantities, parameter_prior, network, datafile, struct('use_sbml_ids', pb_options.use_sbml_ids, 'use_kegg_ids', pb_options.use_kegg_ids, 'use_python_version_defaults', pb_options.use_python_version_defaults));

% Get enzyme specific weight:
try
  parameter_mean  = sbtab_table_get_column(my_sbtab_model.tables.Parameter0,'Mean',1);
  q_type  = sbtab_table_get_column(my_sbtab_model.tables.Parameter0,'QuantityType');
  r_name = sbtab_table_get_column(my_sbtab_model.tables.Parameter0,'Reaction:SBML:reaction:id');
  ind_enzmass = find(strcmp(q_type,'protein molecular mass'));
  i_enz = label_names(r_name(ind_enzmass), network.reaction_names);
  network.protein_molecular_mass = 10000*ones(length(i_enz),1);
  network.protein_molecular_mass(i_enz) = parameter_mean(ind_enzmass);

  catch err
    warnings = [warnings, 'Enzyme masses could not be extracted from model file.'];
end

% Get concentration bounds:
try
  conc_min  = sbtab_table_get_column(my_sbtab_model.tables.ConcentrationConstraint,'Concentration:Min',1);
  conc_max  = sbtab_table_get_column(my_sbtab_model.tables.ConcentrationConstraint,'Concentration:Max',1);  
  compounds  = sbtab_table_get_column(my_sbtab_model.tables.ConcentrationConstraint,'Compound');
  i_met = label_names(compounds, network.metabolites);
  conc_min(i_met) = conc_min*1000; % concentrations are given in M (-> mM)
  conc_max(i_met) = conc_max*1000;
  catch err
    warnings = [warnings, 'Concentration bounds could not be extracted from model file.'];
end

% Get standard Gibbs energies and their standard deviations:
try
  IDs  = sbtab_table_get_column(my_sbtab_model.tables.deltaG,'ID',1);
  mean_G  = sbtab_table_get_column(my_sbtab_model.tables.deltaG,'Mean',1);  
  std_G  = sbtab_table_get_column(my_sbtab_model.tables.deltaG,'Std');
  ind_react = label_names(compounds, network.reaction_names);
  mean_G(ind_react) = mean_G;
  std_G(ind_react) = std_G;  
  catch err
    warnings = [warnings, 'Standard Gibbs energies could not be extracted from model file.'];
end

% initialize kinetics structure:
network.kinetics  = set_kinetics(network, 'cs');

[nr,nm,nx,ind_KM,ind_KA,ind_KI,nKM,nKA,nKI] = network_numbers(network);

kinetics = struct;
kinetics.type = 'cs';
kinetics.h = ones(nr,1);
kinetics.KM = ones(nr,nm)*NaN;
kinetics.KI = zeros(nr,nm);
kinetics.KA = zeros(nr,nm);
kinetics.mu = zeros(nr,nm);
kinetics.c = ones(nm,1)*0.1;
kinetics.u = ones(nr,1)*10^-3;
kinetics.dmu0 = ones(nr,1)*NaN;
kinetics.Kcatf = ones(nr,1)*NaN;
kinetics.Kcatr = ones(nr,1)*NaN;
kinetics.Keq = ones(nr,1)*NaN;
kinetics.A = ones(nr,1)*NaN;
kinetics.Kcatratio = ones(nr,1)*NaN;

kinetics_std = kinetics; % create a duplicate structure for the standard deviations

kinetics_orig = kinetics;
parameter_prior_orig = parameter_prior;
Keq_check = 0;
loop = 0;
% find the index of beta-methylmalonyl-CoA lyase that will we be used to
% correct the C5-dicarboxylic acid platform:
ind_malyl_CoA_lyase = find(strcmp(network.reaction_KEGGID,'R00934'));

loop_M = zeros(nr,1);

% Balance every reaction by itself to avoid numerical problems:
for i=1:nr
    % Create sub-network representing the single reaction i
    [subnetwork, ind_m, ind_r] = network_subnetwork_HL(network, 1:nm, i, 1);
    sub_kinetic_data = kinetics_subkinetics(kinetic_data, i, ind_m);
    sub_kinetic_data_orig = sub_kinetic_data;
    sub_kinetic_data = pb_kinetic_data_adjust(sub_kinetic_data, parameter_prior, subnetwork, pb_options);
    
    % re-iterate until the balanced parameters are consistent with the Keq
    % (maximal 50 iterations):
    while(Keq_check == 0 && loop < 50)
        if(i == ind_malyl_CoA_lyase || sub_kinetic_data.Keq.mean > 10^10 || sub_kinetic_data.Keq.mean < 10^-10)
            pb_options.fix_Keq_in_sampling = 0; % std-deviation of Keq of malyl_CoA_lyase is very high, the estimation from the kinetics is hence probably better
        else
            pb_options.fix_Keq_in_sampling = 1;
        end
        if(i == 94)
            fprintf('break\n');
        end
        task   = parameter_balancing_task(subnetwork, sub_kinetic_data, parameter_prior, model_quantities, basic_quantities, pseudo_quantities);
        result = parameter_balancing_calculation(task, parameter_prior, pb_options);
        % run the actual algorithm
        [r_mode,r_mean,r_std,r_geom_mean,r_geom_std,r_orig,r_samples] = parameter_balancing_output(result, sub_kinetic_data_orig, pb_options);
        
        if(abs(log(r_mode.Keq)-log(r_orig.Keq)) < 0.1 || i == ind_malyl_CoA_lyase || sub_kinetic_data.Keq.mean > 10^10 || sub_kinetic_data.Keq.mean < 10^-10) % std-deviation of Keq of malyl_CoA_lyase is very high, the estimation from the kinetics is hence probably better
            Keq_check = 1;
        else
            % increase the standard deviations of parameter priors and
            % input values if inconsistent Keq values occured:
            [sub_kinetic_data,parameter_prior] = increase_kinetics_std(sub_kinetic_data, parameter_prior, 1, 2, 1, 0);
        end
        loop = loop + 1;
    end
    loop_M(i) = loop;
    
    %store the balanced kinetic parameters in the kinetics structure:
    kinetics = kinetics_write(kinetics,r_geom_mean,i,ind_m);
    kinetics_std = kinetics_write(kinetics_std,r_geom_std,i,ind_m);
    kinetics_orig = kinetics_write(kinetics_orig,r_orig,i,ind_m);
    
    Keq_check = 0;
    loop = 0;
    parameter_prior = parameter_prior_orig;
end

% Adjust reaction in- and effluxes from C5-bodies (to cope with uncertainty
% of the respective reactions: (using the estimated Keq from R00934)
ind_citramalyl_CoA_lyase = find(strcmp(network.reaction_KEGGID,'R00237'));
ind_ccr = find(strcmp(network.reaction_KEGGID,'R09291'));
corr_factor = kinetics.Keq(ind_malyl_CoA_lyase)/kinetic_data.Keq.mean(ind_malyl_CoA_lyase);
kinetics.Keq(ind_citramalyl_CoA_lyase) = kinetics.Keq(ind_citramalyl_CoA_lyase)*corr_factor;
kinetics.Kcatr(ind_citramalyl_CoA_lyase) = kinetics.Kcatr(ind_citramalyl_CoA_lyase)/corr_factor; % kcatr is unknown while kcatf is, so it is fair to reduce the former to adjust the Keq
kinetics.Keq(ind_ccr) = kinetics.Keq(ind_ccr)*corr_factor; % make the reaction a little less irreversible (still will be practically irreversible)
kinetics.Kcatf(ind_ccr) = kinetics.Kcatf(ind_ccr)*corr_factor; % kcatf is unknown while kcatr is, so it is fair to reduce the former to adjust the Keq

%Keq_recalc = recalculate_Keq(network, kinetics);
ind_Keq_err = abs(log(kinetics.Keq)-log(kinetics_orig.Keq)) > 0.1

% calculate geometric mean of catalytic constants:
kinetics.KV = sqrt(kinetics.Kcatf.*kinetics.Kcatr);

% ------------------- Parameter balancing end -----------------------------
% -------------------------------------------------------------------------

%% Start with the ECM-algorithm

% combine pathway stoichiometries of main pathways (e.g. CBB) and
% connecting modules (e.g. GAPtoOAA) in a cell array
% the arrays are defined for three possible products: acetyl-CoA,
% oxaloacetate and glaceraldehyde-3-phosphate (GAP)
cycle_ids_AcCoA =   {{'CBB',0.5,'GAPtoOAA',0.5,'C3to2C2(4HB)(T)',0.5,'-PyrC',-0.5},...
                    {'CBB',0.55,'Photorespiration',0.05,'GAPtoOAA',0.5,'C3to2C2(4HB)(T)',0.5,'-PyrC',-0.5},...
                    {'CBB',1,'GAPtoOAA',1,'PyrKin',1,'PEPCarb',1,'PyrDehCompl',1},...
                    {'HB/HP-Cren',1},{'HB/HP-Thaum',1},...
                    {'HP-Bicycle',0.5,'C3to2C2(4HB)(T)',1},...
                    {'Serine-Cycle',1},{'Serine-Cycle (w/o SHMT)',1},...
                    {'RuMP-Cycle',0.5,'GAPtoOAA',0.5,'C3to2C2(4HB)(T)',0.5,'-PyrC',-0.5},...
                    {'RuMP-Cycle(Formate/THF)',0.5,'GAPtoOAA',0.5,'C3to2C2(4HB)(T)',0.5,'-PyrC',-0.5},...
                    {'rCCC',1},...
                    {'CETCH',1,'b-OH-AspCycle',0.5,'C3to2C2(4HB)(T)',0.5,'-PyrC',-0.5},...
                    {'CETCH',1,'b-OH-AspCycle',0.5,'MCG-like-cycle',0.5},...
                    {'CETCH',1,'RevGlyoxShunt',1},...
                    {'rGlyP',1,'PyrDehCompl',1},...
                    {'rGlyP (w/o SHMT)',1,'PyrDehCompl',1},...
                    {'2-HG-rTCA',1}};

cycle_ids_Oxac =    {{'CBB',1,'GAPtoOAA',1},...
                    {'CBB(Photorespiration)',1.1,'Photorespiration',0.1,'GAPtoOAA',1},...
                    {'CBB(PTS)',1,'GAPtoOAA',1},...
                    {'HB/HP-Cren',1,'rCC-AcCoAtoPyr',1,'PyrC',1},...
                    {'HB/HP-Cren',1,'AcCoAtoOAA(HP)(AMP)',1},...
                    {'HB/HP-Thaum',1,'rCC-AcCoAtoPyr',1,'PyrC',1},...
                    {'HB/HP-Thaum',1,'AcCoAtoOAA(HP)(ADP)',1},...
                    {'HP-Bicycle',1,'PyrC',1},...
                    {'HP-Bicycle(ADP)',1,'PyrC',1},...
                    {'HP-Bicycle(glyox)',2,'b-OH-AspCycle',1},...
                    {'HP-Bicycle(glyox)(ADP)',2,'b-OH-AspCycle',1},...
                    {'Serine-Cycle (w/o SHMT)',1,'EtMalOAAGlyox',0.5,'GlyoxToOAA(Ser)',0.5},...
                    {'Serine-Cycle',1,'EtMalOAAGlyox',0.5,'GlyoxToOAA(Ser)',0.5},...
                    {'RuMP-Cycle',1,'GAPtoOAA',1},...
                    {'RuMP-Cycle(Formate/THF)',1,'GAPtoOAA',1},...
                    {'rCCC',1,'rCC-AcCoAtoPyr',1,'PyrC',1},...
                    {'CETCH',2,'b-OH-AspCycle',1},...
                    {'CETCH',1,'GlyoxToOAA-HPbicycle(AMP)',1},...
                    {'rGlyP',1,'PyrC',1},...
                    {'rGlyP (w/o SHMT)',1,'PyrC',1},...
                    {'2-HG-rTCA',1,'rCC-AcCoAtoPyr',1,'PyrC',1}};

cycle_ids_GAP =     {{'CBB',1},...
                    {'CBB(Photorespiration)',1.1,'Photorespiration',0.1},...
                    {'CBB(PTS)',1},...
                    {'HB/HP-Cren',1,'rCC-AcCoAtoPyr',1,'PyrC',1,'OAAtoGAP',1},...
                    {'HB/HP-Cren',1,'AcCoAtoOAA(HP)(AMP)',1,'OAAtoGAP',1},...
                    {'HB/HP-Thaum',1,'rCC-AcCoAtoPyr',1,'PyrC',1,'OAAtoGAP',1},...
                    {'HB/HP-Thaum',1,'AcCoAtoOAA(HP)(ADP)',1,'OAAtoGAP',1},...
                    {'HP-Bicycle',1,'PyrC',1,'OAAtoGAP',1},...
                    {'HP-Bicycle(ADP)',1,'PyrC',1,'OAAtoGAP',1},...
                    {'HP-Bicycle(glyox)',2,'b-OH-AspCycle',1,'OAAtoGAP',1},...
                    {'HP-Bicycle(glyox)(ADP)',2,'b-OH-AspCycle',1,'OAAtoGAP',1},...
                    {'Serine-Cycle (w/o SHMT)',1,'EtMalOAAGlyox',0.5,'GlyoxToOAA(Ser)',0.5,'OAAtoGAP',1},...
                    {'Serine-Cycle',1,'EtMalOAAGlyox',0.5,'GlyoxToOAA(Ser)',0.5,'OAAtoGAP',1},...
                    {'RuMP-Cycle',1},...
                    {'RuMP-Cycle(Formate/THF)',1,},...
                    {'rCCC',1,'rCC-AcCoAtoPyr',1,'PyrC',1,'OAAtoGAP',1},...
                    {'CETCH',2,'b-OH-AspCycle',1,'OAAtoGAP',1},...
                    {'CETCH',1,'GlyoxToOAA-HPbicycle(AMP)',1,'OAAtoGAP',1},...
                    {'rGlyP',1,'PyrC',1,'OAAtoGAP',1},...
                    {'rGlyP (w/o SHMT)',1,'PyrC',1,'OAAtoGAP',1},...
                    {'2-HG-rTCA',1,'rCC-AcCoAtoPyr',1,'PyrC',1,'OAAtoGAP',1}};

% choose the product for the cycles:
cycle_ids = cycle_ids_AcCoA;
%target_metabolite = 'C_oxaloacetate';
%target_metabolite = 'C_d_glyceraldehyde_3_phosphate';
target_metabolite = 'C_acetyl_coa';
% define target metabolite minimal concentration:
tm_conc = 0.01;

%randomize the kinetic parameters according to std from parameter balancing
n_rand = 3;

conc_max_2 = conc_max;
conc_min_2 = conc_min;

% do the ECM optimization:
[Cap_Cell,Cap_Rev_Cell,Cap_Rev_Sat_Cell,SubNetReactNames_Cell,pathway_activities,pathway_activities_std] = ECM_Compare_Pathways_Std(network,cycle_ids,kinetics,kinetics_std,target_metabolite,tm_conc,conc_min_2,conc_max_2,n_rand,{{'R_glycine_hy_R00945'},{'R_glycine_hy_R00945'},{'R_ribulose_b_R03140'}},{'rGlyP (w/o SHMT)','Serine-Cycle (w/o SHMT)','Photorespiration'});

% Calculate projected ATP-costs for each cycle:
pathway_costs = zeros(numel(cycle_ids),1);
for i_ECM=1:numel(cycle_ids)
    cycle_id = cycle_ids{i_ECM};
    %[v_r, v] = GetPathwayStoichiometry(cycle_id, target_metabolite);
    [v_r, v] = CombinePathways(cycle_id);
    pathway_costs(i_ECM) = CalcProjATPcosts(v_r, v, network);
end

% plotting section
Compare_Pathways_Std_Plot('CO2 fixing pathways',cycle_ids,Cap_Cell,Cap_Rev_Cell,Cap_Rev_Sat_Cell,SubNetReactNames_Cell,pathway_activities,pathway_activities_std,pathway_costs);
switch(target_metabolite)
    case 'C_d_glyceraldehyde_3_phosphate'
        CapRevSat_Plot('',cycle_ids,Cap_Cell,Cap_Rev_Cell,Cap_Rev_Sat_Cell,SubNetReactNames_Cell,[1 16 21],1,3,{'CBB cycle','rCCC','2-HG-rTCA'})
        set(gcf,'Units','centimeters','position',[10 10 9 20]);
    case 'C_acetyl_coa'
        CapRevSat_Plot('',cycle_ids,Cap_Cell,Cap_Rev_Cell,Cap_Rev_Sat_Cell,SubNetReactNames_Cell,[1 11 17],1,3,{'CBB cycle','rCCC','2-HG-rTCA'})
        set(gcf,'Units','centimeters','position',[10 10 9 20]);
end
```
