## Supplementary material for "In-depth computational analysis of natural and artificial carbon fixation pathways": MATLAB files: CombinePathways.m

```
function [reaction_names, fluxes] = CombinePathways(PathwaysAndStoichiometries)

n = numel(PathwaysAndStoichiometries)/2;

[reaction_names, fluxes] = GetPathwayStoichiometry(PathwaysAndStoichiometries{1}, 'none');
fluxes = fluxes*PathwaysAndStoichiometries{2};

for i = 2:n
    [rn, vv] = GetPathwayStoichiometry(PathwaysAndStoichiometries{2*i-1}, 'none');
    vv = vv*PathwaysAndStoichiometries{2*i};
    indices = label_names(rn,reaction_names);
    fluxes(indices(indices~=0)) = fluxes(indices(indices~=0)) + vv(indices~=0);
    reaction_names = [reaction_names, rn(indices==0)];
    fluxes = [fluxes; vv(indices==0)];
    % eliminate reactions with flux 0:
    reaction_names(fluxes == 0) = [];
    fluxes(fluxes == 0) = [];
end

end
```
