## Supplementary material for "In-depth computational analysis of natural and artificial carbon fixation pathways": MATLAB files: Compare_Pathways_Std_Plot.m

```
function Compare_Pathways_Std_Plot(str_title,cycle_ids,Cap_Cell,Cap_Rev_Cell,Cap_Rev_Sat_Cell,SubNetReactNames_Cell,pathway_activities,pathway_activities_std,pathway_costs)

% bf_colormap =   [80 40 0; % Calvin cycle
%                 90 60 0; % HP/HB cycle
%                 95 90 25; % HP bicycle
%                 0 60 50; % Serine cycle
%                 35 70 90; % RuMP cycle
%                 80 60 70; % rCC cycle
%                 0 30 47; % CETCH cycle
%                 20 20 20]/100; % rGC pathw.
bf_colormap =   [51 34 136; % Calvin cycle
                136 204 238; % HP/HB cycle (Cren)
                68 170 153; % HP/HB cycle (Thaum)
                17 119 51; % HP bicycle
                153 153 51; % Serine cycle
                221 204 119; % RuMP cycle
                204 102 119; % rCC cycle
                136 34 85; % CETCH cycle
                170 68 153; % rGC pathw.
                221 221 221]/255; % 2-HA/rTCA
% bf_colormap =   [119 170 221; % Calvin cycle
%                 153 221 255; % HP/HB cycle (Cren)
%                 68 187 153; % HP/HB cycle (Thaum)
%                 187 204 51; % HP bicycle
%                 170 170 0; % Serine cycle
%                 238 221 136; % RuMP cycle
%                 238 136 102; % rCC cycle
%                 255 170 187; % CETCH cycle
%                 221 221 221; % rGC pathw.
%                 50 50 50]/255; % 2-HA/rTCA
pathway_names = {'CBB','HP/HB (Crenarchaea)','HP/HB (Thaumarchaea)','HP Bicycle','Serine Cycle','RuMP Cycle','rCC Cycle','CETCH Cycle','rGlyP','2-HG-rTCA Cycle'};


figure
sgtitle(str_title);
sp_y = round(sqrt(numel(cycle_ids)*3/4)+0.49);
sp_x = round(sp_y*4/3+0.49);

cycle_names = cell(numel(cycle_ids),1);

for i_ECM = 1:numel(cycle_ids)
    cycle = cycle_ids{i_ECM};
    %cycle_names{i_ECM} = ['{\fontfamily{cmss}\selectfont \textbf{' cycle{1} '}'];
    cycle_names{i_ECM} = cycle{1};
    for i = 2:(numel(cycle)/2)
        cycle_names{i_ECM} = [cycle_names{i_ECM} ' | ' cycle{i*2-1}];
    end
    %cycle_names{i_ECM} = [cycle_names{i_ECM} '}'];
    M = [Cap_Cell{i_ECM}, Cap_Rev_Cell{i_ECM} - Cap_Cell{i_ECM}, Cap_Rev_Sat_Cell{i_ECM} - Cap_Rev_Cell{i_ECM}];
    subplot(sp_y, sp_x, i_ECM);
    h = bar(1:length(Cap_Cell{i_ECM}),M, 'stacked','EdgeColor','none','BarWidth',0.8);
    my_colormap = bf_colormap([8 7 6],1:3);%[1-((1-bf_colormap(5,1:3))); 1-((1-bf_colormap(5,1:3))*0.8); 1-((1-bf_colormap(5,1:3))*0.6)];%[0.35 0.35 0.9; 0.8 0.2 0.7; 1 0.3 0.2];
    for it = 1:length(h)
      h(it).FaceColor = 'Flat';
      h(it).CData = my_colormap(it,:);
    end
    ylabel('Enzyme demand [mg/L]');
    set(gca,'Xtick',1:length(Cap_Cell{i_ECM}),'xticklabel',SubNetReactNames_Cell{i_ECM},'TickLabelInterpreter','none');
    xtickangle(45);
    legend('Capacity','Reversibility','Saturation','Location','SouthWest');
    title(cycle_names{i_ECM});
    ax = gca;
    ax.XAxis.TickLength = [0 0];
    ax.XAxis.FontSize = 8;
end

% apply different colors to each primary pathway
pathway_colors = zeros(length(pathway_activities),3);
pathway_indices = zeros(length(pathway_activities),1);
for i=1:length(pathway_activities)
    cycle_id = cycle_ids{i};
    if(strfind(cycle_id{1},'CBB')==1)
        pathway_indices(i) = 1;
        pathway_colors(i,:) = bf_colormap(1,:);
    elseif(strfind(cycle_id{1},'HB/HP-Cren')==1)
        pathway_indices(i) = 2;
        pathway_colors(i,:) = bf_colormap(2,:);
    elseif(strfind(cycle_id{1},'HB/HP-Thaum')==1)
        pathway_indices(i) = 3;
        pathway_colors(i,:) = bf_colormap(3,:);
    elseif(strfind(cycle_id{1},'HP-Bicycle')==1)
        pathway_indices(i) = 4;
        pathway_colors(i,:) = bf_colormap(4,:);
    elseif(strfind(cycle_id{1},'Serine')==1)
        pathway_indices(i) = 5;
        pathway_colors(i,:) = bf_colormap(5,:);
    elseif(strfind(cycle_id{1},'RuMP')==1)
        pathway_indices(i) = 6;
        pathway_colors(i,:) = bf_colormap(6,:);
    elseif(strfind(cycle_id{1},'rCCC')==1)
        pathway_indices(i) = 7;
        pathway_colors(i,:) = bf_colormap(7,:);
    elseif(strfind(cycle_id{1},'CETCH')==1)
        pathway_indices(i) = 8;
        pathway_colors(i,:) = bf_colormap(8,:);
    elseif(strfind(cycle_id{1},'rGlyP')==1)
        pathway_indices(i) = 9;
        pathway_colors(i,:) = bf_colormap(9,:);
    elseif(strfind(cycle_id{1},'2-HG-rTCA')==1)
        pathway_indices(i) = 10;
        pathway_colors(i,:) = bf_colormap(10,:);
    end
end

% % Vertical Bar plot: activities
% legend_unique = zeros(length(pathway_activities),1);
% legend_handles = [];
% figure
% hold on
% % my_colormap = [0.35 0.35 0.9; 0.8 0.2 0.7; 1 0.3 0.2];
% for it = 1:length(pathway_activities)
%   hh = bar(it, pathway_activities(it),'EdgeColor','none','BarWidth',0.8);
%   if(legend_unique(pathway_indices(it)) == 0)
%       legend_handles = [legend_handles hh];
%   end
%   h = errorbar(it, pathway_activities(it),pathway_activities_std(it),pathway_activities_std(it));
%   h.Color = [0 0 0];
%   h.LineStyle = 'none';
%   hh.FaceColor = 'Flat';
%   hh.CData = pathway_colors(it,:);
%   legend_unique(pathway_indices(it)) = legend_unique(pathway_indices(it))+1;
% end
% hold off
% legend(legend_handles,pathway_names);
% ylabel('Pathway activity [�mol/(min mg)]');
% set(gca,'Xtick',1:length(pathway_activities),'xticklabel',cycle_names,'TickLabelInterpreter','none');%'TickLabelInterpreter','latex','FontName','Computer Modern Sans Serif');
% xtickangle(45);
% title(str_title);

% Horizontal Bar plot: activities
legend_unique = zeros(length(pathway_activities),1);
legend_handles = [];
figure('Units','centimeters','position',[10 10 18.5 9]);
hold on
% my_colormap = [0.35 0.35 0.9; 0.8 0.2 0.7; 1 0.3 0.2];
for it = length(pathway_activities):-1:1
  hh = barh(length(pathway_activities)-it+1, pathway_activities(it),'EdgeColor','none','BarWidth',0.8);
  if(legend_unique(pathway_indices(it)) == 0)
      legend_handles = [legend_handles hh];
  end
  h = errorbar(pathway_activities(it),length(pathway_activities)-it+1,pathway_activities_std(it),pathway_activities_std(it),'horizontal','CapSize',3);
  h.Color = [0 0 0];
  h.LineStyle = 'none';
  hh.FaceColor = 'Flat';
  hh.CData = pathway_colors(it,:);
  legend_unique(pathway_indices(it)) = legend_unique(pathway_indices(it))+1;
end
hold off
%legend(legend_handles,pathway_names{numel(pathway_names):-1:1},'FontSize',8,'Location','southeast');
legend(legend_handles(numel(pathway_names):-1:1),pathway_names{1:numel(pathway_names)},'FontSize',8,'Location','northeast');
xlabel('Pathway activity [�mol/(min mg)]','FontSize',10);
%yticks(1:length(pathway_activities));
set(gca,'Ytick',1:length(pathway_activities),'yticklabel',cycle_names(numel(cycle_names):-1:1),'TickLabelInterpreter','none','FontSize',8);%'TickLabelInterpreter','latex','FontName','Computer Modern Sans Serif');
%xtickangle(45);
title(str_title,'FontSize',9);
%hExport('C:\Users\Hannes L�we\Documents\MATLAB\Carbon fixation model\ECM - CF\TestFig.pdf',gcf)

% Scatter plot: activities vs. costs
legend_unique = zeros(length(pathway_activities),1);
legend_handles = [];
figure('Units','centimeters','position',[10 10 18.5 9]);
threshold = 1.2;
bBreak = 0;
if(sum(pathway_activities > threshold) > 0)
    bBreak = 1;
    figureYbreak([0 0.6], 4, [threshold threshold+0.2], 0.05, 0.1, 0.1);
    pathway_activities(pathway_activities > threshold) = pathway_activities(pathway_activities > threshold) -threshold+0.6+0.2;
end
for it = 1:length(pathway_activities)
    
    h = plot(pathway_costs(it),pathway_activities(it),'o','MarkerFaceColor',pathway_colors(it,:),'MarkerEdgeColor',[0 0 0],'MarkerSize',8,'Linewidth', 1);

    if(it ==1)
        hold on
    end
    if(legend_unique(pathway_indices(it)) == 0)
        legend_handles = [legend_handles h];
    end
    cycle_id = cycle_ids{it};
    labelpoints(pathway_costs(it)-0.14,pathway_activities(it)+0.015,cycle_id{1},'FontSize',7);
    legend_unique(pathway_indices(it)) = legend_unique(pathway_indices(it))+1;
end

if(bBreak == 1)
    figureYbreak([0 0.6], 4, [threshold threshold+0.2], 0.05, 0.1, 0.1);
end

legend(legend_handles,pathway_names,'FontSize',8);
xlabel('Pathway costs, ATP/product','FontSize',10);
ylabel('Pathway activity, �mol/(min mg)','FontSize',10);
title(str_title);
xlim([0 11]);
hold off

end
```
