## Supplementary material for "In-depth computational analysis of natural and artificial carbon fixation pathways": MATLAB files: ECM_Compare_Pathways_Std.m

```
function [Cap_Cell,Cap_Rev_Cell,Cap_Rev_Sat_Cell,SubNetReactNames_Cell,pathway_activities,pathway_activities_std] = ECM_Compare_Pathways_Std(network,cycle_ids,kinetics,kinetics_std,target_metabolite,conc_target_metabolite,conc_min,conc_max,n_rand,ignore_reactions,ir_pathways)

Cap_Cell = cell(1,numel(cycle_ids));
Cap_Rev_Cell = cell(1,numel(cycle_ids));
Cap_Rev_Sat_Cell = cell(1,numel(cycle_ids));
SubNetReactNames_Cell = cell(1,numel(cycle_ids));

pathway_activities = zeros(numel(cycle_ids),1);
pathway_activities_std = zeros(numel(cycle_ids),1);

f = waitbar(0,'Optimization running...');

nFinal = ones(numel(cycle_ids),1)*n_rand;

for i_rand = 1:n_rand
kinetics_out = RandomizeParamters(kinetics, kinetics_std, network,i_rand);
network.kinetics = kinetics_out;


%loop through the pathways
for i_ECM = 1:numel(cycle_ids)
    
cycle_id = cycle_ids{i_ECM};

[v_r, v] = CombinePathways(cycle_id);

vv = zeros(length(network.reaction_names),1);

for i = 1:length(v_r)
    index = find(strcmp(network.reaction_names, v_r{i}));
    vv(index(1)) = v(i);
end

[subnetwork, ind_m, ind_r] = network_subnetwork_HL(network,1:length(network.metabolites),find(vv),1);

% For the RuMP-cycle, cofactor concentration ranges need to be changed to
% get a feasible, metabolite profile
if(~isempty(strfind(cycle_id{1},'RuMP')) && ~isempty(conc_min) && ~isempty(conc_max))
    conc_max(3:6) = conc_max(3:6)*10;
    conc_min(3:6) = conc_min(3:6)/10;
end

ecm_options = ecm_default_options(subnetwork, 'My example model');
%ecm_options.c_data = c_data;
%ecm_options.u_data = u_data;
if(~isempty(conc_min))
    ecm_options.conc_min = conc_min(ind_m);
end
if(~isempty(conc_max))
    ecm_options.conc_max = conc_max(ind_m);
end

i_TargMetab = find(strcmp(subnetwork.metabolites,target_metabolite));
ecm_options.conc_min(i_TargMetab) = conc_target_metabolite; % minimal concentration to keep metabolism running
ecm_options.ecm_scores = {'emc4cm'};

ecm_options = ecm_update_options(subnetwork, ecm_options);
subnetwork.protein_molecular_mass = network.protein_molecular_mass(ind_r);
ecm_options.enzyme_cost_weights = subnetwork.protein_molecular_mass;

% Run ECM
% if(i_ECM == 1)
%     fprintf('pause\n');
% end


if(exist('ignore_reactions'))
    test_index = 0;
    for k = 1:(numel(cycle_id)/2)
        for j = 1:numel(ir_pathways)
            if(strcmp(cycle_id{k*2-1},ir_pathways{j}))
                test_index = j;
            end
        end
    end
    if(test_index)
        ir = ignore_reactions{test_index};
        for j = 1:numel(ir)
            ind_SHMT = find(strcmp(subnetwork.reaction_names,ir{j}));
            subnetwork.kinetics.Kcatf(ind_SHMT) = subnetwork.kinetics.Kcatf(ind_SHMT)*10^3;
            subnetwork.kinetics.Kcatr(ind_SHMT) = subnetwork.kinetics.Kcatr(ind_SHMT)*10^3;
            %subnetwork.kinetics.KM(ind_SHMT) = subnetwork.kinetics.KM(ind_SHMT)/100;
        end
    end
end


[c, u, u_cost, up, A_forward, mca_info, c_min, c_max, u_min, u_max, kinetics_new, u_capacity, eta_energetic, eta_saturation] = ecm_enzyme_cost_minimization(subnetwork, subnetwork.kinetics, vv(vv~=0), ecm_options);
%fluxes_at_optimum = modular_velocities('cs', subnetwork.N, subnetwork.regulation_matrix, find(subnetwork.external), u.emc4cm, c.emc4cm, subnetwork.kinetics.KA, subnetwork.kinetics.KI, subnetwork.kinetics.KM, subnetwork.kinetics.KV, subnetwork.kinetics.Keq, subnetwork.kinetics.h);

if(~isempty(c))
    % recalculate u_capacity, eta_saturation and eta_energetic as the
    % original function gives weird results from time to time:
    [u_capacity, eta_saturation, eta_energetic] = calculate_rev_sat_HL('cs', subnetwork.N, subnetwork.regulation_matrix, find(subnetwork.external), u.emc4cm, c.emc4cm, subnetwork.kinetics.KA, subnetwork.kinetics.KI, subnetwork.kinetics.KM, subnetwork.kinetics.KV, subnetwork.kinetics.Keq, subnetwork.kinetics.h);
    if(exist('ignore_reactions'))
        test_index = 0;
        for k = 1:(numel(cycle_id)/2)
            for j = 1:numel(ir_pathways)
                if(strcmp(cycle_id{k*2-1},ir_pathways{j}))
                    test_index = j;
                end
            end
        end
        if(test_index)
            ir = ignore_reactions{test_index};
            for j = 1:numel(ir)
                ind_SHMT = find(strcmp(subnetwork.reaction_names,ir{j}));
                subnetwork.protein_molecular_mass(ind_SHMT) = [];
                u.emc4cm(ind_SHMT) = [];
                u_capacity(ind_SHMT) = [];
                eta_energetic(ind_SHMT) = [];
                eta_saturation(ind_SHMT) = [];
                subnetwork.reaction_names(ind_SHMT) = [];
            end
        end
    end


    if(sum(eta_saturation > 1) > 0)
        fprintf('error!\m')
    end

    % Save the resulting predictions in dedicated matrices and cell arrays
    % and calculate standard deviations:
    if(i_rand == 1)
        Cap_Cell(i_ECM) = {subnetwork.protein_molecular_mass.*u_capacity/nFinal(i_ECM)};
        Cap_Rev_Cell(i_ECM) = {eta_energetic/nFinal(i_ECM)};
        Cap_Rev_Sat_Cell(i_ECM) = {eta_saturation/nFinal(i_ECM)};
        if(eta_saturation > 1)
            fprintf('error!\m')
        end
        test = [u_capacity./eta_energetic./eta_saturation u.emc4cm];
        SubNetReactNames_Cell{i_ECM} = subnetwork.reaction_names;
        pathway_activities(i_ECM) = 60000/sum(subnetwork.protein_molecular_mass.*u.emc4cm)/nFinal(i_ECM);
        pathway_activities_std(i_ECM) = (60000/sum(subnetwork.protein_molecular_mass.*u.emc4cm)).^2/nFinal(i_ECM);
    else
        Cap_Cell(i_ECM) = {Cap_Cell{i_ECM} + subnetwork.protein_molecular_mass.*u_capacity/nFinal(i_ECM)};
        Cap_Rev_Cell(i_ECM) = {Cap_Rev_Cell{i_ECM} + eta_energetic/nFinal(i_ECM)};
        Cap_Rev_Sat_Cell(i_ECM) = {Cap_Rev_Sat_Cell{i_ECM} + eta_saturation/nFinal(i_ECM)};
        pathway_activities(i_ECM) = pathway_activities(i_ECM) + 60000/sum(subnetwork.protein_molecular_mass.*u.emc4cm)/nFinal(i_ECM);
        pathway_activities_std(i_ECM) = pathway_activities_std(i_ECM) + (60000/sum(subnetwork.protein_molecular_mass.*u.emc4cm)).^2/nFinal(i_ECM);
    end
    if(i_rand == n_rand) % last iteration
        pathway_activities_std(i_ECM) = sqrt(pathway_activities_std(i_ECM) - pathway_activities(i_ECM)^2);
        if(sum(Cap_Rev_Cell{i_ECM} > 1) > 0)
            fprintf('error!\m')
        end
        Cap_Rev_Cell(i_ECM) = {Cap_Cell{i_ECM}./Cap_Rev_Cell{i_ECM}};
        if(sum(Cap_Rev_Sat_Cell{i_ECM} > 1) > 0)
            fprintf('error!\m')
        end
        Cap_Rev_Sat_Cell(i_ECM) = {Cap_Rev_Cell{i_ECM}./Cap_Rev_Sat_Cell{i_ECM}};
    end
    
else
    nFinal(i_ECM) = nFinal(i_ECM)-1;
    fff = (nFinal(i_ECM)+1)/nFinal(i_ECM);
    Cap_Cell(i_ECM) = {Cap_Cell{i_ECM} * fff};
    Cap_Rev_Cell(i_ECM) = {Cap_Rev_Cell{i_ECM} * fff};
    Cap_Rev_Sat_Cell(i_ECM) = {Cap_Rev_Sat_Cell{i_ECM} * fff};
    pathway_activities(i_ECM) = pathway_activities(i_ECM) * fff;
    pathway_activities_std(i_ECM) = pathway_activities_std(i_ECM) * fff;
end

end
waitbar(i_rand/n_rand,f,'Optimization running...');
end % Parameter-Randomization loop

close(f);

end
```
