## Supplementary material for "In-depth computational analysis of natural and artificial carbon fixation pathways": MATLAB files: ecm_enzyme_cost_minimization_HL.m

```
function [c, u, u_cost, up, A_forward, mca_info, c_min, c_max, u_min, u_max, r, u_capacity, eta_energetic, eta_saturation, multi] = ecm_enzyme_cost_minimization_HL(network,r,v,ecm_options)

% ECM_ENZYME_COST_MINIMIZATION - Compute optimal flux-specific enzyme costs for given flux distribution
%
% [c, u, u_cost, up, A_forward, mca_info, c_min, c_max, u_min, u_max, r, u_capacity, eta_energetic, eta_saturation, multi] = ecm_enzyme_cost_minimization(network, e, v, ecm_options)
%
% Input 
%   network       metabolic network structure (as in Metabolic Network Toolbox)
%   r             Kinetic constants (from parameter balancing)
%   v             flux mode
%   ecm_options   options struct (for fields and default values, see 'ecm_default_options')
%
% Output
%   c                 metabolite concentrations
%   c.data            data vector (if provided)
%   c.std             data standard deviations vector (if provided)
%   c.fixed           fixed concentration vector
%   c.initial         initial solution vector 
%   c.[SCORE]         1st column: result from optimising the score [SCORE]
%                     other columns: possible sampled solutions
%                     
%   u                 enzyme levels (all enzymes)
%   u.data            data vector
%   u.std             data standard deviations vector (if provided)
%   u.[SCORE]         1st column: result from optimising the score [SCORE]
%                     other columns: possible sampled solutions
%
%   up                enzyme levels (only "scored" enzymes, given by ecm_scores.ind_scored_enzymes)
%   up.[SCORE]        1st column: result from optimising the score [SCORE]
%                     other columns: possible sampled solutions
%
%   u_cost            sum of scored enzyme levels 
%   u_cost.data       data value
%   u_cost.[SCORE]    1st column: result from optimising the score [SCORE]
%                     other columns: possible sampled solutions
%
%   A_forward         reaction affinity (in flux direction)
%   A_forward.initial    initial solution vector 
%   A_forward.[SCORE] 1st column: result from optimising the score [SCORE]
%                     other columns: possible sampled solutions
%
%   mca_info          control analysis results
%   c_min             minimal possible metabolite values (for tolerance calculations)
%   c_max             maximal possible metabolite values (for tolerance calculations)
%   u_min             minimal possible enzyme values (for tolerance calculations)
%   u_max             maximal possible enzyme values (for tolerance calculations)
%   r                 kinetics used
%   u_capacity        enzyme capacities (= abs(v)/kcat_forward)
%   eta_energetic     energy efficiency factors
%   eta_saturation    saturation efficiency factors
%   multi             results from optimisation runs with multiple starting points
%   
% 
% The following fields of ecm_options are used in this function:
%
%    ecm_options.c_data             
%    ecm_options.u_data               
%    ecm_options.conc_min_default   
%    ecm_options.conc_max_default   
%    ecm_options.conc_min
%    ecm_options.conc_max
%    ecm_options.ind_met_fix
%    ecm_options.conc_fix
%    ecm_options.ind_scored_enzymes 
%    ecm_options.enzyme_cost_weights
%    ecm_options.lambda_regularisation
%    ecm_options.ecm_scores         
%    ecm_options.initial_choice
%    ecm_options.x_start
%    ecm_options.multiple_starting_points   flag
%    ecm_options.compute_hessian
%    ecm_options.compute_elasticities
%    ecm_options.compute_tolerance,
%    ecm_options.cost_tolerance_factor

if sum(isnan(v)),
  warning('Flux vector contains nan values; replacing them by zero values');
  v(isnan(v)) = 0;
end

% --------------------------------------------------------------------------------------
% Enzyme cost minimization
% --------------------------------------------------------------------------------------

% * the last time I checked (with 2020), interior-point worked best *

%opt = optimset('MaxFunEvals',10^15,'MaxIter',10^15,'Display','off','TolX',10^-10,'Algorithm','sqp');
%opt = optimset('MaxFunEvals',10^15,'MaxIter',10^15,'TolX',10^-10,'Display','off','Algorithm','active-set');
opt = optimset('MaxFunEvals',10^15,'MaxIter',10^15,'TolX',10^-10,'Display','off','Algorithm','interior-point');
%'Display','iter',

%network.N(network_find_water(network),:) = 0;
network.N(find(ismember(network.metabolites, 'H2O')),:) = 0;

network.kinetics = set_kinetics(network,'cs',r);

% --------------------------------------------------------------------------------------
% initialise 

ecm_options.enzyme_cost_weights  = ecm_options.enzyme_cost_weights(find(v(ecm_options.ind_scored_enzymes)~=0));
ecm_options.ind_scored_enzymes   = ecm_options.ind_scored_enzymes(find(v(ecm_options.ind_scored_enzymes)~=0));

if ecm_options.ignore_regulation,
  network.regulation_matrix = 0*network.regulation_matrix;
end

[Mplus, Mminus, Wplus, Wminus, nm, nr] = make_structure_matrices(network.N,network.regulation_matrix,find(network.external));

c_data                = ecm_options.c_data             ;
u_data                = ecm_options.u_data             ;  
if isfield(ecm_options,'c_std'),
  c_std = ecm_options.c_std;
else
  c_std = nan * c_data;
end
if isfield(ecm_options,'u_std'),
  u_std = ecm_options.u_std;
else
  u_std = nan * u_data;
end
conc_min_default      = ecm_options.conc_min_default   ;
conc_max_default      = ecm_options.conc_max_default   ;
conc_min              = ecm_options.conc_min;
conc_max              = ecm_options.conc_max;
ind_met_fix           = ecm_options.ind_met_fix;
conc_fix              = ecm_options.conc_fix;
ecm_scores            = ecm_options.ecm_scores         ;
enzyme_cost_weights   = ecm_options.enzyme_cost_weights;
lambda_regularisation = ecm_options.lambda_regularisation;
ind_scored_enzymes    = ecm_options.ind_scored_enzymes ;
pp.fluctuations_safety_margin = ecm_options.fluctuations_safety_margin;
pp.cell_volume                = ecm_options.cell_volume;

if find(v(ind_scored_enzymes)==0), 
  error('An enzyme catalysing a 0 flux is scored in the cost function. This can lead to problems')
end
ind_not_scored        = setdiff(1:nr,ind_scored_enzymes);
multiple_conditions_anticipate = ecm_options.multiple_conditions_anticipate;
multiple_conditions_n = ecm_options.multiple_conditions_n;

conc_min(isnan(conc_min)) = conc_min_default;
conc_max(isnan(conc_max)) = conc_max_default;


% --------------------------------------------------------------------------------------
% initialise c, u, u_cost; insert true concentrations and protein levels

c.data      = c_data;
u.data      = u_data;
u_cost.data = nansum(u.data);

if isempty(r),
  r = network.kinetics;
end

% --------------------------------------------------------------------------------------
% adjust some network- and kinetics-related variables

x_min = log(conc_min);
x_max = log(conc_max);
x_fix = nan * ones(nm,1);
x_fix(ind_met_fix) = log(conc_fix); %  , 
ind_fix = find(isfinite(x_fix)); 

x_min(ind_fix) = x_fix(ind_fix);% - 10^-4;
x_max(ind_fix) = x_fix(ind_fix);% + 10^-4;

ind_pos = find(v>=0);
ind_neg = find(v<0);

M_forward(ind_pos,:)       = Mplus(ind_pos,:);
M_forward(ind_neg,:)       = Mminus(ind_neg,:);
Mprod_forward(ind_pos,:)   = Mminus(ind_pos,:);
Mprod_forward(ind_neg,:)   = Mplus(ind_neg,:);
N_forward(:,ind_pos)       =  network.N(:,ind_pos);
N_forward(:,ind_neg)       = -network.N(:,ind_neg);
kc_forward(ind_pos,1)      = r.Kcatf(ind_pos);
kc_forward(ind_neg,1)      = r.Kcatr(ind_neg);
log_Keq_forward(ind_pos,1) = log(r.Keq(ind_pos));
log_Keq_forward(ind_neg,1) = -log(r.Keq(ind_neg));
KM                         = full(r.KM);
KM(network.N'==0)          = 1;
KMf                        = exp(sum(Mplus.*log(KM),2));
KMr                        = exp(sum(Mminus.*log(KM),2));
km_forward(ind_pos,1)      = KMf(ind_pos);
km_forward(ind_neg,1)      = KMr(ind_neg);
kmprod_forward(ind_pos,1)  = KMr(ind_pos);
kmprod_forward(ind_neg,1)  = KMf(ind_neg);


% --------------------------------------------------------------------------
% struct with parameters used in cost score functions

pp.v                     = v                  ; 
pp.network               = network            ;
pp.M_forward             = M_forward          ;
pp.Mprod_forward         = Mprod_forward      ;
pp.N_forward             = N_forward          ;
pp.log_Keq_forward       = log_Keq_forward    ;
pp.kc_forward            = kc_forward         ;
pp.km_forward            = km_forward         ;
pp.kmprod_forward        = kmprod_forward     ;
pp.ind_scored_enzymes    = ind_scored_enzymes ;
pp.enzyme_cost_weights   = enzyme_cost_weights;
pp.ind_not_scored        = ind_not_scored     ;
pp.multiple_conditions_anticipate   = multiple_conditions_anticipate;
pp.multiple_conditions_n = multiple_conditions_n;
[pp.ln_c_data, pp.ln_c_std] = lognormal_normal2log(c_data, c_std,'arithmetic'); 
pp.u_data                = u_data;
pp.u_std                 = u_std;
if isfield(network,'metabolite_mass'),
  pp.metabolite_mass       = network.metabolite_mass;
  pp.metabolite_mass(isnan(pp.metabolite_mass)) = nanmean(pp.metabolite_mass);
else
  pp.metabolite_mass       = ones(size(network.metabolites));
end
if isfield(network,'enzyme_mass'),
  pp.enzyme_mass       = network.enzyme_mass;
  pp.enzyme_mass(isnan(pp.enzyme_mass)) = nanmean(pp.enzyme_mass);
else
  pp.enzyme_mass       = ones(size(network.actions));
end

if multiple_conditions_n < 2, 
  pp.multiple_conditions_anticipate = 0;
end

if pp.multiple_conditions_anticipate,
  display(sprintf('  Running multi-condition optimisation with %d conditions', pp.multiple_conditions_n));
end

% --------------------------------------------------------------------------------------
% find feasible initial solution (as a test for feasibility) and extreme metabolite profiles

% minimal reaction GFE of 10^-10 RT kJ/mol is required;
Theta_min = 10^-10 * double(v~=0);  

if ecm_options.use_linear_cost_constraints,
  if isempty(ecm_options.maximal_u_cost),
    ecm_options.maximal_u_cost = 20 * nansum(ecm_options.enzyme_cost_weights .* abs(v(ecm_options.ind_scored_enzymes)) ./ kc_forward(ecm_options.ind_scored_enzymes));
    display(sprintf('Setting default upper bound %f for enzyme cost. Setting this value manually may speed up the calculation.',ecm_options.maximal_u_cost));
  end
  my_u_max = 10^15 * ones(nr,1); 
  my_u_max(ecm_options.ind_scored_enzymes) = ecm_options.maximal_u_cost ./ ecm_options.enzyme_cost_weights; 
  new_Theta_min =  abs(v) ./ my_u_max ./ kc_forward;
  Theta_min(ecm_options.ind_scored_enzymes) = new_Theta_min(ecm_options.ind_scored_enzymes);
end


try
  [x_start, x1, x2] = find_polytope_centre([],[], N_forward', log_Keq_forward - Theta_min, x_min, x_max, 0*x_min);
catch
  display(sprintf('*** No M-polytope centre found in search for initial solution - maybe concentration constraints are too tight'));
  display(sprintf('*** Flux distribution is not thermodynamically realisable'));
  display(sprintf('*** Use "eba_feasible_lp" to check the thermo-feasibility of your flux distribution (for given bounds on driving forces)'));
  display(sprintf('*** Note that a flux distribution may also be infeasible because of the equilibrium constants and concentration bounds'));
  display(sprintf(''));
  if ecm_options.fix_thermodynamics_by_adjusting_Keq,
    %% Enforce sufficiently large driving forces!
    Theta_min(Theta_min<1) = 1;
    display(sprintf('*** FROM NOW ON; REQUIRING REACTION AFFINITIES LARGER THAT 1 RT - NOTE THAT THIS MAY CAUSE SOME ARBITRARY ENZYME LEVELS'));
    display(sprintf('*** Running MDF to determine Keq adjustments to make this flux distribution feasible'));
    x_mdf = fmincon(@(xx) ecm_mdf(xx,pp), zeros(nm,1),[],[],[],[],x_min,x_max,@(xx) ecm_inequalities(xx,N_forward,log_Keq_forward, Theta_min),opt);
    Theta_mdf = log_Keq_forward-Theta_min-[N_forward' * x_mdf];
    adjustment_log_Keq_forward = -(Theta_mdf-Theta_min);
    adjustment_log_Keq_forward(Theta_mdf>=Theta_min) = 0;
    adjustment_log_Keq = sign(v) .* adjustment_log_Keq_forward;
    display(sprintf('*** Update your Keq vector to fix the problem. The maximal change in |ln Keq| is %f',max(abs(adjustment_log_Keq))));
    display(sprintf('*** Note that the new Keq values may be unrealistic and violate the Wegscheider conditions)'));
    display(sprintf('Please press key to continue'));
    pause
    log_Keq_forward = log_Keq_forward + adjustment_log_Keq_forward;
    pp.log_Keq_forward = log_Keq_forward;
    r.Keq = r.Keq .* exp(adjustment_log_Keq);
    pp.network.kinetics.Keq = r.Keq;
    [x_start, x1, x2] = find_polytope_centre([],[], N_forward', log_Keq_forward - Theta_min, x_min, x_max, 0*x_min);
  else
    error('Flux distribution cannot be thermodynamically realised given the Keq values and metabolite bounds')
    c = [];  u = [];  u_cost = [];  up = [];  A_forward = [];
    return
  end
end

X_extreme = [x1, x2]; % matrix with extreme initial concentration vectors


% --------------------------------------------------------------------------
% choose the starting point

switch ecm_options.initial_choice,
  case 'polytope_center',
    display('Choosing polytope center as the starting point'); 
  case 'interval_center',
    display('Choosing point close to center of allowed intervals as starting point'); 
    x_start = fmincon(@(xx) ecm_regularisation(xx,x_min,x_max, 1), x_start,[],[],[],[],x_min,x_max,@(xx) ecm_inequalities(xx,N_forward,log_Keq_forward, Theta_min),opt);
  case 'mdf',
    display('Choosing MDF solution as the starting point'); 
    x_start = fmincon(@(xx) ecm_mdf(xx,pp) + ecm_regularisation(xx,x_min,x_max,lambda_regularisation), x_start,[],[],[],[],x_min,x_max,@(xx) ecm_inequalities(xx,N_forward,log_Keq_forward, Theta_min),opt);
    if ecm_mdf(x_start,pp) > 0, error('Thermodynamically infeasible reaction'); end
  case 'given_x_start',
    display('Using given solution as a starting point');
    x_start = ecm_options.x_start;
end


% --------------------------------------------------------------------------------------
% run optimisations starting from x_start

display('  o Running optimisation using normal starting point');

x_init         = x_start;
c_init         = exp(x_init);
c.fixed        = exp(x_fix);
c.initial      = c_init;
A_forward_init = RT * [log_Keq_forward  - N_forward' * log(c_init)]; 
A_forward_init(ind_not_scored) = nan;
A_forward.initial = A_forward_init;

if find([A_forward.initial<0] .* [v~=0]),
  error('There is a thermodynamically infeasible reaction in the starting point');
end

if find([A_forward.initial<0.01] .* [v~=0]),
  display('  WARNING: There is a thermodynamically poor reaction (below 0.01 kJ/mol) in the starting point');
end

display(sprintf('Running ECM'));

for it_method = 1:length(ecm_scores),

  ecm_score = ecm_scores{it_method};
  display(sprintf('  o %s',ecm_score));
  
  [my_c, my_u, my_up, my_u_cost, my_A_forward, my_x, my_grad] = ecm_one_run(ecm_score,pp,x_min,x_max,x_init,lambda_regularisation, Theta_min,opt);

  c.(ecm_score)         = my_c;
  u.(ecm_score)         = my_u;
  up.(ecm_score)        = my_up;
  u_cost.(ecm_score)    = my_u_cost;
  A_forward.(ecm_score) = my_A_forward;
  mca_info.(ecm_score).x_gradient = my_grad;
  
  if ecm_options.compute_hessian,  
    my_x = log(c.(ecm_score));
    my_fct = @(xx) ecm_get_score(ecm_score,xx,pp) + ecm_regularisation(xx,x_min,x_max,lambda_regularisation);
    mca_info.(ecm_score).x_hessian = hessian(my_fct,my_x);
    mca_info.(ecm_score).rates     = ecm_get_specific_rates(ecm_score,my_x,pp);
    my_fct_2 = @(xx) log(ecm_get_score(ecm_score,xx,pp) + ecm_regularisation(xx,x_min,x_max,lambda_regularisation));
    log_hessian = hessian(my_fct_2,my_x);
    %% fix negative eigenvalues if they occur
    my_eigs = eig(log_hessian);
    if min(my_eigs)<0, log_hessian = log_hessian - 1.001 * min(my_eigs) * eye(nm);end
    mca_info.(ecm_score).x_log_hessian = log_hessian;
  end

  %% elasticities between the specific rate (i.e. v/u) and x
  if ecm_options.compute_elasticities,
    for it_r = 1:nr,
      mca_info.(ecm_score).elasticities_rate_x(it_r,:) = gradest(@(xx) ecm_get_specific_rates(ecm_score,xx,pp,it_r),my_x)';
    end
  end
  
end


% --------------------------------------------------------------------------------------
% run more optimisations starting from extreme points, or with relaxed optimality condition

c_min = [];
c_max = [];
u_min = [];
u_max = [];


for it_method = 1:length(ecm_scores),

  ecm_score = ecm_scores{it_method};
  if ecm_options.multiple_starting_points + ecm_options.compute_tolerance,
    display(sprintf('   %s',ecm_score));
  end

  %% run more optimisations starting from extreme points  
  if ecm_options.multiple_starting_points,
  
    display(sprintf('     Running optimisation with %d extreme starting points',size(X_extreme,2)));
    
    for itt = 1:size(X_extreme,2),

      %% start from points near the extreme points
      x_init = 0.1 * x_start + 0.9 * X_extreme(:,itt); 
      fprintf('%d ',itt)
      %% same as above
      c_init                         = exp(x_init);
      c.fixed                        = exp(x_fix);
      c.initial(:,1+itt)             = c_init;
      A_forward_init                 = RT * [log_Keq_forward  - N_forward' * log(c_init)]; 
      A_forward_init(ind_not_scored) = nan;
      A_forward.initial(:,1+itt)     = A_forward_init;
      
      [my_c, my_u, my_up, my_u_cost, my_A_forward] = ecm_one_run(ecm_score,pp,x_min,x_max,x_init,lambda_regularisation, Theta_min,opt);
      
      c.(ecm_score)(:,1+itt)         = my_c;
      u.(ecm_score)(:,1+itt)         = my_u;
      up.(ecm_score)(:,1+itt)        = my_up;
      u_cost.(ecm_score)(:,1+itt)    = my_u_cost;
      A_forward.(ecm_score)(:,1+itt) = my_A_forward;
      
    end
    
    score_from_standard_starting_point     = u_cost.(ecm_score)(1)
    best_score_in_multiple_starting_points = min(u_cost.(ecm_score))
    
    %% copy the best result to the first position
    [~,ind_opt] = min(u_cost.(ecm_score));
    
    multi.c.initial             = c.initial            ;  
    multi.c.(ecm_score)         = c.(ecm_score)        ;  
    multi.u.(ecm_score)         = u.(ecm_score)        ;  
    multi.up.(ecm_score)        = up.(ecm_score)       ;  
    multi.u_cost.(ecm_score)    = u_cost.(ecm_score)   ;  
    multi.A_forward.(ecm_score) = A_forward.(ecm_score);
    multi.A_forward.initial     = A_forward.initial;
    
    c.initial             = c.initial(:,ind_opt);  
    c.(ecm_score)         = c.(ecm_score)(:,ind_opt);  
    u.(ecm_score)         = u.(ecm_score)(:,ind_opt);  
    up.(ecm_score)        = up.(ecm_score)(:,ind_opt);  
    u_cost.(ecm_score)    = u_cost.(ecm_score)(ind_opt);  
    A_forward.(ecm_score) = A_forward.(ecm_score)(:,ind_opt);
    A_forward.initial     = A_forward.initial(:,ind_opt);

  end

  %% -------------------------------------------------------------------------
  %% compute tolerance in log concentration profile
  %% (at some maximum allowed percentage increase in cost)

  if ecm_options.compute_tolerance,

    display('    Computing the effects of relaxed optimality assumptions');
  
    x           = log(c.(ecm_score)(:,1));
    u_threshold = ecm_options.cost_tolerance_factor * u_cost.(ecm_score)(1);
  
    my_x_min = x;
    my_x_max = x;
  
    for it = 1:length(x),
      if x(it) ~= x_min(it),
        [my_x, my_x_min(it)] = fmincon(@(xx) xx(it), x,[],[],[],[],x_min,x_max,@(xx) ecm_below_threshold(ecm_score,xx,pp,u_threshold,x_min,x_max,lambda_regularisation), opt);
      end
      if x(it) ~= x_max(it),
        [my_x, my_x_max(it)] = fmincon(@(xx) -xx(it), x,[],[],[],[],x_min,x_max,@(xx) ecm_below_threshold(ecm_score,xx,pp,u_threshold,x_min,x_max,lambda_regularisation), opt);
        my_x_max(it) = -my_x_max(it);
      end
    end
  
    c_min.(ecm_score) = exp(my_x_min);
    c_max.(ecm_score) = exp(my_x_max);
  
    %% compute possible variation in enzyme profile u
    %% (at some maximum allowed percentage increase in cost)
    
    my_u     = u.(ecm_score)(:,1);
    my_u_min = my_u;
    my_u_max = my_u;

    if sum(isfinite(my_u)),
      
    for it = 1:length(u.data),
      [my_x, my_u_min(it),exitflag] = fmincon(@(xx) ecm_get_one_u(it,ecm_score,xx,pp,u_threshold,x_min,x_max,ecm_options), x,[],[],[],[],x_min,x_max,@(xx) ecm_below_threshold(ecm_score, xx, pp, u_threshold, x_min, x_max, lambda_regularisation), opt);
  
      [my_x, my_u_max(it),exitflag] = fmincon(@(xx) -ecm_get_one_u(it,ecm_score,xx,pp,u_threshold,x_min,x_max,ecm_options), x,[],[],[],[],x_min,x_max,@(xx) ecm_below_threshold(ecm_score,xx,pp,u_threshold,x_min,x_max,lambda_regularisation), opt);
  
      if ~exitflag, warning(sprintf('Enzyme variability in %s: No valid solution found',network.actions{it})); 
        my_u_max(it) = nan;
      end 
  
      my_u_max(it) = - my_u_max(it);
    end
  
    u_min.(ecm_score) = my_u_min;
    u_max.(ecm_score) = my_u_max;

    end
  end
  
end


% ---------------------------------------------------------------------
% compute efficiency values

kcat_forward      = r.Kcatf;
kcat_forward(v<0) = r.Kcatr(v<0);
u_capacity        = abs(v)./kcat_forward;
eta_energetic     = struct;
eta_saturation    = struct;

for it_method = 1:length(ecm_options.ecm_scores),

  this_ecm_score   = ecm_options.ecm_scores{it_method};
  my_u             = u.(this_ecm_score);

  switch this_ecm_score,
    case {'mdf','emc1'},
      my_eta_energetic = nan * ones(size(v));
      my_eta_energetic(ecm_options.ind_scored_enzymes) = ones(size(ecm_options.ind_scored_enzymes));
    otherwise,
      my_A_forward     = A_forward.(this_ecm_score);
      my_eta_energetic = 1-exp(-my_A_forward/RT);
      my_eta_energetic(my_eta_energetic<0) = 0;
  end
  
  switch this_ecm_score,
    case {'mdf','emc1','emc2s','emc2sp'},
      my_eta_saturation = nan * ones(size(v));
      my_eta_saturation(ecm_options.ind_scored_enzymes) = ones(size(ecm_options.ind_scored_enzymes));
    otherwise,
      my_eta_saturation = [abs(v) ./ kcat_forward ./ my_eta_energetic] ./ my_u;
  end
  
  eta_energetic.(this_ecm_score)  = my_eta_energetic ;
  eta_saturation.(this_ecm_score) = my_eta_saturation;
  
end

% ------------------------------------------------------
% postprocessing in case of multiple conditions: turn result (metabolite and enzyme) vectors into matrices
%  ecm_options.multiple_conditions_n   -> number of conditions
%  ecm_options.multiple_conditions_anticipate ==1 -> find a single enzyme profile for all conditions

if isfield(ecm_options,'multiple_conditions_n'),

  n_cond = ecm_options.multiple_conditions_n;
  
  fn = fieldnames(c); 
  for it = 1:length(fn),
    if length(c.(fn{it})),
      c.(fn{it}) = reshape(c.(fn{it}),nm/n_cond,n_cond);
    end
  end

  fn = fieldnames(u); 
  for it = 1:length(fn),
    if length(u.(fn{it})),
      u.(fn{it}) = reshape(u.(fn{it}),nr/n_cond,n_cond);
    end
  end

  fn = fieldnames(up); 
  for it = 1:length(fn),
    if length(up.(fn{it})),
      up.(fn{it}) = reshape(up.(fn{it}),nr/n_cond,n_cond);
    end
  end

  fn = fieldnames(A_forward); 
  for it = 1:length(fn),
    if length(A_forward.(fn{it})),
      A_forward.(fn{it}) = reshape(A_forward.(fn{it}),nr/n_cond,n_cond);
    end
  end

  %fn = fieldnames(mca_info);
  %for it = 1:length(fn),
  %  mca_info.(fn{it}).gradient = reshape(mca_info.(fn{it}).gradient,nr/n_cond,n_cond);
  %end

  u_capacity = reshape(u_capacity,nr/n_cond,n_cond);
  fn = fieldnames(eta_energetic); 
  for it = 1:length(fn),
    eta_energetic.(fn{it})  = reshape(eta_energetic.(fn{it}),nr/n_cond,n_cond);
    eta_saturation.(fn{it}) = reshape(eta_saturation.(fn{it}),nr/n_cond,n_cond);
  end


end
```
