## Supplementary material for "In-depth computational analysis of natural and artificial carbon fixation pathways": MATLAB files: figureYbreak.m

```
function figureYbreak(yLimMain, nTicksMain, yLimOutliers, xBreakWidth, yBreakTilt, yBreakDist)
f = gcf;
Interval = (yLimMain(2)-yLimMain(1))/(nTicksMain-1);
nTicksOutliers = round((yLimOutliers(2)-yLimOutliers(1))/Interval+0.49+1);
% yLimOutliers(2) = yLimOutliers(1) + nTicksOutliers*Interval;

yTicks = [0; [1:nTicksMain+nTicksOutliers-1]'*Interval];

ylim([yTicks(1) yTicks(end)]);
yticks(yTicks);

yTicks2 = yTicks;
yLimDiff = yTicks(end)-yTicks(1);
yTicks2(nTicksMain+1:end,1) = yTicks(nTicksMain+1:end,1)+(yLimOutliers(1)-yLimMain(end)-Interval);
yticklabels(num2cell(yTicks2));

%yBreakTilt = Interval*yBreakTilt;

    pos = get(gca, 'position');
    n = length(get(gca, 'yticklabel')) - 1;
        % Find the center along the axis

    spacing = pos(4)/n;
    
    % Find the center along the axis
    cent = pos(2) + 0.5*spacing;
    
    % define the extents of the bar
    X1 = [pos(1) - 0.005 ; 
          pos(1) + 0.005];
    X2 = [pos(1) - xBreakWidth/2; 
          pos(1) + xBreakWidth/2];
    Y = [cent - yBreakDist*spacing/2 + spacing*(nTicksMain-1); 
         cent + yBreakDist*spacing/2 + spacing*(nTicksMain-1)];
    
    yBreakTilt = yBreakTilt*spacing;
    % Make the bar and ticks
    h1 = annotation(f, 'rectangle', [X1(1), Y(1), diff(X1), diff(Y)]);
    annotation(f, 'line', [X2(1) X2(2)], [Y(1)-yBreakTilt Y(1)+yBreakTilt]);
    annotation(f, 'line', [X2(1) X2(2)], [Y(2)-yBreakTilt Y(2)+yBreakTilt]);
    h1.FaceColor = 'w';
    h1.Color = 'w';

end
```
