## Supplementary material for "In-depth computational analysis of natural and artificial carbon fixation pathways": MATLAB files: GetPathwayStoichiometry.m

```
function [reaction_names, fluxes] = GetPathwayStoichiometry(pathway_name, product_name)

% in this function, each pathway name is connected to a set of
% stoichiometries that are returen by the function

reaction_names = {};
fluxes = [];

if(strcmp(product_name,'C_oxaloacetate'))
    switch pathway_name
        case {'CETCH','CETCH-rGlyOx'}
            reaction_names = {'R_aspartate__R07410','R_glycine_ox_R00373','R_l_erythro__R00478',...
                'R_malyl_coa__R00934','R_methylmalo_R00833','R_methylmalo_R02765',...
                'R_RXN_Q00071','R_RXN_Q00072','R_RXN_Q00073','R_RXN_Q00084','R_RXN_Q00085',...
                'R_RXN_R05076','R_RXN_R09281','R_RXN_R09291','R_RXN_R09292','R_RXN_R09979','R_succinate__R09280'};
            fluxes = [0.5, 0.5, -0.5, 1, 1, -1, 1, 1, 1, 1, 0.5, -1, -1, -1, 1, 1, -1]'*2;
        case 'rCCC'
            reaction_names = {'R_3_hydroxyb_R01976','R_acetyl_coa_R00238','R_enoyl_coen_R03026',...
                'R_fumarate_h_R01082','R_fumarate_r_R09490','R_malate_deh_R00342',...
                'R_malyl_coa__R00237','R_RXN_Q00071','R_RXN_Q00072','R_RXN_Q00073',...
                'R_RXN_Q00074','R_RXN_R09281','R_RXN_R09282','R_RXN_R09283'...
                'R_RXN_R09291', 'R_RXN_R09292', 'R_RXN_R09979','R_stk_R00405', 'R_succinate__R09280'};
            fluxes = [-1, 1, 1, 1, -1, -1, ...
                2, 1, 1, 2, 2, -1, -2, 2,...
                -2, 2, 2, 1, -1]';
        case 'rCCC(Malyl-CoA/rGlyOx)'
            reaction_names =   {'R_3_hydroxyb_R01976','R_acetyl_coa_R00238','R_enoyl_coen_R03026',...
                'R_malate_deh_R00342','R_malyl_coa__R00237','R_RXN_Q00073',...
                'R_RXN_Q00074','R_RXN_R09282','R_RXN_R09283'...
                'R_RXN_R09291', 'R_RXN_R09292', 'R_RXN_R09979','R_stk_R00405',...
                'R_RXN_R00406', 'R_malyl_coa__R00473',...
                'R_isocitrate_R00479','R_aconitate__R01900','R_aconitate__R01325',... % from this line: rGlyoxylate-Shunt
                'R_atp_citrat_R00352','R_fumarate_h_R01082','R_fumarate_r_R09490'};
            fluxes = [-1, 1, 1, -1, 1, 1, 1, -1, 1,...
                -1, 1, 1, 0.5, 0.5, 0.5,...
                0.5, 0.5, -0.5, 0.5, 0.5, -0.5]'*2;
        case 'HP-Bicycle'
            reaction_names =   {'R_RXN_Q00091','R_RXN_Q00092','R_RXN_Q00093',...
                    'R_RXN_Q00084', 'R_methylmalo_R02765','R_methylmalo_R00833',...
                    'R_RXN_R00406', 'R_fumarate_r_R09490','R_fumarate_h_R01082',...
                    'R_malyl_coa__R00473', 'R_malyl_coa__R00934', 'R_RXN_R05076',...
                    'R_RXN_R09283','R_RXN_R09282','R_malyl_coa__R00237','R_RXN_Q00074'};
            fluxes = [1 -1 1 0.5 -0.5 0.5 0.5 0.5 -0.5 0.5 -0.5 0.5 0.5 -0.5 0.5 0.5]'*2;
        case 'HP-Bicycle(ADP)'
            reaction_names =   {'R_RXN_Q00091','R_RXN_Q00092','R_3_hydroxyi_R03157','R_RXN_Q00094',...
                    'R_RXN_Q00084', 'R_methylmalo_R02765','R_methylmalo_R00833',...
                    'R_RXN_R00406', 'R_fumarate_r_R09490','R_fumarate_h_R01082',...
                    'R_malyl_coa__R00473', 'R_malyl_coa__R00934', 'R_RXN_R05076',...
                    'R_RXN_R09283','R_RXN_R09282','R_malyl_coa__R00237','R_RXN_Q00074'};
            fluxes = [1 -1 1 -1 0.5 -0.5 0.5 0.5 0.5 -0.5 0.5 -0.5 0.5 0.5 -0.5 0.5 0.5]'*2;
        case 'CBB'
            reaction_names =   {'R_ribulose_b_R00024_2','R_phosphogly_R01512','R_glyceralde_R01063',...
                    'R_triose_pho_R01015','R_fructose_b_R01068','R_fructose_b_R00762',...
                    'R_transketol_R01067','R_fructose_b_R01829','R_sedoheptul_R01845',...
                    'R_transketol_R01641','R_ribulose_p_R01529','R_ribose_5_p_R01056',...
                    'R_phosphorib_R01523','R_RXN_Q00000','R_phosphopyr_R00658',...
                    'R_phosphogly_R01518'};
            fluxes = [-3, 5, -5, 2, -1, 1, 1, -1, 1, 1, -2, 1, 3, -1, 1, -1]';
        case 'rCCC(Malyl-CoA)'
            reaction_names = {'R_3_hydroxyb_R01976','R_acetyl_coa_R00238','R_enoyl_coen_R03026',...
                'R_malate_deh_R00342','R_malyl_coa__R00237','R_RXN_Q00073',...
                'R_RXN_Q00074','R_RXN_R09282','R_RXN_R09283'...
                'R_RXN_R09291', 'R_RXN_R09292', 'R_RXN_R09979','R_stk_R00405',...
                'R_RXN_R00406', 'R_malyl_coa__R00473',...
                'R_glycine_ox_R00373','R_l_erythro__R00478','R_RXN_Q00085','R_aspartate__R07410'}; % Iminoaspartate-Cycle
            fluxes = [-1, 1, 1, -1, 1, 1, 1, -1, 1,...
                -1, 1, 1, 1, 1, 1,...
                0.5, -0.5, 0.5, 0.5]'*2;
        case 'rCCC(Transcarb)'
            reaction_names = {'R_3_hydroxyb_R01976','R_acetyl_coa_R00238','R_enoyl_coen_R03026',...
                'R_fumarate_h_R01082','R_fumarate_r_R09490','R_malate_deh_R00342',...
                'R_malyl_coa__R00237','R_RXN_Q00071','R_RXN_Q00072','R_RXN_Q00073',...
                'R_methylmalo_R00353','R_RXN_R09281','R_RXN_R09282','R_RXN_R09283'...
                'R_RXN_R09291', 'R_RXN_R09292', 'R_RXN_R09979','R_stk_R00405',...
                'R_succinate__R09280','R_RXN_Q00091'};
            fluxes = [-1, 1, 1, 1, -1, -1, ...
                2, 1, 1, 2, 2, -1, -2, 2,...
                -2, 2, 2, 1, -1 2]';
        case 'rCCC(PEP-Synth)'
            reaction_names = {'R_3_hydroxyb_R01976','R_acetyl_coa_R00238','R_enoyl_coen_R03026',...
                'R_fumarate_h_R01082','R_fumarate_r_R09490','R_malate_deh_R00342',...
                'R_malyl_coa__R00237','R_RXN_Q00071','R_RXN_Q00072','R_RXN_Q00073',...
                'R_pyruvate_w_R00199','R_RXN_R09281','R_RXN_R09282','R_RXN_R09283'...
                'R_RXN_R09291', 'R_RXN_R09292', 'R_RXN_R09979','R_stk_R00405',...
                'R_succinate__R09280','R_RXN_Q00000'};
            fluxes = [-1, 1, 1, 1, -1, -1, ...
                2, 1, 1, 2, 2, -1, -2, 2,...
                -2, 2, 2, 1, -1 -2]';
        case 'CBB(PTS)'
            reaction_names =   {'R_ribulose_b_R00024','R_phosphogly_R01512','R_glyceralde_R01063',...
                    'R_triose_pho_R01015','R_fructose_b_R01068','R_diphosphat_R02073',...
                    'R_transketol_R01067','R_fructose_b_R01829','R_RXN_P00011',...
                    'R_transketol_R01641','R_ribulose_p_R01529','R_ribose_5_p_R01056',...
                    'R_phosphorib_R01523','R_RXN_Q00000','R_phosphopyr_R00658',...
                    'R_phosphogly_R01518','R_RXN_P00010'};
            fluxes = [-3, 5, -5, 2, -1, 1, 1, -1, 1, 1, -2, 1, 1, -1, 1, -1, -2]';
        case 'RuMP-Cycle'
            reaction_names =   {'R_methanol_d_R00605','R_3_hexulose_R05338','R_6_phospho__R05339',...
                    'R_6_phosphof_R00756','R_transketol_R01067','R_fructose_b_R01068',...
                    'R_triose_pho_R01015','R_fructose_b_R01829','R_sedoheptul_R01845',...
                    'R_transketol_R01641','R_ribulose_p_R01529','R_ribose_5_p_R01056',...
                    'R_glyceralde_R01061','R_phosphogly_R01512','R_phosphogly_R01518',...
                    'R_phosphopyr_R00658','R_RXN_Q00000'};

            fluxes = [3 -3 3 2 1 2 -1 -1 1 1 -2 1 1 -1 -1 1 -1]';
        case 'RuMP-Cycle(Formate)'
            reaction_names =   {'R_RXN_Q00004','R_3_hexulose_R05338','R_6_phospho__R05339',...
                    'R_6_phosphof_R00756','R_transketol_R01067','R_fructose_b_R01068',...
                    'R_triose_pho_R01015','R_fructose_b_R01829','R_sedoheptul_R01845',...
                    'R_transketol_R01641','R_ribulose_p_R01529','R_ribose_5_p_R01056',...
                    'R_glyceralde_R01061','R_phosphogly_R01512','R_phosphogly_R01518',...
                    'R_phosphopyr_R00658','R_RXN_Q00000','R_RXN_Q00003','R_stk_R00405'};

            fluxes = [-3 -3 3 2 1 2 -1 -1 1 1 -2 1 1 -1 -1 1 -1 -3 3]';
        case 'RuMP-Cycle(Formate/THF)'
            reaction_names =   {'R_RXN_Q00005','R_3_hexulose_R05338','R_6_phospho__R05339',...
                    'R_6_phosphof_R00756','R_transketol_R01067','R_fructose_b_R01068',...
                    'R_triose_pho_R01015','R_fructose_b_R01829','R_sedoheptul_R01845',...
                    'R_transketol_R01641','R_ribulose_p_R01529','R_ribose_5_p_R01056',...
                    'R_glyceralde_R01061','R_phosphogly_R01512','R_phosphogly_R01518',...
                    'R_phosphopyr_R00658','R_RXN_Q00000','R_glycine_hy_R00945',...
                    'R_methylenet_R01220','R_methenylte_R01655','R_formate_te_R00943'};
            fluxes = [3 -3 3 2 1 2 -1 -1 1 1 -2 1 1 -1 -1 1 -1 3 -3 -3 3]';
        case {'Serine-Cycle','Serine-Cycle (w/o SHMT)'}
            reaction_names =   {'R_formate_te_R00943','R_methenylte_R01655','R_methylenet_R01220',...
                    'R_glycine_hy_R00945','R_serine_gly_R00588','R_glyoxylate_R01388',...
                    'R_glycerate__R08572','R_phosphopyr_R00658','R_RXN_Q00000',...
                    'R_malate_deh_R00342','R_RXN_R00406','R_malyl_coa__R00473',...
                    'R_stk_R00405','R_acetyl_coa_R00238','R_3_hydroxyb_R01976',...
                    'R_enoyl_coen_R03026','R_RXN_R09291','R_RXN_R09979',...
                    'R_RXN_R09292','R_RXN_Q00073','R_RXN_R05076','R_malyl_coa__R00934',...
                    'R_RXN_Q00084','R_methylmalo_R02765','R_methylmalo_R00833',...
                    'R_fumarate_r_R09490','R_fumarate_h_R01082'};
            fluxes = [1 -1 -1 3 3 -3 3 3 -3 -1 2 2 1 1 -1,...
                1 -1 1 1 1 -1 1,...
                1 -1 1 1 -1]'/2;
        case {'rGlyP','rGlyP (w/o SHMT)'}
            reaction_names = {'R_formate_te_R00943','R_methenylte_R01655','R_methylenet_R01220',...
                'R_RXN_Q00006','R_RXN_Q00007','R_diaphorase_R01698','R_glycine_hy_R00945',...
                'R_l_serine_d_R00220','R_RXN_Q00074'};
            fluxes = [2 -2 -2 -1 -1 1 1 1 1]';
        case '2-HG-rTCA'
            reaction_names =   {'R_RXN_Q00009','R_RXN_Q00010','R_RXN_Q00004','R_RXN_Q00011',...
                    'R_idp_R00267','R_aconitate__R01900','R_aconitate__R01325',...
                    'R_atp_citrat_R00352','R_fumarate_h_R01082','R_fumarate_r_R09490',...
                    'R_stk_R00405','R_succinate__R09280',...
                    'R_acetyl_coa_R00238','R_3_hydroxyb_R01976','R_enoyl_coen_R03026',... % from this line, the Acetyl-CoA -> Oxaloacetate module starts
                    'R_RXN_R09291','R_RXN_R09979','R_RXN_R09292','R_RXN_Q00073',...
                    'R_RXN_R09283','R_RXN_R09282','R_malyl_coa__R00237','R_RXN_Q00074'};
            fluxes = [1 1 -1 1 -1 1 -1 1 1 -1 1 -1,...
                1 -1 1 -1 1 1 1 1 -1 1 1]';
        case '2-HACL/Mal'
            reaction_names =   {'R_RXN_Q00091','R_malonyl_co_R00740','R_RXN_Q00008',...
                    'R_RXN_Q00004','R_stk_R00405','R_malyl_coa__R00473',...
                    'R_glycine_ox_R00373','R_aspartate__R07410','R_RXN_Q00085',...
                    'R_l_erythro__R00478'};
            fluxes = [1 1 1 -1 1 1 0.5 0.5 0.5 -0.5]'*2;
        case '2-HACL/Ser'
            reaction_names =   {'R_RXN_Q00091','R_malonyl_co_R00740','R_RXN_Q00008',...
                    'R_RXN_Q00004','R_stk_R00405','R_malyl_coa__R00473',...
                    'R_serine_gly_R00588','R_glyoxylate_R01388','R_glycine_hy_R00945',...
                    'R_methylenet_R01220','R_methenylte_R01655','R_formate_te_R00943',...
                    'R_glycerate__R08572','R_phosphopyr_R00658','R_RXN_Q00000'};
            fluxes = [1 1 1 -1 1 1 1 -1 1 -1 -1 1 1 1 -1]';
        case 'HB/HP-Cren'
            reaction_names = {'R_RXN_Q00084','R_methylmalo_R02765','R_methylmalo_R00833',...
                'R_succinate__R09280','R_RXN_R09281','R_RXN_R09279','R_RXN_Q00072',...
                'R_3_hydroxya_R01975','R_acetyl_coa_R00238','R_RXN_Q00091','R_RXN_Q00092',...
                'R_RXN_Q00093','R_enoyl_coen_R03026','R_stk_R00405','R_fumarate_r_R09490',...
                'R_fumarate_h_R01082','R_malate_deh_R00216','R_RXN_Q00074'};
            fluxes = [2 -2 2 -1 -1 1 1 1 -1 2 -2 2 -1 -1 1 -1 1 1]';
        case 'HB/HP-Thaum'
            reaction_names = {'R_RXN_Q00084','R_methylmalo_R02765','R_methylmalo_R00833',...
                'R_succinate__R09280','R_RXN_R09281','R_RXN_Q00071','R_RXN_Q00072',...
                'R_3_hydroxya_R01975','R_acetyl_coa_R00238','R_RXN_Q00091','R_RXN_Q00092',...
                'R_3_hydroxyi_R03157', 'R_RXN_Q00094','R_enoyl_coen_R03026','R_stk_R00405','R_fumarate_r_R09490',...
                'R_fumarate_h_R01082','R_malate_deh_R00216','R_RXN_Q00074'};
            fluxes = [2 -2 2 -1 -1 1 1 1 -1 2 -2 2 -2 -1 -1 1 -1 1 1]';
    end
end

if(strcmp(product_name,'C_acetyl_coa'))
    switch pathway_name
        case 'HB/HP-Cren'
            reaction_names = {'R_RXN_Q00084','R_methylmalo_R02765','R_methylmalo_R00833',...
                'R_succinate__R09280','R_RXN_R09281','R_RXN_R09279','R_RXN_Q00072',...
                'R_3_hydroxya_R01975','R_acetyl_coa_R00238','R_RXN_Q00091','R_RXN_Q00092',...
                'R_RXN_Q00093','R_enoyl_coen_R03026'};
            fluxes = [1 -1 1 -1 -1 1 1 1 -1 1 -1 1 -1]';
        case 'HB/HP-Thaum'
            reaction_names = {'R_RXN_Q00084','R_methylmalo_R02765','R_methylmalo_R00833',...
                'R_succinate__R09280','R_RXN_R09281','R_RXN_Q00071','R_RXN_Q00072',...
                'R_3_hydroxya_R01975','R_acetyl_coa_R00238','R_RXN_Q00091','R_RXN_Q00092',...
                'R_3_hydroxyi_R03157', 'R_RXN_Q00094','R_enoyl_coen_R03026'};
            fluxes = [1 -1 1 -1 -1 1 1 1 -1 1 -1 1 -1 -1]';
        case '2-HG-rTCA'
            reaction_names =   {'R_RXN_Q00009','R_RXN_Q00010','R_RXN_Q00004','R_RXN_Q00011',...
                    'R_idp_R00267','R_aconitate__R01900','R_aconitate__R01325',...
                    'R_atp_citrat_R00352','R_fumarate_h_R01082','R_fumarate_r_R09490',...
                    'R_stk_R00405','R_succinate__R09280'};
            fluxes = [1 1 -1 1 -1 1 -1 1 1 -1 1 -1];
        case {'rGlyP','rGlyP (w/o SHMT)'}
            reaction_names = {'R_formate_te_R00943','R_methenylte_R01655','R_methylenet_R01220',...
                'R_RXN_Q00006','R_RXN_Q00007','R_diaphorase_R01698','R_glycine_hy_R00945',...
                'R_l_serine_d_R00220','R_diaphorase_R00209'};
            fluxes = [2 -2 -2 -1 -1 1 1 1 1]';
        case {'2-HACL/Mal','2-HACL/Ser'}
            reaction_names =   {'R_RXN_Q00091','R_malonyl_co_R00740','R_RXN_Q00008',...
                    'R_RXN_Q00004','R_stk_R00405','R_malyl_coa__R00473',...
                    'R_glycine_ox_R00373','R_aspartate__R07410','R_RXN_Q00085',...
                    'R_l_erythro__R00478','R_malate_deh_R00342','R_RXN_R00406'};
            fluxes = [1 1 1 -1 2 2 1 1 1 -1 -1 1]';
        case {'Serine-Cycle','Serine-Cycle (w/o SHMT)'}
            reaction_names =   {'R_formate_te_R00943','R_methenylte_R01655','R_methylenet_R01220',...
                    'R_glycine_hy_R00945','R_serine_gly_R00588','R_glyoxylate_R01388',...
                    'R_glycerate__R08572','R_phosphopyr_R00658','R_RXN_Q00000',...
                    'R_malate_deh_R00342','R_RXN_R00406','R_malyl_coa__R00473',...
                    'R_stk_R00405'};
            fluxes = [1 -1 -1 1 1 -1 1 1 -1 -1 1 1 1]';
        case 'RuMP-Cycle(Formate/THF)'
            reaction_names =   {'R_RXN_Q00005','R_3_hexulose_R05338','R_6_phospho__R05339',...
                    'R_6_phosphof_R00756','R_transketol_R01067','R_fructose_b_R01068',...
                    'R_triose_pho_R01015','R_fructose_b_R01829','R_sedoheptul_R01845',...
                    'R_transketol_R01641','R_ribulose_p_R01529','R_ribose_5_p_R01056',...
                    'R_glyceralde_R01061','R_phosphogly_R01512','R_phosphogly_R01518',...
                    'R_phosphopyr_R00658','R_RXN_Q00000','R_glycine_hy_R00945',...
                    'R_methylenet_R01220','R_methenylte_R01655','R_formate_te_R00943',...
                    'R_malate_deh_R00342','R_RXN_R00406','R_malyl_coa__R00473',...
                    'R_stk_R00405','R_glycine_ox_R00373','R_l_erythro__R00478',...
                    'R_RXN_Q00085','R_aspartate__R07410'};
            fluxes =    [3 -3 3 2 1 2 -1 -1 1 1 -2 1 1 -1 -1 1 -1 3 -3 -3 3,...
                        -2 2 2 2 1 -1 1 1]'/2;
        case 'RuMP-Cycle'
            reaction_names =   {'R_methanol_d_R00605','R_3_hexulose_R05338','R_6_phospho__R05339',...
                    'R_6_phosphof_R00756','R_transketol_R01067','R_fructose_b_R01068',...
                    'R_triose_pho_R01015','R_fructose_b_R01829','R_sedoheptul_R01845',...
                    'R_transketol_R01641','R_ribulose_p_R01529','R_ribose_5_p_R01056',...
                    'R_glyceralde_R01061','R_phosphogly_R01512','R_phosphogly_R01518',...
                    'R_phosphopyr_R00658','R_RXN_Q00000',...
                    'R_malate_deh_R00342','R_RXN_R00406','R_malyl_coa__R00473',...
                    'R_stk_R00405','R_glycine_ox_R00373','R_l_erythro__R00478',...
                    'R_RXN_Q00085','R_aspartate__R07410'};
            fluxes =    [3 -3 3 2 1 2 -1 -1 1 1 -2 1 1 -1 -1 1 -1,...
                        -2 2 2 2 1 -1 1 1]'/2;
        case 'RuMP-Cycle(Formate)'
            reaction_names =   {'R_RXN_Q00004','R_3_hexulose_R05338','R_6_phospho__R05339',...
                    'R_6_phosphof_R00756','R_transketol_R01067','R_fructose_b_R01068',...
                    'R_triose_pho_R01015','R_fructose_b_R01829','R_sedoheptul_R01845',...
                    'R_transketol_R01641','R_ribulose_p_R01529','R_ribose_5_p_R01056',...
                    'R_glyceralde_R01061','R_phosphogly_R01512','R_phosphogly_R01518',...
                    'R_phosphopyr_R00658','R_RXN_Q00000','R_RXN_Q00003','R_stk_R00405',...
                    'R_malate_deh_R00342','R_RXN_R00406','R_malyl_coa__R00473',...
                    'R_stk_R00405','R_glycine_ox_R00373','R_l_erythro__R00478',...
                    'R_RXN_Q00085','R_aspartate__R07410'};
            fluxes =    [-3 -3 3 2 1 2 -1 -1 1 1 -2 1 1 -1 -1 1 -1 -3 3,...
                        -2 2 2 2 1 -1 1 1]'/2;
        case 'CBB(PTS)'
            reaction_names =   {'R_ribulose_b_R00024','R_phosphogly_R01512','R_glyceralde_R01063',...
                    'R_triose_pho_R01015','R_fructose_b_R01068','R_diphosphat_R02073',...
                    'R_transketol_R01067','R_fructose_b_R01829','R_RXN_P00011',...
                    'R_transketol_R01641','R_ribulose_p_R01529','R_ribose_5_p_R01056',...
                    'R_phosphorib_R01523','R_RXN_Q00000','R_phosphopyr_R00658',...
                    'R_phosphogly_R01518','R_RXN_P00010',...
                    'R_malate_deh_R00342','R_RXN_R00406','R_malyl_coa__R00473',...
                    'R_stk_R00405','R_glycine_ox_R00373','R_l_erythro__R00478',...
                    'R_RXN_Q00085','R_aspartate__R07410'};
            fluxes =    [-3, 5, -5, 2, -1, 1, 1, -1, 1, 1, -2, 1, 1, -1, 1, -1, -2,...
                        -2 2 2 2 1 -1 1 1]'/2;
        case 'CBB'
            reaction_names =   {'R_ribulose_b_R00024','R_phosphogly_R01512','R_glyceralde_R01063',...
                    'R_triose_pho_R01015','R_fructose_b_R01068','R_fructose_b_R00762',...
                    'R_transketol_R01067','R_fructose_b_R01829','R_sedoheptul_R01845',...
                    'R_transketol_R01641','R_ribulose_p_R01529','R_ribose_5_p_R01056',...
                    'R_phosphorib_R01523','R_RXN_Q00000','R_phosphopyr_R00658',...
                    'R_phosphogly_R01518',...
                    'R_malate_deh_R00342','R_RXN_R00406','R_malyl_coa__R00473',...
                    'R_stk_R00405','R_glycine_ox_R00373','R_l_erythro__R00478',...
                    'R_RXN_Q00085','R_aspartate__R07410'};
            fluxes =    [-3, 5, -5, 2, -1, 1, 1, -1, 1, 1, -2, 1, 3, -1, 1, -1,...
                        -2 2 2 2 1 -1 1 1]'/2;
        case 'rCCC(PEP-Synth)'
            reaction_names = {'R_fumarate_h_R01082','R_fumarate_r_R09490','R_malate_deh_R00342',...
                'R_malyl_coa__R00237','R_RXN_Q00071','R_RXN_Q00072','R_RXN_Q00073',...
                'R_pyruvate_w_R00199','R_RXN_R09281','R_RXN_R09282','R_RXN_R09283'...
                'R_RXN_R09291', 'R_RXN_R09292', 'R_RXN_R09979','R_stk_R00405',...
                'R_succinate__R09280','R_RXN_Q00000'};
            fluxes = [1, -1, -1, ...
                1, 1, 1, 1, 1, -1, -1, 1,...
                -1, 1, 1, 1, -1 -1]';
        case 'rCCC(Malyl-CoA)'
            reaction_names = {'R_3_hydroxyb_R01976','R_acetyl_coa_R00238','R_enoyl_coen_R03026',...
                'R_malate_deh_R00342','R_malyl_coa__R00237','R_RXN_Q00073',...
                'R_RXN_Q00074','R_RXN_R09282','R_RXN_R09283'...
                'R_RXN_R09291', 'R_RXN_R09292', 'R_RXN_R09979','R_stk_R00405',...
                'R_RXN_R00406', 'R_malyl_coa__R00473',...
                'R_glycine_ox_R00373','R_l_erythro__R00478','R_RXN_Q00085','R_aspartate__R07410'}; % β-hydroxyaspartate cycle
            fluxes = [-1, 1, 1, -2, 1, 1, 1, -1, 1,...
                -1, 1, 1, 2, 2, 2,...
                1, -1, 1, 1]';
        case 'rCCC(Malyl-CoA/rGlyOx)'
            reaction_names = {'R_3_hydroxyb_R01976','R_acetyl_coa_R00238','R_enoyl_coen_R03026',...
                'R_malate_deh_R00342','R_malyl_coa__R00237','R_RXN_Q00073',...
                'R_RXN_Q00074','R_RXN_R09282','R_RXN_R09283'...
                'R_RXN_R09291', 'R_RXN_R09292', 'R_RXN_R09979','R_stk_R00405',...
                'R_RXN_R00406', 'R_malyl_coa__R00473',...
                'R_isocitrate_R00479','R_aconitate__R01900','R_aconitate__R01325',... % from this line: rGlyoxylate-Shunt
                'R_atp_citrat_R00352','R_fumarate_h_R01082','R_fumarate_r_R09490'}; 
            fluxes = [-1, 1, 1, -2, 1, 1, 1, -1, 1,...
                -1, 1, 1, 1, 1, 1,...
                1, 1, -1, 1, 1, -1]'; %from this line: reverse glyoxylate shunt
        case 'rCCC(Transcarb)'
            reaction_names = {'R_fumarate_h_R01082','R_fumarate_r_R09490','R_malate_deh_R00342',...
                'R_malyl_coa__R00237','R_RXN_Q00071','R_RXN_Q00072','R_RXN_Q00073',...
                'R_methylmalo_R00353','R_RXN_R09281','R_RXN_R09282','R_RXN_R09283'...
                'R_RXN_R09291', 'R_RXN_R09292', 'R_RXN_R09979','R_stk_R00405',...
                'R_succinate__R09280','R_RXN_Q00091'};
            fluxes = [1, -1, -1, ...
                1, 1, 1, 1, 1, -1, -1, 1,...
                -1, 1, 1, 1, -1 1]';
        case 'rCCC'
            reaction_names = {'R_fumarate_h_R01082','R_fumarate_r_R09490','R_malate_deh_R00342',...
                'R_malyl_coa__R00237','R_RXN_Q00071','R_RXN_Q00072','R_RXN_Q00073',...
                'R_RXN_Q00074','R_RXN_R09281','R_RXN_R09282','R_RXN_R09283'...
                'R_RXN_R09291', 'R_RXN_R09292', 'R_RXN_R09979','R_stk_R00405', 'R_succinate__R09280'};
            fluxes = [1, -1, -1, ...
                1, 1, 1, 1, 1, -1, -1, 1,...
                -1, 1, 1, 1, -1]';
        case 'CETCH'
            reaction_names = {'R_aspartate__R07410','R_glycine_ox_R00373','R_l_erythro__R00478',...
                'R_malyl_coa__R00934','R_methylmalo_R00833','R_methylmalo_R02765',...
                'R_RXN_Q00071','R_RXN_Q00072','R_RXN_Q00073','R_RXN_Q00084','R_RXN_Q00085',...
                'R_RXN_R05076','R_RXN_R09281','R_RXN_R09291','R_RXN_R09292','R_RXN_R09979','R_succinate__R09280',...
                'R_malate_deh_R00342','R_RXN_R00406','R_malyl_coa__R00473',...
                'R_stk_R00405'};
            fluxes =    [1, 1, -1, 1, 1, -1, 1, 1, 1, 1, 1, -1, -1, -1, 1, 1, -1,...
                        -1 1 1 1]';
        case 'CETCH-rGlyOx'
            reaction_names = {'R_malyl_coa__R00934','R_methylmalo_R00833','R_methylmalo_R02765',...
                'R_RXN_Q00071','R_RXN_Q00072','R_RXN_Q00073','R_RXN_Q00084',...
                'R_RXN_R05076','R_RXN_R09281','R_RXN_R09291','R_RXN_R09292','R_RXN_R09979','R_succinate__R09280',...
                'R_isocitrate_R00479','R_aconitate__R01900','R_aconitate__R01325',... % from this line: rGlyoxylate-Shunt
                'R_atp_citrat_R00352','R_malate_deh_R00342','R_fumarate_h_R01082','R_fumarate_r_R09490'}; 
            fluxes = [1, 1, -1, 1, 1, 1, 1, -1, -1, -1, 1, 1, -1,...
                     1, 1, -1, 1, -1, 1, -1]'; %from this line: reverse glyoxylate shunt
        case 'HP-Bicycle'
            reaction_names =   {'R_RXN_Q00091','R_RXN_Q00092','R_RXN_Q00093',...
                    'R_RXN_R00406','R_stk_R00405','R_RXN_Q00074','R_malate_deh_R00342',...
                    'R_malyl_coa__R00473', 'R_malyl_coa__R00934', 'R_RXN_R05076',...
                    'R_RXN_R09283','R_RXN_R09282','R_malyl_coa__R00237','R_RXN_Q00074'};
            fluxes = [0.5 -0.5 0.5 0.5 0.5 0.5 -0.5 0.5 -0.5 0.5 0.5 -0.5 0.5 0.5]'*2;
        case 'HP-Bicycle(ADP)'
            reaction_names =   {'R_RXN_Q00091','R_RXN_Q00092','R_3_hydroxyi_R03157','R_RXN_Q00094',...
                    'R_RXN_R00406','R_stk_R00405','R_RXN_Q00074','R_malate_deh_R00342',...
                    'R_malyl_coa__R00473', 'R_malyl_coa__R00934', 'R_RXN_R05076',...
                    'R_RXN_R09283','R_RXN_R09282','R_malyl_coa__R00237','R_RXN_Q00074'};
            fluxes = [0.5 -0.5 0.5 -0.5 0.5 0.5 0.5 -0.5 0.5 -0.5 0.5 0.5 -0.5 0.5 0.5]'*2;
    end
end

if(strcmp(product_name,'none'))
    switch pathway_name
% Acetyl-CoA generating pathways:
        case 'HB/HP-Cren'
            reaction_names = {'R_RXN_Q00084','R_methylmalo_R02765','R_methylmalo_R00833',...
                'R_succinate__R09280','R_RXN_R09281','R_RXN_R09279','R_RXN_Q00072',...
                'R_3_hydroxya_R01975','R_acetyl_coa_R00238','R_RXN_Q00091','R_RXN_Q00092',...
                'R_RXN_Q00093','R_enoyl_coen_R03026'};
            fluxes = [1 -1 1 -1 -1 1 1 1 -1 1 -1 1 -1]';
        case 'HB/HP-Thaum'
            reaction_names = {'R_RXN_Q00084','R_methylmalo_R02765','R_methylmalo_R00833',...
                'R_succinate__R09280','R_RXN_R09281','R_RXN_Q00071','R_RXN_Q00072',...
                'R_3_hydroxya_R01975','R_acetyl_coa_R00238','R_RXN_Q00091','R_RXN_Q00092',...
                'R_3_hydroxyi_R03157', 'R_RXN_Q00094','R_enoyl_coen_R03026'};
            fluxes = [1 -1 1 -1 -1 1 1 1 -1 1 -1 1 -1 -1]';
        case '2-HG-rTCA'
            reaction_names =   {'R_RXN_Q00009','R_RXN_Q00010','R_RXN_Q00004','R_RXN_Q00011',...
                    'R_idp_R00267','R_aconitate__R01900','R_aconitate__R01325',...
                    'R_atp_citrat_R00352','R_fumarate_h_R01082','R_fumarate_r_R09490',...
                    'R_stk_R00405','R_succinate__R09280'};
            fluxes = [1 1 -1 1 -1 1 -1 1 1 -1 1 -1]';
        case 'rCCC(PEP-Synth)'
            reaction_names = {'R_fumarate_h_R01082','R_fumarate_r_R09490','R_malate_deh_R00342',...
                'R_malyl_coa__R00237','R_RXN_Q00071','R_RXN_Q00072','R_RXN_Q00073',...
                'R_pyruvate_w_R00199','R_RXN_R09281','R_RXN_R09282','R_RXN_R09283'...
                'R_RXN_R09291', 'R_RXN_R09292', 'R_RXN_R09979','R_stk_R00405',...
                'R_succinate__R09280','R_RXN_Q00000'};
            fluxes = [1, -1, -1, ...
                1, 1, 1, 1, 1, -1, -1, 1,...
                -1, 1, 1, 1, -1 -1]';
        case 'rCCC(Transcarb)'
            reaction_names = {'R_fumarate_h_R01082','R_fumarate_r_R09490','R_malate_deh_R00342',...
                'R_malyl_coa__R00237','R_RXN_Q00071','R_RXN_Q00072','R_RXN_Q00073',...
                'R_methylmalo_R00353','R_RXN_R09281','R_RXN_R09282','R_RXN_R09283'...
                'R_RXN_R09291', 'R_RXN_R09292', 'R_RXN_R09979','R_stk_R00405',...
                'R_succinate__R09280','R_RXN_Q00091'};
            fluxes = [1, -1, -1, ...
                1, 1, 1, 1, 1, -1, -1, 1,...
                -1, 1, 1, 1, -1 1]';
        case 'rCCC'
            reaction_names = {'R_fumarate_h_R01082','R_fumarate_r_R09490','R_malate_deh_R00342',...
                'R_malyl_coa__R00237','R_RXN_Q00071','R_RXN_Q00072','R_RXN_Q00073',...
                'R_RXN_Q00074','R_RXN_R09281','R_RXN_R09282','R_RXN_R09283'...
                'R_RXN_R09291', 'R_RXN_R09292', 'R_RXN_R09979','R_stk_R00405', 'R_succinate__R09280'};
            fluxes = [1, -1, -1, ...
                1, 1, 1, 1, 1, -1, -1, 1,...
                -1, 1, 1, 1, -1]';
        case {'Serine-Cycle','Serine-Cycle (w/o SHMT)'}
            reaction_names =   {'R_formate_te_R00943','R_methenylte_R01655','R_methylenet_R01220',...
                    'R_glycine_hy_R00945','R_serine_gly_R00588','R_glyoxylate_R01388',...
                    'R_glycerate__R08572','R_phosphopyr_R00658','R_RXN_Q00000',...
                    'R_malate_deh_R00342','R_RXN_R00406','R_malyl_coa__R00473',...
                    'R_stk_R00405'};
            fluxes = [1 -1 -1 1 1 -1 1 1 -1 -1 1 1 1]';
% Glyoxylate-generating pathways:
        case 'rCCC(Malyl-CoA)' % turns it into a glyoxylate-generating pathway
            reaction_names = {'R_3_hydroxyb_R01976','R_acetyl_coa_R00238','R_enoyl_coen_R03026',...
                'R_malate_deh_R00342','R_malyl_coa__R00237','R_RXN_Q00073',...
                'R_RXN_Q00074','R_RXN_R09282','R_RXN_R09283'...
                'R_RXN_R09291', 'R_RXN_R09292', 'R_RXN_R09979','R_stk_R00405',...
                'R_RXN_R00406', 'R_malyl_coa__R00473'};
            fluxes = [-1, 1, 1, -1, 1, 1, 1, -1, 1,...
                -1, 1, 1, 1, 1, 1]';
        case 'HP-Bicycle(glyox)'
            reaction_names =   {'R_RXN_Q00091','R_RXN_Q00092','R_RXN_Q00093',...
                    'R_RXN_Q00084', 'R_methylmalo_R02765','R_methylmalo_R00833',...
                    'R_RXN_R00406', 'R_fumarate_r_R09490','R_fumarate_h_R01082',...
                    'R_malyl_coa__R00473'};
            fluxes = [0.5 -0.5 0.5 0.5 -0.5 0.5 0.5 0.5 -0.5 0.5]'*2;
        case 'HP-Bicycle(glyox)(ADP)'
            reaction_names =   {'R_RXN_Q00091','R_RXN_Q00092','R_3_hydroxyi_R03157','R_RXN_Q00094',...
                    'R_RXN_Q00084', 'R_methylmalo_R02765','R_methylmalo_R00833',...
                    'R_RXN_R00406', 'R_fumarate_r_R09490','R_fumarate_h_R01082',...
                    'R_malyl_coa__R00473'};
            fluxes = [0.5 -0.5 0.5 -0.5 0.5 -0.5 0.5 0.5 0.5 -0.5 0.5]'*2;
        case 'CETCH'
            reaction_names = {'R_malyl_coa__R00934','R_methylmalo_R00833','R_methylmalo_R02765',...
                'R_RXN_Q00071','R_RXN_Q00072','R_RXN_Q00073','R_RXN_Q00084',...
                'R_RXN_R05076','R_RXN_R09281','R_RXN_R09291','R_RXN_R09292','R_RXN_R09979','R_succinate__R09280'};
            fluxes = [1, 1, -1, 1, 1, 1, 1, -1, -1, -1, 1, 1, -1]';
% Pyruvate-generating pathways:
        case 'HP-Bicycle'
            reaction_names =   {'R_RXN_Q00091','R_RXN_Q00092','R_RXN_Q00093',...
                    'R_RXN_Q00084', 'R_methylmalo_R02765','R_methylmalo_R00833',...
                    'R_RXN_R00406', 'R_fumarate_r_R09490','R_fumarate_h_R01082',...
                    'R_malyl_coa__R00473', 'R_malyl_coa__R00934', 'R_RXN_R05076',...
                    'R_RXN_R09283','R_RXN_R09282','R_malyl_coa__R00237'};
            fluxes = [1 -1 1 0.5 -0.5 0.5 0.5 0.5 -0.5 0.5 -0.5 0.5 0.5 -0.5 0.5]'*2;
        case 'HP-Bicycle(ADP)'
            reaction_names =   {'R_RXN_Q00091','R_RXN_Q00092','R_3_hydroxyi_R03157','R_RXN_Q00094',...
                    'R_RXN_Q00084', 'R_methylmalo_R02765','R_methylmalo_R00833',...
                    'R_RXN_R00406', 'R_fumarate_r_R09490','R_fumarate_h_R01082',...
                    'R_malyl_coa__R00473', 'R_malyl_coa__R00934', 'R_RXN_R05076',...
                    'R_RXN_R09283','R_RXN_R09282','R_malyl_coa__R00237'};
            fluxes = [1 -1 1 -1 0.5 -0.5 0.5 0.5 0.5 -0.5 0.5 -0.5 0.5 0.5 -0.5 0.5]'*2;
        case {'rGlyP','rGlyP (w/o SHMT)'}
            reaction_names = {'R_formate_te_R00943','R_methenylte_R01655','R_methylenet_R01220',...
                'R_RXN_Q00006','R_RXN_Q00007','R_diaphorase_R01698','R_glycine_hy_R00945',...
                'R_l_serine_d_R00220'};
            fluxes = [2 -2 -2 -1 -1 1 1 1]';
% GAP-generating pathways:
        case 'CBB(PTS)'
            reaction_names =   {'R_ribulose_b_R00024_2','R_phosphogly_R01512','R_glyceralde_R01063',...
                    'R_triose_pho_R01015','R_fructose_b_R01068','R_diphosphat_R02073',...
                    'R_transketol_R01067','R_fructose_b_R01829','R_RXN_P00011',...
                    'R_transketol_R01641','R_ribulose_p_R01529','R_ribose_5_p_R01056',...
                    'R_phosphorib_R01523','R_RXN_P00010'};
            fluxes = [-3, 6, -6, 2, -1, 1, 1, -1, 1, 1, -2, 1, 1, -2]';
        case 'CBB'
            reaction_names =   {'R_ribulose_b_R00024_2','R_phosphogly_R01512','R_glyceralde_R01063',...
                    'R_triose_pho_R01015','R_fructose_b_R01068','R_fructose_b_R00762',...
                    'R_transketol_R01067','R_fructose_b_R01829','R_sedoheptul_R01845',...
                    'R_transketol_R01641','R_ribulose_p_R01529','R_ribose_5_p_R01056',...
                    'R_phosphorib_R01523'};
            fluxes = [-3, 6, -6, 2, -1, 1, 1, -1, 1, 1, -2, 1, 3]';
        case 'CBB(Photorespiration)'
            reaction_names =   {'R_ribulose_b_R00024_2','R_phosphogly_R01512','R_glyceralde_R01063',...
                    'R_triose_pho_R01015','R_fructose_b_R01068','R_fructose_b_R00762',...
                    'R_transketol_R01067','R_fructose_b_R01829','R_sedoheptul_R01845',...
                    'R_transketol_R01641','R_ribulose_p_R01529','R_ribose_5_p_R01056',...
                    'R_phosphorib_R01523'};
            fluxes = [-3, 6, -6, 2, -1, 1, 1, -1, 1, 1, -2, 1, 3]';
        case 'RuMP-Cycle'
            reaction_names =   {'R_methanol_d_R00605','R_3_hexulose_R05338','R_6_phospho__R05339',...
                    'R_6_phosphof_R00756','R_transketol_R01067','R_fructose_b_R01068',...
                    'R_triose_pho_R01015','R_fructose_b_R01829','R_sedoheptul_R01845',...
                    'R_transketol_R01641','R_ribulose_p_R01529','R_ribose_5_p_R01056'};

            fluxes = [3 -3 3 2 1 2 -1 -1 1 1 -2 1]';
        case 'RuMP-Cycle(Formate)'
            reaction_names =   {'R_RXN_Q00004','R_3_hexulose_R05338','R_6_phospho__R05339',...
                    'R_6_phosphof_R00756','R_transketol_R01067','R_fructose_b_R01068',...
                    'R_triose_pho_R01015','R_fructose_b_R01829','R_sedoheptul_R01845',...
                    'R_transketol_R01641','R_ribulose_p_R01529','R_ribose_5_p_R01056',...
                    'R_RXN_Q00003','R_stk_R00405'};

            fluxes = [-3 -3 3 2 1 2 -1 -1 1 1 -2 1 -3 3]';
        case 'RuMP-Cycle(Formate/THF)'
            reaction_names =   {'R_RXN_Q00005','R_3_hexulose_R05338','R_6_phospho__R05339',...
                    'R_6_phosphof_R00756','R_transketol_R01067','R_fructose_b_R01068',...
                    'R_triose_pho_R01015','R_fructose_b_R01829','R_sedoheptul_R01845',...
                    'R_transketol_R01641','R_ribulose_p_R01529','R_ribose_5_p_R01056',...
                    'R_glycine_hy_R00945','R_methylenet_R01220','R_methenylte_R01655','R_formate_te_R00943'};
            fluxes = [3 -3 3 2 1 2 -1 -1 1 1 -2 1 3 -3 -3 3]';
        case 'Photorespiration'
            reaction_names =    {'R_ribulose_b_R03140','R_phosphogly_R01334','R_s_2_hydrox_R00475',...
                                'R_glycine_ox_R00373','R_l_erythro__R00478','R_RXN_Q00085','R_aspartate__R07410',...
                                'R_phosphoeno_R00341','R_phosphopyr_R00658','R_phosphogly_R01518','R_phosphogly_R01512',...
                                'R_glyceralde_R01063','R_triose_pho_R01015','R_fructose_b_R01068','R_fructose_b_R00762',...
                                'R_transketol_R01067','R_fructose_b_R01829','R_sedoheptul_R01845',...
                                'R_transketol_R01641','R_ribulose_p_R01529','R_ribose_5_p_R01056',...
                                'R_phosphorib_R01523'};
            fluxes =            [3 3 3,...
                                1.5, -1.5, 1.5, 1.5,...
                                1.5 -1.5 1.5 4.5,...
                                -4.5,...
                                2, -1, 1, 1, -1, 1, 1, -2, 1, 3]'*2;
% Include the modules connecting the metabolites as well:
        case 'RevGlyoxShunt'
            reaction_names =    {'R_isocitrate_R00479','R_aconitate__R01900','R_aconitate__R01325',... % from this line: rGlyoxylate-Shunt
                                'R_atp_citrat_R00352','R_malate_deh_R00342','R_fumarate_h_R01082','R_fumarate_r_R09490'}; 
            fluxes =            [1, 1, -1, 1, -1, 1, -1]';
        case 'b-OH-AspCycle'
            reaction_names =    {'R_glycine_ox_R00373','R_l_erythro__R00478','R_RXN_Q00085','R_aspartate__R07410'}; % β-hydroxyaspartate cycle
            fluxes =            [1, -1, 1, 1]';
        case 'OAAtoGAP'
            reaction_names =    {'R_phosphoeno_R00341','R_phosphopyr_R00658','R_phosphogly_R01518','R_phosphogly_R01512','R_glyceralde_R01063'};
            fluxes =            [1, -1, 1, 1, -1]';
        case 'GAPtoOAA'
            reaction_names =    {'R_RXN_Q00000','R_phosphopyr_R00658','R_phosphogly_R01518','R_phosphogly_R01512','R_glyceralde_R01063'};
            fluxes =            [-1, 1, -1, -1, 1]';
        case 'GlyoxToOAA-HPbicycle(AMP)'
            reaction_names =    {'R_malyl_coa__R00934','R_RXN_R05076','R_RXN_R09283','R_RXN_R09282','R_malyl_coa__R00237','R_RXN_Q00074',...
                                'R_RXN_Q00091','R_RXN_Q00092','R_RXN_Q00093'};
            fluxes =            [-1,1,1,-1,1,1,...
                                1,-1,1]';
        case 'GlyoxToOAA-HPbicycle(ADP)'
            reaction_names =    {'R_malyl_coa__R00934','R_RXN_R05076','R_RXN_R09283','R_RXN_R09282','R_malyl_coa__R00237','R_RXN_Q00074',...
                                'R_RXN_Q00091','R_RXN_Q00092','R_3_hydroxyi_R03157','R_RXN_Q00094'};
            fluxes =            [-1,1,1,-1,1,1,...
                                1,-1,1,-1]';
        case 'EthylMalonylGlyox(Mod)'
            reaction_names =    {'R_acetyl_coa_R00238','R_3_hydroxyb_R01976','R_enoyl_coen_R03026','R_RXN_R09291','R_RXN_R09979','R_RXN_R09292',...
                                'R_RXN_Q00073','R_RXN_R05076','R_malyl_coa__R00934','R_RXN_Q00084','R_methylmalo_R02765','R_methylmalo_R00833',...
                                'R_RXN_R00406','R_fumarate_r_R09490','R_fumarate_h_R01082','R_malyl_coa__R00473'};
            fluxes =            [1,-1,1,-1,1,1,...
                                1,-1,1,1,-1,1,...
                                1,1,-1,1]';
        case 'EtMalOAAGlyox'
            reaction_names =    {'R_acetyl_coa_R00238','R_3_hydroxyb_R01976','R_enoyl_coen_R03026','R_RXN_R09291','R_RXN_R09979','R_RXN_R09292',...
                                'R_RXN_Q00073','R_RXN_R05076','R_malyl_coa__R00934','R_RXN_Q00084','R_methylmalo_R02765','R_methylmalo_R00833',...
                                'R_stk_R00405','R_fumarate_r_R09490','R_fumarate_h_R01082','R_malate_deh_R00342'};,...%,'R_malate_deh_R00216','R_RXN_Q00074',...
                                %'R_glycine_ox_R00373','R_l_erythro__R00478','R_RXN_Q00085','R_aspartate__R07410'
            fluxes =            [1,-1,1,-1,1,1,...
                                1,-1,1,1,-1,1,...
                                -1,1,-1,1]';%...%1,1,...
                                %0.5, -0.5, 0.5, 0.5
        case 'rCC-AcCoAtoPyr'
            reaction_names =    {'R_acetyl_coa_R00238','R_3_hydroxyb_R01976','R_enoyl_coen_R03026','R_RXN_R09291','R_RXN_R09979','R_RXN_R09292',...
                                'R_RXN_Q00073','R_RXN_R09283','R_RXN_R09282','R_malyl_coa__R00237'};
            fluxes =            [1,-1,1,-1,1,1,...
                                1,1,-1,1]';
        case 'GlyoxToOAA(Ser)'
            reaction_names =    {'R_formate_te_R00943','R_methenylte_R01655','R_methylenet_R01220',...
                                'R_glycine_hy_R00945','R_serine_gly_R00588','R_glyoxylate_R01388',...
                                'R_glycerate__R08572','R_phosphopyr_R00658','R_RXN_Q00000'};
            fluxes =            [1 -1 -1 1 1 -1 1 1 -1]';
        case 'C3to2C2(4HB)(T)'
            reaction_names =    {'R_RXN_Q00074','R_malate_deh_R00342','R_fumarate_h_R01082','R_fumarate_r_R09490','R_stk_R00405',...
                                'R_succinate__R09280','R_RXN_R09281','R_RXN_Q00071','R_RXN_Q00072','R_enoyl_coen_R03026',...
                                'R_3_hydroxya_R01975','R_acetyl_coa_R00238'};
            fluxes =            [1,-1,1,-1,1,...
                                -1,-1,1,1,-1,...
                                1,-1]';
        case 'C3to2C2(4HB)(C)'
            reaction_names =    {'R_RXN_Q00074','R_malate_deh_R00342','R_fumarate_h_R01082','R_fumarate_r_R09490','R_stk_R00405',...
                                'R_succinate__R09280','R_RXN_R09281','R_RXN_R09279','R_RXN_Q00072','R_enoyl_coen_R03026',...
                                'R_3_hydroxya_R01975','R_acetyl_coa_R00238'};
            fluxes =            [1,-1,1,-1,1,...
                                -1,-1,1,1,-1,...
                                1,-1]';
        case 'PyrDehCompl'
            reaction_names =    {'R_diaphorase_R00209'};
            fluxes =            1;
        case {'PyrC','-PyrC'}
            reaction_names =    {'R_RXN_Q00074'};
            fluxes =            1;
        case 'PyrKin'
            reaction_names =    {'R_pyruvate_k_R00200'};
            fluxes =            1;
        case 'PEPCarb'
            reaction_names =    {'R_RXN_Q00000'};
            fluxes =            1;            
        case 'MCG-like-cycle' % converts 1 Oxaloacetate to 2 Acetyl-CoA (alternative to PyrTo2AcCoA(4-HB)(AMP))
            reaction_names =    {'R_malate_deh_R00342','R_RXN_R00406','R_stk_R00405','R_malyl_coa__R00473',...
                                'R_glycine_ox_R00373','R_l_erythro__R00478','R_RXN_Q00085','R_aspartate__R07410'}; % β-hydroxyaspartate cycle};
            fluxes =            [-2,2,2,2,...
                                1, -1, 1, 1]';
        case 'AcCoAtoOAA(HP)(ADP)'
            reaction_names =    {'R_RXN_Q00091','R_RXN_Q00092','R_3_hydroxyi_R03157','R_RXN_Q00094',...
                                'R_RXN_Q00084','R_methylmalo_R02765','R_methylmalo_R00833','R_stk_R00405',...
                                'R_fumarate_r_R09490','R_fumarate_h_R01082','R_malate_deh_R00342'};%'R_malate_deh_R00216','R_RXN_Q00074'};
            fluxes =            [1,-1,1,-1,...
                                1,-1,1,-1,...
                                1,-1,1]';
        case 'AcCoAtoOAA(HP)(AMP)'
            reaction_names =    {'R_RXN_Q00091','R_RXN_Q00092','R_RXN_Q00093',...
                                'R_RXN_Q00084','R_methylmalo_R02765','R_methylmalo_R00833','R_stk_R00405',...
                                'R_fumarate_r_R09490','R_fumarate_h_R01082','R_malate_deh_R00342'};%'R_malate_deh_R00216','R_RXN_Q00074'};
            fluxes =            [1,-1,1,...
                                1,-1,1,-1,...
                                1,-1,1]';%1,1]';
    end
end

end
```
