## Supplementary material for "In-depth computational analysis of natural and artificial carbon fixation pathways": MATLAB files: kinetics_subkinetics.m

```
function sub_kinetic_data = kinetics_subkinetics(kinetic_data, ind_reaction, ind_m)

sub_kinetic_data = struct;
properties = {'median','mean','std','lower','upper','mean_ln','std_ln','lower_ln','upper_ln'};

for j = 1:numel(properties)
    % Parameter with only rows
    sub_kinetic_data.KV.scaling = kinetic_data.KV.scaling;
    sub_kinetic_data.KV.(properties{j}) = kinetic_data.KV.(properties{j})(ind_reaction);
    % Parameter with rows and columns
    sub_kinetic_data.KM.scaling = kinetic_data.KM.scaling;
    sub_kinetic_data.KM.(properties{j}) = kinetic_data.KM.(properties{j})(ind_reaction,ind_m);
    sub_kinetic_data.KA.scaling = kinetic_data.KA.scaling;
    sub_kinetic_data.KA.(properties{j}) = kinetic_data.KA.(properties{j})(ind_reaction,ind_m);
    sub_kinetic_data.KI.scaling = kinetic_data.KI.scaling;
    sub_kinetic_data.KI.(properties{j}) = kinetic_data.KI.(properties{j})(ind_reaction,ind_m); 
    % Parameter with only rows
    sub_kinetic_data.c.scaling = kinetic_data.c.scaling;
    sub_kinetic_data.c.(properties{j}) = kinetic_data.c.(properties{j})(ind_m); 
    sub_kinetic_data.u.scaling = kinetic_data.u.scaling;
    sub_kinetic_data.u.(properties{j}) = kinetic_data.u.(properties{j})(ind_reaction);   
    sub_kinetic_data.Keq.scaling = kinetic_data.Keq.scaling;
    sub_kinetic_data.Keq.(properties{j}) = kinetic_data.Keq.(properties{j})(ind_reaction);
    sub_kinetic_data.Kcatf.scaling = kinetic_data.Kcatf.scaling;
    sub_kinetic_data.Kcatf.(properties{j}) = kinetic_data.Kcatf.(properties{j})(ind_reaction); 
    sub_kinetic_data.Kcatr.scaling = kinetic_data.Kcatr.scaling;
    sub_kinetic_data.Kcatr.(properties{j}) = kinetic_data.Kcatr.(properties{j})(ind_reaction);
end

end
```
