## Supplementary material for "In-depth computational analysis of natural and artificial carbon fixation pathways": MATLAB files: kinetics_write.m

```
function [kinetics_out] = kinetics_write(kinetics_in, r, ind_row, ind_metabolites)
    if(isfield(r,'KM')), kinetics_in.KM(ind_row, ind_metabolites) = r.KM; end
    if(isfield(r,'KA')),kinetics_in.KA(ind_row, ind_metabolites) = r.KA; end
    if(isfield(r,'KI')),kinetics_in.KI(ind_row, ind_metabolites) = r.KI; end
    if(isfield(r,'c')),kinetics_in.c(ind_metabolites) = r.c; end
    if(isfield(r,'u')),kinetics_in.u(ind_row) = r.u; end
    if(isfield(r,'dmu0')),kinetics_in.dmu0(ind_row) = r.dmu0; end
    if(isfield(r,'mu')),kinetics_in.mu(ind_metabolites) = r.mu; end
    if(isfield(r,'h')),kinetics_in.h(ind_row) = r.h;     end
    if(isfield(r,'Keq')),kinetics_in.Keq(ind_row) = r.Keq; end
    if(isfield(r,'Kcatf')),kinetics_in.Kcatf(ind_row) = r.Kcatf; end
    if(isfield(r,'Kcatr')),kinetics_in.Kcatr(ind_row) = r.Kcatr; end
    if(isfield(r,'A')),kinetics_in.A(ind_row) = r.A; end
    if(isfield(r,'Kcatratio')),kinetics_in.Kcatratio(ind_row) = r.Kcatratio; end
    kinetics_out = kinetics_in;
end
```
