## Supplementary material for "In-depth computational analysis of natural and artificial carbon fixation pathways": MATLAB files: RandomizeParamters.m

```
function kinetics_out = RandomizeParamters(kinetics, kinetics_std, network,rng_seed)

% Seems to work...

kinetics_out = kinetics;

rng(rng_seed+20,'twister'); % avoid strange MATLAB rng behaviour
kinetics_out.Kcatf = normrnd(log(kinetics.Kcatf), log(kinetics_std.Kcatf));
kinetics_out.Kcatr = normrnd(log(kinetics.Kcatr), log(kinetics_std.Kcatr));
kinetics_out.KM = normrnd(log(kinetics.KM), log(kinetics_std.KM));

KM_temp = kinetics_out.KM;
KM_std_temp = log(kinetics_std.KM);
KM_temp(isnan(KM_temp)) = zeros(numel(KM_temp(isnan(KM_temp))),1);
KM_std_temp(isnan(KM_std_temp)) = zeros(numel(KM_std_temp(isnan(KM_std_temp))),1);
std_sum = log(kinetics_std.Kcatf) + log(kinetics_std.Kcatr) + sum(abs(network.N).*KM_std_temp')';

log_Keq = kinetics_out.Kcatf-kinetics_out.Kcatr+sum(network.N'.*KM_temp,2);

diff_Keq = (log_Keq-log(kinetics.Keq));
kinetics_out.Kcatf = kinetics_out.Kcatf - diff_Keq.*log(kinetics_std.Kcatf)./std_sum;
kinetics_out.Kcatr = kinetics_out.Kcatr + diff_Keq.*log(kinetics_std.Kcatr)./std_sum;
kinetics_out.KM = kinetics_out.KM - 1./(network.N').*(diag(diff_Keq./std_sum)*(abs(network.N).*KM_std_temp')');

% KM_temp = kinetics_out.KM;
% KM_temp(isnan(KM_temp)) = zeros(numel(KM_temp(isnan(KM_temp))),1);
% log_Keq = kinetics_out.Kcatf-kinetics_out.Kcatr+sum(network.N'.*KM_temp,2);
% diff_Keq = (log_Keq-log(kinetics.Keq))

kinetics_out.Kcatf = exp(kinetics_out.Kcatf);
kinetics_out.Kcatr = exp(kinetics_out.Kcatr);
kinetics_out.KM = exp(kinetics_out.KM);
kinetics_out.KV = sqrt(kinetics_out.Kcatf.*kinetics_out.Kcatr);

end
```
