## Supplementary material for "In-depth computational analysis of natural and artificial carbon fixation pathways": MATLAB files: sbtab_to_modular_rate_law_std.m

```
function [kinetics, sbtab_table, other_parameters] = sbtab_to_modular_rate_law_std(network, file_kinetic_data, options)

% [kinetics, sbtab_table, other_parameters] = sbtab_to_modular_rate_law(network,file_kinetic_data,options)
%
% Build 'kinetics' data structure from kinetic data (in SBtab file)
%
% Similar functions: 'ms_import_kinetic', 'sbtab_to_modular_rate_law_via_kinetic_data'

eval(default('options','struct'));

options_default = struct('use_sbml_ids',1,'kinetic_law','cs','verbose','0');
options         = join_struct(options_default,options);

switch options.kinetic_law,
  case {'cs','ms','rp','ma','fm'}, % UPDATE rate law names!
     kinetics = set_kinetics(network,options.kinetic_law);
  otherwise, error('Conversion is only possible for modular rate law');
end

if isstr(file_kinetic_data),
  sbtab_table  = sbtab_table_load(file_kinetic_data);
else
  %% assume that file_kinetic_data contains already an sbtab data structure
  sbtab_table  = file_kinetic_data;
end
QuantityType = sbtab_table_get_column(sbtab_table,'QuantityType');
Value        = cell_string2num(sbtab_table_get_column(sbtab_table,'Value'));
Std = [];
if isempty(Value) || isnan(Value)
    Value        = str2double(sbtab_table_get_column(sbtab_table,'Mean'));
    Std        = str2double(sbtab_table_get_column(sbtab_table,'Std'));
end
Compound     = sbtab_table_get_column(sbtab_table,'Compound');
if isempty(Compound)
    Compound     = sbtab_table_get_column(sbtab_table,'Compound:SBML:species:id');
end
Reaction     = sbtab_table_get_column(sbtab_table,'Reaction');
if isempty(Reaction)
    Reaction     = sbtab_table_get_column(sbtab_table,'Reaction:SBML:reaction:id');
end
compound_ind = label_names(Compound,network.metabolites);
reaction_ind = label_names(Reaction,network.actions);

[nr,nm,nx,ind_KM,ind_KA,ind_KI,nKM,nKA,nKI] = network_numbers(network);

ind_Keq = find(strcmp(QuantityType,'equilibrium constant'));
ind_Keq = ind_Keq(label_names(Reaction(ind_Keq),network.actions));
kinetics.Keq = Value(ind_Keq);
if(~isempty(Std) && sum(isnan(Std)) ~= length(Std)) 
    kinetics.dKeq = Std(ind_Keq);
end

ind_KV = find(strcmp(QuantityType,'catalytic rate constant geometric mean'));
ind_KV = ind_KV(label_names(Reaction(ind_KV),network.actions));
kinetics.KV = Value(ind_KV);
if(~isempty(Std) && sum(isnan(Std)) ~= length(Std))
    kinetics.dKV = Std(ind_KV);
end

ind_Kcatf = find(strcmp(QuantityType,'substrate catalytic rate constant'));
i_labels = label_names(Reaction(ind_Kcatf),network.actions);
%ind_Kcatf = ind_Kcatf(label_names(Reaction(ind_Kcatf),network.actions));
kinetics.Kcatf = NaN*ones(nr,1);
kinetics.Kcatf(i_labels) = Value(ind_Kcatf); %Value(ind_Kcatf);
if(~isempty(Std) && sum(isnan(Std)) ~= length(Std))
    kinetics.dKcatf = NaN*ones(nr,1);
    kinetics.dKcatf(i_labels) = Std(ind_Kcatf);
end

ind_Kcatr = find(strcmp(QuantityType,'product catalytic rate constant'));
i_labels = label_names(Reaction(ind_Kcatr),network.actions);
%ind_Kcatf = ind_Kcatf(label_names(Reaction(ind_Kcatf),network.actions));
kinetics.Kcatr = NaN*ones(nr,1);
kinetics.Kcatr(i_labels) = Value(ind_Kcatr); %Value(ind_Kcatf);
if(~isempty(Std) && sum(isnan(Std)) ~= length(Std))
    kinetics.dKcatr = NaN*ones(nr,1);
    kinetics.dKcatr(i_labels) = Std(ind_Kcatr);
end

kinetics.KV = sqrt(kinetics.Kcatr.*kinetics.Kcatf);

ind_KM  = find(strcmp(QuantityType,'Michaelis constant'));
ind_KMc = label_names(Compound(ind_KM),network.metabolites);
ind_KMr = label_names(Reaction(ind_KM),network.actions);
kinetics.KM = zeros(nr,nm);
kinetics.KM(sub2ind([nr,nm],ind_KMr,ind_KMc)) = Value(ind_KM);
if(~isempty(Std) && sum(isnan(Std)) ~= length(Std))
    kinetics.dKM = zeros(nr,nm);
    kinetics.dKM(sub2ind([nr,nm],ind_KMr,ind_KMc)) = Std(ind_KM);
end

ind_KI  = find(strcmp(QuantityType,'inhibitory constant'));
ind_KIc = label_names(Compound(ind_KI),network.metabolites);
ind_KIr = label_names(Reaction(ind_KI),network.actions);
kinetics.KI = zeros(nr,nm);
kinetics.KI(sub2ind([nr,nm],ind_KIr,ind_KIc)) = Value(ind_KI);
if(~isempty(Std) && sum(isnan(Std)) ~= length(Std))
    kinetics.dKI = zeros(nr,nm);
    kinetics.dKI(sub2ind([nr,nm],ind_KIr,ind_KIc)) = Std(ind_KI);
end

ind_KA  = find(strcmp(QuantityType,'activation constant'));
ind_KAc = label_names(Compound(ind_KA),network.metabolites);
ind_KAr = label_names(Reaction(ind_KA),network.actions);
kinetics.KA = zeros(nr,nm);
kinetics.KA(sub2ind([nr,nm],ind_KAr,ind_KAc)) = Value(ind_KA);
if(~isempty(Std) && sum(isnan(Std)) ~= length(Std))
    kinetics.dKA = zeros(nr,nm);
    kinetics.dKA(sub2ind([nr,nm],ind_KIr,ind_KIc)) = Std(ind_KA);
end

ind_c = find(strcmp(QuantityType,'concentration'));
ll    = label_names(Compound(ind_c),network.metabolites);
ind_c = ind_c(find(ll));
kinetics.c = Value(ind_c);
if(~isempty(Std) && sum(isnan(Std)) ~= length(Std))
    kinetics.dc = Std(ind_c);
end

ind_u = find(strcmp(QuantityType,'concentration of enzyme'));
ll    = label_names(Reaction(ind_u),network.actions);
ind_u = ind_u(find(ll));
kinetics.u = Value(ind_u);
if(~isempty(Std) && sum(isnan(Std)) ~= length(Std))
    kinetics.du = Std(ind_u);
end

ind_CompoundMass = find(strcmp(QuantityType,'molecular mass'));
ll = label_names(network.metabolites,Compound(ind_CompoundMass));
other_parameters.metabolite_mass     = nan * ones(nm,1);
other_parameters.metabolite_mass(find(ll)) = Value(ind_CompoundMass(ll(find(ll))));

ind_EnzymeMass = find(strcmp(QuantityType,'protein molecular mass'));
i_labels = label_names(Reaction(ind_EnzymeMass),network.actions);
%ind_Kcatf = ind_Kcatf(label_names(Reaction(ind_Kcatf),network.actions));
other_parameters.enzyme_mass = NaN*ones(nr,1);
other_parameters.enzyme_mass(i_labels) = Value(ind_EnzymeMass);

% ind_EnzymeMass = find(strcmp(QuantityType,'protein molecular mass'));
% ll = label_names(network.actions,Reaction(ind_EnzymeMass));
% other_parameters.enzyme_mass   = nan * ones(nr,1);
% other_parameters.enzyme_mass(find(ll))     = Value(ind_EnzymeMass(ll(find(ll))));

[computed_Kcatf, computed_Kcatr] = modular_KV_Keq_to_kcat(network.N,kinetics,kinetics.KV,kinetics.Keq,kinetics.KM,kinetics.h);

if norm(log(computed_Kcatf) - log(kinetics.Kcatf)) > 0.01 * length(computed_Kcatf), 
  warning('Given Kcat values and Kcat values computed from other parameters do not match'); 
end 

if norm(log(computed_Kcatr) - log(kinetics.Kcatr)) > 0.01 * length(computed_Kcatr), 
  warning('Given Kcat values and Kcat values computed from other parameters do not match'); 
end 
```
