## Supplementary material for "In-depth computational analysis of natural and artificial carbon fixation pathways": MATLAB files: TotalStoichiometryAsString.m

```
function o_str = TotalStoichiometryAsString(network, reaction_names, fluxes)

vv = zeros(length(network.reaction_names),1);

for i = 1:length(reaction_names)
    index = find(strcmp(network.reaction_names, reaction_names{i}));
    vv(index(1)) = fluxes(i);
end

flux = network.N*vv;
o_str = "";
for i=1:length(flux)
    if(flux(i) < -0.000001)
        if(strlength(o_str) ~= 0)
            o_str = o_str + " + ";
        end
        o_str = o_str + sprintf("%.2f %s", -flux(i), network.metabolites{i});
    end
end

o_str = o_str + " -> ";
length0 = strlength(o_str);

for i=1:length(flux)
    if(flux(i) > 0.000001)
        if(strlength(o_str) ~= length0)
            o_str = o_str + " + ";
        end
        o_str = o_str + sprintf("%.2f %s", flux(i), network.metabolites{i});
    end
end

end
```
