## Supplementary material for "In-depth computational analysis of natural and artificial carbon fixation pathways": MATLAB files: append_pdfs.m

```
function append_pdfs(varargin)
%APPEND_PDFS Appends/concatenates multiple PDF files
%
% Example:
%   append_pdfs(outputFilename, inputFilename1, inputFilename2, ...)
%   append_pdfs(outputFilename, inputFilenames_list{:})
%   append_pdfs(outputFilename, inputFilenames_cell_or_string_array)
%   append_pdfs output.pdf input1.pdf input2.pdf
%
% This function appends multiple PDF files to an existing PDF file, or
% concatenates them into a PDF file if the output file doesn't yet exist.
%
% This function requires that you have ghostscript installed on your
% system. Ghostscript can be downloaded from: http://www.ghostscript.com
%
% IN:
%    output - string of output file name (including the extension, .pdf).
%             If it exists it is appended to; if not, it is created.
%    input1 - string of an input file name (including the extension, .pdf).
%             All input files are appended in order.
%    input_list - cell array list of input file name strings. All input
%                 files are appended in order.

% Copyright: Oliver Woodford, 2011-2014, Yair Altman 2015-

%{
% Thanks to Reinhard Knoll for pointing out that appending multiple pdfs in
% one go is much faster than appending them one at a time.

% Thanks to Michael Teo for reporting the issue of a too long command line.
% Issue resolved on 5/5/2011, by passing gs a command file.

% Thanks to Martin Wittmann for pointing out quality issue when appending bitmaps
% Issue resolved (to best of my ability) 1/6/2011, using the prepress setting

% 26/02/15: If temp dir is not writable, use the output folder for temp
%           files when appending (Javier Paredes); sanity check of inputs
% 24/01/18: Fixed error in case of existing output file (append mode)
% 24/01/18: Fixed issue #213: non-ASCII characters in folder names on Windows
% 06/12/18: Avoid an "invalid escape-char" warning upon error
% 22/03/20: Alert if ghostscript.m is not found on Matlab path
% 29/03/20: Accept a cell-array of input files (issue #299); accept both "strings", 'chars'
%}

    if nargin < 2,  return;  end  % sanity check

    % Convert strings => chars; strtrim extra spaces
    varargin = cellfun(@str2char,varargin,'un',false);

    % Convert cell array into individual strings (issue #299)
    if nargin==2 && iscell(varargin{2})
        varargin = {varargin{1} varargin{2}{:}}; %#ok<CCAT>
    end

    % Ensure that ghostscript() exists on the Matlab path
    if ~exist('ghostscript','file')
        error('export_fig:append_pdfs:ghostscript', 'The ghostscript.m function is required by append_pdf.m. Install the complete export_fig package from https://www.mathworks.com/matlabcentral/fileexchange/23629-export_fig or https://github.com/altmany/export_fig')
    end

    % Are we appending or creating a new file
    append = exist(varargin{1}, 'file') == 2;
    output = [tempname '.pdf'];
    try
        % Ensure that the temp dir is writable (Javier Paredes 26/2/15)
        fid = fopen(output,'w');
        fwrite(fid,1);
        fclose(fid);
        delete(output);
        isTempDirOk = true;
    catch
        % Temp dir is not writable, so use the output folder
        [dummy,fname,fext] = fileparts(output); %#ok<ASGLU>
        fpath = fileparts(varargin{1});
        output = fullfile(fpath,[fname fext]);
        isTempDirOk = false;
    end
    if ~append
        output = varargin{1};
        varargin = varargin(2:end);
    end

    % Create the command file
    if isTempDirOk
        cmdfile = [tempname '.txt'];
    else
        cmdfile = fullfile(fpath,[fname '.txt']);
    end
    prepareCmdFile(cmdfile, output, varargin{:});

    % Call ghostscript
    [status, errMsg] = ghostscript(['@"' cmdfile '"']);

    % Check for ghostscript execution errors
    if status && ~isempty(strfind(errMsg,'undefinedfile')) && ispc %#ok<STREMP>
        % Fix issue #213: non-ASCII characters in folder names on Windows
        for fileIdx = 2 : numel(varargin)
            [fpath,fname,fext] = fileparts(varargin{fileIdx});
            varargin{fileIdx} = fullfile(normalizePath(fpath),[fname fext]);
        end
        % Rerun ghostscript with the normalized folder names
        prepareCmdFile(cmdfile, output, varargin{:});
        [status, errMsg] = ghostscript(['@"' cmdfile '"']);
    end

    % Delete the command file
    delete(cmdfile);

    % Check for ghostscript execution errors
    if status
        errMsg = strrep(errMsg,'\','\\');  % Avoid an "invalid escape-char" warning
        error('YMA:export_fig:append_pdf',errMsg);
    end

    % Rename the file if needed
    if append
        movefile(output, varargin{1}, 'f');
    end
end

% Prepare a text file with ghostscript directives
function prepareCmdFile(cmdfile, output, varargin)
    fh = fopen(cmdfile, 'w');
    fprintf(fh, '-q -dNOPAUSE -dBATCH -sDEVICE=pdfwrite -dPDFSETTINGS=/prepress -sOutputFile="%s" -f', output);
    fprintf(fh, ' "%s"', varargin{:});
    fclose(fh);
end

% Convert long/non-ASCII folder names into their short ASCII equivalents
function pathStr = normalizePath(pathStr)
    [fpath,fname,fext] = fileparts(pathStr);
    if isempty(fpath) || strcmpi(fpath,pathStr), return, end
    dirOutput = evalc(['system(''dir /X /AD "' pathStr '*"'')']);
    shortName = strtrim(regexprep(dirOutput,{'.*> *',[fname fext '.*']},''));
    if isempty(shortName)
        shortName = [fname fext];
    end
    fpath = normalizePath(fpath);  %recursive until entire fpath is processed
    pathStr = fullfile(fpath, shortName);
end

% Convert a possible string => char
function value = str2char(value)
    try
        value = controllib.internal.util.hString2Char(value);
    catch
        if isa(value,'string')
            value = char(value);
        end
    end
    value = strtrim(value);
end
```
