## Supplementary material for "In-depth computational analysis of natural and artificial carbon fixation pathways": MATLAB files: export_fig.m

```
function [imageData, alpha] = export_fig(varargin) %#ok<*STRCL1>
%EXPORT_FIG  Exports figures in a publication-quality format
%
% Examples:
%   imageData = export_fig
%   [imageData, alpha] = export_fig
%   export_fig filename
%   export_fig filename -format1 -format2
%   export_fig ... -nocrop
%   export_fig ... -c[<val>,<val>,<val>,<val>]
%   export_fig ... -transparent
%   export_fig ... -native
%   export_fig ... -m<val>
%   export_fig ... -r<val>
%   export_fig ... -a<val>
%   export_fig ... -q<val>
%   export_fig ... -p<val>
%   export_fig ... -d<gs_option>
%   export_fig ... -depsc
%   export_fig ... -<renderer>
%   export_fig ... -<colorspace>
%   export_fig ... -append
%   export_fig ... -bookmark
%   export_fig ... -clipboard<:format>
%   export_fig ... -update
%   export_fig ... -version
%   export_fig ... -nofontswap
%   export_fig ... -font_space <char>
%   export_fig ... -linecaps
%   export_fig ... -noinvert
%   export_fig ... -preserve_size
%   export_fig ... -options <optionsStruct>
%   export_fig ... -silent
%   export_fig(..., handle)
%
% This function saves a figure or single axes to one or more vector and/or
% bitmap file formats, and/or outputs a rasterized version to the workspace,
% with the following properties:
%   - Figure/axes reproduced as it appears on screen
%   - Cropped borders (optional)
%   - Embedded fonts (vector formats)
%   - Improved line and grid line styles
%   - Anti-aliased graphics (bitmap formats)
%   - Render images at native resolution (optional for bitmap formats)
%   - Transparent background supported (pdf, eps, png, tif)
%   - Semi-transparent patch objects supported (png, tif)
%   - RGB, CMYK or grayscale output (CMYK only with pdf, eps, tif)
%   - Variable image compression, including lossless (pdf, eps, jpg)
%   - Optional rounded line-caps (pdf, eps)
%   - Optionally append to file (pdf, tif)
%   - Vector formats: pdf, eps, emf, svg
%   - Bitmap formats: png, tif, jpg, bmp, clipboard, export to workspace
%
% This function is especially suited to exporting figures for use in
% publications and presentations, because of the high quality and
% portability of media produced.
%
% Note that the background color and figure dimensions are reproduced
% (the latter approximately, and ignoring cropping & magnification) in the
% output file. For transparent background (and semi-transparent patch
% objects), use the -transparent option or set the figure 'Color' property
% to 'none'. To make axes transparent set the axes 'Color' property to
% 'none'. PDF, EPS, TIF & PNG are the only formats that support a transparent
% background; only TIF & PNG formats support transparency of patch objects.
%
% The choice of renderer (opengl/zbuffer/painters) has a large impact on the
% output quality. The default value (opengl for bitmaps, painters for vector
% formats) generally gives good results, but if you aren't satisfied
% then try another renderer.  Notes:
%   1) For vector formats (EPS,PDF), only painters generates vector graphics
%   2) For bitmap formats, only opengl correctly renders transparent patches
%   3) For bitmap formats, only painters correctly scales line dash and dot
%      lengths when magnifying or anti-aliasing
%   4) Fonts may be substitued with Courier when using painters
%
% When exporting to vector format (PDF & EPS) and bitmap format using the
% painters renderer, this function requires that ghostscript is installed
% on your system. You can download this from: http://www.ghostscript.com
% When exporting to EPS it additionally requires pdftops, from the Xpdf
% suite of functions. You can download this from: http://xpdfreader.com
%
% SVG output uses the fig2svg (https://github.com/kupiqu/fig2svg) or plot2svg
% (https://github.com/jschwizer99/plot2svg) utilities, or Matlab's built-in
% SVG export if neither of these utilities are available on Matlab's path.
% Note: cropping/padding are not supported in export_fig's SVG and EMF output.
%
% Inputs:
%   filename - string containing the name (optionally including full or
%             relative path) of the file the figure is to be saved as. If
%             a path is not specified, the figure is saved in the current
%             directory. If no name and no output arguments are specified,
%             the default name, 'export_fig_out', is used. If neither a
%             file extension nor a format are specified, a ".png" is added
%             and the figure saved in that format.
%   -<format> - string(s) containing the output file extension(s). Options:
%             '-pdf', '-eps', 'emf', '-svg', '-png', '-tif', '-jpg' and '-bmp'.
%             Multiple formats can be specified, without restriction.
%             For example: export_fig('-jpg', '-pdf', '-png', ...)
%             Note: '-tif','-tiff' are equivalent, and so are '-jpg','-jpeg'.
%   -transparent - option indicating that the figure background is to be made
%             transparent (PNG,PDF,TIF,EPS,EMF formats only). Implies -noinvert.
%   -nocrop - option indicating that empty margins should not be cropped.
%   -c[<val>,<val>,<val>,<val>] - option indicating crop amounts. Must be
%             a 4-element vector of numeric values: [top,right,bottom,left]
%             where NaN/Inf indicates auto-cropping, 0 means no cropping, any
%             other value means cropping in pixel amounts. e.g. '-c7,15,0,NaN'
%             Note: this option is not supported by SVG and EMF formats.
%   -p<val> - option to pad a border of width val to exported files, where
%             val is either a relative size with respect to cropped image
%             size (i.e. p=0.01 adds a 1% border). For EPS & PDF formats,
%             val can also be integer in units of 1/72" points (abs(val)>1).
%             val can be positive (padding) or negative (extra cropping).
%             If used, the -nocrop flag will be ignored, i.e. the image will
%             always be cropped and then padded. Default: 0 (i.e. no padding).
%             Note: this option is not supported by SVG and EMF formats.
%   -m<val> - option val indicates the factor to magnify the figure dimensions
%             when generating bitmap outputs (does not affect vector formats).
%             Default: '-m1' (i.e. val=1). Note: val~=1 slows down export_fig.
%   -r<val> - option val indicates the resolution (in pixels per inch) to
%             export bitmap and vector outputs, without changing dimensions of
%             the on-screen figure. Default: '-r864' (for vector output only).
%             Note: -m option overides -r option for bitmap exports only.
%   -native - option indicating that the output resolution (when outputting
%             a bitmap format) should be such that the vertical resolution
%             of the first suitable image found in the figure is at the
%             native resolution of that image. To specify a particular
%             image to use, give it the tag 'export_fig_native'. 
%             Notes: This overrides any value set with the -m and -r options.
%             It also assumes that the image is displayed front-to-parallel
%             with the screen. The output resolution is approximate and
%             should not be relied upon. Anti-aliasing can have adverse
%             effects on image quality (disable with the -a1 option).
%   -a1, -a2, -a3, -a4 - option indicating the amount of anti-aliasing (AA) to
%             use for bitmap outputs, when GraphicsSmoothing is not available.
%             '-a1'=no AA; '-a4'=max. Default: 3 for HG1, 1 for HG2.
%   -<renderer> - option to force a particular renderer (painters, opengl or
%             [in R2014a or older] zbuffer). Default value: opengl for bitmap
%             formats or figures with patches and/or transparent annotations;
%             painters for vector formats without patches/transparencies.
%   -<colorspace> - option indicating which colorspace color figures should
%             be saved in: RGB (default), CMYK or gray. Usage example: '-gray'.
%             Note: CMYK is only supported in PDF, EPS and TIF formats.
%   -q<val> - option to vary bitmap image quality (PDF, EPS, JPG formats only).
%             A larger val, in the range 0-100, produces higher quality and
%             lower compression. val > 100 results in lossless compression.
%             Default: '-q95' for JPG, ghostscript prepress default for PDF,EPS.
%             Note: lossless compression can sometimes give a smaller file size
%             than the default lossy compression, depending on the image type.
%   -append - option indicating that if the file already exists the figure is to
%             be appended as a new page, instead of being overwritten (default).
%             PDF & TIF output formats only.
%   -bookmark - option to indicate that a bookmark with the name of the
%             figure is to be created in the output file (PDF format only).
%   -clipboard - option to save output as an image on the system clipboard.
%   -clipboard<:format> - copies to clipboard in the specified format:
%             image (default), bitmap, emf, or pdf.
%             Notes: Only -clipboard (or -clipboard:image, which is the same)
%                    applies export_fig parameters such as cropping, padding etc.
%                    Only the emf format supports -transparent background
%             -clipboard:image  create a bitmap image using export_fig processing
%             -clipboard:bitmap create a bitmap image as-is (no auto-cropping etc.)
%             -clipboard:emf is vector format without auto-cropping; Windows-only
%             -clipboard:pdf is vector format without cropping; not universally supported
%   -d<gs_option> - option to indicate a ghostscript setting. For example,
%             -dMaxBitmap=0 or -dNoOutputFonts (Ghostscript 9.15+).
%   -depsc -  option to use EPS level-3 rather than the default level-2 print
%             device. This solves some bugs with Matlab's default -depsc2 device
%             such as discolored subplot lines on images (vector formats only).
%   -update - option to download and install the latest version of export_fig
%   -version - return the current export_fig version, without any figure export
%   -nofontswap - option to avoid font swapping. Font swapping is automatically
%             done in vector formats (only): 11 standard Matlab fonts are
%             replaced by the original figure fonts. This option prevents this.
%   -font_space <char> - option to set a spacer character for font-names that
%             contain spaces, used by EPS/PDF. Default: ''
%   -linecaps - option to create rounded line-caps (vector formats only).
%   -noinvert - option to avoid setting figure's InvertHardcopy property to
%             'off' during output (this solves some problems of empty outputs).
%   -preserve_size - option to preserve the figure's PaperSize property in output
%             file (PDF/EPS formats only; default is to not preserve it).
%   -options <optionsStruct> - format-specific parameters as defined in Matlab's
%             documentation of the imwrite function, contained in a struct under
%             the format name. For example to specify the JPG Comment parameter,
%             pass a struct such as this: options.JPG.Comment='abc'. Similarly,
%             options.PNG.BitDepth=4. Valid only for PNG,TIF,JPG output formats.
%   -silent - option to avoid various warning and informational messages, such
%             as version update checks, transparency or renderer issues, etc.
%   handle -  The handle of the figure, axes or uipanels (can be an array of
%             handles, but the objects must be in the same figure) which is
%             to be saved. Default: gcf (handle of current figure).
%
% Outputs:
%   imageData - MxNxC uint8 image array of the exported image.
%   alpha     - MxN single array of alphamatte values in the range [0,1],
%               for the case when the background is transparent.
%
%   Some helpful examples and tips can be found at:
%      https://github.com/altmany/export_fig
%
%   See also PRINT, SAVEAS, ScreenCapture (on the Matlab File Exchange)

%{
% Copyright (C) Oliver Woodford 2008-2014, Yair Altman 2015-

% The idea of using ghostscript is inspired by Peder Axensten's SAVEFIG
% (fex id: 10889) which is itself inspired by EPS2PDF (fex id: 5782).
% The idea for using pdftops came from the MATLAB newsgroup (id: 168171).
% The idea of editing the EPS file to change line styles comes from Jiro
% Doke's FIXPSLINESTYLE (fex id: 17928).
% The idea of changing dash length with line width came from comments on
% fex id: 5743, but the implementation is mine :)
% The idea of anti-aliasing bitmaps came from Anders Brun's MYAA (fex id:
% 20979).
% The idea of appending figures in pdfs came from Matt C in comments on the
% FEX (id: 23629)

% Thanks to Roland Martin for pointing out the colour MATLAB
% bug/feature with colorbar axes and transparent backgrounds.
% Thanks also to Andrew Matthews for describing a bug to do with the figure
% size changing in -nodisplay mode. I couldn't reproduce it, but included a
% fix anyway.
% Thanks to Tammy Threadgill for reporting a bug where an axes is not
% isolated from gui objects.
%}
%{
% 23/02/12: Ensure that axes limits don't change during printing
% 14/03/12: Fix bug in fixing the axes limits (thanks to Tobias Lamour for reporting it).
% 02/05/12: Incorporate patch of Petr Nechaev (many thanks), enabling bookmarking of figures in pdf files.
% 09/05/12: Incorporate patch of Arcelia Arrieta (many thanks), to keep tick marks fixed.
% 12/12/12: Add support for isolating uipanels. Thanks to michael for suggesting it.
% 25/09/13: Add support for changing resolution in vector formats. Thanks to Jan Jaap Meijer for suggesting it.
% 07/05/14: Add support for '~' at start of path. Thanks to Sally Warner for suggesting it.
% 24/02/15: Fix Matlab R2014b bug (issue #34): plot markers are not displayed when ZLimMode='manual'
% 25/02/15: Fix issue #4 (using HG2 on R2014a and earlier)
% 25/02/15: Fix issue #21 (bold TeX axes labels/titles in R2014b)
% 26/02/15: If temp dir is not writable, use the user-specified folder for temporary EPS/PDF files (Javier Paredes)
% 27/02/15: Modified repository URL from github.com/ojwoodford to /altmany; Indented main function; Added top-level try-catch block to display useful workarounds
% 28/02/15: Enable users to specify optional ghostscript options (issue #36)
% 06/03/15: Improved image padding & cropping thanks to Oscar Hartogensis
% 26/03/15: Fixed issue #49 (bug with transparent grayscale images); fixed out-of-memory issue
% 26/03/15: Fixed issue #42: non-normalized annotations on HG1
% 26/03/15: Fixed issue #46: Ghostscript crash if figure units <> pixels
% 27/03/15: Fixed issue #39: bad export of transparent annotations/patches
% 28/03/15: Fixed issue #50: error on some Matlab versions with the fix for issue #42
% 29/03/15: Fixed issue #33: bugs in Matlab's print() function with -cmyk
% 29/03/15: Improved processing of input args (accept space between param name & value, related to issue #51)
% 30/03/15: When exporting *.fig files, then saveas *.fig if figure is open, otherwise export the specified fig file
% 30/03/15: Fixed edge case bug introduced yesterday (commit #ae1755bd2e11dc4e99b95a7681f6e211b3fa9358)
% 09/04/15: Consolidated header comment sections; initialize output vars only if requested (nargout>0)
% 14/04/15: Workaround for issue #45: lines in image subplots are exported in invalid color
% 15/04/15: Fixed edge-case in parsing input parameters; fixed help section to show the -depsc option (issue #45)
% 21/04/15: Bug fix: Ghostscript croaks on % chars in output PDF file (reported by Sven on FEX page, 15-Jul-2014)
% 22/04/15: Bug fix: Pdftops croaks on relative paths (reported by Tintin Milou on FEX page, 19-Jan-2015)
% 04/05/15: Merged fix #63 (Kevin Mattheus Moerman): prevent tick-label changes during export
% 07/05/15: Partial fix for issue #65: PDF export used painters rather than opengl renderer (thanks Nguyenr)
% 08/05/15: Fixed issue #65: bad PDF append since commit #e9f3cdf 21/04/15 (thanks Robert Nguyen)
% 12/05/15: Fixed issue #67: exponent labels cropped in export, since fix #63 (04/05/15)
% 28/05/15: Fixed issue #69: set non-bold label font only if the string contains symbols (\beta etc.), followup to issue #21
% 29/05/15: Added informative error message in case user requested SVG output (issue #72)
% 09/06/15: Fixed issue #58: -transparent removed anti-aliasing when exporting to PNG
% 19/06/15: Added -update option to download and install the latest version of export_fig
% 07/07/15: Added -nofontswap option to avoid font-swapping in EPS/PDF
% 16/07/15: Fixed problem with anti-aliasing on old Matlab releases
% 11/09/15: Fixed issue #103: magnification must never become negative; also fixed reported error msg in parsing input params
% 26/09/15: Alert if trying to export transparent patches/areas to non-PNG outputs (issue #108)
% 04/10/15: Do not suggest workarounds for certain errors that have already been handled previously
% 01/11/15: Fixed issue #112: use same renderer in print2eps as export_fig (thanks to Jesús Pestana Puerta)
% 10/11/15: Custom GS installation webpage for MacOS. Thanks to Andy Hueni via FEX
% 19/11/15: Fixed clipboard export in R2015b (thanks to Dan K via FEX)
% 21/02/16: Added -c option for indicating specific crop amounts (idea by Cedric Noordam on FEX)
% 08/05/16: Added message about possible error reason when groot.Units~=pixels (issue #149)
% 17/05/16: Fixed case of image YData containing more than 2 elements (issue #151)
% 08/08/16: Enabled exporting transparency to TIF, in addition to PNG/PDF (issue #168)
% 11/12/16: Added alert in case of error creating output PDF/EPS file (issue #179)
% 13/12/16: Minor fix to the commit for issue #179 from 2 days ago
% 22/03/17: Fixed issue #187: only set manual ticks when no exponent is present
% 09/04/17: Added -linecaps option (idea by Baron Finer, issue #192)
% 15/09/17: Fixed issue #205: incorrect tick-labels when Ticks number don't match the TickLabels number
% 15/09/17: Fixed issue #210: initialize alpha map to ones instead of zeros when -transparent is not used
% 18/09/17: Added -font_space option to replace font-name spaces in EPS/PDF (workaround for issue #194)
% 18/09/17: Added -noinvert option to solve some export problems with some graphic cards (workaround for issue #197)
% 08/11/17: Fixed issue #220: axes exponent is removed in HG1 when TickMode is 'manual' (internal Matlab bug)
% 08/11/17: Fixed issue #221: alert if the requested folder does not exist
% 19/11/17: Workaround for issue #207: alert when trying to use transparent bgcolor with -opengl
% 29/11/17: Workaround for issue #206: warn if exporting PDF/EPS for a figure that contains an image
% 11/12/17: Fixed issue #230: use OpenGL renderer when exported image contains transparency (also see issue #206)
% 30/01/18: Updated SVG message to point to https://github.com/kupiqu/plot2svg and display user-selected filename if available
% 27/02/18: Fixed issue #236: axes exponent cropped from output if on right-hand axes
% 29/05/18: Fixed issue #245: process "string" inputs just like 'char' inputs
% 13/08/18: Fixed issue #249: correct black axes color to off-black to avoid extra cropping with -transparent
% 27/08/18: Added a possible file-open reason in EPS/PDF write-error message (suggested by "craq" on FEX page)
% 22/09/18: Xpdf website changed to xpdfreader.com
% 23/09/18: Fixed issue #243: only set non-bold font (workaround for issue #69) in R2015b or earlier; warn if changing font
% 23/09/18: Workaround for issue #241: don't use -r864 in EPS/PDF outputs when -native is requested (solves black lines problem)
% 18/11/18: Issue #261: Added informative alert when trying to export a uifigure (which is not currently supported)
% 13/12/18: Issue #261: Fixed last commit for cases of specifying axes/panel handle as input, rather than a figure handle
% 13/01/19: Issue #72: Added basic SVG output support
% 04/02/19: Workaround for issues #207 and #267: -transparent implies -noinvert
% 08/03/19: Issue #269: Added ability to specify format-specific options for PNG,TIF,JPG outputs; fixed help section
% 21/03/19: Fixed the workaround for issues #207 and #267 from 4/2/19 (-transparent now does *NOT* imply -noinvert; -transparent output should now be ok in all formats)
% 12/06/19: Issue #277: Enabled preservation of figure's PaperSize in output PDF/EPS file
% 06/08/19: Remove warning message about obsolete JavaFrame in R2019b
% 30/10/19: Fixed issue #261: added support for exporting uifigures and uiaxes (thanks to idea by @MarvinILA)
% 12/12/19: Added warning in case user requested anti-aliased output on an aliased HG2 figure (issue #292)
% 15/12/19: Added promo message
% 08/01/20: (3.00) Added check for newer version online (initialized to version 3.00)
% 15/01/20: (3.01) Clarified/fixed error messages; Added error IDs; easier -update; various other small fixes
% 20/01/20: (3.02) Attempted fix for issue #285 (unsupported patch transparency in some Ghostscript versions); Improved suggested fixes message upon error
% 03/03/20: (3.03) Suggest to upload problematic EPS file in case of a Ghostscript error in eps2pdf (& don't delete this file)
% 22/03/20: (3.04) Workaround for issue #15; Alert if ghostscript file not found on Matlab path
% 10/05/20: (3.05) Fix the generated SVG file, based on Cris Luengo's SVG_FIX_VIEWBOX; Don't generate PNG when only SVG is requested
% 02/07/20: (3.06) Significantly improved performance (speed) and fidelity of bitmap images; Return alpha matrix for bitmap images; Fixed issue #302 (-update bug); Added EMF output; Added -clipboard formats (image,bitmap,emf,pdf); Added hints for exportgraphics/copygraphics usage in certain use-cases; Added description of new version features in the update message; Fixed issue #306 (yyaxis cropping); Fixed EPS/PDF auto-cropping with -transparent
% 06/07/20: (3.07) Fixed issue #307 (bug in padding of bitmap images); Fixed axes transparency in -clipboard:emf with -transparent
% 07/07/20: (3.08) Fixed issue #308 (bug in R2019a and earlier)
% 18/07/20: (3.09) Fixed issue #310 (bug with tiny image on HG1); Fixed title cropping bug
% 23/07/20: (3.10) Fixed issues #313,314 (figure position changes if units ~= pixels); Display multiple versions change-log, if relevant; Fixed issue #312 (PNG: only use alpha channel if -transparent was requested)
% 30/07/20: (3.11) Fixed issue #317 (bug when exporting figure with non-pixels units); Potential solve also of issue #303 (size change upon export)
% 14/08/20: (3.12) Fixed some exportgraphics/copygraphics compatibility messages; Added -silent option to suppress non-critical messages; Reduced promo message display rate to once a week; Added progress messages during export_fig('-update')
% 07/10/20: (3.13) Added version info and change-log links to update message (issue #322); Added -version option to return the current export_fig version; Avoid JavaFrame warning message; Improved exportgraphics/copygraphics infomercial message inc. support of upcoming Matlab R2021a
%}

    if nargout
        [imageData, alpha] = deal([]);
    end
    displaySuggestedWorkarounds = true;

    % Ensure the figure is rendered correctly _now_ so that properties like axes limits are up-to-date
    drawnow;
    pause(0.02);  % this solves timing issues with Java Swing's EDT (http://undocumentedmatlab.com/blog/solving-a-matlab-hang-problem)

    % Display promo (just once a week!)
    try promo_time = getpref('export_fig','promo_time'); catch, promo_time=-inf; end
    if abs(now-promo_time) > 7 && ~isdeployed
        programsCrossCheck;
        msg = 'For professional Matlab assistance,  please contact <$>';
        url = 'https://UndocumentedMatlab.com/consulting';
        displayPromoMsg(msg, url);
        setpref('export_fig','promo_time',now)
    end

    % Parse the input arguments
    fig = get(0, 'CurrentFigure');
    argNames = {};
    for idx = nargin:-1:1, argNames{idx} = inputname(idx); end
    [fig, options] = parse_args(nargout, fig, argNames, varargin{:});

    % Check for newer version and exportgraphics/copygraphics compatibility
    currentVersion = 3.13;
    if options.version  % export_fig's version requested - return it and bail out
        imageData = currentVersion;
        return
    end
    if ~options.silent
        % Check for newer version (not too often)
        checkForNewerVersion(3.13);  % ...(currentVersion) is better but breaks in version 3.05- due to regexp limitation in checkForNewerVersion()

        % Hint to users to use exportgraphics/copygraphics in certain cases
        alertForExportOrCopygraphics(options);
        %return
    end

    % Ensure that we have a figure handle
    if isequal(fig,-1)
        return  % silent bail-out
    elseif isempty(fig)
        error('export_fig:NoFigure','No figure found');
    else
        oldWarn = warning('off','MATLAB:HandleGraphics:ObsoletedProperty:JavaFrame');
        warning off MATLAB:ui:javaframe:PropertyToBeRemoved
        uifig = handle(ancestor(fig,'figure'));
        try jf = get(uifig,'JavaFrame_I'); catch, try jf = get(uifig,'JavaFrame'); catch, jf=1; end, end %#ok<JAVFM>
        warning(oldWarn);
        if isempty(jf)  % this is a uifigure
            %error('export_fig:uifigures','Figures created using the uifigure command or App Designer are not supported by export_fig. See %s for details.', hyperlink('https://github.com/altmany/export_fig/issues/261','issue #261'));
            if numel(fig) > 1
                error('export_fig:uifigure:multipleHandles', 'export_fig only supports exporting a single uifigure handle at a time; array of handles is not currently supported.')
            elseif ~any(strcmpi(fig.Type,{'figure','axes'}))
                error('export_fig:uifigure:notFigureOrAxes', 'export_fig only supports exporting a uifigure or uiaxes handle; other handles of a uifigure are not currently supported.')
            end
            % fig is either a uifigure or uiaxes handle
            isUiaxes = strcmpi(fig.Type,'axes');
            if isUiaxes
                % Label the specified axes so that we can find it in the legacy figure
                oldUserData = fig.UserData;
                tempStr = tempname;
                fig.UserData = tempStr;
            end
            try
                % Create an invisible legacy figure at the same position/size as the uifigure
                hNewFig = figure('Units',uifig.Units, 'Position',uifig.Position, 'MenuBar','none', 'ToolBar','none', 'Visible','off');
                % Copy the uifigure contents onto the new invisible legacy figure
                try
                    hChildren = allchild(uifig); %=uifig.Children;
                    copyobj(hChildren,hNewFig);
                catch
                    if ~options.silent
                        warning('export_fig:uifigure:controls', 'Some uifigure controls cannot be exported by export_fig and will not appear in the generated output.');
                    end
                end
                try fig.UserData = oldUserData; catch, end  % restore axes UserData, if modified above
                % Replace the uihandle in the input args with the legacy handle
                if isUiaxes  % uiaxes
                    % Locate the corresponding axes handle in the new legacy figure
                    hAxes = findall(hNewFig,'type','axes','UserData',tempStr);
                    if isempty(hAxes) % should never happen, check just in case
                        hNewHandle = hNewFig;  % export the figure instead of the axes
                    else
                        hNewHandle = hAxes;  % new axes handle found: use it instead of the uiaxes
                    end
                else  % uifigure
                    hNewHandle = hNewFig;
                end
                varargin(cellfun(@(c)isequal(c,fig),varargin)) = {hNewHandle};
                % Rerun export_fig on the legacy figure (with the replaced handle)
                [imageData, alpha] = export_fig(varargin{:});
                % Delete the temp legacy figure and bail out
                try delete(hNewFig); catch, end
                return
            catch err
                % Clean up the temp legacy figure and report the error
                try delete(hNewFig); catch, end
                rethrow(err)
            end
        end
    end

    % Isolate the subplot, if it is one
    cls = all(ismember(get(fig, 'Type'), {'axes', 'uipanel'}));
    if cls
        % Given handles of one or more axes, so isolate them from the rest
        fig = isolate_axes(fig);
    else
        % Check we have a figure
        if ~isequal(get(fig, 'Type'), 'figure')
            error('export_fig:BadHandle','Handle must be that of a figure, axes or uipanel');
        end
        % Get the old InvertHardcopy mode
        old_mode = get(fig, 'InvertHardcopy');
    end
    % from this point onward, fig is assured to be a figure handle

    % Hack the font units where necessary (due to a font rendering bug in print?).
    % This may not work perfectly in all cases.
    % Also it can change the figure layout if reverted, so use a copy.
    magnify = options.magnify * options.aa_factor;
    if isbitmap(options) && magnify ~= 1
        fontu = findall(fig, 'FontUnits', 'normalized');
        if ~isempty(fontu)
            % Some normalized font units found
            if ~cls
                fig = copyfig(fig);
                set(fig, 'Visible', 'off');
                fontu = findall(fig, 'FontUnits', 'normalized');
                cls = true;
            end
            set(fontu, 'FontUnits', 'points');
        end
    end

    try
        % MATLAB "feature": axes limits and tick marks can change when printing
        Hlims = findall(fig, 'Type', 'axes');
        if ~cls
            % Record the old axes limit and tick modes
            Xlims = make_cell(get(Hlims, 'XLimMode'));
            Ylims = make_cell(get(Hlims, 'YLimMode'));
            Zlims = make_cell(get(Hlims, 'ZLimMode'));
            Xtick = make_cell(get(Hlims, 'XTickMode'));
            Ytick = make_cell(get(Hlims, 'YTickMode'));
            Ztick = make_cell(get(Hlims, 'ZTickMode'));
            Xlabel = make_cell(get(Hlims, 'XTickLabelMode')); 
            Ylabel = make_cell(get(Hlims, 'YTickLabelMode')); 
            Zlabel = make_cell(get(Hlims, 'ZTickLabelMode')); 
        end

        % Set all axes limit and tick modes to manual, so the limits and ticks can't change
        % Fix Matlab R2014b bug (issue #34): plot markers are not displayed when ZLimMode='manual'
        set(Hlims, 'XLimMode', 'manual', 'YLimMode', 'manual');
        set_tick_mode(Hlims, 'X');
        set_tick_mode(Hlims, 'Y');
        if ~using_hg2(fig)
            set(Hlims,'ZLimMode', 'manual');
            set_tick_mode(Hlims, 'Z');
        end
    catch
        % ignore - fix issue #4 (using HG2 on R2014a and earlier)
    end

    % Fix issue #21 (bold TeX axes labels/titles in R2014b when exporting to EPS/PDF)
    try
        if using_hg2(fig) && isvector(options)
            % Set the FontWeight of axes labels/titles to 'normal'
            % Fix issue #69: set non-bold font only if the string contains symbols (\beta etc.)
            % Issue #243: only set non-bold font (workaround for issue #69) in R2015b or earlier
            try isPreR2016a = verLessThan('matlab','8.7'); catch, isPreR2016a = true; end
            if isPreR2016a
                texLabels = findall(fig, 'type','text', 'FontWeight','bold');
                symbolIdx = ~cellfun('isempty',strfind({texLabels.String},'\'));
                if ~isempty(symbolIdx)
                    set(texLabels(symbolIdx), 'FontWeight','normal');
                    if ~options.silent
                        warning('export_fig:BoldTexLabels', 'Bold labels with Tex symbols converted into non-bold in export_fig (fix for issue #69)');
                    end
                end
            end
        end
    catch
        % ignore
    end

    % Fix issue #42: non-normalized annotations on HG1 (internal Matlab bug)
    annotationHandles = [];
    try
        if ~using_hg2(fig)
            annotationHandles = findall(fig,'Type','hggroup','-and','-property','Units','-and','-not','Units','norm');
            try  % suggested by Jesús Pestana Puerta (jespestana) 30/9/2015
                originalUnits = get(annotationHandles,'Units');
                set(annotationHandles,'Units','norm');
            catch
            end
        end
    catch
        % should never happen, but ignore in any case - issue #50
    end

    % Fix issue #46: Ghostscript crash if figure units <> pixels
    pos = get(fig, 'Position');  % Fix issues #313, #314
    oldFigUnits = get(fig,'Units');
    set(fig,'Units','pixels');
    pixelpos = get(fig, 'Position'); %=getpixelposition(fig);

    tcol = get(fig, 'Color');
    tcol_orig = tcol;

    % Set to print exactly what is there
    if options.invert_hardcopy
        try set(fig, 'InvertHardcopy', 'off'); catch, end  % fail silently in uifigures
    end

    % Set the renderer
    switch options.renderer
        case 1
            renderer = '-opengl';
        case 2
            renderer = '-zbuffer';
        case 3
            renderer = '-painters';
        otherwise
            renderer = '-opengl'; % Default for bitmaps
    end

    try
        tmp_nam = '';  % initialize

        % Do the bitmap formats first
        if isbitmap(options)
            if abs(options.bb_padding) > 1
                displaySuggestedWorkarounds = false;
                error('export_fig:padding','For bitmap output (png,jpg,tif,bmp) the padding value (-p) must be between -1<p<1')
            end
            % Print large version to array
            [A, tcol, alpha] = getFigImage(fig, magnify, renderer, options, pixelpos);
            % Get the background colour
            if options.transparent && (options.png || options.alpha)
                try %options.aa_factor < 4  % default, faster but lines are not anti-aliased
                    % If all pixels are indicated as opaque (i.e. something went wrong with the Java screen-capture)
                    isBgColor = A(:,:,1) == tcol(1) & ...
                                A(:,:,2) == tcol(2) & ...
                                A(:,:,3) == tcol(3);
                    % Set the bgcolor pixels to be fully-transparent
                    A(repmat(isBgColor,[1,1,3])) = 255; %=white % TODO: more memory efficient without repmat
                    alpha(isBgColor) = 0;
                catch  % older logic - much slower and causes figure flicker
                    if true  % to fold the code below...
                    % Get out an alpha channel
                    % MATLAB "feature": black colorbar axes can change to white and vice versa!
                    hCB = findall(fig, 'Type','axes', 'Tag','Colorbar');
                    if isempty(hCB)
                        yCol = [];
                        xCol = [];
                    else
                        yCol = get(hCB, 'YColor');
                        xCol = get(hCB, 'XColor');
                        if iscell(yCol)
                            yCol = cell2mat(yCol);
                            xCol = cell2mat(xCol);
                        end
                        yCol = sum(yCol, 2);
                        xCol = sum(xCol, 2);
                    end
                    % MATLAB "feature": apparently figure size can change when changing
                    % colour in -nodisplay mode
                    % Set the background colour to black, and set size in case it was
                    % changed internally
                    set(fig, 'Color', 'k', 'Position', pos);
                    % Correct the colorbar axes colours
                    set(hCB(yCol==0), 'YColor', [0 0 0]);
                    set(hCB(xCol==0), 'XColor', [0 0 0]);
                    % Correct black axes color to off-black (issue #249)
                    hAxes = findall(fig, 'Type','axes');
                    [hXs,hXrs] = fixBlackAxle(hAxes, 'XColor');
                    [hYs,hYrs] = fixBlackAxle(hAxes, 'YColor');
                    [hZs,hZrs] = fixBlackAxle(hAxes, 'ZColor');

                    % The following code might cause out-of-memory errors
                    try
                        % Print large version to array
                        B = print2array(fig, magnify, renderer);
                        % Downscale the image
                        B = downsize(single(B), options.aa_factor);
                    catch
                        % This is more conservative in memory, but kills transparency (issue #58)
                        B = single(print2array(fig, magnify/options.aa_factor, renderer));
                    end

                    % Set background to white (and set size)
                    set(fig, 'Color', 'w', 'Position', pos);
                    % Correct the colorbar axes colours
                    set(hCB(yCol==3), 'YColor', [1 1 1]);
                    set(hCB(xCol==3), 'XColor', [1 1 1]);
                    % Revert the black axes colors
                    set(hXs, 'XColor', [0,0,0]);
                    set(hYs, 'YColor', [0,0,0]);
                    set(hZs, 'ZColor', [0,0,0]);
                    set(hXrs, 'Color', [0,0,0]);
                    set(hYrs, 'Color', [0,0,0]);
                    set(hZrs, 'Color', [0,0,0]);

                    % The following code might cause out-of-memory errors
                    try
                        % Print large version to array
                        A = print2array(fig, magnify, renderer);
                        % Downscale the image
                        A = downsize(single(A), options.aa_factor);
                    catch
                        % This is more conservative in memory, but kills transparency (issue #58)
                        A = single(print2array(fig, magnify/options.aa_factor, renderer));
                    end

                    % Workaround for issue #15
                    szA = size(A);
                    szB = size(B);
                    if ~isequal(szA,szB)
                        A = A(1:min(szA(1),szB(1)), 1:min(szA(2),szB(2)), :);
                        B = B(1:min(szA(1),szB(1)), 1:min(szA(2),szB(2)), :);
                        if ~options.silent
                            warning('export_fig:bitmap:sizeMismatch','Problem detected by export_fig generation of a bitmap image; the generated export may look bad. Try to reduce the figure size to fit the screen, or avoid using export_fig''s -transparent option.')
                        end
                    end
                    % Compute the alpha map
                    alpha = round(sum(B - A, 3)) / (255 * 3) + 1;
                    A = alpha;
                    A(A==0) = 1;
                    A = B ./ A(:,:,[1 1 1]);
                    clear B
                    end %folded code...
                end
                %A = uint8(A);
            end
            % Downscale the image if its size was increased (for anti-aliasing)
            if size(A,1) > 1.1 * options.magnify * pixelpos(4) %1.1 to avoid edge-cases
                % Downscale the image
                A     = downsize(A,     options.aa_factor);
                alpha = downsize(alpha, options.aa_factor);
            end
            % Crop the margins based on the bgcolor, if requested
            if options.crop
                %[alpha, v] = crop_borders(alpha, 0, 1, options.crop_amounts);
                %A = A(v(1):v(2),v(3):v(4),:);
                [A, vA, vB] = crop_borders(A, tcol, options.bb_padding, options.crop_amounts);
                if ~any(isnan(vB)) % positive padding
                    sz = size(A); % Fix issue #308
                    B = repmat(uint8(zeros(1,1,size(alpha,3))),sz([1,2])); % Fix issue #307 %=zeros(sz([1,2]),'uint8');
                    B(vB(1):vB(2), vB(3):vB(4), :) = alpha(vA(1):vA(2), vA(3):vA(4), :); % ADDED BY OH
                    alpha = B;
                else  % negative padding
                    alpha = alpha(vA(1):vA(2), vA(3):vA(4), :);
                end
            end
            % Get the non-alpha image (presumably unneeded with Java-based screen-capture)
            %{
            if isbitmap(options)
                % Modify the intensity of the pixels' RGB values based on their alpha transparency
                % TODO: not sure that we want this with Java screen-capture values!
                alph = alpha(:,:,ones(1, size(A, 3)));
                A = uint8(single(A) .* alph + 255 * (1 - alph));
            end
            %}
            % Revert the figure properties back to their original values
            set(fig, 'Units',oldFigUnits, 'Position',pos, 'Color',tcol_orig);
            % Check for greyscale images
            if options.colourspace == 2
                % Convert to greyscale
                A = rgb2grey(A);
            else
                % Return only one channel for greyscale
                A = check_greyscale(A);
            end
            % Change alpha from [0:255] uint8 => [0:1] single from here onward:
            alpha = single(alpha) / 255;
            % Outputs
            if options.im
                imageData = A;
            end
            if options.alpha
                imageData = A;
                %alpha = ones(size(A, 1), size(A, 2), 'single');  %=all pixels opaque
            end
            % Save the images
            if options.png
                % Compute the resolution
                res = options.magnify * get(0, 'ScreenPixelsPerInch') / 25.4e-3;
                % Save the png
                [format_options, bitDepth] = getFormatOptions(options, 'png');  %Issue #269
                pngOptions = {[options.name '.png'], 'ResolutionUnit','meter', 'XResolution',res, 'YResolution',res, format_options{:}}; %#ok<CCAT>
                if options.transparent  % Fix issue #312: only use alpha channel if -transparent was requested
                    pngOptions = [pngOptions 'Alpha',double(alpha)];
                end
                if ~isempty(bitDepth) && bitDepth < 16 && size(A,3) == 3
                    % BitDepth specification requires using a color-map
                    [A, map] = rgb2ind(A, 256);
                    imwrite(A, map, pngOptions{:});
                else
                    imwrite(A, pngOptions{:});
                end
            end
            if options.bmp
                imwrite(A, [options.name '.bmp']);
            end
            if options.jpg
                % Save jpeg with the specified quality
                quality = options.quality;
                if isempty(quality)
                    quality = 95;
                end
                format_options = getFormatOptions(options, 'jpg');  %Issue #269
                if quality > 100
                    imwrite(A, [options.name '.jpg'], 'Mode','lossless', format_options{:});
                else
                    imwrite(A, [options.name '.jpg'], 'Quality',quality, format_options{:});
                end
            end
            if options.tif
                % Save tif images in cmyk if wanted (and possible)
                if options.colourspace == 1 && size(A, 3) == 3
                    A = double(255 - A);
                    K = min(A, [], 3);
                    K_ = 255 ./ max(255 - K, 1);
                    C = (A(:,:,1) - K) .* K_;
                    M = (A(:,:,2) - K) .* K_;
                    Y = (A(:,:,3) - K) .* K_;
                    A = uint8(cat(3, C, M, Y, K));
                    clear C M Y K K_
                end
                append_mode = {'overwrite', 'append'};
                format_options = getFormatOptions(options, 'tif');  %Issue #269
                imwrite(A, [options.name '.tif'], 'Resolution',options.magnify*get(0,'ScreenPixelsPerInch'), 'WriteMode',append_mode{options.append+1}, format_options{:});
            end
        end

        % Now do the vector formats which are based on EPS
        if isvector(options)
            hImages = findall(fig,'type','image');
            % Set the default renderer to painters
            if ~options.renderer
                % Handle transparent patches
                hasTransparency = ~isempty(findall(fig,'-property','FaceAlpha','-and','-not','FaceAlpha',1));
                if hasTransparency
                    % Alert if trying to export transparent patches/areas to non-supported outputs (issue #108)
                    % http://www.mathworks.com/matlabcentral/answers/265265-can-export_fig-or-else-draw-vector-graphics-with-transparent-surfaces
                    % TODO - use transparency when exporting to PDF by not passing via print2eps
                    msg = 'export_fig currently supports transparent patches/areas only in PNG output. ';
                    if options.pdf && ~options.silent
                        warning('export_fig:transparency', '%s\nTo export transparent patches/areas to PDF, use the print command:\n print(gcf, ''-dpdf'', ''%s.pdf'');', msg, options.name);
                    elseif ~options.png && ~options.tif && ~options.silent  % issue #168
                        warning('export_fig:transparency', '%s\nTo export the transparency correctly, try using the ScreenCapture utility on the Matlab File Exchange: http://bit.ly/1QFrBip', msg);
                    end
                elseif ~isempty(hImages)
                    % Fix for issue #230: use OpenGL renderer when exported image contains transparency
                    for idx = 1 : numel(hImages)
                        cdata = get(hImages(idx),'CData');
                        if any(isnan(cdata(:)))
                            hasTransparency = true;
                            break
                        end
                    end
                end
                hasPatches = ~isempty(findall(fig,'type','patch'));
                if hasTransparency || hasPatches
                    % This is *MUCH* slower, but more accurate for patches and transparent annotations (issue #39)
                    renderer = '-opengl';
                else
                    renderer = '-painters';
                end
            end
            options.rendererStr = renderer;  % fix for issue #112
            % Generate some filenames
            tmp_nam = [tempname '.eps'];
            try
                % Ensure that the temp dir is writable (Javier Paredes 30/1/15)
                fid = fopen(tmp_nam,'w');
                fwrite(fid,1);
                fclose(fid);
                delete(tmp_nam);
                isTempDirOk = true;
            catch
                % Temp dir is not writable, so use the user-specified folder
                [dummy,fname,fext] = fileparts(tmp_nam); %#ok<ASGLU>
                fpath = fileparts(options.name);
                tmp_nam = fullfile(fpath,[fname fext]);
                isTempDirOk = false;
            end
            if isTempDirOk
                pdf_nam_tmp = [tempname '.pdf'];
            else
                pdf_nam_tmp = fullfile(fpath,[fname '.pdf']);
            end
            if options.pdf
                pdf_nam = [options.name '.pdf'];
                try copyfile(pdf_nam, pdf_nam_tmp, 'f'); catch, end  % fix for issue #65
            else
                pdf_nam = pdf_nam_tmp;
            end
            % Generate the options for print
            printArgs = {renderer};
            if ~isempty(options.resolution)  % issue #241
                printArgs{end+1} = sprintf('-r%d', options.resolution);
            end
            if options.colourspace == 1  % CMYK
                % Issue #33: due to internal bugs in Matlab's print() function, we can't use its -cmyk option
                %printArgs{end+1} = '-cmyk';
            end
            if ~options.crop
                % Issue #56: due to internal bugs in Matlab's print() function, we can't use its internal cropping mechanism,
                % therefore we always use '-loose' (in print2eps.m) and do our own cropping (in crop_borders)
                %printArgs{end+1} = '-loose';
            end
            if any(strcmpi(varargin,'-depsc'))
                % Issue #45: lines in image subplots are exported in invalid color.
                % The workaround is to use the -depsc parameter instead of the default -depsc2
                printArgs{end+1} = '-depsc';
            end
            % Print to EPS file
            try
                % Remove background if requested (issue #207)
                originalBgColor = get(fig, 'Color');
                [hXs, hXrs, hYs, hYrs, hZs, hZrs] = deal([]);
                if options.transparent %&& ~isequal(get(fig, 'Color'), 'none')
                    if options.renderer == 1 && ~options.silent  % OpenGL
                        warning('export_fig:openglTransparentBG', '-opengl sometimes fails to produce transparent backgrounds; in such a case, try to use -painters instead');
                    end

                    % Fix for issue #207, #267 (corrected)
                    set(fig,'Color','none');

                    % Correct black axes color to off-black (issue #249)
                    hAxes = findall(fig, 'Type','axes');
                    [hXs,hXrs] = fixBlackAxle(hAxes, 'XColor');
                    [hYs,hYrs] = fixBlackAxle(hAxes, 'YColor');
                    [hZs,hZrs] = fixBlackAxle(hAxes, 'ZColor');

                    % Correct black titles to off-black
                    % https://www.mathworks.com/matlabcentral/answers/567027-matlab-export_fig-crops-title
                    try
                        hTitle = get(hAxes, 'Title');
                        for idx = numel(hTitle) : -1 : 1
                            color = get(hTitle,'Color');
                            if isequal(color,[0,0,0]) || isequal(color,'k')
                                set(hTitle(idx), 'Color', [0,0,0.01]); %off-black
                            else
                                hTitle(idx) = [];  % remove from list
                            end
                        end
                    catch
                        hTitle = [];
                    end
                end
                % Generate an eps
                print2eps(tmp_nam, fig, options, printArgs{:});
                % {
                % Remove the background, if desired
                if options.transparent %&& ~isequal(get(fig, 'Color'), 'none')
                    eps_remove_background(tmp_nam, 1 + using_hg2(fig));

                    % Revert the black axes colors
                    set(hXs, 'XColor', [0,0,0]);
                    set(hYs, 'YColor', [0,0,0]);
                    set(hZs, 'ZColor', [0,0,0]);
                    set(hXrs, 'Color', [0,0,0]);
                    set(hYrs, 'Color', [0,0,0]);
                    set(hZrs, 'Color', [0,0,0]);
                    set(hTitle,'Color',[0,0,0]);
                end
                %}
                % Restore the figure's previous background color (if modified)
                try set(fig,'Color',originalBgColor); drawnow; catch, end
                % Fix colorspace to CMYK, if requested (workaround for issue #33)
                if options.colourspace == 1  % CMYK
                    % Issue #33: due to internal bugs in Matlab's print() function, we can't use its -cmyk option
                    change_rgb_to_cmyk(tmp_nam);
                end
                % Add a bookmark to the PDF if desired
                if options.bookmark
                    fig_nam = get(fig, 'Name');
                    if isempty(fig_nam) && ~options.silent
                        warning('export_fig:EmptyBookmark', 'Bookmark requested for figure with no name. Bookmark will be empty.');
                    end
                    add_bookmark(tmp_nam, fig_nam);
                end
                % Generate a pdf
                eps2pdf(tmp_nam, pdf_nam_tmp, 1, options.append, options.colourspace==2, options.quality, options.gs_options);
                % Ghostscript croaks on % chars in the output PDF file, so use tempname and then rename the file
                try
                    % Rename the file (except if it is already the same)
                    % Abbie K's comment on the commit for issue #179 (#commitcomment-20173476)
                    if ~isequal(pdf_nam_tmp, pdf_nam)
                        movefile(pdf_nam_tmp, pdf_nam, 'f');
                    end
                catch
                    % Alert in case of error creating output PDF/EPS file (issue #179)
                    if exist(pdf_nam_tmp, 'file')
                        errMsg = ['Could not create ' pdf_nam ' - perhaps the folder does not exist, or you do not have write permissions, or the file is open in another application'];
                        error('export_fig:PDF:create',errMsg);
                    else
                        error('export_fig:NoEPS','Could not generate the intermediary EPS file.');
                    end
                end
            catch ex
                % Restore the figure's previous background color (in case it was not already restored)
                try set(fig,'Color',originalBgColor); drawnow; catch, end
                % Delete the temporary eps file - NOT! (Yair 3/3/2020)
                %delete(tmp_nam);
                % Rethrow the EPS/PDF-generation error
                rethrow(ex);
            end
            % Delete the eps
            delete(tmp_nam);
            if options.eps || options.linecaps
                try
                    % Generate an eps from the pdf
                    % since pdftops can't handle relative paths (e.g., '..\'), use a temp file
                    eps_nam_tmp = strrep(pdf_nam_tmp,'.pdf','.eps');
                    pdf2eps(pdf_nam, eps_nam_tmp);

                    % Issue #192: enable rounded line-caps
                    if options.linecaps
                        fstrm = read_write_entire_textfile(eps_nam_tmp);
                        fstrm = regexprep(fstrm, '[02] J', '1 J');
                        read_write_entire_textfile(eps_nam_tmp, fstrm);
                        if options.pdf
                            eps2pdf(eps_nam_tmp, pdf_nam, 1, options.append, options.colourspace==2, options.quality, options.gs_options);
                        end
                    end

                    if options.eps
                        movefile(eps_nam_tmp, [options.name '.eps'], 'f');
                    else  % if options.pdf
                        try delete(eps_nam_tmp); catch, end
                    end
                catch ex
                    if ~options.pdf
                        % Delete the pdf
                        delete(pdf_nam);
                    end
                    try delete(eps_nam_tmp); catch, end
                    rethrow(ex);
                end
                if ~options.pdf
                    % Delete the pdf
                    delete(pdf_nam);
                end
            end
            % Issue #206: warn if the figure contains an image
            if ~isempty(hImages) && strcmpi(renderer,'-opengl') && ~options.silent  % see addendum to issue #206
                warnMsg = ['exporting images to PDF/EPS may result in blurry images on some viewers. ' ...
                           'If so, try to change viewer, or increase the image''s CData resolution, or use -opengl renderer, or export via the print function. ' ...
                           'See ' hyperlink('https://github.com/altmany/export_fig/issues/206', 'issue #206') ' for details.'];
                warning('export_fig:pdf_eps:blurry_image', warnMsg);
            end
        end

        % SVG format
        if options.svg
            filename = [options.name '.svg'];
            % Adapted from Dan Joshea's https://github.com/djoshea/matlab-save-figure :
            try %if verLessThan('matlab', '8.4')
                % Try using the fig2svg/plot2svg utilities
                try
                    fig2svg(filename, fig);  %https://github.com/kupiqu/fig2svg
                catch
                    plot2svg(filename, fig); %https://github.com/jschwizer99/plot2svg
                    if ~options.silent
                        warning('export_fig:SVG:plot2svg', 'export_fig used the plot2svg utility for SVG output. Better results may be gotten via the fig2svg utility (https://github.com/kupiqu/fig2svg).');
                    end
                end
            catch %else  % (neither fig2svg nor plot2svg are available)
                % Try Matlab's built-in svg engine (from Batik Graphics2D for java)
                try
                    set(fig,'Units','pixels');   % All data in the svg-file is saved in pixels
                    printArgs = {renderer};
                    if ~isempty(options.resolution)
                        printArgs{end+1} = sprintf('-r%d', options.resolution);
                    end
                    print(fig, '-dsvg', printArgs{:}, filename);
                    if ~options.silent
                        warning('export_fig:SVG:print', 'export_fig used Matlab''s built-in SVG output engine. Better results may be gotten via the fig2svg utility (https://github.com/kupiqu/fig2svg).');
                    end
                catch err  % built-in print() failed - maybe an old Matlab release (no -dsvg)
                    filename = strrep(filename,'export_fig_out','filename');
                    msg = ['SVG output is not supported for your figure: ' err.message '\n' ...
                        'Try one of the following alternatives:\n' ...
                        '  1. saveas(gcf,''' filename ''')\n' ...
                        '  2. fig2svg utility: https://github.com/kupiqu/fig2svg\n' ...  % Note: replaced defunct https://github.com/jschwizer99/plot2svg with up-to-date fork on https://github.com/kupiqu/fig2svg
                        '  3. export_fig to EPS/PDF, then convert to SVG using non-Matlab tools\n'];
                    error('export_fig:SVG:error',msg);
                end
            end
            % SVG output was successful if we reached this point
            % Add warning about unsupported export_fig options with SVG output
            if ~options.silent && (any(~isnan(options.crop_amounts)) || any(options.bb_padding))
                warning('export_fig:SVG:options', 'export_fig''s SVG output does not [currently] support cropping/padding.');
            end

            % Fix the generated SVG file, based on Cris Luengo's SVG_FIX_VIEWBOX:
            % https://www.mathworks.com/matlabcentral/fileexchange/49617-svg_fix_viewbox-in_name-varargin
            try
                % Read SVG file
                s = read_write_entire_textfile(filename);
                % Fix fonts #1: 'SansSerif' doesn't work on my browser, the correct CSS is 'sans-serif'
                s = regexprep(s,'font-family:SansSerif;|font-family:''SansSerif'';','font-family:''sans-serif'';');
                % Fix fonts #1: The document-wide default font is 'Dialog'. What is this anyway?
                s = regexprep(s,'font-family:''Dialog'';','font-family:''sans-serif'';');
                % Replace 'width="xxx" height="yyy"' with 'width="100%" viewBox="0 0 xxx yyy"'
                t = regexp(s,'<svg.* width="(?<width>[0-9]*)" height="(?<height>[0-9]*)"','names');
                if ~isempty(t)
                    relativeWidth = 100;  %TODO - user-settable via input parameter?
                    s = regexprep(s,'(?<=<svg[^\n]*) width="[0-9]*" height="[0-9]*"',sprintf(' width="%d\\%%" viewBox="0 0 %s %s"',relativeWidth,t.width,t.height));
                end
                % Write updated SVG file
                read_write_entire_textfile(filename, s);
            catch
                % never mind - ignore
            end
        end

        % EMF format
        if options.emf
            try
                anythingChanged = false;
                % Handle transparent bgcolor request
                if options.transparent && ~isequal(tcol_orig,'none')
                    anythingChanged = true;
                    set(fig, 'Color','none');
                end
                if ~options.silent
                    if ~ispc
                        warning('export_fig:EMF:NotWindows', 'EMF is only supported on Windows; exporting to EMF format on this machine may result in unexpected behavior.');
                    elseif isequal(renderer,'-painters') && (options.resolution~=864 || options.magnify~=1)
                        warning('export_fig:EMF:Painters', 'export_fig -r and -m options are ignored for EMF export using the -painters renderer.');
                    elseif abs(get(0,'ScreenPixelsPerInch')*options.magnify - options.resolution) > 1e-6
                        warning('export_fig:EMF:Magnify', 'export_fig -m option is ignored for EMF export.');
                    end
                    if ~isequal(options.bb_padding,0) || ~isempty(options.quality)
                        warning('export_fig:EMF:Options', 'export_fig cropping, padding and quality options are ignored for EMF export.');
                    end
                    if ~anythingChanged
                        warning('export_fig:EMF:print', 'For a figure without background transparency, export_fig uses Matlab''s built-in print(''-dmeta'') function without any extra processing, so try using it directly.');
                    end
                end
                printArgs = {renderer};
                if ~isempty(options.resolution)
                    printArgs{end+1} = sprintf('-r%d', options.resolution);
                end
                filename = [options.name '.emf'];
                print(fig, '-dmeta', printArgs{:}, filename);
            catch err  % built-in print() failed - maybe an old Matlab release (no -dsvg)
                msg = ['EMF output is not supported: ' err.message '\n' ...
                       'Try to use export_fig with other formats, such as PDF or EPS.\n'];
                error('export_fig:EMF:error',msg);
            end
        end

        % Revert the figure or close it (if requested)
        if cls || options.closeFig
            % Close the created figure
            close(fig);
        else
            % Reset the hardcopy mode
            try set(fig, 'InvertHardcopy', old_mode); catch, end  % fail silently in uifigures
            % Reset the axes limit and tick modes
            for a = 1:numel(Hlims)
                try
                    set(Hlims(a), 'XLimMode', Xlims{a}, 'YLimMode', Ylims{a}, 'ZLimMode', Zlims{a},... 
                                  'XTickMode', Xtick{a}, 'YTickMode', Ytick{a}, 'ZTickMode', Ztick{a},...
                                  'XTickLabelMode', Xlabel{a}, 'YTickLabelMode', Ylabel{a}, 'ZTickLabelMode', Zlabel{a}); 
                catch
                    % ignore - fix issue #4 (using HG2 on R2014a and earlier)
                end
            end
            % Revert the tex-labels font weights
            try set(texLabels, 'FontWeight','bold'); catch, end
            % Revert annotation units
            for handleIdx = 1 : numel(annotationHandles)
                try
                    oldUnits = originalUnits{handleIdx};
                catch
                    oldUnits = originalUnits;
                end
                try set(annotationHandles(handleIdx),'Units',oldUnits); catch, end
            end
            % Revert figure properties in case they were changed
            try set(fig, 'Units',oldFigUnits, 'Position',pos, 'Color',tcol_orig); catch, end
        end

        % Output to clipboard (if requested)
        if options.clipboard
            % Delete the output file if unchanged from the default name ('export_fig_out.png')
            if strcmpi(options.name,'export_fig_out')
                try
                    fileInfo = dir('export_fig_out.png');
                    if ~isempty(fileInfo)
                        timediff = now - fileInfo.datenum;
                        ONE_SEC = 1/24/60/60;
                        if timediff < ONE_SEC
                            delete('export_fig_out.png');
                        end
                    end
                catch
                    % never mind...
                end
            end

            % Use Java clipboard by default
            if strcmpi(options.clipformat,'image')
                % Save the image in the system clipboard
                % credit: Jiro Doke's IMCLIPBOARD: http://www.mathworks.com/matlabcentral/fileexchange/28708-imclipboard
                try
                    error(javachk('awt', 'export_fig -clipboard output'));
                catch
                    if ~options.silent
                        warning('export_fig:clipboardJava', 'export_fig -clipboard output failed: requires Java to work');
                    end
                    return;
                end
                try
                    % Import necessary Java classes
                    import java.awt.Toolkit                 %#ok<SIMPT>
                    import java.awt.image.BufferedImage     %#ok<SIMPT>
                    import java.awt.datatransfer.DataFlavor %#ok<SIMPT>

                    % Get System Clipboard object (java.awt.Toolkit)
                    cb = Toolkit.getDefaultToolkit.getSystemClipboard();

                    % Add java class (ImageSelection) to the path
                    if ~exist('ImageSelection', 'class')
                        javaaddpath(fileparts(which(mfilename)), '-end');
                    end

                    % Get image size
                    ht = size(imageData, 1);
                    wd = size(imageData, 2);

                    % Convert to Blue-Green-Red format
                    try
                        imageData2 = imageData(:, :, [3 2 1]);
                    catch
                        % Probably gray-scaled image (2D, without the 3rd [RGB] dimension)
                        imageData2 = imageData(:, :, [1 1 1]);
                    end

                    % Convert to 3xWxH format
                    imageData2 = permute(imageData2, [3, 2, 1]);

                    % Append Alpha data (unused - transparency is not supported in clipboard copy)
                    alphaData2 = uint8(permute(255*alpha,[3,2,1])); %=255*ones(1,wd,ht,'uint8')
                    imageData2 = cat(1, imageData2, alphaData2);

                    % Create image buffer
                    imBuffer = BufferedImage(wd, ht, BufferedImage.TYPE_INT_RGB);
                    imBuffer.setRGB(0, 0, wd, ht, typecast(imageData2(:), 'int32'), 0, wd);

                    % Create ImageSelection object from the image buffer
                    imSelection = ImageSelection(imBuffer);

                    % Set clipboard content to the image
                    cb.setContents(imSelection, []);
                catch
                    if ~options.silent
                        warning('export_fig:clipboardFailed', 'export_fig -clipboard output failed: %s', lasterr); %#ok<LERR>
                    end
                end
            else  % use one of print()'s builtin clipboard formats
                % Remove background if requested (EMF format only)
                if options.transparent && strcmpi(options.clipformat,'meta')
                    % Set figure bgcolor to none
                    originalBgColor = get(fig, 'Color');
                    set(fig,'Color','none');

                    % Set axes bgcolor to none
                    hAxes = findall(fig, 'Type','axes');
                    originalAxColor = get(hAxes, 'Color');
                    set(hAxes,'Color','none');

                    drawnow;  %repaint before export
                end

                % Call print() to create the clipboard output
                clipformat = ['-d' options.clipformat];
                printArgs = {renderer};
                if ~isempty(options.resolution)
                    printArgs{end+1} = sprintf('-r%d', options.resolution);
                end
                print(fig, '-clipboard', clipformat, printArgs{:});

                % Restore the original background color
                try set(fig,   'Color',originalBgColor); catch, end
                try set(hAxes, 'Color',originalAxColor); catch, end
                drawnow; 
            end
        end

        % Don't output the data to console unless requested
        if ~nargout
            clear imageData alpha
        end
    catch err
        % Revert figure properties in case they were changed
        try set(fig,'Units',oldFigUnits, 'Position',pos, 'Color',tcol_orig); catch, end
        % Display possible workarounds before the error message
        if displaySuggestedWorkarounds && ~strcmpi(err.message,'export_fig error')
            isNewerVersionAvailable = checkForNewerVersion(currentVersion);  % alert if a newer version exists
            if isempty(regexpi(err.message,'Ghostscript'))
                fprintf(2, 'export_fig error. ');
            end
            fprintf(2, 'Please ensure:\n');
            fprintf(2, ' * that the function you used (%s.m) version %s is from the expected location\n', mfilename('fullpath'), num2str(currentVersion));
            paths = which(mfilename,'-all');
            if iscell(paths) && numel(paths) > 1
                fprintf(2, '    (you appear to have %s of export_fig installed)\n', hyperlink('matlab:which export_fig -all','multiple versions'));
            end
            if isNewerVersionAvailable
                fprintf(2, ' * and that you are using the %s of export_fig (you are not: run %s to update it)\n', ...
                        hyperlink('https://github.com/altmany/export_fig/archive/master.zip','latest version'), ...
                        hyperlink('matlab:export_fig(''-update'')','export_fig(''-update'')'));
            end
            fprintf(2, ' * and that you did not made a mistake in export_fig''s %s\n', hyperlink('matlab:help export_fig','expected input arguments'));
            if isvector(options)
                if ismac
                    url = 'http://pages.uoregon.edu/koch';
                else
                    url = 'http://ghostscript.com';
                end
                fpath = user_string('ghostscript');
                fprintf(2, ' * and that %s is properly installed in %s\n', ...
                        hyperlink(url,'ghostscript'), ...
                        hyperlink(['matlab:winopen(''' fileparts(fpath) ''')'], fpath));
            end
            try
                if options.eps
                    fpath = user_string('pdftops');
                    fprintf(2, ' * and that %s is properly installed in %s\n', ...
                            hyperlink('http://xpdfreader.com/download.html','pdftops'), ...
                            hyperlink(['matlab:winopen(''' fileparts(fpath) ''')'], fpath));
                end
            catch
                % ignore - probably an error in parse_args
            end
            try
                % Alert per issue #149
                if ~strncmpi(get(0,'Units'),'pixel',5)
                    fprintf(2, ' * or try to set groot''s Units property back to its default value of ''pixels'' (%s)\n', hyperlink('https://github.com/altmany/export_fig/issues/149','details'));
                end
            catch
                % ignore - maybe an old MAtlab release
            end
            fprintf(2, '\nIf the problem persists, then please %s.\n', hyperlink('https://github.com/altmany/export_fig/issues','report a new issue'));
            if exist(tmp_nam,'file')
                fprintf(2, 'In your report, please upload the problematic EPS file: %s (you can then delete this file).\n', tmp_nam);
            end
            fprintf(2, '\n');
        end
        rethrow(err)
    end
end

function options = default_options()
    % Default options used by export_fig
    options = struct(...
        'name',            'export_fig_out', ...
        'crop',            true, ...
        'crop_amounts',    nan(1,4), ...  % auto-crop all 4 image sides
        'transparent',     false, ...
        'renderer',        0, ...         % 0: default, 1: OpenGL, 2: ZBuffer, 3: Painters
        'pdf',             false, ...
        'eps',             false, ...
        'emf',             false, ...
        'svg',             false, ...
        'png',             false, ...
        'tif',             false, ...
        'jpg',             false, ...
        'bmp',             false, ...
        'clipboard',       false, ...
        'clipformat',      'image', ...
        'colourspace',     0, ...         % 0: RGB/gray, 1: CMYK, 2: gray
        'append',          false, ...
        'im',              false, ...
        'alpha',           false, ...
        'aa_factor',       0, ...
        'bb_padding',      0, ...
        'magnify',         [], ...
        'resolution',      [], ...
        'bookmark',        false, ...
        'closeFig',        false, ...
        'quality',         [], ...
        'update',          false, ...
        'version',         false, ...
        'fontswap',        true, ...
        'font_space',      '', ...
        'linecaps',        false, ...
        'invert_hardcopy', true, ...
        'format_options',  struct, ...
        'preserve_size',   false, ...
        'silent',          false, ...
        'gs_options',      {{}});
end

function [fig, options] = parse_args(nout, fig, argNames, varargin)
    % Parse the input arguments

    % Convert strings => chars
    varargin = cellfun(@str2char,varargin,'un',false);

    % Set the defaults
    native = false; % Set resolution to native of an image
    options = default_options();
    options.im =    (nout == 1);  % user requested imageData output
    options.alpha = (nout == 2);  % user requested alpha output
    options.handleName = '';  % default handle name

    % Go through the other arguments
    skipNext = false;
    for a = 1:nargin-3
        if skipNext
            skipNext = false;
            continue;
        end
        if all(ishandle(varargin{a}))
            fig = varargin{a};
            options.handleName = argNames{a};
        elseif ischar(varargin{a}) && ~isempty(varargin{a})
            if varargin{a}(1) == '-'
                switch lower(varargin{a}(2:end))
                    case 'nocrop'
                        options.crop = false;
                        options.crop_amounts = [0,0,0,0];
                    case {'trans', 'transparent'}
                        options.transparent = true;
                    case 'opengl'
                        options.renderer = 1;
                    case 'zbuffer'
                        options.renderer = 2;
                    case 'painters'
                        options.renderer = 3;
                    case 'pdf'
                        options.pdf = true;
                    case 'eps'
                        options.eps = true;
                    case {'emf','meta'}
                        options.emf = true;
                    case 'svg'
                        options.svg = true;
                    case 'png'
                        options.png = true;
                    case {'tif', 'tiff'}
                        options.tif = true;
                    case {'jpg', 'jpeg'}
                        options.jpg = true;
                    case 'bmp'
                        options.bmp = true;
                    case 'rgb'
                        options.colourspace = 0;
                    case 'cmyk'
                        options.colourspace = 1;
                    case {'gray', 'grey'}
                        options.colourspace = 2;
                    case {'a1', 'a2', 'a3', 'a4'}
                        options.aa_factor = str2double(varargin{a}(3));
                    case 'append'
                        options.append = true;
                    case 'bookmark'
                        options.bookmark = true;
                    case 'native'
                        native = true;
                    case {'clipboard','clipboard:image'}
                        options.clipboard = true;
                        options.clipformat = 'image';
                        options.im    = true;  %ensure that imageData is created
                        options.alpha = true;
                    case 'clipboard:bitmap'
                        options.clipboard = true;
                        options.clipformat = 'bitmap';
                    case {'clipboard:emf','clipboard:meta'}
                        options.clipboard = true;
                        options.clipformat = 'meta';
                    case 'clipboard:pdf'
                        options.clipboard = true;
                        options.clipformat = 'pdf';
                    case 'update'
                        updateInstalledVersion();
                        fig = -1;  % silent bail-out
                    case 'version'
                        options.version = true;
                        return  % ignore any additional args
                    case 'nofontswap'
                        options.fontswap = false;
                    case 'font_space'
                        options.font_space = varargin{a+1};
                        skipNext = true;
                    case 'linecaps'
                        options.linecaps = true;
                    case 'noinvert'
                        options.invert_hardcopy = false;
                    case 'preserve_size'
                        options.preserve_size = true;
                    case 'options'
                        % Issue #269: format-specific options
                        inputOptions = varargin{a+1};
                        %options.format_options  = inputOptions;
                        if isempty(inputOptions), continue, end
                        formats = fieldnames(inputOptions(1));
                        for idx = 1 : numel(formats)
                            optionsStruct = inputOptions.(formats{idx});
                            %optionsCells = [fieldnames(optionsStruct) struct2cell(optionsStruct)]';
                            formatName = regexprep(lower(formats{idx}),{'tiff','jpeg'},{'tif','jpg'});
                            options.format_options.(formatName) = optionsStruct; %=optionsCells(:)';
                        end
                        skipNext = true;
                    case 'silent'
                        options.silent = true;
                    otherwise
                        try
                            wasError = false;
                            if strcmpi(varargin{a}(1:2),'-d')
                                varargin{a}(2) = 'd';  % ensure lowercase 'd'
                                options.gs_options{end+1} = varargin{a};
                            elseif strcmpi(varargin{a}(1:2),'-c')
                                if strncmpi(varargin{a},'-clipboard:',11)
                                    wasError = true;
                                    error('export_fig:BadOptionValue','option ''%s'' cannot be parsed: only image, bitmap, emf and pdf formats are supported',varargin{a});
                                end
                                if numel(varargin{a})==2
                                    skipNext = true;
                                    vals = str2num(varargin{a+1}); %#ok<ST2NM>
                                else
                                    vals = str2num(varargin{a}(3:end)); %#ok<ST2NM>
                                end
                                if numel(vals)~=4
                                    wasError = true;
                                    error('export_fig:BadOptionValue','option -c cannot be parsed: must be a 4-element numeric vector');
                                end
                                options.crop_amounts = vals;
                                options.crop = true;
                            else  % scalar parameter value
                                val = str2double(regexp(varargin{a}, '(?<=-(m|M|r|R|q|Q|p|P))-?\d*.?\d+', 'match'));
                                if isempty(val) || isnan(val)
                                    % Issue #51: improved processing of input args (accept space between param name & value)
                                    val = str2double(varargin{a+1});
                                    if isscalar(val) && ~isnan(val)
                                        skipNext = true;
                                    end
                                end
                                if ~isscalar(val) || isnan(val)
                                    wasError = true;
                                    error('export_fig:BadOptionValue','option %s is not recognised or cannot be parsed', varargin{a});
                                end
                                switch lower(varargin{a}(2))
                                    case 'm'
                                        % Magnification may never be negative
                                        if val <= 0
                                            wasError = true;
                                            error('export_fig:BadMagnification','Bad magnification value: %g (must be positive)', val);
                                        end
                                        options.magnify = val;
                                    case 'r'
                                        options.resolution = val;
                                    case 'q'
                                        options.quality = max(val, 0);
                                    case 'p'
                                        options.bb_padding = val;
                                end
                            end
                        catch err
                            % We might have reached here by raising an intentional error
                            if wasError  % intentional raise
                                rethrow(err)
                            else  % unintentional
                                error('export_fig:BadOption',['Unrecognized export_fig input option: ''' varargin{a} '''']);
                            end
                        end
                end
            else
                [p, options.name, ext] = fileparts(varargin{a});
                if ~isempty(p)
                    % Issue #221: alert if the requested folder does not exist
                    if ~exist(p,'dir'),  error('export_fig:BadPath',['Folder ' p ' does not exist!']);  end
                    options.name = [p filesep options.name];
                end
                switch lower(ext)
                    case {'.tif', '.tiff'}
                        options.tif = true;
                    case {'.jpg', '.jpeg'}
                        options.jpg = true;
                    case '.png'
                        options.png = true;
                    case '.bmp'
                        options.bmp = true;
                    case '.eps'
                        options.eps = true;
                    case '.emf'
                        options.emf = true;
                    case '.pdf'
                        options.pdf = true;
                    case '.fig'
                        % If no open figure, then load the specified .fig file and continue
                        figFilename = varargin{a};
                        if isempty(fig)
                            fig = openfig(figFilename,'invisible');
                            varargin{a} = fig;
                            options.closeFig = true;
                            options.handleName = ['openfig(''' figFilename ''')'];
                        else
                            % save the current figure as the specified .fig file and exit
                            saveas(fig(1),figFilename);
                            fig = -1;
                            return
                        end
                    case '.svg'
                        options.svg = true;
                    otherwise
                        options.name = varargin{a};
                end
            end
        end
    end

    % Quick bail-out if no figure found
    if isempty(fig),  return;  end

    % Do border padding with repsect to a cropped image
    if options.bb_padding
        options.crop = true;
    end

    % Set default anti-aliasing now we know the renderer
    try isAA = strcmp(get(ancestor(fig, 'figure'), 'GraphicsSmoothing'), 'on'); catch, isAA = false; end
    if isAA
        if options.aa_factor > 1 && ~options.silent
            warning('export_fig:AntiAliasing','You requested anti-aliased export_fig output of a figure that is already anti-aliased - your -a option in export_fig is ignored.')
        end
        options.aa_factor = 1;  % ignore -a option when the figure is already anti-aliased (HG2)
    elseif options.aa_factor == 0  % default
        %options.aa_factor = 1 + 2 * (~(using_hg2(fig) && isAA) | (options.renderer == 3));
        options.aa_factor = 1 + 2 * (~using_hg2(fig));  % =1 in HG2, =3 in HG1
    end
    if options.aa_factor > 1 && ~isAA && using_hg2(fig) && ~options.silent
        warning('export_fig:AntiAliasing','You requested anti-aliased export_fig output of an aliased figure (''GraphicsSmoothing''=''off''). You will see better results if you set your figure''s GraphicsSmoothing property to ''on'' before calling export_fig.')
    end

    % Convert user dir '~' to full path
    if numel(options.name) > 2 && options.name(1) == '~' && (options.name(2) == '/' || options.name(2) == '\')
        options.name = fullfile(char(java.lang.System.getProperty('user.home')), options.name(2:end));
    end

    % Compute the magnification and resolution
    if isempty(options.magnify)
        if isempty(options.resolution)
            options.magnify = 1;
            options.resolution = 864;
        else
            options.magnify = options.resolution ./ get(0, 'ScreenPixelsPerInch');
        end
    elseif isempty(options.resolution)
        options.resolution = 864;
    end

    % Set the format to PNG, if no other format was specified
    if ~isvector(options) && ~isbitmap(options) && ~options.svg && ~options.emf
        options.png = true;
    end

    % Check whether transparent background is wanted (old way)
    if isequal(get(ancestor(fig(1), 'figure'), 'Color'), 'none')
        options.transparent = true;
    end

    % If requested, set the resolution to the native vertical resolution of the
    % first suitable image found
    if native
        if isbitmap(options)
            % Find a suitable image
            list = findall(fig, 'Type','image', 'Tag','export_fig_native');
            if isempty(list)
                list = findall(fig, 'Type','image', 'Visible','on');
            end
            for hIm = list(:)'
                % Check height is >= 2
                height = size(get(hIm, 'CData'), 1);
                if height < 2
                    continue
                end
                % Account for the image filling only part of the axes, or vice versa
                yl = get(hIm, 'YData');
                if isscalar(yl)
                    yl = [yl(1)-0.5 yl(1)+height+0.5];
                else
                    yl = [min(yl), max(yl)];  % fix issue #151 (case of yl containing more than 2 elements)
                    if ~diff(yl)
                        continue
                    end
                    yl = yl + [-0.5 0.5] * (diff(yl) / (height - 1));
                end
                hAx = get(hIm, 'Parent');
                yl2 = get(hAx, 'YLim');
                % Find the pixel height of the axes
                oldUnits = get(hAx, 'Units');
                set(hAx, 'Units', 'pixels');
                pos = get(hAx, 'Position');
                set(hAx, 'Units', oldUnits);
                if ~pos(4)
                    continue
                end
                % Found a suitable image
                % Account for stretch-to-fill being disabled
                pbar = get(hAx, 'PlotBoxAspectRatio');
                pos = min(pos(4), pbar(2)*pos(3)/pbar(1));
                % Set the magnification to give native resolution
                options.magnify = abs((height * diff(yl2)) / (pos * diff(yl)));  % magnification must never be negative: issue #103
                break
            end
        elseif options.resolution == 864  % don't use -r864 in vector mode if user asked for -native
            options.resolution = []; % issue #241 (internal Matlab bug produces black lines with -r864)
        end
    end
end

% Convert a possible string => char (issue #245)
function value = str2char(value)
    if isa(value,'string')
        value = char(value);
    end
end

function A = downsize(A, factor)
    % Downsample an image
    if factor <= 1 || isempty(A) %issue #310
        % Nothing to do
        return
    end
    try
        % Faster, but requires image processing toolbox
        A = imresize(A, 1/factor, 'bilinear');
    catch
        % No image processing toolbox - resize manually
        % Lowpass filter - use Gaussian as is separable, so faster
        % Compute the 1d Gaussian filter
        filt = (-factor-1:factor+1) / (factor * 0.6);
        filt = exp(-filt .* filt);
        % Normalize the filter
        filt = single(filt / sum(filt));
        % Filter the image
        padding = floor(numel(filt) / 2);
        if padding < 1 || isempty(A), return, end  %issue #310
        onesPad = ones(1, padding);
        for a = 1:size(A,3)
            A2 = single(A([onesPad 1:end repmat(end,1,padding)], ...
                          [onesPad 1:end repmat(end,1,padding)], a));
            A(:,:,a) = conv2(filt, filt', A2, 'valid');
        end
        % Subsample
        A = A(1+floor(mod(end-1, factor)/2):factor:end,1+floor(mod(end-1, factor)/2):factor:end,:);
    end
end

function A = rgb2grey(A)
    A = cast(reshape(reshape(single(A), [], 3) * single([0.299; 0.587; 0.114]), size(A, 1), size(A, 2)), class(A)); % #ok<ZEROLIKE>
end

function A = check_greyscale(A)
    % Check if the image is greyscale
    if size(A, 3) == 3 && ...
            all(reshape(A(:,:,1) == A(:,:,2), [], 1)) && ...
            all(reshape(A(:,:,2) == A(:,:,3), [], 1))
        A = A(:,:,1); % Save only one channel for 8-bit output
    end
end

function eps_remove_background(fname, count)
    % Remove the background of an eps file
    % Open the file
    fh = fopen(fname, 'r+');
    if fh == -1
        error('export_fig:EPS:open','Cannot open file %s.', fname);
    end
    % Read the file line by line
    while count
        % Get the next line
        l = fgets(fh);
        if isequal(l, -1)
            break; % Quit, no rectangle found
        end
        % Check if the line contains the background rectangle
        if isequal(regexp(l, ' *0 +0 +\d+ +\d+ +r[fe] *[\n\r]+', 'start'), 1)
            % Set the line to whitespace and quit
            l(1:regexp(l, '[\n\r]', 'start', 'once')-1) = ' ';
            fseek(fh, -numel(l), 0);
            fprintf(fh, l);
            % Reduce the count
            count = count - 1;
        end
    end
    % Close the file
    fclose(fh);
end

function b = isvector(options)
    b = options.pdf || options.eps;
end

function b = isbitmap(options)
    b = options.png || options.tif || options.jpg || options.bmp || options.im || options.alpha;
end

function [A, tcol, alpha] = getFigImage(fig, magnify, renderer, options, pos)
    if options.transparent
        % MATLAB "feature": figure size can change when changing color in -nodisplay mode
        set(fig, 'Color', 'w', 'Position', pos);
        drawnow;  % repaint figure, otherwise Java screencapture will see black bgcolor
    end
    % Print large version to array
    try
        % The following code might cause out-of-memory errors
        [A, tcol, alpha] = print2array(fig, magnify, renderer);
    catch
        % This is more conservative in memory, but perhaps kills transparency(?)
        [A, tcol, alpha] = print2array(fig, magnify/options.aa_factor, renderer);
    end
    % In transparent mode, set the bgcolor to white
    if options.transparent
        % Note: tcol should already be [255,255,255] here, but just in case it's not...
        tcol = uint8([255,255,255]);  %=white
    end
end

function A = make_cell(A)
    if ~iscell(A)
        A = {A};
    end
end

function add_bookmark(fname, bookmark_text)
    % Adds a bookmark to the temporary EPS file after %%EndPageSetup
    % Read in the file
    fh = fopen(fname, 'r');
    if fh == -1
        error('export_fig:bookmark:FileNotFound','File %s not found.', fname);
    end
    try
        fstrm = fread(fh, '*char')';
    catch ex
        fclose(fh);
        rethrow(ex);
    end
    fclose(fh);

    % Include standard pdfmark prolog to maximize compatibility
    fstrm = strrep(fstrm, '%%BeginProlog', sprintf('%%%%BeginProlog\n/pdfmark where {pop} {userdict /pdfmark /cleartomark load put} ifelse'));
    % Add page bookmark
    fstrm = strrep(fstrm, '%%EndPageSetup', sprintf('%%%%EndPageSetup\n[ /Title (%s) /OUT pdfmark',bookmark_text));

    % Write out the updated file
    fh = fopen(fname, 'w');
    if fh == -1
        error('export_fig:bookmark:permission','Unable to open %s for writing.', fname);
    end
    try
        fwrite(fh, fstrm, 'char*1');
    catch ex
        fclose(fh);
        rethrow(ex);
    end
    fclose(fh);
end

function set_tick_mode(Hlims, ax)
    % Set the tick mode of linear axes to manual
    % Leave log axes alone as these are tricky
    M = get(Hlims, [ax 'Scale']);
    if ~iscell(M)
        M = {M};
    end
    %idx = cellfun(@(c) strcmp(c, 'linear'), M);
    idx = find(strcmp(M,'linear'));
    %set(Hlims(idx), [ax 'TickMode'], 'manual');  % issue #187
    %set(Hlims(idx), [ax 'TickLabelMode'], 'manual');  % this hides exponent label in HG2!
    for idx2 = 1 : numel(idx)
        try
            % Fix for issue #187 - only set manual ticks when no exponent is present
            hAxes = Hlims(idx(idx2));
            props = {[ax 'TickMode'],'manual', [ax 'TickLabelMode'],'manual'};
            tickVals = get(hAxes,[ax 'Tick']);
            tickStrs = get(hAxes,[ax 'TickLabel']);
            try % Fix issue #236
                exponents = [hAxes.([ax 'Axis']).SecondaryLabel];
            catch
                exponents = [hAxes.([ax 'Ruler']).SecondaryLabel];
            end
            if isempty([exponents.String])
                % Fix for issue #205 - only set manual ticks when the Ticks number match the TickLabels number
                if numel(tickVals) == numel(tickStrs)
                    set(hAxes, props{:});  % no exponent and matching ticks, so update both ticks and tick labels to manual
                end
            end
        catch  % probably HG1
            % Fix for issue #220 - exponent is removed in HG1 when TickMode is 'manual' (internal Matlab bug)
            if isequal(tickVals, str2num(tickStrs)') %#ok<ST2NM>
                set(hAxes, props{:});  % revert back to old behavior
            end
        end
    end
end

function change_rgb_to_cmyk(fname)  % convert RGB => CMYK within an EPS file
    % Do post-processing on the eps file
    try
        % Read the EPS file into memory
        fstrm = read_write_entire_textfile(fname);

        % Replace all gray-scale colors
        fstrm = regexprep(fstrm, '\n([\d.]+) +GC\n', '\n0 0 0 ${num2str(1-str2num($1))} CC\n');
        
        % Replace all RGB colors
        fstrm = regexprep(fstrm, '\n[0.]+ +[0.]+ +[0.]+ +RC\n', '\n0 0 0 1 CC\n');  % pure black
        fstrm = regexprep(fstrm, '\n([\d.]+) +([\d.]+) +([\d.]+) +RC\n', '\n${sprintf(''%.4g '',[1-[str2num($1),str2num($2),str2num($3)]/max([str2num($1),str2num($2),str2num($3)]),1-max([str2num($1),str2num($2),str2num($3)])])} CC\n');

        % Overwrite the file with the modified contents
        read_write_entire_textfile(fname, fstrm);
    catch
        % never mind - leave as is...
    end
end

function [hBlackAxles, hBlackRulers] = fixBlackAxle(hAxes, axleName)
    hBlackAxles  = [];
    hBlackRulers = [];
    for idx = 1 : numel(hAxes)
        ax = hAxes(idx);
        axleColor = get(ax, axleName);
        if isequal(axleColor,[0,0,0]) || isequal(axleColor,'k')
            hBlackAxles(end+1) = ax; %#ok<AGROW>
            try  % Fix issue #306 - black yyaxis
                if strcmpi(axleName,'Color'), continue, end  %ruler, not axle
                rulerName = strrep(axleName,'Color','Axis');
                hRulers = get(ax, rulerName);
                newBlackRulers = fixBlackAxle(hRulers,'Color');
                hBlackRulers = [hBlackRulers newBlackRulers]; %#ok<AGROW>
            catch
            end
        end
    end
    set(hBlackAxles, axleName, [0,0,0.01]);  % off-black
end

% Issue #269: format-specific options
function [optionsCells, bitDepth] = getFormatOptions(options, formatName)
    bitDepth = [];
    try
        optionsStruct = options.format_options.(lower(formatName));
    catch
        % User did not specify any extra parameters for this format
        optionsCells = {};
        return
    end
    optionNames = fieldnames(optionsStruct);
    optionVals  = struct2cell(optionsStruct);
    optionsCells = [optionNames, optionVals]';
    if nargout < 2, return, end  % bail out if BitDepth is not required
    try
        idx = find(strcmpi(optionNames,'BitDepth'), 1, 'last');
        if ~isempty(idx)
            bitDepth = optionVals{idx};
        end
    catch
        % never mind - ignore
    end
end

% Check for newer version (only once a day)
function isNewerVersionAvailable = checkForNewerVersion(currentVersion)
    persistent lastCheckTime lastVersion
    isNewerVersionAvailable = false;
    if nargin < 1 || isempty(lastCheckTime) || now - lastCheckTime > 1
        url = 'https://raw.githubusercontent.com/altmany/export_fig/master/export_fig.m';
        try
            str = readURL(url);
            [unused,unused,unused,unused,latestVerStrs] = regexp(str, '\n[^:]+: \(([^)]+)\) ([^%]+)(?=\n)'); %#ok<ASGLU>
            latestVersion = str2double(latestVerStrs{end}{1});
            if nargin < 1
                currentVersion = lastVersion;
            else
                currentVersion = currentVersion + 1e3*eps;
            end
            isNewerVersionAvailable = latestVersion > currentVersion;
            if isNewerVersionAvailable
                try
                    verStrs = strtrim(reshape([latestVerStrs{:}],2,[]));
                    verNums = arrayfun(@(c)str2double(c{1}),verStrs(1,:));
                    isValid = verNums > currentVersion;
                    versionDesc = strjoin(flip(verStrs(2,isValid)),';');
                catch
                    % Something bad happened - only display the latest version description
                    versionDesc = latestVerStrs{1}{2};
                end
                try versionDesc = strjoin(strrep(strcat(' ***', strtrim(strsplit(versionDesc,';'))),'***','* '), char(10)); catch, end %#ok<CHARTEN>
                msg = sprintf(['You are using version %.2f of export_fig. ' ...
                               'A newer version (%g) is available, with the following improvements/fixes:\n' ...
                               '%s\n' ...
                               'A change-log of recent releases is available here; the complete change-log is included at the top of the export_fig.m file.\n' ...  % issue #322
                               'You can download the new version from GitHub or Matlab File Exchange, ' ...
                               'or run export_fig(''-update'') to install it directly.' ...
                              ], currentVersion, latestVersion, versionDesc);
                msg = hyperlink('https://github.com/altmany/export_fig', 'GitHub', msg);
                msg = hyperlink('https://www.mathworks.com/matlabcentral/fileexchange/23629-export_fig', 'Matlab File Exchange', msg);
                msg = hyperlink('matlab:export_fig(''-update'')', 'export_fig(''-update'')', msg);
                msg = hyperlink('https://github.com/altmany/export_fig/releases', 'available here', msg);
                msg = hyperlink('https://github.com/altmany/export_fig/blob/master/export_fig.m#L300', 'export_fig.m file', msg);
                warning('export_fig:version',msg);
            end
        catch
            % ignore
        end
        lastCheckTime = now;
        lastVersion = currentVersion;
    end
end

% Update the installed version of export_fig from the latest version online
function updateInstalledVersion()
    % Download the latest version of export_fig into the export_fig folder
    zipFileName = 'https://github.com/altmany/export_fig/archive/master.zip';
    fprintf('Downloading latest version of %s from %s...\n', mfilename, zipFileName);
    folderName = fileparts(which(mfilename('fullpath')));
    targetFileName = fullfile(folderName, datestr(now,'yyyy-mm-dd.zip'));
    try
        folder = hyperlink(['matlab:winopen(''' folderName ''')'], folderName);
    catch  % hyperlink.m is not properly installed
        folder = folderName;
    end
    try
        urlwrite(zipFileName,targetFileName); %#ok<URLWR>
    catch err
        error('export_fig:update:download','Error downloading %s into %s: %s\n',zipFileName,targetFileName,err.message);
    end

    % Unzip the downloaded zip file in the export_fig folder
    fprintf('Extracting %s...\n', targetFileName);
    try
        unzip(targetFileName,folderName);
        % Fix issue #302 - zip file uses an internal folder export_fig-master
        subFolder = fullfile(folderName,'export_fig-master');
        try movefile(fullfile(subFolder,'*.*'),folderName, 'f'); catch, end %All OSes
        try movefile(fullfile(subFolder,'*'),  folderName, 'f'); catch, end %MacOS/Unix
        try movefile(fullfile(subFolder,'.*'), folderName, 'f'); catch, end %MacOS/Unix
        try rmdir(subFolder); catch, end
    catch err
        error('export_fig:update:unzip','Error unzipping %s: %s\n',targetFileName,err.message);
    end

    % Notify the user and rehash
    fprintf('Successfully installed the latest %s version in %s\n', mfilename, folder);
    clear functions %#ok<CLFUNC>
    rehash
end

% Read a file from the web
function str = readURL(url)
    try
        str = char(webread(url));
    catch err %if isempty(which('webread'))
        if isempty(strfind(err.message,'404'))
            v = version;   % '9.6.0.1072779 (R2019a)'
            if v(1) >= '8' % '8.0 (R2012b)'  https://www.mathworks.com/help/matlab/release-notes.html?rntext=urlread&searchHighlight=urlread&startrelease=R2012b&endrelease=R2012b
                str = urlread(url, 'Timeout',5); %#ok<URLRD>
            else
                str = urlread(url); %#ok<URLRD>  % R2012a or older (no Timeout parameter)
            end
        else
            rethrow(err)
        end
    end
    if size(str,1) > 1  % ensure a row-wise string
        str = str';
    end
end

% Display a promo message in the Matlab console
function displayPromoMsg(msg, url)
    %msg = [msg url];
    msg = strrep(msg,'<$>',url);
    link = ['<a href="' url];
    msg = regexprep(msg,url,[link '">$0</a>']);
    %msg = regexprep(msg,{'consulting','training'},[link '/$0">$0</a>']);
    %warning('export_fig:promo',msg);
    disp(['[' 8 msg ']' 8]);
end

% Cross-check existance of other programs
function programsCrossCheck()
    try
        % IQ
        hasTaskList = false;
        if ispc && ~exist('IQML','file')
            hasIQ = exist('C:/Progra~1/DTN/IQFeed','dir') || ...
                    exist('C:/Progra~2/DTN/IQFeed','dir');
            if ~hasIQ
                [status,tasksStr] = system('tasklist'); %#ok<ASGLU>
                tasksStr = lower(tasksStr);
                hasIQ = ~isempty(strfind(tasksStr,'iqconnect')) || ...
                        ~isempty(strfind(tasksStr,'iqlink'));  %#ok<STREMP>
                hasTaskList = true;
            end
            if hasIQ
                displayPromoMsg('To connect Matlab to IQFeed, try the IQML connector <$>', 'https://UndocumentedMatlab.com/IQML');
            end
        end

        % IB
        if ~exist('IBMatlab','file')
            hasIB = false;
            possibleFolders = {'C:/Jts','C:/Programs/Jts','C:/Progra~1/Jts','C:/Progra~2/Jts','~/IBJts','~/IBJts/IBJts'};
            for folderIdx = 1 : length(possibleFolders)
                if exist(possibleFolders{folderIdx},'dir')
                    hasIB = true;
                    break
                end
            end
            if ~hasIB
                if ~hasTaskList
                    if ispc  % Windows
                        [status,tasksStr] = system('tasklist'); %#ok<ASGLU>
                    else  % Linux/MacOS
                        [status,tasksStr] = system('ps -e'); %#ok<ASGLU>
                    end
                    tasksStr = lower(tasksStr);
                end
                hasIB = ~isempty(strfind(tasksStr,'tws')) || ...
                        ~isempty(strfind(tasksStr,'ibgateway'));  %#ok<STREMP>
            end
            if hasIB
                displayPromoMsg('To connect Matlab to IB try the IB-Matlab connector <$>', 'https://UndocumentedMatlab.com/IB-Matlab');
            end
        end
    catch
        % never mind - ignore error
    end
end

% Hint to users to use exportgraphics/copygraphics in certain cases
function alertForExportOrCopygraphics(options)
    %matlabVerNum = str2num(regexprep(version,'(\d+\.\d+).*','$1'));
    try
        % Bail out on R2019b- (copygraphics/exportgraphics not available/reliable)
        if verLessThan('matlab','9.8')  % 9.8 = R2020a
            return
        end

        isNoRendererSpecified = options.renderer == 0;
        isPainters            = options.renderer == 3;
        noResolutionSpecified = isempty(options.resolution) || isequal(options.resolution,864);

        % First check for copygraphics compatibility (export to clipboard)
        params = ',';
        if options.clipboard
            if options.transparent  % -transparent was requested
                if isPainters || isNoRendererSpecified  % painters or default renderer
                    if noResolutionSpecified
                        params = '''BackgroundColor'',''none'',''ContentType'',''vector'',';
                    else  % exception: no message
                        params = ',';
                    end
                else  % opengl/zbuffer renderer
                    if options.invert_hardcopy  % default
                        params = '''BackgroundColor'',''white'',';
                    else  % -noinvert was requested
                        params = '''BackgroundColor'',''current'',';  % 'none' is 'white' when ContentType='image'
                    end
                    params = [params '''ContentType'',''image'','];
                    if ~noResolutionSpecified
                        params = [params '''Resolution'',' num2str(options.resolution) ','];
                    else
                        % don't add a resolution param
                    end
                end
            else  % no -transparent
                if options.invert_hardcopy  % default
                    params = '''BackgroundColor'',''white'',';
                else  % -noinvert was requested
                    params = '''BackgroundColor'',''current'',';
                end
                if isPainters  % painters (but not default!) renderer
                    if noResolutionSpecified
                        params = [params '''ContentType'',''vector'','];
                    else  % exception: no message
                        params = ',';
                    end
                else  % opengl/zbuffer/default renderer
                    params = [params '''ContentType'',''image'','];
                    if ~noResolutionSpecified
                        params = [params '''Resolution'',' num2str(options.resolution) ','];
                    else
                        % don't add a resolution param
                    end
                end
            end

            % If non-RGB colorspace was requested on R2021a+
            if ~verLessThan('matlab','9.10')  % 9.10 = R2021a
                if options.colourspace == 2  % gray
                    params = [params '''Colorspace'',''gray'','];
                end
            end
        end
        displayMsg(params, 'copygraphics', 'clipboard', '');

        % Next check for exportgraphics compatibility (export to file)
        % Note: not <else>, since -clipboard can be combined with file export
        params = ',';
        if ~options.clipboard
            if options.transparent  % -transparent was requested
                if isvector(options)  % vector output
                    if isPainters || isNoRendererSpecified  % painters or default renderer
                        if noResolutionSpecified
                            params = '''BackgroundColor'',''none'',''ContentType'',''vector'',';
                        else  % exception: no message
                            params = ',';
                        end
                    else  % opengl/zbuffer renderer
                        params = '''BackgroundColor'',''none'',''ContentType'',''vector'',';
                    end
                else % non-vector output
                    params = ',';
                end
            else  % no -transparent
                if options.invert_hardcopy  % default
                    params = '''BackgroundColor'',''white'',';
                else  % -noinvert was requested
                    params = '''BackgroundColor'',''current'',';
                end
                if isvector(options)  % vector output
                    if isPainters || isNoRendererSpecified  % painters or default renderer
                        if noResolutionSpecified
                            params = [params '''ContentType'',''vector'','];
                        else  % exception: no message
                            params = ',';
                        end
                    else  % opengl/zbuffer renderer
                        if noResolutionSpecified
                            params = [params '''ContentType'',''image'','];
                        else  % exception: no message
                            params = ',';
                        end
                    end
                else % non-vector output
                    if isPainters  % painters (but not default!) renderer
                       % exception: no message
                       params = ',';
                    else  % opengl/zbuffer/default renderer
                        if ~noResolutionSpecified
                            params = [params '''Resolution'',' num2str(options.resolution) ','];
                        end
                    end
                end
            end

            % If non-RGB colorspace was requested on R2021a+
            if ~verLessThan('matlab','9.10')  % 9.10 = R2021a
                if options.colourspace == 2  % gray
                    params = [params '''Colorspace'',''gray'','];
                elseif options.colourspace == 1 && options.eps % cmyk (eps only)
                    params = [params '''Colorspace'',''cmyk'','];
                end
            end
        end
        filenameParam = 'filename,'; %=[options.name ','];
        displayMsg(params, 'exportgraphics', 'file', filenameParam);
    catch 
        % Ignore errors - do not stop export_fig processing
    end

    % Utility function to display an alert message
    function displayMsg(params, funcName, type, filenameParam)
        if length(params) > 1
            % strip default param values from the message
            params = strrep(params, '''BackgroundColor'',''white'',', '');
            % strip the trailing ,
            if ~isempty(params) && params(end)==',', params(end)=''; end
            % if this message was not already displayed
            try prevParams = getpref('export_fig',funcName); catch, prevParams = ''; end
            if ~strcmpi(params, prevParams)
                % display the message (TODO: perhaps replace warning() with fprintf()?)
                if ~isempty([filenameParam params])
                    filenameParam = [',' filenameParam];
                end
                if ~isempty(filenameParam) && filenameParam(end)==',' && isempty(params)
                    filenameParam(end) = '';
                end
                handleName = options.handleName;
                if isempty(options.handleName) % handle was either not specified, or via gca()/gcf() etc. [i.e. not by variable]
                    handleName = 'hFigure';
                end
                msg = ['In Matlab R2020a+ you can also use ' funcName '(' handleName filenameParam params ') for simple ' type ' export'];
                oldWarn = warning('on','verbose');
                warning(['export_fig:' funcName], msg);
                warning(oldWarn);
                setpref('export_fig',funcName,params);
            end
        end
    end
end
```
