## Supplementary material for "In-depth computational analysis of natural and artificial carbon fixation pathways": MATLAB files: ghostscript.m

```
function varargout = ghostscript(cmd)
%GHOSTSCRIPT  Calls a local GhostScript executable with the input command
%
% Example:
%   [status result] = ghostscript(cmd)
%
% Attempts to locate a ghostscript executable, finally asking the user to
% specify the directory ghostcript was installed into. The resulting path
% is stored for future reference.
% 
% Once found, the executable is called with the input command string.
%
% This function requires a Ghostscript installation on your system.
% You can download Ghostscript from http://ghostscript.com (Windows/Linux)
% or http://pages.uoregon.edu/koch (MacOS).
%
% IN:
%   cmd - Command string to be passed into ghostscript.
%
% OUT:
%   status - 0 iff command ran without problem.
%   result - Output from ghostscript.

% Copyright: Oliver Woodford, 2009-2015, Yair Altman 2015-
%{
% Thanks to Jonas Dorn for the fix for the title of the uigetdir window on Mac OS.
% Thanks to Nathan Childress for the fix to default location on 64-bit Windows systems.
% 27/04/11 - Find 64-bit Ghostscript on Windows. Thanks to Paul Durack and
%            Shaun Kline for pointing out the issue
% 04/05/11 - Thanks to David Chorlian for pointing out an alternative
%            location for gs on linux.
% 12/12/12 - Add extra executable name on Windows. Thanks to Ratish
%            Punnoose for highlighting the issue.
% 28/06/13 - Fix error using GS 9.07 in Linux. Many thanks to Jannick
%            Steinbring for proposing the fix.
% 24/10/13 - Fix error using GS 9.07 in Linux. Many thanks to Johannes 
%            for the fix.
% 23/01/14 - Add full path to ghostscript.txt in warning. Thanks to Koen
%            Vermeer for raising the issue.
% 27/02/15 - If Ghostscript croaks, display suggested workarounds
% 30/03/15 - Improved performance by caching status of GS path check, if ok
% 14/05/15 - Clarified warning message in case GS path could not be saved
% 29/05/15 - Avoid cryptic error in case the ghostscipt path cannot be saved (issue #74)
% 10/11/15 - Custom GS installation webpage for MacOS. Thanks to Andy Hueni via FEX
% 15/01/20 - Various message cleanups/fixes in case of errors
%}

    try
        % Call ghostscript
        [varargout{1:nargout}] = system([gs_command(gs_path()) cmd]);
    catch err
        % Display possible workarounds for Ghostscript croaks
        url1 = 'https://github.com/altmany/export_fig/issues/12#issuecomment-61467998';  % issue #12
        url2 = 'https://github.com/altmany/export_fig/issues/20#issuecomment-63826270';  % issue #20
        hg2_str = ''; if using_hg2, hg2_str = ' or Matlab R2014a'; end
        fprintf(2, 'Ghostscript error. Rolling back to GS 9.10%s may possibly solve this:\n * %s ', hg2_str, hyperlink(url1));
        if using_hg2
            fprintf(2, '(GS 9.10)\n * %s (R2014a)', hyperlink(url2));
        end
        fprintf('\n\n');
        if ismac || isunix
            url3 = 'https://github.com/altmany/export_fig/issues/27';  % issue #27
            fprintf(2, 'Alternatively, this may possibly be due to a font path issue:\n * %s\n\n', hyperlink(url3));
            % issue #20
            % TODO: in Unix/Mac, find a way to automatically determine whether to use "export" (bash) or "setenv" (csh/tcsh)
            if isdeployed
                url = [mfilename '.m'];
            else
                fpath = which(mfilename);
                if isempty(fpath), fpath = [mfilename('fullpath') '.m']; end
                url = ['<a href="matlab:opentoline(''' fpath ''',201)">' fpath '</a>'];
            end
            fprintf(2, 'Alternatively, if you are using csh, modify shell_cmd from "export ..." to "setenv ..."\nat the bottom of %s\n\n', url);
        end
        rethrow(err);
    end
end

function path_ = gs_path
    % Return a valid path
    % Start with the currently set path
    path_ = user_string('ghostscript');
    % Check the path works
    if check_gs_path(path_)
        return
    end
    % Check whether the binary is on the path
    if ispc
        bin = {'gswin32c.exe', 'gswin64c.exe', 'gs'};
    else
        bin = {'gs'};
    end
    for a = 1:numel(bin)
        path_ = bin{a};
        if check_store_gs_path(path_)
            return
        end
    end
    % Search the obvious places
    if ispc
        default_location = 'C:\Program Files\gs\';
        dir_list = dir(default_location);
        if isempty(dir_list)
            default_location = 'C:\Program Files (x86)\gs\'; % Possible location on 64-bit systems
            dir_list = dir(default_location);
        end
        executable = {'\bin\gswin32c.exe', '\bin\gswin64c.exe'};
        ver_num = 0;
        % If there are multiple versions, use the newest
        for a = 1:numel(dir_list)
            ver_num2 = sscanf(dir_list(a).name, 'gs%g');
            if ~isempty(ver_num2) && ver_num2 > ver_num
                for b = 1:numel(executable)
                    path2 = [default_location dir_list(a).name executable{b}];
                    if exist(path2, 'file') == 2
                        path_ = path2;
                        ver_num = ver_num2;
                    end
                end
            end
        end
        if check_store_gs_path(path_)
            return
        end
    else
        executable = {'/usr/bin/gs', '/usr/local/bin/gs'};
        for a = 1:numel(executable)
            path_ = executable{a};
            if check_store_gs_path(path_)
                return
            end
        end
    end
    % Ask the user to enter the path
    while true
        if strncmp(computer, 'MAC', 3) % Is a Mac
            % Give separate warning as the uigetdir dialogue box doesn't have a
            % title on MacOS
            uiwait(warndlg('Ghostscript installation not found - please locate the program.', 'Ghostscript'))
            base = uigetdir('/', 'Ghostcript program location');
        else
            base = uigetdir('/', 'Ghostcript program not found - please locate it');
        end
        if isequal(base, 0)
            % User hit cancel or closed window
            break;
        end
        base = [base filesep]; %#ok<AGROW>
        bin_dir = {'', ['bin' filesep], ['lib' filesep]};
        for a = 1:numel(bin_dir)
            for b = 1:numel(bin)
                path_ = [base bin_dir{a} bin{b}];
                if exist(path_, 'file') == 2
                    if check_store_gs_path(path_)
                        return
                    end
                end
            end
        end
    end
    if ismac
        url = 'http://pages.uoregon.edu/koch';
    else
        url = 'http://ghostscript.com';
    end
    error('Ghostscript:NotFound', 'Ghostscript not found. Have you installed it from %s ?', hyperlink(url));
end

function good = check_store_gs_path(path_)
    % Check the path is valid
    good = check_gs_path(path_);
    if ~good
        return
    end
    % Update the current default path to the path found
    if ~user_string('ghostscript', path_)
        %filename = fullfile(fileparts(which('user_string.m')), '.ignore', 'ghostscript.txt');
        [unused, filename] = user_string('ghostscript'); %#ok<ASGLU>
        warning('Ghostscript:path', 'Path to ghostscript installation could not be saved in %s (perhaps a permissions issue). You can manually create this file and set its contents to %s, to improve performance in future invocations (this warning is safe to ignore).', filename, path_);
        return
    end
end

function good = check_gs_path(path_)
    persistent isOk
    if isempty(path_)
        isOk = false;
    elseif ~isequal(isOk,true)
        % Check whether the path is valid
        [status, message] = system([gs_command(path_) '-h']); %#ok<ASGLU>
        isOk = status == 0;
    end
    good = isOk;
end

function cmd = gs_command(path_)
    % Initialize any required system calls before calling ghostscript
    % TODO: in Unix/Mac, find a way to automatically determine whether to use "export" (bash) or "setenv" (csh/tcsh)
    shell_cmd = '';
    if isunix
        shell_cmd = 'export LD_LIBRARY_PATH=""; '; % Avoids an error on Linux with GS 9.07
    end
    if ismac
        shell_cmd = 'export DYLD_LIBRARY_PATH=""; ';  % Avoids an error on Mac with GS 9.07
    end
    % Construct the command string
    cmd = sprintf('%s"%s" ', shell_cmd, path_);
end
```
