## Supplementary material for "In-depth computational analysis of natural and artificial carbon fixation pathways": MATLAB files: hyperlink.m

```
function str = hyperlink(url, label, msg)
%HYPERLINK  create a string that is displayable as hyperlink in Matlab console
%
% Usage examples:
%   fprintf('Search on %s\n', hyperlink('http://google.com','Google'));
%   fprintf(hyperlink('http://google.com','Google','Search on Google\n'));
%
% HYPERLINK converts the specified URL and text label into a string that is
% displayed as a hyperlink in the Matlab console (Command Window). 
% In a deployed (compiled)  program, the URL and text label (if different
% from the URL) are displayed in a non-hyperlinked plain-text manner.
% If the optional 3rd input argument (msg) is specified, then all instances of
% the specified label within msg will be handled as above (hyperlinked etc.)
%
% IN:
%   url   - (mandatory) URL of webpage or Matlab command (e.g., 'matlab:which(...')
%   label - (optional)  text label of the hyperlink. Default: url
%   msg   - (optional)  string in which all label instances should be hyperlinked
%
% OUT:
%   str - string output

% Copyright: Yair Altman 2020-
%{
% 15/01/20 - Initial version
%}

    error(nargchk(1,3,nargin)); %#ok<NCHKN> narginchk is only available in R2011b+
    if nargin > 2 % msg was specified
        % replace all instances of label within msg with corresponding hyperlink
        str = strrep(msg, label, hyperlink(url,label));
        return
    end
    if nargin < 2, label = url; end
    isWebpage = strncmpi(url,'http',4);

    % Only hyperlink in interactive Matlab sessions, not in a deployed program
    if ~isdeployed  % interactive Matlab session
        if isWebpage  % open in a web browser
            str = ['<a href="matlab:web(''-browser'',''' url ''');">' label '</a>'];
        else  % Matlab command e.g. 'matlab:which(...'
            str = ['<a href="' url '">' label '</a>'];
        end
    else  % deployed (compiled)
        if isWebpage && ~strcmp(label,url)  % display label next to url
            str = [label ' (' url ')'];
        elseif isWebpage  % text==url - display only the url
            str = url;
        else  % Matlab command (not a webpage) - just display the label
            str = label;
        end
    end
end
```
