## Supplementary material for "In-depth computational analysis of natural and artificial carbon fixation pathways": MATLAB files: im2gif.m

```
%IM2GIF Convert a multiframe image to an animated GIF file
%
% Examples:
%   im2gif infile
%   im2gif infile outfile
%   im2gif(A, outfile)
%   im2gif(..., '-nocrop')
%   im2gif(..., '-nodither')
%   im2gif(..., '-ncolors', n)
%   im2gif(..., '-loops', n)
%   im2gif(..., '-delay', n) 
%   
% This function converts a multiframe image to an animated GIF.
%
% To create an animation from a series of figures, export to a multiframe
% TIFF file using export_fig, then convert to a GIF, as follows:
%
%    for a = 2 .^ (3:6)
%       peaks(a);
%       export_fig test.tif -nocrop -append
%    end
%    im2gif('test.tif', '-delay', 0.5);
%
%IN:
%   infile - string containing the name of the input image.
%   outfile - string containing the name of the output image (must have the
%             .gif extension). Default: infile, with .gif extension.
%   A - HxWxCxN array of input images, stacked along fourth dimension, to
%       be converted to gif.
%   -nocrop - option indicating that the borders of the output are not to
%             be cropped.
%   -nodither - option indicating that dithering is not to be used when
%               converting the image.
%   -ncolors - option pair, the value of which indicates the maximum number
%              of colors the GIF can have. This can also be a quantization
%              tolerance, between 0 and 1. Default/maximum: 256.
%   -loops - option pair, the value of which gives the number of times the
%            animation is to be looped. Default: 65535.
%   -delay - option pair, the value of which gives the time, in seconds,
%            between frames. Default: 1/15.

% Copyright (C) Oliver Woodford 2011

%{
% 14/02/18: Merged issue #235: reduced memory usage, improved performance (thanks to @numb7rs)
% 30/11/19: Merged issue #288: Fix im2gif.m for greyscale TIFF images (thanks @Blackbelt1221)
%}

function im2gif(A, varargin)

    % Parse the input arguments
    [A, options] = parse_args(A, varargin{:});

    if options.crop ~= 0
        % Crop
        A = crop_borders(A, A(ceil(end/2),1,:,1));
    end

    % Convert to indexed image
    [h, w, c, n] = size(A);

    % Issue #235: Using unique(A,'rows') on the whole image stack at once causes
    % massive memory usage when dealing with large images (at least on Matlab 2017b).
    % Running unique(...) on individual frames, then again on the results drastically
    % reduces the memory usage & slightly improves the execution time (@numb7rs).
    uns = cell(1,size(A,4));
    for nn=1:size(A,4)
        uns{nn}=unique(reshape(A(:,:,:,nn), h*w, c),'rows');
    end
    map=unique(cell2mat(uns'),'rows');

    A = reshape(permute(A, [1 2 4 3]), h, w*n, c);

    if size(map, 1) > 256
        dither_str = {'dither', 'nodither'};
        dither_str = dither_str{1+(options.dither==0)};
        if options.ncolors <= 1
            [B, map] = rgb2ind(A, options.ncolors, dither_str);
            if size(map, 1) > 256
                [B, map] = rgb2ind(A, 256, dither_str);
            end
        else
            [B, map] = rgb2ind(A, min(round(options.ncolors), 256), dither_str);
        end
    else
        if max(map(:)) > 1
            map = double(map) / 255;
            A = double(A) / 255;
        end
        B = rgb2ind(im2double(A), map);
    end
    B = reshape(B, h, w, 1, n);

    % Bug fix to rgb2ind
    map(B(1)+1,:) = im2double(A(1,1,:));

    % Save as a gif
    imwrite(B, map, options.outfile, 'LoopCount', round(options.loops(1)), 'DelayTime', options.delay);
end

%% Parse the input arguments
function [A, options] = parse_args(A, varargin)
    % Set the defaults
    options = struct('outfile', '', ...
                     'dither', true, ...
                     'crop', true, ...
                     'ncolors', 256, ...
                     'loops', 65535, ...
                     'delay', 1/15);

    % Go through the arguments
    a = 0;
    n = numel(varargin);
    while a < n
        a = a + 1;
        if ischar(varargin{a}) && ~isempty(varargin{a})
            if varargin{a}(1) == '-'
                opt = lower(varargin{a}(2:end));
                switch opt
                    case 'nocrop'
                        options.crop = false;
                    case 'nodither'
                        options.dither = false;
                    otherwise
                        if ~isfield(options, opt)
                            error('Option %s not recognized', varargin{a});
                        end
                        a = a + 1;
                        if ischar(varargin{a}) && ~ischar(options.(opt))
                            options.(opt) = str2double(varargin{a});
                        else
                            options.(opt) = varargin{a};
                        end
                end
            else
                options.outfile = varargin{a};
            end
        end
    end

    if isempty(options.outfile)
        if ~ischar(A)
            error('No output filename given.');
        end
        % Generate the output filename from the input filename
        [path, outfile] = fileparts(A);
        options.outfile = fullfile(path, [outfile '.gif']);
    end

    if ischar(A)
        % Read in the image
        A = imread_rgb(A);
    end
end

%% Read image to uint8 rgb array
function [A, alpha] = imread_rgb(name)
    % Get file info
    info = imfinfo(name);
    % Special case formats
    switch lower(info(1).Format)
        case 'gif'
            [A, map] = imread(name, 'frames', 'all');
            if ~isempty(map)
                map = uint8(map * 256 - 0.5); % Convert to uint8 for storage
                A = reshape(map(uint32(A)+1,:), [size(A) size(map, 2)]); % Assume indexed from 0
                A = permute(A, [1 2 5 4 3]);
            end
        case {'tif', 'tiff'}
            A = cell(numel(info), 1);
            for a = 1:numel(A)
                [A{a}, map] = imread(name, 'Index', a, 'Info', info);
                if ~isempty(map)
                    map = uint8(map * 256 - 0.5); % Convert to uint8 for storage
                    A{a} = reshape(map(uint32(A{a})+1,:), [size(A) size(map, 2)]); % Assume indexed from 0
                end
                if size(A{a}, 3) == 4
                    % TIFF in CMYK colourspace - convert to RGB
                    if isfloat(A{a})
                        A{a} = A{a} * 255;
                    else
                        A{a} = single(A{a});
                    end
                    A{a} = 255 - A{a};
                    A{a}(:,:,4) = A{a}(:,:,4) / 255;
                    A{a} = uint8(A(:,:,1:3) .* A{a}(:,:,[4 4 4]));
                elseif size(A{a}, 3) < 3 %Check whether TIFF has been read in as greyscale
                    %Convert from greyscale to RGB colorspace (issue #288)
                    A{a} = cat(3, A{a}, A{a}, A{a});
                end
            end
            A = cat(4, A{:});
        otherwise
            [A, map, alpha] = imread(name);
            A = A(:,:,:,1); % Keep only first frame of multi-frame files
            if ~isempty(map)
                map = uint8(map * 256 - 0.5); % Convert to uint8 for storage
                A = reshape(map(uint32(A)+1,:), [size(A) size(map, 2)]); % Assume indexed from 0
            elseif size(A, 3) == 4
                % Assume 4th channel is an alpha matte
                alpha = A(:,:,4);
                A = A(:,:,1:3);
            end
    end
end
```
