## Supplementary material for "In-depth computational analysis of natural and artificial carbon fixation pathways": MATLAB files: pdf2eps.m

```
%PDF2EPS  Convert a pdf file to eps format using pdftops
%
% Examples:
%   pdf2eps source dest
%
% This function converts a pdf file to eps format.
%
% This function requires that you have pdftops, from the Xpdf suite of
% functions, installed on your system. This can be downloaded from:
% http://xpdfreader.com
%
% Inputs:
%   source - filename of the source pdf file to convert. The filename is
%            assumed to already have the extension ".pdf".
%   dest - filename of the destination eps file. The filename is assumed to
%          already have the extension ".eps".

% Copyright (C) Oliver Woodford 2009-2010, Yair Altman 2015-

% Thanks to Aldebaro Klautau for reporting a bug when saving to
% non-existant directories.

% 22/09/2018 - Xpdf website changed to xpdfreader.com

function pdf2eps(source, dest)
    % Construct the options string for pdftops
    options = ['-q -paper match -eps -level2 "' source '" "' dest '"'];

    % Convert to eps using pdftops
    [status, message] = pdftops(options);

    % Check for error
    if status
        % Report error
        if isempty(message)
            error('Unable to generate eps. Check destination directory is writable.');
        else
            error(message);
        end
    end

    % Fix the DSC error created by pdftops
    fid = fopen(dest, 'r+');
    if fid == -1
        % Cannot open the file
        return
    end
    fgetl(fid); % Get the first line
    str = fgetl(fid); % Get the second line
    if strcmp(str(1:min(13, end)), '% Produced by')
        fseek(fid, -numel(str)-1, 'cof');
        fwrite(fid, '%'); % Turn ' ' into '%'
    end
    fclose(fid);
end
```
