## Supplementary material for "In-depth computational analysis of natural and artificial carbon fixation pathways": MATLAB files: pdftops.m

```
function varargout = pdftops(cmd)
%PDFTOPS  Calls a local pdftops executable with the input command
%
% Example:
%   [status result] = pdftops(cmd)
%
% Attempts to locate a pdftops executable, finally asking the user to
% specify the directory pdftops was installed into. The resulting path is
% stored for future reference.
% 
% Once found, the executable is called with the input command string.
%
% This function requires that you have pdftops (from the Xpdf package)
% installed on your system. You can download this from: http://xpdfreader.com
%
% IN:
%   cmd - Command string to be passed into pdftops (e.g. '-help').
%
% OUT:
%   status - 0 iff command ran without problem.
%   result - Output from pdftops.

% Copyright: Oliver Woodford, 2009-2010

% Thanks to Jonas Dorn for the fix for the title of the uigetdir window on Mac OS.
% Thanks to Christoph Hertel for pointing out a bug in check_xpdf_path under linux.
% 23/01/2014 - Add full path to pdftops.txt in warning.
% 27/05/2015 - Fixed alert in case of missing pdftops; fixed code indentation
% 02/05/2016 - Added possible error explanation suggested by Michael Pacer (issue #137)
% 02/05/2016 - Search additional possible paths suggested by Jonas Stein (issue #147)
% 03/05/2016 - Display the specific error message if pdftops fails for some reason (issue #148)
% 22/09/2018 - Xpdf website changed to xpdfreader.com; improved popup logic
% 03/02/2019 - Fixed one-off 'pdftops not found' error after install (Mac/Linux) (issue #266)
% 15/01/2020 - Fixed reported path of pdftops.txt file in case of error; added warning ID
% 23/07/2020 - Fixed issue #311 (confusion regarding Xpdf-tools download/installation); silent check of pdftops installation in case no input arg specified

    % If no command parameter specified, just check pdftops installation and bail out
    if nargin < 1
        xpdf_path();  % this will error if pdftops is not found
        return        % silent bail-out if pdftops was successfully located
    end

    % Call pdftops
    [varargout{1:nargout}] = system([xpdf_command(xpdf_path()) cmd]);
end

function path_ = xpdf_path
    % Return a valid path
    % Start with the currently set path
    path_ = user_string('pdftops');
    % Check the path works
    if check_xpdf_path(path_)
        return
    end
    % Check whether the binary is on the path
    if ispc
        bin = 'pdftops.exe';
    else
        bin = 'pdftops';
    end
    if check_store_xpdf_path(bin)
        path_ = bin;
        return
    end
    % Search the obvious places
    if ispc
        paths = {'C:\Program Files\xpdf\pdftops.exe', 'C:\Program Files (x86)\xpdf\pdftops.exe'};
    else
        paths = {'/usr/bin/pdftops', '/usr/local/bin/pdftops'};
    end
    for a = 1:numel(paths)
        path_ = paths{a};
        if check_store_xpdf_path(path_)
            return
        end
    end

    % Ask the user to enter the path
    errMsg1 = 'Pdftops utility not found. Please locate the program, or install xpdf-tools from ';
    url1 = 'http://xpdfreader.com/download.html'; %='http://foolabs.com/xpdf';
    fprintf(2, '%s%s ("Xpdf command line tools" section)\n', errMsg1, hyperlink(url1));
    errMsg1 = [errMsg1 url1];
    %if strncmp(computer,'MAC',3) % Is a Mac
    %    % Give separate warning as the MacOS uigetdir dialogue box doesn't have a title
    %    uiwait(warndlg(errMsg1))
    %end

    % Provide an alternative possible explanation as per issue #137
    errMsg2 = 'If pdftops is installed, maybe Matlab is shaddowing it, as described in ';
    url2 = 'https://github.com/altmany/export_fig/issues/137';
    fprintf(2, '%s%s\n', errMsg2, hyperlink(url2,'issue #137'));
    errMsg2 = [errMsg2 url2];

    % Provide an alternative possible explanation as per issue #311
    errMsg3 = 'Or perhaps you installed XpdfReader but not xpdf-tools, as described in ';
    url3 = 'https://github.com/altmany/export_fig/issues/311';
    fprintf(2, '%s%s\n', errMsg3, hyperlink(url3,'issue #311'));
    errMsg3 = [errMsg3 url3];

    state = 1;
    while 1
        if state
            option1 = 'Install pdftops';
        else
            option1 = 'Issue #137';
        end
        answer = questdlg({errMsg1,'',errMsg2,'',errMsg3},'Pdftops error',option1,'Locate pdftops','Cancel','Cancel');
        drawnow;  % prevent a Matlab hang: http://undocumentedmatlab.com/blog/solving-a-matlab-hang-problem
        switch answer
            case 'Install pdftops'
                web('-browser',url1);
                state = 0;
            case 'Issue #137'
                web('-browser',url2);
                state = 1;
            case 'Locate pdftops'
                base = uigetdir('/', errMsg1);
                if isequal(base, 0)
                    % User hit cancel or closed window
                    break
                end
                base = [base filesep]; %#ok<AGROW>
                bin_dir = {'', ['bin' filesep], ['lib' filesep]};
                for a = 1:numel(bin_dir)
                    path_ = [base bin_dir{a} bin];
                    if exist(path_, 'file') == 2
                        break
                    end
                end
                if check_store_xpdf_path(path_)
                    return
                end

            otherwise  % User hit Cancel or closed window
                break
        end
    end
    error('pdftops executable not found.');
end

function good = check_store_xpdf_path(path_)
    % Check the path is valid
    good = check_xpdf_path(path_);
    if ~good
        return
    end
    % Update the current default path to the path found
    if ~user_string('pdftops', path_)
        %filename = fullfile(fileparts(which('user_string.m')), '.ignore', 'pdftops.txt');
        [unused, filename] = user_string('pdftops'); %#ok<ASGLU>
        warning('export_fig:pdftops','Path to pdftops executable could not be saved. Enter it manually in %s.', filename);
        return
    end
end

function good = check_xpdf_path(path_)
    % Check the path is valid
    [good, message] = system([xpdf_command(path_) '-h']); %#ok<ASGLU>
    % system returns good = 1 even when the command runs
    % Look for something distinct in the help text
    good = ~isempty(strfind(message, 'PostScript')); %#ok<STREMP>

    % Display the error message if the pdftops executable exists but fails for some reason
    % Note: on Mac/Linux, exist('pdftops','file') will always return 2 due to pdftops.m => check for '/','.' (issue #266)
    if ~good && exist(path_,'file') && ~isempty(regexp(path_,'[/.]')) %#ok<RGXP1> % file exists but generates an error
        fprintf('Error running %s:\n', path_);
        fprintf(2,'%s\n\n',message);
    end
end

function cmd = xpdf_command(path_)
    % Initialize any required system calls before calling ghostscript
    % TODO: in Unix/Mac, find a way to determine whether to use "export" (bash) or "setenv" (csh/tcsh)
    shell_cmd = '';
    if isunix
        % Avoids an error on Linux with outdated MATLAB lib files
        % R20XXa/bin/glnxa64/libtiff.so.X
        % R20XXa/sys/os/glnxa64/libstdc++.so.X
        shell_cmd = 'export LD_LIBRARY_PATH=""; ';
    end
    if ismac
        shell_cmd = 'export DYLD_LIBRARY_PATH=""; ';
    end
    % Construct the command string
    cmd = sprintf('%s"%s" ', shell_cmd, path_);
end
```
