## Supplementary material for "In-depth computational analysis of natural and artificial carbon fixation pathways": MATLAB files: print2array.m

```
function [A, bcol, alpha] = print2array(fig, res, renderer, gs_options)
%PRINT2ARRAY  Exports a figure to a bitmap RGB image array
%
% Examples:
%   A = print2array
%   A = print2array(figure_handle)
%   A = print2array(figure_handle, resolution)
%   A = print2array(figure_handle, resolution, renderer)
%   A = print2array(figure_handle, resolution, renderer, gs_options)
%   [A, bcol, alpha] = print2array(...)
%
% This function outputs a bitmap image of a figure, at the desired resolution.
%
% When resolution==1, fast Java screen-capture is attempted first.
% If the Java screen-capture fails or if resolution~=1, the builtin print()
% function is used to create a temp TIF file, which is then loaded and reported.
% If this fails, print() is used to create a temp EPS file which is converted to
% a TIF file using Ghostcript (http://www.ghostscript.com), loaded and reported.
%
% Inputs:
%   figure_handle - The handle of the figure to be exported. Default: gcf.
%   resolution - Output resolution as a factor of screen resolution. Default: 1
%                Note: resolution ~= 1 uses a slow print to/from image file
%   renderer   - The renderer to be used by print() function. Default: '-opengl'
%                Note: only used when resolution ~= 1
%   gs_options - optional ghostscript parameters (e.g.: '-dNoOutputFonts').
%                Enclose multiple options in a cell array, e.g. {'-a','-b'}
%                Note: only used when resolution ~= 1 and basic print() fails
%
% Outputs:
%   A     - MxNx3 uint8 bitmap image of the figure (MxN pixels x 3 RGB values)
%   bcol  - 1x3 uint8 vector of the background RGB color
%   alpha - MxN uint8 array of alpha values (between 0=transparent, 255=opaque)

% Copyright (C) Oliver Woodford 2008-2014, Yair Altman 2015-
%{
% 05/09/11: Set EraseModes to normal when using opengl or zbuffer
%           renderers. Thanks to Pawel Kocieniewski for reporting the issue.
% 21/09/11: Bug fix: unit8 -> uint8! Thanks to Tobias Lamour for reporting it.
% 14/11/11: Bug fix: stop using hardcopy(), as it interfered with figure size
%           and erasemode settings. Makes it a bit slower, but more reliable.
%           Thanks to Phil Trinh and Meelis Lootus for reporting the issues.
% 09/12/11: Pass font path to ghostscript.
% 27/01/12: Bug fix affecting painters rendering tall figures. Thanks to
%           Ken Campbell for reporting it.
% 03/04/12: Bug fix to median input. Thanks to Andy Matthews for reporting it.
% 26/10/12: Set PaperOrientation to portrait. Thanks to Michael Watts for
%           reporting the issue.
% 26/02/15: If temp dir is not writable, use the current folder for temp
%           EPS/TIF files (Javier Paredes)
% 27/02/15: Display suggested workarounds to internal print() error (issue #16)
% 28/02/15: Enable users to specify optional ghostscript options (issue #36)
% 10/03/15: Fixed minor warning reported by Paul Soderlind; fixed code indentation
% 28/05/15: Fixed issue #69: patches with LineWidth==0.75 appear wide (internal bug in Matlab's print() func)
% 07/07/15: Fixed issue #83: use numeric handles in HG1
% 11/12/16: Fixed cropping issue reported by Harry D.
% 29/09/18: Fixed issue #254: error in print2array>read_tif_img
% 22/03/20: Alert if ghostscript.m is required but not found on Matlab path
% 24/05/20: Significant performance speedup; added alpha values (where possible)
% 07/07/20: Fixed issue #308: bug in R2019a and earlier
% 07/10/20: Use JavaFrame_I where possible, to avoid evoking a JavaFrame warning
%}

    % Generate default input arguments, if needed
    if nargin < 1,  fig = gcf;  end
    if nargin < 2,  res = 1;    end

    % Get the figure size in pixels
    old_mode = get(fig, 'Units');
    set(fig, 'Units', 'pixels');
    px = get(fig, 'Position');
    set(fig, 'Units', old_mode);

    % Retrieve the background colour
    bcol = get(fig, 'Color');
    try
        % Try a direct Java screen-capture first - *MUCH* faster than print() to file
        % Note: we could also use A=matlab.graphics.internal.getframeWithDecorations(fig,false) but it (1) returns no alpha and (2) does not exist in older Matlabs
        if res == 1
            [A, alpha] = getJavaImage(fig);
        else
            error('magnify/downscale via print() to image file and then import');
        end
    catch err  %#ok<NASGU>
        % Warn if output is large
        npx = prod(px(3:4)*res)/1e6;
        if npx > 30
            % 30M pixels or larger!
            warning('MATLAB:LargeImage', 'print2array generating a %.1fM pixel image. This could be slow and might also cause memory problems.', npx);
        end
        % Set the resolution parameter
        res_str = ['-r' num2str(ceil(get(0, 'ScreenPixelsPerInch')*res))];
        % Generate temporary file name
        tmp_nam = [tempname '.tif'];
        try
            % Ensure that the temp dir is writable (Javier Paredes 26/2/15)
            fid = fopen(tmp_nam,'w');
            fwrite(fid,1);
            fclose(fid);
            delete(tmp_nam);  % cleanup
            isTempDirOk = true;
        catch
            % Temp dir is not writable, so use the current folder
            [dummy,fname,fext] = fileparts(tmp_nam); %#ok<ASGLU>
            fpath = pwd;
            tmp_nam = fullfile(fpath,[fname fext]);
            isTempDirOk = false;
        end
        % Enable users to specify optional ghostscript options (issue #36)
        if nargin > 3 && ~isempty(gs_options)
            if iscell(gs_options)
                gs_options = sprintf(' %s',gs_options{:});
            elseif ~ischar(gs_options)
                error('gs_options input argument must be a string or cell-array of strings');
            else
                gs_options = [' ' gs_options];
            end
        else
            gs_options = '';
        end
        if nargin > 2 && strcmp(renderer, '-painters')
            % First try to print directly to image file
            try
                % Print the file into a temporary image file and read it into array A
                [A, alpha, err, ex] = getPrintImage(fig, res_str, renderer, tmp_nam);
                if err, rethrow(ex); end
            catch  % error - try to print to EPS and then using Ghostscript to TIF
                % Ensure that ghostscript() exists on the Matlab path
                if ~exist('ghostscript','file') && isempty(which('ghostscript'))
                    error('export_fig:print2array:ghostscript', 'The ghostscript.m function is required by print2array.m. Install the complete export_fig package from https://www.mathworks.com/matlabcentral/fileexchange/23629-export_fig or https://github.com/altmany/export_fig')
                end
                % Print to eps file
                if isTempDirOk
                    tmp_eps = [tempname '.eps'];
                else
                    tmp_eps = fullfile(fpath,[fname '.eps']);
                end
                print2eps(tmp_eps, fig, 0, renderer, '-loose');
                try
                    % Initialize the command to export to tiff using ghostscript
                    cmd_str = ['-dEPSCrop -q -dNOPAUSE -dBATCH ' res_str ' -sDEVICE=tiff24nc'];
                    % Set the font path
                    fp = font_path();
                    if ~isempty(fp)
                        cmd_str = [cmd_str ' -sFONTPATH="' fp '"'];
                    end
                    % Add the filenames
                    cmd_str = [cmd_str ' -sOutputFile="' tmp_nam '" "' tmp_eps '"' gs_options];
                    % Execute the ghostscript command
                    ghostscript(cmd_str);
                catch me
                    % Delete the intermediate file
                    delete(tmp_eps);
                    rethrow(me);
                end
                % Delete the intermediate file
                delete(tmp_eps);
                % Read in the generated bitmap
                A = imread(tmp_nam);
                % Delete the temporary bitmap file
                delete(tmp_nam);
            end
        else
            if nargin < 3
                renderer = '-opengl';
            end
            % Print the file into a temporary image file and read it into array A
            [A, alpha, err, ex] = getPrintImage(fig, res_str, renderer, tmp_nam);
            % Throw any error that occurred
            if err
                % Display suggested workarounds to internal print() error (issue #16)
                fprintf(2, 'An error occured with Matlab''s builtin print function.\nTry setting the figure Renderer to ''painters'' or use opengl(''software'').\n\n');
                rethrow(ex);
            end
        end
    end

    % Set the background color
    if isequal(bcol, 'none')
        bcol = squeeze(A(1,1,:));
        if ~all(bcol==0)  %if not black
            bcol = [255,255,255]; %=white  %=[];
        end
    else
        if all(bcol <= 1)
            bcol = bcol * 255;
        end
        if ~isequal(bcol, round(bcol))
            bcol = squeeze(A(1,1,:));
            %{
            % Set border pixels to the correct colour
            for l = 1:size(A, 2)
                if ~all(reshape(A(:,l,:) == 255, [], 1))
                    break;
                end
            end
            for r = size(A, 2):-1:l
                if ~all(reshape(A(:,r,:) == 255, [], 1))
                    break;
                end
            end
            for t = 1:size(A, 1)
                if ~all(reshape(A(t,:,:) == 255, [], 1))
                    break;
                end
            end
            for b = size(A, 1):-1:t
                if ~all(reshape(A(b,:,:) == 255, [], 1))
                    break;
                end
            end
            bcol = median(single([reshape(A(:,[l r],:), [], size(A, 3)); ...
                                  reshape(A([t b],:,:), [], size(A, 3))]), 1));
            for c = 1:size(A, 3)
                A(:,[1:l-1, r+1:end],c) = bcol(c);
                A([1:t-1, b+1:end],:,c) = bcol(c);
            end
            %}
        end
    end
    bcol = uint8(bcol);

    % Ensure that the output size is correct
    if isequal(res, round(res))
        px = round([px([4 3])*res 3]);  % round() to avoid an indexing warning below
        if any(size(A) > px) %~isequal(size(A), px)
            A = A(1:min(end,px(1)),1:min(end,px(2)),:);
        end
        if any(size(alpha) > px(1:2))
            alpha = alpha(1:min(end,px(1)),1:min(end,px(2)));
        end
    end
end

% Get the Java-based screen-capture of the figure's JFrame content-panel
function [imgData, alpha] = getJavaImage(hFig)
    % Get the figure's underlying Java frame
    oldWarn = warning('off','MATLAB:HandleGraphics:ObsoletedProperty:JavaFrame');
    warning('off','MATLAB:ui:javaframe:PropertyToBeRemoved');
    try
        jf = get(handle(hFig),'JavaFrame_I');
    catch
        jf = get(handle(hFig),'JavaFrame'); %#ok<JAVFM>
    end
    warning(oldWarn);

    % Get the Java frame's root frame handle
    %jframe = jf.getFigurePanelContainer.getComponent(0).getRootPane.getParent;
    try
        jClient = jf.fHG2Client;  % This works from R2014b and up
    catch
        try
            jClient = jf.fHG1Client;  % This works from R2008b-R2014a
        catch
            jClient = jf.fFigureClient;  % This works up to R2011a
        end
    end

    % Get the content-pane
    try
        jPanel = jClient.getContentPane;
    catch
        jPanel = jClient.getFigurePanelContainer;
    end
    jPanel.repaint;
    w = jPanel.getWidth;
    h = jPanel.getHeight;

    % Create a BufferedImage and paint the content-pane into it
    % (https://coderanch.com/t/470601/java/screenshot-JPanel)
    % Note: contrary to documentation and common-sense, it turns out that TYPE_INT_RGB
    % ^^^^  returns non-opaque alpha, while TYPE_INT_ARGB only returns 255s in the alpha channel 
    jOriginalGraphics = jPanel.getGraphics;
    import java.awt.image.BufferedImage
    try TYPE_INT_RGB = BufferedImage.TYPE_INT_RGB; catch, TYPE_INT_RGB = 1; end
    jImage = BufferedImage(w, h, TYPE_INT_RGB);
    jPanel.paint(jImage.createGraphics);
    jPanel.paint(jOriginalGraphics);  % repaint original figure to avoid a blank window

    % Extract the RGB pixels from the BufferedImage (see screencapture.m)
    pixelsData = reshape(typecast(jImage.getData.getDataStorage, 'uint8'), 4, w, h);
    imgData = cat(3, ...
                  transpose(reshape(pixelsData(3, :, :), w, h)), ...
                  transpose(reshape(pixelsData(2, :, :), w, h)), ...
                  transpose(reshape(pixelsData(1, :, :), w, h)));

    % And now also the alpha channel (if available)
    alpha   =     transpose(reshape(pixelsData(4, :, :), w, h));

    % Ensure that the results are the expected size, otherwise raise an error
    figSize = getpixelposition(gcf);
    expectedSize = [figSize(4), figSize(3), 3];
    if ~isequal(expectedSize, size(imgData))
        error('bad Java screen-capture size!')
    end
end

% Export an image file of the figure using print() and then read it into an array
function [imgData, alpha, err, ex] = getPrintImage(fig, res_str, renderer, tmp_nam)
    imgData = [];  % fix for issue #254
    err     = false;
    ex      = [];
    alpha   = [];
    % Temporarily set the paper size
    old_pos_mode    = get(fig, 'PaperPositionMode');
    old_orientation = get(fig, 'PaperOrientation');
    set(fig, 'PaperPositionMode','auto', 'PaperOrientation','portrait');
    try
        % Workaround for issue #69: patches with LineWidth==0.75 appear wide (internal bug in Matlab's print() function)
        fp = [];  % in case we get an error below
        fp = findall(fig, 'Type','patch', 'LineWidth',0.75);
        set(fp, 'LineWidth',0.5);
        try %if using_hg2(fig)  % HG2 (R2014b or newer)
            % Use print('-RGBImage') directly (a bit faster than via temp image file)
            imgData = print(fig, renderer, res_str, '-RGBImage');
        catch %else  % HG1 (R2014a or older)
            % Fix issue #83: use numeric handles in HG1
            fig = double(fig);
            % Print to image file
            print(fig, renderer, res_str, '-dtiff', tmp_nam);
            imgData = imread(tmp_nam);
            % Delete the temporary file
            delete(tmp_nam);
        end
        imgSize = size(imgData); imgSize = imgSize([1,2]);  % Fix issue #308
        alpha = 255 * ones(imgSize, 'uint8');  % =all pixels opaque
    catch ex
        err = true;
    end
    set(fp, 'LineWidth',0.75);  % restore original figure appearance
    % Reset the paper size
    set(fig, 'PaperPositionMode',old_pos_mode, 'PaperOrientation',old_orientation);
end

% Return (and create, where necessary) the font path (for use by ghostscript)
function fp = font_path()
    fp = user_string('gs_font_path');
    if ~isempty(fp)
        return
    end
    % Create the path
    % Start with the default path
    fp = getenv('GS_FONTPATH');
    % Add on the typical directories for a given OS
    if ispc
        if ~isempty(fp)
            fp = [fp ';'];
        end
        fp = [fp getenv('WINDIR') filesep 'Fonts'];
    else
        if ~isempty(fp)
            fp = [fp ':'];
        end
        fp = [fp '/usr/share/fonts:/usr/local/share/fonts:/usr/share/fonts/X11:/usr/local/share/fonts/X11:/usr/share/fonts/truetype:/usr/local/share/fonts/truetype'];
    end
    user_string('gs_font_path', fp);
end
```
