## Supplementary material for "In-depth computational analysis of natural and artificial carbon fixation pathways": MATLAB files: print2eps.m

```
function print2eps(name, fig, export_options, varargin)
%PRINT2EPS  Prints figures to eps with improved line styles
%
% Examples:
%   print2eps filename
%   print2eps(filename, fig_handle)
%   print2eps(filename, fig_handle, export_options)
%   print2eps(filename, fig_handle, export_options, print_options)
%
% This function saves a figure as an eps file, with two improvements over
% MATLAB's print command. First, it improves the line style, making dashed
% lines more like those on screen and giving grid lines a dotted line style.
% Secondly, it substitutes original font names back into the eps file,
% where these have been changed by MATLAB, for up to 11 different fonts.
%
% Inputs:
%   filename - string containing the name (optionally including full or
%              relative path) of the file the figure is to be saved as. A
%              ".eps" extension is added if not there already. If a path is
%              not specified, the figure is saved in the current directory.
%   fig_handle - The handle of the figure to be saved. Default: gcf().
%   export_options - array or struct of optional scalar values:
%       bb_padding - Scalar value of amount of padding to add to border around
%                    the cropped image, in points (if >1) or percent (if <1).
%                    Can be negative as well as positive; Default: 0
%       crop       - Cropping flag. Deafult: 0
%       fontswap   - Whether to swap non-default fonts in figure. Default: true
%       preserve_size - Whether to preserve the figure's PaperSize. Default: false
%       font_space - Character used to separate font-name terms in the EPS output
%                    e.g. "Courier New" => "Courier-New". Default: ''
%                    (available only via the struct alternative)
%       renderer   - Renderer used to generate bounding-box. Default: 'opengl'
%                    (available only via the struct alternative)
%       crop_amounts - 4-element vector of crop amounts: [top,right,bottom,left]
%                    (available only via the struct alternative)
%   print_options - Additional parameter strings to be passed to the print command

%{
% Copyright (C) Oliver Woodford 2008-2014, Yair Altman 2015-

% The idea of editing the EPS file to change line styles comes from Jiro
% Doke's FIXPSLINESTYLE (fex id: 17928)
% The idea of changing dash length with line width came from comments on
% fex id: 5743, but the implementation is mine :)
%}
%{
% 14/11/11: Fix a MATLAB bug rendering black or white text incorrectly.
%           Thanks to Mathieu Morlighem for reporting the issue and
%           obtaining a fix from TMW.
% 08/12/11: Added ability to correct fonts. Several people have requested
%           this at one time or another, and also pointed me to printeps
%           (fex id: 7501), so thank you to them. My implementation (which
%           was not inspired by printeps - I'd already had the idea for my
%           approach) goes slightly further in that it allows multiple
%           fonts to be swapped.
% 14/12/11: Fix bug affecting font names containing spaces. Thanks to David
%           Szwer for reporting the issue.
% 25/01/12: Add a font not to be swapped. Thanks to Anna Rafferty and Adam
%           Jackson for reporting the issue. Also fix a bug whereby using a
%           font alias can lead to another font being swapped in.
% 10/04/12: Make the font swapping case insensitive.
% 26/10/12: Set PaperOrientation to portrait. Thanks to Michael Watts for
%           reporting the issue.
% 26/10/12: Fix issue to do with swapping fonts changing other fonts and
%           sizes we don't want, due to listeners. Thanks to Malcolm Hudson
%           for reporting the issue.
% 22/03/13: Extend font swapping to axes labels. Thanks to Rasmus Ischebeck
%           for reporting the issue.
% 23/07/13: Bug fix to font swapping. Thanks to George for reporting the
%           issue.
% 13/08/13: Fix MATLAB feature of not exporting white lines correctly.
%           Thanks to Sebastian Hesslinger for reporting it.
% 24/02/15: Fix for Matlab R2014b bug (issue #31): LineWidths<0.75 are not
%           set in the EPS (default line width is used)
% 25/02/15: Fixed issue #32: BoundingBox problem caused uncropped EPS/PDF files
% 05/03/15: Fixed issue #43: Inability to perform EPS file post-processing
% 06/03/15: Improved image padding & cropping thanks to Oscar Hartogensis
% 21/03/15: Fixed edge-case of missing handles having a 'FontName' property
% 26/03/15: Attempt to fix issue #45: white lines in subplots do not print correctly
% 27/03/15: Attempt to fix issue #44: white artifact lines appearing in patch exports
% 30/03/15: Fixed issue #52: improved performance on HG2 (R2014b+)
% 09/04/15: Comment blocks consolidation and minor code cleanup (no real code change)
% 12/04/15: Fixed issue #56: bad cropping
% 14/04/15: Workaround for issue #45: lines in image subplots are exported in invalid color
% 07/07/15: Added option to avoid font-swapping in EPS/PDF
% 07/07/15: Fixed issue #83: use numeric handles in HG1
% 22/07/15: Fixed issue #91 (thanks to Carlos Moffat)
% 28/09/15: Fixed issue #108 (thanks to JacobD10)
% 01/11/15: Fixed issue #112: optional renderer for bounding-box computation (thanks to Jes�s Pestana Puerta)
% 21/02/16: Enabled specifying non-automated crop amounts
% 22/02/16: Better support + backward compatibility for transparency (issue #108)
% 10/06/16: Fixed issue #159: text handles get cleared by Matlab in the print() command
% 12/06/16: Improved the fix for issue #159 (in the previous commit)
% 12/06/16: Fixed issue #158: transparent patch color in PDF/EPS
% 18/09/17: Fixed issue #194: incorrect fonts in EPS/PDF output
% 18/09/17: Fixed issue #195: relaxed too-tight cropping in EPS/PDF
% 14/11/17: Workaround for issue #211: dashed/dotted lines in 3D axes appear solid
% 15/11/17: Updated issue #211: only set SortMethod='ChildOrder' in HG2, and when it looks the same onscreen; support multiple figure axes
% 18/11/17: Fixed issue #225: transparent/translucent dashed/dotted lines appear solid in EPS/PDF
% 24/03/18: Fixed issue #239: black title meshes with temporary black background figure bgcolor, causing bad cropping
% 21/03/19: Improvement for issue #258: missing fonts in output EPS/PDF (still *NOT* fully solved)
% 21/03/19: Fixed issues #166,#251: Arial font is no longer replaced with Helvetica but rather treated as a non-standard user font
% 14/05/19: Made Helvetica the top default font-swap, replacing Courier
% 12/06/19: Issue #277: Enabled preservation of figure's PaperSize in output PDF/EPS file
% 06/08/19: Issue #281: only fix patch/textbox color if it's not opaque
% 15/01/20: Added warning ID for easier suppression by users
% 20/01/20: Added comment about unsupported patch transparency in some Ghostscript versions (issue #285)
%}

    options = {'-loose'};
    if nargin > 3
        options = [options varargin];
    elseif nargin < 3
        export_options = 0;
        if nargin < 2
            fig = gcf();
        end
    end

    % Retrieve padding, crop & font-swap values
    crop_amounts = nan(1,4);  % auto-crop all 4 sides by default
    if isstruct(export_options)
        try preserve_size = export_options.preserve_size; catch, preserve_size = false; end
        try fontswap      = export_options.fontswap;      catch, fontswap = true;       end
        try font_space    = export_options.font_space;    catch, font_space = '';       end
        font_space(2:end) = '';
        try bb_crop       = export_options.crop;          catch, bb_crop = 0;           end
        try crop_amounts  = export_options.crop_amounts;  catch,                        end
        try bb_padding    = export_options.bb_padding;    catch, bb_padding = 0;        end
        try renderer      = export_options.rendererStr;   catch, renderer = 'opengl';   end  % fix for issue #110
        if renderer(1)~='-',  renderer = ['-' renderer];  end
    else
        if numel(export_options) > 3  % preserve_size
            preserve_size = export_options(4);
        else
            preserve_size = false;
        end
        if numel(export_options) > 2  % font-swapping
            fontswap = export_options(3);
        else
            fontswap = true;
        end
        if numel(export_options) > 1  % cropping
            bb_crop = export_options(2);
        else
            bb_crop = 0;  % scalar value, so use default bb_crop value of 0
        end
        if numel(export_options) > 0  % padding
            bb_padding = export_options(1);
        else
            bb_padding = 0;
        end
        renderer = '-opengl';
        font_space = '';
    end

    % Construct the filename
    if numel(name) < 5 || ~strcmpi(name(end-3:end), '.eps')
        name = [name '.eps']; % Add the missing extension
    end

    % Set paper size
    old_pos_mode    = get(fig, 'PaperPositionMode');
    old_orientation = get(fig, 'PaperOrientation');
    old_paper_units = get(fig, 'PaperUnits');
    set(fig, 'PaperPositionMode','auto', 'PaperOrientation','portrait', 'PaperUnits','points');

    % Find all the used fonts in the figure
    font_handles = findall(fig, '-property', 'FontName');
    fonts = get(font_handles, 'FontName');
    if isempty(fonts)
        fonts = {};
    elseif ~iscell(fonts)
        fonts = {fonts};
    end

    % Map supported font aliases onto the correct name
    fontsl = lower(fonts);
    for a = 1:numel(fonts)
        f = fontsl{a};
        f(f==' ') = [];
        switch f
            case {'times', 'timesnewroman', 'times-roman'}
                fontsl{a} = 'times';
            %case {'arial', 'helvetica'}  % issues #166, #251
            %    fontsl{a} = 'helvetica';
            case {'newcenturyschoolbook', 'newcenturyschlbk'}
                fontsl{a} = 'newcenturyschlbk';
            otherwise
        end
    end
    fontslu = unique(fontsl);

    % Determine the font swap table
    if fontswap
        % Issue #258: Rearrange standard fonts list based on decending "problematicness"
        % The issue is still *NOT* fully solved because I cannot figure out how to force
        % the EPS postscript engine to look for the user's font on disk
        % Also see: https://stat.ethz.ch/pipermail/r-help/2005-January/064374.html
        matlab_fonts = {'Helvetica', 'Times', 'Courier', 'Symbol', 'ZapfDingbats', ...
                        'Palatino', 'Bookman', 'ZapfChancery', 'AvantGarde', ...
                        'NewCenturySchlbk', 'Helvetica-Narrow'};
        matlab_fontsl = lower(matlab_fonts);
        require_swap = find(~ismember(fontslu, matlab_fontsl));
        unused_fonts = find(~ismember(matlab_fontsl, fontslu));
        font_swap = cell(3, min(numel(require_swap), numel(unused_fonts)));
        fonts_new = fonts;
        for a = 1:size(font_swap, 2)
            font_swap{1,a} = find(strcmp(fontslu{require_swap(a)}, fontsl));
            font_swap{2,a} = matlab_fonts{unused_fonts(a)};
            font_swap{3,a} = fonts{font_swap{1,a}(1)};
            fonts_new(font_swap{1,a}) = font_swap(2,a);
        end
    else
        font_swap = [];
    end

    % Swap the fonts
    if ~isempty(font_swap)
        fonts_size = get(font_handles, 'FontSize');
        if iscell(fonts_size)
            fonts_size = cell2mat(fonts_size);
        end
        M = false(size(font_handles));

        % Loop because some changes may not stick first time, due to listeners
        c = 0;
        update = zeros(1000, 1);
        for b = 1:10 % Limit number of loops to avoid infinite loop case
            for a = 1:numel(M)
                M(a) = ~isequal(get(font_handles(a), 'FontName'), fonts_new{a}) || ~isequal(get(font_handles(a), 'FontSize'), fonts_size(a));
                if M(a)
                    set(font_handles(a), 'FontName', fonts_new{a}, 'FontSize', fonts_size(a));
                    c = c + 1;
                    update(c) = a;
                end
            end
            if ~any(M)
                break;
            end
        end

        % Compute the order to revert fonts later, without the need of a loop
        [update, M] = unique(update(1:c));
        [dummy, M] = sort(M); %#ok<ASGLU>
        update = reshape(update(M), 1, []);
    end

    % MATLAB bug fix - black and white text can come out inverted sometimes
    % Find the white and black text
    black_text_handles = findall(fig, 'Type', 'text', 'Color', [0 0 0]);
    white_text_handles = findall(fig, 'Type', 'text', 'Color', [1 1 1]);
    % Set the font colors slightly off their correct values
    set(black_text_handles, 'Color', [0 0 0] + eps);
    set(white_text_handles, 'Color', [1 1 1] - eps);

    % MATLAB bug fix - white lines can come out funny sometimes
    % Find the white lines
    white_line_handles = findall(fig, 'Type', 'line', 'Color', [1 1 1]);
    % Set the line color slightly off white
    set(white_line_handles, 'Color', [1 1 1] - 0.00001);

    % MATLAB bug fix (issue #211): dashed/dotted lines in 3D axes appear solid
    % Note: this "may limit other functionality in plotting such as hidden line/surface removal"
    % reference: Technical Support Case #02838114, https://mail.google.com/mail/u/0/#inbox/15fb7659f70e7bd8
    hAxes = findall(fig, 'Type', 'axes');
    if using_hg2 && ~isempty(hAxes)  % issue #211 presumably happens only in HG2, not HG1
        try
            % If there are any axes using SortMethod~='ChildOrder'
            oldSortMethods = get(hAxes,{'SortMethod'});  % use {'SortMethod'} to ensure we get a cell array, even for single axes
            if any(~strcmpi('ChildOrder',oldSortMethods))  % i.e., any oldSortMethods=='depth'
                % Check if the axes look visually different onscreen when SortMethod='ChildOrder'
                imgBefore = print2array(fig);
                set(hAxes,'SortMethod','ChildOrder');
                imgAfter  = print2array(fig);
                if isequal(imgBefore, imgAfter)
                    % They look the same, so use SortMethod='ChildOrder' when generating the EPS
                else
                    % They look different, so revert SortMethod and issue a warning message
                    warning('YMA:export_fig:issue211', ...
                            ['You seem to be using axes that have overlapping/hidden graphic elements. ' 10 ...
                             'Setting axes.SortMethod=''ChildOrder'' may solve potential problems in EPS/PDF export. ' 10 ...
                             'Additional info: https://github.com/altmany/export_fig/issues/211'])
                    set(hAxes,{'SortMethod'},oldSortMethods);
                end
            end
        catch err
            % ignore
            a=err;  %#ok<NASGU> % debug breakpoint
        end
    end

    % Workaround for issue #45: lines in image subplots are exported in invalid color
    % In this case the -depsc driver solves the problem, but then all the other workarounds
    % below (for all the other issues) will fail, so it's better to let the user decide by
    % just issuing a warning and accepting the '-depsc' input parameter
    epsLevel2 = ~any(strcmpi(options,'-depsc'));
    if epsLevel2
        % Use -depsc2 (EPS color level-2) if -depsc (EPS color level-3) was not specifically requested
        options{end+1} = '-depsc2';
        % Issue a warning if multiple images & lines were found in the figure, and HG1 with painters renderer is used
        isPainters = any(strcmpi(options,'-painters'));
        if isPainters && ~using_hg2 && numel(findall(fig,'Type','image'))>1 && ~isempty(findall(fig,'Type','line'))
            warning('YMA:export_fig:issue45', ...
                    ['Multiple images & lines detected. In such cases, the lines might \n' ...
                     'appear with an invalid color due to an internal MATLAB bug (fixed in R2014b). \n' ...
                     'Possible workaround: add a ''-depsc'' or ''-opengl'' parameter to the export_fig command.']);
        end
    end

    % Fix issue #83: use numeric handles in HG1
    if ~using_hg2(fig),  fig = double(fig);  end

    % Workaround for when transparency is lost through conversion fig>EPS>PDF (issue #108)
    % Replace transparent patch RGB values with an ID value (rare chance that ID color is being used already)
    if using_hg2
        origAlphaColors = eps_maintainAlpha(fig);
    end

    % Print to eps file
    print(fig, options{:}, name);

    % Restore the original axes SortMethods (if updated)
    try set(hAxes,{'SortMethod'},oldSortMethods); catch, end

    % Do post-processing on the eps file
    try
        % Read the EPS file into memory
        fstrm = read_write_entire_textfile(name);
    catch
        fstrm = '';
    end

    % Restore colors for transparent patches/lines and apply the
    % setopacityalpha setting in the EPS file (issue #108)
    if using_hg2
        [~,fstrm,foundFlags] = eps_maintainAlpha(fig, fstrm, origAlphaColors);

        % If some of the transparencies were not found in the EPS file, then rerun the
        % export with only the found transparencies modified (backward compatibility)
        if ~isempty(fstrm) && ~all(foundFlags)
            foundIdx = find(foundFlags);
            for objIdx = 1 : sum(foundFlags)
                colorsIdx = foundIdx(objIdx);
                colorsData = origAlphaColors{colorsIdx};
                hObj     = colorsData{1};
                propName = colorsData{2};
                newColor = colorsData{4};
                hObj.(propName).ColorData = newColor;
            end
            delete(name);
            print(fig, options{:}, name);
            fstrm = read_write_entire_textfile(name);
            [~,fstrm] = eps_maintainAlpha(fig, fstrm, origAlphaColors(foundFlags));
        end
    end

    % Bail out if EPS post-processing is not possible
    if isempty(fstrm)
        warning('YMA:export_fig:EPS','Loading EPS file failed, so unable to perform post-processing. This is usually because the figure contains a large number of patch objects. Consider exporting to a bitmap format in this case.');
        return
    end

    % Fix for Matlab R2014b bug (issue #31): LineWidths<0.75 are not set in the EPS (default line width is used)
    try
        if using_hg2(fig)
            % Convert miter joins to line joins
            %fstrm = regexprep(fstrm, '\n10.0 ML\n', '\n1 LJ\n');
            % This is faster (the original regexprep could take many seconds when the axes contains many lines):
            fstrm = strrep(fstrm, sprintf('\n10.0 ML\n'), sprintf('\n1 LJ\n'));

            % In HG2, grid lines and axes Ruler Axles have a default LineWidth of 0.5 => replace en-bulk (assume that 1.0 LineWidth = 1.333 LW)
            %   hAxes=gca; hAxes.YGridHandle.LineWidth, hAxes.YRuler.Axle.LineWidth
            %fstrm = regexprep(fstrm, '(GC\n2 setlinecap\n1 LJ)\nN', '$1\n0.667 LW\nN');
            % This is faster:
            fstrm = strrep(fstrm, sprintf('GC\n2 setlinecap\n1 LJ\nN'), sprintf('GC\n2 setlinecap\n1 LJ\n0.667 LW\nN'));

            % This is more accurate but *MUCH* slower (issue #52)
            %{
            % Modify all thin lines in the figure to have 10x LineWidths
            hLines = findall(fig,'Type','line');
            hThinLines = [];
            for lineIdx = 1 : numel(hLines)
                thisLine = hLines(lineIdx);
                if thisLine.LineWidth < 0.75 && strcmpi(thisLine.Visible,'on')
                    hThinLines(end+1) = thisLine; %#ok<AGROW>
                    thisLine.LineWidth = thisLine.LineWidth * 10;
                end
            end

            % If any thin lines were found
            if ~isempty(hThinLines)
                % Prepare an EPS with large-enough line widths
                print(fig, options{:}, name);
                % Restore the original LineWidths in the figure
                for lineIdx = 1 : numel(hThinLines)
                    thisLine = handle(hThinLines(lineIdx));
                    thisLine.LineWidth = thisLine.LineWidth / 10;
                end

                % Compare the original and the new EPS files and correct the original stream's LineWidths
                fstrm_new = read_write_entire_textfile(name);
                idx = 500;  % skip heading with its possibly-different timestamp
                markerStr = sprintf('10.0 ML\nN');
                markerLen = length(markerStr);
                while ~isempty(idx) && idx < length(fstrm)
                    lastIdx = min(length(fstrm), length(fstrm_new));
                    delta = fstrm(idx+1:lastIdx) - fstrm_new(idx+1:lastIdx);
                    idx = idx + find(delta,1);
                    if ~isempty(idx) && ...
                            isequal(fstrm(idx-markerLen+1:idx), markerStr) && ...
                            ~isempty(regexp(fstrm_new(idx-markerLen+1:idx+12),'10.0 ML\n[\d\.]+ LW\nN')) %#ok<RGXP1>
                        value = str2double(regexprep(fstrm_new(idx:idx+12),' .*',''));
                        if isnan(value), break; end  % something's wrong... - bail out
                        newStr = sprintf('%0.3f LW\n',value/10);
                        fstrm = [fstrm(1:idx-1) newStr fstrm(idx:end)];
                        idx = idx + 12;
                    else
                        break;
                    end
                end
            end
            %}

            % This is much faster although less accurate: fix all non-gray lines to have a LineWidth of 0.75 (=1 LW)
            % Note: This will give incorrect LineWidth of 075 for lines having LineWidth<0.75, as well as for non-gray grid-lines (if present)
            %       However, in practice these edge-cases are very rare indeed, and the difference in LineWidth should not be noticeable
            %fstrm = regexprep(fstrm, '([CR]C\n2 setlinecap\n1 LJ)\nN', '$1\n1 LW\nN');
            % This is faster (the original regexprep could take many seconds when the axes contains many lines):
            fstrm = strrep(fstrm, sprintf('\n2 setlinecap\n1 LJ\nN'), sprintf('\n2 setlinecap\n1 LJ\n1 LW\nN'));
        end
    catch err
        fprintf(2, 'Error fixing LineWidths in EPS file: %s\n at %s:%d\n', err.message, err.stack(1).file, err.stack(1).line);
    end

    % Reset the font and line colors
    try
        set(black_text_handles, 'Color', [0 0 0]);
        set(white_text_handles, 'Color', [1 1 1]);
    catch
        % Fix issue #159: redo findall() '*text_handles'
        black_text_handles = findall(fig, 'Type', 'text', 'Color', [0 0 0]+eps);
        white_text_handles = findall(fig, 'Type', 'text', 'Color', [1 1 1]-eps);
        set(black_text_handles, 'Color', [0 0 0]);
        set(white_text_handles, 'Color', [1 1 1]);
    end
    set(white_line_handles, 'Color', [1 1 1]);

    % Preserve the figure's PaperSize in the output file, if requested (issue #277)
    if preserve_size
        % https://stackoverflow.com/questions/19646329/postscript-document-size
        paper_size = get(fig, 'PaperSize');  % in [points]
        fstrm = sprintf('<< /PageSize [%d %d] >> setpagedevice\n%s', paper_size, fstrm);
    end

    % Reset paper size
    set(fig, 'PaperPositionMode',old_pos_mode, 'PaperOrientation',old_orientation, 'PaperUnits',old_paper_units);

    % Reset the font names in the figure
    if ~isempty(font_swap)
        for a = update
            set(font_handles(a), 'FontName', fonts{a}, 'FontSize', fonts_size(a));
        end
    end

    % Replace the font names
    if ~isempty(font_swap)
        for a = 1:size(font_swap, 2)
            fontName = font_swap{3,a};
            %fontName = fontName(~isspace(font_swap{3,a}));
            if length(fontName) > 29
                warning('YMA:export_fig:font_name','Font name ''%s'' is longer than 29 characters. This might cause problems in some EPS/PDF readers. Consider using a different font.',fontName);
            end
            if isempty(font_space)
                fontName(fontName==' ') = '';
            else
                fontName(fontName==' ') = char(font_space);
            end

            % Replace all instances of the standard Matlab fonts with the original user's font names
            %fstrm = regexprep(fstrm, [font_swap{1,a} '-?[a-zA-Z]*\>'], fontName);
            %fstrm = regexprep(fstrm, [font_swap{2,a} '([ \n])'], [fontName '$1']);
            %fstrm = regexprep(fstrm, font_swap{2,a}, fontName);  % also replace -Bold, -Italic, -BoldItalic

            % Times-Roman's Bold/Italic fontnames don't include '-Roman'
            fstrm = regexprep(fstrm, [font_swap{2,a} '(\-Roman)?'], fontName);
        end
    end

    % Move the bounding box to the top of the file (HG2 only), or fix the line styles (HG1 only)
    if using_hg2(fig)
        % Move the bounding box to the top of the file (HG2 only)
        [s, e] = regexp(fstrm, '%%BoundingBox: [^%]*%%');
        if numel(s) == 2
            fstrm = fstrm([1:s(1)-1 s(2):e(2)-2 e(1)-1:s(2)-1 e(2)-1:end]);
        end
    else
        % Fix the line styles (HG1 only)
        fstrm = fix_lines(fstrm);
    end

    % Apply the bounding box padding & cropping, replacing Matlab's print()'s bounding box
    if bb_crop
        % Calculate a new bounding box based on a bitmap print using crop_border.m
        % 1. Determine the Matlab BoundingBox and PageBoundingBox
        [s,e] = regexp(fstrm, '%%BoundingBox: [^%]*%%'); % location BB in eps file
        if numel(s)==2, s=s(2); e=e(2); end
        aa = fstrm(s+15:e-3); % dimensions bb - STEP1
        bb_matlab = cell2mat(textscan(aa,'%f32%f32%f32%f32'));  % dimensions bb - STEP2

        [s,e] = regexp(fstrm, '%%PageBoundingBox: [^%]*%%'); % location bb in eps file
        if numel(s)==2, s=s(2); e=e(2); end
        aa = fstrm(s+19:e-3); % dimensions bb - STEP1
        pagebb_matlab = cell2mat(textscan(aa,'%f32%f32%f32%f32'));  % dimensions bb - STEP2

        % 1b. Fix issue #239: black title meshes with temporary black background figure bgcolor, causing bad cropping
        hTitles = [];
        if isequal(get(fig,'Color'),'none')
            hAxes = findall(fig,'type','axes');
            for idx = 1 : numel(hAxes)
                hAx = hAxes(idx);
                try
                    hTitle = hAx.Title;
                    oldColor = hTitle.Color;
                    if all(oldColor < 5*eps) || (ischar(oldColor) && lower(oldColor(1))=='k')
                        hTitles(end+1) = hTitle; %#ok<AGROW>
                        hTitle.Color = [0,0,.01];
                    end
                catch
                end
            end
        end

        % 2. Create a bitmap image and use crop_borders to create the relative
        %    bb with respect to the PageBoundingBox
        [A, bcol] = print2array(fig, 1, renderer);
        [aa, aa, aa, bb_rel] = crop_borders(A, bcol, bb_padding, crop_amounts); %#ok<ASGLU>

        try set(hTitles,'Color','k'); catch, end

        % 3. Calculate the new Bounding Box
        pagew = pagebb_matlab(3)-pagebb_matlab(1);
        pageh = pagebb_matlab(4)-pagebb_matlab(2);
        %bb_new = [pagebb_matlab(1)+pagew*bb_rel(1) pagebb_matlab(2)+pageh*bb_rel(2) ...
        %          pagebb_matlab(1)+pagew*bb_rel(3) pagebb_matlab(2)+pageh*bb_rel(4)];
        bb_new = pagebb_matlab([1,2,1,2]) + [pagew,pageh,pagew,pageh].*bb_rel;  % clearer
        bb_offset = (bb_new-bb_matlab) + [-2,-2,2,2];  % 2px margin so that cropping is not TOO tight (issue #195)

        % Apply the bounding box padding
        if bb_padding
            if abs(bb_padding)<1
                bb_padding = round((mean([bb_new(3)-bb_new(1) bb_new(4)-bb_new(2)])*bb_padding)/0.5)*0.5; % ADJUST BB_PADDING
            end
            add_padding = @(n1, n2, n3, n4) sprintf(' %.0f', str2double({n1, n2, n3, n4}) + bb_offset + bb_padding*[-1,-1,1,1]); %#ok<NASGU>
        else
            add_padding = @(n1, n2, n3, n4) sprintf(' %.0f', str2double({n1, n2, n3, n4}) + bb_offset); %#ok<NASGU> % fix small but noticeable bounding box shift
        end
        fstrm = regexprep(fstrm, '%%BoundingBox:[ ]+([-]?\d+)[ ]+([-]?\d+)[ ]+([-]?\d+)[ ]+([-]?\d+)', '%%BoundingBox:${add_padding($1, $2, $3, $4)}');
    end

    % Fix issue #44: white artifact lines appearing in patch exports
    % Note: the problem is due to the fact that Matlab's print() function exports patches
    %       as a combination of filled triangles, and a white line appears where the triangles touch
    % In the workaround below, we will modify such dual-triangles into a filled rectangle.
    % We are careful to only modify regexps that exactly match specific patterns - it's better to not
    % correct some white-line artifacts than to change the geometry of a patch, or to corrupt the EPS.
    %   e.g.: '0 -450 937 0 0 450 3 MP PP 937 0 0 -450 0 450 3 MP PP' => '0 -450 937 0 0 450 0 0 4 MP'
    fstrm = regexprep(fstrm, '\n([-\d.]+ [-\d.]+) ([-\d.]+ [-\d.]+) ([-\d.]+ [-\d.]+) 3 MP\nPP\n\2 \1 \3 3 MP\nPP\n','\n$1 $2 $3 0 0 4 MP\nPP\n');
    fstrm = regexprep(fstrm, '\n([-\d.]+ [-\d.]+) ([-\d.]+ [-\d.]+) ([-\d.]+ [-\d.]+) 3 MP\nPP\n\2 \3 \1 3 MP\nPP\n','\n$1 $2 $3 0 0 4 MP\nPP\n');
    fstrm = regexprep(fstrm, '\n([-\d.]+ [-\d.]+) ([-\d.]+ [-\d.]+) ([-\d.]+ [-\d.]+) 3 MP\nPP\n\3 \1 \2 3 MP\nPP\n','\n$1 $2 $3 0 0 4 MP\nPP\n');
    fstrm = regexprep(fstrm, '\n([-\d.]+ [-\d.]+) ([-\d.]+ [-\d.]+) ([-\d.]+ [-\d.]+) 3 MP\nPP\n\3 \2 \1 3 MP\nPP\n','\n$1 $2 $3 0 0 4 MP\nPP\n');

    % Write out the fixed eps file
    read_write_entire_textfile(name, fstrm);
end

function [StoredColors, fstrm, foundFlags] = eps_maintainAlpha(fig, fstrm, StoredColors)
    if nargin == 1  % in: convert transparency in Matlab figure into unique RGB colors
        hObjs = findall(fig); %findobj(fig,'Type','Area');
        StoredColors = {};
        propNames = {'Face','Edge'};
        for objIdx = 1:length(hObjs)
            hObj = hObjs(objIdx);
            for propIdx = 1 : numel(propNames)
                try
                    propName = propNames{propIdx};
                    if strcmp(hObj.(propName).ColorType, 'truecoloralpha')
                        oldColor = hObj.(propName).ColorData;
                        if numel(oldColor)>3 && oldColor(4)~=255  % issue #281: only fix patch/textbox color if it's not opaque
                            nColors = length(StoredColors);
                            newColor = uint8([101; 102+floor(nColors/255); mod(nColors,255); 255]);
                            StoredColors{end+1} = {hObj, propName, oldColor, newColor}; %#ok<AGROW>
                            hObj.(propName).ColorData = newColor;
                        end
                    end
                catch
                    % Never mind - ignore (either doesn't have the property or cannot change it)
                end
            end
        end
    else  % restore transparency in Matlab figure by converting back from the unique RGBs
        %Find the transparent patches
        wasError = false;
        nColors = length(StoredColors);
        foundFlags = false(1,nColors);
        for objIdx = 1 : nColors
            colorsData = StoredColors{objIdx};
            hObj      = colorsData{1};
            propName  = colorsData{2};
            origColor = colorsData{3};
            newColor  = colorsData{4};
            try
                %Restore the EPS files patch color
                colorID   = num2str(round(double(newColor(1:3)') /255,3),'%.3g %.3g %.3g'); %ID for searching
                origRGB   = num2str(round(double(origColor(1:3)')/255,3),'%.3g %.3g %.3g'); %Replace with original color
                origAlpha = num2str(round(double(origColor(end)) /255,3),'%.3g'); %Convert alpha value for EPS

                %Find and replace the RGBA values within the EPS text fstrm
                %Note: .setopacityalpha is an unsupported PS extension that croaks in some GS versions (issue #285, https://bugzilla.redhat.com/show_bug.cgi?id=1632030)
                %      (such cases are caught in eps2pdf.m and corrected by adding the -dNOSAFER Ghosscript option or by removing the .setopacityalpha line)
                if strcmpi(propName,'Face')
                    oldStr = sprintf(['\n' colorID ' RC\n']);  % ...N\n (removed to fix issue #225)
                    newStr = sprintf(['\n' origRGB ' RC\n' origAlpha ' .setopacityalpha true\n']);  % ...N\n
                else  %'Edge'
                    oldStr = sprintf(['\n' colorID ' RC\n']);  % ...1 LJ\n (removed to fix issue #225)
                    newStr = sprintf(['\n' origRGB ' RC\n' origAlpha ' .setopacityalpha true\n']);
                end
                foundFlags(objIdx) = ~isempty(strfind(fstrm, oldStr)); %#ok<STREMP>
                fstrm = strrep(fstrm, oldStr, newStr);

                %Restore the figure object's original color
                hObj.(propName).ColorData = origColor;
            catch err
                % something is wrong - cannot restore transparent color...
                if ~wasError
                    fprintf(2, 'Error maintaining transparency in EPS file: %s\n at %s:%d\n', err.message, err.stack(1).file, err.stack(1).line);
                    wasError = true;
                end
            end
        end
    end
end
```
