## Supplementary material for "In-depth computational analysis of natural and artificial carbon fixation pathways": MATLAB files: README.md

export_fig
==========

A toolbox for exporting figures from MATLAB to standard image and document formats nicely.

### Overview
Exporting a figure from MATLAB the way you want it (hopefully the way it looks on screen), can be a real headache for the unitiated, thanks to all the settings that are required, and also due to some eccentricities (a.k.a. features and bugs) of functions such as `print`. The first goal of export_fig is to make transferring a plot from screen to document, just the way you expect (again, assuming that's as it appears on screen), a doddle.
  
The second goal is to make the output media suitable for publication, allowing you to publish your results in the full glory that you originally intended. This includes embedding fonts, setting image compression levels (including lossless), anti-aliasing, cropping, setting the colourspace, alpha-blending and getting the right resolution.

Perhaps the best way to demonstrate what export_fig can do is with some examples.

*Note: `export_fig` currently supports only figures created with the `figure` function, or GUIDE. Figures created using `uifigure` or AppDesigner are only partially supported. See issues [#287](https://github.com/altmany/export_fig/issues/287), [#261](https://github.com/altmany/export_fig/issues/261) for details.*
  
### Examples
**Visual accuracy** - MATLAB's exporting functions, namely `saveas` and `print`, change many visual properties of a figure, such as size, axes limits and ticks, and background colour, in unexpected and unintended ways. Export_fig aims to faithfully reproduce the figure as it appears on screen. For example:  
```Matlab
plot(cos(linspace(0, 7, 1000)));
set(gcf, 'Position', [100 100 150 150]);
saveas(gcf, 'test.png');
export_fig test2.png
```
generates the following:

| Figure: | test.png: | test2.png: |
|:-------:|:---------:|:----------:|
|![](https://farm6.staticflickr.com/5616/15589249291_16e485c29a_o_d.png)|![](https://farm4.staticflickr.com/3944/15406302850_4d2e1c7afa_o_d.png)|![](https://farm6.staticflickr.com/5607/15568225476_8ce9bd5f6b_o_d.png)|

Note that the size and background colour of test2.png (the output of export_fig) are the same as those of the on screen figure, in contrast to test.png. Of course, if you want the figure background to be white (or any other colour) in the exported file then you can set this prior to exporting using:
```Matlab
set(gcf, 'Color', 'w');
```
  
Notice also that export_fig crops and anti-aliases (smooths, for bitmaps only) the output by default. However, these options can be disabled; see the Tips section below for details.
  
**Resolution** - by default, export_fig exports bitmaps at screen resolution. However, you may wish to save them at a different resolution. You can do this using either of two options: `-m<val>`, where <val> is a positive real number, magnifies the figure by the factor <val> for export, e.g. `-m2` produces an image double the size (in pixels) of the on screen figure; `-r<val>`, again where <val> is a positive real number, specifies the output bitmap to have <val> pixels per inch, the dimensions of the figure (in inches) being those of the on screen figure. For example, using:  
```Matlab
export_fig test.png -m2.5
```
on the figure from the example above generates:

![](https://farm4.staticflickr.com/3937/15591910915_dc7040c477_o_d.png)

Sometimes you might have a figure with an image in. For example:
```Matlab
imshow(imread('cameraman.tif'))
hold on
plot(0:255, sin(linspace(0, 10, 256))*127+128);
set(gcf, 'Position', [100 100 150 150]);
```
generates this figure:

![](https://farm4.staticflickr.com/3942/15589249581_ff87a56a3f_o_d.png)
  
Here the image is displayed in the figure at resolution lower than its native resolution. However, you might want to export the figure at a resolution such that the image is output at its native (i.e. original) size (in pixels). Ordinarily this would require some non-trivial computation to work out what that resolution should be, but export_fig has an option to do this for you. Using:
```Matlab
export_fig test.png -native
```
produces:

![](https://farm6.staticflickr.com/5604/15589249591_da2b2652e4_o_d.png)

with the image being the size (in pixels) of the original image. Note that if you want an image to be a particular size, in pixels, in the output (other than its original size) then you can resize it to this size and use the `-native` option to achieve this.

All resolution options (`-m<val>`, `-q<val>` and `-native`) correctly set the resolution information in PNG and TIFF files, as if the image were the dimensions of the on screen figure.

**Shrinking dots & dashes** - when exporting figures with dashed or dotted lines using either the ZBuffer or OpenGL (default for bitmaps) renderers, the dots and dashes can appear much shorter, even non-existent, in the output file, especially if the lines are thick and/or the resolution is high. For example:  
```Matlab
plot(sin(linspace(0, 10, 1000)), 'b:', 'LineWidth', 4);
hold on
plot(cos(linspace(0, 7, 1000)), 'r--', 'LineWidth', 3);
grid on
export_fig test.png
```
generates:

![](https://farm4.staticflickr.com/3956/15592747732_f943d4aa0a_o_d.png)

This problem can be overcome by using the painters renderer. For example:
```Matlab
export_fig test.png -painters
```
used on the same figure generates:

![](https://farm4.staticflickr.com/3945/14971168504_77692f11f5_o_d.png)

Note that not only are the plot lines correct, but the grid lines are too.

**Transparency** - sometimes you might want a figure and axes' backgrounds to be transparent, so that you can see through them to a document (for example a presentation slide, with coloured or textured background) that the exported figure is placed in. To achieve this, first (optionally) set the axes' colour to 'none' prior to exporting, using:  
```Matlab
set(gca, 'Color', 'none'); % Sets axes background
```
    
then use export_fig's `-transparent` option when exporting:
```Matlab
export_fig test.png -transparent
```

This will make the background transparent in PDF, EPS and PNG outputs. You can additionally save fully alpha-blended semi-transparent patch objects to the PNG format. For example:

```Matlab
logo;
alpha(0.5);
```

generates a figure like this:

![](https://farm4.staticflickr.com/3933/15405290339_b08de33528_o_d.png)

If you then export this to PNG using the `-transparent` option you can then put the resulting image into, for example, a presentation slide with fancy, textured background, like so:

![](https://farm6.staticflickr.com/5599/15406302920_59beaefff1_o_d.png)

and the image blends seamlessly with the background.

**Image quality** - when publishing images of your results, you want them to look as good as possible. By default, when outputting to lossy file formats (PDF, EPS and JPEG), export_fig uses a high quality setting, i.e. low compression, for images, so little information is lost. This is in contrast to MATLAB's print and saveas functions, whose default quality settings are poor. For example:
```Matlab
A = im2double(imread('peppers.png'));
B = randn(ceil(size(A, 1)/6), ceil(size(A, 2)/6), 3) * 0.1;
B = cat(3, kron(B(:,:,1), ones(6)), kron(B(:,:,2), ones(6)), kron(B(:,:,3), ones(6)));
B = A + B(1:size(A, 1),1:size(A, 2),:);
imshow(B);
print -dpdf test.pdf
```
generates a PDF file, a sub-window of which looks (when zoomed in) like this:

![](https://farm6.staticflickr.com/5613/15405290309_881b2774d6_o_d.png)

while the command

```Matlab
export_fig test.pdf
```
on the same figure produces this:

![](https://farm4.staticflickr.com/3947/14971168174_687473133f_o_d.png)

While much better, the image still contains some compression artifacts (see the low level noise around the edge of the pepper). You may prefer to export with no artifacts at all, i.e. lossless compression. Alternatively, you might need a smaller file, and be willing to accept more compression. Either way, export_fig has an option that can suit your needs: `-q<val>`, where <val> is a number from 0-100, will set the level of lossy image compression (again in PDF, EPS and JPEG outputs only; other formats are lossless), from high compression (0) to low compression/high quality (100). If you want lossless compression in any of those formats then specify a <val> greater than 100. For example:
```Matlab
export_fig test.pdf -q101
```
again on the same figure, produces this:

![](https://farm6.staticflickr.com/5608/15405803908_934512c1fe_o_d.png)

Notice that all the noise has gone.

### Tips
**Anti-aliasing** - the anti-aliasing which export_fig applies to bitmap outputs by default makes the images look nice, but it can also blur images and increase exporting time and memory requirements, so you might not always want it. You can set the level of anti-aliasing by using the `-a<val>` option, where <val> is 1 (no anti-aliasing), 2, 3 (default) or 4 (maximum anti-aliasing).  
  
**Cropping** - by default, export_fig crops its output to minimize the amount of empty space around the figure. If you'd prefer the figure to be uncropped, and instead have the same appearance (in terms of border width) as the on screen figure, then use the `-nocrop` option.  
  
**Colourspace** - by default, export_fig generates files in the RGB [colourspace](https://en.wikipedia.org/wiki/Color_space). However, you can also export in greyscale or the CMYK colourspace, using the `-grey` (or `-gray`) and `-cmyk` options respectively. The CMYK option is useful for publishers who require documents in this colourspace, but the option is only supported for PDF, EPS and TIFF files.

**Specifying a target directory** - you can get export_fig to save output files to any directory (for which you have write permission), simply by specifying the full or relative path in the filename. For example:
```Matlab
export_fig ../subdir/fig.png;
export_fig('C:/Users/Me/Documents/figures/myfig', '-pdf', '-png');
```

**Variable file names** - often you might want to save a series of figures in a for loop, each with a different name. For this you can use the functional form of input arguments, i.e. `export_fig(arg1, arg2)`,  and construct the filename string in a variable. Here's an example of this:  
```Matlab
for a = 1:5
    plot(rand(5, 2));
    export_fig(sprintf('plot%d.png', a));
end
```
When using the functional form like this, be sure to put string variables in quotes:
```Matlab
export_fig(sprintf('plot%d', a), '-a1', '-pdf', '-png');
```

**Specifying the figure/axes** - if you have multiple figures open you can specify which figure to export using its handle:  
```Matlab
export_fig(figure_handle, filename);
```
Equally, if your figure contains several subplots then you can export just one of them by giving export_fig the handle to the relevant axes:
```Matlab
export_fig(axes_handle, filename);
```

**Multiple formats** - save time by exporting to multiple formats simultaneously. E.g.: 
```Matlab
export_fig filename -pdf -eps -png -jpg -tiff
```

**Other file formats** - if you'd like to save your figure to a bitmap format that is not supported by export_fig, e.g. animated GIF, PPM file or a frame in a movie, then you can use export_fig to output the image, and optionally an alpha-matte, to the workspace. E.g.:  
```Matlab
frame = export_fig;
```
or
```Matlab
[frame, alpha] = export_fig;
```
These variables can then be saved to other image formats using other functions, such as imwrite.

**Appending to a file** - you can use the `-append` option to append the figure to the end of an image/document, if it already exists. This is supported for PDF and TIFF files only. Note that if you wish to append a lot of figures consecutively to a PDF, it can be more efficient to save all the figures to PDF separately then append them all in one go at the end (e.g. using [append_pdfs](http://www.mathworks.com/matlabcentral/fileexchange/31215-appendpdfs)).  
  
**Output to clipboard** - you can use the `-clipboard` option to copy the specified figure or axes to the system clipboard, for easy paste into other documents (e.g., Word or PowerPoint). Note that the image is copied in bitmap (not vector) format.  
  
**Font size** - if you want to place an exported figure in a document with the font a particular size then you need to set the font to that size in the figure, and not resize the output of export_fig in the document. To avoid resizing, simply make sure that the on screen figure is the size you want the output to be in the document before exporting.  
  
**Renderers** - MATLAB has three renderers for displaying and exporting figures: painters, OpenGL and ZBuffer. The different renderers have different [features](http://www.mathworks.com/access/helpdesk/help/techdoc/creating_plots/f3-84337.html#f3-102410), so if you aren't happy with the result from one renderer try another. By default, vector formats (i.e. PDF and EPS outputs) use the painters renderer, while other formats use the OpenGL renderer. Non-default renderers can be selected by using one of these three export_fig input options: `-painters`, `-opengl`, `-zbuffer`:  
```Matlab
export_fig test.png -painters
```
  
**Artifacts** - sometimes the output that you get from export_fig is not what you expected. If an output file contains artifacts that aren't in the on screen figure then make sure that the renderer used for rendering the figure on screen is the same as that used for exporting. To set the renderer used to display the figure, use:  
```Matlab
set(figure_handle, 'Renderer', 'opengl');
```
After matching the two renderers, if the artifact appears in the on screen figure then you'll need to fix that before exporting. Alternatively you can try changing the renderer used by export_fig. Finally check that it isn't one of the known issues mentioned in the section below.

**Smoothed/interpolated images in output PDF** - if you produce a PDF using export_fig and images in the PDF look overly smoothed or interpolated, this is because the software you are using to view the PDF is smoothing or interpolating the image data. The image is not smoothed in the PDF file itself. If the software has an option to disable this feature, you should select it. Alternatively, use another PDF viewer that doesn't exhibit this problem.  
  
**Locating Ghostscript/pdftops** - You may find a dialogue box appears when using export_fig, asking you to locate either [Ghostscript](http://www.ghostscript.com) or [pdftops (part of the Xpdf package)](http://www.xpdfreader.com). These are separate applications which export_fig requires to perform certain functions. If such a dialogue appears it is because export_fig can't find the application automatically. This is because you either haven't installed it, or it isn't in the normal place. Make sure you install the applications correctly first. They can be downloaded from the following places:  
 1. Ghostscript:     [www.ghostscript.com](http://www.ghostscript.com)
 2. pdftops (install the Xpdf package): [www.xpdfreader.com](http://www.xpdfreader.com)

If you choose to install them in a non-default location then point export_fig
to this location using the dialogue box.

**Undefined function errors** - If you download and run export_fig and get an error similar to this:  
```
??? Undefined function or method 'print2array' for input arguments of type 'double'.
```
then you are missing one or more of the files that come in the export_fig package. Make sure that you click the "Get from GitHub" button at the top-right of the download [page](http://www.mathworks.co.uk/matlabcentral/fileexchange/23629-exportfig), then extract all the files in the zip file to the same directory. You should then have all the necessary files.
  
### Known issues
There are lots of problems with MATLAB's exporting functions, especially `print`. Export_fig is simply a glorified wrapper for MATLAB's `print` function, and doesn't solve all of its bugs (yet?). Some of the problems I know about are:
  
**Fonts** - when using the painters renderer, MATLAB can only export a small number of fonts, details of which can be found [here](http://www.mathworks.com/help/releases/R2014a/matlab/creating_plots/choosing-a-printer-driver.html#f3-96545). Export_fig attempts to correct font names in the resulting EPS file (up to a maximum of 11 different fonts in one figure), but this is not always guaranteed to work. In particular, the text positions will be affected. It also does not work for text blocks where the 'Interpreter' property is set to 'latex'.

Also, when using the painters renderer, ghostscript will sometimes throw an error such as `Error: /undefined in /findfont`. This suggests that ghostscript could not find a definition file for one of your fonts. One possible fix for this is to make sure the file `EXPORT_FIG_PATH/.ignore/gs_font_path.txt` exists and contains a list of paths to the folder(s) containing the necessary font definitions (make sure that they are TrueType definitions!), separated by a semicolon.

**RGB color data not yet supported in Painter's mode** - you will see this as a warning if you try to export a figure which contains patch objects whose face or vertex colors are specified as an RGB colour, rather than an index into the colormap, using the painters renderer (the default renderer for vector output). This problem can arise if you use `pcolor`, for example. This is a problem with MATLAB's painters renderer, which also affects `print`; there is currently no fix available in export_fig (other than to export to bitmap). The suggested workaround is to avoid colouring patches using RGB. First, try to use colours in the figure's colourmap (instructions [here](http://www.mathworks.co.uk/support/solutions/en/data/1-6OTPQE/)) - change the colourmap, if necessary. If you are using `pcolor`, try using [uimagesc](http://www.mathworks.com/matlabcentral/fileexchange/11368) (on the file exchange) instead.  

**Dashed contour lines appear solid** - when using the painters renderer, MATLAB cannot generate dashed lines using the `contour` function (either on screen or in exported PDF and EPS files). Details can be found [here](http://www.mathworks.com/support/solutions/en/data/1-14PPHB/?solution=1-14PPHB).  
  
**Text size** - when using the OpenGL or ZBuffer renderers, large text can be resized relative to the figure when exporting at non-screen-resolution (including using anti-alising at screen resolution). This is a feature of MATLAB's `print `function. In this case, try using the `-painters` option.  
  
**Lighting and transparency** - when using the painters renderer, transparency and lighting effects are not supported. Sorry, but this is an inherent feature of MATLAB's painters renderer. To find out more about the capabilities of each rendering method, see [here](http://www.mathworks.com/access/helpdesk/help/techdoc/creating_plots/f3-84337.html#f3-102410). You can still export transparent objects to vector format (SVG) using the excellent [plot2svg](http://www.mathworks.com/matlabcentral/fileexchange/7401) package, then convert this to PDF, for example using [Inkscape](http://inkscape.org/). However, it can't handle lighting.  
  
**Lines in patch objects** - when exporting patch objects to PDF using the painters renderer (default), sometimes the output can appear to have lines across the middle of rectangular patches; these lines are the colour of the background, as if there is a crack in the patch, allowing you to see through. This appears to be due to bugs in MATLAB's internal vector rendering code. These lines can often be removed from the PDF using software such as [InkScape](https://inkscape.org). Sometimes disabling anti-aliasing in the PDF-reader software can get rid of the lines ([discussion](https://github.com/altmany/export_fig/issues/44)).  
  
**Out of memory** - if you run into memory issues when using export_fig, some ways to get round this are:  
 1. Reduce the level of anti-aliasing.
 2. Reduce the size of the figure.
 3. Reduce the export resolution (dpi). 
 4. Change the renderer to painters or ZBuffer.  
  
**Errors** - the other common type of errors people get with export_fig are OpenGL errors. This isn't a fault of export_fig, but either a bug in MATLAB's `print`, or your graphics driver getting itself into a state. Always make sure your graphics driver is up-to-date. If it still doesn't work, try using the ZBuffer renderer.  
  
### Raising issues
If you think you have found a genuine error or issue with export_fig **that is not listed above**, first ensure that the figure looks correct on screen when rendered using the renderer that export_fig is set to use (e.g. if exporting to PDF or EPS, does the figure look correct on screen using the painters renderer, or if exporting to bitmap, does the figure look correct on screen using the OpenGL renderer?). If it looks wrong then the problem is there, and I cannot help (other than to suggest you try exporting using a different renderer).

Secondly, if exporting to bitmap, do try all the renderers (i.e. try the options `-opengl`, `-zbuffer` and `-painters` separately), to see if one of them does produce an acceptable output, and if so, use that.

If this still does not help, then ensure that you are using the latest version of export_fig, which is available [here](https://github.com/altmany/export_fig/archive/master.zip).  
 
If the figure looks correct on screen, but an error exists in the exported output (which cannot be solved using a different renderer) then please feel free to raise an [issue](https://github.com/altmany/export_fig/issues). Please be sure to include the .fig file, the export_fig command you use, the output you get, and a description of what you expected. I can't promise anything, but if it's easy to fix I may indeed do it. Often I will find that the error is due to a bug in MATLAB's `print` function, in which case I will suggest you submit it as a bug to TheMathWorks, and inform me of any fix they suggest. Also, if there's a feature you'd like that isn't supported please tell me what it is and I'll consider implementing it.

### And finally...

![](https://farm4.staticflickr.com/3956/15591911455_b9008bd77e_o_d.jpg)

If you've ever wondered what's going on in the logo on the export_fig download page (reproduced here), then this explanantion is for you. The logo is designed to demonstrate as many of export_fig's features as possible: 
 
Given a figure containing a translucent mesh (top right), export_fig can export to pdf (bottom centre), which allows the figure to be zoomed-in without losing quality (because it's a vector graphic), but isn't able to reproduce the translucency. Also, depending on the PDF viewer program, small gaps appear between the patches, which are seen here as thin white lines. 
 
By contrast, when exporting to png (top left), translucency is preserved (see how the graphic below shows through), and the figure is anti-aliased. However, zooming-in does not reveal more detail since png is a bitmap format. Also, lines appear less sharp than in the pdf output.
