## Supplementary material for "In-depth computational analysis of natural and artificial carbon fixation pathways": MATLAB files: user_string.m

```
function [string, file_name] = user_string(string_name, string)
%USER_STRING  Get/set a user specific string
%
% Examples:
%   string  = user_string(string_name)
%   isSaved = user_string(string_name, new_string)
%   [value, file_name] = user_string(...)
%
% Function to get and set a string in a system or user specific file. This
% enables, for example, system specific paths to binaries to be saved.
%
% The specified string will be saved in a file named <string_name>.txt,
% either in a subfolder named .ignore under this file's folder, or in the
% user's prefdir folder (in case this file's folder is non-writable).
%
% IN:
%   string_name - String containing the name of the string required, which
%                 sets the filename storing the string: <string_name>.txt
%   new_string  - The new string to be saved in the <string_name>.txt file
%
% OUT:
%   string  - The currently saved string. Default: ''
%   isSaved - Boolean indicating whether the save was succesful
%   file_name - path of the .txt file used to contain the requested string

% Copyright (C) Oliver Woodford 2011-2014, Yair Altman 2015-

% This method of saving paths avoids changing .m files which might be in a
% version control system. Instead it saves the user dependent paths in
% separate files with a .txt extension, which need not be checked in to
% the version control system. Thank you to Jonas Dorn for suggesting this
% approach.

% 10/01/2013 - Access files in text, not binary mode, as latter can cause
%              errors. Thanks to Christian for pointing this out.
% 29/05/2015 - Save file in prefdir if current folder is non-writable (issue #74)
% 09/01/2018 - Fix issue #232: if the string looks like a file/folder path, ensure it actually exists
% 15/01/2020 - Added file_name output argument

    if ~ischar(string_name)
        error('string_name must be a string.');
    end
    % Create the full filename
    fname = [string_name '.txt'];
    dname = fullfile(fileparts(mfilename('fullpath')), '.ignore');
    file_name = fullfile(dname, fname);
    default_file_name = file_name;
    if nargin > 1
        % Set string
        if ~ischar(string)
            error('new_string must be a string.');
        end
        % Make sure the save directory exists
        %dname = fileparts(file_name);
        if ~exist(dname, 'dir')
            % Create the directory
            try
                if ~mkdir(dname)
                    string = false;
                    return
                end
            catch
                string = false;
                return
            end
            % Make it hidden
            try
                fileattrib(dname, '+h');
            catch
            end
        end
        % Write the file
        fid = fopen(file_name, 'wt');
        if fid == -1
            % file cannot be created/updated - use prefdir if file does not already exist
            % (if file exists but is simply not writable, don't create a duplicate in prefdir)
            if ~exist(file_name,'file')
                file_name = fullfile(prefdir, fname);
                fid = fopen(file_name, 'wt');
            end
            if fid == -1
                string = false;
                file_name = default_file_name;
                return
            end
        end
        try
            fprintf(fid, '%s', string);
        catch
            fclose(fid);
            string = false;
            return
        end
        fclose(fid);
        string = true;
    else
        % Get string
        fid = fopen(file_name, 'rt');
        if fid == -1
            % file cannot be read, try to read the file in prefdir
            file_name = fullfile(prefdir, fname);
            fid = fopen(file_name, 'rt');
            if fid == -1
                string = '';
                file_name = default_file_name;
                return
            end
        end
        string = fgetl(fid);
        fclose(fid);

        % Fix issue #232: if the string looks like a file/folder path, ensure it actually exists
        if ~isempty(string) && any(string=='\' | string=='/') && ~exist(string) %#ok<EXIST>
            string = '';
        end
    end
end
```
