## Supplementary material for "In-depth computational analysis of natural and artificial carbon fixation pathways": MATLAB files: Contents.html

Description of Contents


Home > enzyme-cost-minimization > Contents.m

### Contents

#### PURPOSE

**Functions for Enzyme Cost Minimization**

#### SYNOPSIS

**This is a script file.**

#### DESCRIPTION

```
 Functions for Enzyme Cost Minimization

 Main functions 
   ecm_paramater_balancing      - Run parameter balancing
   ecm_enzyme_cost_minimization - Run enzyme cost minimization
 
 Demos
   See subdirectory 'demo'

 MATLAB Toolboxes required
   Metabolic Network Toolbox - (https://github.com/wolframliebermeister/mnt)
   SBMLtoolbox               - SBML import / export  (see http://sbml.org/Software/SBMLToolbox)
   SBtab toolbox             - SBtab format (https://github.com/wolframliebermeister/sbtab-matlab)
   efmtool                   - Elementary flux modes (see http://www.csb.ethz.ch/tools/efmtool)

 (C) 2015
 Wolfram Liebermeister  <>
```

#### CROSS-REFERENCE INFORMATION

This function calls:


This function is called by:


---

Generated on Mon 30-Jan-2017 18:21:00 by **m2html** © 2003
