## Supplementary material for "In-depth computational analysis of natural and artificial carbon fixation pathways": MATLAB files: Contents.html

Description of Contents


Home > enzyme-cost-minimization > demo > Contents.m

### Contents

#### PURPOSE

**Demo files for Enzyme Cost Minimisation**

#### SYNOPSIS

**This is a script file.**

#### DESCRIPTION

```
Demo files for Enzyme Cost Minimisation
```

#### CROSS-REFERENCE INFORMATION

This function calls:


This function is called by:


---

Generated on Mon 30-Jan-2017 18:21:00 by **m2html** © 2003
