## Supplementary material for "In-depth computational analysis of natural and artificial carbon fixation pathways": MATLAB files: demo_escherichia_coli_ccm.html

Description of demo\_escherichia\_coli\_ccm


Home > enzyme-cost-minimization > demo > demo\_escherichia\_coli\_ccm.m

### demo\_escherichia\_coli\_ccm

#### PURPOSE

**Demo script for Parameter Balancing and Enzyme Cost Minimization**

#### SYNOPSIS

**This is a script file.**

#### DESCRIPTION

```
 Demo script for Parameter Balancing and Enzyme Cost Minimization

 Prepared model files for E coli central metabolism.are used.
 
 The MATLAB function 'ecm_simple' is a wrapper function 
 that reads the data, sets some default parameters, and calls 
 dedicated functions for Parameter Balancing or Enzyme Cost Minimization.

 The input files (in SBtab format) are given in the subdirectory 'data'
 The results (in SBtab format) are written to the subdirectory 'results'
```

#### CROSS-REFERENCE INFORMATION

This function calls:

- ecm\_BASEDIR ECM\_BASEDIR - Return ECM toolbox directory
- ecm\_simple ECM\_SIMPLE - Wrapper script for Parameter Balancing or Enzyme Cost Minimisation

This function is called by:


---

Generated on Mon 30-Jan-2017 18:21:00 by **m2html** © 2003
