## Supplementary material for "In-depth computational analysis of natural and artificial carbon fixation pathways": MATLAB files: index.html

Index for Directory ./enzyme-cost-minimization/demo


|  |  |
| --- | --- |
| Master index | Index for ./enzyme-cost-minimization/demo |

### Index for ./enzyme-cost-minimization/demo

#### Matlab files in this directory:

|  |  |
| --- | --- |
| Contents | Demo files for Enzyme Cost Minimisation |
| demo\_escherichia\_coli\_ccm | Demo script for Parameter Balancing and Enzyme Cost Minimization |

#### Subsequent directories:

- data
- results

---

Generated on Mon 30-Jan-2017 18:21:00 by **m2html** © 2003
