## Supplementary material for "In-depth computational analysis of natural and artificial carbon fixation pathways": MATLAB files: ecm_ecf4cmr.html

Description of ecm\_ecf4cmr


Home > enzyme-cost-minimization > ecf\_scores > ecm\_ecf4cmr.m

### ecm\_ecf4cmr

#### PURPOSE

**[u\_tot, u] = ecm\_ecf4cmr(x,pp)**

#### SYNOPSIS

**function [u\_tot, u, w] = ecm\_ecf4cmr(x,pp)**

#### DESCRIPTION

```
 [u_tot, u] = ecm_ecf4cmr(x,pp)
```

#### CROSS-REFERENCE INFORMATION

This function calls:


This function is called by:

- ecm\_ecf4geom [u\_tot, u] = ecm\_ecf4geom(x,pp)
- ecm\_get\_score
- ecm\_get\_specific\_rates


---

Generated on Tue 29-Mar-2016 12:27:24 by **m2html** © 2003
