## Supplementary material for "In-depth computational analysis of natural and artificial carbon fixation pathways": MATLAB files: ecm_mdf.html

Description of ecm\_mdf


Home > enzyme-cost-minimization > ecf\_scores > ecm\_mdf.m

### ecm\_mdf

#### PURPOSE

**[f,u] = ecm\_mdf(x,pp)**

#### SYNOPSIS

**function [f,u,w] = ecm\_mdf(x,pp)**

#### DESCRIPTION

```
 [f,u] = ecm_mdf(x,pp)
```

#### CROSS-REFERENCE INFORMATION

This function calls:


This function is called by:

- ecm\_enzyme\_cost\_minimization ECM\_ENZYME\_COST\_MINIMIZATION - Compute optimal flux-specific enzyme costs for given flux distribution
- ecm\_get\_score
- ecm\_get\_specific\_rates


---

Generated on Tue 29-Mar-2016 12:27:24 by **m2html** © 2003
