## Supplementary material for "In-depth computational analysis of natural and artificial carbon fixation pathways": MATLAB files: ecm_min_ecf2sp.html

Description of ecm\_min\_ecf2sp


Home > enzyme-cost-minimization > ecf\_scores > ecm\_min\_ecf2sp.m

### ecm\_min\_ecf2sp

#### PURPOSE

**[f, u] = ecm\_min\_ecf2sp(x,pp)**

#### SYNOPSIS

**function [f, u, w] = ecm\_min\_ecf2sp(x,pp)**

#### DESCRIPTION

```
 [f, u] = ecm_min_ecf2sp(x,pp)
```

#### CROSS-REFERENCE INFORMATION

This function calls:


This function is called by:

- ecm\_get\_score
- ecm\_get\_specific\_rates


---

Generated on Tue 29-Mar-2016 12:27:24 by **m2html** © 2003
