## Supplementary material for "In-depth computational analysis of natural and artificial carbon fixation pathways": MATLAB files: index.html

Index for Directory ./enzyme-cost-minimization/ecf\_scores


|  |  |
| --- | --- |
| Master index | Index for ./enzyme-cost-minimization/ecf\_scores |

### Index for ./enzyme-cost-minimization/ecf\_scores

#### Matlab files in this directory:

|  |  |
| --- | --- |
| ecm\_ecf1 | [u\_tot, u] = ecm\_ecf1(x,pp) |
| ecm\_ecf2s | [u\_tot, u] = ecm\_ecf2s(x,pp) |
| ecm\_ecf2sp | [u\_tot, u] = ecm\_ecf2sp(x,pp) |
| ecm\_ecf3s | [u\_tot, u] = ecm\_ecf3s(x,pp) |
| ecm\_ecf3sp | [u\_tot, u] = ecm\_ecf3sp(x,pp) |
| ecm\_ecf4cmr | [u\_tot, u] = ecm\_ecf4cmr(x,pp) |
| ecm\_ecf4dmr | [u\_tot, u] = ecm\_ecf4cmr(x,pp) |
| ecm\_ecf4geom | [u\_tot, u] = ecm\_ecf4geom(x,pp) |
| ecm\_ecf4smr | [u\_tot, u] = ecm\_ecf4cmr(x,pp) |
| ecm\_mdf | [f,u] = ecm\_mdf(x,pp) |
| ecm\_min\_ecf2s | [f, u] = ecm\_ecf2s(x,pp) |
| ecm\_min\_ecf2sp | [f, u] = ecm\_min\_ecf2sp(x,pp) |

---

Generated on Tue 29-Mar-2016 12:27:23 by **m2html** © 2003
