## Supplementary material for "In-depth computational analysis of natural and artificial carbon fixation pathways": MATLAB files: ecm_BASEDIR.html

Description of ecm\_BASEDIR


Home > enzyme-cost-minimization > ecm\_BASEDIR.m

### ecm\_BASEDIR

#### PURPOSE

#### SYNOPSIS

**function d = ecm\_BASEDIR()**

#### DESCRIPTION

#### CROSS-REFERENCE INFORMATION

This function calls:


This function is called by:

- demo\_escherichia\_coli\_ccm --------------------------------------------------------
- ecm\_setup

#### SOURCE CODE

```
0001 function d = ecm_BASEDIR()
0002 
0003 d = [fileparts(which(mfilename)) '/'];
```

---

Generated on Thu 12-Feb-2015 14:18:00 by **m2html** © 2003
