## Supplementary material for "In-depth computational analysis of natural and artificial carbon fixation pathways": MATLAB files: ecm_enzyme_cost_minimization.html

Description of ecm\_enzyme\_cost\_minimization


Home > enzyme-cost-minimization > ecm\_enzyme\_cost\_minimization.m

### ecm\_enzyme\_cost\_minimization

#### PURPOSE

**ECM\_ENZYME\_COST\_MINIMIZATION - Compute optimal flux-specific enzyme costs for given flux distribution**

#### SYNOPSIS

**function [c, u, u\_cost, up, A\_forward, mca\_info, c\_min, c\_max, u\_min, u\_max, r, u\_capacity, eta\_energetic, eta\_saturation] = ecm\_enzyme\_cost\_minimization(network,r,v,ecm\_options)**

#### DESCRIPTION

```
 ECM_ENZYME_COST_MINIMIZATION - Compute optimal flux-specific enzyme costs for given flux distribution

 [c, u, u_cost, up, A_forward, mca_info, c_min, c_max, u_min, u_max, r, u_capacity, eta_energetic, eta_saturation] = ecm_enzyme_cost_minimization(network, e, v, ecm_options)

 Input 
   network       metabolic network structure (as in Metabolic Network Toolbox)
   r             Kinetic constants (from parameter balancing)
   v             flux mode
   ecm_options   options struct (for fields and default values, see 'ecm_default_options')

 Output
   c                 metabolite concentrations
   c.data            data vector (if provided)
   c.fixed           fixed concentration vector
   c.initial         initial solution vector 
   c.[SCORE]         1st column: result from optimising the score [SCORE]
                     other columns: possible sampled solutions
                     
   u                 enzyme levels (all enzymes)
   u.data            data vector
   u.[SCORE]         1st column: result from optimising the score [SCORE]
                     other columns: possible sampled solutions

   up                enzyme levels (only scored enzymes as defined by 
                                    ecm_scores.ind_scored_enzymes)
   up.[SCORE]        1st column: result from optimising the score [SCORE]
                     other columns: possible sampled solutions

   u_cost            sum of scored enzyme levels 
   u_cost.data       data value
   u_cost.[SCORE]    1st column: result from optimising the score [SCORE]
                     other columns: possible sampled solutions

   A_forward         reaction affinity (in flux direction)
   A_forward.initial    initial solution vector 
   A_forward.[SCORE] 1st column: result from optimising the score [SCORE]
                     other columns: possible sampled solutions
```

#### CROSS-REFERENCE INFORMATION

This function calls:

- ecm\_below\_threshold ECM\_BELOW\_THRESHOLD - Helper function for enzyme cost minimization
- ecm\_get\_one\_u ECM\_GET\_ONE\_U - Helper function for enzyme variability calculation after enzyme cost minimization:
- ecm\_get\_score ECM\_GET\_SCORE - Helper function
- ecm\_get\_specific\_rates ECM\_GET\_SPECIFIC\_RATES - Helper function
- ecm\_inequalities MEASURES\_FOR\_ENZYME\_COSTS\_INEQUALITIES - Helper function
- ecm\_one\_run ECM\_ONE\_RUN - Perform one ECM run
- ecm\_regularisation ECM\_REGULARISATION - Regularisation term for ECM
- ecm\_mdf [f,u] = ecm\_mdf(x,pp)

This function is called by:

- ecm\_simple ECM\_SIMPLE - Wrapper script for Parameter Balancing or Enzyme Cost Minimisation


---

Generated on Mon 30-Jan-2017 18:21:00 by **m2html** © 2003
