## Supplementary material for "In-depth computational analysis of natural and artificial carbon fixation pathways": MATLAB files: ecm_BASEDIR.html

Description of ecm\_BASEDIR


Home > enzyme-cost-minimization > ecm\_functions > ecm\_BASEDIR.m

### ecm\_BASEDIR

#### PURPOSE

**ECM\_BASEDIR - Return ECM toolbox directory**

#### SYNOPSIS

**function d = ecm\_BASEDIR()**

#### DESCRIPTION

```
 ECM_BASEDIR - Return ECM toolbox directory

 d = ecm_BASEDIR()
```

#### CROSS-REFERENCE INFORMATION

This function calls:


This function is called by:

- demo\_escherichia\_coli\_ccm Demo script for Parameter Balancing and Enzyme Cost Minimization
- ecm\_setup ECM\_SETUP - Determine data directory


---

Generated on Mon 30-Jan-2017 18:21:00 by **m2html** © 2003
