## Supplementary material for "In-depth computational analysis of natural and artificial carbon fixation pathways": MATLAB files: ecm_below_threshold.html

Description of ecm\_below\_threshold


Home > enzyme-cost-minimization > ecm\_functions > ecm\_below\_threshold.m

### ecm\_below\_threshold

#### PURPOSE

**ECM\_BELOW\_THRESHOLD - Helper function for enzyme cost minimization**

#### SYNOPSIS

**function [ineq\_constraints, eq\_constraints] = ecm\_below\_threshold(ecm\_score,xx,pp,u\_threshold,x\_min,x\_max,ecm\_options)**

#### DESCRIPTION

```
 ECM_BELOW_THRESHOLD - Helper function for enzyme cost minimization
 
 function [ineq_constraints, eq_constraints] = ecm_below_threshold(ecm_score,xx,pp,u_threshold,x_min,x_max,ecm_options)
```

#### CROSS-REFERENCE INFORMATION

This function calls:

- ecm\_get\_score ECM\_GET\_SCORE - Helper function
- ecm\_inequalities MEASURES\_FOR\_ENZYME\_COSTS\_INEQUALITIES - Helper function
- ecm\_regularisation ECM\_REGULARISATION - Regularisation term for ECM

This function is called by:

- ecm\_enzyme\_cost\_minimization ECM\_ENZYME\_COST\_MINIMIZATION - Compute optimal flux-specific enzyme costs for given flux distribution


---

Generated on Mon 30-Jan-2017 18:21:00 by **m2html** © 2003
