## Supplementary material for "In-depth computational analysis of natural and artificial carbon fixation pathways": MATLAB files: ecm_check_parameter_balancing.html

Description of ecm\_check\_parameter\_balancing


Home > enzyme-cost-minimization > ecm\_functions > ecm\_check\_parameter\_balancing.m

### ecm\_check\_parameter\_balancing

#### PURPOSE

**ECM\_CHECK\_PARAMETER\_BALANCING - Checks for balanced parameters**

#### SYNOPSIS

**function ecm\_check\_parameter\_balancing(r, r\_orig, network, quantity\_info\_used, show\_graphics)**

#### DESCRIPTION

```
 ECM_CHECK_PARAMETER_BALANCING - Checks for balanced parameters

 ecm_check_parameter_balancing(r, r_orig, network, quantity_info_used, show_graphics)
```

#### CROSS-REFERENCE INFORMATION

This function calls:


This function is called by:


---

Generated on Mon 30-Jan-2017 18:21:00 by **m2html** © 2003
