## Supplementary material for "In-depth computational analysis of natural and artificial carbon fixation pathways": MATLAB files: ECM_data_dir.html

Description of ECM\_data\_dir


Home > enzyme-cost-minimization > ecm\_functions > ECM\_data\_dir.m

### ECM\_data\_dir

#### PURPOSE

#### SYNOPSIS

**function f = ecm\_data\_dir()**

#### DESCRIPTION

#### CROSS-REFERENCE INFORMATION

This function calls:

- ecm\_setup

This function is called by:


#### SOURCE CODE

```
0001 function f = ecm_data_dir()
0002 
0003 ecm_info = ecm_setup;
0004 
0005 f = ecm_info.data_dir;
```

---

Generated on Thu 12-Feb-2015 14:14:13 by **m2html** © 2003
