## Supplementary material for "In-depth computational analysis of natural and artificial carbon fixation pathways": MATLAB files: ecm_default_options.html

Description of ecm\_default\_options


Home > enzyme-cost-minimization > ecm\_functions > ecm\_default\_options.m

### ecm\_default\_options

#### PURPOSE

**ECM\_DEFAULT\_OPTIONS - Defaults for ECM**

#### SYNOPSIS

**function ecm\_options = ecm\_default\_options(network, model\_name)**

#### DESCRIPTION

```
 ECM_DEFAULT_OPTIONS -  Defaults for ECM
 
 ecm_options = ecm_default_options(network, model_name)

 ecm options and their default values

 model
   ecm_options.model_name               = model_name         ; 
   ecm_options.run_id                   = 'RUN';
   ecm_options.model_id                 = 'MODEL';
   ecm_options.network_CoHid            = network;
 
 metabolite constraints
   ecm_options.fix_metabolites          = {}; % metabolites with fixed concentrations (overrides values from input model)
   ecm_options.fix_metabolite_values    = [];
   ecm_options.conc_min_default         = 0.001              ; % mM
   ecm_options.conc_max_default         = 10                 ; % mM
   ecm_options.conc_min                 = [];% 0.001 * ones(nm,1) ; % mM
   ecm_options.conc_max                 = [];% 10 * ones(nm,1);   ; % mM
   ecm_options.conc_fix                 = [];
   ecm_options.met_fix                  = [];
   ecm_options.replace_cofactors        = {};
 
 given data
   ecm_options.c_data                   = [];
   ecm_options.u_data                   = [];
   ecm_options.kinetic_data             = [];
 
 kinetic data
   ecm_options.reaction_column_names    = []; % column names (in data file) for loading of kinetic data
   ecm_options.compound_column_names    = [];
   ecm_options.KM_lower                 = []; % mM
   ecm_options.Keq_upper                = [];
   ecm_options.flag_given_kinetics      = 0;
   ecm_options.kcat_usage               = 'use';
   ecm_options.kcat_upper               = 10000;  % 1/s
   ecm_options.kcat_lower               = 0.1;    % 1/s
   ecm_options.kcatr_lower              = 0.0001; % 1/s
   ecm_options.kcat_prior_median        = 10;     % for ccm this value need to be modified
   ecm_options.kcat_prior_log10_std     = 0.2;   % reduce spread of kcat values
   ecm_options.GFE_fixed                = 1;     % flag
   ecm_options.insert_Keq_from_data     = 0;     % flag
 
 parameter balancing
   ecm_options.n_samples = 0;
   ecm_options.use_pseudo_values = 1;
 
 enzyme cost weights
   ecm_options.ind_scored_enzymes       = 1:length(network.actions);
   ecm_options.enzyme_cost_weights      = ones(length(ecm_options.ind_scored_enzymes),1);
   ecm_options.use_cost_weights         = 'none';
 
 ecm
   ecm_options.initial_choice           = 'mdf'; 
   ecm_options.multiple_starting_points = 0;
   ecm_options.ecm_scores               = {'emc3sp'}           ;
   ecm_options.lambda_regularisation    = 10^-3; 
   ecm_options.lambda_reg_factor        = 0.01;
   ecm_options.quantity_info_file       = [];
   ecm_options.compute_hessian          = 0;
   ecm_options.compute_elasticities     = 0;
   ecm_options.compute_tolerance        = 0;
   ecm_options.cost_tolerance_factor    = 1.01; % one percent
   ecm_options.tolerance_from_hessian   = 0;
   ecm_options.fix_Keq_in_sampling      = 0;
 
 graphics
   ecm_options.print_graphics           = 0;
   ecm_options.show_graphics            = 1;
   ecm_options.show_metabolites         = network.metabolites;
```

#### CROSS-REFERENCE INFORMATION

This function calls:


This function is called by:

- ecm\_update\_options ECM\_UPDATE\_OPTIONS - Helper function for ECM options (struct 'ecm\_options')
- ecm\_parameter\_balancing ECM\_PARAMETER\_BALANCING - Prepare and run parameter balancing
- ecm\_simple ECM\_SIMPLE - Wrapper script for Parameter Balancing or Enzyme Cost Minimisation


---

Generated on Mon 30-Jan-2017 18:21:00 by **m2html** © 2003
