## Supplementary material for "In-depth computational analysis of natural and artificial carbon fixation pathways": MATLAB files: ecm_dependencies.html

Description of ecm\_dependencies


Home > enzyme-cost-minimization > ecm\_functions > ecm\_dependencies.m

### ecm\_dependencies

#### PURPOSE

**ECM\_DEPENDENCIES - Check whether dependencies of ECM toolbox are satisfied**

#### SYNOPSIS

**function ecm\_dependencies()**

#### DESCRIPTION

```
 ECM_DEPENDENCIES - Check whether dependencies of ECM toolbox are satisfied
```

#### CROSS-REFERENCE INFORMATION

This function calls:


This function is called by:


---

Generated on Mon 30-Jan-2017 18:21:00 by **m2html** © 2003
