## Supplementary material for "In-depth computational analysis of natural and artificial carbon fixation pathways": MATLAB files: ecm_display.html

Description of ecm\_display


Home > enzyme-cost-minimization > ecm\_functions > ecm\_display.m

### ecm\_display

#### PURPOSE

**ECM\_DISPLAY - Display results of ECM**

#### SYNOPSIS

**function ecm\_display(ecm\_options,graphics\_options,network,v,c,u,u\_tot,up,A\_forward,r,kinetic\_data,c\_min,c\_max,u\_min,u\_max,u\_capacity,eta\_energetic,eta\_saturation)**

#### DESCRIPTION

```
 ECM_DISPLAY - Display results of ECM

 ecm_display(network,options,ecm_options,v,c,u,u_tot,up,A_forward,r,kinetic_data,c_min,c_max,u_min,u_max)

 ECM_DISPLAY - Display results of Enzyme Cost Minimization

 Options in 'graphics_options':

  graphics_options.few_graphics                    Omit many graphics
  graphics_options.show_proteomaps                 Show proteomaps
  graphics_options.show_original_data              Show graphs with original data
  graphics_options.show_network_graphics           Show network graphs
  graphics_options.show_matrix_graphics            Show matrix graphs
  graphics_options.reaction_order_file             Filename for reordering reactions
  graphics_options.metabolites_order_file          Filename for reordering compounds
```

#### CROSS-REFERENCE INFORMATION

This function calls:


This function is called by:

- ecm\_simple ECM\_SIMPLE - Wrapper script for Parameter Balancing or Enzyme Cost Minimisation


---

Generated on Mon 30-Jan-2017 18:21:00 by **m2html** © 2003
