## Supplementary material for "In-depth computational analysis of natural and artificial carbon fixation pathways": MATLAB files: ecm_get_one_u.html

Description of ecm\_get\_one\_u


Home > enzyme-cost-minimization > ecm\_functions > ecm\_get\_one\_u.m

### ecm\_get\_one\_u

#### PURPOSE

**ECM\_GET\_ONE\_U - Helper function for enzyme variability calculation after enzyme cost minimization:**

#### SYNOPSIS

**function my\_u\_it = ecm\_get\_one\_u(it,ecm\_score,xx,pp,u\_threshold,x\_min,x\_max,ecm\_options)**

#### DESCRIPTION

```
 ECM_GET_ONE_U - Helper function for enzyme variability calculation after enzyme cost minimization:

 function my_u_it = ecm_get_one_u(it,ecm_score,xx,pp,u_threshold,x_min,x_max,ecm_options)

 given log metabolite profile xx, evaluate enzyme levels 
 and return the levels of the it'th enzyme
```

#### CROSS-REFERENCE INFORMATION

This function calls:

- ecm\_get\_score ECM\_GET\_SCORE - Helper function

This function is called by:

- ecm\_enzyme\_cost\_minimization ECM\_ENZYME\_COST\_MINIMIZATION - Compute optimal flux-specific enzyme costs for given flux distribution


---

Generated on Mon 30-Jan-2017 18:21:00 by **m2html** © 2003
