## Supplementary material for "In-depth computational analysis of natural and artificial carbon fixation pathways": MATLAB files: ecm_get_score.html

Description of ecm\_get\_score


Home > enzyme-cost-minimization > ecm\_functions > ecm\_get\_score.m

### ecm\_get\_score

#### PURPOSE

**ECM\_GET\_SCORE - Helper function**

#### SYNOPSIS

**function [u\_cost, u] = ecm\_get\_score(ecm\_score,x,pp)**

#### DESCRIPTION

```
 ECM_GET_SCORE - Helper function
 
 function [u_cost, u] = ecm_get_score(ecm_score,x,pp)
```

#### CROSS-REFERENCE INFORMATION

This function calls:

- ecm\_emc1 [u\_tot, u] = ecm\_emc1(x,pp)
- ecm\_emc2s [u\_tot, u] = ecm\_emc2s(x,pp)
- ecm\_emc2sp [u\_tot, u] = ecm\_emc2sp(x,pp)
- ecm\_emc3s [u\_tot, u] = ecm\_emc3s(x,pp)
- ecm\_emc3sp [u\_tot, u] = ecm\_emc3sp(x,pp)
- ecm\_emc4cm [u\_tot, u] = ecm\_emc4cmr(x,pp)
- ecm\_emc4dm [u\_tot, u] = ecm\_emc4cmr(x,pp)
- ecm\_emc4geom [u\_tot, u] = ecm\_emc4geom(x,pp)
- ecm\_emc4sm [u\_tot, u] = ecm\_emc4cmr(x,pp)
- ecm\_mdf [f,u] = ecm\_mdf(x,pp)
- ecm\_memc2s [f, u] = ecm\_emc2s(x,pp)
- ecm\_memc2sp [f, u] = ecm\_min\_emc2sp(x,pp)

This function is called by:

- ecm\_enzyme\_cost\_minimization ECM\_ENZYME\_COST\_MINIMIZATION - Compute optimal flux-specific enzyme costs for given flux distribution
- ecm\_below\_threshold ECM\_BELOW\_THRESHOLD - Helper function for enzyme cost minimization
- ecm\_get\_one\_u ECM\_GET\_ONE\_U - Helper function for enzyme variability calculation after enzyme cost minimization:
- ecm\_one\_run ECM\_ONE\_RUN - Perform one ECM run


---

Generated on Mon 30-Jan-2017 18:21:00 by **m2html** © 2003
