## Supplementary material for "In-depth computational analysis of natural and artificial carbon fixation pathways": MATLAB files: ecm_get_specific_rates.html

Description of ecm\_get\_specific\_rates


Home > enzyme-cost-minimization > ecm\_functions > ecm\_get\_specific\_rates.m

### ecm\_get\_specific\_rates

#### PURPOSE

**ECM\_GET\_SPECIFIC\_RATES - Helper function**

#### SYNOPSIS

**function rates = ecm\_get\_specific\_rates(ecm\_score,x,pp,ind)**

#### DESCRIPTION

```
 ECM_GET_SPECIFIC_RATES - Helper function

 function rates = ecm_get_specific_rates(ecm_score,x,pp,ind)
```

#### CROSS-REFERENCE INFORMATION

This function calls:

- ecm\_emc1 [u\_tot, u] = ecm\_emc1(x,pp)
- ecm\_emc2s [u\_tot, u] = ecm\_emc2s(x,pp)
- ecm\_emc2sp [u\_tot, u] = ecm\_emc2sp(x,pp)
- ecm\_emc3s [u\_tot, u] = ecm\_emc3s(x,pp)
- ecm\_emc3sp [u\_tot, u] = ecm\_emc3sp(x,pp)
- ecm\_emc4cm [u\_tot, u] = ecm\_emc4cmr(x,pp)
- ecm\_emc4dm [u\_tot, u] = ecm\_emc4cmr(x,pp)
- ecm\_emc4geom [u\_tot, u] = ecm\_emc4geom(x,pp)
- ecm\_emc4sm [u\_tot, u] = ecm\_emc4cmr(x,pp)
- ecm\_mdf [f,u] = ecm\_mdf(x,pp)
- ecm\_memc2s [f, u] = ecm\_emc2s(x,pp)
- ecm\_memc2sp [f, u] = ecm\_min\_emc2sp(x,pp)

This function is called by:

- ecm\_enzyme\_cost\_minimization ECM\_ENZYME\_COST\_MINIMIZATION - Compute optimal flux-specific enzyme costs for given flux distribution


---

Generated on Mon 30-Jan-2017 18:21:00 by **m2html** © 2003
