## Supplementary material for "In-depth computational analysis of natural and artificial carbon fixation pathways": MATLAB files: ecm_inequalities.html

Description of ecm\_inequalities


Home > enzyme-cost-minimization > ecm\_functions > ecm\_inequalities.m

### ecm\_inequalities

#### PURPOSE

**MEASURES\_FOR\_ENZYME\_COSTS\_INEQUALITIES - Helper function**

#### SYNOPSIS

**function [delta\_G\_by\_RT,eq\_cons] = measures\_for\_enzyme\_costs\_inequalities(x,N\_forward,log\_Keq\_forward)**

#### DESCRIPTION

```
 MEASURES_FOR_ENZYME_COSTS_INEQUALITIES - Helper function 

 [delta_G_by_RT,eq_cons] = measures_for_enzyme_costs_inequalities(x,N_forward,log_Keq_forward)
```

#### CROSS-REFERENCE INFORMATION

This function calls:


This function is called by:

- ecm\_enzyme\_cost\_minimization ECM\_ENZYME\_COST\_MINIMIZATION - Compute optimal flux-specific enzyme costs for given flux distribution
- ecm\_below\_threshold ECM\_BELOW\_THRESHOLD - Helper function for enzyme cost minimization
- ecm\_one\_run ECM\_ONE\_RUN - Perform one ECM run


---

Generated on Mon 30-Jan-2017 18:21:00 by **m2html** © 2003
