## Supplementary material for "In-depth computational analysis of natural and artificial carbon fixation pathways": MATLAB files: ecm_kegg_compound_id_to_name.html

Description of ecm\_kegg\_compound\_id\_to\_name


Home > enzyme-cost-minimization > ecm\_functions > ecm\_kegg\_compound\_id\_to\_name.m

### ecm\_kegg\_compound\_id\_to\_name

#### PURPOSE

**ECM\_KEGG\_COMPOUND\_ID\_TO\_NAME - Name conversion for compounds**

#### SYNOPSIS

**function names = ecm\_kegg\_compound\_id\_to\_name(ids,kegg\_conversion\_file)**

#### DESCRIPTION

```
 ECM_KEGG_COMPOUND_ID_TO_NAME - Name conversion for compounds

 function names = ecm_kegg_compound_id_to_name(ids,kegg_conversion_file)
```

#### CROSS-REFERENCE INFORMATION

This function calls:

- ecm\_setup ECM\_SETUP - Determine data directory

This function is called by:

- ecm\_sbtab2mnt [network, v, conc\_fix, kinetic\_data] = ecm\_sbtab2mnt(model\_filename, model\_dir, matlab\_dir,kinetic\_data\_file\_names)


---

Generated on Mon 30-Jan-2017 18:21:00 by **m2html** © 2003
