## Supplementary material for "In-depth computational analysis of natural and artificial carbon fixation pathways": MATLAB files: ecm_load_model_and_data_sbtab.html

Description of ecm\_load\_model\_and\_data\_sbtab


Home > enzyme-cost-minimization > ecm\_functions > ecm\_load\_model\_and\_data\_sbtab.m

### ecm\_load\_model\_and\_data\_sbtab

#### PURPOSE

**ECM\_LOAD\_MODEL\_AND\_DATA\_SBTAB - Load data from SBtab file**

#### SYNOPSIS

**function [network,v,c\_data,u\_data, conc\_min, conc\_max, met\_fix, conc\_fix,positions, enzyme\_cost\_weights, warnings] = ecm\_load\_model\_and\_data\_sbtab(filename, tmp\_dir)**

#### DESCRIPTION

```
 ECM_LOAD_MODEL_AND_DATA_SBTAB - Load data from SBtab file

 [network,v,c_data,u_data, conc_min, conc_max, positions, warnings] = ecm_load_model_and_data_sbtab(filename, tmp_dir)

Load SBtab file containing (model and data) information for Enzyme Cost Minimization

For saving an SBtab file, see 'help ecm_save_model_and_data_sbtab'

Arguments
 filename               filename for SBtab output
 tmp_dir                a directory to which a temporary file can be written (needed for technical reasons)

Output
 network                (struct describing model, see mnt toolbox)
 v                      (nr x 1 vector of reaction rates)
 r                      (struct describing model kinetics, see mnt toolbox)
 c_data                 (nm x 1 vector of measured concentrations (only for information))
 u_data                 (nr x 1 vector of measured enzyme concentrations (only for information))
 kinetic_data           (OPTIONAL: struct with kinetic data; only to give original dmu0 values)
 conc_min               (nm x 1 vector of minimal concentrations)
 conc_max               (nm x 1 vector of maximal concentrations)
 met_fix                (OPTIONAL: list of metabolites with fixed concentrations)
 conc_fix               (OPTIONAL: fixed concentrations corresponding to met_fix)
 enzyme_cost_weights    ( nr x 1 vector of enzyme cost weights; default [])
 save_single_tables     (flag for saving SBtab tables in single files; default 0)
```

#### CROSS-REFERENCE INFORMATION

This function calls:


This function is called by:

- ecm\_simple ECM\_SIMPLE - Wrapper script for Parameter Balancing or Enzyme Cost Minimisation


---

Generated on Mon 30-Jan-2017 18:21:00 by **m2html** © 2003
