## Supplementary material for "In-depth computational analysis of natural and artificial carbon fixation pathways": MATLAB files: ecm_one_run.html

Description of ecm\_one\_run


Home > enzyme-cost-minimization > ecm\_functions > ecm\_one\_run.m

### ecm\_one\_run

#### PURPOSE

**ECM\_ONE\_RUN - Perform one ECM run**

#### SYNOPSIS

**function [my\_c, my\_u, my\_up, my\_u\_cost, my\_A\_forward, my\_x, my\_grad, my\_lambda] = ecm\_one\_run(ecm\_score,pp,x\_min,x\_max,x\_init,ecm\_options,opt)**

#### DESCRIPTION

```
 ECM_ONE_RUN - Perform one ECM run
 
 function [my_c, my_u, my_up, my_u_cost, my_A_forward, my_x, my_grad, my_lambda] = ecm_one_run(ecm_score,pp,x_min,x_max,x_init,ecm_options,opt)
```

#### CROSS-REFERENCE INFORMATION

This function calls:

- ecm\_get\_score ECM\_GET\_SCORE - Helper function
- ecm\_inequalities MEASURES\_FOR\_ENZYME\_COSTS\_INEQUALITIES - Helper function
- ecm\_regularisation ECM\_REGULARISATION - Regularisation term for ECM

This function is called by:

- ecm\_enzyme\_cost\_minimization ECM\_ENZYME\_COST\_MINIMIZATION - Compute optimal flux-specific enzyme costs for given flux distribution


---

Generated on Mon 30-Jan-2017 18:21:00 by **m2html** © 2003
