## Supplementary material for "In-depth computational analysis of natural and artificial carbon fixation pathways": MATLAB files: ecm_read_options.html

Description of ecm\_read\_options


Home > enzyme-cost-minimization > ecm\_functions > ecm\_read\_options.m

### ecm\_read\_options

#### PURPOSE

**ECM\_READ\_OPTIONS - Read ECM options from file**

#### SYNOPSIS

**function options = ecm\_read\_options(options\_file, default\_options)**

#### DESCRIPTION

```
 ECM_READ_OPTIONS - Read ECM options from file
 
 options = ecm_read_options(options_file, default_options)
```

#### CROSS-REFERENCE INFORMATION

This function calls:


This function is called by:


---

Generated on Mon 30-Jan-2017 18:21:00 by **m2html** © 2003
