## Supplementary material for "In-depth computational analysis of natural and artificial carbon fixation pathways": MATLAB files: ecm_regularisation.html

Description of ecm\_regularisation


Home > enzyme-cost-minimization > ecm\_functions > ecm\_regularisation.m

### ecm\_regularisation

#### PURPOSE

**ECM\_REGULARISATION - Regularisation term for ECM**

#### SYNOPSIS

**function f = ecm\_regularisation(x,x\_min,x\_max,lambda)**

#### DESCRIPTION

```
 ECM_REGULARISATION - Regularisation term for ECM 
 
 function f = ecm_regularisation(x,x_min,x_max,lambda)
```

#### CROSS-REFERENCE INFORMATION

This function calls:


This function is called by:

- ecm\_enzyme\_cost\_minimization ECM\_ENZYME\_COST\_MINIMIZATION - Compute optimal flux-specific enzyme costs for given flux distribution
- ecm\_below\_threshold ECM\_BELOW\_THRESHOLD - Helper function for enzyme cost minimization
- ecm\_one\_run ECM\_ONE\_RUN - Perform one ECM run


---

Generated on Mon 30-Jan-2017 18:21:00 by **m2html** © 2003
