## Supplementary material for "In-depth computational analysis of natural and artificial carbon fixation pathways": MATLAB files: ecm_save_model_and_data_gams.html

Description of ecm\_save\_model\_and\_data\_gams


Home > enzyme-cost-minimization > ecm\_functions > ecm\_save\_model\_and\_data\_gams.m

### ecm\_save\_model\_and\_data\_gams

#### PURPOSE

**ECM\_SAVE\_MODEL\_AND\_DATA\_GAMS - Save input files for ECM by GAMS solvers**

#### SYNOPSIS

**function ecm\_save\_model\_and\_data\_gams(filename,network,v,r,c\_data,u\_data,enzyme\_cost\_weights,ecm\_options)**

#### DESCRIPTION

```
 ECM_SAVE_MODEL_AND_DATA_GAMS - Save input files for ECM by GAMS solvers
 
 ecm_save_model_and_data_gams(filename,network,v,r,c_data,u_data,enzyme_cost_weights,ecm_options)
 
 Convert data for Enzyme Cost Minimization (model and data) from SBtab format to GAMS input format
 
 For generating the input file (SBtab format), see 'help ecm_save_model_and_data_sbtab'
```

#### CROSS-REFERENCE INFORMATION

This function calls:


This function is called by:


---

Generated on Mon 30-Jan-2017 18:21:00 by **m2html** © 2003
