## Supplementary material for "In-depth computational analysis of natural and artificial carbon fixation pathways": MATLAB files: ecm_save_model_and_data_gams_OLD.html

Description of ecm\_save\_model\_and\_data\_gams\_OLD


Home > enzyme-cost-minimization > ecm\_functions > ecm\_save\_model\_and\_data\_gams\_OLD.m

### ecm\_save\_model\_and\_data\_gams\_OLD

#### PURPOSE

#### SYNOPSIS

**function ecm\_save\_model\_and\_data\_gams(filename,network,v,r,c\_data,u\_data,ecm\_options)**

#### DESCRIPTION

#### CROSS-REFERENCE INFORMATION

This function calls:


This function is called by:


---

Generated on Wed 30-Mar-2016 18:45:01 by **m2html** © 2003
