## Supplementary material for "In-depth computational analysis of natural and artificial carbon fixation pathways": MATLAB files: ecm_save_result_sbtab.html

Description of ecm\_save\_result\_sbtab


Home > enzyme-cost-minimization > ecm\_functions > ecm\_save\_result\_sbtab.m

### ecm\_save\_result\_sbtab

#### PURPOSE

**ECM\_SAVE\_RESULT\_SBTAB - Save ECM results in SBtab format**

#### SYNOPSIS

**function ecm\_save\_result\_sbtab(filename,network,c,u,A\_forward,options,c\_min,c\_max,u\_min,u\_max,u\_capacity,eta\_energetic,eta\_saturation)**

#### DESCRIPTION

```
ECM_SAVE_RESULT_SBTAB - Save ECM results in SBtab format
 
ecm_save_result_sbtab(filename,network,c,u,A_forward,options,c_min,c_max,u_min,u_max,u_capacity,eta_energetic,eta_saturation)
```

#### CROSS-REFERENCE INFORMATION

This function calls:


This function is called by:

- ecm\_simple ECM\_SIMPLE - Wrapper script for Parameter Balancing or Enzyme Cost Minimisation


---

Generated on Mon 30-Jan-2017 18:21:00 by **m2html** © 2003
