## Supplementary material for "In-depth computational analysis of natural and artificial carbon fixation pathways": MATLAB files: ecm_sbtab2mnt.html

Description of ecm\_sbtab2mnt


Home > enzyme-cost-minimization > ecm\_functions > ecm\_sbtab2mnt.m

### ecm\_sbtab2mnt

#### PURPOSE

**[network, v, conc\_fix, kinetic\_data] = ecm\_sbtab2mnt(model\_filename, model\_dir, matlab\_dir,kinetic\_data\_file\_names)**

#### SYNOPSIS

**function [network, v, conc\_min, conc\_max, kinetic\_data] = ecm\_sbtab2mnt(model\_filename, filenames,kinetic\_data\_file\_names,options,organism\_long,kegg\_conversion\_file,position\_file,reaction\_column\_name,compound\_column\_name,use\_kegg\_ids)**

#### DESCRIPTION

```
 [network, v, conc_fix, kinetic_data] = ecm_sbtab2mnt(model_filename, model_dir, matlab_dir,kinetic_data_file_names)

 Read network structure (from SBtab files), collect dG0' values (from network_thermo)
 and convert them into MNT network format
 the arguments kegg_conversion_file, position_file can be given 
 explicitly, or as fields of "filenames" (which can remain empty otherwise)
```

#### CROSS-REFERENCE INFORMATION

This function calls:

- ecm\_kegg\_compound\_id\_to\_name ECM\_KEGG\_COMPOUND\_ID\_TO\_NAME - Name conversion for compounds

This function is called by:


---

Generated on Mon 30-Jan-2017 18:21:00 by **m2html** © 2003
