## Supplementary material for "In-depth computational analysis of natural and artificial carbon fixation pathways": MATLAB files: ecm_setup.html

Description of ecm\_setup


Home > enzyme-cost-minimization > ecm\_functions > ecm\_setup.m

### ecm\_setup

#### PURPOSE

**ECM\_SETUP - Determine data directory**

#### SYNOPSIS

**function ecm\_info = ecm\_setup()**

#### DESCRIPTION

```
 ECM_SETUP - Determine data directory
```

#### CROSS-REFERENCE INFORMATION

This function calls:

- ecm\_BASEDIR ECM\_BASEDIR - Return ECM toolbox directory

This function is called by:

- ecm\_kegg\_compound\_id\_to\_name ECM\_KEGG\_COMPOUND\_ID\_TO\_NAME - Name conversion for compounds


---

Generated on Mon 30-Jan-2017 18:21:00 by **m2html** © 2003
