## Supplementary material for "In-depth computational analysis of natural and artificial carbon fixation pathways": MATLAB files: ecm_tolerances.html

Description of ecm\_tolerances


Home > enzyme-cost-minimization > ecm\_functions > ecm\_tolerances.m

### ecm\_tolerances

#### PURPOSE

**ECM\_TOLERANCES - Compute approximate tolerance ranges for metabolites and enzymes**

#### SYNOPSIS

**function [c\_min, c\_max, u\_min, u\_max] = ecm\_tolerances(c,u,v,ecm\_options,mca\_info)**

#### DESCRIPTION

```
ECM_TOLERANCES - Compute approximate tolerance ranges for metabolites and enzymes
 
[c_min, c_max, u_min, u_max] = ecm_tolerances(c,u,v,ecm_options,mca_info)
```

#### CROSS-REFERENCE INFORMATION

This function calls:


This function is called by:


---

Generated on Mon 30-Jan-2017 18:21:00 by **m2html** © 2003
