## Supplementary material for "In-depth computational analysis of natural and artificial carbon fixation pathways": MATLAB files: ecm_update_options.html

Description of ecm\_update\_options


Home > enzyme-cost-minimization > ecm\_functions > ecm\_update\_options.m

### ecm\_update\_options

#### PURPOSE

**ECM\_UPDATE\_OPTIONS - Helper function for ECM options (struct 'ecm\_options')**

#### SYNOPSIS

**function ecm\_options = ecm\_update\_options(network, ecm\_options);**

#### DESCRIPTION

```
 ECM_UPDATE_OPTIONS - Helper function for ECM options (struct 'ecm_options')

 ecm_options = ecm_update_options(network, ecm_options)

 Update ECM options for a given network:
  - adapt metabolite constraints
  - insert protein cost weights
  - adjust Keq values
```

#### CROSS-REFERENCE INFORMATION

This function calls:

- ecm\_default\_options ECM\_DEFAULT\_OPTIONS - Defaults for ECM

This function is called by:

- ecm\_simple ECM\_SIMPLE - Wrapper script for Parameter Balancing or Enzyme Cost Minimisation


---

Generated on Mon 30-Jan-2017 18:21:00 by **m2html** © 2003
