## Supplementary material for "In-depth computational analysis of natural and artificial carbon fixation pathways": MATLAB files: index.html

Index for Directory ./enzyme-cost-minimization/ecm\_functions


|  |  |
| --- | --- |
| Master index | Index for ./enzyme-cost-minimization/ecm\_functions |

### Index for ./enzyme-cost-minimization/ecm\_functions

#### Matlab files in this directory:

|  |  |
| --- | --- |
| Contents | Internal functions for Enzyme Cost Minimisation |
| ecm\_BASEDIR | ECM\_BASEDIR - Return ECM toolbox directory |
| ecm\_below\_threshold | ECM\_BELOW\_THRESHOLD - Helper function for enzyme cost minimization |
| ecm\_check\_parameter\_balancing | ECM\_CHECK\_PARAMETER\_BALANCING - Checks for balanced parameters |
| ecm\_default\_options | ECM\_DEFAULT\_OPTIONS - Defaults for ECM |
| ecm\_dependencies | ECM\_DEPENDENCIES - Check whether dependencies of ECM toolbox are satisfied |
| ecm\_display | ECM\_DISPLAY - Display results of ECM |
| ecm\_get\_one\_u | ECM\_GET\_ONE\_U - Helper function for enzyme variability calculation after enzyme cost minimization: |
| ecm\_get\_score | ECM\_GET\_SCORE - Helper function |
| ecm\_get\_specific\_rates | ECM\_GET\_SPECIFIC\_RATES - Helper function |
| ecm\_inequalities | MEASURES\_FOR\_ENZYME\_COSTS\_INEQUALITIES - Helper function |
| ecm\_kegg\_compound\_id\_to\_name | ECM\_KEGG\_COMPOUND\_ID\_TO\_NAME - Name conversion for compounds |
| ecm\_load\_model\_and\_data\_sbtab | ECM\_LOAD\_MODEL\_AND\_DATA\_SBTAB - Load data from SBtab file |
| ecm\_one\_run | ECM\_ONE\_RUN - Perform one ECM run |
| ecm\_read\_options | ECM\_READ\_OPTIONS - Read ECM options from file |
| ecm\_regularisation | ECM\_REGULARISATION - Regularisation term for ECM |
| ecm\_save\_model\_and\_data\_gams | ECM\_SAVE\_MODEL\_AND\_DATA\_GAMS - Save input files for ECM by GAMS solvers |
| ecm\_save\_model\_and\_data\_sbtab | ECM\_SAVE\_MODEL\_AND\_DATA\_SBTAB - Write SBtab file containing (model and data) information for Enzyme Cost Minimization |
| ecm\_save\_result\_sbtab | ECM\_SAVE\_RESULT\_SBTAB - Save ECM results in SBtab format |
| ecm\_sbtab2mnt | [network, v, conc\_fix, kinetic\_data] = ecm\_sbtab2mnt(model\_filename, model\_dir, matlab\_dir,kinetic\_data\_file\_names) |
| ecm\_setup | ECM\_SETUP - Determine data directory |
| ecm\_tolerances | ECM\_TOLERANCES - Compute approximate tolerance ranges for metabolites and enzymes |
| ecm\_update\_options | ECM\_UPDATE\_OPTIONS - Helper function for ECM options (struct 'ecm\_options') |

---

Generated on Mon 30-Jan-2017 18:21:00 by **m2html** © 2003
