## Supplementary material for "In-depth computational analysis of natural and artificial carbon fixation pathways": MATLAB files: ecm_parameter_balancing.html

Description of ecm\_parameter\_balancing


Home > enzyme-cost-minimization > ecm\_parameter\_balancing.m

### ecm\_parameter\_balancing

#### PURPOSE

**ECM\_PARAMETER\_BALANCING - Prepare and run parameter balancing**

#### SYNOPSIS

**function [r, r\_orig, kinetic\_data, r\_samples, ecm\_options, quantity\_info\_used, r\_std] = ecm\_parameter\_balancing(network, ecm\_options, kinetic\_data);**

#### DESCRIPTION

```
 ECM_PARAMETER_BALANCING - Prepare and run parameter balancing

 [r, r_orig, kinetic_data, ecm_options, quantity_info_used, r_std] = ecm_parameter_balancing(network, ecm_options, kinetic_data);

 Output
   r        Kinetic constants (posterior model values; used as input in parameter balancing)
   r_orig   Original kinetic constants (used as input in parameter balancing)
   r_std    Kinetic constants (posterior standard deviations)
 
 Uses (potentially) the following options from ecm_options.
  ecm_options.flag_given_kinetics
  ecm_options.reaction_column_name (only if no kinetic data are given)
  ecm_options.compound_column_name (only if no kinetic data are given)
  ecm_options.kcat_usage  {'use','none','forward'} (default: 'use')
  ecm_options.kcat_prior_median
  ecm_options.kcat_prior_log10_std
  ecm_options.kcat_lower
  ecm_options.kcatr_lower
  ecm_options.kcat_upper
  ecm_options.KM_lower
  ecm_options.Keq_upper
  ecm_options.quantity_info_file
  ecm_options.GFE_fixed
  ecm_options.use_pseudo_values
  ecm_options.fix_Keq_in_sampling
```

#### CROSS-REFERENCE INFORMATION

This function calls:

- ecm\_default\_options ECM\_DEFAULT\_OPTIONS - Defaults for ECM

This function is called by:

- ecm\_simple ECM\_SIMPLE - Wrapper script for Parameter Balancing or Enzyme Cost Minimisation


---

Generated on Mon 30-Jan-2017 18:21:00 by **m2html** © 2003
