## Supplementary material for "In-depth computational analysis of natural and artificial carbon fixation pathways": MATLAB files: ecm_emc2sp.html

Description of ecm\_emc2sp


Home > enzyme-cost-minimization > ecm\_scores > ecm\_emc2sp.m

### ecm\_emc2sp

#### PURPOSE

**[u\_tot, u] = ecm\_emc2sp(x,pp)**

#### SYNOPSIS

**function [u\_tot, u, w ] = ecm\_emc2sp(x,pp)**

#### DESCRIPTION

```
 [u_tot, u] = ecm_emc2sp(x,pp)
```

#### CROSS-REFERENCE INFORMATION

This function calls:


This function is called by:

- ecm\_get\_score ECM\_GET\_SCORE - Helper function
- ecm\_get\_specific\_rates ECM\_GET\_SPECIFIC\_RATES - Helper function


---

Generated on Mon 30-Jan-2017 18:21:00 by **m2html** © 2003
