## Supplementary material for "In-depth computational analysis of natural and artificial carbon fixation pathways": MATLAB files: ecm_simple.html

Description of ecm\_simple


Home > enzyme-cost-minimization > ecm\_simple.m

### ecm\_simple

#### PURPOSE

**ECM\_SIMPLE - Wrapper script for Parameter Balancing or Enzyme Cost Minimisation**

#### SYNOPSIS

**function [report, errors] = ecm\_simple(model\_data\_file, outdir, options)**

#### DESCRIPTION

```
 ECM_SIMPLE - Wrapper script for Parameter Balancing or Enzyme Cost Minimisation

 [report, errors] = ecm_simple(model_data_file, outdir, options)

 This function reads an input file [model_data_file] in SBtab format, performs either
 Parameter Balancing or ECM, and saves the results to an SBtab file in [outdir]

 Fields of struct 'options':
   options.actions      string {'ecm_standard','parameter_balancing'} - default: 'ecm_standard'
   options.make_report  flag   - show graphics (only used with 'ecm_standard') - default: 0

 Possible actions: (in options.actions) 

 'parameter_balancing': 
    run parameter_balancing using standard settings, 
    input file: prepared model with data ("ModelData")
 
 'ecm_standard':
    run an ECM using standard settings, 
    input file: prepared model with data ("ModelData")

 The function can also be called via the python script 'ecm.py'
```

#### CROSS-REFERENCE INFORMATION

This function calls:

- ecm\_enzyme\_cost\_minimization ECM\_ENZYME\_COST\_MINIMIZATION - Compute optimal flux-specific enzyme costs for given flux distribution
- ecm\_default\_options ECM\_DEFAULT\_OPTIONS - Defaults for ECM
- ecm\_display ECM\_DISPLAY - Display results of ECM
- ecm\_load\_model\_and\_data\_sbtab ECM\_LOAD\_MODEL\_AND\_DATA\_SBTAB - Load data from SBtab file
- ecm\_save\_result\_sbtab ECM\_SAVE\_RESULT\_SBTAB - Save ECM results in SBtab format
- ecm\_update\_options ECM\_UPDATE\_OPTIONS - Helper function for ECM options (struct 'ecm\_options')
- ecm\_parameter\_balancing ECM\_PARAMETER\_BALANCING - Prepare and run parameter balancing

This function is called by:

- demo\_escherichia\_coli\_ccm Demo script for Parameter Balancing and Enzyme Cost Minimization


---

Generated on Mon 30-Jan-2017 18:21:00 by **m2html** © 2003
