## Supplementary material for "In-depth computational analysis of natural and artificial carbon fixation pathways": MATLAB files: index.html

Index for Directory ./enzyme-cost-minimization


|  |  |
| --- | --- |
| Master index | Index for ./enzyme-cost-minimization |

### Index for ./enzyme-cost-minimization

#### Matlab files in this directory:

|  |  |
| --- | --- |
| Contents | Functions for Enzyme Cost Minimization |
| ecm\_enzyme\_cost\_minimization | ECM\_ENZYME\_COST\_MINIMIZATION - Compute optimal flux-specific enzyme costs for given flux distribution |
| ecm\_parameter\_balancing | ECM\_PARAMETER\_BALANCING - Prepare and run parameter balancing |
| ecm\_simple | ECM\_SIMPLE - Wrapper script for Parameter Balancing or Enzyme Cost Minimisation |

#### Subsequent directories:

- demo
- ecm\_functions
- ecm\_scores

---

Generated on Mon 30-Jan-2017 18:21:00 by **m2html** © 2003
