## Supplementary material for "In-depth computational analysis of natural and artificial carbon fixation pathways": MATLAB files: index.html

Matlab Index


### Matlab Index

#### Matlab Directories

- ./enzyme-cost-minimization
- ./enzyme-cost-minimization/demo
- ./enzyme-cost-minimization/ecm\_functions
- ./enzyme-cost-minimization/ecm\_scores

#### Matlab Files found in these Directories

|  |  |  |  |
| --- | --- | --- | --- |
| Contents | ecm\_emc1 | ecm\_get\_score | ecm\_regularisation |
| Contents | ecm\_emc2s | ecm\_get\_specific\_rates | ecm\_save\_model\_and\_data\_gams |
| Contents | ecm\_emc2sp | ecm\_inequalities | ecm\_save\_model\_and\_data\_sbtab |
| Contents | ecm\_emc3s | ecm\_kegg\_compound\_id\_to\_name | ecm\_save\_result\_sbtab |
| demo\_escherichia\_coli\_ccm | ecm\_emc3sp | ecm\_load\_model\_and\_data\_sbtab | ecm\_sbtab2mnt |
| ecm\_BASEDIR | ecm\_emc4cm | ecm\_mdf | ecm\_setup |
| ecm\_below\_threshold | ecm\_emc4dm | ecm\_memc2s | ecm\_simple |
| ecm\_check\_parameter\_balancing | ecm\_emc4geom | ecm\_memc2sp | ecm\_tolerances |
| ecm\_default\_options | ecm\_emc4sm | ecm\_one\_run | ecm\_update\_options |
| ecm\_dependencies | ecm\_enzyme\_cost\_minimization | ecm\_parameter\_balancing |  |
| ecm\_display | ecm\_get\_one\_u | ecm\_read\_options |  |

---

Generated on Mon 30-Jan-2017 18:21:00 by **m2html** © 2003
