## Supplementary material for "In-depth computational analysis of natural and artificial carbon fixation pathways": MATLAB files: Contents.m

```
% --------------------------
%  Enzyme Cost Minimization
% --------------------------
%
% Demo scripts
%
%   demo_ecm_simple
%   demo_ecm_ecoli_noor_2016
%
% Main functions 
%
%   ecm_paramater_balancing      - Run parameter balancing with standard settings
%   ecm_enzyme_cost_minimization - Run enzyme cost minimization
%   ecm_simple                   - Wrapper function for parameter balancing or enzyme cost minimization
% 
% For a list of options for customising the ECM algorithm, see 'help ecm_default_options'
%
% For a list of possible cost scores (minimisation of enzyme and metabolites levels, as well as fitting of 
% enzyme and metabolite data), see 'help ecm_get_score'
%
% Matlab Toolboxes required
%
%   Metabolic Network Toolbox    - Metabolic networks    (https://github.com/wolframliebermeister/mnt)
%   SBMLtoolbox                  - SBML import / export  (http://sbml.org/Software/SBMLToolbox)
%   SBtab toolbox                - SBtab format          (https://github.com/wolframliebermeister/sbtab-matlab)
%   efmtool                      - Elementary flux modes (http://www.csb.ethz.ch/tools/efmtool)
%
% (C) 2019 Wolfram Liebermeister  <>
```
