## Supplementary material for "In-depth computational analysis of natural and artificial carbon fixation pathways": MATLAB files: demo_ecm_e_coli_noor_2016.m

```
% ---------------------------------------------------------------------------------------
% Enzyme cost minimization demo script
% ---------------------------------------------------------------------------------------


% ---------------------------------------------------------------------------------------
% Set location of your ECM Model file; you can choose a different location.
% (by default, matlab will search the files on the matlab search path)

data_dir                 = [ecm_RESOURCEDIR filesep 'models' filesep 'e_coli_noor_2016'];
filename_model           = [data_dir filesep 'e_coli_noor_2016_ECM_Model.tsv'];
filename_validation_data = [data_dir filesep 'e_coli_noor_2016_ECM_ValidationData.tsv'];


% ---------------------------------------------------------------------------------------
% Load model and data from the ECM Model file and translate them into matlab data structures 
% (see documentation of the Metabolic Network Toolbox for details)

display(sprintf('Reading model and data file %s', filename_model));
display(sprintf('Reading validation data file %s', filename_validation_data));

[network,v,c_data,u_data, conc_min, conc_max, met_fix, conc_fix,positions, enzyme_cost_weights, warnings] = load_model_and_data_sbtab(filename_model, filename_validation_data);


% ---------------------------------------------------------------------------------------
% Define default options for ECM; to change the options, refer to the documentation

ecm_options            = ecm_default_options(network, 'E. coli central carbon metabolism');
ecm_options.c_data     = c_data;
ecm_options.u_data     = u_data;
ecm_options.ecm_scores = {'emc4cm'};
ecm_options            = ecm_update_options(network, ecm_options);


% ---------------------------------------------------------------------------------------
% Run ECM

[c, u, u_cost, up, A_forward, mca_info, c_min, c_max, u_min, u_max, r, u_capacity, eta_energetic, eta_saturation] = ecm_enzyme_cost_minimization(network, network.kinetics, v, ecm_options);


% ---------------------------------------------------------------------------------------
% Save results to SBtab files (and options in JSON file) (here: in your tmp directory)

document_name = 'E. coli central carbon metabolism - ECM result';

outfile_name          = [tempdir 'demo_ecm_e_coli_noor_2016'];
outfile_options_json  = [tempdir 'demo_ecm_e_coli_noor_2016_options.json'];
outfile_options_sbtab = [tempdir 'demo_ecm_e_coli_noor_2016_options.tsv' ];

options = struct('r', network.kinetics, 'method', 'emc4cm', 'document_name', document_name, 'save_tolerance_ranges', 1);

ecm_save_result_sbtab(outfile_name, network, c, u, A_forward, options, c_min, c_max, u_min, u_max, u_capacity, eta_energetic, eta_saturation, v);

% Write ECM options to JSON file
% ecm_options_save(outfile_options_json,ecm_options);

% Write ECM options to SBtab file
options_to_sbtab(ecm_options,struct('filename',outfile_options_sbtab,'TableName','Options for ECM','TableID','OptionsECM','Method','enzyme-cost-minimization'));


% ---------------------------------------------------------------------------------------
% Display results as graphics

kinetic_data                           = [];
graphics_options.print_graphics        = 1;
graphics_options.few_graphics          = 1;
graphics_options.metabolite_order_file = [];
graphics_options.reaction_order_file   = [];
graphics_options.enzyme_colors         = sunrise_colors(length(ecm_options.ind_scored_enzymes));

ecm_display(ecm_options, graphics_options, network,v,c,u,u_cost,up,A_forward,r,kinetic_data,c_min,c_max,u_min,u_max,u_capacity,eta_energetic,eta_saturation);
```
