## Supplementary material for "In-depth computational analysis of natural and artificial carbon fixation pathways": MATLAB files: demo_ecm_simple.m

```
% --------------------------------------------------------------------------------------------------
% Run ECM workflow on E. coli model Noor et al 2016
% (standard version of ECM, called through the wrapper function ecm_simple) 
% 
% The MATLAB function 'ecm_simple' is a wrapper function that reads the data, 
% sets default parameters, and calls functions for Parameter Balancing or Enzyme Cost Minimization.

% Use the prepared model and data files 
%   enzyme-cost-minimization/matlab/enzyme-cost-minimization/demo/data/ecoli_noor_2016_ECM_Model.tsv
%   enzyme-cost-minimization/matlab/enzyme-cost-minimization/demo/data/ecoli_noor_2016_ECM_ValidationData.tsv

% Results are written to subdirectory "enzyme-cost-minimization/matlab/enzyme-cost-minimization/demo/results"
%
% To see the workflow in more detail, have a look at demo_ecm_ecoli_noor_2016.m
% --------------------------------------------------------------------------------------------------

display(sprintf('\n---------------------------------------------------------------------------'));
display(sprintf('Enzyme cost minimization demo'));
display(sprintf('---------------------------------------------------------------------------\n'));
display(sprintf(' This script calls the wrapper function "ecm_simple".'));
display(sprintf(' It reads a prepared model and data file and determines balanced model parameters.'));
display(sprintf(' Then it performs ECM for a number of enzyme cost scores.'))
display(sprintf(' Graphics are generated, and the results are saved to files.\n'))


% --------------------------------------------------------
% Set some filenames for this demo script
% --------------------------------------------------------

filenames = filenames_demo_ecm_simple;


% --------------------------------------------------------
% Run Parameter Balancing with standard settings
% --------------------------------------------------------

% Note that ecm_simple has its owon little options data structure 
% (different from "ecm_options" used generally in this toolbox

% set verbose = 1 to see more output

options = struct('actions', 'parameter balancing', 'verbose', 0);

ecm_simple(filenames.model_data_file, filenames.validation_data_file, filenames.result_dir, options);


% --------------------------------------------------------
% Run Enzyme Cost Minimization with standard settings
% --------------------------------------------------------

% set options.verbose = 1 to see more output

options                            = struct('actions', 'ecm', 'generate_report', 1, 'verbose', 0);
options.replace_cofactors          = {'ATP','ADP','Orthophosphate','NADH', 'NAD+', 'NADPH','NADP+','Ubiquinone', 'Ubiquinol'};
options.fluctuations_safety_margin = 3;           % safety margin (# std dev) to counter protein number fluctuations
options.cell_volume                = 1.1*10^-18;  % in m^3, default value for E coli (needed for safety margin) 
options.verbose                    = 0;

ecm_simple(filenames.model_data_file, filenames.validation_data_file, filenames.result_dir, options);
```
