## Supplementary material for "In-depth computational analysis of natural and artificial carbon fixation pathways": MATLAB files: filenames_demo_ecm_simple.m

```
function filenames = filenames_demo_ecm_simple()
  
filenames.data_dir             = [ecm_BASEDIR filesep 'demo' filesep 'data' filesep];
filenames.model_data_file      = [filenames.data_dir 'ecoli_noor_2016_ECM_Model.tsv'];
filenames.validation_data_file = [filenames.data_dir 'ecoli_noor_2016_ECM_ValidationData.tsv'];
filenames.result_dir           = [tempdir 'demo_ecm_simple'];

[~,~] = mkdir(filenames.result_dir);
```
