## Supplementary material for "In-depth computational analysis of natural and artificial carbon fixation pathways": MATLAB files: ecm_below_threshold.m

```
% ECM_BELOW_THRESHOLD - Helper function for enzyme cost minimization
% 
% function [ineq_constraints, eq_constraints] = ecm_below_threshold(ecm_score,xx,pp,u_threshold,x_min,x_max,lambda_regularisation)

function [ineq_constraints, eq_constraints] = ecm_below_threshold(ecm_score,xx,pp,u_threshold,x_min,x_max,lambda_regularisation)


a1 = ecm_get_score(ecm_score,xx,pp) + ecm_regularisation(xx,x_min,x_max,lambda_regularisation) - u_threshold;
 
a2 = ecm_inequalities(xx,pp.N_forward,pp.log_Keq_forward);

ineq_constraints = [a1; a2];
eq_constraints   = [];
```
