## Supplementary material for "In-depth computational analysis of natural and artificial carbon fixation pathways": MATLAB files: ecm_default_options.m

```
function ecm_options = ecm_default_options(network, model_name)

% ECM_DEFAULT_OPTIONS -  Default options for ECM algorithm
% 
% ecm_options = ecm_default_options(network, model_name)
%
% ECM options and their default values
%
% General options and model name
% ------------------------------
%   ecm_options.verbose                (Boolean)
%   ecm_options.model_name             (string) model_name
%   ecm_options.model_id               (string) model id (for file names)
%   ecm_options.run_id                 (string) simulation run id (for file names)
% 
% Metabolite constraints
% ----------------------
%   ecm_options.conc_min               (float vector)      minimal metabolite concentrations (mM), with nans 
%   ecm_options.conc_max               (float vector)      maximal metabolite concentrations (mM), with nans 
%   ecm_options.conc_min_default       (float)             default minimal metabolite concentration (mM), default 0.01
%   ecm_options.conc_max_default       (float)             default maximal metabolite concentration (mM), default 10
%   ecm_options.fix_metabolites        (Boolean vector)    metabolites with fixed concentrations (overrides values from input model)
%   ecm_options.fix_metabolite_values  (vector)            fixed metabolite concentrations (see ecm_options.fix_metabolites)
%   ecm_options.met_fix                                    metabolites with fixed concentrations
%   ecm_options.conc_fix               (float vector)      fixed metabolite concentrations (mM) (see ecm_options.met_fix)
%   ecm_options.replace_cofactors      (string cell array) names of cofactors; this is only used in ecm_update_options.m - 
%                                                          bounds for cofactor concentrations are automatically set based on 
%                                                          median value from concentration data
% Kinetic data
% ------------
%   ecm_options.kinetic_data                     matlab struct with kinetic data; see data structures in mnt toolbox
%   ecm_options.reaction_column_names            reaction column names (in data file) for loading of kinetic data
%   ecm_options.compound_column_names            compound column names (in data file) for loading of kinetic data
%   ecm_options.KM_lower               (float)   lower bound on KM values (mM)
%   ecm_options.Keq_upper              (float)   uppe bound on Keq values
%   ecm_options.flag_given_kinetics    (Boolean) use given kinetics
%   ecm_options.kcat_usage             (string)  use kcat data values (default: 'use')
%   ecm_options.kcat_upper             (float)   upper bound on Kcat values (default 10000, 1/s)
%   ecm_options.kcat_lower             (float)   lower bound on Kcat values (default 0.1, 1/s)
%   ecm_options.kcat_lower             (float)   lower bound on reverse Kcat values (default 0.0001, 1/s)
%   ecm_options.kcat_prior_median      (float)   median for Kcat values (default 10, 1/s)
%   ecm_options.kcat_prior_log10_std   (float)   std deviation of log10 Kcat values (default 0.2, 1/s)
% 
% Parameter balancing
% -------------------
%   ecm_options.use_pseudo_values      (Boolean) use pseudo values during parameter balancing
%   ecm_options.GFE_fixed              (Boolean) keep GFE values fixed during parameter balancing (default true)
%   ecm_options.insert_Keq_from_data   (Boolean) insert again Keq values from data after parameter balancing (default false)
%   ecm_options.n_samples              (integer) number of samples in posterior sampling
% 
% Enzyme cost weights
% -------------------
%   ecm_options.ind_scored_enzymes     (integer vector) indices of reactions with enzymes to be scored in cost function  (default [1:length(network.actions)]')
%   ecm_options.enzyme_cost_weights    (float vector)   numerical enzyme cost weights (default ones(length(ecm_options.ind_scored_enzymes),1))
%   ecm_options.use_cost_weights       (string)         usage of cost weights (default 'none')
%   ecm_options.parameter_prior_file   (string)         parameter prior file filename, optional (default [])
% 
% ECM calculation
% ---------------
%   ecm_options.ignore_regulation           (Boolean)      disregard enzyme regulation; default = 0
%   ecm_options.initial_choice              (string)       method for computing inital solution {'mdf' 'polytope_center', 'interval_center',
%                                                          'given_x_start'}, default: 'mdf'
%   ecm_options.x_start                     (float vector) initial solution vector, only for ecm_options.initial_choice='given_x_start'
%   ecm_options.multiple_conditions_n       (int)          number of experimental conditions to be considered
%   ecm_options.multiple_conditions_anticipate  (boolean)      optimise a single enzyme profile for multiple conditions? default false
%   ecm_options.multiple_starting_points    (boolean)      default false
%   ecm_options.ecm_scores                  (string cell array) list of ecm scores to be used (default 'emc3sp'); see 'help ecm_scores'
%   ecm_options.lambda_regularisation       (float)        coefficient for metabolite concentration regularisation term (default 10^-3)
%   ecm_options.lambda_reg_factor           (float)        regularisation factor for metabolite concentration regularisation term (default 10^-3)
%                                                          (parameter for choosing ecm_options.lambda_regularisation automatically, in ecm_update_options; set to [] in order to not use it)
%   ecm_options.use_linear_cost_constraints (Boolean)      tighter constraints derived upper bound on enzyme cost 
%   ecm_options.fluctuations_safety_margin  (float)        safety margin (# std dev) to counter protein number fluctuations, default 0
%   ecm_options.cell_volume                 (float)        default value for E coli (needed for safety margin), in m^3, default 1.1*10^-18
%   ecm_options.maximal_u_cost              (float)        used with option "use_linear_cost_constraints", default []
%   ecm_options.compute_hessian             (Boolean)      default false
%   ecm_options.compute_elasticities        (Boolean)      default false
%   ecm_options.compute_tolerance           (Boolean)      default false
%   ecm_options.cost_tolerance_factor       (float)        factor defining the cost tolerance, default 1.01
%   ecm_options.tolerance_from_hessian      (Boolean)      default false
%   ecm_options.fix_Keq_in_sampling         (Boolean)      default false
%   ecm_options.fix_thermodynamics_by_adjusting_Keq  (Boolean) enforce thermodynamically feasibile fluxes by adjusting Keq
% 
% Metabolic data (for comparison)
% -------------------------------
%   ecm_options.c_data  (float vector) metabolite concentration data, with nans
%   ecm_options.u_data  (float vector) enzyme concentration data, with nans
% 
% Graphics
% --------
%   ecm_options.print_graphics         (Boolean)           default false
%   ecm_options.show_graphics          (Boolean)           default true
%   ecm_options.show_metabolites       (string cell array) default network.metabolites;
%   ecm_options.metabolite_order_file  (string)            filename of file containing metabolite order for graphics, default []
%   ecm_options.reaction_order_file    (string)            filename of file containing reaction order for graphics, default []

% Argument 'model_name' is optional

  eval(default('network','[]','model_name','''model'''));

if isempty(network),
  network = network_construct;%(N,reversible,ind_external,metabolites,actions,no_graphics,regulation_matrix,flag_kinetics)
end

[nm,nr] = size(network.N);

ecm_options = struct;

% model
ecm_options.model_name               = model_name         ; 
ecm_options.run_id                   = 'RUN';
ecm_options.model_id                 = 'MODEL';

% given data
ecm_options.c_data                   = [];
ecm_options.u_data                   = [];
ecm_options.kinetic_data             = [];

% add parameter balancing options
ecm_options = join_struct(ecm_options, parameter_balancing_options);
ecm_options.kcat_prior_median        = 10;     % for ccm this value need to be modified
ecm_options.kcat_prior_log10_std     = 0.2;   % reduce spread of kcat values
ecm_options.kcat_upper               = 10000;  % 1/s
ecm_options.kcat_lower               = 0.1;    % 1/s
ecm_options.kcatr_lower              = 0.0001; % 1/s
ecm_options.GFE_fixed                = 1;     % flag

% metabolite constraints
ecm_options.fix_metabolites          = {}; % metabolites with fixed concentrations (overrides values from input model)
ecm_options.fix_metabolite_values    = [];
ecm_options.conc_min_default         = 0.001; % mM
ecm_options.conc_max_default         = 10;  % mM
ecm_options.conc_min                 = [];
ecm_options.conc_max                 = [];
ecm_options.conc_fix                 = [];
ecm_options.ind_met_fix              = [];
ecm_options.met_fix                  = [];
ecm_options.replace_cofactors        = {};

% kinetic data
ecm_options.reaction_column_names    = []; % column names (in data file) for loading of kinetic data
ecm_options.compound_column_names    = [];
ecm_options.flag_given_kinetics      = 0;
ecm_options.insert_Keq_from_data     = 0;     % flag

% enzyme cost weights
ecm_options.ind_scored_enzymes       = [1:length(network.actions)]';
ecm_options.enzyme_cost_weights      = ones(length(ecm_options.ind_scored_enzymes),1);
ecm_options.use_cost_weights         = 'none';

% ecm
ecm_options.ignore_regulation           = 0;
ecm_options.initial_choice              = 'polytope_center';
ecm_options.x_start                     = []; 
ecm_options.fix_thermodynamics_by_adjusting_Keq = 1;
ecm_options.multiple_conditions_anticipate = 0;
ecm_options.multiple_conditions_n       = 1;
ecm_options.multiple_starting_points    = 0;
ecm_options.ecm_scores                  = {'emc3sp'}           ;
ecm_options.lambda_regularisation       = 10^-3; 
ecm_options.lambda_reg_factor           = 0.01;
ecm_options.parameter_prior_file        = [];
ecm_options.use_linear_cost_constraints = 1; 
ecm_options.fluctuations_safety_margin  = 0; % safety margin, to counter protein number fluctuations
ecm_options.cell_volume                 = 1.1*10^-18;  % in m^3, default value for E coli (needed for safety margin) 
ecm_options.maximal_u_cost              = []; 
ecm_options.compute_hessian             = 0;
ecm_options.compute_elasticities        = 0;
ecm_options.compute_tolerance           = 0;
ecm_options.cost_tolerance_factor       = 1.01; % one percent
ecm_options.tolerance_from_hessian      = 0;
ecm_options.fix_Keq_in_sampling         = 0;

% graphics
ecm_options.print_graphics           = 0;
ecm_options.show_graphics            = 1;
ecm_options.show_metabolites         = network.metabolites;
ecm_options.metabolite_order_file    = [];
ecm_options.reaction_order_file      = [];

% general
ecm_options.verbose                  = 0;
```
