## Supplementary material for "In-depth computational analysis of natural and artificial carbon fixation pathways": MATLAB files: ecm_dependencies.m

```
% ECM_DEPENDENCIES - Check whether dependencies of ECM toolbox are satisfied

function ecm_dependencies()

if ~exist('mnt_version','file'),
  error('Please install the Metabolic Network Toolbox (https://github.com/wolframliebermeister/mnt)');
end

if ~exist('TranslateSBML','file'),
  warning('Please install the SBML Toolbox (http://sbml.org/Software/SBMLToolbox) - Otherwise SBML import and export will not work');
end
```
