## Supplementary material for "In-depth computational analysis of natural and artificial carbon fixation pathways": MATLAB files: ecm_display.m

```
function ecm_display(ecm_options,graphics_options,network,v,c,u,u_tot,up,A_forward,r,kinetic_data,c_min,c_max,u_min,u_max,u_capacity,eta_energetic,eta_saturation,network_CoHid)

% ECM_DISPLAY - Display results of ECM
%
% ecm_display(network,options,ecm_options,v,c,u,u_tot,up,A_forward,r,kinetic_data,c_min,c_max,u_min,u_max)
%
% ECM_DISPLAY - Display results of Enzyme Cost Minimization
%
% Options in 'graphics_options':
%
%  graphics_options.few_graphics                    Omit many graphics
%  graphics_options.show_proteomaps                 Show proteomaps
%  graphics_options.show_original_data              Show graphs with original data
%  graphics_options.show_network_graphics           Show network graphs
%  graphics_options.show_matrix_graphics            Show matrix graphs
%  graphics_options.reaction_order_file             Filename for reordering reactions
%  graphics_options.metabolites_order_file          Filename for reordering compounds

if ~exist('network_CoHid','var'),
  network_CoHid = network;
end
  
%----------------------------------------------------------------------------------
% Options

graphics_options_default.few_graphics       = 0;
graphics_options_default.show_proteomaps    = 1;
graphics_options_default.show_original_data = 1;
graphics_options_default.show_network_graphics = 1;
graphics_options_default.show_matrix_graphics = 1;
graphics_options_default.show_metabolite_graphics = 1;

graphics_options = join_struct(graphics_options_default,graphics_options);

if graphics_options.few_graphics,
  graphics_options.show_proteomaps       = 0;
  graphics_options.show_original_data    = 0;
  graphics_options.show_network_graphics = 0;
  graphics_options.show_matrix_graphics  = 0;
  graphics_options.show_metabolite_graphics = 0;
end

%----------------------------------------------------------------------------------

if isfield(graphics_options,'reaction_order_file'),
  if length(graphics_options.reaction_order_file),
    ecm_options.reaction_order   = load_any_table(graphics_options.reaction_order_file);
    ecm_options.metabolite_order = load_any_table(graphics_options.metabolite_order_file);
  end
end

% -----------------------------------
% info

if ecm_options.verbose,
if length(ecm_options.met_fix),
  display('Fixed concentrations')
  pm(ecm_options.conc_fix,ecm_options.met_fix)
end
end

% -----------------------------------

[nm,nr] = size(network.N);

value_colors  = flipud(colormapRGBmatrices); % rb_colors;

%cm            = jet(length(ecm_options.ecm_scores));
ca

if ~isfield(graphics_options,'psfile_dir'),
  graphics_options.print_graphics = 0;
  else
    if isempty(graphics_options.psfile_dir),
      graphics_options.print_graphics = 0;
    end
end

if isempty(c.data),
  c.data = nan * ones(nm,1);
end

if isempty(u.data),
  u.data = nan * ones(nr,1);
end


% -----------------------------------

reaction_names = network.actions;

if isfield(network,'genes'),
  gene_names     = network.genes;
else
  gene_names     = network.actions;
end
  
if ~isfield(ecm_options,'reaction_order'),
  ecm_options.reaction_order = reaction_names(ecm_options.ind_scored_enzymes);
else,
  if isempty(ecm_options.reaction_order),
    ecm_options.reaction_order = {};
  end
  ll = label_names(ecm_options.reaction_order,reaction_names);
  ecm_options.reaction_order = ecm_options.reaction_order(find(ll));
  ecm_options.reaction_order = [column(ecm_options.reaction_order); column(setdiff(reaction_names,ecm_options.reaction_order))];
end

ecm_options.reaction_order = ecm_options.reaction_order(find(label_names(ecm_options.reaction_order,reaction_names(ecm_options.ind_scored_enzymes))));

ind_enzymes           = label_names(reaction_names,ecm_options.reaction_order);
ind_enzymes_rev       = label_names(ecm_options.reaction_order,reaction_names(ecm_options.ind_scored_enzymes));
ind_enzymes_scored    = ecm_options.ind_scored_enzymes(ind_enzymes_rev(find(ind_enzymes_rev)));

% enzyme colors (according to reactions in network model)
my_colors  = ones(nr,3);
my_colors(ecm_options.ind_scored_enzymes,:) = graphics_options.enzyme_colors(ind_enzymes_rev,:);


% -----------------------------------

if ~isfield(ecm_options,'metabolite_order'),
  ecm_options.metabolite_order = network.metabolites;
else,
  ecm_options.metabolite_order = [column(ecm_options.metabolite_order); column(setdiff(network.metabolites,ecm_options.metabolite_order))];
end

ind_show_met = [];
for it = 1:length(ecm_options.metabolite_order),
  my_metabolite = intersect(ecm_options.metabolite_order{it},ecm_options.show_metabolites);
  if length(my_metabolite),
    ind_show_met = [ ind_show_met, label_names(my_metabolite, network.metabolites)];
  end
end
ind_show_met = ind_show_met(find(ind_show_met));


% -----------------------------------
% GENERAL PLOTS 
% -----------------------------------

if length(kinetic_data),
  kcat_forward_original      = kinetic_data.Kcatf.median;
  kcat_forward_original(v<0) = kinetic_data.Kcatr.median(v<0);
else
  kcat_forward_original      = nan * ones(size(network.actions));
end

kcat_forward      = r.Kcatf;
kcat_forward(v<0) = r.Kcatr(v<0);

gp = struct('arrowsize',0.05,'arrowstyle','fluxes','actstyle','none','arrowvalues',v);

if graphics_options.show_network_graphics,

  figure(27); netgraph_concentrations(network_CoHid,[],v,1,gp);

 figure(4); clf; 
 gp = struct('arrowsize',0.05,'actstyle','none','colorbar',1,'showsign',0,'actprintnames',1,'metprintnames',0,'flag_edges',1,'colormap', value_colors);
 gp.actvaluesmin = log10(10);
 gp.actvaluesmax = log10(10000);
 gp.actstyle     = 'fixed';
 gp.arrowstyle   = 'none';
 netgraph_concentrations(network_CoHid,[],log10(kcat_forward),1,gp); 
 title('log10 Kcat+, balanced (1/s)');
 
 if graphics_options.show_original_data,
 figure(14); clf;
 if length(kinetic_data),
   gp = struct('arrowsize',0.05,'actstyle','none','colorbar',1,'showsign',0,'actprintnames',1,'metprintnames',0,'flag_edges',1,'colormap', value_colors);
   gp.actvaluesmin = log10(10);
   gp.actvaluesmax = log10(10000);
   gp.actstyle     = 'fixed';
   gp.arrowstyle   = 'none';
   netgraph_concentrations(network_CoHid,[],log10(kcat_forward_original),1,gp); 
   title('log10 Kcat+, original (1/s)');
 else
   text(1,1,'No original Kcat data available');
 end
 end

 figure(11); clf; 
 gp.actstyle     = 'none';
 gp.metvaluesmin = log10(ecm_options.conc_min_default);
 gp.metvaluesmax = log10(100);% (ecm_options.conc_max_default);
 netgraph_concentrations(network,log10(c.initial(:,1)),[],1,gp); title('Initial state: log10 Concentrations (mM)');
 
 figure(12); clf; 
 gp.arrowstyle   ='none'; gp.actstyle = 'fixed';
 gp.actvaluesmin = log10(1);
 gp.actvaluesmax = log10(100);
 gp.metvaluesmin = [];
 gp.metvaluesmax = [];
 netgraph_concentrations(network_CoHid,[],log10(A_forward.initial),1,gp); title('Inital state: log10 Reaction affinities (kJ/mol)');
 
 figure(8); clf
 gp = struct('arrowstyle','none','colorbar',1,'showsign',0,'actprintnames',0,'flag_edges',1,'colormap', value_colors);
 gp.metvaluesmin = log10(ecm_options.conc_min_default);
 gp.metvaluesmax = log10(100); % ecm_options.conc_max_default);
 netgraph_concentrations(network,log10(c.data(:,1)),[],1,gp); title('Experimental data (1st file): log10 Concentrations (mM)');
 
 figure(28); clf
 gp = struct('arrowstyle','none','colorbar',1,'showsign',0,'actprintnames',0,'flag_edges',1,'colormap', value_colors);
 gp.metvaluesmin = log10(ecm_options.conc_min_default);
 gp.metvaluesmax = log10(100); % ecm_options.conc_max_default);
 netgraph_concentrations(network,log10(c.fixed),[],1,gp); title('Fixed values: log10 Concentrations (mM)');
 
 figure(9); clf
 gp = struct('arrowstyle','none','colorbar',1,'showsign',0,'actprintnames',1,'flag_edges',1,'colormap', value_colors);
 gp.actnames     = gene_names;
 gp.actvaluesmin = log10(nanmin(u.data));
 gp.actvaluesmax = log10(nanmax(u.data));
 gp.metvaluesmin = [];
 gp.metvaluesmax = [];
 netgraph_concentrations(network_CoHid,[],log10(u.data),1,gp); title('Experimental data: log10 enzyme levels (mM)');

end

% --------------------------------------
% concentration predictions

% everything in one picture

if graphics_options.show_metabolite_graphics,

figure(18); clf; hold on;
set(gcf,'Position',[100 300 1000 500])
xtick = 0:length(ind_show_met)-1;
h     = [];
my_ecm_scores = [{'initial'},ecm_options.ecm_scores];
for it = 1:length(my_ecm_scores),
  ecm_score = my_ecm_scores{it};
  h(it)     = plot(xtick,c.(ecm_score)(ind_show_met,1) * exp(0.03*it),'-','MarkerSize',20); %  * exp(0.001*it)
end
line_colors(h,'sunrise_colors');

plot(xtick,c.fixed(ind_show_met),'r.','MarkerSize',30);
if length(c.data),
  n_c_data_sets = size(c.data,2);
  plot(repmat(column(xtick),n_c_data_sets,1),reshape(c.data(ind_show_met,:),n_c_data_sets * length(ind_show_met),1),'k.','MarkerSize',20); 
  methods = [my_ecm_scores,{'fixed','data'}];
else
  methods = [my_ecm_scores,{'fixed'}];
end

metnames_show = strrep(network.metabolites(ind_show_met),'D-','');
metnames_show = strrep(metnames_show,'_','.');

text(xtick + 0.2,c.data(ind_show_met),metnames_show,'FontSize',8);
legend(methods,'Location','NorthEastOutside','FontSize',10);
axis tight; set(gca,'XTick',xtick,'XTickLabel',metnames_show,'YScale','Log');
my_xticklabel([],ecm_options.conc_min_default,[],6);
title('Comparison: log10 Concentration profiles (mM)');

end

% --------------------------------------
% reaction affinity predictions

if graphics_options.show_metabolite_graphics,

figure(19); clf; hold on;
set(gcf,'Position',[200 400 1000 500])
xtick = 0:length(network.actions);
h = [];

my_ecm_scores = [{'initial'},ecm_options.ecm_scores];

for it = 1:length(my_ecm_scores),
  ecm_score = my_ecm_scores{it};
  h(it)     = plot(xtick,[0; -cumsum(v .* A_forward.(ecm_score)(:,1))] + (0.002*it),'-'); %  
end
line_colors(h,'sunrise_colors');

legend(my_ecm_scores,'Location','NorthEast','FontSize',12);
axis tight; set(gca,'XTick',xtick,'XTickLabel',[{''};strrep(network.actions,'_', ' ')]);
my_xticklabel([],[],[],10);
title('Cumulative entropy production (v * A)');
end

% --------------------------------------
% c over KM

if graphics_options.show_matrix_graphics,
figure(20); subplot('Position',[0.15 0.05 0.8 0.7]);
c_over_KM = repmat(c.initial(:,1)',length(network.actions),1)./r.KM;
c_over_KM(r.KM==0) = nan;
im(log10(c_over_KM(:,ind_show_met)),[-3,3],network.actions,metnames_show); colormap([0.95 0.95 0.95; value_colors]); colorbar
my_xticklabel
%title('log10 c (initial solution) / KM');
end

% --------------------------------------
% KM

% data

if graphics_options.show_original_data,
figure(21); subplot('Position',[0.15 0.05 0.8 0.7]);
if length(kinetic_data),
  my_KM = kinetic_data.KM.median;
  my_KM(my_KM==0) = nan;
  im(log10(my_KM(:,ind_show_met)),[-3,2],network.actions,metnames_show); colormap([0.95 0.95 0.95; value_colors]); colorbar
  my_xticklabel
else
  text(1,1,'No original KM data available');
end
end

if graphics_options.show_matrix_graphics,
% balanced
figure(22); subplot('Position',[0.15 0.05 0.8 0.7]);
my_KM = r.KM;
my_KM(my_KM==0) = nan;
im(log10(my_KM(:,ind_show_met)),[-3,2],network.actions,metnames_show); colormap([0.95 0.95 0.95; value_colors]); colorbar
my_xticklabel
end

% --------------------------------------
% enzyme level predictions

% barplot for all methods

if graphics_options.few_graphics ==0, 

  figure(33); clf; 
  ug      = [];
  methods = ecm_options.ecm_scores;
  
  for it = 1:length(ecm_options.ecm_scores),
    ecm_score = ecm_options.ecm_scores{it};
    %if ~strcmp(ecm_score,'mdf'),
      ug_meth   = up.(ecm_score)(ecm_options.ind_scored_enzymes,1);
      ug_meth   = ug_meth/sum(ug_meth);
      ug        = [ug, ug_meth];
    %end
  end
  
  if length(u.data),
    ug_data = u.data(ecm_options.ind_scored_enzymes) / nansum(u.data(ecm_options.ind_scored_enzymes));
    ug      = [ug, ug_data];
    methods = [methods,{'data'}];
  end
  
  h = bar(flipud(ug)','stacked'); 
  my_colormap = flipud(graphics_options.enzyme_colors);
  for it = 1:length(h)
    h(it).FaceColor = 'Flat';
    h(it).CData = my_colormap(it,:);
  end

  axis([0,3+length(ecm_options.ecm_scores), 0, 1.02]); 
  legend(fliplr(h),gene_names(ind_enzymes_scored),'Fontsize',8,'Location','East')
  set(gca,'FontSize',12);
  set(gca,'XTickLabel',methods); ylabel('Relative protein amount');

end

% color legend

figure(34); clf; 
mmm = nan * ones(nr,1); 
mmm(ecm_options.ind_scored_enzymes) = ind_enzymes(ecm_options.ind_scored_enzymes);
gpp = struct('actprintnames',1,'arrowstyle','none','showsign',0,'colormap',graphics_options.enzyme_colors,'text_offset',[0.03,0]);
gpp.actnames = gene_names;
netgraph_concentrations(network_CoHid,[],column(mmm),1,gpp); 
title('Enzyme colour legend');

% -------------------------------------------------------------------------
% proteomaps
% -------------------------------------------------------------------------

% proteomap from data

if  graphics_options.show_proteomaps,

  figure(350)
  if length(u.data),
    ud = u.data(ecm_options.ind_scored_enzymes);
    ud = ud / nansum(ud);
    %ud = ud(ind_enzymes_scored);
   proteomap_draw_treemap(gene_names(ind_enzymes_scored),ud,{''},struct('color_file',graphics_options.enzyme_colors,'show_level',1,'window_position',[250,120,600,600],'fignums',350,'fontsize',12)); % title('');
  end
  
  % proteomap from predictions
  
  n_methods = length(ecm_options.ecm_scores);
  for it_method = 1:n_methods,
    this_ecm_score = ecm_options.ecm_scores{it_method};
    ug = up.(this_ecm_score)(ecm_options.ind_scored_enzymes,1);
    ug = ug / nansum(ug);
    proteomap_draw_treemap(gene_names(ind_enzymes_scored),ug,{this_ecm_score},struct('color_file',graphics_options.enzyme_colors,'show_level',1,'window_position',[250,120,600,600], 'fignums', 350+it_method));
    if graphics_options.print_graphics,
      cd(graphics_options.psfile_dir);
      figure(350+it_method);
      print([ ecm_options.model_id '_' ecm_options.run_id '_' this_ecm_score '_proteomap.eps'],['-f' num2str(350+it_method)],'-depsc');
    end
  end

end

% -----------------------------------
% PLOTS FOR MEASURED DATA ONLY 
% -----------------------------------

% ---------------------------------------------
% plot flux (data) versus enzyme level (data)

figure(44); clf; hold on
is_finite  = isfinite(u.data .* v);
ind_finite = find(is_finite);
for it = 1:nr,
  if isfinite(u.data(it)),
    plot(1000*u.data(it),abs(v(it)),'.','Color',my_colors(it,:),'MarkerSize',20);
  end
end
text(1.25*1000*u.data(ind_finite),1.05*v(ind_finite),gene_names(ind_finite),'FontSize',12);
set(gca,'XScale','Log','YScale','Log','Fontsize',18);
xlabel('Measured enzyme level (uM)','Fontsize',18); ylabel('Measured flux (mM/s)','Fontsize',18);
if length(ind_finite),
  [cc, pvalue] = corr(log(u.data(ind_finite)),log(abs(v(ind_finite))));
  [cc_spear, pvalue_spear] = corr(log(u.data(ind_finite)),log(abs(v(ind_finite))),'type','Spearman');
  rmse = sqrt(mean([log10(u.data(ind_finite)) - log10(v(ind_finite))].^2));
  %title(sprintf('RMSE=%2.2f, r^2=%2.2f',rmse, cc^2));
  %display('Flux vs enzyme levels:')
  %sprintf('RMSE: %2.2f\n r (Pearson correlation): %2.2f\n r (Spearman rank corr): %2.2f',rmse, cc, cc_spear);
  %  [p-value %2.3f]  [p-value %2.3f] pvalue, pvalue_spear
end
axis tight;% aa = axis; axis([min(aa([1,3])), max(aa([2,4])), min(aa([1,3])), max(aa([2,4]))]); 


% --------------------------------------------------------
% plot flux (data) versus enzyme level * kcat (both data)

figure(45); clf; hold on

is_finite  = isfinite(u.data .* kcat_forward .* v);
ind_finite = find(is_finite);
%amin = min([u.data; abs(v)./kcat_forward]);
%amax = max([u.data; abs(v)./kcat_forward]);
amin = 0.01; amax = 100;
plot([amin, amax],[amin,amax],'--k');

for it = 1:nr,
  if is_finite(it),
    plot(1000*u.data(it),1000*abs(v(it))/kcat_forward(it),'.','Color',my_colors(it,:),'MarkerSize',20);
  end
end
text(1.25*1000*u.data(ind_finite),1.05*1000*v(ind_finite)  ./ kcat_forward(ind_finite),gene_names(ind_finite),'FontSize',12);
set(gca,'XScale','Log','YScale','Log','Fontsize',18);
xlabel('Measured enzyme level (uM)','Fontsize',18); 
ylabel('Measured flux / kcat (uM, data)','Fontsize',18);
if length(ind_finite),
  [cc, pvalue] = corr(log(u.data(ind_finite)),log(abs(v(ind_finite))./kcat_forward(ind_finite)));
  [cc_spear, pvalue_spear] = corr(log(u.data(ind_finite)),log(abs(v(ind_finite))./kcat_forward(ind_finite)),'type','Spearman');
  rmse = sqrt(mean([log10(u.data(ind_finite)) - log10(v(ind_finite)./kcat_forward(ind_finite))].^2));
  text(0.02,90,sprintf('RMSE=%2.2f',rmse),'Fontsize',18);
  text(0.02,40,sprintf('r^2=%2.2f', cc^2),'Fontsize',18);
  text(0.02,15,sprintf('Typical fold error=%2.2f', 10^rmse),'Fontsize',18);
  %title(sprintf('RMSE=%2.2f, r^2=%2.2f',rmse, cc^2));
  %display('Fluxes vs enzyme levels/kcat');
  %sprintf('RMS error: %2.2f\n r (Pearson correlation): %2.2f\n r (Spearman rank corr): %2.2f',rmse, cc, cc_spear); 
  %%  [p-value %2.3f]  [p-value %2.3f] pvalue, pvalue_spear
end
%axis tight; aa = axis; axis([min(aa([1,3])), max(aa([2,4])), min(aa([1,3])), max(aa([2,4]))]); axis square;
axis([amin amax amin amax]); axis square;

% --------------------------------------------------------
% barplot

figure(46); clf; 
n_methods = length(ecm_options.ecm_scores);

for it_method = 1:n_methods,
  this_ecm_score = ecm_options.ecm_scores{it_method};
  ug   = up.(this_ecm_score)(:,1);%(ecm_options.ind_scored_enzymes,1);
  subplot(n_methods+1,1,it_method); hold on;set(gca,'FontSize',10);
  for it = 1:length(ind_enzymes_scored);
    h = bar(it,ug(ind_enzymes_scored(it))); set(h,'FaceColor',graphics_options.enzyme_colors(it,:)); 
  end
  ylabel(sprintf('%s',this_ecm_score));
  set(gca,'XTick',[]); 
end

ud   = u.data;%(ecm_options.ind_scored_enzymes,1);
subplot(n_methods+1,1,it_method+1); hold on
for it = 1:length(ind_enzymes_scored);
  h= bar(it,ud(ind_enzymes_scored(it))); set(h,'FaceColor',graphics_options.enzyme_colors(it,:)); 
end
ylabel('Data');
set(gca,'FontSize',6);
set(gca,'XTick',1:length(ind_enzymes_scored),'XTickLabel',gene_names(ind_enzymes_scored)); 


% --------------------------------------------------------
% SAVE GRAPHICS
% --------------------------------------------------------

if graphics_options.print_graphics,
  cd(graphics_options.psfile_dir);
  if graphics_options.show_network_graphics,
    print([ ecm_options.model_id '_' ecm_options.run_id '_fluxes.eps'],'-f27','-depsc');
    print([ ecm_options.model_id '_' ecm_options.run_id '_kcat_balanced.eps'],'-f4','-depsc');
    print([ ecm_options.model_id '_' ecm_options.run_id '_init_concentrations.eps'],'-f11','-depsc');
    print([ ecm_options.model_id '_' ecm_options.run_id '_init_affinities.eps'],'-f12','-depsc');
    print([ ecm_options.model_id '_' ecm_options.run_id '_data_concentrations.eps'],'-f8','-depsc');
    print([ ecm_options.model_id '_' ecm_options.run_id '_data_enzymes.eps'],'-f9','-depsc');
    print([ ecm_options.model_id '_' ecm_options.run_id '_fix_concentrations.eps'],'-f28','-depsc');
    if graphics_options.show_original_data,
      print([ ecm_options.model_id '_' ecm_options.run_id '_kcat_original.eps'],'-f14','-depsc');
    end
  end
  if graphics_options.show_original_data,
    print([ ecm_options.model_id '_' ecm_options.run_id '_KM_original.eps'],'-f21','-depsc');
  end
  if graphics_options.few_graphics ==0, 
    print([ ecm_options.model_id '_' ecm_options.run_id '_barplot.eps'],'-f33','-depsc');
  end
  print([ ecm_options.model_id '_' ecm_options.run_id '_barplot_legend.eps'],'-f34','-depsc');
  if graphics_options.show_metabolite_graphics,
    print([ ecm_options.model_id '_' ecm_options.run_id '_all_conc_profiles.eps'],'-f18','-depsc');
    print([ ecm_options.model_id '_' ecm_options.run_id '_affinity_profiles.eps'],'-f19','-depsc');
  end
  if graphics_options.show_matrix_graphics,
    print([ ecm_options.model_id '_' ecm_options.run_id '_c_by_KM_initial.eps'],'-f20','-depsc');
    print([ ecm_options.model_id '_' ecm_options.run_id '_KM_balanced.eps'],'-f22','-depsc');
  end
  print([ ecm_options.model_id '_' ecm_options.run_id '_scatter_enzyme_vs_flux.eps'],'-f44','-depsc');
  print([ ecm_options.model_id '_' ecm_options.run_id '_scatter_enzyme_kcat_vs_flux.eps'],'-f45','-depsc');
  print([ ecm_options.model_id '_' ecm_options.run_id '_enzyme_multiple_barplots.eps'],'-f46','-depsc');
  if  graphics_options.show_proteomaps,
    print([ ecm_options.model_id '_' ecm_options.run_id '_proteomap_data.eps'],'-f350','-depsc');
  end
end


% -----------------------------------
% PLOTS FOR INDIVIDUAL SCORES
% -----------------------------------

for it_method = 1:length(ecm_options.ecm_scores),

  fig_offset = 1000 * it_method;
  
  this_ecm_score = ecm_options.ecm_scores{it_method};
  
  my_c = c.(this_ecm_score)(:,1);
  my_A_forward = A_forward.(this_ecm_score)(:,1);
  my_up = up.(this_ecm_score)(:,1);

  if graphics_options.show_network_graphics,

    figure(fig_offset + 1); clf; 
    gp.actstyle = 'none';
    gp.metvaluesmin = log10(ecm_options.conc_min_default);
    gp.metvaluesmax = log10(100);%ecm_options.conc_max_default);
    netgraph_concentrations(network,log10(my_c),[],1,gp); title('Optimised state: log10 Concentrations (mM)');
    
    figure(fig_offset + 2); clf; 
    gp.arrowstyle='none'; 
    gp.actstyle = 'fixed';
    gp.actvaluesmin = log10(0.01);
    gp.actvaluesmax = log10(100);
    gp.metvaluesmin = [];
    gp.metvaluesmax = [];
    my_A_forward(my_A_forward<0) = nan;
    netgraph_concentrations(network_CoHid,[],log10(my_A_forward),1,gp); title('Optimised state: log10 Reaction affinities (kJ/mol)');
  
    figure(fig_offset + 3); clf; 
    gp.actnames  = gene_names;
    gp.actvaluesmin = log10(10^-6);
    gp.actvaluesmax = log10(10^-2);
    gp.metvaluesmin = [];
    gp.metvaluesmax = [];
    netgraph_concentrations(network_CoHid,[],log10(my_up),1,gp); title('Optimised state: log10 Enzyme levels (mM)');

  end

  % --------------------------------------------------------

if graphics_options.show_metabolite_graphics,

  figure(fig_offset + 101); clf; hold on;
  xtick      = 0:length(ind_show_met)-1;
  this_c     = c.(this_ecm_score)(ind_show_met,1);
  if length(c_min),
    this_c_min = c_min.(this_ecm_score)(ind_show_met);
    this_c_max = c_max.(this_ecm_score)(ind_show_met);
  else
    this_c_min = 0;
    this_c_max = 0;
  end
  ind_fixed  = find(label_names(metnames_show, ecm_options.met_fix));

  h = errorbar(xtick,this_c,this_c-this_c_min,this_c_max-this_c,'o','LineWidth',2);
  h = plot(xtick(ind_fixed),this_c(ind_fixed),'r.','MarkerSize',30);
  if length(c.data),
    n_c_data_sets = size(c.data,2);
    plot(repmat(column(xtick),n_c_data_sets,1),reshape(c.data(ind_show_met,:),n_c_data_sets * length(ind_show_met),1),'k.','MarkerSize',20); 
    methods = {this_ecm_score,'fixed','data'};
  else
    methods = {this_ecm_score,'fixed'};
  end
  metnames_show = strrep(metnames_show,'D-','');
  text(xtick + 0.2,c.(this_ecm_score)(ind_show_met),metnames_show,'FontSize',10);
  legend(methods,'Location','NorthEastOutside','FontSize',10);
  ylabel('Concentration [mM]');
  axis tight; set(gca,'YScale','Log','XTick',[]); 
end  
  
  % --------------------------------------
  % enzyme predictions scatter plot
  
  figure(fig_offset + 102); clf; hold on;
  set(gcf,'Position',[50 50 600 600])
  is_finite  = isfinite(u.data) .* [v~=0];
  ind_finite = find(is_finite);

  amin = 0.01; amax = 1000;
  plot([amin, amax],[amin,amax],'--k');

  if ~isfield(network,'genes'),
    network.genes = network.actions;
  end
  
  for it = 1:nr,
    if is_finite(it),
      plot(1000*u.data(it),1000*u.(this_ecm_score)(it,1),'.','Markersize',20,'Color',my_colors(it,:)); 
      text(1.25*1000*u.data(it),1.05*1000*u.(this_ecm_score)(it,1),network.genes(it),'FontSize',12);
      if length(u_min),
        if isfield(u_min,this_ecm_score)
          plot(1000*u.data(it) * [1 1], 1000*[u_min.(this_ecm_score)(it),u_max.(this_ecm_score)(it)],'-','Color',my_colors(it,:)); 
        end
      end
    end
  end

  title(this_ecm_score,'Fontsize',18);
  xlabel('Measured enzyme level (uM)','Fontsize',18);
  ylabel(sprintf('Predicted enzyme level (uM)'),'Fontsize',16); %  %s',this_ecm_score
  set(gca,'XScale','Log','YScale','Log','Fontsize',18);
  axis([amin amax amin amax]); axis square;

  if length(ind_finite),
    [cc, pvalue] = corr(log(u.data(ind_finite)),log(u.(this_ecm_score)(ind_finite,1)));
    [cc_spear, pvalue_spear] = corr(log(u.data(ind_finite)),log(u.(this_ecm_score)(ind_finite,1)),'type','Spearman');
    rmse = sqrt(mean([log10(u.data(ind_finite)) - log10(u.(this_ecm_score)(ind_finite,1))].^2));
  %title(sprintf('RMSE=%2.2f, r^2=%2.2f',rmse, cc^2));
  text(0.02,90,sprintf('RMSE=%2.2f',rmse),'Fontsize',18);
  text(0.02,40,sprintf('r^2=%2.2f', cc^2),'Fontsize',18);
  text(0.02,15,sprintf('Typical fold error=%2.2f', 10^rmse),'Fontsize',18);

  %display(sprintf('Score: %s',this_ecm_score))
  %sprintf('RMS error: %2.2f\n r (Pearson correlation): %2.2f\n r (Spearman rank corr): %2.2f',rmse, cc, cc_spear); 
  %[p-value %2.3f]  [p-value %2.3f] pvalue, pvalue_spear

  end  
  
  % --------------------------------------
  % concentration predictions scatter plot
  
  figure(fig_offset + 103); clf; hold on; set(gcf,'Position',[700 50 600 600])
  
  my_c_min = min(nanmin(c.data(:)),nanmin(c.(this_ecm_score)(:)));
  my_c_max = max(nanmax(c.data(:)),nanmax(c.(this_ecm_score)(:)));
  
  plot([my_c_min,my_c_max],[my_c_min,my_c_max],'--k');
  ind_non_fixed = find(label_names(network.metabolites, ecm_options.met_fix)==0);
  
  for itt = 1:length(ind_non_fixed),
    it = ind_non_fixed(itt);
    plot([nanmin(c.data(it,:)),nanmax(c.data(it,:))],c.(this_ecm_score)(it,1) * [1 1],'-','Color',[.7 .7 .7]);
    if [1-isempty(c_min)],
      if isfield(c_min,this_ecm_score),
      plot(nanmean(c.data(it,:))* [1 1],[c_min.(this_ecm_score)(it),c_max.(this_ecm_score)(it)],'k','Color',[.5 0 0]);
      end
    end
    plot(nanmean(c.data(it,:)),c.(this_ecm_score)(it,1),'r.','Markersize',20);
    text(1.1*nanmean(c.data(it,:)),1.1*c.(this_ecm_score)(it,1),strrep(network.metabolites{it},'_','.'),'FontSize',6);  
  end
  
  title(this_ecm_score,'Fontsize',18);
  xlabel('Median experimental concentration (mM)','Fontsize',16);
  ylabel(sprintf('Predicted concentration (mM) %s',this_ecm_score),'Fontsize',16);
  set(gca,'XScale','Log','YScale','Log','Fontsize',16);
  axis tight; 
  aa = axis; axis([min(aa([1,3])), max(aa([2,4])), min(aa([1,3])), max(aa([2,4]))]); axis square;
  
  my_nanmean_c_data   = nanmean(c.data,2);
  ind_finite          = find(isfinite(my_nanmean_c_data));
if length(ind_finite),
  [cc, pvalue] = corr(log(my_nanmean_c_data(ind_finite)),log(c.(this_ecm_score)(ind_finite,1)));
  [cc_spear, pvalue_spear] = corr(log(my_nanmean_c_data(ind_finite)),log(c.(this_ecm_score)(ind_finite,1)),'type','Spearman');
  rmse = sqrt(mean([log10(my_nanmean_c_data(ind_finite)) - log10(c.(this_ecm_score)(ind_finite,1))].^2));
  %%title(sprintf('RMSE=%2.2f, r^2=%2.2f, Typical fold error=%f',rmse, cc^2, 10^rmse));
  %%title(sprintf('RMSE=%2.2f, r^2=%2.2f, Typical fold error=%f',rmse, cc^2, 10^rmse));

  text(0.005,2,sprintf('RMSE=%2.2f',rmse),'Fontsize',18);
  text(0.005,1.1,sprintf('r^2=%2.2f', cc^2),'Fontsize',18);
  text(0.005,0.5,sprintf('Typical fold error=%2.2f', 10^rmse),'Fontsize',18);

 % display(sprintf('Concentration prediction %s',this_ecm_score))
 %sprintf('RMS error (log10 scale): %2.2f\n r (Pearson correlation): %2.2f\n r (Spearman rank corr): %2.2f',rmse, cc, cc_spear);
 % [p-value %2.3f]  [p-value %2.3f] pvalue, pvalue_spear

end

  % --------------------------------------------------------
  % barplot
  
  ug   = up.(this_ecm_score)(:,1);
  ud   = u.data(:,1);
  
  figure(fig_offset + 104); clf; 
  subplot(2,1,1);
  for it = 1:length(ind_enzymes_scored);
    h = bar(it,ug(ind_enzymes_scored(it))); set(h,'FaceColor',graphics_options.enzyme_colors(it,:)); hold on
  end
  ylabel(sprintf('Predicted enzyme level [mM] %s',this_ecm_score));
  title(sprintf('Total predicted enzyme level (%s): %f mM',this_ecm_score, sum(ug(ind_enzymes_scored))));

  subplot(2,1,2);
  for it = 1:length(ind_enzymes_scored);
    h= bar(it,ud(ind_enzymes_scored(it))); set(h,'FaceColor',graphics_options.enzyme_colors(it,:)); hold on
  end
  set(gca,'FontSize',8);
  set(gca,'XTick',1:length(ind_enzymes_scored),'XTickLabel',gene_names(ind_enzymes_scored)); 
  ylabel('Measured enzyme level [mM]');
  set(gca,'FontSize',8);

  % --------------------------------------
% c over KM

if graphics_options.show_matrix_graphics,

figure(fig_offset + 105); subplot('Position',[0.15 0.05 0.8 0.7]);
c_over_KM = repmat(c.(this_ecm_score)(:,1)',length(network.actions),1)./r.KM;
c_over_KM(r.KM==0) = nan;
im(log10(c_over_KM(:,ind_show_met)),[-3,3],network.actions,metnames_show); colormap([0.95 0.95 0.95; value_colors]); colorbar
my_xticklabel
%title('log10 c (initial solution) / KM');

end

% --------------------------------------------------------
%% arrow cost diagram

my_v              = abs(v(ecm_options.ind_scored_enzymes));
my_u_data         = u.data(ecm_options.ind_scored_enzymes);
my_u              = u.(this_ecm_score)(ecm_options.ind_scored_enzymes);
my_h              = ecm_options.enzyme_cost_weights;
my_eta_energetic  = eta_energetic.(this_ecm_score)(ecm_options.ind_scored_enzymes);
my_eta_saturation = eta_saturation.(this_ecm_score)(ecm_options.ind_scored_enzymes); 
my_u_capacity     = u_capacity(ecm_options.ind_scored_enzymes);

if graphics_options.few_graphics == 0,

  figure(fig_offset + 500); clf; set(gca,'FontSize',14);
  plot(my_v,'-','Color',[0.3 0.3 0.3],'Linewidth',2); hold on
  plot(my_u_capacity,'-','Color',[0 0 0.9],'Linewidth',2);
  plot(my_u_capacity./my_eta_energetic,'-','Color',[0.8 0 0.8],'Linewidth',2);
  plot(my_u,'-','Color',[1 0 0],'Linewidth',2);
  plot(my_u .* my_h,'--','Color',[0.9 0.6 0],'Linewidth',2);
  plot(my_u_data,'k.','Markersize',20);
  axis([1 length(u.data) 10^-5 100]);
  set(gca, 'YScale','Log'); hold off
  title(this_ecm_score)
  my_xticklabel(1:length(ecm_options.ind_scored_enzymes),10^-5,strrep(network.genes(ecm_options.ind_scored_enzymes),'_','-')')
  xlabel('Enzymes'); ylabel('Enzyme cost measures');
  legend('Flux','Flux/k_{cat}','Flux/(k_{cat}*\eta^{energy})','Enzyme','Enzyme cost','Data');

end

figure(fig_offset + 501); clf; 
set(gcf,'Position',[1350 50 600 600])
subplot('Position',[0.15,0.8,0.8,0.15]); set(gca, 'Fontsize',12); hold on;
for itt = 1:length(my_v),
  plot(itt,5+log10(my_v(itt)),'k.','Markersize',15);
end
ylabel('Flux [mM/s]'); 
set(gca,'Ytick',0:6,'YTicklabel',{'0.00001','0.0001','0.001','0.01','0.1','1','10'},'Xtick',[]);
subplot('Position',[0.15,0.1,0.8,0.65]); set(gca, 'Fontsize',12); 
hold on;
M = [5+log10(my_u_capacity), -log10(my_eta_energetic), -log10(my_eta_saturation)];
h = bar(M,'stacked'); 
my_colormap = [0.35 0.35 0.9; 0.8 0.2 0.7; 1 0.3 0.2];
for it = 1:length(h)
  h(it).FaceColor = 'Flat';
  h(it).CData = my_colormap(it,:);
end
hold on; 
h(4) = plot(5+log10(my_u_data),'.','Color',[1 0.7 0],'Markersize',20);
for itt = 1:length(my_v),
  my_xticklabel(itt,-0.5,strrep(network.genes(ecm_options.ind_scored_enzymes(itt)),'_','-')',12,graphics_options.enzyme_colors(itt,:))
end
axis([0,length(my_u_capacity)+1,0,4.1])
ylabel('Enzyme demand [uM]'); set(gca,'Ytick',0:4,'YTicklabel',{'0.01','0.1','1','10','100'});

switch this_ecm_score,
  case 'emc4cm',
    legend(h([4 3 2 1]),'Data','Saturation','Reversibility','Capacity','Location','SouthWest');
end

%title(this_ecm_score)

if graphics_options.show_network_graphics,
figure(fig_offset + 502); clf
dum = nan * v;
dum(ecm_options.ind_scored_enzymes) = my_eta_energetic;
netgraph_concentrations(network_CoHid, [], dum, 1, struct('actvalues',dum,'actstyle','fixed','showsign',0)); title('\eta^{\rm therm}');
figure(fig_offset + 503); clf
dum(ecm_options.ind_scored_enzymes) = my_eta_saturation;
netgraph_concentrations(network_CoHid, [], dum, 1, struct('actvalues',dum,'actstyle','fixed','showsign',0)); title('\eta^{\rm kin}');
end

  % -------------------------------------------------------------------------------------
  
  if graphics_options.print_graphics,
    cd(graphics_options.psfile_dir);
    if graphics_options.show_network_graphics,
      print([ ecm_options.model_id '_' ecm_options.run_id '_' this_ecm_score '_opt_concentrations.eps'],sprintf('-f%d',fig_offset + 1),'-depsc');
      print([ ecm_options.model_id '_' ecm_options.run_id '_' this_ecm_score '_opt_affinities.eps'],    sprintf('-f%d',fig_offset + 2),'-depsc');
      print([ ecm_options.model_id '_' ecm_options.run_id '_' this_ecm_score '_opt_enzymes.eps'],       sprintf('-f%d',fig_offset + 3),'-depsc');
      print([ ecm_options.model_id '_' ecm_options.run_id '_' this_ecm_score '_cost_diagram_eta_therm.eps'],sprintf('-f%d',fig_offset + 502),'-depsc');
      print([ ecm_options.model_id '_' ecm_options.run_id '_' this_ecm_score '_cost_diagram_eta_kin.eps'],  sprintf('-f%d',fig_offset + 503),'-depsc');
    end
    if graphics_options.show_metabolite_graphics,
      print([ ecm_options.model_id '_' ecm_options.run_id '_' this_ecm_score '_conc_prediction.eps'],  sprintf('-f%d',fig_offset + 101),'-depsc');
    end
    print([ ecm_options.model_id '_' ecm_options.run_id '_' this_ecm_score '_enzyme_scatter.eps'], sprintf('-f%d',fig_offset + 102),'-depsc');
    print([ ecm_options.model_id '_' ecm_options.run_id '_' this_ecm_score '_conc_scatter.eps'], sprintf('-f%d',fig_offset + 103),'-depsc');
    print([ ecm_options.model_id '_' ecm_options.run_id '_' this_ecm_score '_enzyme_prediction.eps'],sprintf('-f%d',fig_offset + 104),'-depsc');
    if    graphics_options.show_matrix_graphics,
      print([ ecm_options.model_id '_' ecm_options.run_id '_' this_ecm_score '_c_by_KM.eps'],sprintf('-f%d',fig_offset + 105),'-depsc');
    end
    if graphics_options.few_graphics == 0,
      print([ ecm_options.model_id '_' ecm_options.run_id '_' this_ecm_score '_arrow_cost_diagram.eps'],sprintf('-f%d',fig_offset + 500),'-depsc');
    end
    print([ ecm_options.model_id '_' ecm_options.run_id '_' this_ecm_score '_arrow_cost_diagram_bar.eps'],sprintf('-f%d',fig_offset + 501),'-depsc');
  end
  
end

if graphics_options.print_graphics,
  display(sprintf('Graphics files written to directory %s',graphics_options.psfile_dir));
end
```
