## Supplementary material for "In-depth computational analysis of natural and artificial carbon fixation pathways": MATLAB files: ecm_get_one_u.m

```
function my_u_it = ecm_get_one_u(it, ecm_score, xx, pp)

% ECM_GET_ONE_U - Helper function for enzyme variability calculation after enzyme cost minimization:
%
% function my_u_it = ecm_get_one_u(it,ecm_score,xx,pp)
%
% given log metabolite profile xx, evaluate enzyme levels and return the levels of the it'th enzyme

[~, my_u] = ecm_get_score(ecm_score, xx, pp);

my_u_it   = my_u(it);
```
