## Supplementary material for "In-depth computational analysis of natural and artificial carbon fixation pathways": MATLAB files: ecm_get_score.m

```
% ECM_GET_SCORE - Helper function
% 
% function [u_cost, u, u_preemptive] = ecm_get_score(ecm_score,x,pp,no_warnings)
%
% Possible cost functions: 
%  Max-min driving force
%   mdf
%   
%  Enzyme cost based on different rate law approximations (see Noor et al 2016)
%   emc1
%   emc2s
%   emc2sp        
%   emc3s         
%   emc3sp        
%   emc4dm        
%   emc4sm        
%   emc4cm        
%   emc4geom      
%   
%  Maximal cost among enzymes
%   max_1
%   max_2s        
%   max_2sp
%   max_3s        
%   max_3sp        
%   max_4cm 
%   
%  Optimal fit (sum of squared residuals) for metabolite and  enzymes data
%   fitting_1     
%   fitting_2s    
%   fitting_2sp   
%   fitting_3s    
%   fitting_3sp   
%   fitting_4cm   
%   
%  Sum of enzyme concentrations with safety tolerance for enzyme expression
%   robust_enzconc_1  
%   robust_enzconc_2s 
%   robust_enzconc_2sp
%   robust_enzconc_3s 
%   robust_enzconc_3sp
%   robust_enzconc_4cm
%   
%  Sum of metabolite and enzyme concentrations
%   metenzconc_1  
%   metenzconc_2s 
%   metenzconc_2sp
%   metenzconc_3s 
%   metenzconc_3sp
%   metenzconc_4cm
%   
%  Sum of metabolite and enzyme concentrations with safety tolerance for enzyme expression
%   robust_metenzconc_1  
%   robust_metenzconc_2s 
%   robust_metenzconc_2sp
%   robust_metenzconc_3s 
%   robust_metenzconc_3sp
%   robust_metenzconc_4cm
%   
%  Sum of mass-weighted metabolite and enzyme concentrations
%   metenzmass_1  
%   metenzmass_2s 
%   metenzmass_2sp
%   metenzmass_3s 
%   metenzmass_3sp
%   metenzmass_4cm

function [u_cost, u, u_preemptive] = ecm_get_score(ecm_score,x,pp,no_warnings)

  eval(default('no_warnings','0'));
  
switch ecm_score,
  case 'mdf',                   [u_cost, u] = ecm_mdf(x,pp);      % only max Delta G counts  
  case 'emc1',                  [u_cost, u] = ecm_emc1(x,pp);     % sum values
  case 'emc2s',                 [u_cost, u] = ecm_emc2s(x,pp);
  case 'emc2sp',                [u_cost, u] = ecm_emc2sp(x,pp);
  case 'emc3s',                 [u_cost, u] = ecm_emc3s(x,pp);
  case 'emc3sp',                [u_cost, u] = ecm_emc3sp(x,pp);
  case 'emc4dm',                [u_cost, u] = ecm_emc4dm(x,pp);
  case 'emc4sm',                [u_cost, u] = ecm_emc4sm(x,pp);
  case 'emc4cm',                [u_cost, u] = ecm_emc4cm(x,pp,no_warnings);
  case 'emc4geom',              [u_cost, u] = ecm_emc4geom(x,pp);  
  case 'max_1',                 [u_cost, u] = ecm_max_enzyme_cost(x,pp,'emc1');   % max values
  case 'max_2s',                [u_cost, u] = ecm_max_enzyme_cost(x,pp,'emc2s');
  case 'max_2sp',               [u_cost, u] = ecm_max_enzyme_cost(x,pp,'emc2sp');
  case 'max_3s',                [u_cost, u] = ecm_max_enzyme_cost(x,pp,'emc3s');
  case 'max_3sp',               [u_cost, u] = ecm_max_enzyme_cost(x,pp,'emc3sp');
  case 'max_4cm',               [u_cost, u] = ecm_max_enzyme_cost(x,pp,'emc4cm');
  case 'fitting_1',             [u_cost, u] = ecm_fitting(x,pp,'emc1');
  case 'fitting_2s',            [u_cost, u] = ecm_fitting(x,pp,'emc2s');
  case 'fitting_2sp',           [u_cost, u] = ecm_fitting(x,pp,'emc2sp');
  case 'fitting_3s',            [u_cost, u] = ecm_fitting(x,pp,'emc3s');
  case 'fitting_3sp',           [u_cost, u] = ecm_fitting(x,pp,'emc3sp');
  case 'fitting_4cm',           [u_cost, u] = ecm_fitting(x,pp,'emc4cm');
  case 'metenzconc_1',          [u_cost, u] = ecm_metenz_conc(x,pp,'emc1');
  case 'metenzconc_2s',         [u_cost, u] = ecm_metenz_conc(x,pp,'emc2s');
  case 'metenzconc_2sp',        [u_cost, u] = ecm_metenz_conc(x,pp,'emc2sp');
  case 'metenzconc_3s',         [u_cost, u] = ecm_metenz_conc(x,pp,'emc3s');
  case 'metenzconc_3sp',        [u_cost, u] = ecm_metenz_conc(x,pp,'emc3sp');
  case 'metenzconc_4cm',        [u_cost, u] = ecm_metenz_conc(x,pp,'emc4cm');
  case 'robust_enzconc_1',      [u_cost, u] = ecm_robust_enz(x,pp,'emc1');
  case 'robust_enzconc_2s',     [u_cost, u] = ecm_robust_enz(x,pp,'emc2s');
  case 'robust_enzconc_2sp',    [u_cost, u] = ecm_robust_enz(x,pp,'emc2sp');
  case 'robust_enzconc_3s',     [u_cost, u] = ecm_robust_enz(x,pp,'emc3s');
  case 'robust_enzconc_3sp',    [u_cost, u] = ecm_robust_enz(x,pp,'emc3sp');
  case 'robust_enzconc_4cm',    [u_cost, u] = ecm_robust_enz(x,pp,'emc4cm');
  case 'robust_metenzconc_1',   [u_cost, u] = ecm_robust_metenz(x,pp,'emc1');
  case 'robust_metenzconc_2s',  [u_cost, u] = ecm_robust_metenz(x,pp,'emc2s');
  case 'robust_metenzconc_2sp', [u_cost, u] = ecm_robust_metenz(x,pp,'emc2sp');
  case 'robust_metenzconc_3s',  [u_cost, u] = ecm_robust_metenz(x,pp,'emc3s');
  case 'robust_metenzconc_3sp', [u_cost, u] = ecm_robust_metenz(x,pp,'emc3sp');
  case 'robust_metenzconc_4cm', [u_cost, u] = ecm_robust_metenz(x,pp,'emc4cm');
  case 'metenzmass_1',          [u_cost, u] = ecm_metenz_mass(x,pp,'emc1');
  case 'metenzmass_2s',         [u_cost, u] = ecm_metenz_mass(x,pp,'emc2s');
  case 'metenzmass_2sp',        [u_cost, u] = ecm_metenz_mass(x,pp,'emc2sp');
  case 'metenzmass_3s',         [u_cost, u] = ecm_metenz_mass(x,pp,'emc3s');
  case 'metenzmass_3sp',        [u_cost, u] = ecm_metenz_mass(x,pp,'emc3sp');
  case 'metenzmass_4cm',        [u_cost, u] = ecm_metenz_mass(x,pp,'emc4cm');
  
  otherwise,             error('Unknown ecm_score');
end

u(pp.v==0) = 0;

% Safety margin, to counter fluctuations in protein levels

if pp.fluctuations_safety_margin,
  u = u + pp.fluctuations_safety_margin / sqrt(pp.cell_volume * 6.02214086 * 10^23) * sqrt(u);
end

% Enzyme cost functions for multiple conditions
% Assume that all reaction vectors are actually a concatenation of several (condition-specific) vectors

if pp.multiple_conditions_anticipate,
  %% Recompute enzyme values and total cost
  nu             = length(u) / pp.multiple_conditions_n;
  u_matrix       = reshape(u,nu,pp.multiple_conditions_n);
  u_preemptive   = repmat(nanmax(u_matrix,[],2),pp.multiple_conditions_n,1);
  switch ecm_score,
    case {'emc1', 'emc2s', 'emc2sp', 'emc3s',  'emc3sp', 'emc4dm', 'emc4sm', 'emc4cm', 'emc4geom'},
      %for simple cost functions: sum over weighted enzyme concentrations
      u_cost = nansum(pp.enzyme_cost_weights .* u_preemptive(pp.ind_scored_enzymes));
    otherwise, 
      %more complicated: sum over weighted enzyme concentrations
      %plus other terms -> adjust the original cost! substract the enzyme cost and add the
      enz_cost_old = nansum(pp.enzyme_cost_weights .* u(pp.ind_scored_enzymes) );
      enz_cost_new = nansum(pp.enzyme_cost_weights .* u_preemptive(pp.ind_scored_enzymes) );
      u_cost       = u_cost - enz_cost_old + enz_cost_new;
  end
else
  u_preemptive = [];
end
```
