## Supplementary material for "In-depth computational analysis of natural and artificial carbon fixation pathways": MATLAB files: ecm_get_specific_rates.m

```
% ECM_GET_SPECIFIC_RATES - Helper function
%
% function rates = ecm_get_specific_rates(ecm_score,x,pp,ind)

function rates = ecm_get_specific_rates(ecm_score,x,pp,ind)

switch ecm_score,
  case 'mdf',      [~, ~, rates] = ecm_mdf(x,pp);
  case 'memc2s',   [~, ~, rates] = ecm_memc2s(x,pp);
  case 'memc2',    [~, ~, rates] = ecm_memc2sp(x,pp);
  case 'emc1',     [~, ~, rates] = ecm_emc1(x,pp);
  case 'emc2s',    [~, ~, rates] = ecm_emc2s(x,pp);
  case 'emc2sp',   [~, ~, rates] = ecm_emc2sp(x,pp);
  case 'emc3s',    [~, ~, rates] = ecm_emc3s(x,pp);
  case 'emc3sp',   [~, ~, rates] = ecm_emc3sp(x,pp);
  case 'emc4dm',   [~, ~, rates] = ecm_emc4dm(x,pp);
  case 'emc4sm',   [~, ~, rates] = ecm_emc4sm(x,pp);
  case 'emc4cm',   [~, ~, rates] = ecm_emc4cm(x,pp);
  case 'emc4geom', [~, ~, rates] = ecm_emc4geom(x,pp);
  case 'fitting_emc3sp', [~, ~, rates] = ecm_fitting_emc3sp(x,pp);
  otherwise,       error('Unknown ecm score');
end

if exist('ind','var'),  rates = rates(ind);  end
```
