## Supplementary material for "In-depth computational analysis of natural and artificial carbon fixation pathways": MATLAB files: ecm_inequalities.m

```
% MEASURES_FOR_ENZYME_COSTS_INEQUALITIES - Helper function 
%
% [delta_G_by_RT,eq_cons] = measures_for_enzyme_costs_inequalities(x,N_forward,log_Keq_forward)
%

function [delta_G_by_RT,eq_cons] = measures_for_enzyme_costs_inequalities(x,N_forward,log_Keq_forward,Theta_min)

delta_G_by_RT = - [log_Keq_forward-Theta_min-[N_forward' * x]];

eq_cons = [];
```
