## Supplementary material for "In-depth computational analysis of natural and artificial carbon fixation pathways": MATLAB files: ecm_kegg_compound_id_to_name.m

```
% ECM_KEGG_COMPOUND_ID_TO_NAME - Name conversion for compounds
%
% function names = ecm_kegg_compound_id_to_name(ids,kegg_conversion_file)

function names = ecm_kegg_compound_id_to_name(ids,kegg_conversion_file)

if ~exist('kegg_conversion_file','var'),
  ecm_info = ecm_setup;
  kegg_conversion_file = [ecm_info.data_dir 'name-conversion/kegg_compound_names.tsv'];
end
  
T = sbtab_table_load(kegg_conversion_file);

kegg_id   = sbtab_table_get_column(T, '!Identifiers:kegg.compound');
kegg_name = sbtab_table_get_column(T, '!Compound:Name');
ll = label_names(ids,kegg_id);
names = repmat({''},length(ids),1);
names(find(ll)) = kegg_name(ll(find(ll)));
```
