## Supplementary material for "In-depth computational analysis of natural and artificial carbon fixation pathways": MATLAB files: ecm_load_result_sbtab.m

```
function [c_emc4cm, u_emc4cm] = ecm_load_result_sbtab(network, state_runs_file_sbtab, emc_score);

M = sbtab_document_load(state_runs_file_sbtab);
predConc = sbtab_document_get_table(M,'Predictedconcentations');
predEnz  = sbtab_document_get_table(M,'Predictedenzymelevels');
 
ll = label_names(sbtab_table_get_column(predConc,'Compound'),network.metabolites);
my_c = cell_string2num(predConc.uncontrolled.data(:,find(strcmp(predConc.uncontrolled.headers,emc_score))));
c_emc4cm = my_c(ll);

ll = label_names(sbtab_table_get_column(predEnz,'Reaction'), network.actions);
my_u = cell_string2num(predEnz.uncontrolled.data(:,find(strcmp(predEnz.uncontrolled.headers,emc_score))));
u_emc4cm = my_u(ll);
```
