## Supplementary material for "In-depth computational analysis of natural and artificial carbon fixation pathways": MATLAB files: ecm_multiple_network.m

```
function [network_multi, r_multi, ecm_options_multi] = ecm_multiple_network(network,r,n_conditions, ecm_options)

% [network_multi, r_multi, ecm_options_multi] = ecm_multiple_network(network,r,n_conditions,ecm_options)
%
% build network and kinetics structure with multiple copies of the original model 
% (associated with the different conditions to be simulated)
%
% This is only needed for multiple condition ECM

  
% ------------------------------------------------------------------
% Network
  
[nm,nr] = size(network.N);
network_multi.metabolites = {};
network_multi.actions     = {};
for it = 1:n_conditions,
  network_multi.metabolites = [network_multi.metabolites; numbered_names_simple(['M' num2str(it) '_'],nm)];
  network_multi.actions = [network_multi.actions; numbered_names_simple(['R' num2str(it) '_'],nr)];
end

network_multi.N                 = blockrep(network.N,         n_conditions);
network_multi.reversible        = repmat(network.reversible,  n_conditions,1);
network_multi.external          = repmat(network.external,    n_conditions,1);
network_multi.regulation_matrix = blockrep(network.regulation_matrix,        n_conditions);

% ------------------------------------------------------------------
% Kinetics
  

r_multi.type  =  r.type;
r_multi.u     =  [];
r_multi.c     =  [];
r_multi.KA    =  blockrep(r.KA,  n_conditions);
r_multi.KI    =  blockrep(r.KI,  n_conditions);
r_multi.KM    =  blockrep(r.KM,  n_conditions);
r_multi.KV    =  repmat(r.KV,    n_conditions, 1);
r_multi.Keq   =  repmat(r.Keq,   n_conditions, 1);
r_multi.h     =  repmat(r.h,     n_conditions, 1);
r_multi.Kcatf =  repmat(r.Kcatf, n_conditions, 1);
r_multi.Kcatr =  repmat(r.Kcatr, n_conditions, 1);


% ------------------------------------------------------------------
% Options
  
ecm_options_multi = ecm_options;%ecm_default_options(network_multi,'E coli multi-condition test model');
ecm_options_multi = ecm_update_options(network_multi, ecm_options_multi);
ecm_options_multi.multiple_conditions_n = n_conditions;
ecm_options_multi.conc_min            = repmat(ecm_options.conc_min,n_conditions,1);
ecm_options_multi.conc_max            = repmat(ecm_options.conc_max,n_conditions,1);
ecm_options_multi.enzyme_cost_weights = repmat(ecm_options.enzyme_cost_weights,n_conditions,1);
for it = 1:n_conditions-1;
  ecm_options_multi.ind_scored_enzymes = [ecm_options_multi.ind_scored_enzymes; it * nr + ecm_options.ind_scored_enzymes];
end
ecm_options_multi.show_metabolites  = repmat(ecm_options.show_metabolites,n_conditions,1);
ecm_options_multi.N_subunits_in_complex  = repmat(ecm_options.N_subunits_in_complex,n_conditions,1);
ecm_options_multi.N_cat_sites_in_complex = repmat(ecm_options.N_cat_sites_in_complex,n_conditions,1);
```
