## Supplementary material for "In-depth computational analysis of natural and artificial carbon fixation pathways": MATLAB files: ecm_one_run.m

```
% ECM_ONE_RUN - Perform one ECM run
% 
% function [my_c, my_u, my_up, my_u_cost, my_A_forward, my_x, my_grad, my_lambda] = ecm_one_run(ecm_score,pp,x_min,x_max,x_init,lambda_regularisation,opt)
%
% my_up is the same as my_u, except for the fact that the levels of unscored enzymes are replaced by nan
% In the case of multiple-condition ECM, my_up contains the preemptive protein levels (i.e., only one value per reaction)

function [my_c, my_u, my_up, my_u_cost, my_A_forward, my_x, my_grad, my_lambda] = ecm_one_run(ecm_score,pp,x_min,x_max,x_init,lambda_regularisation, Theta_min, opt)

%% optimize log metabolite concentration profile

%% Set global variables to speed up function modular_velocities
global global_structure_matrices 
global_structure_matrices = 1;
global Mplus Mminus Wplus Wminus nm nr ind_M ind_Wp ind_Wm
N = pp.network.N; W = pp.network.regulation_matrix; ind_ext = find(pp.network.external); h = pp.network.kinetics.h;
[Mplus, Mminus, Wplus, Wminus, nm, nr, N_int,ind_M,ind_Wp,ind_Wm] = make_structure_matrices(N,W,ind_ext,h);
%% END Set global variables

try
    [my_x, my_fval,my_exitflag,my_output,my_lambda,my_grad] = fmincon(@(xx) ecm_get_score(ecm_score,xx,pp) + ecm_regularisation(xx,x_min,x_max,lambda_regularisation), x_init,[],[],[],[],x_min,x_max,@(xx) ecm_inequalities(xx,pp.N_forward,pp.log_Keq_forward, Theta_min),opt);
catch
    my_exitflag = -10;
end

if my_exitflag < 1
    my_c = [];
    my_u = [];
    my_up = [];
    my_u_cost = [];
    my_A_forward = [];
    my_x = [];
    my_grad = [];
    my_lambda = [];
    return;
    %error('optimisation unsuccessful'); 
end 

my_c         = exp(my_x);
my_A_forward = RT * [pp.log_Keq_forward - pp.N_forward' * log(my_c)]; 
my_A_forward(pp.ind_not_scored) = nan;

%% compute resulting enzyme profile

[my_u_cost, my_u, my_u_preemptive] = ecm_get_score(ecm_score, my_x, pp);

my_up                    = my_u;

if pp.multiple_conditions_anticipate, 
  my_up = my_u_preemptive; 
end

my_up(pp.ind_not_scored) = nan;
```
