## Supplementary material for "In-depth computational analysis of natural and artificial carbon fixation pathways": MATLAB files: ecm_options_read.m

```
function ecm_options = ecm_options_read(filename)

% ECM_OPTIONS_READ - Read options struct from JSON file
%
% ecm_options = ecm_options_read(filename)

X           = load_any_table(filename);
ecm_options = jsondecode(X{1,1});
```
