## Supplementary material for "In-depth computational analysis of natural and artificial carbon fixation pathways": MATLAB files: ecm_options_save.m

```
function ecm_options_save(filename,ecm_options,verbose)

% ECM_OPTIONS_SAVE - Save options struct to JSON file
%
% ecm_options_save(filename, ecm_options)

% remove all fields that are structs (because they cannot be exported)
% remove all fields that are strings containing "/" characters, such as filenames (because they cannot be exported)

  eval(default('verbose','1'));
  
fn = fieldnames(ecm_options);
for it = 1:length(fn),
  if isstruct(ecm_options.(fn{it})), 
    ecm_options = rmfield(ecm_options,fn{it});
  elseif isstr(ecm_options.(fn{it})), 
    if findstr(ecm_options.(fn{it}),'/'),
      ecm_options = rmfield(ecm_options,fn{it});
    end
  end
end
  
fileID = fopen(filename,'w');

fprintf(fileID,jsonencode(ecm_options));

if verbose,
  display(sprintf('Writing file %s',filename));
end
```
