## Supplementary material for "In-depth computational analysis of natural and artificial carbon fixation pathways": MATLAB files: ecm_read_options.m

```
% ECM_READ_OPTIONS - Read ECM options from file
% 
% options = ecm_read_options(options_file, default_options)

function options = ecm_read_options(options_file, default_options)

eval(default('default_options','struct'));

options = struct('model_name','');
fid     = fopen(options_file);
c       = fgetl(fid);

while c ~= -1,
  eval(c);
  c = fgetl(fid);
end

options = join_struct(default_options, options);
```
