## Supplementary material for "In-depth computational analysis of natural and artificial carbon fixation pathways": MATLAB files: ecm_regularisation.m

```
% ECM_REGULARISATION - Regularisation term for ECM 
% 
% function f = ecm_regularisation(x,x_min,x_max,lambda)

function f = ecm_regularisation(x,x_min,x_max,lambda)

f = lambda * 0.5 * sum([x - 0.5 * [x_min + x_max]].^2);
```
