## Supplementary material for "In-depth computational analysis of natural and artificial carbon fixation pathways": MATLAB files: ecm_save_model_and_data_neos.m

```
function ecm_save_model_and_data_neos(filename,network,v,r,c_data,u_data,enzyme_cost_weights,conc_min,conc_max, met_fix,conc_fix)

% ECM_SAVE_MODEL_AND_DATA_NEOS - Save input files for ECM by NEOS solvers
% 
% ecm_save_model_and_data_neos(filename,network,v,r,c_data,u_data,enzyme_cost_weights,conc_min,conc_max, met_fix,conc_fix)
% 
% Convert data for Enzyme Cost Minimization (model and data) from SBtab format to NEOS input format
% 
% For generating the input file (SBtab format), see 'help ecm_save_model_and_data_sbtab'

display(sprintf('\nWriting input files for the NEOS optimization server to directory %s', filename))

% adjust names for sbml output

[network.metabolites,network.actions] = network_adjust_names_for_sbml_export(network.metabolites,network.actions,1);

% ecm_options.fix_metabolites = network_adjust_names_for_sbml_export(ecm_options.fix_metabolites,[],1);

% turn network + kinetics into sbtab format

network.kinetics = r;
network.kinetics.type = 'cs';
network_sbtab = network_to_sbtab(network, struct('use_sbml_ids',0,'verbose',0,'write_concentrations',0,'write_enzyme_concentrations',0));

quantity_type = network_sbtab.tables.Parameter.column.column.QuantityType;
if isfield(network_sbtab.tables.Parameter.column.column,'Value'),
  value         = network_sbtab.tables.Parameter.column.column.Value;
else
  value         = network_sbtab.tables.Parameter.column.column.Mode;
end
compound      = network_sbtab.tables.Parameter.column.column.Compound;
reaction      = network_sbtab.tables.Parameter.column.column.Reaction;

[compound,reaction] = network_adjust_names_for_sbml_export(compound,reaction,1);
%display('Writing NEOS files');

% -----------------------------------------------------------
% Write neos input files

% write table stoich.csv

ind = find(strcmp('Michaelis constant', quantity_type));

stoich = [];
for it = 1:length(ind),
  i_met = find(strcmp(compound{ind(it)},network.metabolites));
  i_rea = find(strcmp(reaction{ind(it)},network.actions));
  stoich(it,1) = network.N(i_met,i_rea);
end

mytable([reaction(ind), compound(ind), num2cell([stoich])],'comma',[filename 'stoich.csv']);

% write table kms.csv

ind = find(strcmp('Michaelis constant', quantity_type));

mytable([reaction(ind), compound(ind), num2cell([value(ind)])],'comma',[filename 'kms.csv']);

% write table metfixed.csv

mytable([column(met_fix), num2cell(column(conc_fix))],'comma',[filename 'metfixed.csv']);

% write table boundse.csv

[nm,nr] = size(network.N);

mytable([[network.actions, repmat({'lo','0'},nr,1)]; ...
         [network.actions, repmat({'up','100'},nr,1)]],'comma',[filename 'boundse.csv']);

% write table boundsx.csv

mytable([[network.metabolites, repmat({'lo'},nm,1), num2cell(conc_min)]; ...
         [network.metabolites, repmat({'up'},nm,1), num2cell(conc_max)]],'comma',[filename 'boundsx.csv']);

% write table rates.csv

mytable([[{'0'}, network.actions']; [{'1'}, num2cell(v)']],'comma',[filename 'rates.csv']);

% write table weights.csv

enzyme_cost_weights(~isfinite(enzyme_cost_weights)) = 0;
mytable([network.actions, num2cell(enzyme_cost_weights)],'comma',[filename 'weights.csv']);

% write table kcats.csv

ind = find(strcmp('substrate catalytic rate constant', quantity_type));
mytable([reaction(ind), num2cell(value(ind))],'comma',[filename 'kcats.csv']);

% write table keqs.csv

ind = find(strcmp('equilibrium constant', quantity_type));
mytable([reaction(ind), num2cell(value(ind))],'comma',[filename 'keqs.csv']);

% write table moietymet.csv

%mytable({[]},'comma',[filename 'moietymet.csv']);

% write table moietyval.csv

%mytable({[]},'comma',[filename 'moietyval.csv']);

% write table alpha.csv

%mytable({[]},'comma',[filename 'alpha.csv']);

% write table a.csv

%mytable({[]},'comma',[filename 'a.csv']);

% write table newmet.csv (DEPRECATED)
% mytable({[]},'comma',[filename 'newmet.csv']);

% write table activators.csv

%ind = find(strcmp('activation constant', quantity_type));
%mytable([compound(ind), reaction(ind), num2cell(value(ind))],'comma',[filename 'activators.csv']);

% write table inhibitors.csv

%ind = find(strcmp('inhibition constant', quantity_type));
%mytable([compound(ind), reaction(ind), num2cell(value(ind))],'comma',[filename 'inhibitors.csv']);

% write table efms.csv (DEPRECATED)
% mytable([ [{'efms'}; network.actions], [{1}; num2cell(v)] ]','comma',[filename 'efms.csv']);

%eval(sprintf(' cd %s; zip MODEL.zip *tsv;', filename));

display(sprintf('To run an optimisation with these files, please use the NEOS server at https://proto.neos-server.org/neos/solvers/application:MER/csv.html'))


% -----------------------------------------------------------------
% Build neos sbtab file

document_name = 'NEOS input file';

% write table stoich.csv
ind = find(strcmp('Michaelis constant', quantity_type));
table_stoich = sbtab_table_construct(struct('TableName','stoich','TableType','Stoichiometries','Document',document_name),{'Reaction','Compound','StoichiometricCoefficient'},{reaction(ind), compound(ind), num2cell([stoich])});

% write table metfixed.csv
%mytable([column(met_fix), num2cell(column(conc_fix))],'comma',[filename 'metfixed.csv']);
table_metfixed = sbtab_table_construct(struct('TableName','metfixed','TableType','Compound','Document',document_name),{'Compound','IsFixed'},{column(met_fix), num2cell(column(conc_fix))});

% write table kcats.csv
ind = find(strcmp('substrate catalytic rate constant', quantity_type));
%mytable([reaction(ind), num2cell(value(ind))],'comma',[filename 'kcats.csv']);
table_kcats = sbtab_table_construct(struct('TableName','kcats','TableType','Reaction','Document',document_name),{'Reaction','Value'},{reaction(ind), num2cell(value(ind))});

% write table keqs.csv
ind = find(strcmp('equilibrium constant', quantity_type));
%mytable([reaction(ind), num2cell(value(ind))],'comma',[filename 'keqs.csv']);
table_keqs = sbtab_table_construct(struct('TableName','keqs','TableType','Reaction','Document',document_name),{'Reaction','Value'},{reaction(ind), num2cell(value(ind))});

% write table kms.csv
ind = find(strcmp('Michaelis constant', quantity_type));
%mytable([reaction(ind), compound(ind), num2cell([value(ind)])],'comma',[filename 'kms.csv']);
table_kms = sbtab_table_construct(struct('TableName','kms','TableType','Quantity','Document',document_name),{'Reaction','Compound','Value'},{reaction(ind), compound(ind), num2cell([value(ind)])});

% write table weights.csv
enzyme_cost_weights(~isfinite(enzyme_cost_weights)) = 0;
%mytable([network.actions, num2cell(enzyme_cost_weights)],'comma',[filename 'weights.csv']);
table_weights = sbtab_table_construct(struct('TableName','weights','TableType','Reaction','Document',document_name),{'Reaction','Value'},{network.actions, num2cell(enzyme_cost_weights)});

% write table rates.csv
% mytable([[{'0'}, network.actions']; [{'1'}, num2cell(v)']],'comma',[filename 'rates.csv']);
table_rates = sbtab_table_construct(struct('TableName','rates','TableType','Reaction','Document',document_name),{'Counter'},{{'0','1'}});
for it = 1:length(network.actions),
  table_rates = sbtab_table_add_column(table_rates,network.actions{it},{v(it)});
end

% write table boundse.csv
%mytable([[network.actions, repmat({'lo','0'},nr,1)]; ...
%         [network.actions, repmat({'up','100'},nr,1)]],'comma',[filename 'boundse.csv']);
table_boundse = sbtab_table_construct(struct('TableName','boundse','TableType','Enzyme','Document',document_name),{'Reaction','BoundType','Value'},{[network.actions; network.actions], [repmat({'lo'},nr,1);repmat({'up'},nr,1)], [repmat({'0'},nr,1);repmat({'100'},nr,1)]});

% write table boundsx.csv
%mytable([[network.metabolites, repmat({'lo'},nm,1), num2cell(conc_min)]; ...
%         [network.metabolites, repmat({'up'},nm,1), num2cell(conc_max)]],'comma',[filename 'boundsx.csv']);
table_boundsx = sbtab_table_construct(struct('TableName','boundsx','TableType','Compound','Document',document_name),{'Compound','BoundType','Value'},{[network.metabolites; network.metabolites], [repmat({'lo'},nm,1); repmat({'up'},nm,1)] , [num2cell(conc_min); num2cell(conc_max)] });


%sbtab_table_show(stoichiometry_table)
neos_sbtab = sbtab_document_construct(struct,{'stoich','metfixed','kcats', 'keqs','kms', 'weights', 'rates', 'boundse', 'boundsx'},{table_stoich,table_metfixed, table_kcats, table_keqs, table_kms, table_weights, table_rates, table_boundse, table_boundsx});

% sbtab_document_show(neos_sbtab)

sbtab_document_save_to_one(neos_sbtab,[filename 'NEOS_input_sbtab.tsv']);
```
