## Supplementary material for "In-depth computational analysis of natural and artificial carbon fixation pathways": MATLAB files: ecm_save_model_and_data_sbtab.m

```
function ecm_save_model_and_data_sbtab(filename, network, v, r, c_data, u_data, kinetic_data, conc_min, conc_max, met_fix, conc_fix, enzyme_cost_weights, document_name, save_single_tables,verbose)

%ECM_SAVE_MODEL_AND_DATA_SBTAB - Write SBtab file containing model and validation data for Enzyme Cost Minimization
%
% ecm_save_model_and_data_sbtab(filename, network, v, r, c_data, u_data, kinetic_data, conc_min, conc_max, met_fix, conc_fix, enzyme_cost_weights, document_name, save_single_tables,verbose)
%
%The data refer to a single metabolic state; data describing several metabolic states, see 'help ecm_save_result_sbtab'
% 
%For loading such an SBtab file, see 'help load_model_and_data_sbtab'
%
%Arguments
% filename               filename for SBtab output
% network                (struct describing model, see mnt toolbox)
% v                      (nr x 1 vector of reaction rates)
% r                      (struct describing model kinetics, see mnt toolbox)
% c_data                 (nm x 1 vector of measured concentrations (only for information))
% u_data                 (nr x 1 vector of measured enzyme concentrations (only for information))
% kinetic_data           (OPTIONAL: struct with kinetic data; only to give original dmu0 values)
% conc_min               (nm x 1 vector of minimal concentrations)
% conc_max               (nm x 1 vector of maximal concentrations)
% met_fix                (OPTIONAL: list of metabolites with fixed concentrations)
% conc_fix               (OPTIONAL: fixed concentrations corresponding to met_fix)
% enzyme_cost_weights    ( nr x 1 vector of enzyme cost weights; default [])
% document_name          (string; document name to be mentioned in SBtab)
% save_single_tables     (flag for saving SBtab tables in single files; default 0)
% write_delta_G0         flag, determining whether standard reaction Gibbs free energies should be included in the file

eval(default('v','[]','r','[]','c_data','[]','u_data','[]','kinetic_data','[]','conc_min','[]','conc_max','[]','enzyme_cost_weights','[]','document_name','[]', 'save_single_tables','0','write_delta_G0','0','verbose','0'));

% -------------------------------------------------------
% prepare data

[network.metabolites,network.actions] = network_adjust_names_for_sbml_export(network.metabolites,network.actions);

[nm,nr] = size(network.N);

formulae = network_print_formulae(network);

network.kinetics      = r;
network.kinetics.type = 'cs';
network.kinetics.c    = nan * ones(size(network.metabolites));
network.kinetics.u    = nan * ones(size(network.actions));

% model tables ('Compound', 'Reaction', 'QuantityData')
sbtab_document = network_to_sbtab(network, struct('use_sbml_ids',0,'verbose',0,'modular_rate_law_kinetics', 0, 'write_concentrations',0,'write_enzyme_concentrations',0,'document_name',document_name));

% manually add column 'NameForPlots' in table 'Reaction'
reaction_table = sbtab_document.tables.Reaction;
if ~sbtab_table_has_column(reaction_table,'NameForPlots'),
  if sbtab_table_has_column(reaction_table,'Gene'),
    gene_names     = sbtab_table_get_column(reaction_table,'Gene');
  else
    gene_names     = sbtab_table_get_column(reaction_table,'ID');
  end
  reaction_table = sbtab_table_add_column(reaction_table,'NameForPlots',lower(gene_names));
  sbtab_document.tables.Reaction  = reaction_table;
end

% manually add column 'NameForPlots' in table 'Compound'
compound_table = sbtab_document.tables.Compound;
if ~sbtab_table_has_column(compound_table,'NameForPlots'),
  compound_names = sbtab_table_get_column(compound_table,'Name');
  compound_table = sbtab_table_add_column(compound_table,'NameForPlots',compound_names);
  sbtab_document.tables.Compound  = compound_table;
end

if save_single_tables,
  sbtab_document_save(sbtab_document,filename,0,1);
end
  
% flux table
if length(v),      
  flux_table = sbtab_table_construct(struct('TableName','Flux', 'TableType','Quantity','TableID','Quantity','Unit','mM/s'),{'QuantityType','Reaction','Reaction:Identifiers:kegg.reaction','Value'},{repmat({'flux'},nr,1),network.actions,network.reaction_KEGGID,v});
  if save_single_tables,
    sbtab_table_save(flux_table, struct('filename',[ filename '_Flux.tsv'])); 
  end
  sbtab_document = sbtab_document_add_table(sbtab_document,'Flux',flux_table);
end

% compound GFE of formation
% if length(r),
%   GFE_table = sbtab_table_construct(struct('TableName','GibbsEnergyOfFormation','TableType','Quantity','Unit','kJ/mol','StandardConcentration','1mM'),{'QuantityType','Compound','Compound:Identifiers:kegg.compound','Value'},{repmat({'standard Gibbs energy of formation'},nm,1),network.metabolites, network.metabolite_KEGGID, r.mu0});
%   if save_single_tables,
%   sbtab_table_save(GFE_table, struct('filename',[ filename '_FormationGFE.tsv'])); 
% end
%   sbtab_document = sbtab_document_add_table(sbtab_document,'GibbsEnergyOfFormation',GFE_table);
% end

% reaction GFE table
if write_delta_G0,
if length(r),
  delta_mu0      = network.N' * r.mu0;
  if length(kinetic_data),
    delta_mu0_orig = kinetic_data.dmu0.median;
  else
    delta_mu0_orig = nan*delta_mu0;
  end
  if sum(isfinite(delta_mu0)), 
  dGFE_table = sbtab_table_construct(struct('TableName','GibbsEnergyOfReaction','TableType','Quantity','Unit','kJ/mol','StandardConcentration','1mM'),{'QuantityType','Reaction','Reaction:Identifiers:kegg.reaction','Value','OriginalValue'},{repmat({'standard Gibbs energy of reaction'},nr,1),network.actions, network.reaction_KEGGID, delta_mu0, delta_mu0_orig});
  if save_single_tables,
    sbtab_table_save(dGFE_table, struct('filename',[ filename '_StandardReactionGFE.tsv'])); 
  end
  sbtab_document = sbtab_document_add_table(sbtab_document,'GibbsEnergyOfReaction',dGFE_table);
  end
end
end

% metabolite concentration table
if length(c_data), 
  concentration_table = sbtab_table_construct(struct('TableName','ConcentrationData', ...
                                                    'TableType','Quantity','TableID','Quantity','Unit','mM'),...
                                              {'QuantityType','Compound','Compound:Identifiers:kegg.compound','Value'},...
                                              {repmat({'concentration'},nm,1),network.metabolites, network.metabolite_KEGGID, c_data(:,1)});
  if save_single_tables,
    sbtab_table_save(concentration_table, struct('filename',[ filename '_Concentration.tsv'])); 
  end
end

% enzyme concentration table
if length(u_data),
  if sum(isfinite(u_data(:,1))),
  enzyme_table = sbtab_table_construct(struct('TableName','EnzymeData','TableType','Quantity','TableID','Quantity',...
                                              'Unit','mM'),{'QuantityType','Reaction','Reaction:Identifiers:kegg.reaction','Value'},{repmat({'concentration of enzyme'},nr,1),network.actions, network.reaction_KEGGID, u_data(:,1)});
  if save_single_tables,
    sbtab_table_save(enzyme_table, struct('filename',[ filename '_EnzymeConcentration.tsv']));
  end
  end
end

if length(met_fix),
  ind = label_names(met_fix,network.metabolite_names);
  conc_min(ind) = conc_fix;
  conc_max(ind) = conc_fix;
end

% metabolite constraint table
if length(conc_min),
  constraint_table = sbtab_table_construct(struct('TableName','ConcentrationConstraint', 'TableType','Quantity','TableID','Quantity','Unit','mM'),{'QuantityType','Compound','Compound:Identifiers:kegg.compound','Concentration:Min','Concentration:Max'},{repmat({'concentration'},nm,1),network.metabolites, network.metabolite_KEGGID, conc_min, conc_max});
  if save_single_tables,
    sbtab_table_save(constraint_table, struct('filename',[ filename '_ConcentrationConstraint.tsv']));
  end
  sbtab_document = sbtab_document_add_table(sbtab_document,'ConcentrationConstraint',constraint_table);
end

% enzyme cost weight table
if length(enzyme_cost_weights),
  enzyme_cost_weight_table = sbtab_table_construct(struct('TableName','EnzymeCostWeight','TableType','Quantity','TableID','Quantity','Unit','arbitrary unit'),{'QuantityType','Reaction','Reaction:Identifiers:kegg.reaction','Value'},{repmat({'enzyme cost weight'},nr,1),network.actions, network.reaction_KEGGID, enzyme_cost_weights(:,1)});
  if save_single_tables,
    sbtab_table_save(enzyme_cost_weight_table, struct('filename',[ filename '_EnzymeCostWeight.tsv']));  
  end
  sbtab_document = sbtab_document_add_table(sbtab_document,'EnzymeCostWeight',enzyme_cost_weight_table);
end

% position table
if isfield(network,'graphics_par'),
  x = network.graphics_par.x(1,:)';
  y = network.graphics_par.x(2,:)';
  position_table = sbtab_table_construct(struct('TableName','Position', 'TableType','Position','TableID','Position'),{'Element','PositionX','PositionY'},{[network.metabolites; network.actions],x,y});
  if isfield(network.graphics_par,'metinvisible'),
    position_table = sbtab_table_add_column(position_table,'IsInvisible',[column(network.graphics_par.metinvisible); column(network.graphics_par.actinvisible)],1);
  end
  if save_single_tables,
    sbtab_table_save(position_table, struct('filename',[ filename '_Position.tsv']));
  end
  sbtab_document = sbtab_document_add_table(sbtab_document,'Position',position_table);
end

% % Include validation data to model itself?
% 
% if exist('concentration_table','var'),
%   sbtab_document = sbtab_document_add_table(sbtab_document,'Concentration',concentration_table);
% end
% if exist('enzyme_table','var'),
%   sbtab_document = sbtab_document_add_table(sbtab_document,'EnzymeConcentration',enzyme_table);
% end

sbtab_document_save_to_one(sbtab_document,[filename, '_ECM_Model.tsv']);


%% Separate file for validation data

if exist('concentration_table','var'),
  if exist('enzyme_table','var'),
    sbtab_document_validation_data = sbtab_document_construct(struct,{'Concentration','EnzymeConcentration'},{concentration_table,enzyme_table});
  else
    sbtab_document_validation_data = sbtab_document_construct(struct,{'Concentration'},{concentration_table});
  end
  sbtab_document_save_to_one(sbtab_document_validation_data,[filename, '_ECM_ValidationData.tsv']);
end

if verbose, display(sprintf('Wrote model files (sbtab format) with basename\n%s', filename)); end
```
