## Supplementary material for "In-depth computational analysis of natural and artificial carbon fixation pathways": MATLAB files: ecm_save_result_sbtab.m

```
%ECM_SAVE_RESULT_SBTAB - Save ECM result in SBtab format
% 
% ecm_save_result_sbtab(filename, network, c, u, A_forward, options, c_min, c_max, u_min, u_max, u_capacity, eta_energetic, eta_saturation, v)
%
% The function arguments are the result of an ECM calculation, which may refer to several metabolic states
% To save data files for a single metabolic state, see  'help ecm_save_model_and_data_sbtab'
%
% Model document (default filename: [filename 'ModelState.tsv']) containing:
%   Metabolic network (compounds and reactions) and kinetic rate laws (parameters and formulae)
%
% Metabolic state document (default filename: [filename 'StateRuns.tsv']) containing:
%  o metabolite concentrations
%  o enzyme levels
% and optionally:
%  o enzyme maximal capacities
%  o enzyme thermodynamic efficiencies
%  o enzyme saturation efficiencies
%  o metabolic fluxes
%
%Arguments
% filename               filename for SBtab output
% network                (struct describing model, see mnt toolbox)
% c                      (struct) containing concentration vectors
% u                      (struct) containing enzyme concentration vectors
% A_forward              (struct) containing reaction affinity vectors
% options                (struct) options, see below
% c_min                  (nm x 1 vector of minimal concentrations)
% c_max                  (nm x 1 vector of maximal concentrations)
% u_min                  (nr x 1 vector of minimal enzyme concentrations)
% u_max                  (nr x 1 vector of maximal enzyme concentrations)
% u_capacity             (struct) containing enzyme capacities
% eta_energetic          (struct) containing energetic efficiency factors
% eta_saturation         (struct) containing saturation efficiency factors
% v                      (nr x 1 vector of reaction rates)
%
% Options (with default values)
%   .r                           = (mandatory) network_kinetics struct
%   .flag_one_file               = 1; flag: save tables in a single SBtab file
%   .document_name               = 'Model';
%   .save_tolerance_ranges       = 0; flag; add columns for minimal and maximal c values to concentration table
%   .sbtab_attributes            = struct(); additional document attributes to be put in the SBtab document
%   .filename_model_state        = 'ModelState';
%   .filename_state_runs         = 'StateRuns';
%   .write_concentrations        = 0       flag: show metabolite concentrations (first state) in model parameter table?
%   .write_enzyme_concentrations = 0 flag: show enzyme concentrations (first state) in model parameter table?
%   .modular_rate_law_kinetics   = 0   flag: show kinetic rate law formulae in table
%   .use_measurement_table       = 0;

function ecm_save_result_sbtab(filename, network, c, u, A_forward, options, c_min, c_max, u_min, u_max, u_capacity, eta_energetic, eta_saturation,v)

eval(default('options','struct','c_min', '[]', 'c_max', '[]', 'u_min', '[]', 'u_max', '[]', 'u_capacity', '[]', 'eta_energetic', '[]', 'eta_saturation', '[]','v','[]'));

if strcmp(filename(end-3:end),'.tsv'), 
  filename(1:end-4);
end
  
options_default.omit_kegg_ids = 1;
options_default.flag_one_file = 1;
if isfield(network,'kinetics'),
  options_default.r = network.kinetics;
else
  options_default.r = struct;
end
options_default.use_measurement_table  = 0;
options_default.document_id = 'Model';
options_default.document_name = 'ECM metabolic state';
options_default.save_tolerance_ranges = 0;
options_default.sbtab_attributes = struct('Document', options_default.document_id,'DocumentName',options_default.document_name);
options_default.filename_model_state  = 'ModelState';
options_default.filename_state_runs   = 'StateRuns';
options_default.write_concentrations = 0;
options_default.write_enzyme_concentrations = 0;
options_default.modular_rate_law_kinetics = 0;

options = join_struct(options_default, options);

[network.metabolites,network.actions] = network_adjust_names_for_sbml_export(network.metabolites,network.actions);

[nm,nr] = size(network.N);

if ~isfield(network,'metabolite_KEGGID'),
  if isfield(network,'Identifiers_kegg_compound'),
    network.metabolite_KEGGID = network.Identifiers_kegg_compound;
  else
    network.metabolite_KEGGID = [];
  end
end

if isfield(network,'metabolite_KEGGID'),
  metabolite_KEGGID = network.metabolite_KEGGID;
else
  metabolite_KEGGID = [];
end

if ~isfield(network,'reaction_KEGGID'),
  if isfield(network,'Identifiers_kegg_reaction'),
    network.reaction_KEGGID = network.Identifiers_kegg_reaction;    
  else
    network.reaction_KEGGID = [];
  end
end

if isfield(network,'reaction_KEGGID'),
  reaction_KEGGID = network.reaction_KEGGID;
else
  reaction_KEGGID = [];
end

if options.omit_kegg_ids,
  metabolite_KEGGID = [];
  reaction_KEGGID = [];
end

% -------------------------------------------------------
% model with optimized metabolite and enzyme levels
% -------------------------------------------------------

[network.metabolites,network.actions] = network_adjust_names_for_sbml_export(network.metabolites,network.actions);

%formulae = network_print_formulae(network);

% don't show metabolite and enzyme levels in parameter table 
my_opt = struct('use_sbml_ids',0,'verbose',0,'write_concentrations',options.write_concentrations,'write_enzyme_concentrations',options.write_enzyme_concentrations,'modular_rate_law_kinetics',options.modular_rate_law_kinetics,'document_name', options.document_name,'omit_kegg_ids',1);

% try to insert results from enzyme prediction with common modular rate law
[nm,nr]          = size(network.N);
network.kinetics = options.r;

if isfield(c,'emc4cm'),
  network.kinetics.type = 'cs';
  network.kinetics.c    = c.emc4cm;
  network.kinetics.u    = u.emc4cm;
  sbtab_document = network_to_sbtab(network, join_struct(my_opt,struct('c',c.emc4cm,'modular_rate_law_parameter_id', 0)));
elseif isfield(c,'emc3sp'),
  network.kinetics.type = 'ds';
  network.kinetics.c    = c.emc3sp;
  network.kinetics.u    = u.emc3sp;
  sbtab_document = network_to_sbtab(network, join_struct(my_opt,struct('c',c.emc3sp,'modular_rate_law_parameter_id', 0)));
else
  network.kinetics.type = 'cs';
  sbtab_document = network_to_sbtab(network, my_opt);
end

sbtab_document.attributes = join_struct(sbtab_document.attributes,options.sbtab_attributes);

if strcmp(filename(end), filesep),
  sbtab_document_save_to_one(sbtab_document,[filename options.filename_model_state '.tsv'],1);
else
  sbtab_document_save_to_one(sbtab_document,[filename '_' options.filename_model_state '.tsv'],1);
end

% -----------------------------------------------
% metabolic states
% -----------------------------------------------


% -----------------------------------------------
% predicted concentrations

fn = fieldnames(c); 

attributes = struct('TableName','Metabolite concentrations', 'TableID','MetaboliteConcentration', 'TableType','QuantityMatrix', 'Unit','mM');

c_table = sbtab_table_construct(attributes, {'QuantityType','Compound'}, {repmat({'concentration'},nm,1),network.metabolites});

if length(metabolite_KEGGID),
  c_table = sbtab_table_add_column(c_table,'Compound:Identifiers:kegg.compound',metabolite_KEGGID,1);
end

for it = 1:length(fn),
  if length(c.(fn{it})),
    if options.use_measurement_table,
      c_table = sbtab_table_add_column(c_table, ['>' fn{it}], c.(fn{it})(:,1), 1);
    else
      c_table = sbtab_table_add_column(c_table, fn{it}, c.(fn{it})(:,1), 0);
    end
  end
end

% predicted concentration tolerance ranges
if options.save_tolerance_ranges,
  for it = 1:length(fn),
   if isfield(c_min,fn{it}),
     c_table = sbtab_table_add_column(c_table,[fn{it} ':Lower'], c_min.(fn{it})(:,1), 1);
     c_table = sbtab_table_add_column(c_table,[fn{it} ':Upper'], c_max.(fn{it})(:,1), 1);
   end
  end
end

% -----------------------------------------------
% predicted enzyme levels

fn = fieldnames(u); 

u_table = sbtab_table_construct(struct('TableName','Enzyme concentrations','TableID','EnzymeConcentration','TableType','QuantityMatrix','Unit','mM'),{'QuantityType','Reaction'},{repmat({'concentration of enzyme'},nr,1),network.actions});

if length(reaction_KEGGID),
  u_table = sbtab_table_add_column(u_table,'Reaction:Identifiers:kegg.reaction',reaction_KEGGID,1);
end

for it = 1:length(fn),
  if length(u.(fn{it})),
    if options.use_measurement_table,
      u_table = sbtab_table_add_column(u_table, ['>' fn{it}], u.(fn{it})(:,1), 1);
    else
      u_table = sbtab_table_add_column(u_table, fn{it}, u.(fn{it})(:,1), 0);
    end
  end
end

% predicted concentration tolerance ranges
if options.save_tolerance_ranges,
  for it = 1:length(fn),
   if isfield(u_min,fn{it}),
    u_table = sbtab_table_add_column(u_table,[fn{it} ':Lower'], u_min.(fn{it})(:,1), 1);
    u_table = sbtab_table_add_column(u_table,[fn{it} ':Upper'], u_max.(fn{it})(:,1), 1);
   end
  end
end

% -----------------------------------------------
% predicted reaction driving forces

fn = fieldnames(A_forward); 

A_forward_table = sbtab_table_construct(struct('TableName','Gibbs free energies of reaction','TableID','ReactionGibbsFreeEnergy', 'TableType','QuantityMatrix','Unit','kJ/mol'),{'QuantityType','Reaction'},{repmat({'Gibbs energy of reaction'},nr,1),network.actions});

if length(reaction_KEGGID),
  A_forward_table = sbtab_table_add_column(A_forward_table,'Reaction:Identifiers:kegg.reaction',reaction_KEGGID,1);
end

for it = 1:length(fn),
  if options.use_measurement_table,
    A_forward_table = sbtab_table_add_column(A_forward_table, ['>' fn{it}], A_forward.(fn{it})(:,1),1);
  else
    A_forward_table = sbtab_table_add_column(A_forward_table, fn{it}, A_forward.(fn{it})(:,1),0);
  end
end

% -----------------------------------------------
% predicted enzyme capacity (v/kcat) and efficiencies

if length(u_capacity),

  u_capacity_table = sbtab_table_construct(struct('TableName','Enzyme capacities','TableID','EnzymeCapacity','TableType','QuantityMatrix','Unit','mM'),{'QuantityType','Reaction','Value'},{repmat({'enzyme capacity'},nr,1),network.actions, u_capacity});
  
  if length(reaction_KEGGID),
    u_capacity_table = sbtab_table_add_column(u_capacity_table,'Reaction:Identifiers:kegg.reaction',reaction_KEGGID,1);
  end
  
  fn = fieldnames(eta_energetic); 
  
  eta_energetic_table = sbtab_table_construct(struct('TableName','Energetic efficiencies','TableID','EnergeticEfficiency','TableType','QuantityMatrix','Unit','dimensionless'),{'QuantityType','Reaction'},{repmat({'energetic efficiency'},nr,1),network.actions});
  
  if length(reaction_KEGGID),
    eta_energetic_table = sbtab_table_add_column(eta_energetic_table,'Reaction:Identifiers:kegg.reaction',reaction_KEGGID,1);
  end

  
  for it = 1:length(fn),
    if options.use_measurement_table,
      eta_energetic_table = sbtab_table_add_column(eta_energetic_table, ['>' fn{it}], eta_energetic.(fn{it})(:,1),1);
    else
      eta_energetic_table = sbtab_table_add_column(eta_energetic_table, fn{it}, eta_energetic.(fn{it})(:,1),0);
    end
  end
  
  fn = fieldnames(eta_saturation); 
  
  eta_saturation_table = sbtab_table_construct(struct('TableName','Saturation efficiencies','TableID','SaturationEfficiency','TableType','QuantityMatrix','Unit','dimensionless'),{'QuantityType','Reaction'},{repmat({'saturation efficiency'},nr,1),network.actions});
  
  if length(reaction_KEGGID),
    eta_saturation_table = sbtab_table_add_column(eta_saturation_table,'Reaction:Identifiers:kegg.reaction',reaction_KEGGID,1);
  end

  for it = 1:length(fn),
    if options.use_measurement_table,
      eta_saturation_table = sbtab_table_add_column(eta_saturation_table, ['>' fn{it}], eta_saturation.(fn{it})(:,1),1);
    else
      eta_saturation_table = sbtab_table_add_column(eta_saturation_table, fn{it}, eta_saturation.(fn{it})(:,1),0);
    end
  end

end

if length(v),
  
  if ~isstruct(v),
    % if v is just a vector
    v = struct('data',v);
  end
  fn = fieldnames(v); 
  v_table = sbtab_table_construct(struct('TableName','Metabolic fluxes','TableID','MetabolicFlux','TableType','QuantityMatrix','Unit','mM/s'),{'QuantityType','Reaction'},{repmat({'rate of reaction'},nr,1),network.actions});
  
  if length(reaction_KEGGID),
    v_table = sbtab_table_add_column(v_table,'Reaction:Identifiers:kegg.reaction',reaction_KEGGID,1);
  end

  for it = 1:length(fn),
    if options.use_measurement_table,
      v_table = sbtab_table_add_column(v_table, ['>' fn{it}], v.(fn{it})(:,1),1);
    else
      v_table = sbtab_table_add_column(v_table, fn{it}, v.(fn{it})(:,1),0);
    end
  end
end

% -----------------------------------------------
% sample table

if options.use_measurement_table,
  sample_table = sbtab_table_construct(struct('TableName','Metabolic states','TableID','MetabolicStates','TableType','Measurement'),{'ID','Name'},{fieldnames(c),fieldnames(c)});
end

% -----------------------------------------------
% save tables

switch options.flag_one_file, 
  
  case 0,
    %display(sprintf('Writing model files (SBtab format) with basename\n%s', filename))
    sbtab_table_save(c_table, struct('filename',[ filename '_Concentration.tsv']));
    sbtab_table_save(u_table, struct('filename',[ filename '_EnzymeConcentration.tsv']));
    sbtab_table_save(A_forward_table, struct('filename',[ filename '_ReactionGibbsFreeEnergy.tsv']));
    if length(u_capacity),
      sbtab_table_save(u_capacity_table, struct('filename',[ filename '_EnzymeCapacity.tsv']));
      sbtab_table_save(eta_energetic_table, struct('filename',[ filename '_EnergeticEfficiency.tsv']));
      sbtab_table_save(eta_saturation_table, struct('filename',[ filename '_SaturationEfficiency.tsv']));
    end
    if length(v),
      sbtab_table_save(v_table, struct('filename',[ filename '_MetabolicFlux.tsv']));
    end

  case 1,
    sbtab_document = sbtab_document_construct(options.sbtab_attributes);
    sbtab_document = sbtab_document_add_table(sbtab_document,'Concentration',c_table);
    sbtab_document = sbtab_document_add_table(sbtab_document,'EnzymeConcentration',u_table);
    sbtab_document = sbtab_document_add_table(sbtab_document,'ReactionAffinity',A_forward_table);
    if length(u_capacity),
      sbtab_document = sbtab_document_add_table(sbtab_document,'EnzymeCapacity',u_capacity_table);
      sbtab_document = sbtab_document_add_table(sbtab_document,'EnergeticEfficiency',eta_energetic_table);
      sbtab_document = sbtab_document_add_table(sbtab_document,'SaturationEfficiency',eta_saturation_table);
    end
    if length(v),
      sbtab_document = sbtab_document_add_table(sbtab_document,'MetabolicFlux',v_table);
    end
    if options.use_measurement_table,
      sbtab_document = sbtab_document_add_table(sbtab_document,'MetabolicStates',sample_table);
    end
    if strcmp(filename(end), filesep),
      sbtab_document_save_to_one(sbtab_document, [ filename options.filename_state_runs '.tsv'],1);
    else
      sbtab_document_save_to_one(sbtab_document, [ filename '_' options.filename_state_runs '.tsv'],1);
    end
    
end
```
