## Supplementary material for "In-depth computational analysis of natural and artificial carbon fixation pathways": MATLAB files: ecm_sbtab2mnt.m

```
function [network, v, conc_min, conc_max, kinetic_data] = ecm_sbtab2mnt(model_filename, filenames,kinetic_data_file_names,options,organism_long,kegg_conversion_file,layout_file,reaction_column_name,compound_column_name,use_kegg_ids)

% [network, v, conc_fix, kinetic_data] = ecm_sbtab2mnt(model_filename, model_dir, matlab_dir,kinetic_data_file_names)
%
% Read network structure (from SBtab files), collect dG0' values (from network_thermo)
% and convert them into MNT network format
% the arguments kegg_conversion_file, layout_file can be given 
% explicitly, or as fields of "filenames" (which can remain empty otherwise)

% I use mM and convert everything from Elad's convention within this script

% if organism_long is given, the kinetic data are filtered for the organism

eval(default('options','struct','filenames.kegg_conversion_file','layout_file','filenames.layout_file'));

if ~exist('reaction_column_name','var'),  reaction_column_name = []; end
if ~exist('compound_column_name','var'),  compound_column_name = []; end
if ~exist('use_kegg_ids','var'),          use_kegg_ids = []; end

options_default = struct();
options = join_struct(options_default,options);

network            = sbtab_to_network(model_filename, struct('load_quantity_table',0));
v                  = cell_string2num(network.Flux);
network.external   = [network.N * v ~= 0];
network.metabolite_names = ecm_kegg_compound_id_to_name(network.metabolite_KEGGID,kegg_conversion_file);

% -----------------------------------------------------------
% omit protons from network model

ind_proton = find(strcmp('C00080',network.metabolite_KEGGID));

if ind_proton,
  display('Omitting protons as a compound in the network model');
  network = network_remove(network,ind_proton);
end


% -----------------------------------------------------------
% extract kegg reaction IDs

for it=1:length(network.actions),
   dum = Strsplit('_',network.actions{it});
   network.reaction_KEGGID{it,1} = dum{2};
end

network.Identifiers_kegg_reaction = network.reaction_KEGGID;


% -----------------------------------------------------------
% convert from M to mM

conc_min  = 1000 * cell_string2num(network.Concentration_Min);
conc_max  = 1000 * cell_string2num(network.Concentration_Max);

ind_water = network_find_water(network);
conc_min(ind_water) = 1;
conc_max(ind_water) = 1;

network                       = netgraph_make_graph(network);
network.graphics_par.metnames = network.metabolite_names;

network.reaction_formula_KEGGID = strrep(network.formulae,' ','_');

network = netgraph_read_positions(network,layout_file,[0,0],1,1,network.reaction_formula_KEGGID);


% -----------------------------------------------------------
% make data structure 'kinetic_data' for all kinetic constants
% and insert values of equilibrium constants

import_quantity_list = {'standard Gibbs energy of reaction', 'standard chemical potential','Michaelis constant',...
                    'activation constant', 'inhibitory constant',...
                    'equilibrium constant','substrate catalytic rate constant', ...
                    'product catalytic rate constant'};

kinetic_data = data_integration_load_kinetic_data(import_quantity_list, [], network, kinetic_data_file_names, 0, use_kegg_ids,1,0,'Organism',organism_long,reaction_column_name,compound_column_name);
```
