## Supplementary material for "In-depth computational analysis of natural and artificial carbon fixation pathways": MATLAB files: ecm_tolerances.m

```
%ECM_TOLERANCES - Compute approximate tolerance ranges for metabolites and enzymes
% 
%[c_min, c_max, u_min, u_max] = ecm_tolerances(c,u,v,ecm_options,mca_info)

function [c_min, c_max, u_min, u_max] = ecm_tolerances(c,u,v,ecm_options,mca_info)

% rough estimate, assuming that 1/2 deltaxi^2 Hii = ytolerance

if ecm_options.tolerance_from_hessian,

  for it = 1:length(ecm_options.ecm_scores),
    my_cost_score     = ecm_options.ecm_scores{it};
    my_x              = log(c.(my_cost_score));
    my_cost_score     = ecm_options.ecm_scores{it};
    my_cost_tolerance = log(ecm_options.cost_tolerance_factor);
    my_hessian        = mca_info.(my_cost_score).x_log_hessian;
    if strcmp(my_cost_score,'mdf'),
      %% expansion does not make sense for MDF (function is 
      %% non-differentiable at optimum)
      my_hessian = nan * my_hessian;
    end
    %% assume that all other metabolites are ideally co-adjusted
    %% my_x_tolerance = sqrt(2 * my_cost_tolerance * diag(inv(my_hessian))); 
    %% assume that all other metabolites remain fixed
    my_x_tolerance = sqrt(2 * my_cost_tolerance * 1./diag(my_hessian));
    my_x_min = max(my_x-my_x_tolerance, log(ecm_options.conc_min));
    my_x_max = min(my_x+my_x_tolerance, log(ecm_options.conc_max));
    c_min.(my_cost_score) = exp(my_x_min);
    c_max.(my_cost_score) = exp(my_x_max);
    
    %if 0,
    %  figure(1024);
    %  errorbar(1:length(my_x), exp(my_x), exp(my_x)-exp(my_x_min), exp(my_x_max)-exp(my_x),'.');
    %  set(gca,'YScale','Log'); title(my_cost_score)
    %  xlabel('Metabolite number'); ylabel('Tolerance(log metabolite level)');
    %end

    %% usual scaled elasticities (in matrix my_E_sc)
    my_u           = u.(my_cost_score);
    my_E_sc        = diag(1./[v+10^-10]) * mca_info.(my_cost_score).elasticities_rate_x;
    my_u_tolerance = my_u.^2 .* sqrt(2 * my_cost_tolerance * diag(my_E_sc * inv(my_hessian) * my_E_sc'));
    my_u_max = my_u + my_u_tolerance;
    my_u_min = my_u.^2 ./ my_u_max;
    u_min.(my_cost_score) = my_u_min;
    u_max.(my_cost_score) = my_u_max;
    %if 0,
    %  figure(1024);
    %  errorbar(1:length(my_u), my_u, my_u - my_u_min, my_u_max-my_u,'.');
    %  set(gca,'YScale','Log'); title(my_cost_score)
    %  xlabel('Enzyme number'); ylabel('Tolerance(enzyme level)');
    %end
    
  end
  %% to avoid complex nan values: 
  if isfield(c_min,'mdf'),
    c_min.mdf(find(~isfinite(  c_min.mdf)))=nan;
  end

else
  display('  No tolerances are computed');
  c_min = c;
  c_max = c;
  u_min = u;
  u_max = u;
end

% Hessian is not defined for MDF (not differentiable)

if isfield(c_min,'mdf'),
  c_min.mdf = nan * c_min.mdf;
  c_max.mdf = nan * c_max.mdf;
  u_min.mdf = nan * real(u_min.mdf);
  u_max.mdf = nan * real(u_max.mdf);
end
```
