## Supplementary material for "In-depth computational analysis of natural and artificial carbon fixation pathways": MATLAB files: ecm_update_options.m

```
function ecm_options = ecm_update_options(network, ecm_options);

% ECM_UPDATE_OPTIONS - Helper function for ECM options (struct 'ecm_options')
%
% ecm_options = ecm_update_options(network, ecm_options)
%
% Update ECM options for a given network:
%  - adapt metabolite constraints
%  - insert protein cost weights
%  - adjust Keq values

display('Updating the parameters for ECM');

% Insert default options, if not yet specified

ecm_options_default = ecm_default_options(network);
ecm_options         = join_struct(ecm_options_default,ecm_options);

% Adjust concentration constraints

conc_min = ecm_options.conc_min;
conc_max = ecm_options.conc_max;
c_data   = ecm_options.c_data  ;

[nm,nr] = size(network.N);
if isempty(conc_min), conc_min = ecm_options.conc_min_default * ones(nm,1); end
if isempty(conc_max), conc_max = ecm_options.conc_max_default * ones(nm,1); end

% Replace cofactor concentrations

if length(c_data),
  display('  o Replacing cofactor concentrations');
  if 1,
    display('  o Using first data values');
    c_data_median = c_data(:,1);
  else
    display('  o Using median data values');
    c_data_median = nanmedian(c_data,2);
  end    
  ll = label_names(ecm_options.replace_cofactors,network.metabolites);
  ll = ll(find(ll));
  ll = ll(find(isfinite(c_data_median(ll))));
  conc_min(ll) = c_data_median(ll);
  conc_max(ll) = c_data_median(ll);
else
  display('  o Not replacing any cofactor concentrations');
end

% Fix predefined concentrations

if length(ecm_options.fix_metabolites),
  display('  o Fixing some metabolite concentrations');
  for it = 1:length(ecm_options.fix_metabolites),
    ll = label_names(ecm_options.fix_metabolites(it),network.metabolites);
    conc_min(ll) = ecm_options.fix_metabolite_values(it);    
    conc_max(ll) = ecm_options.fix_metabolite_values(it);
  end
else
  display('  o Not fixing any metabolite concentrations');
end

ecm_options.conc_min = conc_min;
ecm_options.conc_max = conc_max;

ind_conc_fix = find(ecm_options.conc_min == ecm_options.conc_max);
met_fix      = network.metabolites(ind_conc_fix);
conc_fix     = ecm_options.conc_min(ind_conc_fix);
if prod(size(met_fix))==0, met_fix={}; conc_fix=[]; end

ind_met_fix = label_names(met_fix, network.metabolites);

if setdiff(find(network.external),ind_met_fix),
  display('  o Some external metabolites in network will not be treated as fixed');
end

% 

ecm_options.conc_min    = conc_min;
ecm_options.conc_max    = conc_max;
ecm_options.conc_fix    = conc_fix;
ecm_options.met_fix     = met_fix;
ecm_options.ind_met_fix = ind_met_fix;


% -------------------------------------------
% set enzyme cost weights
if isempty(ecm_options.use_cost_weights),
  if ecm_options.verbose,
    display('Using predefined enzyme cost weights');
  end
else  
  switch ecm_options.use_cost_weights,
    case 'none',
      ecm_options.enzyme_cost_weights = ones(length(ecm_options.ind_scored_enzymes),1);
    case 'protein_size',
      ecm_options.enzyme_cost_weights = network.enzyme_size(ecm_options.ind_scored_enzymes);
    case 'protein_mass',
      ecm_options.enzyme_cost_weights = network.enzyme_mass(ecm_options.ind_scored_enzymes);
    case 'aa_composition',
      ecm_options.enzyme_cost_weights = network.akashi_protein_cost(ecm_options.ind_scored_enzymes);
  end
  %% scale to median = 1
  ecm_options.enzyme_cost_weights = ecm_options.enzyme_cost_weights / nanmedian(ecm_options.enzyme_cost_weights);
end

% -----------------------------
% adapt lambda_regularistion to typical magnitude of enzyme cost

median_cost = nanmedian(ecm_options.enzyme_cost_weights);

if isnan(median_cost), median_cost = 1; end 

if length(ecm_options.lambda_reg_factor),
  ecm_options.lambda_regularisation = ecm_options.lambda_reg_factor * median_cost;
end

% Adapt print_graphics

if ecm_options.show_graphics == 0, ecm_options.print_graphics = 0;         end
if ecm_options.compute_tolerance,  ecm_options.tolerance_from_hessian = 0; end

% -----------------------------

if ~isfield(ecm_options,'N_subunits_in_complex'),
  ecm_options.N_subunits_in_complex  = ones(size(ecm_options.enzyme_cost_weights));
end

if ~isfield(ecm_options,'N_cat_sites_in_complex'),
  ecm_options.N_cat_sites_in_complex = ones(size(ecm_options.enzyme_cost_weights));
end
```
