## Supplementary material for "In-depth computational analysis of natural and artificial carbon fixation pathways": MATLAB files: ecm_parameter_balancing.m

```
function [r, r_orig, kinetic_data, r_samples, pb_options, parameter_prior, r_mean, r_std] = ecm_parameter_balancing(network, pb_options, kinetic_data);

% ECM_PARAMETER_BALANCING - Run parameter balancing with standard options
%
% [r, r_orig, kinetic_data, r_samples, pb_options, parameter_prior, r_std] = ecm_parameter_balancing(network, pb_options, kinetic_data);
%
% Input
%   network         Metabolic network data structure
%   kinetic_data    Kinetic_data used
%   pb_options         Parameter balancing options used
%
% Output
%   r               Kinetic constants (posterior mode values, respecting the linear constraints)
%   r_orig          Original kinetic constants (used as input in parameter balancing)
%   r_mean          Kinetic constants (posterior means, ignoring the  linear constraints)
%   r_std           Kinetic constants (posterior standard deviations, ignoring the  linear constraints)
%   r_samples       Kinetic constants sampled from the posterior
%   pb_options      Parameter balancing options used (for details, see parameter_balancing_kinetic.m)
%   kinetic_data    Kinetic_data used (for details, see parameter_balancing_kinetic.m)
%   parameter_prior Prior distributions used

pb_options = join_struct(parameter_balancing_options,pb_options);

% extra options used only in this function  
pb_options_default.flag_given_kinetics = 0;
pb_options_default.insert_Keq_from_data = 0;

pb_options = join_struct(pb_options_default,pb_options);


% --------------------------------------------------------------------------------------

if pb_options.flag_given_kinetics,
  %% Instead of running parameter balancing, simply return the parameters found in the model

  switch network.kinetics.type 
    case {'cs','ms','ds'},
      display('  Using kinetic constants found in network.kinetics');
      r.Keq     = network.kinetics.Keq;
      r.KM      = network.kinetics.KM;
      [r.Kcatf, r.Kcatr] = modular_KV_Keq_to_kcat(network.N,network.kinetics);
    otherwise error('Kinetics cannot be handled');
  end

else
  %% Use kinetic parameters from "kinetic_data" function argument and run parameter balancing

   [r, r_orig, kinetic_data, r_samples, parameter_prior, r_mean, r_std] = parameter_balancing_kinetic(network, kinetic_data, pb_options);

end

% -----------------------------
% if desired, insert original Keq values

if pb_options.insert_Keq_from_data,
  if length(pb_options.Keq_given),
    display('Inserting predefined equilibrium constants'); 
    ind_finite = find(isfinite(pb_options.Keq_given));
    %r.Keq(ind_finite) = r_orig.Keq(ind_finite);
    r.Keq(ind_finite) = pb_options.Keq_given(ind_finite);
  end
end
```
