## Supplementary material for "In-depth computational analysis of natural and artificial carbon fixation pathways": MATLAB files: ecm_emc1.m

```
function [u_cost, u, w] = ecm_emc1(x,pp)

% [u_cost, u] = ecm_emc1(x,pp)

u = abs(pp.v) ./ pp.kc_forward;
u(pp.v==0) = 0;
w = pp.kc_forward;

u_cost = sum(pp.enzyme_cost_weights .* u(pp.ind_scored_enzymes));
```
