## Supplementary material for "In-depth computational analysis of natural and artificial carbon fixation pathways": MATLAB files: ecm_emc3sp.m

```
function [u_cost, u, w] = ecm_emc3sp(x,pp)

% [u_cost, u] = ecm_emc3sp(x,pp)

delta_G_by_RT = pp.N_forward' * x - pp.log_Keq_forward;

w =  [ pp.kc_forward .* [1 - exp(delta_G_by_RT)] ./ [[  pp.km_forward  ./ exp(abs(pp.M_forward) * x)] .* [1 + 1./pp.km_forward .* exp(abs(pp.M_forward) * x) + 1./pp.kmprod_forward .* exp(abs(pp.Mprod_forward) * x)]  ]];

u = abs(pp.v) ./ w;
u(pp.v==0) = 0;

if sum(delta_G_by_RT>0),
  u(find(delta_G_by_RT>0)) = 10^20 * max(delta_G_by_RT);
  warning('Unfeasible metabolite profile');
end

u_cost = sum(pp.enzyme_cost_weights .* u(pp.ind_scored_enzymes));
```
