## Supplementary material for "In-depth computational analysis of natural and artificial carbon fixation pathways": MATLAB files: ecm_emc4cm.m

```
function [u_cost, u, w] = ecm_emc4cm(x,pp,no_warnings)

  eval(default('no_warnings','0'));
  
% [u_cost, u] = ecm_emc4cmr(x,pp)

delta_G_by_RT = sign(pp.v) .* [pp.network.N' * x - log(pp.network.kinetics.Keq)];

kinetics      = pp.network.kinetics;
kinetics.type = 'cs';
kinetics.u    = ones(size(pp.network.N,2),1); 

w = network_velocities(exp(x),pp.network,kinetics);
u = pp.v./w;
u(pp.v==0) = 0;

if sum(delta_G_by_RT>0),
  u(find(delta_G_by_RT>0)) = 10^10 * max(delta_G_by_RT);
  if no_warnings == 0,
    warning('Unfeasible metabolite profile');
  end
end

u_cost = sum(pp.enzyme_cost_weights.*u(pp.ind_scored_enzymes));
```
