## Supplementary material for "In-depth computational analysis of natural and artificial carbon fixation pathways": MATLAB files: ecm_emc4dm.m

```
function [u_cost, u, w] = ecm_emc4cmr(x,pp)

% [u_cost, u] = ecm_emc4cmr(x,pp)

network = pp.network;

delta_G_by_RT =  sign(pp.v) .* [network.N' * x - log(pp.network.kinetics.Keq)];

network.kinetics.type = 'ds';
network.kinetics.u    = ones(size(network.N,2),1); 

w = network_velocities(exp(x),network);

u = pp.v ./ w;
u(pp.v==0) = 0;

if sum(delta_G_by_RT>0),
  u(find(delta_G_by_RT>0)) = 10^20 * max(delta_G_by_RT);
  warning('Unfeasible metabolite profile');
end

u_cost = sum(pp.enzyme_cost_weights.* u(pp.ind_scored_enzymes));
```
